# MAIT cells have a negative impact on glioblastoma

**DOI:** 10.1101/2022.07.17.499189

**Authors:** Seketoulie Keretsu, Taijun Hana, Alexander Lee, Noemi Kedei, Nargis Malik, Hye Kim, Jo Spurgeon, Guzal Khayrullina, Benjamin Ruf, Ayaka Hara, Morgan Coombs, Matthew Watowich, Ananth Hari, Michael K.B. Ford, Cenk Sahinalp, Masashi Watanabe, George Zaki, Mark R. Gilbert, Patrick. J. Cimino, Robert Prins, Masaki Terabe

## Abstract

Glioblastoma (GBM) is the most aggressive primary brain cancer in adults and remains incurable. Our study revealed an immunosuppressive role of mucosal-associated invariant T (MAIT) cells in GBM. In bulk RNA sequencing data analysis of GBM tissues, MAIT cell gene signature significantly correlated with poor patient survival. A scRNA-seq of CD45^+^ cells from 23 GBM tissue samples showed 15 (65.2%) were positive for MAIT cells and the enrichment of MAIT17. The MAIT cell signature significantly correlated with the activity of tumor-associated neutrophils (TANs) and myeloid-derived suppressor cells (MDSCs). Multiple immune suppressive genes known to be used by TANs/MDSCs were upregulated in MAIT-positive tumors. Spatial imaging analysis of GBM tissues showed that all specimens were positive for both MAIT cells and TANs and localized enrichment of TANs. These findings highlight the MAIT-TAN/MDSC axis as a novel therapeutic target to modulate GBM’s immunosuppressive tumor microenvironment.

## Introduction

GBM is the most aggressive primary brain tumor in adults, and the five-year survival of GBM is approximately 5%, underscoring the urgent need for better treatment^1,2,3^. Immunotherapies have shown promise in treatments for other cancer types but, to date, have not yet demonstrated clinical efficacy for GBM^4^. The lack of efficacy has been attributed to multiple factors, including a highly immunosuppressive microenvironment, including a high immunosuppressive myeloid cell content^5^. However, the factors that induce those immunosuppressive myeloid cells still require further investigation.

Recently, it was reported that a transcriptomic expression level of major histocompatibility complex class I-related (*MR1*) significantly correlated to the poor survival of glioma patients^6^. However, the underlying mechanism still needs to be determined. MR1 presents antigens to mucosal-associated invariant T (MAIT) cells, which express a semi-invariant TCRα chain utilizing the TCR gene segment combinations Vα7.2-Jα33, Vα7.2-Jα12, or Vα7.2-Jα20 in humans^7^. They are highly prevalent in humans. Upon activation, MAIT cells secrete multiple cytokines, such as Th1 and Th17-cytokines. Their cytokine profile is altered by the local tissue environment^8^. In humans, MAIT cells have two functional subsets, IFN-γ-producing MAIT1 and IL-17-producing MAIT17. Most MAIT cells in tissues can produce IL-17^9–11^. Th17-type cytokines promote the local attraction and activation of specific myeloid cells, such as neutrophils, through the induction of chemokines^12,13^.

The role of MAIT cells in cancer immunity is unclear^8^. MAIT cells can potentially lyse cancer cells, including glioma-derived cells, under certain in vitro conditions, but their ability to target cancers in vivo has not been confirmed^14,15^. They were reported to facilitate tumor immunity in liver cancer models when stimulated with CpG + 5-OP-RU, MAIT cell agonist^16^.

However, they were shown to be suppressed by myeloid cells in tumor tissue^17^. Conversely, MAIT cells promoted tumor growth and metastasis in mouse tumor models^15,18^. An inverse correlation between the number of MAIT cells in cancer tissues and patient survival has been reported in liver and colorectal cancer^19,20^. The discrepancy in the role of MAIT cells among tumor immunological studies suggests that their surrounding environment could modify MAIT cell functions. The interaction between GBM and MAIT cells has not been well studied. An earlier study proposed the existence of MAIT cells in clinical GBM samples, but their clinical impact or function remained to be elucidated^21^.

In this study, we explored the role of MAIT cells in GBM by using multi-omics and flow cytometry. By utilizing TCGA-GBM datasets, we found a significant correlation between the expression levels of MAIT cell signature genes and worsened survival in a patient cohort. Patients with MAIT TCR transcripts showed a higher expression of *IL17A,* which could induce chemokines that recruit myeloid cells^22^. In a single-cell RNA-seq study of tumor-infiltrating leukocytes (TILs) from 23 patients, 15 patients (65.2%) were found positive for MAIT cells, accounting for 1.46% of all T cells. MAIT cells clustered with cell populations expressing a high level of transcription factor gene *RORC*, suggesting the enrichment of MAIT17 subset. Consistent with these observations, further analysis indicated that MAIT cells in GBM correlated with the activity of TANs/MDSCs. Spatial analysis of the MAIT-neutrophil relationship indicated neutrophils are enriched in the tissue and could be found around MAIT cells. Based on the findings, we propose that MAIT cells play a tumor-promoting role in GBM through the production of IL-17, which activates TANs/MDSCs.

## Results

### MAIT cells are decreased in peripheral blood from patients with GBM

To investigate if MAIT cells change in quantity and quality in GBM patients, we quantified the MAIT cell frequency (Vα7.2^+^CD161^+^ T cells) in the peripheral blood of GBM patients by flow cytometry (Extended data Figure 1A). The frequency of circulating MAIT cells among T cells in GBM (median % of total T cells = 0.45%) was significantly lower than that of the healthy controls (median % of total T cells = 2%), suggesting a preferential reduction of MAIT cells in the circulation of GBM patients (Figure 1A). To examine the functions of MAIT cells, PBMCs were stimulated with PMA and ionomycin and stained for TNF-α, IL-17, IFN-γ, and IL-4. We could not detect any IL-4-producing MAIT cells. The frequencies of MAIT cells that are single producer of IFN-γ or TNF-α (IFN-γ ^+^IL-17^-^TNF-α^-^ or IFN-γ ^-^IL-17^-^TNF-α^+^) were significantly higher in the healthy donor samples than in the GBM patients (Figure 1B). Interestingly, the frequencies of MAIT cells producing both IL-17 and IFN-γ were significantly higher in the GBM patients regardless of the TNF-α production status (IFN-γ^+^IL-17^+^TNF-α^-^ or IFN-γ^+^IL-17^+^TNF-α^+^). These results showed a decrease in circulating MAIT cells in GBM patients compared to healthy donors, and the MAIT cells from GBM patients tend to produce more IL-17.

**Figure 1.**
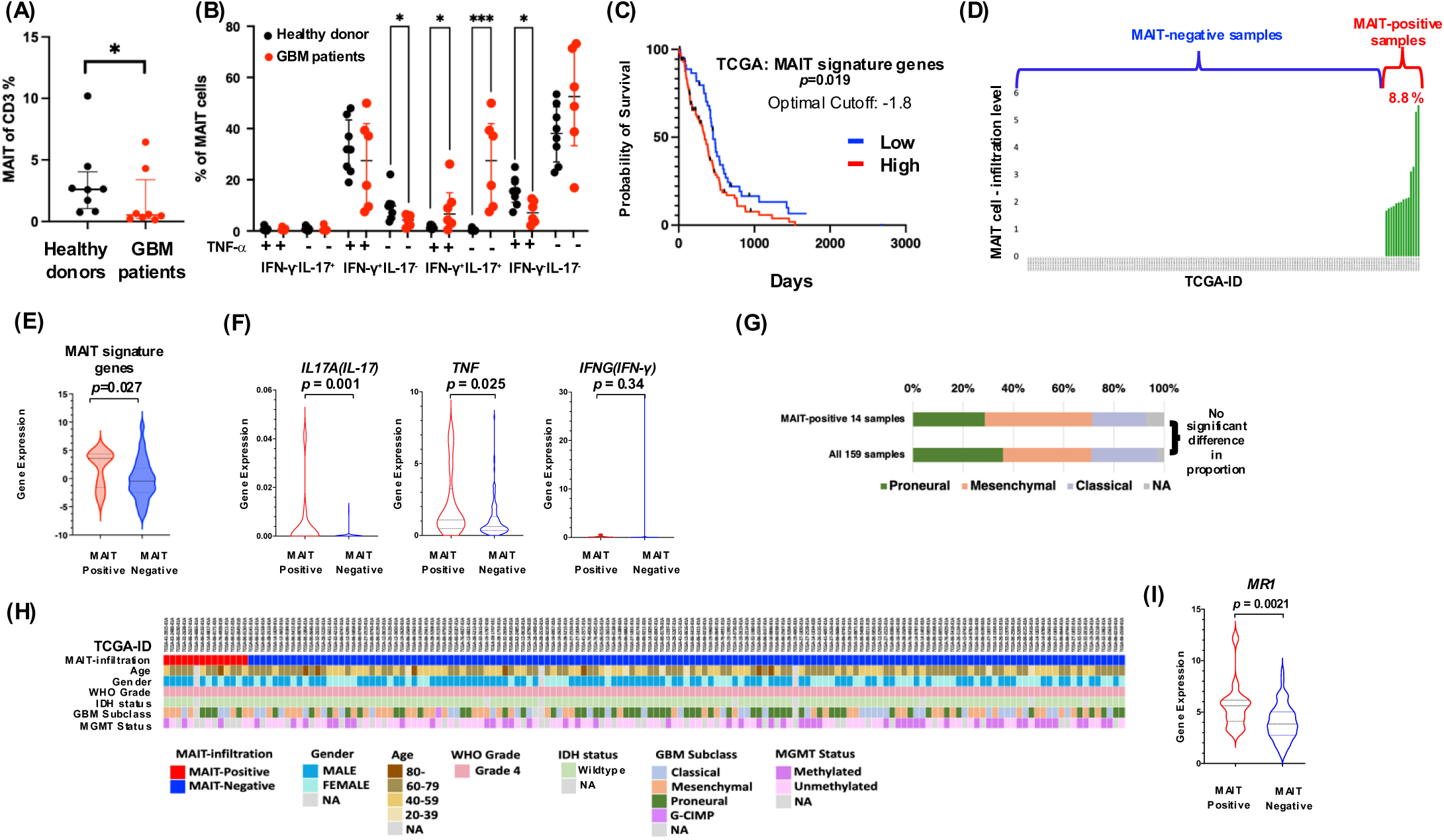
MAIT cells in GBM patient samples and their correlation with clinical prognosis. **(A)** MAIT cell frequency in T cells of GBM patients’ PBMC was compared with healthy control. Statistical significance was calculated using two-way ANOVA. ***p < .001, *p < .01. (B) Percentage of PBMC-derived MAIT cells secreting IFN-γ, IL17 or TNF-α. **(C)** The correlation between expression of MAIT cell signature genes and GBM survival in TCGA. The statistical significance of the correlation between the expression of MAIT signature genes and patient survival was tested using the Log-rank (Mantel-Cox) test. **(D)** The levels of MAIT TCR transcripts in TCGA-GBM samples. The expression of the MAIT cell signature genes **(E)** and *IL17A*, *TNF,* and *IFNG* **(F)** between MAIT positive and negative samples. **(G)** Color map showing multiple clinical/molecular indicators. Molecular indicators’ information was obtained from a previously reported database, and the neural samples were integrated into the proneural samples. **(H)** The distribution of GBM subtypes among samples. **(I)** MR1 expression in MAIT positive and –negative patient samples. *P*-values were calculated by a non-parametric Mann-Whitney test.

### MR1 expression levels correlated with worse overall survival in GBM patients

Since MAIT cells are tissue-resident cells, we focused on MAIT cells and their antigen-presenting molecule MR1 in GBM tissues. MR1 has been reported to be expressed in multiple types of cancer. However, the exact role of MR1 in cancer immunity has not been well-studied^6^. Bulk transcriptomics data of GBM patients were collected from the TCGA^23^. The samples’ *MR1* expression levels in TCGA-GBM were heterogenous (Extended data Figure 1B). We observed significant negative correlations between *MR1* expression levels and GBM survival using TCGA-GBM (Extended data Figure 1C), which agrees with the previously reported prognostic value of *MR1* expression in glioma^6^. We also investigated whether the observed association could be attributed to functional mutations in *MR1.* We found a very low frequency (0.06%) of genomic mutations in 3391 glioma samples, none of which had the reported loss of function mutation MR1-R31H^24^ (Extended data Figure 1D). Additionally, we found that the *MR1* promotor was active across GBM using H3K27ac ChIP data^25^ (Extended data Figure 1E). Given the bulk tissue nature of TCGA-GBM samples, we considered whether the *MR1* expression in a subset of the samples could be due to the contribution of micro-environment components. We utilized single-cell RNA-seq data^26^ and found that *MR1* was widely expressed in both glioma cells and TILs in GBM and that *MR1* expression was relatively upregulated in the tumor core compared to the tumor periphery (Extended data Figure 1F). All these observations indicated that intact *MR1* is widely transcribed in GBM tissue. Furthermore, the high expression of *MR1* was associated with poor prognosis in GBM patients.

### Expression levels of MAIT cell gene signature correlated with poor disease prognosis in GBM

MR1 is critical for the activation of MAIT cells. However, MR1 expression in GBM tumors is not direct evidence of MAIT cell activity because MR1 expression is necessary but not sufficient to prove MAIT cell activation. To understand the clinical significance of MAIT cells in GBM, we next conducted correlation analyses of MAIT cell signature genes, *ME1, RORC, ZBTB16, SLC4A10, MAF, KLRB1, COLQ, DUSP2, ARL4C and IL23R,* and patient survival using TCGA-GBM. These genes have been reported as markers for MAIT cells in recent transcriptomics studies^17,27–29^. The expression level of MAIT signature genes showed a negative correlation with the probability of survival in GBM patients in TCGA-GBM (Figure 1C). These suggested that MAIT cells have a negative impact on the survival of GBM patients.

### MAIT cells were identified in GBM patient tumors

Since there was minimal information on MAIT cells in GBM tissues, we sought more direct evidence for MAIT cells with the TCGA-GBM dataset. To detect MAIT cells in bulk transcriptomic data, we took advantage of the expression of a unique semi-invariant TCR α- chain by MAIT cells (details are in Methods). MAIT cells specifically express the TCR α-chain using the Vα gene segment *Vα7.2 (TRAV1-2)* in combination with the Jα gene segment *Jα33 (TRAJ33), Jα20 (TRAJ20), or Jα12 (TRAJ12)* (Supplementary Table 1). Fourteen out of the 159 (8.8%) TCGA-GBM samples were identified as MAIT-positive (Figure 1D, Supplementary Table 2, 3). The expression level of the MAIT signature genes (*ME1, RORC, ZBTB16, SLC4A10, MAF, KLRB1, COLQ, DUSP2, ARL4C,* and *IL23R*) was significantly higher in the MAIT-positive samples (Figure 1E). All MAIT cell surface receptor genes, *CCR5*, *CCR6*, *CXCR6*, *CD161*, *IL-12R,* and *IL-18R,* also had significantly higher expression levels in MAIT- positive samples (Extended data Figure 1G). Among typical cytokines produced by MAIT cells, the expression levels of *IL17A* and *TNF* were significantly higher in MAIT-positive samples, while no difference in *IFNG* expression was detected (Figure 1F). These data supported our hypothesis that samples containing transcripts of the invariant TCRα gene segment were positive for MAIT cells. The cytokine gene expression data also raised the possibility that the MAIT cell population in GBM tissue may be skewed toward MAIT17.

There was no statistically significant deviation in the distribution regarding age, gender, and MGMT status among the MAIT-positive/negative samples. The MAIT-positive samples were statistically evenly distributed in multiple GBM subtypes^30^ (Figure 1G, 1H). These findings indicate that MAIT-positive samples were not enriched in any conventional attributes of the GBM. We then found that MAIT-positive samples had a significantly higher expression level of *MR1* than MAIT-negative samples (Figure 1I). We also performed a correlation analysis of the *MR1* with the *Vα7.*2, the gene segment of the TCR α-chain expressed by MAIT cells, using previously reported scRNA-seq data of glioma patient data^31^. *MR1* showed a moderate but significant correlation with *Vα7.*2 (*correlation coefficient= 0.4, p=0.0091*) (Supplementary Table 4), suggesting that MR1 density in the tumor microenvironment may correlate with the density of *Vα7.*2^+^ MAIT cells. These findings showed that the *MR1* and MAIT cell signature genes are highly expressed in MAIT-positive patients.

To further confirm the existence of MAIT cells in GBM tissues, we performed single-cell RNA sequencing (scRNA-seq) and 5’ TCR/Gene expression analysis of CD45^+^ leukocytes from 23 GBM patient samples to identify and characterize MAIT cells in GBM tumors. Initial clustering of 89622 cells resulted in one lymphoid cluster, one myeloid cluster, and one cluster having tumor/normal brain cells (Figure 2A). The lymphoid cluster was positive for *CD3D/E/G*. The myeloid cell cluster showed high expression of myeloid cell makers, including *CD14*, *CD16, TMEM119,* and *P2RY12,* and was negative for *CD3D/E/G* expression. One cluster had a high expression of GFAP and SOX2, which could be expressed in tumor or normal brain cells.

**Figure 2.**
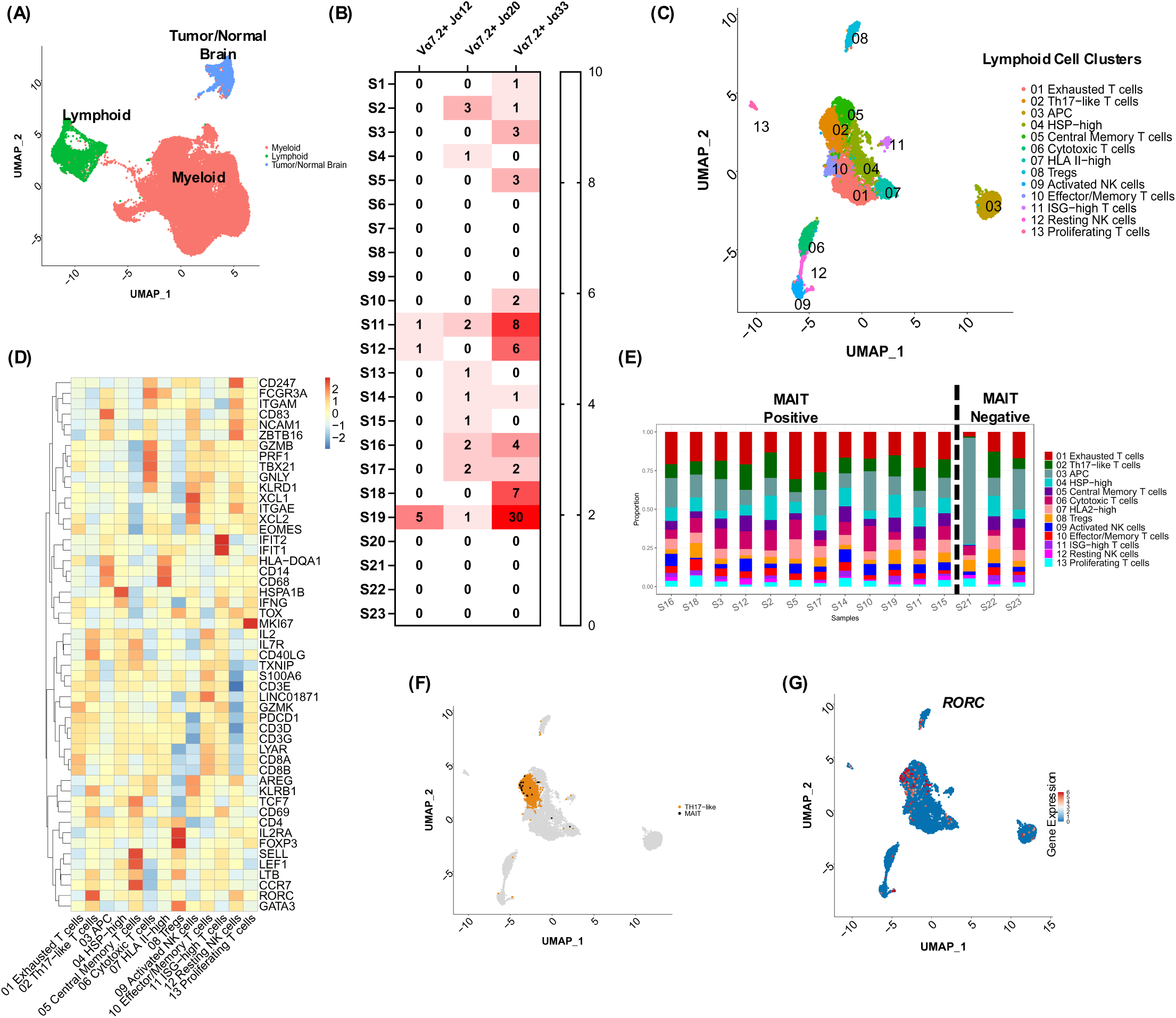
Single-cell RNA sequencing data analysis of GBM patient samples and identification of MAIT cells. Unsupervised clustering of the CD45^+^ cells from 23 samples. **(A)** UMAP of CD45^+^ cells. **(B)** Heat map of MAIT cell counts identified per sample by TCR sequencing. **(C)** UMAP of lymphoid cell population. **(D)** Heat map of marker genes for lymphoid cell clusters. **(E)** Distribution of lymphoid cell populations across samples. **(F)** MAIT cells (Black) identified by TCR sequencing on UMAP. Th17-like T cells are shown in orange. **(G)** *RORC* gene expression projected on UMAP.

MAIT cells express a unique semi-invariant TCRα chain; hence, using the TCR sequencing, we were able to identify MAIT cells. MAIT cells were identified by the expression of semi-invariant TCR gene utilizing the combinations of *V*α*7.2-J*α*33, V*α*7.2-J*α*12, or V*α*7.2-J*α*20*. In the cohort of 23 GBM patients, 15 (65.2%) patient samples contained MAIT cells (Figure 2B). Putative MAIT cells in GBM samples preferentially used the TCR chain with the combination of *V*α*7.2* and *J*α*33*. In MAIT-containing samples, we identified 89 MAIT cells (1.46% of total T cells) from 6113 T cells that successfully provided the TCR VDJ sequence. MAIT cells have limited TCRα but diverse TCRβ repertoire. The TRBV expression was varied but biased toward *TRBV6-1, TRBV6-2, TRBV6-4,* and *TRBV20-1* (Extended data Figure 2, Supplementary Table 5), similar to the previous report^32^.

### MAIT cells clustered with Th17-like T cells

We further analyzed the lymphoid cluster to delineate the lymphoid cell populations in GBM. Unsupervised clustering of the lymphoid cell population resulted in 13 subclusters (Figure 2C, 2D, Supplementary Table 6), which included: one cytotoxic T cell cluster (*GZMB*, *PRF1*, *GNLY*, *KLRD1);* one central memory T cell cluster (*IL7R*, *CD40LG*, *SELL*, *LEF1*, *CCR7)*; one exhausted T cell cluster (high *CD8A/B, TOX)*; one resting NK cell cluster (high *ZBTB16)*; one activated NK cell cluster (*XCL1, XCL2)*; one Tregs cluster (*FOXP3)*; one T cell cluster highly enriched in interferon stimulated genes (ISG) like *ISG15* and *ISG20* annotated as ISG-high T cells; one effector/memory T cell cluster which expressed markers of both effector T cells (*GZMB*, *PRF1, GNLY*) and memory T cells (*CCL5, IL7R*) (Extended data Figure 3); one proliferating T cell cluster (*MKI67*); one HSP-high cluster highly enriched in genes encoding heat shock proteins like *HSPA1B ;* and Th17-like T cell cluster (*RORC*, *KLRB1)*; one HLA II-high cluster (*HLA-DQA1, HLA-DQB1, HLA_DQB2, HLA-DMA, HLA-DMB, HLA-DRA)*; one antigen presenting cell (APC) cluster with relatively higher transcriptomic content and a high expression of MHC-II genes, *CD14*, *CD83, and CD68*. These results showed the heterogeneity of the T cell population within the GBM tumor. We could not find a significant difference in the distribution of the lymphoid subpopulations concerning the MAIT negative/positive status of GBM patients (Figure 2E).

**Figure 3.**
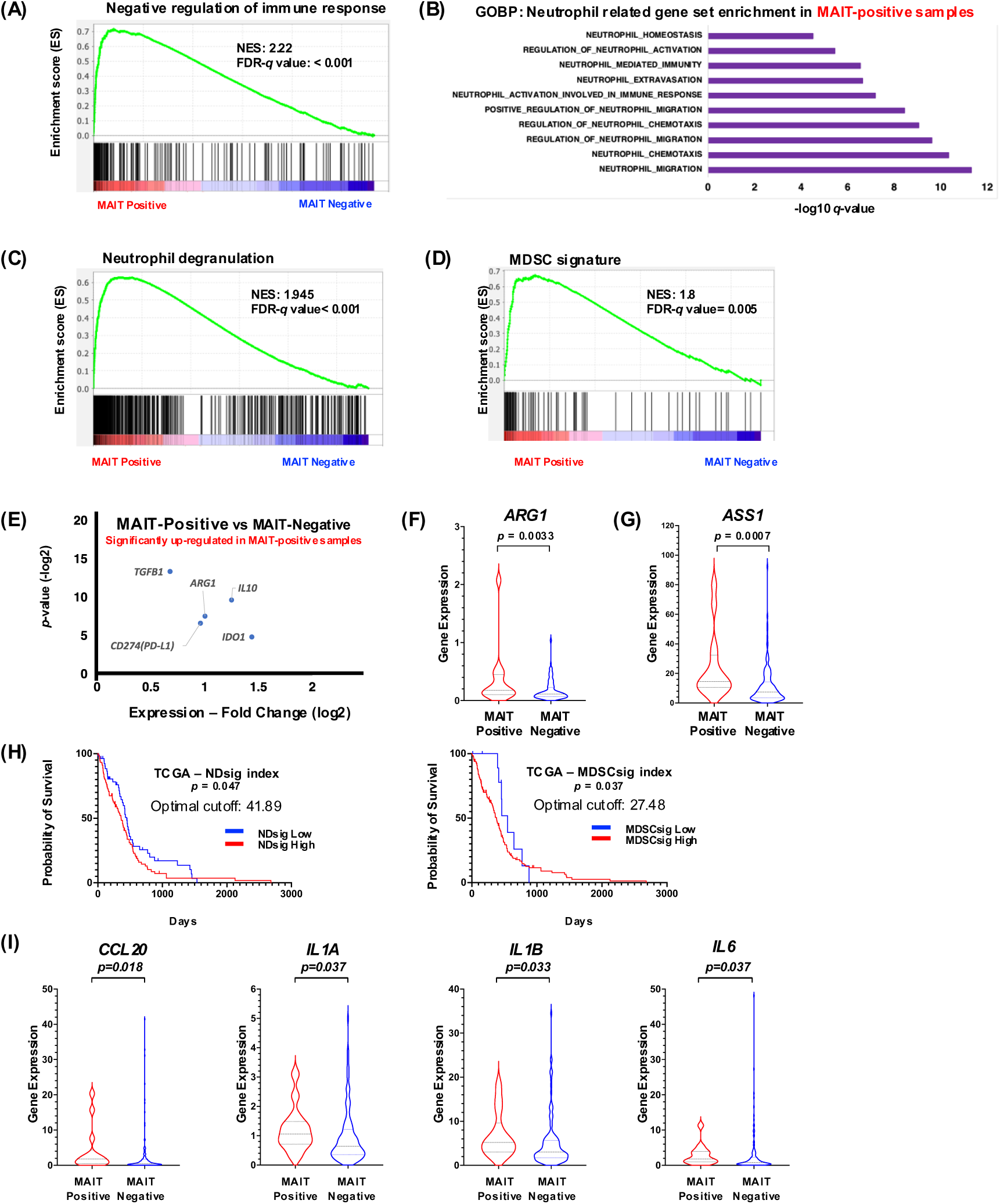
Evaluations of the activity of TANs/MDSCs in MAIT-positive/negative samples. The GSEA Enrichment score of pathways was analyzed. **(A)** Negative regulation of immune response. **(B)** Neutrophil-related genes enriched in MAIT-positive samples. **(C)** Neutrophil Degranulation and **(D)** MDSC signature. **(E)** Differential expression levels of the immunosuppressive genes linked to TANs/MDSCs. Violin plots of ARG1 **(F)** and ASS1 **(G)** in MAIT-positive/negative samples. **(H)** Prognostic correlations of NDsig-index and MDSCsig-index with TCGA-GBM clinical data. Cutoff: NDsig-index = 41.89, MDSCsig-index = 27.48. Statistical significance was tested with a Gehan- Breslow-Wilcoxon test. **(I)** Expression of *CCL20*, *IL1A*, *IL1B,* and *IL6* involved in IL17-mediated TAN/MDSC recruitment was upregulated in MAIT-positive patients. *P*-values were calculated by a non-parametric Mann-Whitney test.

Most MAIT cells identified from TCR sequencing were mapped to the Th17-like T cell cluster (Figure 2F). Cells in this cluster have a high expression of the transcription factor *RORC* that regulates *IL17* expression (Figure 2G). The cluster also showed high expression of MAIT marker genes *IL7R*, *CD161* (*KLRB1*), and *SLC4A10* compared to the rest of the lymphoid cell clusters (Extended data Figure 4). The high expression of *RORC* indicated that these cells have the potential to produce IL-17. Differential gene expression analysis showed that the Th17-like T cell cluster has a higher expression of the *KLRB1*, *RORA*, *RORC*, *CCR6,* and *CXCR6* genes expressed by IL-17-producing T cells^33^. The cells in the cluster had low expression of the transcription factor gene *TBX21* and a low expression of the cytokine *IFNG*. The identification of MAIT cells with high *RORC* and low *TBX21* expressions suggested that MAIT17 cells were a predominant MAIT cell subset in GBM tumors.

**Figure 4.**
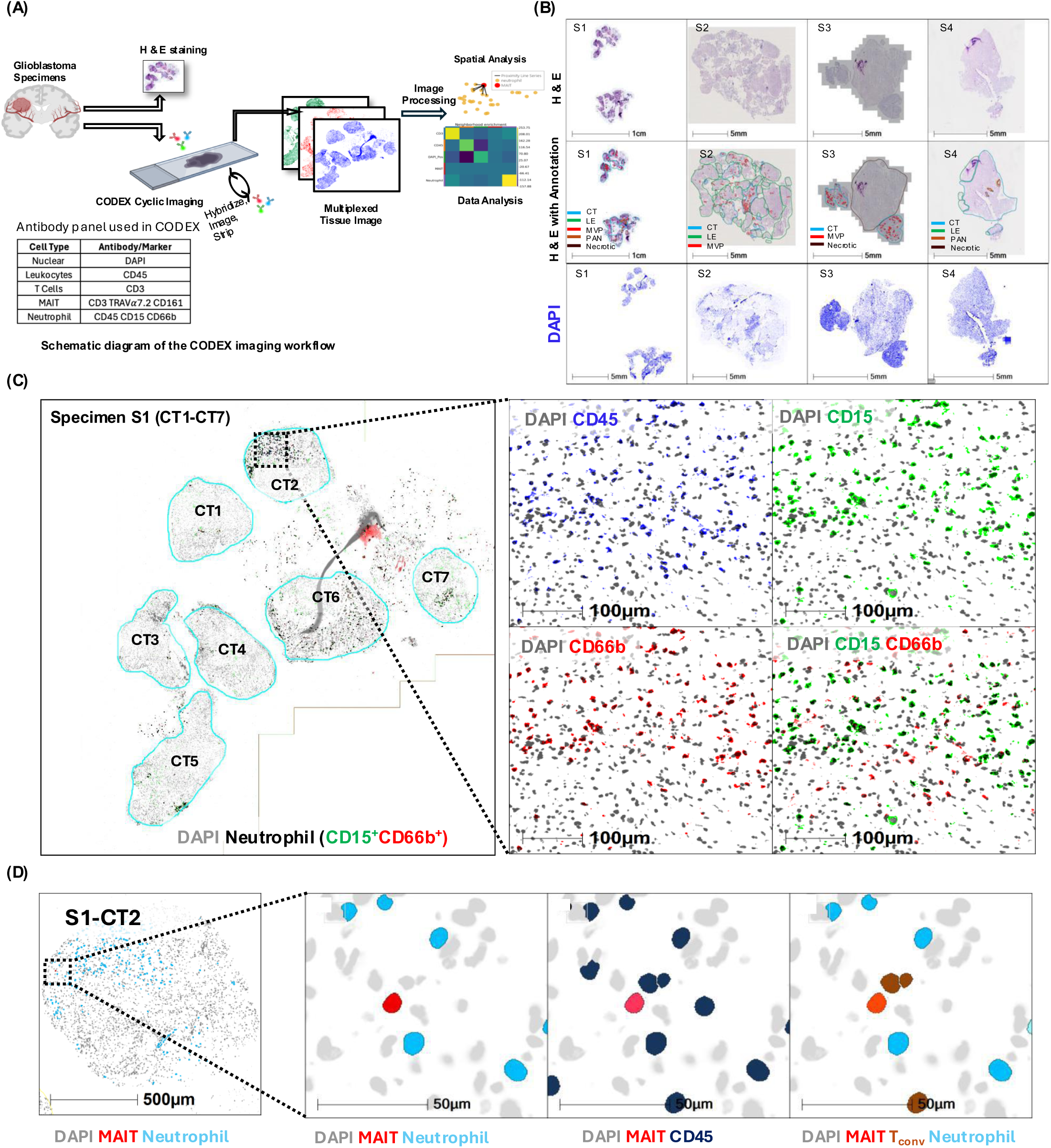
**Multiplexed Tissue Imaging of GBM specimens with CODEX technology**. **(A)** Schematic diagram showing the CODEX Phenocycler fusion imaging workflow and the antibodies used for staining. **(B)** H&E-stain slides of the 4 specimens with histopathological annotations CT, LE, MVP, PAN, and necrotic regions are shown in blue, green, red, orange and brown, respectively. The first row of the panel shows the four H&E images without annotation. The second row shows the same H&E images with the annotations. The third row shows the DAPI stain from CODEX. **(C)** Multiple CT regions (CT1-CT7) from one specimen (S1). Representative heterogeneous cellularity is shown with DAPI (grey), CD15 (green), and CD66b (red). Neutrophils are CD15^+^CD66b^+^ cells with green and red overlapped, shown in dark (black). **(D)** Neutrophil aggregation observed in S1-CT2. Neutrophil (cyan) aggregation was observed in proximity to a MAIT cell (red). Conventional T (T_conv_) cells are indicated in brown.

### Multiple pathways of TANs/MDSC-migration and activation enriched in MAIT-positive patient samples

To further elucidate the transcriptomic differences between MAIT-positive and -negative samples defined with the bulk sequencing data analysis (Figure 1D), we conducted a GSEA with the bulk RNA-seq data from TCGA-GBM. Interestingly, the most significantly enriched process among all Gene Ontology Biological Process (GOBP)-processes, including non-immunological processes in MAIT-positive samples, was an immunological process, the negative regulation of immune response (GO:0050777, Figure 3A). Since the negative regulation of immune response is an integrated pathway including many immunosuppressive processes, this significant enrichment suggests that multiple immunosuppressive mechanisms are more active in MAIT-positive samples than in MAIT-negative samples. Multiple neutrophil migration/chemotaxis and activation-related pathways were significantly enriched in MAIT- positive samples (Figure 3B). The enrichment of these gene sets in MAIT-positive samples went along with the finding that a majority of MAIT cells in GBM expressed *RORC,* suggesting their ability to produce IL-17, and IL-17 induces chemotaxis and activation of neutrophils^22^.

The biological characteristics of tumor-infiltrated neutrophils, called tumor-associated neutrophils (TANs), and blood-circulating neutrophils are significantly different^34^. Since we focused on tumor-derived transcriptomic data, all neutrophils in this study will be called TANs. Recently, it was reported that Neutrophil Degranulation (ND)-pathway genes were up-regulated in activated MDSCs^35^. Thus, the observed up-regulation of the ND-pathway can also be an indication of MDSC activation. Additionally, it is known that TANs and MDSCs share many morphological, functional, and transcriptomic characteristics and are difficult to distinguish from each other^36,37^. Therefore, the activities of TANs and MDSCs were evaluated concurrently in the current study. The signature of Neutrophil Degranulation (NDsig) and MDSC (MDSCsig) in GSEA (Supplementary Table 7) was significantly enriched in MAIT-positive samples (Figure 3C, 3D). These results indicated that TANs/MDSCs are highly activated in MAIT- positive samples compared with MAIT-negative samples.

TANs/MDSCs are known to suppress cancer immunity through various mechanisms^38,39^. The high *ARG1* expression by TANs/MDSCs depletes the local arginine^40^, and the low arginine level impairs T cells by down-regulating CD3ζ chains^41^. TANs/MDSCs also express PD-L1 in cancer, inhibiting T cell functions^42^ The up-regulation of indoleamine 2,3-dioxygenase (IDO) depletes the local tryptophan and produces kynurenine, which inhibits T cell proliferation. The IL-10 and TGF-β produced by TANs/MDSCs also suppress T cell proliferation/activity^43–45^. We found that all those genes were significantly up-regulated in MAIT-positive samples compared with MAIT-negative samples (Figure 3E). Consistent with *ARG1* up-regulation, the arginine reporter gene, *ASS1* (up-regulated upon the local arginine depletion), was up-regulated in MAIT-positive samples (Figure 3F, 3G). These data suggested that the multiple immunosuppressive pathways mediated by TANs/MDSCs were active in the MAIT-positive samples. Significantly negative correlations of TAN/MDSC activity with the overall survival indicated that their activities have an unfavorable impact on the GBM malignancy (Figure 3H).

The enrichment of genes for TANs/MDSCs in MAIT-positive patients aligns with our data, which shows that MAIT cells in GBM are MAIT17 since IL-17 is known to induce mechanisms to attract neutrophils. IL-17 triggers the production of CCL20 and G-CSF, which can synergize with other proinflammatory cytokines such as IL-6, IL-1α, and IL-1β to recruit neutrophils^22^. MAIT-positive samples were higher in the expression of the genes *CCL20, IL1A, IL1B,* and *IL6* in the TCGA-GBM data, suggesting that immunosuppressive myeloid cells could be recruited into the GBM tumor environment via an IL-17-dependent manner (Figure 3I). Altogether, these data strongly suggested that MAIT cells could activate and recruit TANs/MDSCs through IL- 17-mediated signaling, which exerts multiple immunosuppressive mechanisms in GBM.

To confirm the observation with a bulk RNA-seq dataset, TCGA-GBM, we analyzed myeloid cells from scRNA-seq data (Figure 3A). Unsupervised clustering of the myeloid cell population resulted in 13 clusters of microglia, macrophage, and DC (Extended data Figure 5A, 5B, Supplementary Table 8). We could not find a significant difference in the distribution of the myeloid cell populations in MAIT-positive and MAIT-negative patients (Extended data Figure 5C). Further clustering of the DC cell population resulted in 10 subclusters (Extended data Figure 5D, 5E, Supplementary Table 9). There was no significant difference in the distribution of the DC subpopulations between MAIT-positive and MAIT-negative groups.

**Figure 5.**
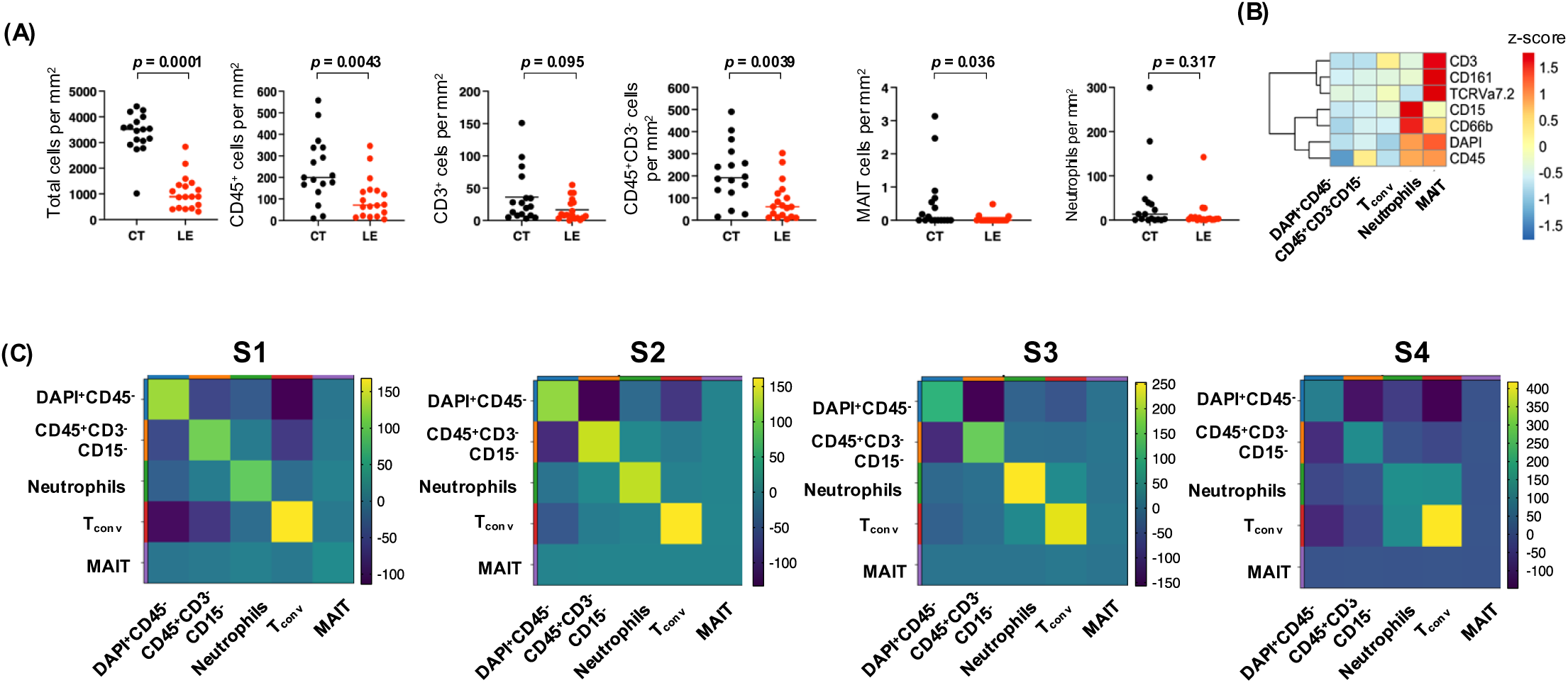
Analysis of MAIT cells and Neutrophils in GBM tissue specimens. **(A)** Comparison of cell counts normalized by section area (mm^2^) between CT and LE: total cells (DAPI^+^ cells), CD45^+^, CD3^+^, CD45^+^CD3^-^, MAIT cells, and neutrophils. A non-parametric Mann-Whitney test was performed to compare cell populations in CT and LE regions. **(B)** Hierarchical heatmap showing the intensities of the markers of five annotated cell types. **(C)** The neighborhood enrichment matrix showing the relationships between the various cell types in the four specimens **S1, S2, S3,** and **S4**. The neighborhood enrichment determines if two cell types are neighbors more often than expected by chance. Conventional T (T_conv_) cells are CD3^+^ TCR Vα7.2^-^CD161^-^ cells

We also checked the expression of neutrophil marker genes *NAMPT*, *NEAT1*, *CD29*, *PAD14,* and *LTF* in the scRNA-seq data. However, we could not find clusters with expression of neutrophil markers. Neutrophils are known to have a low transcriptomic content, expressing only 300 to 350 genes and have a short life span, which results in low representation in sequencing^46–48^. We have also tried implementing the recommended protocol to improve neutrophil capturing by including low-UMI barcodes; however, even with this technique, we could not detect neutrophils.

### Spatial analysis of MAIT cells and neutrophils in GBM tissues

From our scRNA-seq data, we have shown evidence of MAIT cells in the GBM tumor. Subsequently, we have shown a potential MAIT-TAN/MDSC association and how this could play an immunosuppressive role in GBM. However, direct detection of the neutrophil was challenging as neutrophil transcripts could not be detected in the scRNA-seq data.

Next, we performed multiplexed tissue imaging with co-detection by indexing (CODEX) to study the spatial interactions between neutrophils and MAIT cells (Figure 4A). Previous classifications of glioblastoma (GBM) tissues have delineated five distinct anatomical features: cellular tumor (CT), infiltrating tumor (IT), leading edge (LE), microvascular proliferation (MVP), and palisading around necrosis (PAN) ^49^. However, the area within the MVP and IT are relatively small and did not harbor MAIT cells (Supplementary Table 10,11, 12). Therefore, we focus our analysis on the CT, LE, and PAN tumor regions. This resulted in 36 annotated tissue regions consisting of 17 CT, 18 LE, and 1 PAN from four specimens.

There was a high degree of heterogeneity among four GBM specimens (Figure 4B). Interestingly, we observed a high intratumor heterogeneity when we looked at the neutrophil markers (CD15, CD66b). Neutrophils (CD45^+^CD15^+^CD66b^+^), when found in GBM tissues, tend to cluster together in specific regions; however, these neutrophil clusters were rare events and were only observed in the specific regions of the GBM tissues (Figure 4C). We found MAIT cells in all the four specimens we analyzed (Extended data Figure 6). The number of MAIT cells (CD45^+^CD3^+^TCR Vα7.2^+^CD161^+^) in the tumor tissue was low, with the highest number of MAIT cells (9 cells) found in the PAN. Further analysis showed that neutrophils and MAIT cells could be in proximity and directly interact (Figure 4D). MAIT cells can secrete multiple cytokines, including IL-17, and could mediate the activation of neutrophils. The observation of the neutrophils near MAIT cells suggests that MAIT cells could directly interact with neutrophils or indirectly regulate the activation of neutrophils through secreted cytokines.

**Figure 6.**
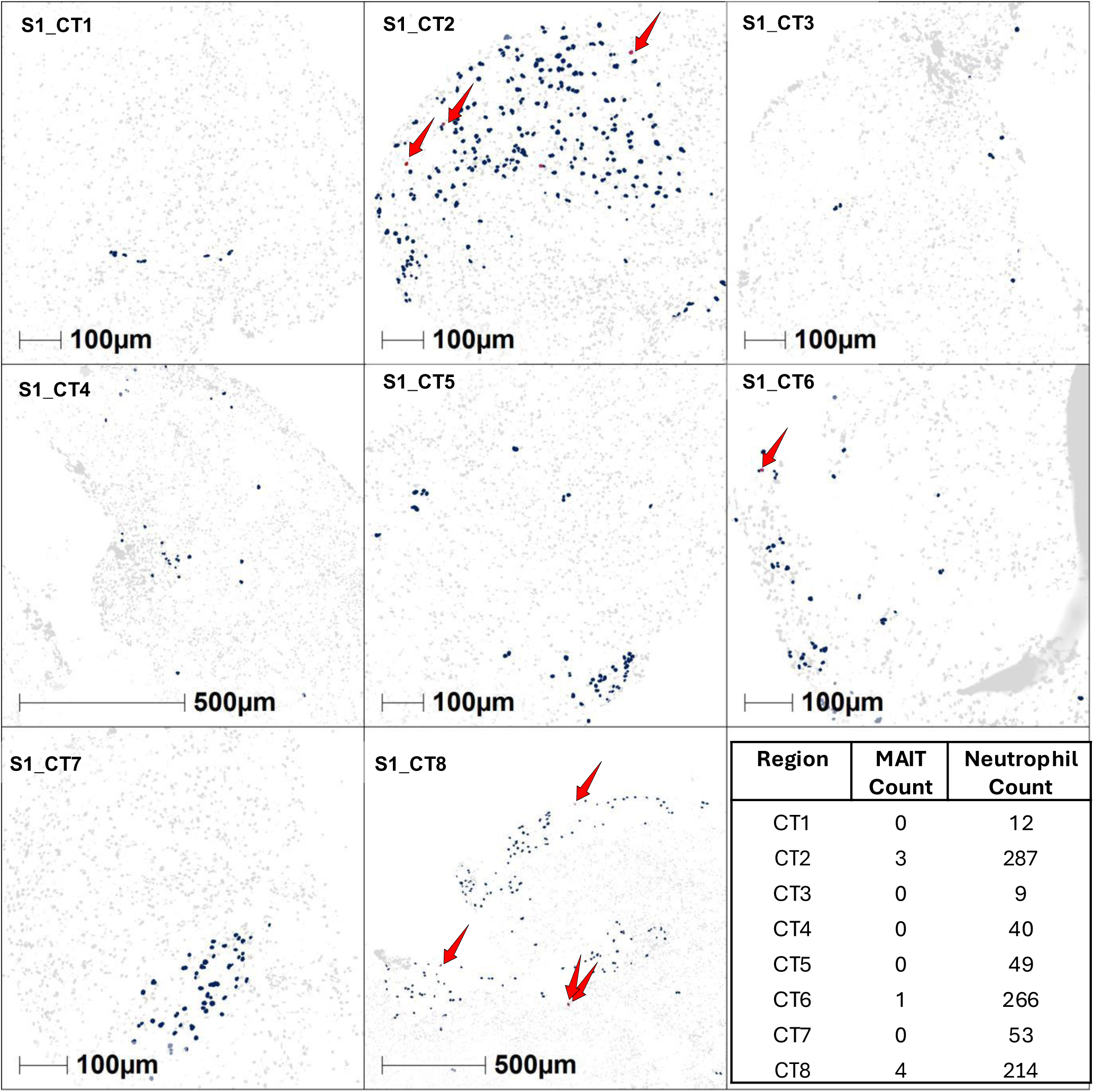
Heterogeneous distribution of Neutrophils across CT regions. MAIT cells and neutrophils are shown from multiple CT regions of specimen S1. CT2, CT6, and CT8 are positive for MAIT cells and showed a higher number of neutrophils. MAIT cells are shown with red arrows. DAPI stain is shown in grey. MAIT cells and neutrophils are indicated in red and blue, respectively.

GBM is characterized by high intra- and inter-tumor heterogeneity in cell type and composition. Hence, we next analyzed the cellular composition of various tumors to investigate cells enriched in the CT, LE, and PAN tumor regions. 280,302 cells were analyzed from CT, LE, and PAN regions. CT regions had a higher number of total DAPI^+^ cells and CD45^+^ leukocytes than LE regions (Figure 5A). Within the CD45^+^ cells, we observed a higher population of the CD45^+^CD3^-^ cells in CT regions. CT regions showed a higher MAIT cell enrichment than LEs (*p*=0.036); however, there was no difference in the neutrophil count between the two groups. Only one PAN region was positive for MAIT cells; hence, it was not included in the statistical analysis. These results showed that CT, known to have a higher population of neoplastic tumor cells than LE, also harbors a higher number of CD45^+^ leukocytes. Previous studies of GBM showed that the tumor core has a higher enrichment of the immune-suppressive macrophages and microglia^50^. Although we did not stain macrophage and microglia cell markers, a higher number of the CD45^+^CD3^-^ cells in the CT could be due to the high number of immune-suppressive macrophage and microglia populations.

Spatial relationships between cell types can be established by utilizing the x/y coordinates of each cell in the tissue. We used Spatial Single-Cell Datasets (SPAC) workflow for data processing the marker intensities and spatial analysis^51^. Analyzing the cellular neighborhoods can reveal the spatial organization of the tumor microenvironment, which could be used to investigate cell types that preferentially cluster together. Neighborhood enrichment analysis was carried out to explore the spatial relationship between MAIT cells and neutrophils (Figure 5B, 5C). In our analysis, due to the low MAIT cell count, we did not observe a pattern in the neighborhood pairing with MAIT cells. Consequently, we could not make deductions on the likelihood of MAIT-neutrophil pairings. Briefly, cells from the same cell type tend to form neighboring pairs with the same cell types. The conventional T cells (T_conv_, CD3^+^TCR Va7.2^-^ CD161^-^) showed the highest neighboring pairs with T_conv_ cells, suggesting the tendency of these cells to be close neighbors. Similarly, the DAPI^+^CD45^-^ and CD45^+^CD3^-^ cells showed high neighboring pairs between the same cell types. The pattern of neighboring pairs between the same cell type was observed in all four samples. Although cellular and tissue compositions may vary between samples, the cells of the same type tend to be neighbors to each other.

The spatial analysis of MAIT and neutrophils was challenging due to the low MAIT cell count, and we did not observe a direct correlation between the number of MAIT cells and neutrophils. It is widely accepted that GBM is characterized by high tumor heterogeneity, and various factors, including treatment history could change the cellularity of the tissue. Due to the small sample size, we could not dichotomize the specimens to account for all the confounds in our analysis. Therefore, a MAIT-neutrophil association could not be ruled out. Despite the variability between specimens observed in GBM, analyzing each specimen individually can yield vital insights into the tumor’s cellular composition. Thus, it is crucial to conduct specimen-specific analyses. Interestingly, in one GBM sample (specimen S1), we found multiple neutrophil clusters at various CT regions of the tissue. When we analyzed the MAIT cells and neutrophils in this specimen, the regions with MAIT cells showed a relatively higher number of neutrophils (Figure 6), and MAIT cell counts correlated with neutrophils (spearman r = 0.73, p=0.035).

We also estimated the propensity of MAIT cells and neutrophils to cluster together using Ripley L statistics. When we applied these to the CT8 region from specimen S1, we found that the MAIT cells and neutrophils showed a higher Ripley L statistic value compared to the value from random simulation at a radius of approximately 20 pixels (the average diameter of a cell is approximately 15-16 pixels) demonstrating clustering between MAIT cells and neutrophils (Extended data Figure 7).

**Figure 7.**
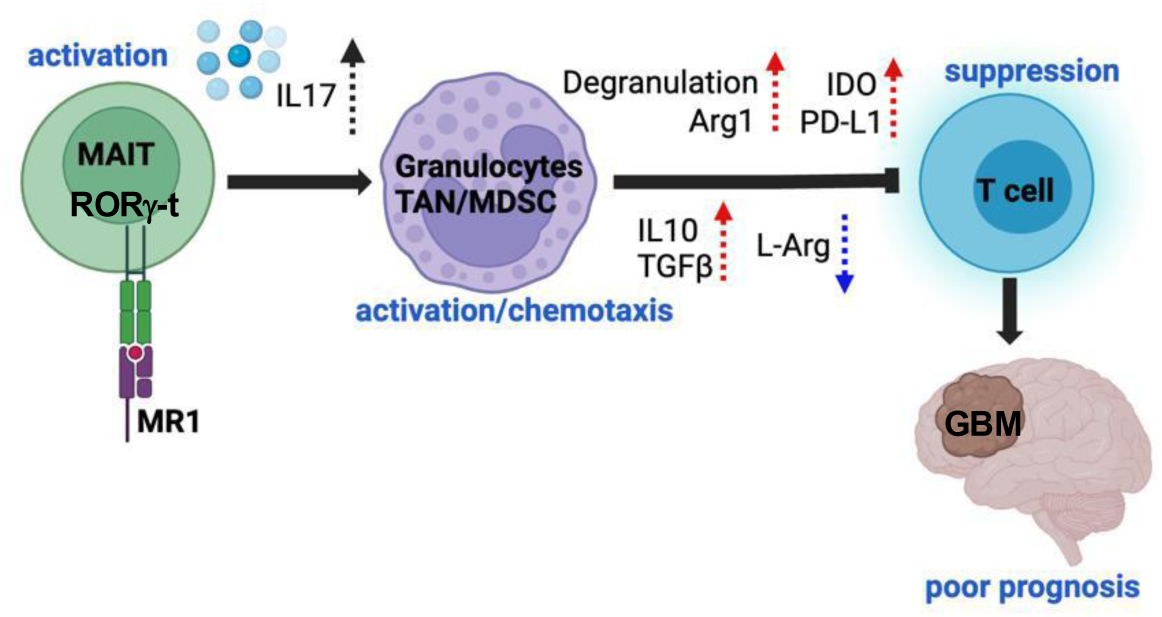
MAIT cells in GBM have a potential to produce IL17 to activate immunosuppressive TAN/MDSC. RORγ-t expressing MAIT cells (MAIT17) enriched in GBM could produce IL-17, which activates TANs and MDSCs. Activated TANs and MDSCs express multiple immune suppressive molecules such as Arg-1, IDO, PD-L1, IL-10, and TGF-β that suppress T cells, leading to a poor prognosis of the disease.

## Discussion

The current study found that MAIT gene signature expression levels correlated with poor outcomes in GBM patients. We identified MAIT cells in the GBM tissue by a single-cell TCR VDJ sequencing and found that more than half of GBM patient tumor tissues contained MAIT cells and by a multiplex tissue staining. The immunological role of MAIT cells in cancer is unsettled given that MAIT cells either promote or suppress tumor growth, which varies depending on the cancer type. In the context of GBM, a limited number of studies on MAIT cells have been conducted. MAIT TCR transcripts were reported in GBM samples^52^, and MAIT cells have the potential to lyse GBM in the presence of 5-OP-RU^53^. To our knowledge, this is the first study demonstrating the existence of MAIT cells in GBM tissue related to the disease prognosis.

The distribution of functionally distinct MAIT cell subtypes varies across different tissues^54^ Interestingly, most MAIT cells in GBM tissue expressed *RORC* (encoding RORγ-t), indicating that they were MAIT17 (Figure 2G). Corroborating with the data from scRNA-seq, we also found with TCGA-GBM data analysis that the expression levels of *IL17A* and *TNF* were significantly higher in MAIT-positive samples, while there was no statistical difference in *IFNG* expression between MAIT-positive and MAIT-negative samples. The skewing of MAIT function toward IL-17 production in cancer has been reported in other studies. Yan et al., reported that tumor-infiltrating MAIT cells enhanced the production of IL-17 and TNF-a while the IFN-γ production level was not changed^18^. In colon adenocarcinomas, MAIT cells showed reduced IFN-γ production^55^. In hepatocellular carcinoma (HCC), dysfunctional MAIT cells were reported with altered transcriptional factor profiles, switching MAIT cells from a Th1-like to Th17-like phenotype with increased RORγ-t expression and a significantly reduced T-bet expression^17^. In multiple myeloma (MM), decreased MAIT proportion in PBMC compared with healthy control and a decrease in effector functional capacity of circulating MAIT cells, characterized by reduced IFN-γ production, was reported^14^.

The skewing mechanism of the MAIT subsets is not well understood; however, the decrease in IFN-γ expression and increased IL-17 could have various implications for cancer. An effective mechanism for cancer to evade immune response is the dysfunction of the effector T-cells^56^. Tumor cells can employ multiple mechanisms to suppress IFN-γ signaling and render T cells ineffective^57^. In murine models, MAIT cell dysfunction characterized by high expression of PD-1 and TIM-3 was reported^17^. These MAIT cells showed significantly lower expression of IFN-γ. Over time, the increase in the dysfunction of MAIT cells was accompanied by loss of IFN-γ production. Therefore, the decrease in the IFN-γ expression in MAIT cells in GBM could be a sign of the dysfunction of these cells. IL-17 is a cytokine associated with the development and pro-tumor function of immuno-suppressive myeloid cells^58–60^. Taken together, the skewing of MAIT cell function could effectively promote immune suppression in GBM.

We found that the frequency of MAIT cells among T cells in GBM tissues that contain MAIT cells was 1.46%. In contrast, the frequency of MAIT cells among T cells in peripheral blood samples of GBM patients was 0.45% (median frequency from 8 patients), significantly lower than that of healthy donors. Although the GBM tissue samples and peripheral blood samples were from different sets of GBM patients, this observation strongly suggested the enrichment of MAIT cells in GBM tissues. Like our observation made in this study, the significant reduction of MAIT cells in the circulation in cancer patients has been reported in other types of cancer, including colorectal, lung, and liver^19,61,62^. In colorectal cancer, a decrease in circulating MAIT cells was associated with increased infiltrated MAIT cells^20,62^. More interestingly, the frequency of MAIT cells in cancer tissues correlated with the advancement of the disease in colorectal cancer patients^62^. Together with our observation of the negative correlations between MAIT signature gene expression levels and patient survival in the TCGA-GBM data set, MAIT cells in cancer tissues may contribute to forming an immunosuppressive tumor microenvironment in certain types of cancer.

The level of TAN infiltration has been reported to correlate with the grades of glioma, and some tumor-promoting mechanisms by TANs in GBM have been suggested^63,64^. However, the mechanisms attracting TANs to the GBM tissue and their activation mechanisms still need to be better understood. Our study observed a significant correlation between the expression level of MAIT cell signature genes and the activation level of TANs/MDSCs. MAIT cells rapidly secrete a copious amount of cytokines, including IL-17^9^, which is important in this context as IL-17 promotes the local attraction and activation of TANs through the induction of chemokines^12,13^. We observed that MAIT cells express *RORC* by scRNA-seq and a significant up-regulation of *IL17A* in MAIT-positive samples in TCGA-GBM data. Concurrently, the activation of TANs/MDSCs and multiple neutrophil attraction pathways were up-regulated in MAIT-positive samples. These observations suggested the contribution of MAIT cells to recruit TANs in GBM tissues.

The transcriptomic quantification of neutrophils with RNA-seq was challenging. Hence, we applied multiplexed tissue imaging to study the MAIT cells and neutrophils in GBM tumor tissue. The tissue imaging of the GBM tumor tissue from four patient samples provided several important insights. While the number of MAIT cells in the GBM tumor tissues was low, neutrophils were present in significant numbers, exhibiting a pattern of localized enrichment throughout the tissue. Although statistical significance could not be established due to the low number of samples we examined, our observation showed a pattern of higher MAIT cells in the CT regions of GBM tissue. In specimen S1 with multiple CT regions (CT1-CT8), CT regions with MAIT cells showed higher neutrophil enrichment, which may suggest an association between MAIT cells and neutrophil enrichment (Figure 6). The simultaneous presence of MAIT cells and granulocytes has been documented in hepatocellular carcinoma^65^. However, to our knowledge, this is the first study to report the coexistence of MAIT cells and neutrophils in GBM tissues.

Neutrophils can exert strong cytotoxicity; therefore, they may play anti-tumor roles. However, the biological function of neutrophils can be modified by the tissue environment, especially by the local TGF-β status. Under a TGF-β depleted environment, neutrophils tend to acquire the anti-tumor phenotype, called N1, whereas, under the TGF-β sufficient environment, they tend to acquire the pro-tumor phenotype, N2^39^. The immunosuppressive and tumor-promoting role of N2-TAN has been reported in multiple cancer types. Since the expression level of *TGFB1* in MAIT-positive samples was significantly up-regulated (Figure 3E), the TME of MAIT-positive patients is the preferred environment for TANs to acquire N2 phenotype. Thus, the TANs in MAIT-positive GBM samples are considered to play a relatively immunosuppressive role and can cause GBM’s poor prognosis.

MDSCs suppress cancer immunity^38^. In GBM, their immunosuppressive role and the correlation between their infiltration and poor survival have been reported^66^. He et al., showed that MDSCs have phenotypic plasticity in their immunosuppressive function using *in vivo* cancer model, and IL-17 was required for the development of MDSCs in tumor-bearing mice and for exerting MDSC’s pro-tumor functions^60^. Guan et al., also observed a significant correlation between the tumor IL-17 level and the MDSC infiltration level in renal cancer^67^. These reports support our hypothesis that the IL-17 secreted from MAIT cells can attract MDSCs and promote their immunosuppressive function in GBM.

Although we found MAIT cells in patients’ GBM tissues, it is unclear how they get activated in the cancer tissue to express their functions. MAIT cells recognize bacteria metabolites, riboflavin. Recently, Naghavian et al. reported that bacteria-specific peptide antigens were presented in GBM tissues^68^. This study raises the possibility that MAIT cells in GBM could be activated by bacteria metabolites that could reach the brain. Antigens recognized by MAIT cells are not fully understood. A study by Zhang et al., reporting the role of MAIT cells residing in the meninges contributing to memory formation by producing antioxidant molecules, suggested that MAIT cells get activated by recognizing unknown antigens in the brain^69^. MAIT cells in GBM may recognize the same antigen to express their function.

In conclusion, we propose a novel MAIT-TAN/MDSC pathway that could contribute to the poor prognosis of GBM (Figure 7). MAIT cells may be an underappreciated link between GBM and TAN/MDSC’s activation and infiltration. The current study sheds light on the possible immunosuppressive role of MAIT cells in GBM, suggesting that modulatory strategies may enhance the efficacy of immunotherapies.

## Methods

### PBMC collection

Blood samples were collected from patients with newly diagnosed GBM recruited as part of a clinical trial at the Neuro-Oncology Branch at the National Cancer Institute [NCT04817254]. GBM patients’ blood-derived Peripheral blood mononuclear cells (PBMCs) were collected using CPT tubes (BD Vacutainer CPT with sodium citrate, cat #362761) and stored in a liquid nitrogen freezer prior to conducting assays.

PBMCs from healthy donors were isolated from buffy coat samples obtained from the NIH Blood Bank using Ficoll-Paque Plus (Millipore Sigma). All patients and healthy donors signed an institutional review board–approved, written informed consent form for collection of blood samples.

### PBMC Sample preparation and cytokine stimulation

PBMCs were stimulated as a single-cell suspension at a concentration of 4-5 x 10^6^ cells/mL with the Cell Activation Cocktail (Biolegend, cat # 423301) and incubated at 37°C with 5% CO_2_. Brefeldin A was added after 1 hr, and cells were further incubated at 37°C with 5% CO_2_ for an additional 5 hrs before the harvest for flow cytometry.

### Flow cytometry

Approximately 2 x 10^6^ cells per sample were stained with LIVE/DEAD™ Fixable Aqua Dead Cell Stain Kit according to manufacturer’s instructions (Invitrogen LIVE/DEAD™ Fixable Aqua Dead Cell Stain Kit, for 405 nm excitation, Thermo Fischer) for 30 min at room temperature (RT) in the dark. Cells were then washed (5 min, 350 x *g*) with HBSS containing 0.1% bovine serum albumin and 0.05% sodium azide. Cells were subsequently incubated with a cocktail containing Human TruStainFcX (Biolegend) and 5-OP-RU-loaded MR1 tetramers (NIH Tetramer Facility) for 20 mins and subsequently stained with the surface antibodies against CD3 (Biolegend cat #: 300324), CD4 (BD cat #:611887), CD8 (BD cat #: 563795), Vα7.2 (Biolegend cat #: 351730), CD161 (Biolegend cat #: 339916) (for 30 mins at 4°C. Cells were washed twice with HBSS containing 0.1% bovine serum albumin and 0.05% sodium azide. Intracellular cytokine staining (ICS) was performed according to the manufacturer’s instructions (Biolegend cat#420801; Cat#421022) and stained with antibodies specific for IFN-γ (Biolegend cat #: 502538), TNF-α (Biolegend cat #: 502912), IL-4 (Biolegend cat #: 500818), and IL-17A (Biolegend cat #: 512322). Samples were analyzed using the Cytek®Aurora flow cytometer (Cytek Biosciences, Fremont, CA, USA).

### The definition of GBM in this study

With WHO classification 2021, isocitrate dehydrogenase (IDH) mutant grade 4 astrocytoma was excluded from the previously defined “glioblastoma”^70^. Therefore, we removed data from patients with mutant IDH from the TCGA dataset used.

### Gene expression analysis and the predictive examination for GBM

All bulk RNA-seq data analyzed in this study are in fragments per kilobase of transcript per million mapped fragments (FPKM) unit. The TCGA-GBM core 159 samples’ (Supplementary Table 2) transcriptomic data was downloaded from the Genomic Data Commons Data Portal (https://portal.gdc.cancer.gov/). All the transcriptomic analyses were conducted using this set of data. Prognostic examinations were done with the data of TCGA-GBM 144 samples (Supplementary Table 3) with available clinical data using the Log-rank test, and all cut-off thresholds were decided using the TCGA survival data analyzer^71^. For TCGA analysis, transcriptomic/clinical data were obtained from The Human Protein Atlas^72^. When comparing specific gene expressions between two groups, a two-tailed unpaired Student’s t-test was used, and *p* < 0.05 was considered significant. When fold changes in gene expression were calculated, the mean FPKM of both groups was used.

### Survival analysis of MAIT signature genes

MAIT signature genes were reported from recent bulk and scRNA-seq studies ^73–76^. From the reported genes, 10 MAIT signature genes (*ME1, RORC, ZBTB16, SLC4A10, MAF, KLRB1, COLQ, DUSP2, ARL4C,* and *IL23R*) which were highly differentially expressed in MAIT cells were selected. Of the selected 10 genes, seven genes were reported as highly differentially expressed in MAIT cells in two or more independent studies. The expression value of each gene in FPKM was scaled across samples using the R base scale method. This was followed by summing up the expression values of the 10 MAIT signature genes for each sample. The samples were bifurcated into a MAIT-high or MAIT-low group based on the expression value of the aggregated MAIT signature genes. The optimal cutoff is defined as the point with the most significant (log-rank test) split using the *CutoffFinder*^71^. The survival analysis with Cox proportional hazard regression was performed on Prism.

### Detecting the MAIT-positive samples in TCGA-GBM

All FASTQ files were downloaded from the GDC-Data Portal Legacy Archive (https://portal.gdc.cancer.gov/legacy-archive/). Reads with the characteristic T cell receptor sequences (Supplementary Table 1) of MAIT cells were searched with the FASTQ files. The samples containing the reads of TRAV1-2(Vα7.2) and TRAJ33(Jα33) or TRAJ12(Jα12) or TRAJ20(Jα20) were classified as the samples with the MAIT cell infiltration and were defined as MAIT-positive. The MAIT-infiltration level of each sample was calculated with the formula given below:

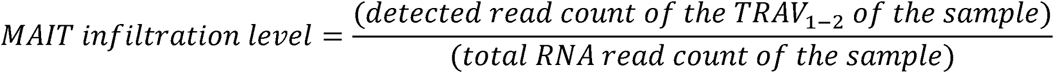

All these analyses were conducted on the NIH-high performance computer, the Biowulf cluster system.

It was confirmed that all the searched sequences (Supplementary Table 1) were unique in the whole genome, and no other loci have the same sequence using the GGGenome-ultrafast sequence search (https://gggenome.dbcls.jp/en/).

### Analyses using the signature of neutrophil degranulation and MDSC

The gene set of Neutrophil degranulation (REACTOME: R-HSA-6798695, 479 genes) was used to generate the neutrophil degranulation signature genes. The MDSC characteristic signature gene set was obtained from a previous study^77^. Since the gene set was originally composed of 105 murine genes, we extracted 80 genes from them that exist in humans, and the 80 genes were defined as MDSC signature genes (Supplementary Table 7).

### Gene expression heatmap and Gene Set Enrichment Analysis (GSEA)

GSEAs of this study were conducted using the expression data of core 159 TCGA-GBM samples. GSEA_4.2.3 was used to run the 1000 permutation analyses using the signatures of c2.cp. Reactome.v7.5.1.symbols.gmt, c5.go.bp.v7.5.1.symbols.gmt, and c2.cp.kegg.v7.5.1.symbols.gmt^78,79^. GSEA with the signature of neutrophil degranulation and MDSC was conducted using our customized “gmx” signature files made from their characteristic signatures.

### Identification of MR1 mutational frequency, the epigenetic transcriptional status, and the transcriptional distribution in GBM tissue

The mutational frequency was confirmed using *cBioportal* (https://www.cbioportal.org/). A total of 3391 multi-grade glioma samples were examined. The H3K27ac ChIP data of GSM1824806/1824808 were used for the ChIP analysis. Single-cell RNA-sequencing (RNA-seq) analyzing web tool, GBM-seq, was used to generate the tSNE plot of *MR1* expression in GBM.

### Examining correlation between MR1 and TRAV1-2 gene expression in scRNA-seq data from glioma patients

We analyzed the 10X Genomics 3’ scRNA-seq dataset from Abdelfattah et al., ^31^, who sequenced 44 tumor fragment samples from 18 patients with low-grade glioma, newly diagnosed GBM, and recurrent GBM (GEO: GSE182109). Out of the 201986 cell barcodes that pass quality control, the authors labeled 19570 of them as T cells (*CD3E^+^*cells). From all samples’ feature-barcode matrices output by *CellRanger*, we extracted the number of non-T cells expressing MR1 and T cells expressing TRAV1-2 into two-column vectors (Supplementary Table 4). We used the latter as a proxy for the count of MAIT cells since the expression of TRAJ12/33/20 could not be captured with the limited resolution of the 3’ dataset. The number of cells expressing the *Vα7.2* gene was adjusted based on the total number of T cells in the sample, and the number of cells expressing the *MR1* gene was adjusted based on the total number of non-T cells (Supplementary Table 4). We then computed Spearman’s rank correlation coefficient between the two vectors.

### GBM patient tissue collection and single-cell isolation

The UCLA Medical Institutional Review Board 2 (IRB#10-000655-AM-00059) approved all protocols related to patient tumor specimen collection at UCLA. All GBM patients (newly diagnosed and recurrent) were treated with standard-of-care therapies. All patients provided written informed consent, and this study was conducted under an institutional review board-approved protocol. Patients with recurrent GBMs who received neoadjuvant anti-PD1 treatment were given an intravenous infusion of off-label, off-trial pembrolizumab 14 +/- 5 days before surgery, similar to a previously published study^80^.

The GBM patient cohort comprised nine patients newly diagnosed with GBM, seven patients with recurrent GBM who have not been treated with any immunotherapy, and seven recurrent GBM patients who were treated with neoadjuvant anti-PD1 therapy (Supplementary Table 5).

Tumor-infiltrating CD45^+^ immune cell populations were isolated from GBM tumors of patients undergoing surgical resection at the University of California, Los Angeles. Resected tumor tissue not needed for diagnosis was digested using the Miltenyi Brain Tumor Dissociation kit (Miltenyi Biotec, cat. 130-095-42) and gentleMACS dissociator (Miltenyi Biotec, cat. 130-093- 235). After treatment with Myelin Removal Beads II (Miltenyi Biotec, cat. 130-096-433), cells were labeled with CD45^+^ microbeads (Miltenyi Biotec, cat. 130-045-801). CD45^+^ cells were positively selected from the tumor digest with Miltenyi LS columns (Miltenyi Biotec, cat. 130- 042-401) and MidiMACS separator (Miltenyi Biotec, cat. 130-042-302) and immediately frozen for batched analysis in Bambanker (Fisher Scientific, cat. 302-14681) and stored in liquid nitrogen.

### Single-cell library preparation and sequencing

On the day of single-cell library preparation, CD45^+^ cells collected from GBM patient tumors were thawed, counted, and resuspended in PBS with 0.4% BSA (Gemini, cat. 700-101P- 100). Single-cell library construction was performed by the Technology Center for Genomics and Bioinformatics Core, UCLA, utilizing the Chromium single-cell 5′ and VDJ library construction (10x Genomics), per the manufacturer’s protocol. The constructed single-cell libraries were sequenced on a Novaseq S4 2x100bp flow cell (Illumina).

### Single-cell RNA-seq data processing and analysis

Data were demultiplexed and aligned with Cell Ranger version 7.1 (10X Genomics) and aligned to the Genome Reference Consortium Human Build 38 (GRCh38-2020). Transcriptions were quantified using *cellranger count* with *--force-cells: 8000* and *--include-introns: true*. *CellRanger aggr* was used to aggregate the samples for downstream analysis. TCR transcripts were quantified using *cellranger vdj.* The quality-checking reports from Cell Ranger were analyzed to assess sequencing quality and determine cell and gene counts per sample. Single-cell sequencing data were analyzed in R using the *Seurat* pipeline^81^. The preliminary analysis was conducted for each sample to filter out empty cells and doublets. Cells with a minimum of 200 genes and genes found in a minimum of 20 cells were included for downstream analysis. Cells with a percentage of mitochondrial gene counts higher than 5% of total gene counts were also excluded. After a filtering process for quality assurance, a total of 89622 cells were used for further downstream statistical analysis. Normalization was performed with *NormalizeData* and with the normalization method set to *LogNormalize* and the scaling factor set to *1e6*. The gene expression values were scaled with *ScaleData* function with variables to regress accounting for mitochondrial features, ribosomal features, and cell cycle score. Principal component analysis (PCA) was performed using *RunPCA,* with the number of principal components set to 50. The significance of the principal components was evaluated based on the ‘scree test’ method, and 30 principal components were selected. Sample integration was performed with *harmony*. Unsupervised clustering based on uniform manifold approximation and projection (UMAP) with 30 principal components was used to generate the clusters. The Seurat objects from the individual samples were integrated into one *Seurat* object.

Further analysis of the clusters was carried out by separating the cells of the clusters of interest and repeating the abovementioned analysis process. The *FindAllMarkers* function with default parameters (Wilcoxon Rank Sum test) was used to find the genes differentially expressed in each cluster. Clusters were annotated based on marker genes reported in previous studies^82,83^. The TCR VDJ profile data were also integrated with the gene expression data in R based on the matching cell barcodes. MAIT cells were identified by the expression of the TCR *a*-chain gene segments *Va7.2* and *Ja33/Ja22/Ja12*. The expression of the markers for tumor/normal, lymphoid and myeloid cells are provided in Extended data Figure 8, 9, and 10.

### Differential expression analysis

Differential expression analysis was performed using the bulk RNA-seq analysis pipeline, using limma, glimma, and edgeR, as described by Law et al., 2016 ^84^.

### Human GBM specimen for CODEX imaging

The use of human subject material was performed in accordance with the World Medical Association Declaration of Helsinki and with the approval of the National Institutes of Health Review Board (Protocol #03N0164). Neurosurgical tissue from the Surgical Neurology Branch of the National Institute of Neurological Disorders and Stroke (NINDS) was immediately frozen and blocked in optimal cutting temperature (OCT) compound and stored at -80C. For each specimen, the final diagnosis of Glioblastoma, IDH-wildtype, CNS WHO grade 4 was confirmed by a board-certified neuropathologist (PJC).

### GBM tissue multiplex IF staining with CODEX technology on Phenocycler Fusion

Tissue sections from fresh, frozen specimens collected from GBM patients were stained with a panel of six oligonucleotide-conjugated antibodies (Extended data Figure 11, Supplementary Table 10). Four specimens, which simultaneously stained for markers of both MAIT cells and neutrophil, was used for the study (Supplementary Table 11).

Freshly frozen tissue specimens from 4 GBM patients embedded in OCT were used for imaging with CODEX technology on Phenocycler Fusion (Akoya Biosciences). Eight-meter-thick tissue sections were mounted onto positively charged slides (Plus Gold) and kept at -80 °C until staining.

The tissue fixation and staining steps were performed by following the recommended protocol from Akoya Biosciences using commercial staining reagents (Akoya Biosciences Cat. No. 7000008).

The tissue was fixed for 5 min in acetone (Macron Chemicals, Cat.No. MK244310), air dried for 2 min, followed by hydration 2 × 2 min in hydration buffer (Akoya Biosciences Cat. No. 7000008 Part No. 232105). The tissue was fixed for 10 min in 1.6% paraformaldehyde diluted in hydration buffer and kept in staining buffer for no longer than 30 min before staining. Staining was done in staining buffer supplemented with blocker N, blocker J, blocker S, and blocker G (dilution factor 1:40) (Akoya Biosciences Cat. No. 7000008) with commercial and custom-conjugated antibodies at 1:200 (CD45, CD3, CD15, CD66b) and 1:100 dilution (CD161, TCR Va7.2) (Supplementary Table 10). The conjugation of CD15, CD66b, CD161, and TCR Va7.2 was performed as previously described^18^. The tissue was stained at RT for 3 h, followed by 2 × 2 min wash in staining buffer, followed by fixation in 1.6% paraformaldehyde diluted in storage buffer (Akoya Biosciences Cat. No. 7000008 Part No. 232106). The tissue was rinsed 3 X in PBS and fixed in ice-cold methanol for 5 min. After rinsing in PBS 3 times, the fixative (Akoya Biosciences Cat. No. 7000008 Part No. 232107) diluted in PBS was applied for 20 min, followed by rinsing in PBS. The stained samples were kept at 4°C in a storage buffer for less than two weeks or imaged immediately.

### Whole slide imaging using Phenocycler Fusion

Imaging of stained slides was performed on the Phenocycler Fusion system using DAPI, CY3, CY5, and CY7 filters with a 20X objective with NA of 0.75 following a recommendation from Akoya Biosciences. Imaging was done in multiple cycles using exposure times listed in Supplementary Table 10 using software version 2.1.0. When the run was finished, the generated qptiff composite images were loaded into HALO 2D image analysis software (Indica Labs) for visualization and analysis.

After imaging, the slides were immersed in Xylene (Sigma-Aldrich) overnight to remove the flow cell, and after rehydration in PBS, were stained for hematoxylin-eosin (H&E) using DAKO reagents and recommended protocol on Parhelia’s Spatial Station autostainer. The stained slides were cover slipped using Permount (Fisher Chemical) mounting medium and scanned on Phenoimager (Akoya Biosciences) using 20X magnification. The H&E slides were loaded into HALO for visualization and tissue annotation by a board-certified pathologist (PJC).

### Image analysis in HALO

The histological features of the tissues were annotated on the hematoxylin and eosin (H&E)–stain images. For further analysis, we applied manual cell segmentation to quantify the nucleated cells. The cell segmentation results from the four images were then merged and further analyzed.

The HALO HighPlex FL module 4.2.14 was used for cell segmentation, signal thresholding, and manual phenotyping. Cell segmentation was performed using a traditional DAPI-controlled watershed nuclear segmentation algorithm implemented in HALO. Digital overlays of the cell-segmentation masks were used to visually control the performance of single-cell detection.

For manual gating of cell types, marker positivity of each marker was assessed visually, and positivity thresholds were set for each marker for each tissue section. This results in the classification of each cell as either positive or negative for a respective marker. MAIT cells were defined as TCR Va7.2 and CD161 double-positive cells.

The single-cell analysis results were exported as CSV files for further downstream analysis.

### Spatial analysis with SPAC

Downstream analysis and visualization were performed within the NIH Integrated Analysis Platform (NIDAP) using the SPAC python package (https://github.com/FNLCR-DMAP/SCSAWorkflow)^51^ developed by a team of NCI data scientists on the Foundry platform (Palantir Technologies). We employed the neighborhood enrichment method implemented in *Squidpy* to analyze spatial interactions between cells^85^. The neighborhood of a cell was defined by the k-nearest neighboring (where *k*=6) cells. Based on this neighborhood definition, we determined if two cell types form neighboring pairs more often than expected by chance. A spatial connectivity graph was constructed using six nearest neighbors to define the local microenvironment of each cell. To assess the significance of observed cell-type associations, we performed 100 random permutations of cell-type labels while preserving the spatial structure. For each pair of cell types, enrichment was quantified using z-scores, which reflect the deviation of observed interactions from the random permutations. The resulting z-scores highlight statistically enriched or depleted cell-type interactions within the tissue microenvironment.

To assess spatial clustering across multiple phenotypes, we utilized a modified version of Ripley’s L statistic, originally implemented in squidpy. Our modification, integrated into the SPAC Python package, extends the squidpy method to handle the Ripley’s L measure between two phenotypes. The spatial region’s area was estimated by fitting a polygon encompassing all observed cells. As in Squidpy’s implementation, cell positions were randomized using a Poisson point process to generate null distributions, with 100 simulations performed for each analysis. The results were visualized as the Ripley’s L measure across various distances R from the center phenotypes. The Ripley L statistics were calculated for multiple radii and compared with a random simulation of the MAIT and neutrophil to see if the observed MAIT cell and neutrophils cluster more than that of the random simulation.

### Statistical Analysis

The significance of variables was assessed using a t-test for two independent groups. A one-way ANOVA followed by multiple comparisons involving more than two independent groups. A p-value of ≤ 0.05 was considered statistically significant. In cases where the assumptions for parametric tests were not met, the Mann-Whitney U test for two independent groups was used. In survival analysis, the Cox proportional hazards regression model was used to calculate hazard ratios (HR) and corresponding 95% confidence intervals (CI). All statistical analyses were performed in Prism (version 10.4.1) unless otherwise specified.

### Ethics

Access to TCGA-controlled data and studies using them were approved by the National Cancer Institute (#28406). Human peripheral blood samples were obtained with approval from the institutional review board of NCI. The UCLA Medical Institutional Review Board 2 (IRB#10-000655-AM-00059) approved all protocols related to patient tumor specimen collection at UCLA.

### Data availability

The TCGA-GBM data used in this study are publicly available. The glioblastoma TCGA data can be downloaded from https://www.cbioportal.org/.

## Acknowledgement

The results published here are in part based upon data generated by the TCGA Research Network: https://www.cancer.ov/tcga. CODEX tissue images were analyzed in part utilizing the NCI HALO Image Analysis Resource. Single cell RNA-seq data analysis was performed on NIH HPC Biowulf. Human PBMC samples were obtained from the Department of Transfusion Medicine at the NIH 146 Clinical Center. We sincerely thank the SPAC development team, including Rui He, and Fang Liu, for their outstanding support in function development and for training us on the use of the corresponding SPAC package.

## Authorship Statement

Conceptualization: S.K., T.H, M.T.; Investigation: S.K., T.H., M.T., J.S., N.K., N.M., A.L., R.P., P.J.C., A.H., M.K.B.R., C.S., H.K, M.C, A.H, G.K., M. Watowich, M. Watanabe, M.G; Writing and Editing: All authors; Funding acquisition: M.T., R.P., M.R.G. All authors read and approved the manuscript.

## Notes

### Competing Interest Statement

The authors have declared no competing interest.

### Summary of Updates

Results from Multiplexed tissue imaging of glioblastoma patient specimens with Co-detection by indexing (CODEX) was included. The tissue imaging experiment showed that MAIT cells and neutrophils are found in the tumor of glioblastoma patients. In addition, to be consistent with the glioblastoma definition by WHO classification 2021, patient samples with reported IDH mutation were removed from TCGA dataset and the revised results were updated. MAIT cell signature genes were updated to include more MAIT cell marker genes which were reported by multiple independent studies.

