## Supplementary Material for "MAIT cells have a negative impact on glioblastoma"

^1^Neuro-Oncology Branch, Center for Cancer Research, National Cancer Institute, National Institutes of Health, USA, ^2^Department of Neurosurgery, UCLA, USA, ^3^Parker Institute for Cancer Immunotherapy, ^4^Collaborative Protein Technology Resource, Office of Science and Technology, ^5^Thoracic and Gastrointestinal Malignancy Branch, ^6^Cancer Data Science Laboratory, Center for Cancer Research, National Cancer Institute, National Institutes of Health, Bethesda, MD, USA, ^7^Surgical Neurology Branch, National Institute of Neurological Disorders and Stroke, National Institutes of Health, Bethesda, MD 20892, USA, ^8^Biomedical and Computational Science Directorate, Frederick National Laboratory for Cancer Research, Frederick, Maryland, USA

* These authors contributed equally

**Corresponding:**

Masaki Terabe, Ph.D, Neuro-Oncology Branch, National Cancer Institute, NIH, Building-37, Room-1016, 37 Convent Drive, Bethesda, MD 20892, USA

**Identification of MR1 mutational frequency, the epigenetic transcriptional status, the transcriptional distribution in GBM tissue**

Examined cBioportal’s datasets for the 3391 multi-grade glioma samples were Glioblastoma (TCGA, Cell 2013), Low-Grade Gliomas (UCSF, Science 2014), Merged Cohort of LGG and GBM (TCGA, Cell 2016), Anaplastic Oligodendroglioma and Anaplastic Oligoastrocytoma (MSKCC, Neuro Oncol 2017), Glioma (MSK, Nature 2019), Glioma (MSKCC, Clin Cancer Res 2019), Brain Lower Grade Glioma (TCGA, PanCancer Atlas). The H3K27ac ChIP data of GBM2w (IDHwt GBM, GSM1824806), AA15m (IDHmut-R132H, Grade3 Astrocytoma, GSM1824808) were used for the ChIP analysis. IgV_2.11.9 was used to visualize them.

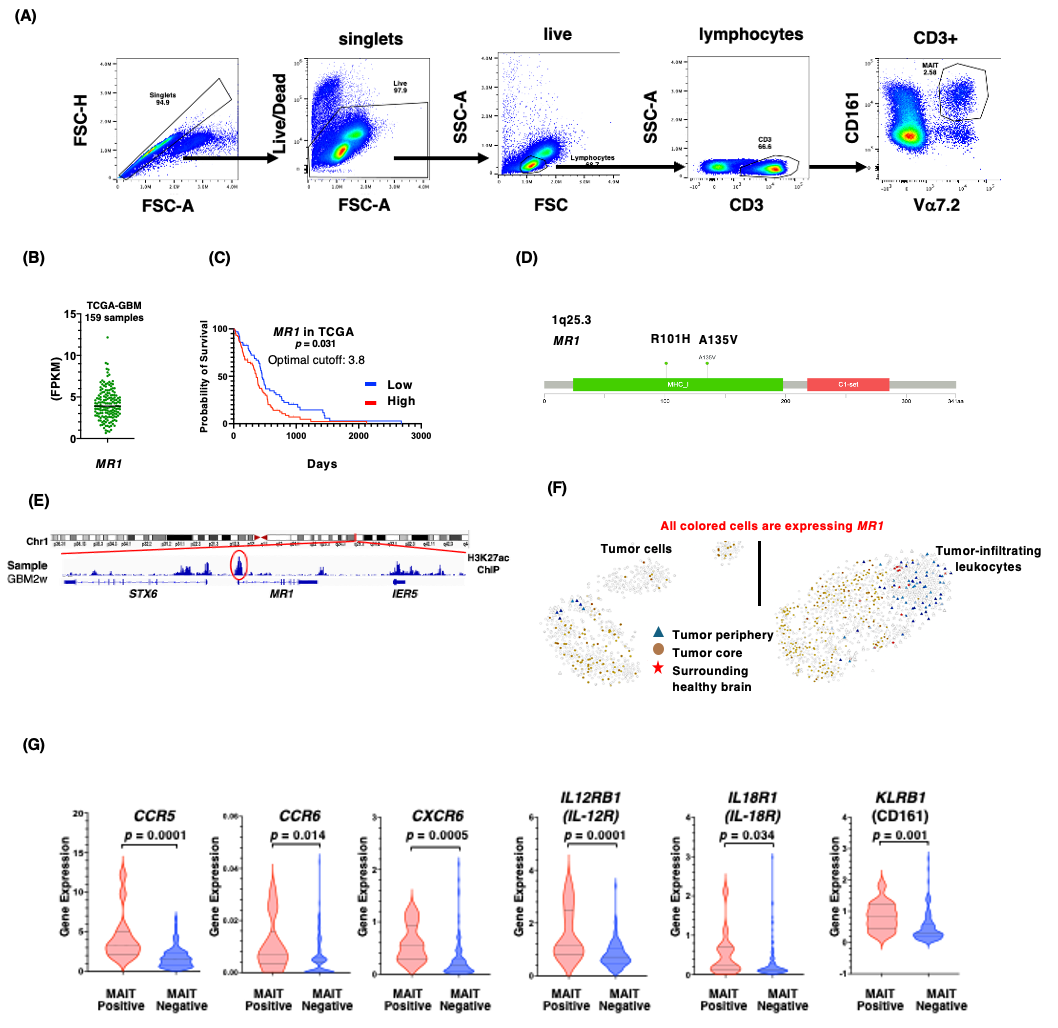

**Extended data Figure 1. (A)** Gating strategy to identify MAIT cells in PBMCs. **(B)** Expression of MR1 in the TCGA-GBM data set. **(C)** The correlation between MR1 expression levels and GBM patient survival in the TCGA -GBM dataset. The correlation between MR1 expression and patient survival was tested with a Log-Rank (Mantel-Cox) test. **(D)** The mutational schema in MR1 locus. **(E)** H3K27ac ChIP peaks in MR1 locus of GBM (GBM2w, IDHwt) samples. **(F)** tSNE plots of single-cell RNA-seq data from GBM tissues of four patients. Colored dots represent MR1-expressing cells. **(G)** Transcriptomic expression of MAIT cell surface receptors, *CCR5*, *CCR6*, *CXCR6*, *IL-12R, IL-18R, and CD161.* Statistical *p*-values were calculated by a non-parametric Mann-Whitney test.

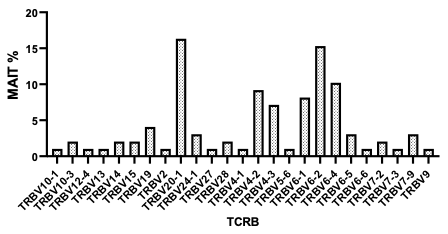

**Extended data Figure 2. MAIT cell TCR beta chain usage in GBM obtained from single-cell TCR sequencing.**

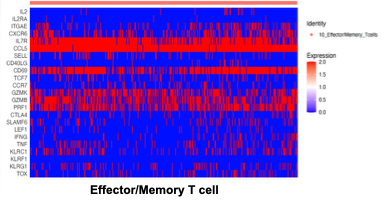

**Extended data Figure 3. Heatmap showing the effector T cell markers and memory T cell markers in the Effector/Memory T cell cluster.** Each vertical line in the heat map represents a cell in the Effector/Memory T cell cluster.

**(B)**

**(A)**

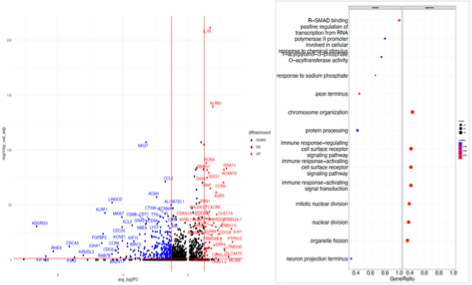

**Extended data Figure 4. Differential gene expression analysis of Th17-like cluster.** **(A)** A volcano plot of up- and down-regulated genes in Th17-like cell population. (B) GSEA analysis of gene differentially expressed in Th17-like cell population. Differential expression analysis was performed in R with *limma* and *edger* packages.

**(C)**

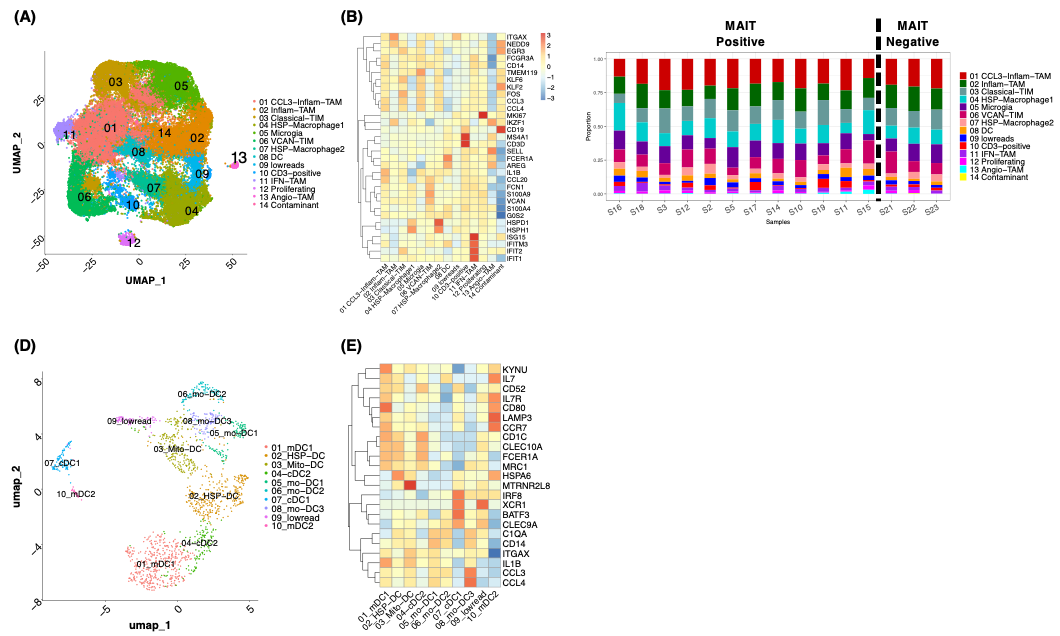

**Extended data Figure 5. Analysis of myeloid cell populations. (A)** UMAP of the myeloid cell population **(B)** Heatmap of markers for the myeloid clusters **(C)** Distribution of cell types between MAIT-Positive and MAIT-Negative samples. The myeloid clusters were characterized in the following ways: one inflammatory tumor-associated macrophages (TAMs) with high expression of *IKZF1, and NFKB1* (Inflam-TAM); one *CCL3* high inflammatory TAM (CCL3-Inflam-TAM); Classical tumor-infiltrating monocytes (TIMs) with high *CD14* and *EGR1* expression (classical-TIM); two clusters of heat shock protein (HSP) enriched macrophages showed high expression for *HSPE1*, *HSPH1* and *HSPD1* (HSP-Macrophage1 and HSP-Macrophage2); one microglia enriched clusters with high *TMEM119* (Microglia); one cluster with high *VCAN, SERPINB2, CXCL2/3/5/8, S100A9* and *LYZ* expression (VCAN-TIM); DC cluster with high *LGALS2*, *FCER1A* and MHC II expression (DC); interferon-primed TAMs (IFN-TAM) with high *IFIT1/IFIT2, IFITM3*, and *ISG15*; proliferating cell cluster (Proliferating) with high *MKI67* expression; angio TAM with *GZMB, CXCR3, CXCR4* and *AREG* expression; One cluster of a mixture of myeloid and lymphoid cells with high *CD19* expression (Contaminant); one CD3-positive cluster with *CD3E*, *CD3D* expression within the myeloid population (CD3-positive). A cluster with mixed cell populations positive for myeloid cell markers or T-cell markers in a tumor tissue has been reported in another study[1]. **(D)** UMAP of Dendritic cell populations in GBM patient samples **(E)** Heatmap of the markers for the different DC clusters. The cluster annotation in the DC population in **(D)** and **(E)** are as follows: Two migratory DC (mDC1 & mDC2), DC cluster with high expression of heat shock proteins (HSP-DC), DC cluster with high expression of mitochondrial genes (mito-DC), two cDC clusters (cDC1 & cDC2) and three monocytic DC clusters (mo-DC1 & mo-DC2). The DC clusters are characterized as follows: one activated mDC1 (migratory DC) with high expression of *CCR7, LAMP3, KYNU, IL7R* and *CLEC10A*; one migratory DC (mDC2) with high expression of *LAMP3, IL7* and *CCR7*; one cluster enriched in heat shock proteins (HSP-DC); one cluster with high mitochondrial gene expression (mito-DC); one cDC1 expressing *IRF8, BATF3, XCR1*; cDC2 with high *FCER1A* and *CD1C*. There were three monocyte-like DCs: mo-DC1 with high *CD14* and *C1QA* expression; Mo-DC2 expressing high *CD14, C1QA, CLEC9A* and Mo-DC3 with high expression of *CD14, C1QA, CCL3/4.*

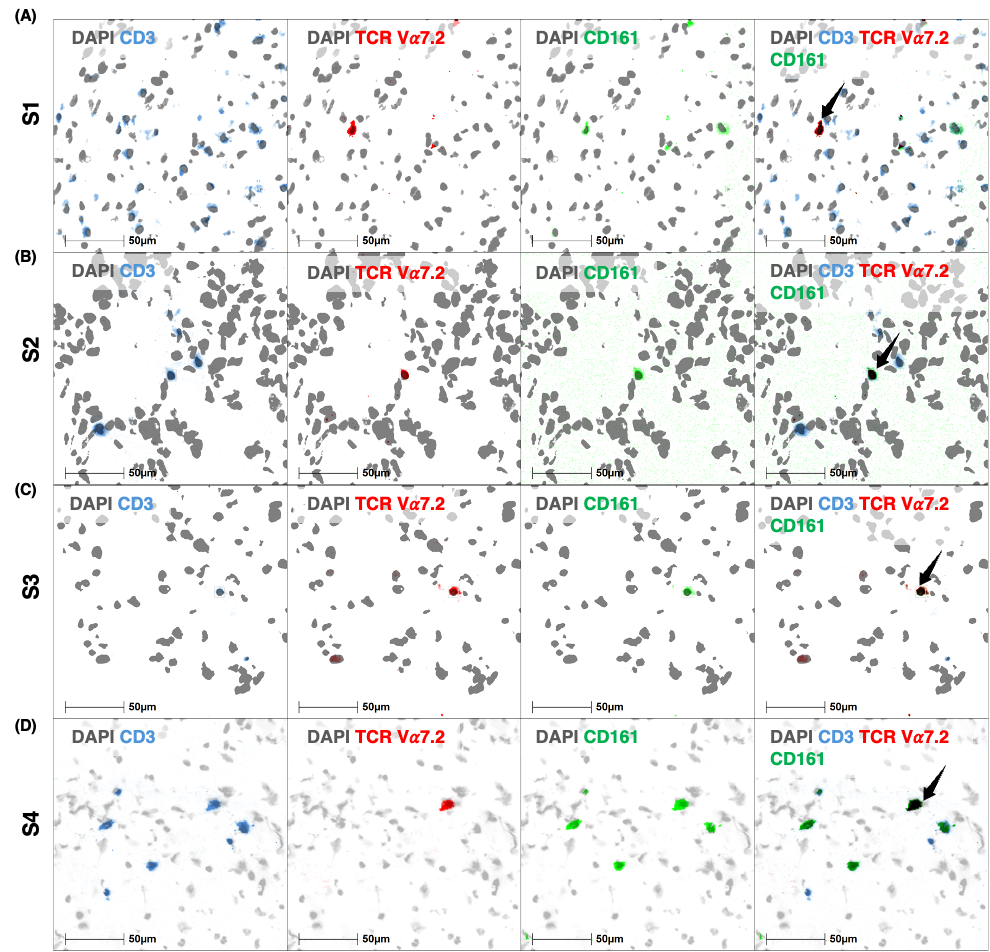

**Extended data Figure 6. MAIT cells were identified based on the expression of CD3, TCR V**𝛼**7.2, and CD161. (A-D)** Representative MAIT cells in four specimens, S1, S2, S3, and S4. DAPI (nuclear staining), CD3, TCR V𝛼7.2, CD161, are represented in grey, blue, red, and green, respectively. MAIT cells with the overlaid stains (DAPI, CD3, TCR V𝛼7.2 and CD161) are shown with black arrows.

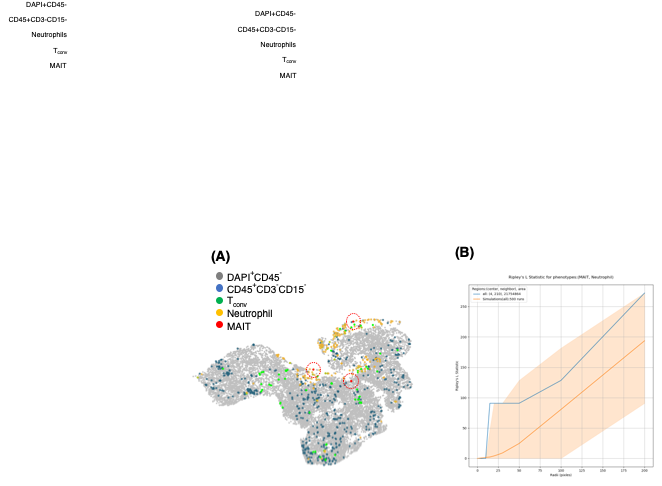

**Figures 7.** **Spatial relationship between MAIT cells and neutrophils.** **(A)** Neutrophils (yellow) aggregation is localized near the MAIT cells (red). MAIT cells are shown within red dotted circles. Other cell-types including T_conv_ cells (green) and CD45^+^CD3^-^CD15^-^ cells (blue) and DAPI^+^CD45^-^ cells (grey) do not aggregate near the MAIT cells. **(B)** Ripley L statistical score for MAIT cell-neutrophil for multiple radii ranging from 0 to 200 pixel is shown. Orange line represents the Ripley L score from simulation while the blue line represents the observed Ripley L score. The error margin is shown in light brown color.

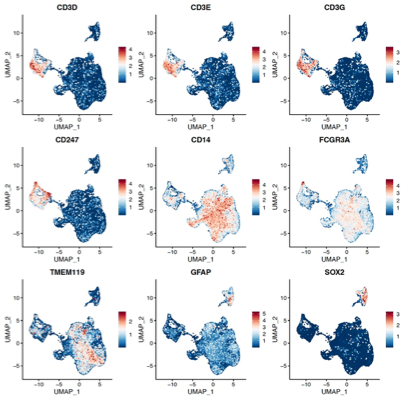

**Extended data Figure 8. Overview of the expression of marker genes used to identify cell clusters.** Marker genes used to identify cell clusters: lymphoid cells (*CD3D/E/G and CD247*), myeloid cells (*CD14, ITGAX, FCGR3A*, and *TMEM119*), normal/tumor brain cells (*GFAP* and *SOX2*).

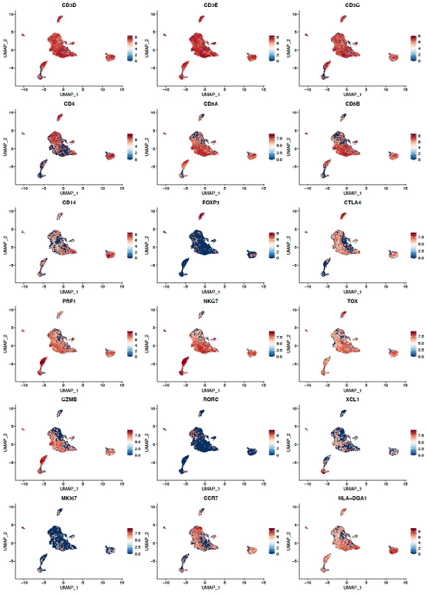

**Extended data Figure 9. Overview of markers used to identify lymphoid cell clusters.**

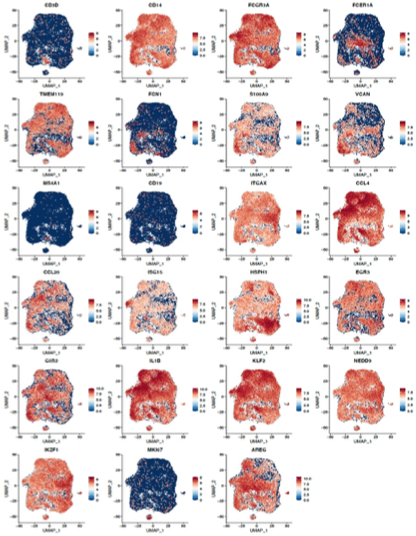

**Extended data Figure 10. Overview of markers used to identify myeloid cell clusters**.

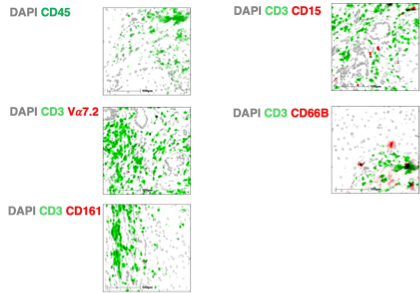

**Extended data Figure 11.** Validation of CODEX antibody panel. The antibody-oligonucleotide conjugates were tested on human liver tissue.
