## Supplementary Table 1 for "MAIT cells have a negative impact on glioblastoma"

| Supplementary Table 1: Searched characteristic sequences of MAIT-TCR gene segments | |
|---|---|
| Gene segment | Sequence |
| TRAV1-2 | GAGCTCCAGATGAAAGACTCTG |
| TRAJ12 | AAATTGATCTTCGGGAGTGGGA |
| TRAJ20 | CAAGCTCAGCTTTGGAGCCGGA |
| TRAJ33 | TCAGTTAATCTGGGGCGCTGGG |
