## Supplementary Table 2 for "MAIT cells have a negative impact on glioblastoma"

| Supplementary Table 2: TCGA-GBM core 159 sample IDs | |
|---|---|
| TCGA-02-2486-01A | MAIT-positive |
| TCGA-06-0130-01A | MAIT-positive |
| TCGA-06-0132-01A | MAIT-positive |
| TCGA-06-0152-02A | MAIT-positive |
| TCGA-06-0171-02A | MAIT-positive |
| TCGA-06-0190-01A | MAIT-positive |
| TCGA-06-0211-01A | MAIT-positive |
| TCGA-06-0649-01B | MAIT-positive |
| TCGA-06-2557-01A | MAIT-positive |
| TCGA-14-0817-01A | MAIT-positive |
| TCGA-19-4065-01A | MAIT-positive |
| TCGA-32-2638-01A | MAIT-positive |
| TCGA-41-3915-01A | MAIT-positive |
| TCGA-76-4928-01B | MAIT-positive |
| TCGA-02-0047-01A | MAIT-negative |
| TCGA-02-0055-01A | MAIT-negative |
| TCGA-02-2485-01A | MAIT-negative |
| TCGA-06-0125-01A | MAIT-negative |
| TCGA-06-0125-02A | MAIT-negative |
| TCGA-06-0138-01A | MAIT-negative |
| TCGA-06-0139-01A | MAIT-negative |
| TCGA-06-0141-01A | MAIT-negative |
| TCGA-06-0156-01A | MAIT-negative |
| TCGA-06-0157-01A | MAIT-negative |
| TCGA-06-0158-01A | MAIT-negative |
| TCGA-06-0168-01A | MAIT-negative |
| TCGA-06-0174-01A | MAIT-negative |
| TCGA-06-0178-01A | MAIT-negative |
| TCGA-06-0184-01A | MAIT-negative |
| TCGA-06-0187-01A | MAIT-negative |
| TCGA-06-0190-02A | MAIT-negative |
| TCGA-06-0210-01A | MAIT-negative |
| TCGA-06-0210-02A | MAIT-negative |
| TCGA-06-0211-01B | MAIT-negative |
| TCGA-06-0211-02A | MAIT-negative |
| TCGA-06-0219-01A | MAIT-negative |
| TCGA-06-0221-02A | MAIT-negative |
| TCGA-06-0238-01A | MAIT-negative |
| TCGA-06-0644-01A | MAIT-negative |
| TCGA-06-0645-01A | MAIT-negative |
| TCGA-06-0646-01A | MAIT-negative |
| TCGA-06-0686-01A | MAIT-negative |

| | |
|---|---|
| TCGA-06-0743-01A | MAIT-negative |
| TCGA-06-0744-01A | MAIT-negative |
| TCGA-06-0745-01A | MAIT-negative |
| TCGA-06-0747-01A | MAIT-negative |
| TCGA-06-0749-01A | MAIT-negative |
| TCGA-06-0750-01A | MAIT-negative |
| TCGA-06-0878-01A | MAIT-negative |
| TCGA-06-0882-01A | MAIT-negative |
| TCGA-06-1804-01A | MAIT-negative |
| TCGA-06-2558-01A | MAIT-negative |
| TCGA-06-2559-01A | MAIT-negative |
| TCGA-06-2561-01A | MAIT-negative |
| TCGA-06-2562-01A | MAIT-negative |
| TCGA-06-2563-01A | MAIT-negative |
| TCGA-06-2564-01A | MAIT-negative |
| TCGA-06-2565-01A | MAIT-negative |
| TCGA-06-2567-01A | MAIT-negative |
| TCGA-06-2569-01A | MAIT-negative |
| TCGA-06-5408-01A | MAIT-negative |
| TCGA-06-5410-01A | MAIT-negative |
| TCGA-06-5411-01A | MAIT-negative |
| TCGA-06-5412-01A | MAIT-negative |
| TCGA-06-5413-01A | MAIT-negative |
| TCGA-06-5414-01A | MAIT-negative |
| TCGA-06-5418-01A | MAIT-negative |
| TCGA-06-5856-01A | MAIT-negative |
| TCGA-06-5858-01A | MAIT-negative |
| TCGA-06-5859-01A | MAIT-negative |
| TCGA-08-0386-01A | MAIT-negative |
| TCGA-12-0616-01A | MAIT-negative |
| TCGA-12-0618-01A | MAIT-negative |
| TCGA-12-0619-01A | MAIT-negative |
| TCGA-12-0821-01A | MAIT-negative |
| TCGA-12-1597-01B | MAIT-negative |
| TCGA-12-3650-01A | MAIT-negative |
| TCGA-12-3652-01A | MAIT-negative |
| TCGA-12-3653-01A | MAIT-negative |
| TCGA-12-5295-01A | MAIT-negative |
| TCGA-12-5299-01A | MAIT-negative |
| TCGA-14-0736-02A | MAIT-negative |
| TCGA-14-0781-01B | MAIT-negative |
| TCGA-14-0787-01A | MAIT-negative |
| TCGA-14-0789-01A | MAIT-negative |

| | |
|---|---|
| TCGA-14-0790-01B | MAIT-negative |
| TCGA-14-0871-01A | MAIT-negative |
| TCGA-14-1034-01A | MAIT-negative |
| TCGA-14-1034-02B | MAIT-negative |
| TCGA-14-1402-02A | MAIT-negative |
| TCGA-14-1823-01A | MAIT-negative |
| TCGA-14-1825-01A | MAIT-negative |
| TCGA-14-1829-01A | MAIT-negative |
| TCGA-14-2554-01A | MAIT-negative |
| TCGA-15-0742-01A | MAIT-negative |
| TCGA-15-1444-01A | MAIT-negative |
| TCGA-16-0846-01A | MAIT-negative |
| TCGA-16-1045-01B | MAIT-negative |
| TCGA-19-0957-02A | MAIT-negative |
| TCGA-19-1389-02A | MAIT-negative |
| TCGA-19-1390-01A | MAIT-negative |
| TCGA-19-2619-01A | MAIT-negative |
| TCGA-19-2620-01A | MAIT-negative |
| TCGA-19-2624-01A | MAIT-negative |
| TCGA-19-2625-01A | MAIT-negative |
| TCGA-19-4065-02A | MAIT-negative |
| TCGA-19-5960-01A | MAIT-negative |
| TCGA-26-5132-01A | MAIT-negative |
| TCGA-26-5133-01A | MAIT-negative |
| TCGA-26-5134-01A | MAIT-negative |
| TCGA-26-5135-01A | MAIT-negative |
| TCGA-26-5136-01B | MAIT-negative |
| TCGA-26-5139-01A | MAIT-negative |
| TCGA-27-1830-01A | MAIT-negative |
| TCGA-27-1831-01A | MAIT-negative |
| TCGA-27-1832-01A | MAIT-negative |
| TCGA-27-1834-01A | MAIT-negative |
| TCGA-27-1835-01A | MAIT-negative |
| TCGA-27-1837-01A | MAIT-negative |
| TCGA-27-2519-01A | MAIT-negative |
| TCGA-27-2523-01A | MAIT-negative |
| TCGA-27-2524-01A | MAIT-negative |
| TCGA-27-2526-01A | MAIT-negative |
| TCGA-27-2528-01A | MAIT-negative |
| TCGA-28-1747-01C | MAIT-negative |
| TCGA-28-1753-01A | MAIT-negative |
| TCGA-28-2499-01A | MAIT-negative |
| TCGA-28-2509-01A | MAIT-negative |

| | |
|---|---|
| TCGA-28-2510-01A | MAIT-negative |
| TCGA-28-2513-01A | MAIT-negative |
| TCGA-28-2514-01A | MAIT-negative |
| TCGA-28-5204-01A | MAIT-negative |
| TCGA-28-5207-01A | MAIT-negative |
| TCGA-28-5208-01A | MAIT-negative |
| TCGA-28-5209-01A | MAIT-negative |
| TCGA-28-5213-01A | MAIT-negative |
| TCGA-28-5215-01A | MAIT-negative |
| TCGA-28-5216-01A | MAIT-negative |
| TCGA-28-5218-01A | MAIT-negative |
| TCGA-28-5220-01A | MAIT-negative |
| TCGA-32-1970-01A | MAIT-negative |
| TCGA-32-1980-01A | MAIT-negative |
| TCGA-32-1982-01A | MAIT-negative |
| TCGA-32-2615-01A | MAIT-negative |
| TCGA-32-2616-01A | MAIT-negative |
| TCGA-32-2632-01A | MAIT-negative |
| TCGA-32-2634-01A | MAIT-negative |
| TCGA-32-4213-01A | MAIT-negative |
| TCGA-32-5222-01A | MAIT-negative |
| TCGA-41-2571-01A | MAIT-negative |
| TCGA-41-2572-01A | MAIT-negative |
| TCGA-41-4097-01A | MAIT-negative |
| TCGA-41-5651-01A | MAIT-negative |
| TCGA-76-4925-01A | MAIT-negative |
| TCGA-76-4926-01B | MAIT-negative |
| TCGA-76-4927-01A | MAIT-negative |
| TCGA-76-4929-01A | MAIT-negative |
| TCGA-76-4931-01A | MAIT-negative |
| TCGA-76-4932-01A | MAIT-negative |
