## Supplementary Table 3 for "MAIT cells have a negative impact on glioblastoma"

### Supplementary Table 3: The IDs of 144 T

### TCGA Dataset

TCGA-02-0047-01A
TCGA-02-0055-01A
TCGA-02-2485-01A
TCGA-02-2486-01A
TCGA-06-0125-01A
TCGA-06-0130-01A
TCGA-06-0132-01A
TCGA-06-0138-01A
TCGA-06-0139-01A
TCGA-06-0141-01A
TCGA-06-0156-01A
TCGA-06-0157-01A
TCGA-06-0158-01A
TCGA-06-0168-01A
TCGA-06-0174-01A
TCGA-06-0178-01A
TCGA-06-0184-01A
TCGA-06-0187-01A
TCGA-06-0190-01A
TCGA-06-0210-01A
TCGA-06-0211-01B
TCGA-06-0219-01A
TCGA-06-0238-01A
TCGA-06-0644-01A
TCGA-06-0645-01A
TCGA-06-0646-01A
TCGA-06-0649-01B
TCGA-06-0686-01A
TCGA-06-0743-01A
TCGA-06-0744-01A
TCGA-06-0745-01A
TCGA-06-0747-01A
TCGA-06-0749-01A
TCGA-06-0750-01A
TCGA-06-0878-01A
TCGA-06-0882-01A
TCGA-06-1804-01A
TCGA-06-2557-01A
TCGA-06-2558-01A

TCGA-06-2559-01A
TCGA-06-2561-01A
TCGA-06-2562-01A
TCGA-06-2563-01A
TCGA-06-2564-01A
TCGA-06-2565-01A
TCGA-06-2567-01A
TCGA-06-2569-01A
TCGA-06-5408-01A
TCGA-06-5410-01A
TCGA-06-5411-01A
TCGA-06-5412-01A
TCGA-06-5413-01A
TCGA-06-5414-01A
TCGA-06-5418-01A
TCGA-06-5856-01A
TCGA-06-5858-01A
TCGA-06-5859-01A
TCGA-08-0386-01A
TCGA-12-0616-01A
TCGA-12-0618-01A
TCGA-12-0619-01A
TCGA-12-0821-01A
TCGA-12-1597-01B
TCGA-12-3650-01A
TCGA-12-3652-01A
TCGA-12-3653-01A
TCGA-12-5295-01A
TCGA-12-5299-01A
TCGA-14-0781-01B
TCGA-14-0787-01A
TCGA-14-0789-01A
TCGA-14-0790-01B
TCGA-14-0817-01A
TCGA-14-0871-01A
TCGA-14-1034-01A
TCGA-14-1823-01A
TCGA-14-1825-01A
TCGA-14-1829-01A
TCGA-14-2554-01A
TCGA-15-1444-01A
TCGA-16-0846-01A
TCGA-16-1045-01B

TCGA-19-1390-01A
TCGA-19-2619-01A
TCGA-19-2620-01A
TCGA-19-2624-01A
TCGA-19-2625-01A
TCGA-19-4065-01A
TCGA-19-5960-01A
TCGA-26-5132-01A
TCGA-26-5133-01A
TCGA-26-5134-01A
TCGA-26-5135-01A
TCGA-26-5136-01B
TCGA-26-5139-01A
TCGA-27-1830-01A
TCGA-27-1831-01A
TCGA-27-1832-01A
TCGA-27-1834-01A
TCGA-27-1835-01A
TCGA-27-1837-01A
TCGA-27-2519-01A
TCGA-27-2523-01A
TCGA-27-2524-01A
TCGA-27-2526-01A
TCGA-27-2528-01A
TCGA-28-1747-01C
TCGA-28-1753-01A
TCGA-28-2499-01A
TCGA-28-2509-01A
TCGA-28-2513-01A
TCGA-28-2514-01A
TCGA-28-5204-01A
TCGA-28-5207-01A
TCGA-28-5208-01A
TCGA-28-5209-01A
TCGA-28-5213-01A
TCGA-28-5215-01A
TCGA-28-5216-01A
TCGA-28-5218-01A
TCGA-28-5220-01A
TCGA-32-1970-01A
TCGA-32-1980-01A
TCGA-32-1982-01A
TCGA-32-2615-01A

TCGA-32-2616-01A
TCGA-32-2632-01A
TCGA-32-2634-01A
TCGA-32-2638-01A
TCGA-32-4213-01A
TCGA-32-5222-01A
TCGA-41-2571-01A
TCGA-41-2572-01A
TCGA-41-3915-01A
TCGA-41-4097-01A
TCGA-41-5651-01A
TCGA-76-4925-01A
TCGA-76-4926-01B
TCGA-76-4927-01A
TCGA-76-4928-01B
TCGA-76-4929-01A
TCGA-76-4931-01A
TCGA-76-4932-01A

**CGA-GBM samples used for the prognostic analysis**
