## Supplementary Table 4 for "MAIT cells have a negative impact on glioblastoma"

| Supplementary Table 4. Correlation analysis between MR1 and TRAV1-2 gene expression in scRNA-seq data from glioma patients | | | | | | | | | |
|---|---|---|---|---|---|---|---|---|---|
| **S.No.** | **GSMID** | **Fragment** | **Subtype** | **MR1** | **TRAV1-2** | **TRAJ12/20/33** | **Num cells** | **Num Tcells** | **Num non-Tcells** |
| **0** | GSM5518605 | ndGBM-11-B | ndGBM | 17 | 0 | 0 | 271 | 14 | 257 |
| **1** | GSM5518604 | ndGBM-11-A | ndGBM | 31 | 0 | 0 | 357 | 19 | 338 |
| **2** | GSM5518607 | ndGBM-11-D | ndGBM | 65 | 0 | 0 | 1041 | 25 | 1016 |
| **3** | GSM5518614 | rGBM-02-4 | rGBM | 80 | 1 | 0 | 780 | 539 | 241 |
| **4** | GSM5518606 | ndGBM-11-C | ndGBM | 160 | 0 | 0 | 2310 | 29 | 2281 |
| **5** | GSM5518600 | ndGBM-01-A | ndGBM | 165 | 0 | 0 | 3348 | 32 | 3316 |
| **6** | GSM5518632 | LGG-04-3 | LGG | 175 | 0 | 0 | 4815 | 33 | 4782 |
| **7** | GSM5518601 | ndGBM-01-C | ndGBM | 192 | 0 | 0 | 7503 | 45 | 7458 |
| **8** | GSM5518602 | ndGBM-01-D | ndGBM | 204 | 0 | 0 | 5059 | 3 | 5056 |
| **9** | GSM5518635 | ndGBM-06 | ndGBM | 334 | 0 | 0 | 3167 | 47 | 3120 |
| **10** | GSM5518603 | ndGBM-01-F | ndGBM | 340 | 0 | 0 | 4450 | 50 | 4400 |
| **11** | GSM5518617 | rGBM-03-2 | rGBM | 371 | 0 | 0 | 1600 | 181 | 1419 |
| **12** | GSM5518612 | rGBM-02-2 | rGBM | 439 | 5 | 0 | 2106 | 950 | 1156 |
| **13** | GSM5518610 | ndGBM-02-4 | ndGBM | 475 | 1 | 0 | 3126 | 41 | 3085 |
| **14** | GSM5518616 | rGBM-03-1 | rGBM | 481 | 0 | 0 | 3629 | 44 | 3585 |
| **15** | GSM5518631 | LGG-04-2 | LGG | 531 | 0 | 0 | 2201 | 39 | 2162 |
| **16** | GSM5518630 | LGG-04-1 | LGG | 544 | 0 | 0 | 5032 | 94 | 4938 |
| **17** | GSM5518624 | ndGBM-03-2 | ndGBM | 567 | 0 | 0 | 2387 | 155 | 2232 |
| **18** | GSM5518629 | ndGBM-10 | ndGBM | 588 | 0 | 0 | 9873 | 152 | 9721 |
| **19** | GSM5518611 | ndGBM-02-5 | ndGBM | 594 | 0 | 0 | 3182 | 25 | 3157 |
| **20** | GSM5518628 | rGBM-05-3 | rGBM | 595 | 0 | 0 | 1837 | 280 | 1557 |
| **21** | GSM5518623 | ndGBM-03-1 | ndGBM | 623 | 0 | 0 | 3541 | 368 | 3173 |
| **22** | GSM5518619 | rGBM-04-1 | rGBM | 703 | 0 | 0 | 6165 | 311 | 5854 |
| **23** | GSM5518608 | ndGBM-02-1 | ndGBM | 714 | 0 | 0 | 4196 | 21 | 4175 |
| **24** | GSM5518638 | LGG-03 | LGG | 794 | 0 | 0 | 7608 | 64 | 7544 |
| **25** | GSM5518618 | rGBM-03-3 | rGBM | 798 | 0 | 0 | 3995 | 222 | 3773 |
| **26** | GSM5518598 | rGBM-01-C | rGBM | 821 | 2 | 0 | 4533 | 130 | 4403 |
| **27** | GSM5518620 | rGBM-04-2 | rGBM | 829 | 1 | 0 | 6132 | 185 | 5947 |
| **28** | GSM5518621 | rGBM-04-3 | rGBM | 837 | 0 | 0 | 7400 | 250 | 7150 |
| **29** | GSM5518596 | rGBM-01-A | rGBM | 839 | 1 | 0 | 3339 | 175 | 3164 |
| **30** | GSM5518625 | ndGBM-03-3 | ndGBM | 862 | 1 | 0 | 3099 | 242 | 2857 |
| **31** | GSM5518609 | ndGBM-02-2 | ndGBM | 899 | 0 | 0 | 4030 | 313 | 3717 |
| **32** | GSM5518627 | rGBM-05-2 | rGBM | 923 | 0 | 0 | 3003 | 165 | 2838 |
| **33** | GSM5518599 | rGBM-01-D | rGBM | 1027 | 0 | 0 | 4695 | 50 | 4645 |
| **34** | GSM5518633 | ndGBM-04 | ndGBM | 1049 | 1 | 0 | 7411 | 111 | 7300 |
| **35** | GSM5518626 | rGBM-05-1 | rGBM | 1079 | 1 | 0 | 4696 | 520 | 4176 |
| **36** | GSM5518639 | ndGBM-09 | ndGBM | 1112 | 4 | 0 | 4939 | 1041 | 3898 |
| **37** | GSM5518634 | ndGBM-05 | ndGBM | 1119 | 0 | 0 | 6727 | 249 | 6478 |

| | | | | | | | | | |
|---|---|---|---|---|---|---|---|---|---|
| **38** | GSM5518615 | rGBM-02-5 | rGBM | 1194 | 16 | 0 | 9922 | 4964 | 4958 |
| **39** | GSM5518597 | rGBM-01-B | rGBM | 1299 | 6 | 0 | 8585 | 221 | 8364 |
| **40** | GSM5518613 | rGBM-02-3 | rGBM | 1301 | 7 | 0 | 9091 | 4438 | 4653 |
| **41** | GSM5518637 | ndGBM-08 | ndGBM | 1385 | 0 | 0 | 5497 | 134 | 5363 |
| **42** | GSM5518622 | rGBM-04-4 | rGBM | 1403 | 0 | 0 | 6372 | 376 | 5996 |
| **43** | GSM5518636 | ndGBM-07 | ndGBM | 2699 | 2 | 0 | 8936 | 2223 | 6713 |
