## Supplementary Table 5 for "MAIT cells have a negative impact on glioblastoma"

**Supplementary Table 5: Single-cell RNA Sequencing data analysis of 23 Glioblastomat patients. Clinical status and the MAIT TRAJ usage of the the glioblastoma patients. MAIT cells were identified based on the TCR gene segment combinations Vα7.2-Jα33, Vα7.2-Jα12, or Vα7.2-Jα20**

| Sample ID | Treatment | MAIT status | TRAJ_12 | TRAJ_20 | TRAJ_33 | Total MAIT Cells | Lymphoid Cluster Analysis | No. of Myeloid Cells | No. of Tumor/Normal Cells | No. of Lymphoid Cells | TCR seq. ID | Cell Count (TCR sequencing) |
|---|---|---|---|---|---|---|---|---|---|---|---|---|
| **S17** | recurrent GBM neoadjuvant anti-PD-1 | MAIT | 0 | 2 | 2 | 4 | Yes | 1296 | 419 | 898 | 18 | 492 |
| **S10** | recurrent GBM immunotherapy naïve | MAIT | 0 | 0 | 2 | 2 | Yes | 2642 | 545 | 184 | 7 | 209 |
| **S16** | recurrent GBM immunotherapy naïve | MAIT | 0 | 2 | 4 | 6 | Yes | 2087 | 397 | 693 | 1 | 477 |
| **S18** | recurrent GBM neoadjuvant anti-PD-1 | MAIT | 0 | 0 | 7 | 7 | Yes | 1547 | 359 | 464 | 5 | 413 |
| **S15** | recurrent GBM immunotherapy naïve | MAIT | 0 | 1 | 0 | 1 | Yes | 723 | 200 | 238 | 14 | 116 |
| **S14** | recurrent GBM immunotherapy naïve | MAIT | 0 | 1 | 1 | 2 | Yes | 1403 | 379 | 164 | 22 | 152 |
| **S5** | newly diagnosed GBM | MAIT | 0 | 0 | 3 | 3 | Yes | 2221 | 563 | 252 | 17 | 89 |
| **S2** | newly diagnosed GBM | MAIT | 0 | 3 | 1 | 4 | Yes | 6608 | 1251 | 975 | 15 | 577 |
| **S12** | recurrent GBM immunotherapy naïve | MAIT | 1 | 0 | 6 | 7 | Yes | 3558 | 1804 | 815 | 13 | 355 |
| **S19** | recurrent GBM neoadjuvant anti-PD-1 | MAIT | 5 | 1 | 30 | 36 | Yes | 3884 | 1338 | 2564 | 11 | 1668 |
| **S11** | recurrent GBM immunotherapy naïve | MAIT | 1 | 2 | 8 | 11 | Yes | 9470 | 2899 | 4194 | 4 | 897 |
| **S3** | newly diagnosed GBM | MAIT | 0 | 0 | 3 | 3 | Yes | 4798 | 698 | 624 | 6 | 435 |
| **S13** | recurrent GBM immunotherapy naïve | MAIT | 0 | 1 | 0 | 1 | No | 331 | 91 | 35 | 10 | 109 |
| **S4** | newly diagnosed GBM | MAIT | 0 | 1 | 0 | 1 | No | 370 | 73 | 71 | 8 | 113 |
| **S1** | newly diagnosed GBM | MAIT | 0 | 0 | 1 | 1 | No | 422 | 70 | 35 | 19 | 11 |
| **S21** | recurrent GBM neoadjuvant anti-PD-1 | non-MAIT | 0 | 0 | 0 | 0 | Yes | 4356 | 991 | 840 | 2 | 36 |
| **S23** | recurrent GBM neoadjuvant anti-PD-1 | non-MAIT | 0 | 0 | 0 | 0 | Yes | 1525 | 214 | 280 | 12 | 80 |
| **S22** | recurrent GBM neoadjuvant anti-PD-1 | non-MAIT | 0 | 0 | 0 | 0 | Yes | 7803 | 2908 | 1238 | 21 | 2 |
| **S20** | recurrent GBM neoadjuvant anti-PD-1 | non-MAIT | 0 | 0 | 0 | 0 | No | 791 | 200 | 99 | 16 | 21 |
| **S6** | newly diagnosed GBM | non-MAIT | 0 | 0 | 0 | 0 | No | 1002 | 180 | 113 | 3 | 29 |
| **S8** | newly diagnosed GBM | non-MAIT | 0 | 0 | 0 | 0 | No | 374 | 214 | 155 | 23 | 39 |
| **S7** | newly diagnosed GBM | non-MAIT | 0 | 0 | 0 | 0 | No | 1049 | 269 | 116 | 9 | 54 |
| **S9** | newly diagnosed GBM | non-MAIT | 0 | 0 | 0 | 0 | No | 109 | 69 | 75 | 20 | 55 |
