## Supplementary Table 6 for "MAIT cells have a negative impact on glioblastoma"

| Supplementary Table 6. Seurat based clustering of the Lymphoid population in the scRNA-seq analysis | | | | | |
|---|---|---|---|---|---|
| **Gene** | **avg_log2FC** | **pct.1** | **pct.2** | **p_val_adj** | **Cluster** |
| CD8A | 1.728216427 | 0.766 | 0.31 | 0 | 01_Exhausted_TCells |
| GZMK | 1.806354991 | 0.829 | 0.382 | 0 | 01_Exhausted_TCells |
| CCL5 | 1.095982852 | 0.934 | 0.635 | 1.07E-264 | 01_Exhausted_TCells |
| CD8B | 1.690659565 | 0.594 | 0.215 | 1.20E-261 | 01_Exhausted_TCells |
| CST7 | 1.199916268 | 0.869 | 0.559 | 7.91E-240 | 01_Exhausted_TCells |
| CRTAM | 2.245054326 | 0.364 | 0.101 | 4.32E-196 | 01_Exhausted_TCells |
| TUBA4A | 1.076889464 | 0.769 | 0.527 | 1.27E-145 | 01_Exhausted_TCells |
| GZMH | 1.106671161 | 0.505 | 0.257 | 1.05E-104 | 01_Exhausted_TCells |
| LYST | 1.032815641 | 0.549 | 0.312 | 9.97E-103 | 01_Exhausted_TCells |
| KLRB1 | -1.977129592 | 0.13 | 0.338 | 2.04E-75 | 01_Exhausted_TCells |
| IL7R | 1.443507286 | 0.907 | 0.505 | 6.51E-252 | 02_Th17-like_TCells |
| KLRB1 | 1.710512241 | 0.621 | 0.247 | 6.85E-196 | 02_Th17-like_TCells |
| CCR6 | 2.322857065 | 0.338 | 0.073 | 1.18E-189 | 02_Th17-like_TCells |
| CD40LG | 2.236561181 | 0.277 | 0.057 | 2.25E-158 | 02_Th17-like_TCells |
| ADAM19 | 1.874841662 | 0.384 | 0.117 | 4.25E-147 | 02_Th17-like_TCells |
| RORA | 1.098112432 | 0.694 | 0.383 | 7.01E-124 | 02_Th17-like_TCells |
| LTB | 1.145295704 | 0.595 | 0.293 | 3.04E-107 | 02_Th17-like_TCells |
| AQP3 | 1.628165574 | 0.341 | 0.114 | 5.67E-107 | 02_Th17-like_TCells |
| NKG7 | -2.079901646 | 0.309 | 0.551 | 3.32E-87 | 02_Th17-like_TCells |
| ERN1 | 1.132854776 | 0.533 | 0.277 | 4.89E-85 | 02_Th17-like_TCells |
| TOB1 | 1.075382248 | 0.487 | 0.247 | 7.81E-75 | 02_Th17-like_TCells |
| DUSP5 | 1.12340092 | 0.449 | 0.224 | 1.96E-70 | 02_Th17-like_TCells |
| ANKRD28 | 1.226114455 | 0.389 | 0.181 | 2.08E-69 | 02_Th17-like_TCells |
| AOAH | -1.802507128 | 0.263 | 0.487 | 1.63E-68 | 02_Th17-like_TCells |
| CXCR6 | 1.086992782 | 0.426 | 0.207 | 3.45E-67 | 02_Th17-like_TCells |
| FKBP11 | 1.061802451 | 0.414 | 0.2 | 1.46E-65 | 02_Th17-like_TCells |
| KLRK1 | -2.034244541 | 0.131 | 0.34 | 7.58E-58 | 02_Th17-like_TCells |
| CD8A | -1.3791096 | 0.218 | 0.426 | 1.41E-50 | 02_Th17-like_TCells |
| PLXDC2 | 4.093155492 | 0.719 | 0.13 | 0 | 03_APC |
| SLC1A3 | 4.373001956 | 0.663 | 0.09 | 0 | 03_APC |
| DOCK4 | 4.245486741 | 0.696 | 0.125 | 0 | 03_APC |
| LRMDA | 4.316157417 | 0.6 | 0.053 | 0 | 03_APC |
| RAB31 | 3.700108868 | 0.603 | 0.094 | 0 | 03_APC |
| SLC11A1 | 4.09973121 | 0.562 | 0.059 | 0 | 03_APC |
| ACSL1 | 3.559230673 | 0.636 | 0.135 | 0 | 03_APC |
| FMNL2 | 4.106327832 | 0.563 | 0.068 | 0 | 03_APC |
| FGD4 | 4.145510605 | 0.539 | 0.055 | 0 | 03_APC |
| RBM47 | 4.253596432 | 0.515 | 0.043 | 0 | 03_APC |
| FRMD4A | 3.484646316 | 0.583 | 0.116 | 0 | 03_APC |
| OLR1 | 3.422756122 | 0.576 | 0.112 | 0 | 03_APC |
| FMN1 | 4.129709629 | 0.519 | 0.057 | 0 | 03_APC |
| ARHGAP24 | 4.277003085 | 0.499 | 0.038 | 0 | 03_APC |
| IRAK3 | 3.274471144 | 0.499 | 0.057 | 0 | 03_APC |
| TLR2 | 3.48797807 | 0.488 | 0.052 | 0 | 03_APC |

| | | | | | |
|---|---|---|---|---|---|
| SLC8A1 | 4.056915404 | 0.468 | 0.042 | 0 | 03_APC |
| MSR1 | 3.348426413 | 0.487 | 0.065 | 0 | 03_APC |
| SRGAP1 | 4.098173768 | 0.455 | 0.034 | 0 | 03_APC |
| PLAUR | 3.270143657 | 0.551 | 0.134 | 0 | 03_APC |
| CSF2RA | 3.595664019 | 0.455 | 0.043 | 0 | 03_APC |
| SRGAP2 | 2.972303499 | 0.494 | 0.083 | 0 | 03_APC |
| ABCA1 | 4.073762961 | 0.452 | 0.043 | 0 | 03_APC |
| NHSL1 | 4.379032041 | 0.409 | 0.03 | 0 | 03_APC |
| GAB2 | 2.731515366 | 0.447 | 0.074 | 0 | 03_APC |
| TBXAS1 | 2.717087727 | 0.451 | 0.081 | 0 | 03_APC |
| IFNGR2 | 2.858728966 | 0.442 | 0.073 | 0 | 03_APC |
| UBE2E2 | 3.812841846 | 0.399 | 0.031 | 0 | 03_APC |
| C5AR1 | 2.877387696 | 0.443 | 0.078 | 0 | 03_APC |
| MEF2C | 3.174227922 | 0.423 | 0.06 | 0 | 03_APC |
| RNF144B | 3.64170746 | 0.407 | 0.044 | 0 | 03_APC |
| SRGAP2B | 3.116937631 | 0.413 | 0.055 | 0 | 03_APC |
| KCNMA1 | 3.850775318 | 0.391 | 0.038 | 0 | 03_APC |
| FAM49A | 2.985844392 | 0.403 | 0.052 | 0 | 03_APC |
| ITGAX | 3.500566378 | 0.392 | 0.044 | 0 | 03_APC |
| ABL2 | 2.841880242 | 0.41 | 0.064 | 0 | 03_APC |
| MAML3 | 2.676341966 | 0.397 | 0.059 | 0 | 03_APC |
| MERTK | 3.667308839 | 0.37 | 0.032 | 0 | 03_APC |
| RAB20 | 3.097780362 | 0.374 | 0.044 | 0 | 03_APC |
| FNIP2 | 3.114141655 | 0.384 | 0.056 | 0 | 03_APC |
| IRAK2 | 3.037743758 | 0.378 | 0.053 | 0 | 03_APC |
| TRIO | 3.11447171 | 0.37 | 0.046 | 0 | 03_APC |
| LHFPL2 | 2.953496337 | 0.373 | 0.049 | 0 | 03_APC |
| CSF3R | 4.610210313 | 0.339 | 0.017 | 0 | 03_APC |
| ZFHX3 | 2.944888474 | 0.347 | 0.041 | 0 | 03_APC |
| CD86 | 3.445645137 | 0.341 | 0.038 | 0 | 03_APC |
| TNS3 | 3.650119575 | 0.315 | 0.025 | 0 | 03_APC |
| SDK1 | 4.388481529 | 0.308 | 0.019 | 0 | 03_APC |
| KCNQ3 | 4.251491203 | 0.312 | 0.024 | 0 | 03_APC |
| ITSN1 | 3.677164315 | 0.307 | 0.022 | 0 | 03_APC |
| PALD1 | 3.558222033 | 0.29 | 0.026 | 0 | 03_APC |
| NRP2 | 3.924841904 | 0.283 | 0.021 | 0 | 03_APC |
| LNCAROD | 4.543537155 | 0.271 | 0.02 | 0 | 03_APC |
| PHACTR1 | 3.9602376 | 0.247 | 0.015 | 0 | 03_APC |
| ARHGAP6 | 3.901749283 | 0.248 | 0.016 | 0 | 03_APC |
| NEAT1 | 2.053923959 | 0.883 | 0.655 | 0 | 03_APC |
| MCTP1 | 3.591422271 | 0.273 | 0.022 | 3.29E-303 | 03_APC |
| PLA2G4A | 3.654874566 | 0.239 | 0.015 | 2.19E-301 | 03_APC |
| CD83 | 2.629431251 | 0.629 | 0.211 | 1.17E-300 | 03_APC |
| STAB1 | 3.246035633 | 0.305 | 0.032 | 5.74E-299 | 03_APC |
| TANC2 | 2.923441894 | 0.371 | 0.053 | 5.28E-297 | 03_APC |
| KYNU | 3.665053932 | 0.272 | 0.024 | 2.01E-294 | 03_APC |

| | | | | | |
|---|---|---|---|---|---|
| B3GNT5 | 2.883461833 | 0.348 | 0.047 | 2.08E-293 | 03_APC |
| FCGR2A | 2.667749528 | 0.42 | 0.074 | 6.04E-289 | 03_APC |
| SH3RF3 | 3.689038348 | 0.242 | 0.017 | 2.59E-288 | 03_APC |
| VEGFA | 3.258439548 | 0.321 | 0.039 | 3.54E-288 | 03_APC |
| PDGFB | 3.216855541 | 0.304 | 0.034 | 6.18E-288 | 03_APC |
| CPM | 3.146724681 | 0.34 | 0.046 | 1.78E-287 | 03_APC |
| CLEC7A | 3.244974711 | 0.328 | 0.042 | 1.08E-286 | 03_APC |
| SLC43A2 | 3.534113994 | 0.253 | 0.02 | 5.09E-286 | 03_APC |
| KLF4 | 3.233574375 | 0.307 | 0.035 | 1.52E-285 | 03_APC |
| PADI2 | 3.597409372 | 0.327 | 0.043 | 2.37E-285 | 03_APC |
| ADAM28 | 2.724864564 | 0.302 | 0.033 | 3.40E-283 | 03_APC |
| JDP2 | 2.989351733 | 0.34 | 0.047 | 2.54E-279 | 03_APC |
| EPB41L2 | 2.715700514 | 0.42 | 0.078 | 5.08E-277 | 03_APC |
| TCF4 | 3.308051283 | 0.284 | 0.03 | 1.62E-274 | 03_APC |
| MS4A7 | 2.334910944 | 0.523 | 0.13 | 1.74E-274 | 03_APC |
| BNC2 | 2.573510364 | 0.351 | 0.051 | 6.25E-273 | 03_APC |
| GNAQ | 2.393358048 | 0.531 | 0.138 | 2.75E-272 | 03_APC |
| NLRP3 | 3.112027764 | 0.457 | 0.103 | 4.77E-270 | 03_APC |
| TNFAIP2 | 3.611488728 | 0.216 | 0.013 | 2.53E-269 | 03_APC |
| EPB41L3 | 3.04169567 | 0.299 | 0.036 | 6.42E-269 | 03_APC |
| AXL | 3.35789943 | 0.253 | 0.023 | 2.28E-267 | 03_APC |
| RAPH1 | 3.109909792 | 0.273 | 0.028 | 1.44E-266 | 03_APC |
| AZIN1-AS1 | 4.024257754 | 0.287 | 0.034 | 4.66E-265 | 03_APC |
| SLCO2B1 | 2.682158702 | 0.331 | 0.047 | 8.87E-265 | 03_APC |
| CTTNBP2 | 3.769919547 | 0.217 | 0.014 | 1.34E-264 | 03_APC |
| LYN | 2.338319557 | 0.462 | 0.1 | 3.70E-264 | 03_APC |
| MMP19 | 4.336169448 | 0.248 | 0.023 | 1.75E-263 | 03_APC |
| CD163 | 2.94244236 | 0.388 | 0.072 | 2.00E-261 | 03_APC |
| IL1RAP | 3.15466593 | 0.314 | 0.043 | 1.50E-260 | 03_APC |
| AL078590.2 | 3.268865867 | 0.238 | 0.02 | 1.63E-257 | 03_APC |
| MTSS1 | 2.623153047 | 0.432 | 0.091 | 2.07E-257 | 03_APC |
| MYO1E | 3.11783978 | 0.325 | 0.047 | 5.20E-257 | 03_APC |
| MEF2A | 2.366447118 | 0.543 | 0.154 | 1.01E-254 | 03_APC |
| MITF | 3.681810596 | 0.229 | 0.019 | 2.32E-254 | 03_APC |
| NFKBID | 2.222875199 | 0.416 | 0.081 | 2.52E-252 | 03_APC |
| SGK1 | 2.461221387 | 0.6 | 0.203 | 4.07E-251 | 03_APC |
| FHIT | 2.542612233 | 0.375 | 0.067 | 1.07E-249 | 03_APC |
| QKI | 2.261817978 | 0.623 | 0.213 | 7.38E-249 | 03_APC |
| MANBA | 2.201820721 | 0.485 | 0.122 | 1.27E-243 | 03_APC |
| RIN2 | 3.545134463 | 0.229 | 0.02 | 1.70E-242 | 03_APC |
| C3 | 1.970455398 | 0.548 | 0.161 | 1.45E-240 | 03_APC |
| CSF1R | 2.272376252 | 0.376 | 0.07 | 1.50E-236 | 03_APC |
| BASP1 | 2.754978931 | 0.348 | 0.062 | 1.07E-234 | 03_APC |
| VASH1 | 2.821352191 | 0.273 | 0.035 | 9.78E-232 | 03_APC |
| SOCS6 | 2.978039124 | 0.252 | 0.029 | 4.23E-231 | 03_APC |
| MAFB | 2.610592559 | 0.389 | 0.081 | 3.78E-228 | 03_APC |

| | | | | | |
|---|---|---|---|---|---|
| ETS2 | 2.226875263 | 0.456 | 0.113 | 1.39E-227 | 03_APC |
| APOE | 2.01707425 | 0.782 | 0.458 | 7.60E-227 | 03_APC |
| CEP170 | 2.462610924 | 0.344 | 0.061 | 3.18E-226 | 03_APC |
| LPCAT2 | 2.833122518 | 0.282 | 0.04 | 5.26E-223 | 03_APC |
| RFX2 | 2.765210557 | 0.268 | 0.035 | 1.70E-222 | 03_APC |
| KLF7 | 2.809143403 | 0.263 | 0.034 | 2.25E-221 | 03_APC |
| PEAK1 | 2.270516003 | 0.394 | 0.086 | 1.67E-219 | 03_APC |
| DENND5A | 2.783525132 | 0.288 | 0.043 | 8.82E-218 | 03_APC |
| ALCAM | 2.626616323 | 0.362 | 0.074 | 7.86E-215 | 03_APC |
| TTYH3 | 2.70663051 | 0.246 | 0.03 | 3.19E-214 | 03_APC |
| FPR1 | 2.523744286 | 0.31 | 0.052 | 2.36E-213 | 03_APC |
| MAP4K3 | 2.328337812 | 0.375 | 0.079 | 2.53E-213 | 03_APC |
| PPARD | 2.34268783 | 0.335 | 0.061 | 6.93E-212 | 03_APC |
| RHBDF2 | 1.991524391 | 0.449 | 0.114 | 7.05E-212 | 03_APC |
| LILRB4 | 2.332004699 | 0.32 | 0.056 | 1.03E-211 | 03_APC |
| DST | 3.133901114 | 0.25 | 0.032 | 1.94E-211 | 03_APC |
| ST6GALNAC3 | 3.14176569 | 0.257 | 0.035 | 7.26E-211 | 03_APC |
| SPTLC2 | 2.270442743 | 0.477 | 0.136 | 4.12E-210 | 03_APC |
| GNG7 | 3.036522451 | 0.242 | 0.03 | 7.48E-210 | 03_APC |
| ADAP2 | 2.717462974 | 0.261 | 0.036 | 5.22E-208 | 03_APC |
| SERPINE1 | 3.716685378 | 0.389 | 0.099 | 4.29E-206 | 03_APC |
| SPI1 | 2.092152148 | 0.412 | 0.098 | 5.29E-206 | 03_APC |
| CHKA | 2.114341241 | 0.315 | 0.055 | 4.65E-205 | 03_APC |
| MS4A4A | 2.37595032 | 0.325 | 0.061 | 2.30E-204 | 03_APC |
| HCK | 3.07054697 | 0.238 | 0.03 | 2.85E-203 | 03_APC |
| LIMS1 | 2.128337861 | 0.608 | 0.25 | 1.84E-202 | 03_APC |
| MIR181A1HG | 2.280417866 | 0.313 | 0.056 | 2.38E-202 | 03_APC |
| GK | 2.244686806 | 0.367 | 0.08 | 4.56E-202 | 03_APC |
| ALOX5 | 2.88611165 | 0.245 | 0.033 | 6.66E-201 | 03_APC |
| DSE | 2.145475004 | 0.39 | 0.092 | 5.88E-200 | 03_APC |
| SWAP70 | 2.444876416 | 0.279 | 0.044 | 1.44E-198 | 03_APC |
| SH2B3 | 2.114872223 | 0.341 | 0.068 | 4.24E-198 | 03_APC |
| CTSB | 1.870460579 | 0.673 | 0.326 | 1.74E-197 | 03_APC |
| LGMN | 2.465643095 | 0.325 | 0.064 | 5.36E-196 | 03_APC |
| SLC25A37 | 2.33176593 | 0.335 | 0.068 | 1.64E-195 | 03_APC |
| TBC1D16 | 2.845258011 | 0.238 | 0.031 | 2.67E-195 | 03_APC |
| ZMIZ1 | 2.31356392 | 0.313 | 0.059 | 2.15E-192 | 03_APC |
| TRIB1 | 2.656412468 | 0.244 | 0.034 | 2.61E-192 | 03_APC |
| SLC31A2 | 2.439851603 | 0.286 | 0.049 | 9.43E-192 | 03_APC |
| CYFIP1 | 2.130917009 | 0.35 | 0.076 | 9.69E-189 | 03_APC |
| IL18 | 2.247083758 | 0.294 | 0.053 | 2.86E-188 | 03_APC |
| IL6R | 2.502258335 | 0.267 | 0.043 | 1.34E-186 | 03_APC |
| LDLRAD4 | 1.69783442 | 0.725 | 0.409 | 1.35E-186 | 03_APC |
| ADAMTSL4-A | 2.047632657 | 0.452 | 0.13 | 2.13E-186 | 03_APC |
| CD14 | 2.194580475 | 0.394 | 0.101 | 7.67E-183 | 03_APC |
| NR4A1 | 1.758308225 | 0.625 | 0.271 | 8.42E-183 | 03_APC |

| | | | | | |
|---|---|---|---|---|---|
| RHOB | 2.293956271 | 0.573 | 0.235 | 6.15E-182 | 03_APC |
| GSN | 1.957452795 | 0.442 | 0.126 | 1.12E-181 | 03_APC |
| ITGAV | 2.016858243 | 0.41 | 0.108 | 7.42E-181 | 03_APC |
| C1QC | 1.716528491 | 0.672 | 0.328 | 1.79E-180 | 03_APC |
| MARCKS | 1.937142925 | 0.42 | 0.115 | 1.10E-178 | 03_APC |
| SH3TC1 | 1.962040127 | 0.319 | 0.065 | 1.54E-176 | 03_APC |
| CSGALNACT1 | 2.028441901 | 0.389 | 0.1 | 2.35E-176 | 03_APC |
| HLA-DRA | 1.757727305 | 0.766 | 0.492 | 4.18E-175 | 03_APC |
| ATG7 | 1.990351836 | 0.423 | 0.118 | 1.19E-174 | 03_APC |
| IFI30 | 2.00247728 | 0.532 | 0.195 | 1.51E-174 | 03_APC |
| IL13RA1 | 2.465730577 | 0.254 | 0.042 | 1.94E-173 | 03_APC |
| CTSL | 2.365825739 | 0.32 | 0.07 | 2.98E-172 | 03_APC |
| DAGLB | 2.471668628 | 0.25 | 0.042 | 4.77E-171 | 03_APC |
| CPVL | 2.137374745 | 0.279 | 0.052 | 5.41E-171 | 03_APC |
| IER3 | 2.087613699 | 0.483 | 0.163 | 8.74E-171 | 03_APC |
| ATP8B4 | 1.914570717 | 0.273 | 0.049 | 1.57E-170 | 03_APC |
| CXCL16 | 1.81646381 | 0.382 | 0.101 | 5.25E-163 | 03_APC |
| SPP1 | 1.742372485 | 0.802 | 0.539 | 1.23E-162 | 03_APC |
| OGFRL1 | 1.929344204 | 0.366 | 0.095 | 2.49E-160 | 03_APC |
| C9orf72 | 1.927046228 | 0.379 | 0.102 | 9.41E-160 | 03_APC |
| ABR | 1.766818287 | 0.455 | 0.146 | 9.22E-158 | 03_APC |
| SBF2 | 2.071946853 | 0.361 | 0.094 | 1.40E-156 | 03_APC |
| SRGAP2C | 2.099993553 | 0.311 | 0.071 | 2.25E-156 | 03_APC |
| MPP1 | 2.122464425 | 0.271 | 0.054 | 5.63E-155 | 03_APC |
| PDE8A | 2.154812898 | 0.369 | 0.102 | 7.25E-155 | 03_APC |
| CYBB | 2.097857935 | 0.299 | 0.066 | 1.02E-153 | 03_APC |
| TREM2 | 1.818830842 | 0.38 | 0.106 | 9.42E-153 | 03_APC |
| NAMPT | 1.911442902 | 0.618 | 0.3 | 4.61E-151 | 03_APC |
| PRKAG2 | 1.887383427 | 0.357 | 0.096 | 3.57E-150 | 03_APC |
| ICAM1 | 2.067815919 | 0.349 | 0.095 | 2.57E-147 | 03_APC |
| CST3 | 1.698233183 | 0.633 | 0.317 | 9.41E-147 | 03_APC |
| ADGRE2 | 2.043940568 | 0.245 | 0.045 | 1.29E-146 | 03_APC |
| ELL2 | 1.672651607 | 0.727 | 0.496 | 1.93E-146 | 03_APC |
| HIF1A | 1.607659653 | 0.618 | 0.299 | 3.10E-146 | 03_APC |
| RASSF4 | 2.392235019 | 0.254 | 0.05 | 1.65E-145 | 03_APC |
| CUX1 | 1.676792496 | 0.439 | 0.142 | 1.67E-145 | 03_APC |
| ARHGAP26 | 1.606980192 | 0.658 | 0.329 | 1.76E-145 | 03_APC |
| MFSD1 | 1.846166392 | 0.369 | 0.106 | 1.03E-143 | 03_APC |
| MS4A6A | 1.855958076 | 0.385 | 0.115 | 8.08E-143 | 03_APC |
| ARHGAP21 | 2.278026497 | 0.249 | 0.049 | 5.92E-142 | 03_APC |
| FCER1G | 1.365643017 | 0.605 | 0.273 | 7.94E-142 | 03_APC |
| MTHFD1L | 2.064233732 | 0.255 | 0.052 | 9.19E-141 | 03_APC |
| BMP2K | 1.949546967 | 0.313 | 0.078 | 3.09E-140 | 03_APC |
| C1QB | 1.449928047 | 0.636 | 0.314 | 2.02E-139 | 03_APC |
| CPEB4 | 2.115149397 | 0.304 | 0.075 | 5.45E-139 | 03_APC |
| ATP13A3 | 1.841112676 | 0.385 | 0.118 | 2.46E-138 | 03_APC |

| | | | | | |
|---|---|---|---|---|---|
| C1QA | 1.570499082 | 0.598 | 0.285 | 5.22E-138 | 03_APC |
| SDCCAG8 | 1.940681354 | 0.336 | 0.091 | 1.72E-137 | 03_APC |
| ATF6 | 1.949565597 | 0.442 | 0.154 | 4.46E-137 | 03_APC |
| ELMO1 | 1.642998032 | 0.649 | 0.354 | 4.07E-136 | 03_APC |
| NUMB | 1.921510239 | 0.38 | 0.117 | 1.89E-134 | 03_APC |
| PDK4 | 2.491145326 | 0.335 | 0.096 | 2.04E-134 | 03_APC |
| PARVB | 1.940802206 | 0.306 | 0.078 | 5.77E-134 | 03_APC |
| GNA12 | 2.10205859 | 0.262 | 0.058 | 1.99E-133 | 03_APC |
| APOC1 | 1.761898722 | 0.583 | 0.278 | 3.47E-132 | 03_APC |
| MYO9B | 1.548420677 | 0.549 | 0.235 | 2.79E-131 | 03_APC |
| GRASP | 1.201509069 | 0.417 | 0.136 | 9.81E-131 | 03_APC |
| RAPGEF1 | 1.536618371 | 0.602 | 0.294 | 1.36E-129 | 03_APC |
| PDE4DIP | 2.087388773 | 0.405 | 0.14 | 1.58E-129 | 03_APC |
| PLTP | 1.933011419 | 0.348 | 0.104 | 1.38E-128 | 03_APC |
| KIF1B | 1.820057798 | 0.352 | 0.105 | 1.29E-127 | 03_APC |
| DENND3 | 1.782255459 | 0.292 | 0.074 | 6.64E-127 | 03_APC |
| HLA-DRB1 | 1.41974161 | 0.767 | 0.522 | 5.93E-126 | 03_APC |
| ADAM17 | 1.646752685 | 0.343 | 0.1 | 1.11E-124 | 03_APC |
| CCL3L1 | 2.708451881 | 0.422 | 0.166 | 2.67E-124 | 03_APC |
| ZBTB16 | 1.492175531 | 0.511 | 0.207 | 5.83E-124 | 03_APC |
| CEBPD | 1.70425818 | 0.584 | 0.295 | 9.85E-124 | 03_APC |
| KLHL6 | 1.641134401 | 0.346 | 0.103 | 2.54E-122 | 03_APC |
| DLEU1 | 1.747785201 | 0.404 | 0.141 | 7.96E-122 | 03_APC |
| IRS2 | 1.747896531 | 0.32 | 0.09 | 9.55E-122 | 03_APC |
| IFITM1 | -1.224217619 | 0.584 | 0.799 | 2.28E-121 | 03_APC |
| HMOX1 | 2.151515075 | 0.301 | 0.083 | 1.17E-120 | 03_APC |
| CCL3 | 2.503015535 | 0.651 | 0.427 | 2.41E-120 | 03_APC |
| FCGR1A | 1.96756642 | 0.268 | 0.066 | 2.76E-120 | 03_APC |
| RNF130 | 1.510120406 | 0.409 | 0.141 | 3.45E-120 | 03_APC |
| HLA-DQA1 | 1.623141491 | 0.492 | 0.207 | 6.76E-119 | 03_APC |
| TCF12 | 1.598137804 | 0.525 | 0.232 | 5.66E-117 | 03_APC |
| ZSWIM6 | 1.355291534 | 0.616 | 0.307 | 9.98E-117 | 03_APC |
| CXCL8 | 2.49939369 | 0.309 | 0.093 | 9.48E-116 | 03_APC |
| CD68 | 1.600593106 | 0.395 | 0.138 | 2.11E-115 | 03_APC |
| PSAP | 1.296833285 | 0.688 | 0.424 | 2.73E-115 | 03_APC |
| DISC1 | 1.912239776 | 0.281 | 0.075 | 3.53E-115 | 03_APC |
| USP53 | 1.939904915 | 0.317 | 0.095 | 1.99E-114 | 03_APC |
| SIPA1L1 | 1.516318151 | 0.629 | 0.356 | 3.57E-114 | 03_APC |
| FCGBP | 2.232272132 | 0.275 | 0.075 | 1.30E-113 | 03_APC |
| SNX29 | 1.671423737 | 0.346 | 0.108 | 3.32E-113 | 03_APC |
| CHSY1 | 1.579514418 | 0.311 | 0.09 | 5.06E-113 | 03_APC |
| NR4A3 | 1.730829571 | 0.556 | 0.278 | 1.89E-112 | 03_APC |
| CH25H | 2.033289277 | 0.281 | 0.078 | 2.91E-112 | 03_APC |
| NCOR2 | 1.599493784 | 0.366 | 0.121 | 1.09E-111 | 03_APC |
| TYROBP | 1.11256294 | 0.677 | 0.377 | 1.27E-111 | 03_APC |
| TEX14 | 2.070334607 | 0.409 | 0.157 | 2.13E-109 | 03_APC |

| | | | | | |
|---|---|---|---|---|---|
| FCGRT | 1.526289923 | 0.354 | 0.116 | 3.41E-109 | 03_APC |
| UBASH3B | 1.516078991 | 0.395 | 0.144 | 2.29E-108 | 03_APC |
| RABGEF1 | 1.489372735 | 0.468 | 0.195 | 1.02E-107 | 03_APC |
| RHOQ | 1.711786684 | 0.288 | 0.082 | 2.97E-106 | 03_APC |
| TFRC | 1.997226255 | 0.36 | 0.127 | 4.02E-106 | 03_APC |
| ANKS1A | 1.644098495 | 0.279 | 0.078 | 5.50E-105 | 03_APC |
| ETV6 | 1.332176937 | 0.475 | 0.2 | 1.86E-103 | 03_APC |
| FCHSD2 | 1.638251794 | 0.375 | 0.136 | 2.83E-101 | 03_APC |
| MED13L | 1.37168929 | 0.51 | 0.226 | 5.17E-101 | 03_APC |
| PELI1 | 1.470452955 | 0.494 | 0.221 | 8.88E-101 | 03_APC |
| FOXO3 | 1.41467019 | 0.4 | 0.15 | 4.18E-99 | 03_APC |
| HLA-DQB1 | 1.323048162 | 0.558 | 0.287 | 1.07E-98 | 03_APC |
| PLEKHG2 | 1.730436038 | 0.289 | 0.088 | 2.08E-97 | 03_APC |
| ASAH1 | 1.450119711 | 0.411 | 0.163 | 8.48E-97 | 03_APC |
| SOD2 | 1.73881024 | 0.534 | 0.28 | 1.41E-96 | 03_APC |
| GRAMD1B | 1.52062644 | 0.398 | 0.155 | 3.54E-96 | 03_APC |
| ITPR2 | 1.304728732 | 0.473 | 0.202 | 7.45E-95 | 03_APC |
| SUSD6 | 1.258359446 | 0.431 | 0.174 | 3.49E-94 | 03_APC |
| TET2 | 1.750800041 | 0.365 | 0.136 | 5.16E-94 | 03_APC |
| FNDC3B | 1.279323816 | 0.431 | 0.177 | 1.63E-92 | 03_APC |
| DIAPH2 | 1.418700882 | 0.389 | 0.149 | 4.41E-92 | 03_APC |
| GLUL | 1.362484849 | 0.559 | 0.288 | 4.36E-91 | 03_APC |
| ZFAND3 | 1.107339909 | 0.603 | 0.318 | 1.11E-90 | 03_APC |
| NPC2 | 1.221465723 | 0.556 | 0.293 | 6.56E-90 | 03_APC |
| MAN1A1 | 1.464151582 | 0.385 | 0.15 | 1.20E-89 | 03_APC |
| GRN | 1.472690461 | 0.345 | 0.127 | 2.32E-89 | 03_APC |
| RIN3 | 1.233164842 | 0.361 | 0.133 | 2.99E-89 | 03_APC |
| APLP2 | 1.308696689 | 0.359 | 0.133 | 5.53E-87 | 03_APC |
| SPIDR | 1.242444363 | 0.462 | 0.206 | 9.95E-87 | 03_APC |
| AP1B1 | 1.405870093 | 0.326 | 0.116 | 5.12E-86 | 03_APC |
| RB1 | 1.205491447 | 0.446 | 0.193 | 1.43E-84 | 03_APC |
| HLA-DMA | 1.280703683 | 0.424 | 0.185 | 4.56E-84 | 03_APC |
| ST6GAL1 | 1.48660371 | 0.396 | 0.164 | 4.70E-84 | 03_APC |
| ATF3 | 1.527949057 | 0.426 | 0.192 | 1.27E-83 | 03_APC |
| AIF1 | 1.202524991 | 0.434 | 0.191 | 9.02E-82 | 03_APC |
| EGR1 | 1.635011163 | 0.383 | 0.162 | 9.76E-82 | 03_APC |
| RAB1A | 1.097895848 | 0.475 | 0.216 | 1.17E-81 | 03_APC |
| BCL2A1 | 1.554793573 | 0.337 | 0.128 | 1.86E-81 | 03_APC |
| MAML2 | 1.231080713 | 0.615 | 0.353 | 7.77E-81 | 03_APC |
| RERE | 1.12718154 | 0.439 | 0.19 | 1.11E-80 | 03_APC |
| SPAG9 | 1.261306336 | 0.441 | 0.2 | 1.44E-79 | 03_APC |
| GNA13 | 1.174758157 | 0.514 | 0.266 | 4.28E-79 | 03_APC |
| ARL8B | 1.317010664 | 0.371 | 0.15 | 2.97E-78 | 03_APC |
| HLA-DRB5 | 1.27005305 | 0.573 | 0.322 | 2.70E-77 | 03_APC |
| PRKCE | 1.138648516 | 0.353 | 0.137 | 6.55E-77 | 03_APC |
| DPYD | 1.294586764 | 0.577 | 0.331 | 7.62E-77 | 03_APC |

| | | | | | |
|---|---|---|---|---|---|
| SLC9A9 | 1.279985512 | 0.367 | 0.148 | 8.47E-77 | 03_APC |
| SFMBT2 | 1.074730051 | 0.533 | 0.278 | 1.29E-76 | 03_APC |
| A2M | 1.172101332 | 0.408 | 0.177 | 1.29E-75 | 03_APC |
| HAVCR2 | 1.176917893 | 0.351 | 0.141 | 5.23E-73 | 03_APC |
| PLEK | 1.450346015 | 0.363 | 0.154 | 1.64E-72 | 03_APC |
| TOM1 | 1.225690117 | 0.347 | 0.139 | 2.26E-72 | 03_APC |
| PLSCR1 | 1.075755234 | 0.381 | 0.161 | 5.92E-72 | 03_APC |
| TSC22D2 | 1.33880163 | 0.431 | 0.203 | 1.26E-70 | 03_APC |
| CTNNB1 | 1.064897678 | 0.483 | 0.243 | 4.23E-70 | 03_APC |
| IL1B | 2.377246252 | 0.457 | 0.256 | 7.03E-70 | 03_APC |
| CTSZ | 1.227016322 | 0.356 | 0.15 | 1.08E-68 | 03_APC |
| PIK3R5 | 1.10034043 | 0.543 | 0.302 | 2.57E-68 | 03_APC |
| SNX9 | 1.117926709 | 0.545 | 0.307 | 4.74E-68 | 03_APC |
| RASGEF1B | 1.06868616 | 0.554 | 0.313 | 1.07E-67 | 03_APC |
| AC020916.1 | 1.031611834 | 0.481 | 0.244 | 2.60E-65 | 03_APC |
| DLEU2 | 1.083537864 | 0.456 | 0.224 | 2.41E-64 | 03_APC |
| B4GALT1 | 1.086157162 | 0.556 | 0.322 | 9.46E-64 | 03_APC |
| SERPINB9 | 1.07369255 | 0.565 | 0.347 | 1.11E-62 | 03_APC |
| MGAT1 | 1.213945758 | 0.387 | 0.183 | 1.53E-60 | 03_APC |
| MGAT5 | 1.124810424 | 0.465 | 0.24 | 1.54E-59 | 03_APC |
| TYMP | 1.105554713 | 0.398 | 0.192 | 4.89E-59 | 03_APC |
| TAOK3 | 1.043442914 | 0.488 | 0.265 | 2.77E-57 | 03_APC |
| PTPN1 | 1.07725828 | 0.506 | 0.288 | 3.39E-56 | 03_APC |
| CEBPB | 1.038942859 | 0.501 | 0.29 | 2.06E-55 | 03_APC |
| RAB7A | 1.163324111 | 0.545 | 0.342 | 7.81E-44 | 03_APC |
| HSPA1B | 2.58860749 | 0.708 | 0.391 | 7.31E-204 | 04_HSP_high |
| HSPA1A | 2.439603258 | 0.71 | 0.465 | 1.96E-155 | 04_HSP_high |
| METRNL | -1.652148717 | 0.114 | 0.365 | 3.82E-57 | 04_HSP_high |
| MAP3K8 | -1.161618911 | 0.141 | 0.376 | 1.95E-45 | 04_HSP_high |
| FOSL2 | -1.075986119 | 0.14 | 0.374 | 2.43E-45 | 04_HSP_high |
| CCR7 | 2.896402198 | 0.5 | 0.108 | 4.78E-225 | 05_Central_Memory_Tcells |
| IL7R | 1.390357866 | 0.952 | 0.526 | 1.41E-178 | 05_Central_Memory_Tcells |
| SELL | 2.561934888 | 0.457 | 0.119 | 1.13E-160 | 05_Central_Memory_Tcells |
| LEF1 | 2.659806925 | 0.359 | 0.074 | 9.05E-157 | 05_Central_Memory_Tcells |
| MAL | 2.879252769 | 0.261 | 0.044 | 1.18E-132 | 05_Central_Memory_Tcells |
| NKG7 | -3.848663174 | 0.108 | 0.553 | 1.37E-120 | 05_Central_Memory_Tcells |
| CCL5 | -2.191746007 | 0.372 | 0.72 | 4.23E-110 | 05_Central_Memory_Tcells |
| KLF2 | 1.474800601 | 0.726 | 0.398 | 4.43E-107 | 05_Central_Memory_Tcells |
| CST7 | -1.960523873 | 0.28 | 0.647 | 5.07E-91 | 05_Central_Memory_Tcells |
| TCF7 | 1.844961003 | 0.357 | 0.118 | 2.39E-82 | 05_Central_Memory_Tcells |
| LTB | 1.329434552 | 0.602 | 0.312 | 1.69E-75 | 05_Central_Memory_Tcells |
| ANK3 | 1.621321728 | 0.498 | 0.221 | 2.42E-75 | 05_Central_Memory_Tcells |
| ZEB2 | -2.74025967 | 0.119 | 0.462 | 3.77E-74 | 05_Central_Memory_Tcells |
| GZMA | -1.85390469 | 0.228 | 0.571 | 4.08E-73 | 05_Central_Memory_Tcells |
| AOAH | -2.453281488 | 0.143 | 0.483 | 1.17E-71 | 05_Central_Memory_Tcells |
| GZMH | -4.534409668 | 0.027 | 0.329 | 1.38E-65 | 05_Central_Memory_Tcells |

| | | | | | |
|---|---|---|---|---|---|
| TOB1 | 1.47089398 | 0.519 | 0.26 | 2.35E-64 | 05_Central_Memory_Tcells |
| SESN3 | 2.228693911 | 0.316 | 0.114 | 3.69E-64 | 05_Central_Memory_Tcells |
| CCL4 | -2.050819708 | 0.566 | 0.773 | 1.01E-60 | 05_Central_Memory_Tcells |
| ANXA1 | 1.062307453 | 0.803 | 0.577 | 2.56E-59 | 05_Central_Memory_Tcells |
| BACH2 | 1.176780254 | 0.502 | 0.246 | 7.79E-58 | 05_Central_Memory_Tcells |
| PRF1 | -2.621228048 | 0.091 | 0.379 | 1.06E-54 | 05_Central_Memory_Tcells |
| SOCS3 | 1.536617298 | 0.388 | 0.175 | 3.57E-54 | 05_Central_Memory_Tcells |
| SLC2A3 | 1.011191584 | 0.799 | 0.586 | 4.50E-53 | 05_Central_Memory_Tcells |
| CD8A | -1.986958799 | 0.136 | 0.42 | 2.50E-51 | 05_Central_Memory_Tcells |
| CRYBG1 | 1.053281232 | 0.744 | 0.52 | 2.59E-51 | 05_Central_Memory_Tcells |
| HLA-DRB1 | -1.819656425 | 0.344 | 0.573 | 1.48E-49 | 05_Central_Memory_Tcells |
| CD55 | 1.092643989 | 0.62 | 0.4 | 3.52E-47 | 05_Central_Memory_Tcells |
| GZMB | -4.121475073 | 0.024 | 0.253 | 3.46E-44 | 05_Central_Memory_Tcells |
| GZMK | -1.400819758 | 0.23 | 0.489 | 8.97E-43 | 05_Central_Memory_Tcells |
| HLA-DPA1 | -1.486737895 | 0.305 | 0.542 | 9.57E-41 | 05_Central_Memory_Tcells |
| TOX | -1.972890503 | 0.099 | 0.342 | 1.49E-39 | 05_Central_Memory_Tcells |
| GPR183 | 1.08468455 | 0.558 | 0.34 | 1.76E-39 | 05_Central_Memory_Tcells |
| LMNA | 1.130516869 | 0.489 | 0.281 | 2.79E-37 | 05_Central_Memory_Tcells |
| HLA-DRA | -1.783842571 | 0.339 | 0.545 | 5.59E-37 | 05_Central_Memory_Tcells |
| LYST | -1.813311201 | 0.149 | 0.376 | 2.02E-34 | 05_Central_Memory_Tcells |
| CTSW | -1.519082305 | 0.251 | 0.473 | 4.77E-34 | 05_Central_Memory_Tcells |
| HLA-DPB1 | -1.428440344 | 0.269 | 0.481 | 1.58E-32 | 05_Central_Memory_Tcells |
| KLRK1 | -1.659712993 | 0.112 | 0.328 | 7.58E-32 | 05_Central_Memory_Tcells |
| AKNA | -1.133004262 | 0.325 | 0.531 | 7.55E-29 | 05_Central_Memory_Tcells |
| METRNL | -1.783863412 | 0.149 | 0.352 | 1.59E-28 | 05_Central_Memory_Tcells |
| CTSC | -1.377192375 | 0.182 | 0.395 | 2.39E-28 | 05_Central_Memory_Tcells |
| GLCCI1 | -1.665598107 | 0.133 | 0.333 | 1.99E-27 | 05_Central_Memory_Tcells |
| GZMB | 3.460354678 | 0.873 | 0.182 | 0 | 06_Cytotoxic_Tcells |
| GNLY | 3.456716525 | 0.798 | 0.153 | 0 | 06_Cytotoxic_Tcells |
| FGFBP2 | 5.829982876 | 0.588 | 0.019 | 0 | 06_Cytotoxic_Tcells |
| KLRD1 | 2.663766525 | 0.719 | 0.163 | 0 | 06_Cytotoxic_Tcells |
| PRF1 | 2.860002709 | 0.845 | 0.316 | 0 | 06_Cytotoxic_Tcells |
| SPON2 | 3.46422579 | 0.69 | 0.173 | 0 | 06_Cytotoxic_Tcells |
| NKG7 | 2.718405948 | 0.945 | 0.483 | 0 | 06_Cytotoxic_Tcells |
| S1PR5 | 4.292371543 | 0.394 | 0.028 | 0 | 06_Cytotoxic_Tcells |
| KLRF1 | 3.976430582 | 0.352 | 0.023 | 0 | 06_Cytotoxic_Tcells |
| PRSS23 | 4.375751784 | 0.22 | 0.012 | 5.16E-251 | 06_Cytotoxic_Tcells |
| IGFBP7 | 3.360090508 | 0.322 | 0.034 | 8.32E-247 | 06_Cytotoxic_Tcells |
| FCGR3A | 2.84297225 | 0.569 | 0.15 | 1.07E-223 | 06_Cytotoxic_Tcells |
| OSBPL5 | 3.58816562 | 0.239 | 0.021 | 3.64E-205 | 06_Cytotoxic_Tcells |
| C1orf21 | 2.554508277 | 0.38 | 0.07 | 6.83E-180 | 06_Cytotoxic_Tcells |
| CLIC3 | 2.711074522 | 0.454 | 0.107 | 2.15E-178 | 06_Cytotoxic_Tcells |
| ADGRG1 | 2.71826707 | 0.401 | 0.084 | 2.02E-175 | 06_Cytotoxic_Tcells |
| CTSW | 1.801089563 | 0.808 | 0.426 | 5.84E-166 | 06_Cytotoxic_Tcells |
| GZMH | 1.885448455 | 0.681 | 0.274 | 1.34E-152 | 06_Cytotoxic_Tcells |
| TTC38 | 2.98993729 | 0.274 | 0.044 | 2.72E-147 | 06_Cytotoxic_Tcells |

| | | | | | |
|---|---|---|---|---|---|
| CST7 | 1.339532269 | 0.885 | 0.596 | 3.20E-144 | 06_Cytotoxic_Tcells |
| RAP1GAP2 | 2.792515306 | 0.238 | 0.034 | 1.06E-139 | 06_Cytotoxic_Tcells |
| PLAC8 | 2.500930028 | 0.37 | 0.09 | 8.30E-133 | 06_Cytotoxic_Tcells |
| EFHD2 | 2.009978501 | 0.553 | 0.258 | 5.21E-102 | 06_Cytotoxic_Tcells |
| ZEB2 | 1.177728275 | 0.748 | 0.409 | 4.81E-91 | 06_Cytotoxic_Tcells |
| MCTP2 | 1.695554436 | 0.431 | 0.152 | 4.85E-90 | 06_Cytotoxic_Tcells |
| CD247 | 1.34355867 | 0.76 | 0.54 | 4.86E-90 | 06_Cytotoxic_Tcells |
| PTPN12 | 2.166267975 | 0.275 | 0.074 | 2.91E-80 | 06_Cytotoxic_Tcells |
| ITGB2 | 1.273884237 | 0.721 | 0.499 | 1.55E-77 | 06_Cytotoxic_Tcells |
| HOPX | 2.056500934 | 0.382 | 0.144 | 1.73E-74 | 06_Cytotoxic_Tcells |
| COTL1 | -1.647460178 | 0.233 | 0.596 | 7.91E-73 | 06_Cytotoxic_Tcells |
| HSH2D | 1.980066334 | 0.3 | 0.096 | 1.32E-69 | 06_Cytotoxic_Tcells |
| IL7R | -1.594803775 | 0.238 | 0.586 | 1.49E-69 | 06_Cytotoxic_Tcells |
| CD7 | 1.03272896 | 0.779 | 0.545 | 9.26E-69 | 06_Cytotoxic_Tcells |
| TPST2 | 1.861093104 | 0.347 | 0.132 | 1.46E-64 | 06_Cytotoxic_Tcells |
| APMAP | 1.467447022 | 0.492 | 0.245 | 5.22E-64 | 06_Cytotoxic_Tcells |
| GZMK | -1.701528337 | 0.16 | 0.494 | 8.70E-61 | 06_Cytotoxic_Tcells |
| FLNA | 1.298320737 | 0.589 | 0.349 | 1.01E-60 | 06_Cytotoxic_Tcells |
| CAMK4 | -2.064471036 | 0.107 | 0.436 | 1.09E-60 | 06_Cytotoxic_Tcells |
| LDLRAD4 | -2.336700811 | 0.165 | 0.475 | 3.79E-60 | 06_Cytotoxic_Tcells |
| PXN | 1.629665771 | 0.352 | 0.142 | 6.58E-57 | 06_Cytotoxic_Tcells |
| AREG | 1.400993963 | 0.562 | 0.328 | 3.01E-54 | 06_Cytotoxic_Tcells |
| LTB | -2.897658904 | 0.071 | 0.356 | 4.40E-54 | 06_Cytotoxic_Tcells |
| BIN2 | 1.380594671 | 0.514 | 0.293 | 2.04E-53 | 06_Cytotoxic_Tcells |
| SAMSN1 | -1.325019486 | 0.382 | 0.653 | 4.58E-52 | 06_Cytotoxic_Tcells |
| PLEK | 1.401329119 | 0.377 | 0.166 | 2.16E-50 | 06_Cytotoxic_Tcells |
| ICOS | -2.861445839 | 0.07 | 0.342 | 8.75E-50 | 06_Cytotoxic_Tcells |
| PDE4B | -1.320182283 | 0.281 | 0.578 | 1.23E-49 | 06_Cytotoxic_Tcells |
| DUSP4 | -2.060342082 | 0.104 | 0.388 | 2.35E-49 | 06_Cytotoxic_Tcells |
| METRNL | 1.294985681 | 0.543 | 0.319 | 2.93E-48 | 06_Cytotoxic_Tcells |
| GPR183 | -2.213834123 | 0.107 | 0.377 | 1.30E-46 | 06_Cytotoxic_Tcells |
| PBX4 | -2.130255001 | 0.103 | 0.372 | 6.03E-45 | 06_Cytotoxic_Tcells |
| CD44 | -1.021327521 | 0.476 | 0.719 | 4.61E-44 | 06_Cytotoxic_Tcells |
| GZMM | 1.213971447 | 0.499 | 0.294 | 1.47E-43 | 06_Cytotoxic_Tcells |
| INPP4B | -2.278187236 | 0.079 | 0.318 | 2.98E-39 | 06_Cytotoxic_Tcells |
| RGS1 | -1.175802059 | 0.438 | 0.646 | 4.23E-39 | 06_Cytotoxic_Tcells |
| SYNE1 | 1.254963387 | 0.463 | 0.26 | 1.68E-38 | 06_Cytotoxic_Tcells |
| TC2N | -1.521547763 | 0.202 | 0.452 | 3.81E-38 | 06_Cytotoxic_Tcells |
| SNX9 | -2.100177206 | 0.121 | 0.357 | 1.97E-37 | 06_Cytotoxic_Tcells |
| CXCR6 | -2.145318669 | 0.051 | 0.252 | 1.72E-31 | 06_Cytotoxic_Tcells |
| RNF19A | -1.07410261 | 0.402 | 0.613 | 7.20E-31 | 06_Cytotoxic_Tcells |
| CDC14A | -2.024634471 | 0.113 | 0.323 | 5.64E-30 | 06_Cytotoxic_Tcells |
| RGCC | -1.046148132 | 0.341 | 0.556 | 1.47E-28 | 06_Cytotoxic_Tcells |
| BCL11B | -1.016129649 | 0.288 | 0.522 | 2.28E-27 | 06_Cytotoxic_Tcells |
| PFKFB3 | -1.421212477 | 0.159 | 0.369 | 4.18E-26 | 06_Cytotoxic_Tcells |
| OXNAD1 | -1.375280302 | 0.137 | 0.341 | 1.16E-25 | 06_Cytotoxic_Tcells |

| | | | | | |
|---|---|---|---|---|---|
| BCL2 | -1.265874841 | 0.216 | 0.421 | 1.96E-24 | 06_Cytotoxic_Tcells |
| ATXN1 | -1.032017138 | 0.312 | 0.525 | 3.33E-24 | 06_Cytotoxic_Tcells |
| C1QB | 2.631186494 | 0.824 | 0.327 | 1.76E-223 | 07_HLA_2_high |
| C1QA | 2.434906524 | 0.787 | 0.297 | 3.58E-203 | 07_HLA_2_high |
| C1QC | 2.467435893 | 0.817 | 0.346 | 4.58E-199 | 07_HLA_2_high |
| APOE | 2.277043712 | 0.859 | 0.478 | 1.14E-157 | 07_HLA_2_high |
| CST3 | 2.039593646 | 0.767 | 0.333 | 3.16E-155 | 07_HLA_2_high |
| HLA-DRA | 1.931099919 | 0.868 | 0.507 | 5.71E-150 | 07_HLA_2_high |
| APOC1 | 2.17541287 | 0.71 | 0.293 | 2.40E-146 | 07_HLA_2_high |
| AIF1 | 2.181458905 | 0.61 | 0.199 | 2.97E-144 | 07_HLA_2_high |
| NPC2 | 2.098998929 | 0.71 | 0.304 | 6.71E-143 | 07_HLA_2_high |
| APOC2 | 2.642531713 | 0.508 | 0.15 | 4.54E-133 | 07_HLA_2_high |
| TREM2 | 2.422597031 | 0.465 | 0.122 | 1.02E-131 | 07_HLA_2_high |
| CD68 | 2.137571603 | 0.511 | 0.15 | 3.83E-127 | 07_HLA_2_high |
| TYROBP | 1.622715004 | 0.797 | 0.392 | 1.03E-126 | 07_HLA_2_high |
| SPP1 | 2.402358643 | 0.849 | 0.556 | 2.99E-125 | 07_HLA_2_high |
| C3 | 2.414569532 | 0.556 | 0.191 | 3.13E-120 | 07_HLA_2_high |
| FCGR1A | 2.385465937 | 0.356 | 0.076 | 7.48E-119 | 07_HLA_2_high |
| FOLR2 | 2.470758282 | 0.349 | 0.074 | 3.50E-118 | 07_HLA_2_high |
| PLTP | 2.378154754 | 0.432 | 0.118 | 1.75E-112 | 07_HLA_2_high |
| FCER1G | 1.684645129 | 0.689 | 0.293 | 1.82E-111 | 07_HLA_2_high |
| TMEM176B | 2.661317302 | 0.324 | 0.068 | 2.23E-111 | 07_HLA_2_high |
| A2M | 1.991904003 | 0.54 | 0.187 | 2.07E-106 | 07_HLA_2_high |
| FCGRT | 2.136572514 | 0.438 | 0.129 | 8.17E-106 | 07_HLA_2_high |
| HLA-DPA1 | 1.654362593 | 0.814 | 0.504 | 8.44E-104 | 07_HLA_2_high |
| VSIG4 | 2.322074786 | 0.341 | 0.08 | 4.34E-102 | 07_HLA_2_high |
| MS4A6A | 2.084763749 | 0.443 | 0.132 | 1.55E-101 | 07_HLA_2_high |
| SERPINA1 | 2.309583339 | 0.324 | 0.075 | 9.42E-99 | 07_HLA_2_high |
| CD14 | 2.092639254 | 0.414 | 0.123 | 3.33E-93 | 07_HLA_2_high |
| TMIGD3 | 2.474101669 | 0.29 | 0.065 | 1.22E-90 | 07_HLA_2_high |
| LY86 | 2.578017226 | 0.275 | 0.062 | 3.74E-85 | 07_HLA_2_high |
| CTSB | 1.479211033 | 0.686 | 0.352 | 1.70E-81 | 07_HLA_2_high |
| HLA-DRB5 | 1.71899688 | 0.662 | 0.336 | 3.32E-81 | 07_HLA_2_high |
| HTRA1 | 2.359848939 | 0.302 | 0.079 | 5.25E-78 | 07_HLA_2_high |
| HLA-DPB1 | 1.483382158 | 0.743 | 0.446 | 2.57E-77 | 07_HLA_2_high |
| HLA-DRB1 | 1.34789889 | 0.811 | 0.538 | 3.10E-77 | 07_HLA_2_high |
| CXCL16 | 1.862960697 | 0.387 | 0.123 | 7.58E-76 | 07_HLA_2_high |
| GPR34 | 1.954136384 | 0.325 | 0.092 | 1.56E-73 | 07_HLA_2_high |
| CSF1R | 2.053728137 | 0.327 | 0.097 | 2.06E-69 | 07_HLA_2_high |
| RNASET2 | 1.438776657 | 0.648 | 0.37 | 4.68E-68 | 07_HLA_2_high |
| GSN | 1.632045492 | 0.419 | 0.152 | 1.83E-66 | 07_HLA_2_high |
| MARCKS | 1.792484146 | 0.394 | 0.14 | 3.92E-66 | 07_HLA_2_high |
| PSAP | 1.509996879 | 0.692 | 0.444 | 3.43E-65 | 07_HLA_2_high |
| SPI1 | 1.658555202 | 0.37 | 0.125 | 2.02E-64 | 07_HLA_2_high |
| LILRB4 | 2.07709361 | 0.281 | 0.079 | 5.57E-64 | 07_HLA_2_high |
| FCGR3A | 1.462197497 | 0.437 | 0.165 | 6.72E-64 | 07_HLA_2_high |

| | | | | | |
|---|---|---|---|---|---|
| IFI30 | 1.463185056 | 0.511 | 0.223 | 2.88E-63 | 07_HLA_2_high |
| HLA-DMA | 1.66207949 | 0.475 | 0.201 | 9.34E-63 | 07_HLA_2_high |
| HLA-DQA1 | 1.584742491 | 0.514 | 0.228 | 8.38E-62 | 07_HLA_2_high |
| S100A11 | 1.045265047 | 0.822 | 0.61 | 3.96E-59 | 07_HLA_2_high |
| CREG1 | 1.767689574 | 0.325 | 0.105 | 1.83E-58 | 07_HLA_2_high |
| MS4A7 | 1.270169466 | 0.417 | 0.167 | 7.39E-52 | 07_HLA_2_high |
| HLA-DQB1 | 1.241437107 | 0.581 | 0.306 | 4.65E-49 | 07_HLA_2_high |
| GRN | 1.586489815 | 0.362 | 0.143 | 3.78E-48 | 07_HLA_2_high |
| IFITM3 | 1.454273389 | 0.456 | 0.23 | 8.95E-40 | 07_HLA_2_high |
| CD63 | 1.005309801 | 0.584 | 0.361 | 7.82E-34 | 07_HLA_2_high |
| CTLA4 | 3.644078324 | 0.751 | 0.13 | 0 | 08_Tregs |
| IKZF2 | 4.431818638 | 0.684 | 0.069 | 0 | 08_Tregs |
| FOXP3 | 6.57852227 | 0.587 | 0.011 | 0 | 08_Tregs |
| RTKN2 | 5.103813226 | 0.426 | 0.023 | 0 | 08_Tregs |
| IL2RA | 4.902916353 | 0.43 | 0.034 | 0 | 08_Tregs |
| LAYN | 5.51161911 | 0.273 | 0.008 | 0 | 08_Tregs |
| CCR8 | 7.107983672 | 0.251 | 0.003 | 0 | 08_Tregs |
| BATF | 3.344124394 | 0.758 | 0.181 | 1.88E-287 | 08_Tregs |
| TBC1D4 | 3.223093018 | 0.623 | 0.117 | 4.53E-258 | 08_Tregs |
| AL136456.1 | 4.116074169 | 0.484 | 0.068 | 3.86E-251 | 08_Tregs |
| AC093865.1 | 4.266444447 | 0.292 | 0.02 | 6.79E-250 | 08_Tregs |
| LINC02694 | 4.229327968 | 0.415 | 0.053 | 5.52E-229 | 08_Tregs |
| STAM | 2.891157511 | 0.585 | 0.12 | 2.29E-215 | 08_Tregs |
| MAST4 | 3.736821743 | 0.422 | 0.061 | 1.53E-205 | 08_Tregs |
| LINC01943 | 3.701988078 | 0.341 | 0.043 | 2.17E-186 | 08_Tregs |
| TNFRSF18 | 3.137541444 | 0.487 | 0.092 | 2.94E-184 | 08_Tregs |
| TNFRSF4 | 3.330442389 | 0.495 | 0.097 | 2.64E-180 | 08_Tregs |
| ICA1 | 3.982435145 | 0.255 | 0.026 | 2.45E-162 | 08_Tregs |
| TIGIT | 2.633990159 | 0.551 | 0.138 | 2.09E-158 | 08_Tregs |
| CRADD | 4.067219115 | 0.386 | 0.069 | 4.55E-153 | 08_Tregs |
| CARD16 | 2.569807891 | 0.644 | 0.216 | 1.35E-146 | 08_Tregs |
| TNFRSF9 | 2.336688693 | 0.475 | 0.104 | 4.08E-146 | 08_Tregs |
| ICOS | 2.293432919 | 0.742 | 0.297 | 2.10E-142 | 08_Tregs |
| DUSP16 | 2.39375935 | 0.653 | 0.223 | 1.79E-131 | 08_Tregs |
| SLAMF1 | 2.584172211 | 0.462 | 0.113 | 8.32E-129 | 08_Tregs |
| UGP2 | 2.343818861 | 0.671 | 0.257 | 2.43E-127 | 08_Tregs |
| IL12RB2 | 2.65293475 | 0.406 | 0.096 | 2.08E-114 | 08_Tregs |
| CCL5 | -2.795092705 | 0.229 | 0.719 | 2.09E-110 | 08_Tregs |
| PHACTR2 | 2.288631478 | 0.581 | 0.216 | 1.19E-100 | 08_Tregs |
| ENTPD1 | 2.732680846 | 0.336 | 0.074 | 6.86E-100 | 08_Tregs |
| PHTF2 | 2.158667684 | 0.549 | 0.202 | 2.82E-93 | 08_Tregs |
| PELI1 | 1.888559836 | 0.585 | 0.239 | 2.10E-83 | 08_Tregs |
| GK | 2.434700213 | 0.374 | 0.104 | 6.64E-83 | 08_Tregs |
| TRAF3 | 2.13403762 | 0.48 | 0.169 | 5.11E-80 | 08_Tregs |
| NKG7 | -3.979953915 | 0.119 | 0.541 | 2.70E-79 | 08_Tregs |
| KAT2B | 2.062282128 | 0.556 | 0.224 | 1.95E-78 | 08_Tregs |

| | | | | | |
|---|---|---|---|---|---|
| PLCL1 | 2.553447094 | 0.318 | 0.08 | 4.65E-78 | 08_Tregs |
| ZNRF1 | 2.393638191 | 0.296 | 0.071 | 6.19E-76 | 08_Tregs |
| HPGD | 2.380885822 | 0.301 | 0.074 | 8.97E-76 | 08_Tregs |
| MIR4435-2HG | 1.804173841 | 0.491 | 0.183 | 4.85E-74 | 08_Tregs |
| DUSP4 | 1.387576487 | 0.699 | 0.347 | 5.87E-73 | 08_Tregs |
| ANXA1 | -2.308395912 | 0.215 | 0.617 | 5.19E-72 | 08_Tregs |
| ACP5 | 2.358699092 | 0.3 | 0.076 | 6.07E-72 | 08_Tregs |
| BIRC3 | 1.747039884 | 0.61 | 0.279 | 4.61E-69 | 08_Tregs |
| BTG3 | 1.665495882 | 0.525 | 0.222 | 2.78E-66 | 08_Tregs |
| MAP3K5 | 1.780402504 | 0.522 | 0.226 | 4.05E-62 | 08_Tregs |
| SNX9 | 1.416852075 | 0.653 | 0.321 | 1.33E-61 | 08_Tregs |
| GZMA | -2.045269033 | 0.184 | 0.564 | 1.74E-61 | 08_Tregs |
| GBP5 | 1.648850907 | 0.527 | 0.229 | 4.63E-61 | 08_Tregs |
| CD28 | 1.569081793 | 0.455 | 0.181 | 5.37E-56 | 08_Tregs |
| CTSW | -2.9750172 | 0.126 | 0.474 | 8.07E-56 | 08_Tregs |
| PMAIP1 | 1.731898587 | 0.455 | 0.189 | 3.50E-55 | 08_Tregs |
| STAT3 | 1.334700123 | 0.738 | 0.45 | 3.74E-53 | 08_Tregs |
| PHLDA1 | 1.676841222 | 0.444 | 0.183 | 1.01E-52 | 08_Tregs |
| CD4 | 1.611944151 | 0.383 | 0.14 | 7.09E-52 | 08_Tregs |
| SAMSN1 | 1.344492848 | 0.841 | 0.621 | 9.10E-51 | 08_Tregs |
| CTSC | 1.515332503 | 0.63 | 0.364 | 1.84E-49 | 08_Tregs |
| ARID5B | 1.150822809 | 0.735 | 0.455 | 2.13E-49 | 08_Tregs |
| NAMPT | 1.154468208 | 0.626 | 0.326 | 1.17E-48 | 08_Tregs |
| LTB | 1.614874422 | 0.592 | 0.32 | 6.10E-48 | 08_Tregs |
| TTN | 1.770996521 | 0.31 | 0.101 | 7.93E-48 | 08_Tregs |
| TRAF1 | 1.736617378 | 0.374 | 0.14 | 8.88E-48 | 08_Tregs |
| AOAH | -2.533453961 | 0.146 | 0.474 | 1.31E-47 | 08_Tregs |
| CD8A | -2.784816999 | 0.09 | 0.415 | 4.65E-47 | 08_Tregs |
| CYTOR | 1.205102549 | 0.625 | 0.34 | 1.45E-46 | 08_Tregs |
| TOX | 1.624809997 | 0.587 | 0.308 | 3.01E-46 | 08_Tregs |
| EPSTI1 | 1.696064332 | 0.401 | 0.165 | 7.89E-46 | 08_Tregs |
| SIRPG | 1.862002124 | 0.303 | 0.103 | 9.06E-46 | 08_Tregs |
| USP15 | 1.318334423 | 0.711 | 0.464 | 1.01E-44 | 08_Tregs |
| CASK | 1.553832583 | 0.532 | 0.269 | 2.41E-44 | 08_Tregs |
| GZMK | -2.070330696 | 0.175 | 0.485 | 1.12E-43 | 08_Tregs |
| THADA | 2.284995159 | 0.352 | 0.14 | 6.79E-43 | 08_Tregs |
| GPHN | 1.844621523 | 0.33 | 0.124 | 1.26E-41 | 08_Tregs |
| PRDM1 | 1.272698022 | 0.648 | 0.402 | 2.74E-41 | 08_Tregs |
| SGMS1 | 1.953830498 | 0.408 | 0.181 | 3.08E-41 | 08_Tregs |
| KLRK1 | -3.736474486 | 0.049 | 0.326 | 3.72E-39 | 08_Tregs |
| FRMD4B | 1.448059944 | 0.412 | 0.184 | 1.06E-37 | 08_Tregs |
| IL6ST | 1.311252475 | 0.5 | 0.258 | 6.66E-37 | 08_Tregs |
| CD27 | 1.269339466 | 0.464 | 0.229 | 1.24E-36 | 08_Tregs |
| RHBDD2 | 1.414806985 | 0.435 | 0.208 | 2.11E-36 | 08_Tregs |
| TNIK | 1.296573305 | 0.522 | 0.278 | 3.63E-36 | 08_Tregs |
| FOXO1 | 1.273542498 | 0.634 | 0.38 | 9.44E-36 | 08_Tregs |

| | | | | | |
|---|---|---|---|---|---|
| TC2N | -2.117168782 | 0.171 | 0.448 | 1.20E-35 | 08_Tregs |
| CXCR6 | 1.209984843 | 0.453 | 0.224 | 3.69E-35 | 08_Tregs |
| MALT1 | 1.309918301 | 0.538 | 0.291 | 6.58E-35 | 08_Tregs |
| ZEB2 | -2.162846263 | 0.184 | 0.449 | 1.30E-34 | 08_Tregs |
| GBP2 | 1.285064416 | 0.444 | 0.221 | 5.71E-34 | 08_Tregs |
| TSPAN5 | 1.460460715 | 0.426 | 0.209 | 1.15E-32 | 08_Tregs |
| TUBA4A | -1.556477965 | 0.356 | 0.586 | 1.16E-32 | 08_Tregs |
| ZNF292 | 1.324860009 | 0.487 | 0.261 | 2.15E-32 | 08_Tregs |
| SKAP1 | 1.123855718 | 0.7 | 0.495 | 9.25E-32 | 08_Tregs |
| GLCCI1 | 1.142002836 | 0.542 | 0.305 | 9.96E-32 | 08_Tregs |
| TRPS1 | 1.410221223 | 0.424 | 0.213 | 2.14E-30 | 08_Tregs |
| TANK | 1.318225063 | 0.523 | 0.302 | 6.86E-30 | 08_Tregs |
| VAV3 | 1.542947923 | 0.477 | 0.264 | 9.21E-30 | 08_Tregs |
| LDLRAD4 | 1.030742222 | 0.697 | 0.437 | 3.00E-28 | 08_Tregs |
| GZMH | -2.622925292 | 0.085 | 0.318 | 6.67E-28 | 08_Tregs |
| PBXIP1 | 1.111855897 | 0.504 | 0.288 | 9.53E-28 | 08_Tregs |
| ZBTB38 | 1.028412849 | 0.478 | 0.258 | 2.65E-27 | 08_Tregs |
| LITAF | -1.32751089 | 0.352 | 0.569 | 3.52E-27 | 08_Tregs |
| AUTS2 | -2.630020424 | 0.088 | 0.318 | 3.93E-27 | 08_Tregs |
| CORO1B | 1.158069692 | 0.509 | 0.301 | 3.97E-27 | 08_Tregs |
| BHLHE40 | -2.45086183 | 0.106 | 0.334 | 6.94E-27 | 08_Tregs |
| CD8B | -2.872224451 | 0.079 | 0.3 | 2.36E-26 | 08_Tregs |
| ADGRE5 | -1.253522506 | 0.379 | 0.586 | 9.75E-26 | 08_Tregs |
| CD55 | -1.616881824 | 0.195 | 0.43 | 1.34E-25 | 08_Tregs |
| NSD3 | 1.089569943 | 0.554 | 0.349 | 1.35E-24 | 08_Tregs |
| IPCEF1 | 1.098552267 | 0.48 | 0.274 | 1.02E-23 | 08_Tregs |
| NOP58 | 1.089127988 | 0.525 | 0.325 | 4.56E-23 | 08_Tregs |
| MYADM | -1.543314022 | 0.186 | 0.388 | 2.77E-20 | 08_Tregs |
| ALOX5AP | -1.186562335 | 0.294 | 0.496 | 5.96E-19 | 08_Tregs |
| P2RY8 | -1.308559638 | 0.2 | 0.401 | 1.45E-17 | 08_Tregs |
| XCL1 | 4.300694518 | 0.552 | 0.061 | 0 | 09_Activated_NKcells |
| KRT86 | 5.604929142 | 0.442 | 0.02 | 0 | 09_Activated_NKcells |
| KRT81 | 6.720866103 | 0.234 | 0.004 | 0 | 09_Activated_NKcells |
| KLRD1 | 2.577342445 | 0.774 | 0.177 | 3.16E-249 | 09_Activated_NKcells |
| XCL2 | 3.48095975 | 0.536 | 0.088 | 1.95E-226 | 09_Activated_NKcells |
| TRDC | 4.082885279 | 0.246 | 0.016 | 1.14E-201 | 09_Activated_NKcells |
| KIR2DL4 | 4.282075029 | 0.22 | 0.013 | 2.71E-196 | 09_Activated_NKcells |
| KLRC2 | 3.353193068 | 0.341 | 0.043 | 1.65E-170 | 09_Activated_NKcells |
| IGFBP2 | 3.661531053 | 0.27 | 0.027 | 1.02E-163 | 09_Activated_NKcells |
| KLRC1 | 3.365028681 | 0.343 | 0.047 | 5.13E-160 | 09_Activated_NKcells |
| SH2D1B | 3.429278655 | 0.226 | 0.022 | 7.33E-138 | 09_Activated_NKcells |
| CD7 | 1.830380786 | 0.881 | 0.547 | 1.57E-125 | 09_Activated_NKcells |
| TYROBP | 1.716987198 | 0.831 | 0.396 | 4.20E-121 | 09_Activated_NKcells |
| TMIGD2 | 3.245733071 | 0.284 | 0.044 | 1.18E-113 | 09_Activated_NKcells |
| MCTP2 | 2.190434579 | 0.536 | 0.155 | 2.32E-113 | 09_Activated_NKcells |
| AREG | 2.211184924 | 0.738 | 0.326 | 1.26E-111 | 09_Activated_NKcells |

| | | | | | |
|---|---|---|---|---|---|
| TXK | 2.28072124 | 0.456 | 0.116 | 2.40E-108 | 09_Activated_NKcells |
| CLIC3 | 2.58552934 | 0.435 | 0.118 | 2.26E-97 | 09_Activated_NKcells |
| NKG7 | 1.33388876 | 0.877 | 0.5 | 2.05E-93 | 09_Activated_NKcells |
| CTSW | 1.684114984 | 0.768 | 0.44 | 9.72E-82 | 09_Activated_NKcells |
| KLRB1 | 1.574150573 | 0.657 | 0.28 | 7.28E-81 | 09_Activated_NKcells |
| CD3D | -2.336453189 | 0.141 | 0.582 | 5.83E-67 | 09_Activated_NKcells |
| IL2RB | 2.009215543 | 0.532 | 0.233 | 8.25E-67 | 09_Activated_NKcells |
| MATK | 2.18842598 | 0.409 | 0.137 | 2.79E-66 | 09_Activated_NKcells |
| LAT2 | 2.551243117 | 0.28 | 0.073 | 2.61E-60 | 09_Activated_NKcells |
| PLCG2 | 2.345778321 | 0.288 | 0.082 | 3.49E-54 | 09_Activated_NKcells |
| IL7R | -2.182819836 | 0.194 | 0.578 | 4.60E-54 | 09_Activated_NKcells |
| HOPX | 1.849363005 | 0.379 | 0.152 | 1.43E-43 | 09_Activated_NKcells |
| TNFRSF18 | 1.893676749 | 0.306 | 0.104 | 3.02E-43 | 09_Activated_NKcells |
| BCL11B | -1.794642378 | 0.165 | 0.521 | 2.22E-41 | 09_Activated_NKcells |
| GRASP | 2.127249431 | 0.381 | 0.163 | 3.11E-40 | 09_Activated_NKcells |
| ITGAE | 1.910269986 | 0.375 | 0.163 | 2.71E-38 | 09_Activated_NKcells |
| CD3G | -2.667331723 | 0.077 | 0.383 | 1.13E-37 | 09_Activated_NKcells |
| CAMK4 | -2.180016319 | 0.117 | 0.426 | 4.18E-36 | 09_Activated_NKcells |
| THEMIS | -2.618281929 | 0.065 | 0.364 | 1.76E-35 | 09_Activated_NKcells |
| METRNL | 1.61837113 | 0.544 | 0.326 | 1.35E-34 | 09_Activated_NKcells |
| ANXA1 | -1.562143107 | 0.335 | 0.608 | 1.91E-32 | 09_Activated_NKcells |
| CD8A | -2.343248584 | 0.135 | 0.411 | 3.41E-32 | 09_Activated_NKcells |
| ARID5B | -1.648144685 | 0.188 | 0.485 | 1.39E-31 | 09_Activated_NKcells |
| MAP3K8 | 1.238318914 | 0.558 | 0.338 | 2.64E-31 | 09_Activated_NKcells |
| CD52 | -1.125006775 | 0.359 | 0.622 | 3.20E-29 | 09_Activated_NKcells |
| KLRK1 | 1.087300128 | 0.524 | 0.301 | 4.75E-29 | 09_Activated_NKcells |
| PBX4 | -2.040568449 | 0.091 | 0.365 | 4.95E-29 | 09_Activated_NKcells |
| CD6 | -2.138835871 | 0.095 | 0.367 | 5.66E-29 | 09_Activated_NKcells |
| CD8B | -3.036138086 | 0.056 | 0.3 | 4.04E-27 | 09_Activated_NKcells |
| FCER1G | 1.007138619 | 0.53 | 0.307 | 1.26E-25 | 09_Activated_NKcells |
| ICOS | -2.440032885 | 0.091 | 0.333 | 4.14E-25 | 09_Activated_NKcells |
| PDE3B | -1.42603356 | 0.274 | 0.529 | 2.10E-23 | 09_Activated_NKcells |
| INPP4B | -2.346353042 | 0.079 | 0.311 | 2.96E-23 | 09_Activated_NKcells |
| TOB1 | -2.461212907 | 0.065 | 0.291 | 6.24E-23 | 09_Activated_NKcells |
| GPR183 | -1.721014038 | 0.127 | 0.368 | 3.34E-22 | 09_Activated_NKcells |
| CD5 | -2.865465149 | 0.056 | 0.268 | 6.07E-22 | 09_Activated_NKcells |
| KLF2 | -1.578886237 | 0.2 | 0.435 | 2.50E-21 | 09_Activated_NKcells |
| SPOCK2 | -1.007426614 | 0.268 | 0.523 | 3.17E-20 | 09_Activated_NKcells |
| ZNF831 | -1.749128116 | 0.111 | 0.334 | 6.15E-20 | 09_Activated_NKcells |
| SERPINB9 | -1.350750376 | 0.155 | 0.387 | 3.18E-19 | 09_Activated_NKcells |
| CD2 | -1.002531246 | 0.321 | 0.551 | 3.25E-18 | 09_Activated_NKcells |
| MGAT4A | -1.569217861 | 0.133 | 0.35 | 3.89E-18 | 09_Activated_NKcells |
| SYNE2 | -1.035569075 | 0.284 | 0.513 | 6.32E-17 | 09_Activated_NKcells |
| ELMO1 | -1.457075503 | 0.194 | 0.404 | 7.49E-17 | 09_Activated_NKcells |
| RNASET2 | -1.244031323 | 0.183 | 0.397 | 3.44E-16 | 09_Activated_NKcells |
| FOXO1 | -1.309449608 | 0.2 | 0.403 | 1.06E-14 | 09_Activated_NKcells |

| | | | | | |
|---|---|---|---|---|---|
| ZNF683 | 2.670318608 | 0.316 | 0.052 | 4.29E-102 | 10_Effector/Memory_Tcells |
| CD8B | 1.532280618 | 0.744 | 0.269 | 9.78E-95 | 10_Effector/Memory_Tcells |
| CD8A | 1.348499766 | 0.848 | 0.379 | 2.29E-84 | 10_Effector/Memory_Tcells |
| LINC02446 | 2.412230156 | 0.309 | 0.067 | 1.44E-70 | 10_Effector/Memory_Tcells |
| GZMB | 1.478368014 | 0.599 | 0.22 | 1.14E-68 | 10_Effector/Memory_Tcells |
| LINC01871 | 1.967722022 | 0.478 | 0.157 | 1.69E-67 | 10_Effector/Memory_Tcells |
| GNLY | 1.56197653 | 0.51 | 0.189 | 3.02E-53 | 10_Effector/Memory_Tcells |
| TOB1 | 1.458478964 | 0.587 | 0.267 | 2.30E-48 | 10_Effector/Memory_Tcells |
| HOPX | 1.525905661 | 0.406 | 0.152 | 1.01E-39 | 10_Effector/Memory_Tcells |
| IFIT3 | 4.589471443 | 0.452 | 0.034 | 6.56E-200 | 11_ISG_TCells |
| RSAD2 | 4.376793052 | 0.47 | 0.038 | 6.86E-199 | 11_ISG_TCells |
| IFIT1 | 4.628725547 | 0.379 | 0.025 | 1.11E-186 | 11_ISG_TCells |
| CMPK2 | 4.245339427 | 0.388 | 0.032 | 4.35E-156 | 11_ISG_TCells |
| MX1 | 3.434528755 | 0.799 | 0.181 | 6.03E-145 | 11_ISG_TCells |
| IFIT2 | 4.522658617 | 0.37 | 0.034 | 3.83E-135 | 11_ISG_TCells |
| OAS3 | 3.85338108 | 0.429 | 0.05 | 3.20E-127 | 11_ISG_TCells |
| LAMP3 | 4.656813088 | 0.251 | 0.016 | 1.54E-126 | 11_ISG_TCells |
| IFI44L | 3.342844203 | 0.667 | 0.133 | 9.14E-124 | 11_ISG_TCells |
| IFI6 | 3.361549346 | 0.808 | 0.232 | 6.31E-119 | 11_ISG_TCells |
| OAS1 | 3.480229647 | 0.447 | 0.063 | 2.23E-108 | 11_ISG_TCells |
| HERC5 | 3.184453447 | 0.47 | 0.083 | 3.49E-88 | 11_ISG_TCells |
| STAT1 | 2.652977224 | 0.776 | 0.263 | 3.42E-85 | 11_ISG_TCells |
| ISG15 | 3.324344225 | 0.721 | 0.236 | 7.32E-85 | 11_ISG_TCells |
| USP18 | 3.836521739 | 0.26 | 0.026 | 1.25E-83 | 11_ISG_TCells |
| MX2 | 2.947684448 | 0.575 | 0.136 | 3.95E-82 | 11_ISG_TCells |
| OASL | 3.012933059 | 0.553 | 0.13 | 1.02E-78 | 11_ISG_TCells |
| EIF2AK2 | 2.735527429 | 0.594 | 0.153 | 6.45E-78 | 11_ISG_TCells |
| IFI44 | 3.174197083 | 0.498 | 0.106 | 2.72E-77 | 11_ISG_TCells |
| GBP1 | 3.165488546 | 0.502 | 0.111 | 2.34E-76 | 11_ISG_TCells |
| DDX60L | 3.042671969 | 0.493 | 0.113 | 1.62E-68 | 11_ISG_TCells |
| XAF1 | 2.715271847 | 0.525 | 0.13 | 1.60E-67 | 11_ISG_TCells |
| HELZ2 | 3.084826935 | 0.324 | 0.05 | 6.94E-67 | 11_ISG_TCells |
| HERC6 | 3.066052309 | 0.315 | 0.051 | 9.24E-61 | 11_ISG_TCells |
| EPSTI1 | 2.539235772 | 0.575 | 0.169 | 2.44E-60 | 11_ISG_TCells |
| SAMD9L | 2.933026803 | 0.457 | 0.111 | 3.97E-59 | 11_ISG_TCells |
| IFIH1 | 3.084646924 | 0.347 | 0.063 | 1.17E-58 | 11_ISG_TCells |
| TRIM22 | 2.15350045 | 0.763 | 0.325 | 5.66E-58 | 11_ISG_TCells |
| OAS2 | 2.829823052 | 0.397 | 0.084 | 7.79E-58 | 11_ISG_TCells |
| IRF7 | 2.428387908 | 0.58 | 0.178 | 1.38E-57 | 11_ISG_TCells |
| IFI35 | 2.790380987 | 0.443 | 0.111 | 5.64E-54 | 11_ISG_TCells |
| PARP9 | 2.620365877 | 0.411 | 0.096 | 1.26E-52 | 11_ISG_TCells |
| PNPT1 | 3.015180925 | 0.301 | 0.054 | 4.27E-50 | 11_ISG_TCells |
| PLSCR1 | 2.296488652 | 0.562 | 0.183 | 2.10E-49 | 11_ISG_TCells |
| STAT2 | 2.780501909 | 0.374 | 0.085 | 4.12E-49 | 11_ISG_TCells |
| LY6E | 2.031516421 | 0.753 | 0.47 | 2.07E-42 | 11_ISG_TCells |
| LAG3 | 2.314062786 | 0.53 | 0.188 | 1.71E-41 | 11_ISG_TCells |

| | | | | | |
|---|---|---|---|---|---|
| RNF213 | 1.837221854 | 0.822 | 0.505 | 5.51E-41 | 11_ISG_TCells |
| ISG20 | 1.469915458 | 0.872 | 0.669 | 3.86E-40 | 11_ISG_TCells |
| PARP12 | 2.446677391 | 0.361 | 0.091 | 1.88E-39 | 11_ISG_TCells |
| DDX60 | 2.775258897 | 0.32 | 0.077 | 9.59E-37 | 11_ISG_TCells |
| DDX58 | 2.819459254 | 0.315 | 0.078 | 2.18E-35 | 11_ISG_TCells |
| HSH2D | 2.23480096 | 0.374 | 0.106 | 6.27E-34 | 11_ISG_TCells |
| LGALS9 | 2.20935016 | 0.397 | 0.125 | 1.05E-32 | 11_ISG_TCells |
| BST2 | 1.79651293 | 0.621 | 0.297 | 1.28E-32 | 11_ISG_TCells |
| TNFSF10 | 2.488185566 | 0.356 | 0.103 | 8.37E-32 | 11_ISG_TCells |
| PARP14 | 2.219679542 | 0.498 | 0.199 | 1.20E-31 | 11_ISG_TCells |
| ZBP1 | 2.313420339 | 0.356 | 0.106 | 4.17E-30 | 11_ISG_TCells |
| GBP4 | 2.568143811 | 0.365 | 0.115 | 3.64E-29 | 11_ISG_TCells |
| LAP3 | 1.903174058 | 0.461 | 0.178 | 7.43E-27 | 11_ISG_TCells |
| SAMD9 | 2.209551991 | 0.365 | 0.12 | 8.21E-27 | 11_ISG_TCells |
| SP110 | 1.69995739 | 0.562 | 0.271 | 1.15E-26 | 11_ISG_TCells |
| XRN1 | 1.955266866 | 0.571 | 0.285 | 7.79E-26 | 11_ISG_TCells |
| IFI16 | 1.276962639 | 0.712 | 0.427 | 2.99E-24 | 11_ISG_TCells |
| SHFL | 1.870258567 | 0.457 | 0.186 | 1.01E-23 | 11_ISG_TCells |
| UBE2L6 | 1.839635115 | 0.502 | 0.233 | 1.95E-23 | 11_ISG_TCells |
| GBP5 | 1.70286826 | 0.507 | 0.24 | 2.89E-21 | 11_ISG_TCells |
| C5orf56 | 1.578985625 | 0.479 | 0.225 | 9.70E-19 | 11_ISG_TCells |
| SP100 | 1.290712869 | 0.717 | 0.515 | 2.23E-18 | 11_ISG_TCells |
| PHF11 | 1.662412844 | 0.402 | 0.171 | 4.53E-18 | 11_ISG_TCells |
| ADAR | 1.501937551 | 0.539 | 0.287 | 7.59E-18 | 11_ISG_TCells |
| NT5C3A | 1.567664095 | 0.406 | 0.181 | 2.42E-16 | 11_ISG_TCells |
| DRAP1 | 1.248946853 | 0.603 | 0.37 | 4.42E-16 | 11_ISG_TCells |
| PSMB9 | 1.224403965 | 0.639 | 0.407 | 1.74E-15 | 11_ISG_TCells |
| APOL6 | 1.746405915 | 0.425 | 0.203 | 2.71E-15 | 11_ISG_TCells |
| PLAAT4 | 1.217352266 | 0.68 | 0.462 | 4.18E-15 | 11_ISG_TCells |
| RBCK1 | 1.626441585 | 0.388 | 0.175 | 4.52E-15 | 11_ISG_TCells |
| HELB | 1.770893531 | 0.37 | 0.164 | 1.95E-14 | 11_ISG_TCells |
| PPM1K | 1.82711291 | 0.384 | 0.178 | 2.34E-14 | 11_ISG_TCells |
| NUB1 | 1.702351527 | 0.361 | 0.157 | 2.79E-14 | 11_ISG_TCells |
| UTRN | 1.247994561 | 0.689 | 0.451 | 5.88E-14 | 11_ISG_TCells |
| ERAP2 | 1.610631297 | 0.384 | 0.175 | 2.27E-13 | 11_ISG_TCells |
| CHST12 | 1.374965387 | 0.443 | 0.225 | 1.78E-12 | 11_ISG_TCells |
| SLFN5 | 1.352943118 | 0.47 | 0.259 | 1.24E-11 | 11_ISG_TCells |
| TAP1 | 1.207627021 | 0.507 | 0.293 | 2.62E-11 | 11_ISG_TCells |
| MCTP2 | 3.383463912 | 0.594 | 0.165 | 9.03E-77 | 12_Resting_NKcells |
| TMSB4X | -1.756294732 | 0.756 | 0.97 | 1.15E-68 | 12_Resting_NKcells |
| RPS27 | -2.009017033 | 0.512 | 0.921 | 5.64E-64 | 12_Resting_NKcells |
| RPL28 | -1.711048321 | 0.641 | 0.944 | 6.70E-61 | 12_Resting_NKcells |
| RPS15A | -1.839093032 | 0.539 | 0.925 | 2.36E-60 | 12_Resting_NKcells |
| EEF1A1 | -1.634327002 | 0.71 | 0.96 | 2.56E-59 | 12_Resting_NKcells |
| RPS14 | -1.840815194 | 0.512 | 0.907 | 5.95E-59 | 12_Resting_NKcells |
| RPS12 | -1.798563622 | 0.599 | 0.937 | 1.80E-58 | 12_Resting_NKcells |

| | | | | | |
|---|---|---|---|---|---|
| RPL10 | -1.60195573 | 0.668 | 0.952 | 3.77E-58 | 12_Resting_NKcells |
| RPS3 | -1.753039549 | 0.53 | 0.913 | 1.74E-56 | 12_Resting_NKcells |
| RPS18 | -1.84644687 | 0.447 | 0.905 | 1.07E-55 | 12_Resting_NKcells |
| RPL30 | -1.696396852 | 0.544 | 0.923 | 1.48E-55 | 12_Resting_NKcells |
| RPL13 | -1.53201719 | 0.682 | 0.95 | 3.29E-54 | 12_Resting_NKcells |
| RPLP1 | -1.477741731 | 0.724 | 0.96 | 8.76E-54 | 12_Resting_NKcells |
| RPL32 | -1.719922525 | 0.507 | 0.913 | 9.95E-54 | 12_Resting_NKcells |
| RPL14 | -1.786067813 | 0.447 | 0.887 | 1.09E-53 | 12_Resting_NKcells |
| RPL19 | -1.632974878 | 0.548 | 0.919 | 4.53E-53 | 12_Resting_NKcells |
| RPL11 | -1.633218827 | 0.521 | 0.918 | 5.04E-53 | 12_Resting_NKcells |
| RPS15 | -1.589519942 | 0.507 | 0.912 | 6.72E-53 | 12_Resting_NKcells |
| RPS19 | -1.593930116 | 0.567 | 0.929 | 7.00E-53 | 12_Resting_NKcells |
| HLA-B | -1.356083039 | 0.728 | 0.963 | 7.94E-53 | 12_Resting_NKcells |
| RPL18A | -1.748603941 | 0.47 | 0.896 | 8.16E-53 | 12_Resting_NKcells |
| FAU | -1.66157992 | 0.47 | 0.896 | 9.62E-53 | 12_Resting_NKcells |
| HIPK2 | 3.015829165 | 0.484 | 0.138 | 1.32E-52 | 12_Resting_NKcells |
| HLA-A | -1.457550274 | 0.677 | 0.948 | 1.65E-52 | 12_Resting_NKcells |
| RPL26 | -1.664128973 | 0.502 | 0.903 | 5.95E-52 | 12_Resting_NKcells |
| RPS27A | -1.597667642 | 0.544 | 0.925 | 2.00E-51 | 12_Resting_NKcells |
| RPS25 | -1.737411912 | 0.479 | 0.884 | 2.50E-51 | 12_Resting_NKcells |
| RPS28 | -1.718829215 | 0.512 | 0.896 | 6.37E-51 | 12_Resting_NKcells |
| RPL18 | -1.635496079 | 0.424 | 0.894 | 1.20E-49 | 12_Resting_NKcells |
| RPL12 | -1.611804405 | 0.479 | 0.91 | 4.83E-49 | 12_Resting_NKcells |
| RPS4X | -1.70488578 | 0.452 | 0.889 | 5.59E-49 | 12_Resting_NKcells |
| RPL7A | -1.632591535 | 0.447 | 0.895 | 1.47E-48 | 12_Resting_NKcells |
| RPS7 | -1.561372476 | 0.525 | 0.9 | 3.06E-48 | 12_Resting_NKcells |
| RPL23A | -1.833881245 | 0.369 | 0.847 | 4.36E-48 | 12_Resting_NKcells |
| RPS8 | -1.492574996 | 0.594 | 0.927 | 4.67E-48 | 12_Resting_NKcells |
| RPL34 | -1.552366953 | 0.479 | 0.906 | 2.30E-47 | 12_Resting_NKcells |
| RPS16 | -1.767623762 | 0.378 | 0.857 | 5.91E-47 | 12_Resting_NKcells |
| RPS2 | -1.687539955 | 0.47 | 0.874 | 1.65E-46 | 12_Resting_NKcells |
| RPL17 | -1.716078347 | 0.438 | 0.859 | 1.67E-46 | 12_Resting_NKcells |
| RPS23 | -1.455930759 | 0.53 | 0.914 | 4.07E-46 | 12_Resting_NKcells |
| IL32 | -2.519797433 | 0.253 | 0.762 | 5.81E-46 | 12_Resting_NKcells |
| RPL6 | -1.59582779 | 0.484 | 0.881 | 8.10E-46 | 12_Resting_NKcells |
| HLA-C | -1.413867453 | 0.618 | 0.924 | 9.24E-46 | 12_Resting_NKcells |
| RPL29 | -1.559869613 | 0.456 | 0.884 | 5.52E-45 | 12_Resting_NKcells |
| RPL39 | -1.858569943 | 0.387 | 0.858 | 5.93E-45 | 12_Resting_NKcells |
| RPL37 | -1.685581127 | 0.429 | 0.87 | 6.98E-45 | 12_Resting_NKcells |
| RPL10A | -1.880206304 | 0.318 | 0.809 | 8.21E-45 | 12_Resting_NKcells |
| CD3E | -2.880412307 | 0.143 | 0.68 | 1.04E-44 | 12_Resting_NKcells |
| TPT1 | -1.324778672 | 0.696 | 0.937 | 1.30E-44 | 12_Resting_NKcells |
| RPS24 | -1.457780264 | 0.535 | 0.9 | 2.30E-44 | 12_Resting_NKcells |
| RPS6 | -1.531775279 | 0.507 | 0.888 | 2.44E-44 | 12_Resting_NKcells |
| TMSB10 | -1.398034978 | 0.627 | 0.941 | 2.96E-44 | 12_Resting_NKcells |
| RPLP0 | -1.662129382 | 0.465 | 0.86 | 6.40E-44 | 12_Resting_NKcells |

| | | | | | |
|---|---|---|---|---|---|
| RPL3 | -1.516371569 | 0.456 | 0.881 | 6.78E-44 | 12_Resting_NKcells |
| RPS3A | -1.535493362 | 0.493 | 0.877 | 1.29E-43 | 12_Resting_NKcells |
| RAP1GAP2 | 3.790524538 | 0.253 | 0.045 | 1.66E-43 | 12_Resting_NKcells |
| RPS13 | -1.472576901 | 0.535 | 0.894 | 1.71E-43 | 12_Resting_NKcells |
| PLCG2 | 3.1719472 | 0.355 | 0.086 | 3.15E-43 | 12_Resting_NKcells |
| RPL8 | -1.513045857 | 0.433 | 0.88 | 7.52E-43 | 12_Resting_NKcells |
| RPL15 | -1.547091565 | 0.396 | 0.859 | 9.78E-43 | 12_Resting_NKcells |
| RPL35A | -1.5402647 | 0.442 | 0.87 | 2.97E-42 | 12_Resting_NKcells |
| RPS26 | -1.579689104 | 0.461 | 0.883 | 6.98E-42 | 12_Resting_NKcells |
| RPL36 | -1.61330618 | 0.415 | 0.848 | 9.93E-42 | 12_Resting_NKcells |
| RPS9 | -1.49614189 | 0.438 | 0.868 | 6.61E-41 | 12_Resting_NKcells |
| RPSA | -1.711618689 | 0.364 | 0.818 | 3.40E-40 | 12_Resting_NKcells |
| ATP8B4 | 3.361455853 | 0.318 | 0.074 | 4.25E-40 | 12_Resting_NKcells |
| RPL35 | -1.650269167 | 0.373 | 0.813 | 3.52E-39 | 12_Resting_NKcells |
| NACA | -1.578310585 | 0.355 | 0.816 | 4.15E-39 | 12_Resting_NKcells |
| LINC00299 | 3.312997493 | 0.318 | 0.076 | 4.39E-39 | 12_Resting_NKcells |
| RPL21 | -1.545753881 | 0.419 | 0.849 | 1.34E-38 | 12_Resting_NKcells |
| RPLP2 | -1.381061691 | 0.502 | 0.883 | 9.60E-38 | 12_Resting_NKcells |
| UBC | -1.213805856 | 0.548 | 0.914 | 1.17E-37 | 12_Resting_NKcells |
| C1orf21 | 3.102873815 | 0.346 | 0.089 | 1.26E-37 | 12_Resting_NKcells |
| RPS5 | -1.519581281 | 0.406 | 0.828 | 6.09E-36 | 12_Resting_NKcells |
| AOAH | 2.211674535 | 0.733 | 0.451 | 6.25E-36 | 12_Resting_NKcells |
| SH3BGRL3 | -1.608978059 | 0.406 | 0.807 | 8.72E-36 | 12_Resting_NKcells |
| RPS21 | -1.474734027 | 0.41 | 0.844 | 2.36E-35 | 12_Resting_NKcells |
| RPL13A | -1.433796715 | 0.433 | 0.838 | 2.65E-35 | 12_Resting_NKcells |
| RPL9 | -1.50459896 | 0.369 | 0.806 | 6.19E-35 | 12_Resting_NKcells |
| CALM1 | -1.502074199 | 0.419 | 0.821 | 8.10E-35 | 12_Resting_NKcells |
| RACK1 | -1.487765727 | 0.429 | 0.828 | 1.08E-34 | 12_Resting_NKcells |
| EIF1 | -1.136344498 | 0.562 | 0.913 | 1.50E-34 | 12_Resting_NKcells |
| JUNB | -1.552424606 | 0.521 | 0.865 | 1.60E-34 | 12_Resting_NKcells |
| CD3D | -3.153456032 | 0.092 | 0.571 | 2.08E-34 | 12_Resting_NKcells |
| BTG1 | -1.247388613 | 0.636 | 0.92 | 4.02E-34 | 12_Resting_NKcells |
| UBA52 | -1.449291179 | 0.396 | 0.812 | 4.31E-34 | 12_Resting_NKcells |
| CFL1 | -1.498562006 | 0.415 | 0.821 | 7.35E-34 | 12_Resting_NKcells |
| RPL36A | -1.798826142 | 0.272 | 0.733 | 8.88E-34 | 12_Resting_NKcells |
| RPL37A | -1.612129965 | 0.327 | 0.786 | 9.79E-34 | 12_Resting_NKcells |
| CD247 | 2.124651408 | 0.756 | 0.553 | 9.94E-34 | 12_Resting_NKcells |
| RPL5 | -1.45005411 | 0.41 | 0.828 | 1.92E-33 | 12_Resting_NKcells |
| RPS29 | -1.855358755 | 0.267 | 0.724 | 5.40E-33 | 12_Resting_NKcells |
| GAPDH | -1.441475772 | 0.433 | 0.838 | 5.41E-33 | 12_Resting_NKcells |
| NCALD | 2.701906699 | 0.447 | 0.166 | 3.09E-32 | 12_Resting_NKcells |
| TXK | 3.066924544 | 0.387 | 0.127 | 1.83E-31 | 12_Resting_NKcells |
| PFN1 | -1.43648398 | 0.516 | 0.853 | 6.40E-31 | 12_Resting_NKcells |
| RPL24 | -1.289905292 | 0.429 | 0.837 | 6.78E-31 | 12_Resting_NKcells |
| IFITM1 | -1.683403804 | 0.378 | 0.779 | 1.24E-30 | 12_Resting_NKcells |
| VAV3 | 2.361016061 | 0.567 | 0.269 | 1.48E-30 | 12_Resting_NKcells |

| | | | | | |
|---|---|---|---|---|---|
| S100A4 | -1.740073452 | 0.267 | 0.706 | 7.81E-30 | 12_Resting_NKcells |
| BTF3 | -1.827025204 | 0.249 | 0.681 | 1.96E-29 | 12_Resting_NKcells |
| CXCR4 | -1.339983587 | 0.452 | 0.833 | 3.46E-29 | 12_Resting_NKcells |
| ATP5F1E | -1.5766592 | 0.286 | 0.727 | 3.89E-29 | 12_Resting_NKcells |
| CDK17 | 1.848947038 | 0.668 | 0.401 | 4.00E-29 | 12_Resting_NKcells |
| RPL27 | -1.551121042 | 0.295 | 0.729 | 5.41E-29 | 12_Resting_NKcells |
| CARD11 | 2.619468789 | 0.53 | 0.26 | 5.35E-28 | 12_Resting_NKcells |
| ZEB2 | 1.98220416 | 0.687 | 0.429 | 5.92E-28 | 12_Resting_NKcells |
| ZBTB16 | 2.38262929 | 0.525 | 0.242 | 9.96E-28 | 12_Resting_NKcells |
| EEF1B2 | -1.487420377 | 0.3 | 0.717 | 1.40E-26 | 12_Resting_NKcells |
| EEF2 | -1.461550655 | 0.3 | 0.714 | 2.44E-26 | 12_Resting_NKcells |
| ACTG1 | -1.241384192 | 0.465 | 0.821 | 3.54E-26 | 12_Resting_NKcells |
| RPL38 | -1.869442497 | 0.207 | 0.632 | 1.24E-25 | 12_Resting_NKcells |
| GZMK | -3.426664188 | 0.074 | 0.477 | 1.27E-25 | 12_Resting_NKcells |
| PFDN5 | -1.295265945 | 0.318 | 0.747 | 2.55E-25 | 12_Resting_NKcells |
| ATP5MG | -1.780098617 | 0.184 | 0.626 | 2.60E-25 | 12_Resting_NKcells |
| EEF1D | -1.196124347 | 0.387 | 0.804 | 2.92E-25 | 12_Resting_NKcells |
| SMYD3 | 2.213481577 | 0.516 | 0.253 | 1.97E-24 | 12_Resting_NKcells |
| NPM1 | -1.515417715 | 0.263 | 0.681 | 2.58E-24 | 12_Resting_NKcells |
| RPL36AL | -1.447523749 | 0.267 | 0.693 | 4.22E-24 | 12_Resting_NKcells |
| RPL31 | -1.576990457 | 0.249 | 0.668 | 7.15E-24 | 12_Resting_NKcells |
| RPL4 | -1.502736853 | 0.295 | 0.696 | 7.88E-24 | 12_Resting_NKcells |
| BRAF | 2.206906146 | 0.553 | 0.295 | 8.70E-24 | 12_Resting_NKcells |
| RPL7 | -1.558378407 | 0.217 | 0.649 | 1.49E-23 | 12_Resting_NKcells |
| HSPA8 | -1.217758727 | 0.373 | 0.771 | 1.81E-23 | 12_Resting_NKcells |
| NFKB1 | 2.067645217 | 0.691 | 0.479 | 2.36E-23 | 12_Resting_NKcells |
| MIF | -1.367619019 | 0.24 | 0.662 | 3.23E-23 | 12_Resting_NKcells |
| CD2 | -2.173569457 | 0.152 | 0.549 | 5.15E-23 | 12_Resting_NKcells |
| COX7C | -1.822912533 | 0.207 | 0.606 | 6.52E-23 | 12_Resting_NKcells |
| ARHGDIB | -1.101480324 | 0.493 | 0.819 | 6.72E-23 | 12_Resting_NKcells |
| CD52 | -1.515425535 | 0.203 | 0.618 | 9.03E-23 | 12_Resting_NKcells |
| RPL22 | -1.262368583 | 0.382 | 0.764 | 1.10E-22 | 12_Resting_NKcells |
| FNDC3B | 2.394725026 | 0.452 | 0.206 | 1.28E-22 | 12_Resting_NKcells |
| COX4I1 | -1.228855507 | 0.313 | 0.723 | 2.62E-22 | 12_Resting_NKcells |
| HLA-E | -1.014949881 | 0.535 | 0.864 | 3.01E-22 | 12_Resting_NKcells |
| YES1 | 2.640748662 | 0.336 | 0.118 | 3.46E-22 | 12_Resting_NKcells |
| RPL27A | -1.531503993 | 0.24 | 0.647 | 3.49E-22 | 12_Resting_NKcells |
| CHCHD2 | -1.479805457 | 0.217 | 0.609 | 1.24E-21 | 12_Resting_NKcells |
| ATP5MC2 | -1.517180269 | 0.272 | 0.652 | 1.40E-21 | 12_Resting_NKcells |
| TOMM7 | -1.458646452 | 0.207 | 0.617 | 1.82E-21 | 12_Resting_NKcells |
| HCST | -1.686500378 | 0.235 | 0.613 | 2.05E-21 | 12_Resting_NKcells |
| JARID2 | 1.927766866 | 0.571 | 0.334 | 2.81E-21 | 12_Resting_NKcells |
| MYL6 | -1.087270879 | 0.438 | 0.791 | 4.02E-21 | 12_Resting_NKcells |
| OAZ1 | -1.274847386 | 0.359 | 0.738 | 6.08E-21 | 12_Resting_NKcells |
| GFOD1 | 2.68804559 | 0.346 | 0.128 | 1.07E-20 | 12_Resting_NKcells |
| NFAT5 | 1.760109976 | 0.544 | 0.302 | 1.28E-20 | 12_Resting_NKcells |

| | | | | | |
|---|---|---|---|---|---|
| LAPTM5 | -1.290829079 | 0.341 | 0.705 | 2.44E-20 | 12_Resting_NKcells |
| RPL41 | -1.755842769 | 0.447 | 0.785 | 2.47E-20 | 12_Resting_NKcells |
| LYN | 2.397144375 | 0.364 | 0.144 | 3.35E-20 | 12_Resting_NKcells |
| CST7 | -1.650159642 | 0.249 | 0.626 | 4.01E-20 | 12_Resting_NKcells |
| ZFAND3 | 1.843995218 | 0.576 | 0.351 | 6.72E-20 | 12_Resting_NKcells |
| HINT1 | -1.437896336 | 0.217 | 0.612 | 7.18E-20 | 12_Resting_NKcells |
| PLAAT4 | -2.237855132 | 0.106 | 0.474 | 7.68E-20 | 12_Resting_NKcells |
| XYLT1 | 2.133651905 | 0.41 | 0.181 | 2.05E-19 | 12_Resting_NKcells |
| PPIA | -1.035388248 | 0.465 | 0.798 | 2.11E-19 | 12_Resting_NKcells |
| PLCB1 | 2.420065117 | 0.35 | 0.137 | 2.18E-19 | 12_Resting_NKcells |
| MYL12A | -1.422351577 | 0.323 | 0.676 | 3.16E-19 | 12_Resting_NKcells |
| EXT1 | 2.785649551 | 0.313 | 0.112 | 4.94E-19 | 12_Resting_NKcells |
| SRP14 | -1.426208988 | 0.212 | 0.598 | 6.35E-19 | 12_Resting_NKcells |
| RIN3 | 2.487190791 | 0.378 | 0.159 | 6.59E-19 | 12_Resting_NKcells |
| MYL12B | -1.410511906 | 0.258 | 0.621 | 1.25E-18 | 12_Resting_NKcells |
| LY6E | -2.144131808 | 0.129 | 0.483 | 1.67E-18 | 12_Resting_NKcells |
| PSME1 | -1.656901783 | 0.194 | 0.56 | 1.75E-18 | 12_Resting_NKcells |
| RPS11 | -1.133684739 | 0.276 | 0.682 | 1.86E-18 | 12_Resting_NKcells |
| APBA2 | 2.404275435 | 0.364 | 0.153 | 1.96E-18 | 12_Resting_NKcells |
| GPATCH8 | 1.833192931 | 0.512 | 0.293 | 5.23E-18 | 12_Resting_NKcells |
| ARPC3 | -1.324536921 | 0.253 | 0.614 | 5.69E-18 | 12_Resting_NKcells |
| C12orf57 | -2.094445064 | 0.111 | 0.465 | 6.28E-18 | 12_Resting_NKcells |
| GZMA | -1.703388105 | 0.198 | 0.551 | 6.64E-18 | 12_Resting_NKcells |
| TUBA4A | -1.623922728 | 0.226 | 0.581 | 9.55E-18 | 12_Resting_NKcells |
| UQCRB | -1.758155985 | 0.152 | 0.51 | 9.73E-18 | 12_Resting_NKcells |
| EIF3K | -1.55024557 | 0.203 | 0.563 | 1.06E-17 | 12_Resting_NKcells |
| S100A10 | -1.280015455 | 0.244 | 0.625 | 1.12E-17 | 12_Resting_NKcells |
| PPDPF | -1.295056106 | 0.263 | 0.624 | 1.37E-17 | 12_Resting_NKcells |
| ITM2B | -1.081526931 | 0.456 | 0.77 | 1.49E-17 | 12_Resting_NKcells |
| AUTS2 | 1.889469163 | 0.53 | 0.301 | 1.61E-17 | 12_Resting_NKcells |
| CORO1A | -1.344309381 | 0.304 | 0.65 | 1.64E-17 | 12_Resting_NKcells |
| LEPROTL1 | -1.306070657 | 0.3 | 0.662 | 1.66E-17 | 12_Resting_NKcells |
| S100A6 | -1.198485963 | 0.276 | 0.637 | 1.88E-17 | 12_Resting_NKcells |
| COMMD6 | -1.817779468 | 0.171 | 0.518 | 2.76E-17 | 12_Resting_NKcells |
| MAPK1 | 1.995942 | 0.465 | 0.246 | 3.28E-17 | 12_Resting_NKcells |
| PLCL2 | 2.21128539 | 0.429 | 0.217 | 9.34E-17 | 12_Resting_NKcells |
| TRBC2 | -2.156168337 | 0.088 | 0.418 | 9.93E-17 | 12_Resting_NKcells |
| HIGD2A | -2.379378573 | 0.106 | 0.432 | 1.62E-16 | 12_Resting_NKcells |
| PPP3CA | 1.907032226 | 0.479 | 0.256 | 1.78E-16 | 12_Resting_NKcells |
| CCL5 | -1.429943505 | 0.41 | 0.699 | 2.44E-16 | 12_Resting_NKcells |
| CRIP1 | -1.153499966 | 0.281 | 0.645 | 2.66E-16 | 12_Resting_NKcells |
| SLC25A6 | -1.368468903 | 0.207 | 0.572 | 2.68E-16 | 12_Resting_NKcells |
| NDUFS5 | -1.630743178 | 0.134 | 0.477 | 3.73E-16 | 12_Resting_NKcells |
| RPS20 | -1.492625133 | 0.194 | 0.546 | 4.01E-16 | 12_Resting_NKcells |
| OST4 | -1.74425705 | 0.175 | 0.516 | 4.05E-16 | 12_Resting_NKcells |
| CLASP1 | 2.091766428 | 0.433 | 0.221 | 4.27E-16 | 12_Resting_NKcells |

| | | | | | |
|---|---|---|---|---|---|
| UBB | -1.053376761 | 0.295 | 0.677 | 5.82E-16 | 12_Resting_NKcells |
| UBL5 | -2.055898539 | 0.106 | 0.437 | 8.59E-16 | 12_Resting_NKcells |
| SSR4 | -1.384633923 | 0.189 | 0.539 | 9.27E-16 | 12_Resting_NKcells |
| GMFG | -1.596611959 | 0.166 | 0.514 | 9.65E-16 | 12_Resting_NKcells |
| H2AFZ | -1.77643203 | 0.157 | 0.488 | 1.04E-15 | 12_Resting_NKcells |
| PCED1B-AS1 | -1.749112982 | 0.152 | 0.493 | 1.06E-15 | 12_Resting_NKcells |
| JAZF1 | 2.148232988 | 0.442 | 0.235 | 1.16E-15 | 12_Resting_NKcells |
| LDHB | -1.811056163 | 0.143 | 0.473 | 1.19E-15 | 12_Resting_NKcells |
| RPS4Y1 | -1.663595557 | 0.217 | 0.539 | 1.52E-15 | 12_Resting_NKcells |
| COTL1 | -1.449026381 | 0.24 | 0.575 | 1.63E-15 | 12_Resting_NKcells |
| CALM2 | -1.139093426 | 0.281 | 0.622 | 3.70E-15 | 12_Resting_NKcells |
| APRT | -2.106242577 | 0.101 | 0.413 | 7.60E-15 | 12_Resting_NKcells |
| COX6A1 | -1.190127184 | 0.184 | 0.52 | 7.66E-15 | 12_Resting_NKcells |
| MAP3K8 | 1.526792323 | 0.553 | 0.344 | 9.07E-15 | 12_Resting_NKcells |
| CD8A | -2.782424449 | 0.111 | 0.404 | 1.05E-14 | 12_Resting_NKcells |
| CLIC1 | -1.249443824 | 0.249 | 0.581 | 3.10E-14 | 12_Resting_NKcells |
| COX5B | -1.607105272 | 0.111 | 0.438 | 3.12E-14 | 12_Resting_NKcells |
| CD48 | -1.982557639 | 0.097 | 0.41 | 3.52E-14 | 12_Resting_NKcells |
| YWHAB | -1.193618029 | 0.276 | 0.603 | 5.17E-14 | 12_Resting_NKcells |
| TLE5 | -1.62553203 | 0.171 | 0.487 | 5.40E-14 | 12_Resting_NKcells |
| DYRK1A | 1.86463043 | 0.438 | 0.237 | 6.09E-14 | 12_Resting_NKcells |
| RAB8B | 1.908528462 | 0.475 | 0.273 | 6.41E-14 | 12_Resting_NKcells |
| ITM2A | -1.852215185 | 0.147 | 0.453 | 9.03E-14 | 12_Resting_NKcells |
| SUMO2 | -1.290073439 | 0.226 | 0.549 | 1.17E-13 | 12_Resting_NKcells |
| AKT3 | 1.819269896 | 0.438 | 0.229 | 1.41E-13 | 12_Resting_NKcells |
| RAC2 | -1.309918611 | 0.212 | 0.528 | 2.27E-13 | 12_Resting_NKcells |
| TMA7 | -1.08122482 | 0.267 | 0.602 | 2.53E-13 | 12_Resting_NKcells |
| LIME1 | -2.38924266 | 0.069 | 0.359 | 2.71E-13 | 12_Resting_NKcells |
| SLC25A5 | -1.696647815 | 0.124 | 0.432 | 2.96E-13 | 12_Resting_NKcells |
| YBX1 | -1.327176302 | 0.226 | 0.541 | 5.65E-13 | 12_Resting_NKcells |
| SNRPD2 | -1.223088813 | 0.18 | 0.506 | 5.77E-13 | 12_Resting_NKcells |
| IL7R | -1.289890234 | 0.258 | 0.566 | 5.81E-13 | 12_Resting_NKcells |
| RPL23 | -1.316277555 | 0.184 | 0.493 | 6.51E-13 | 12_Resting_NKcells |
| CD99 | -1.038358245 | 0.332 | 0.662 | 9.57E-13 | 12_Resting_NKcells |
| EEF1G | -1.035400626 | 0.332 | 0.644 | 1.48E-12 | 12_Resting_NKcells |
| COX7A2 | -1.723202517 | 0.134 | 0.424 | 1.95E-12 | 12_Resting_NKcells |
| ATP6V0E1 | -1.563150529 | 0.138 | 0.438 | 2.00E-12 | 12_Resting_NKcells |
| PRDX1 | -2.002327198 | 0.101 | 0.389 | 2.34E-12 | 12_Resting_NKcells |
| CIB1 | -1.475852734 | 0.166 | 0.469 | 2.53E-12 | 12_Resting_NKcells |
| PPIB | -1.099287758 | 0.249 | 0.572 | 3.14E-12 | 12_Resting_NKcells |
| LSP1 | -1.410628312 | 0.217 | 0.521 | 3.19E-12 | 12_Resting_NKcells |
| CD3G | -2.204558489 | 0.092 | 0.374 | 3.27E-12 | 12_Resting_NKcells |
| ISG20 | -1.123755943 | 0.364 | 0.68 | 3.37E-12 | 12_Resting_NKcells |
| RGS10 | -1.965869101 | 0.074 | 0.356 | 3.50E-12 | 12_Resting_NKcells |
| TPI1 | -1.363137286 | 0.166 | 0.472 | 4.86E-12 | 12_Resting_NKcells |
| MZT2B | -1.872219048 | 0.101 | 0.391 | 6.01E-12 | 12_Resting_NKcells |

| | | | | | |
|---|---|---|---|---|---|
| KLRD1 | 1.516266334 | 0.406 | 0.201 | 8.41E-12 | 12_Resting_NKcells |
| RGS1 | -1.215221241 | 0.373 | 0.635 | 1.08E-11 | 12_Resting_NKcells |
| PCBP1 | -1.417180476 | 0.129 | 0.431 | 1.42E-11 | 12_Resting_NKcells |
| ICAM3 | -2.124531878 | 0.06 | 0.332 | 1.55E-11 | 12_Resting_NKcells |
| SPOCK2 | -1.421259911 | 0.226 | 0.517 | 1.70E-11 | 12_Resting_NKcells |
| ELOB | -1.206945155 | 0.138 | 0.439 | 1.73E-11 | 12_Resting_NKcells |
| PSMB8 | -1.791344637 | 0.088 | 0.366 | 1.81E-11 | 12_Resting_NKcells |
| COX6B1 | -1.387194307 | 0.166 | 0.457 | 2.15E-11 | 12_Resting_NKcells |
| CD8B | -3.076784139 | 0.041 | 0.293 | 2.48E-11 | 12_Resting_NKcells |
| RPS17 | -1.276967058 | 0.147 | 0.44 | 2.69E-11 | 12_Resting_NKcells |
| RABAC1 | -1.917589862 | 0.101 | 0.375 | 3.28E-11 | 12_Resting_NKcells |
| GUK1 | -1.18082771 | 0.217 | 0.524 | 3.28E-11 | 12_Resting_NKcells |
| S100A11 | -1.001465867 | 0.323 | 0.63 | 4.79E-11 | 12_Resting_NKcells |
| UQCR11 | -1.152598202 | 0.171 | 0.472 | 6.03E-11 | 12_Resting_NKcells |
| NDUFA4 | -1.253013975 | 0.157 | 0.455 | 7.12E-11 | 12_Resting_NKcells |
| HLA-DRB1 | -1.418525018 | 0.276 | 0.561 | 8.50E-11 | 12_Resting_NKcells |
| GSTK1 | -1.317038301 | 0.198 | 0.483 | 9.09E-11 | 12_Resting_NKcells |
| KRTCAP2 | -1.405967932 | 0.111 | 0.392 | 9.25E-11 | 12_Resting_NKcells |
| PSMA7 | -1.216701875 | 0.194 | 0.494 | 9.74E-11 | 12_Resting_NKcells |
| CNN2 | -1.938673672 | 0.111 | 0.374 | 1.08E-10 | 12_Resting_NKcells |
| ATP5IF1 | -1.544431482 | 0.111 | 0.388 | 1.33E-10 | 12_Resting_NKcells |
| COX8A | -1.368971 | 0.166 | 0.447 | 1.70E-10 | 12_Resting_NKcells |
| SNHG29 | -1.07308013 | 0.23 | 0.533 | 1.94E-10 | 12_Resting_NKcells |
| EIF3F | -1.19772421 | 0.194 | 0.476 | 2.07E-10 | 12_Resting_NKcells |
| TRMT112 | -1.433479935 | 0.129 | 0.405 | 2.31E-10 | 12_Resting_NKcells |
| SAP18 | -1.477731152 | 0.111 | 0.385 | 2.41E-10 | 12_Resting_NKcells |
| ARL6IP5 | -1.317230842 | 0.175 | 0.467 | 4.39E-10 | 12_Resting_NKcells |
| SPCS1 | -1.728637661 | 0.101 | 0.367 | 5.37E-10 | 12_Resting_NKcells |
| GIMAP7 | -1.960441711 | 0.069 | 0.323 | 5.95E-10 | 12_Resting_NKcells |
| ATP5F1D | -1.324401722 | 0.147 | 0.423 | 6.24E-10 | 12_Resting_NKcells |
| ANAPC16 | -1.410007945 | 0.129 | 0.398 | 8.42E-10 | 12_Resting_NKcells |
| EDF1 | -1.070851474 | 0.184 | 0.478 | 9.79E-10 | 12_Resting_NKcells |
| PSMB9 | -1.09845216 | 0.134 | 0.418 | 1.30E-09 | 12_Resting_NKcells |
| CD27 | -3.581523315 | 0.023 | 0.247 | 1.31E-09 | 12_Resting_NKcells |
| COX6C | -1.142418458 | 0.203 | 0.491 | 1.42E-09 | 12_Resting_NKcells |
| RAN | -1.192752192 | 0.18 | 0.452 | 2.01E-09 | 12_Resting_NKcells |
| NOP53 | -1.119010141 | 0.207 | 0.485 | 2.24E-09 | 12_Resting_NKcells |
| C9orf16 | -1.325127007 | 0.124 | 0.386 | 2.47E-09 | 12_Resting_NKcells |
| CORO1B | -2.200566792 | 0.078 | 0.318 | 3.31E-09 | 12_Resting_NKcells |
| ZYX | -1.503018224 | 0.083 | 0.334 | 3.54E-09 | 12_Resting_NKcells |
| NDUFA13 | -1.270822946 | 0.198 | 0.472 | 4.23E-09 | 12_Resting_NKcells |
| GABARAP | -1.045970756 | 0.267 | 0.549 | 4.51E-09 | 12_Resting_NKcells |
| ALDOA | -1.189253757 | 0.166 | 0.441 | 4.72E-09 | 12_Resting_NKcells |
| C19orf53 | -1.889929111 | 0.083 | 0.328 | 5.08E-09 | 12_Resting_NKcells |
| ERP29 | -1.347103609 | 0.101 | 0.354 | 6.36E-09 | 12_Resting_NKcells |
| DAZAP2 | -1.014227024 | 0.244 | 0.53 | 7.74E-09 | 12_Resting_NKcells |

| | | | | | |
|---|---|---|---|---|---|
| HLA-DPB1 | -1.3520467 | 0.203 | 0.47 | 8.69E-09 | 12_Resting_NKcells |
| TUBA1B | -1.124522638 | 0.221 | 0.513 | 9.32E-09 | 12_Resting_NKcells |
| SAT1 | -1.161771691 | 0.512 | 0.739 | 1.10E-08 | 12_Resting_NKcells |
| UQCRH | -1.091473101 | 0.147 | 0.408 | 1.18E-08 | 12_Resting_NKcells |
| ICOS | -2.379175751 | 0.092 | 0.326 | 1.24E-08 | 12_Resting_NKcells |
| SNHG6 | -1.30487275 | 0.147 | 0.406 | 1.35E-08 | 12_Resting_NKcells |
| DRAP1 | -1.358965105 | 0.124 | 0.381 | 1.68E-08 | 12_Resting_NKcells |
| HLA-DPA1 | -1.257719646 | 0.272 | 0.528 | 2.07E-08 | 12_Resting_NKcells |
| PGAM1 | -1.675356015 | 0.088 | 0.326 | 2.50E-08 | 12_Resting_NKcells |
| NDUFA1 | -1.699274413 | 0.088 | 0.328 | 2.56E-08 | 12_Resting_NKcells |
| PRR13 | -1.250084967 | 0.138 | 0.39 | 2.69E-08 | 12_Resting_NKcells |
| EIF3G | -1.034856643 | 0.161 | 0.431 | 2.92E-08 | 12_Resting_NKcells |
| GZMM | -1.835855292 | 0.083 | 0.314 | 2.97E-08 | 12_Resting_NKcells |
| GPR183 | -1.586306803 | 0.12 | 0.362 | 3.16E-08 | 12_Resting_NKcells |
| SNRPB | -1.604651467 | 0.106 | 0.347 | 3.66E-08 | 12_Resting_NKcells |
| CD6 | -1.57865147 | 0.115 | 0.359 | 3.77E-08 | 12_Resting_NKcells |
| LGALS1 | -1.340050719 | 0.171 | 0.422 | 4.90E-08 | 12_Resting_NKcells |
| PEBP1 | -1.471391846 | 0.097 | 0.337 | 4.90E-08 | 12_Resting_NKcells |
| TERF2IP | -1.284072017 | 0.18 | 0.431 | 7.19E-08 | 12_Resting_NKcells |
| RBM3 | -1.101570287 | 0.235 | 0.492 | 7.29E-08 | 12_Resting_NKcells |
| ABRACL | -1.498021705 | 0.101 | 0.336 | 7.68E-08 | 12_Resting_NKcells |
| PRDM1 | -1.178831559 | 0.161 | 0.421 | 8.35E-08 | 12_Resting_NKcells |
| OCIAD2 | -3.101745343 | 0.032 | 0.234 | 9.51E-08 | 12_Resting_NKcells |
| CAMK4 | -1.051248548 | 0.161 | 0.416 | 1.11E-07 | 12_Resting_NKcells |
| ARL6IP4 | -1.338372762 | 0.124 | 0.362 | 1.13E-07 | 12_Resting_NKcells |
| CIAO2B | -1.679875361 | 0.074 | 0.294 | 1.32E-07 | 12_Resting_NKcells |
| ANXA1 | -1.02232652 | 0.346 | 0.6 | 1.35E-07 | 12_Resting_NKcells |
| GABARAPL2 | -1.660410792 | 0.088 | 0.31 | 1.40E-07 | 12_Resting_NKcells |
| GIMAP4 | -2.174241935 | 0.06 | 0.273 | 1.60E-07 | 12_Resting_NKcells |
| ISG15 | -2.832789557 | 0.046 | 0.25 | 1.63E-07 | 12_Resting_NKcells |
| LTB | -1.691319135 | 0.106 | 0.339 | 1.69E-07 | 12_Resting_NKcells |
| UQCR10 | -1.027294529 | 0.115 | 0.356 | 1.70E-07 | 12_Resting_NKcells |
| C4orf3 | -1.44478183 | 0.097 | 0.327 | 1.80E-07 | 12_Resting_NKcells |
| CYCS | -1.291500853 | 0.12 | 0.363 | 1.86E-07 | 12_Resting_NKcells |
| GAREM1 | -1.75018537 | 0.092 | 0.308 | 2.56E-07 | 12_Resting_NKcells |
| NEDD8 | -1.184604813 | 0.097 | 0.328 | 2.61E-07 | 12_Resting_NKcells |
| TOMM6 | -1.569389185 | 0.078 | 0.295 | 3.06E-07 | 12_Resting_NKcells |
| PRDX2 | -2.292340659 | 0.037 | 0.238 | 3.69E-07 | 12_Resting_NKcells |
| ATP5MPL | -2.013373932 | 0.055 | 0.264 | 3.72E-07 | 12_Resting_NKcells |
| BAX | -2.004613511 | 0.06 | 0.268 | 3.78E-07 | 12_Resting_NKcells |
| TMEM258 | -1.684422602 | 0.083 | 0.302 | 3.82E-07 | 12_Resting_NKcells |
| HLA-DQB1 | -1.465433697 | 0.106 | 0.328 | 4.77E-07 | 12_Resting_NKcells |
| ATP5MF | -1.590803035 | 0.083 | 0.302 | 5.47E-07 | 12_Resting_NKcells |
| CITED2 | -1.665488603 | 0.101 | 0.318 | 6.20E-07 | 12_Resting_NKcells |
| DAD1 | -1.192371879 | 0.092 | 0.313 | 6.37E-07 | 12_Resting_NKcells |
| PKM | -1.167559268 | 0.134 | 0.366 | 8.04E-07 | 12_Resting_NKcells |

| | | | | | |
|---|---|---|---|---|---|
| JTB | -1.317336536 | 0.106 | 0.332 | 8.31E-07 | 12_Resting_NKcells |
| HLA-DRA | -1.201198538 | 0.281 | 0.534 | 8.71E-07 | 12_Resting_NKcells |
| MZT2A | -2.01465635 | 0.06 | 0.262 | 9.76E-07 | 12_Resting_NKcells |
| PBX4 | -1.292573537 | 0.129 | 0.357 | 1.11E-06 | 12_Resting_NKcells |
| NDUFB10 | -1.400055055 | 0.078 | 0.288 | 1.14E-06 | 12_Resting_NKcells |
| CD5 | -1.86397599 | 0.06 | 0.262 | 1.33E-06 | 12_Resting_NKcells |
| BLOC1S1 | -1.614397265 | 0.06 | 0.264 | 1.42E-06 | 12_Resting_NKcells |
| ARHGDIA | -1.069561496 | 0.157 | 0.391 | 1.45E-06 | 12_Resting_NKcells |
| SF3B5 | -1.392431573 | 0.083 | 0.301 | 1.62E-06 | 12_Resting_NKcells |
| ARF5 | -1.344972955 | 0.097 | 0.319 | 1.70E-06 | 12_Resting_NKcells |
| GPSM3 | -1.074907681 | 0.166 | 0.402 | 1.71E-06 | 12_Resting_NKcells |
| TOMM20 | -1.087772996 | 0.12 | 0.344 | 2.03E-06 | 12_Resting_NKcells |
| ATP5PO | -1.226063176 | 0.111 | 0.337 | 2.05E-06 | 12_Resting_NKcells |
| ATP6V1F | -1.505563018 | 0.097 | 0.311 | 2.18E-06 | 12_Resting_NKcells |
| HMGN1 | -1.008890481 | 0.152 | 0.386 | 2.34E-06 | 12_Resting_NKcells |
| RHOG | -1.076086868 | 0.134 | 0.363 | 2.43E-06 | 12_Resting_NKcells |
| SMDT1 | -1.49860107 | 0.092 | 0.305 | 2.45E-06 | 12_Resting_NKcells |
| ISCU | -1.543222725 | 0.078 | 0.286 | 2.61E-06 | 12_Resting_NKcells |
| EVI2B | -1.406184026 | 0.097 | 0.312 | 2.73E-06 | 12_Resting_NKcells |
| CSTB | -1.150136268 | 0.129 | 0.357 | 3.22E-06 | 12_Resting_NKcells |
| PSME2 | -1.126671555 | 0.175 | 0.405 | 3.44E-06 | 12_Resting_NKcells |
| MICOS10 | -1.51153797 | 0.078 | 0.282 | 4.20E-06 | 12_Resting_NKcells |
| ATP5MD | -1.325367401 | 0.101 | 0.314 | 5.41E-06 | 12_Resting_NKcells |
| CUTA | -1.089710836 | 0.138 | 0.361 | 5.43E-06 | 12_Resting_NKcells |
| NDUFB11 | -1.13460666 | 0.138 | 0.367 | 5.90E-06 | 12_Resting_NKcells |
| TUBB | -1.401271566 | 0.106 | 0.319 | 5.97E-06 | 12_Resting_NKcells |
| JPT1 | -1.120642621 | 0.106 | 0.32 | 7.73E-06 | 12_Resting_NKcells |
| TAF10 | -1.583097885 | 0.106 | 0.309 | 7.93E-06 | 12_Resting_NKcells |
| HMGN2 | -1.200961236 | 0.161 | 0.385 | 1.00E-05 | 12_Resting_NKcells |
| PLP2 | -1.314237486 | 0.134 | 0.346 | 1.19E-05 | 12_Resting_NKcells |
| FKBP8 | -1.159888154 | 0.101 | 0.31 | 1.34E-05 | 12_Resting_NKcells |
| NDUFC2 | -1.352231239 | 0.097 | 0.3 | 1.41E-05 | 12_Resting_NKcells |
| SNHG8 | -1.297687161 | 0.111 | 0.322 | 1.57E-05 | 12_Resting_NKcells |
| SCAND1 | -1.25085597 | 0.097 | 0.301 | 1.69E-05 | 12_Resting_NKcells |
| PARK7 | -1.053582338 | 0.189 | 0.424 | 1.88E-05 | 12_Resting_NKcells |
| POLR2L | -1.17392135 | 0.115 | 0.32 | 2.00E-05 | 12_Resting_NKcells |
| CYTOR | -1.176046994 | 0.147 | 0.359 | 2.03E-05 | 12_Resting_NKcells |
| DUSP4 | -1.355820258 | 0.166 | 0.37 | 2.15E-05 | 12_Resting_NKcells |
| BRK1 | -1.345982343 | 0.129 | 0.335 | 2.16E-05 | 12_Resting_NKcells |
| COX7B | -1.057608929 | 0.101 | 0.308 | 2.52E-05 | 12_Resting_NKcells |
| GTF3A | -1.128221339 | 0.101 | 0.304 | 3.02E-05 | 12_Resting_NKcells |
| HLA-DRB5 | -1.507836399 | 0.157 | 0.36 | 3.79E-05 | 12_Resting_NKcells |
| C11orf58 | -1.047889147 | 0.147 | 0.364 | 5.54E-05 | 12_Resting_NKcells |
| CSNK2B | -1.078341046 | 0.143 | 0.352 | 7.37E-05 | 12_Resting_NKcells |
| NSA2 | -1.039694547 | 0.106 | 0.306 | 8.20E-05 | 12_Resting_NKcells |
| ANXA2 | -1.05373442 | 0.143 | 0.349 | 9.81E-05 | 12_Resting_NKcells |

| | | | | | |
|---|---|---|---|---|---|
| SOCS1 | -1.099693889 | 0.138 | 0.338 | 0.00013 | 12_Resting_NKcells |
| HMGB2 | -1.180937576 | 0.147 | 0.35 | 0.00013 | 12_Resting_NKcells |
| CTSC | -1.020995013 | 0.171 | 0.383 | 0.000184 | 12_Resting_NKcells |
| MKI67 | 6.022656514 | 0.565 | 0.018 | 0 | 13_Proliferating_TCells |
| RRM2 | 6.529848104 | 0.531 | 0.009 | 0 | 13_Proliferating_TCells |
| TYMS | 6.175499628 | 0.531 | 0.011 | 0 | 13_Proliferating_TCells |
| PCLAF | 5.757557626 | 0.478 | 0.011 | 0 | 13_Proliferating_TCells |
| TOP2A | 5.587294055 | 0.478 | 0.016 | 0 | 13_Proliferating_TCells |
| ASPM | 5.992616334 | 0.415 | 0.008 | 0 | 13_Proliferating_TCells |
| TPX2 | 5.09113654 | 0.406 | 0.013 | 0 | 13_Proliferating_TCells |
| CLSPN | 4.944873266 | 0.391 | 0.013 | 0 | 13_Proliferating_TCells |
| UBE2C | 7.142353002 | 0.362 | 0.004 | 0 | 13_Proliferating_TCells |
| ZWINT | 6.377311109 | 0.348 | 0.005 | 0 | 13_Proliferating_TCells |
| GTSE1 | 6.588172489 | 0.3 | 0.003 | 0 | 13_Proliferating_TCells |
| PKMYT1 | 6.645948609 | 0.271 | 0.003 | 0 | 13_Proliferating_TCells |
| BIRC5 | 6.495429894 | 0.271 | 0.003 | 0 | 13_Proliferating_TCells |
| CDT1 | 6.336194733 | 0.266 | 0.003 | 0 | 13_Proliferating_TCells |
| MELK | 6.04934265 | 0.261 | 0.004 | 0 | 13_Proliferating_TCells |
| AURKB | 6.466193325 | 0.242 | 0.002 | 0 | 13_Proliferating_TCells |
| TROAP | 7.947030881 | 0.237 | 0.001 | 0 | 13_Proliferating_TCells |
| POLQ | 5.87241859 | 0.237 | 0.002 | 0 | 13_Proliferating_TCells |
| CDCA5 | 6.841528568 | 0.213 | 0.002 | 0 | 13_Proliferating_TCells |
| SPC25 | 6.666471511 | 0.208 | 0.002 | 0 | 13_Proliferating_TCells |
| CCNB2 | 5.608495365 | 0.242 | 0.004 | 4.20E-296 | 13_Proliferating_TCells |
| DIAPH3 | 5.903164931 | 0.251 | 0.004 | 1.14E-295 | 13_Proliferating_TCells |
| KIF11 | 5.150071194 | 0.353 | 0.012 | 1.15E-281 | 13_Proliferating_TCells |
| ASF1B | 4.837661099 | 0.295 | 0.007 | 1.52E-279 | 13_Proliferating_TCells |
| UHRF1 | 5.437064108 | 0.28 | 0.007 | 2.73E-276 | 13_Proliferating_TCells |
| KIF15 | 5.801646327 | 0.217 | 0.003 | 9.05E-275 | 13_Proliferating_TCells |
| DTL | 5.068063249 | 0.304 | 0.008 | 1.46E-274 | 13_Proliferating_TCells |
| CDCA8 | 5.536579503 | 0.237 | 0.004 | 4.68E-267 | 13_Proliferating_TCells |
| NCAPG | 5.130979025 | 0.242 | 0.005 | 2.86E-260 | 13_Proliferating_TCells |
| CDK1 | 4.559588115 | 0.372 | 0.015 | 4.08E-256 | 13_Proliferating_TCells |
| KNL1 | 4.597314487 | 0.367 | 0.015 | 1.25E-249 | 13_Proliferating_TCells |
| CCNA2 | 5.14165072 | 0.251 | 0.006 | 6.89E-247 | 13_Proliferating_TCells |
| CDCA2 | 5.852799251 | 0.203 | 0.003 | 1.36E-244 | 13_Proliferating_TCells |
| KIFC1 | 5.002642013 | 0.271 | 0.008 | 6.49E-230 | 13_Proliferating_TCells |
| KIF23 | 5.086517461 | 0.237 | 0.006 | 9.30E-227 | 13_Proliferating_TCells |
| RAD51AP1 | 5.116239559 | 0.246 | 0.006 | 4.72E-226 | 13_Proliferating_TCells |
| ANLN | 4.99854175 | 0.237 | 0.006 | 8.28E-220 | 13_Proliferating_TCells |
| HIST1H1B | 5.05924743 | 0.329 | 0.014 | 3.26E-212 | 13_Proliferating_TCells |
| NUSAP1 | 4.308975068 | 0.498 | 0.038 | 4.10E-210 | 13_Proliferating_TCells |
| TK1 | 5.086674942 | 0.246 | 0.007 | 1.08E-206 | 13_Proliferating_TCells |
| CENPW | 4.623816369 | 0.266 | 0.009 | 3.17E-201 | 13_Proliferating_TCells |
| CENPF | 4.416221961 | 0.493 | 0.04 | 1.01E-193 | 13_Proliferating_TCells |
| ORC6 | 5.051959159 | 0.227 | 0.007 | 2.28E-186 | 13_Proliferating_TCells |

| | | | | | |
|---|---|---|---|---|---|
| FANCI | 3.989473528 | 0.382 | 0.024 | 4.91E-182 | 13_Proliferating_TCells |
| STIL | 4.488078899 | 0.232 | 0.008 | 3.20E-179 | 13_Proliferating_TCells |
| HIST1H2AJ | 4.792800982 | 0.213 | 0.006 | 8.59E-177 | 13_Proliferating_TCells |
| PRC1 | 3.940661587 | 0.333 | 0.019 | 1.90E-172 | 13_Proliferating_TCells |
| CIT | 4.96626862 | 0.208 | 0.006 | 3.62E-170 | 13_Proliferating_TCells |
| CDKN3 | 4.383681993 | 0.266 | 0.012 | 3.50E-169 | 13_Proliferating_TCells |
| STMN1 | 4.433940944 | 0.758 | 0.136 | 5.77E-166 | 13_Proliferating_TCells |
| ECT2 | 4.261405567 | 0.222 | 0.008 | 4.18E-162 | 13_Proliferating_TCells |
| CENPE | 4.140248255 | 0.3 | 0.018 | 1.24E-151 | 13_Proliferating_TCells |
| NUF2 | 4.632525649 | 0.213 | 0.008 | 6.92E-149 | 13_Proliferating_TCells |
| MAD2L1 | 3.846149441 | 0.329 | 0.022 | 7.95E-146 | 13_Proliferating_TCells |
| SGO2 | 3.971589997 | 0.213 | 0.009 | 1.05E-141 | 13_Proliferating_TCells |
| CENPM | 3.232822843 | 0.401 | 0.036 | 1.08E-137 | 13_Proliferating_TCells |
| CKS1B | 3.549905408 | 0.367 | 0.031 | 6.38E-136 | 13_Proliferating_TCells |
| NCAPG2 | 3.978245261 | 0.309 | 0.024 | 3.16E-122 | 13_Proliferating_TCells |
| CENPK | 2.951743149 | 0.391 | 0.042 | 9.79E-111 | 13_Proliferating_TCells |
| BUB1 | 3.752455309 | 0.217 | 0.013 | 7.29E-108 | 13_Proliferating_TCells |
| SMC2 | 3.253294857 | 0.329 | 0.031 | 6.17E-106 | 13_Proliferating_TCells |
| ATAD2 | 3.260114088 | 0.473 | 0.066 | 1.16E-105 | 13_Proliferating_TCells |
| MCM7 | 3.168961002 | 0.406 | 0.05 | 6.16E-103 | 13_Proliferating_TCells |
| NCAPH | 3.256326572 | 0.29 | 0.026 | 3.83E-98 | 13_Proliferating_TCells |
| DTYMK | 3.010944566 | 0.329 | 0.035 | 7.76E-95 | 13_Proliferating_TCells |
| CENPN | 3.051908179 | 0.3 | 0.029 | 4.05E-92 | 13_Proliferating_TCells |
| KIF22 | 2.683206226 | 0.507 | 0.087 | 2.65E-90 | 13_Proliferating_TCells |
| NSD2 | 2.606719562 | 0.493 | 0.085 | 4.00E-85 | 13_Proliferating_TCells |
| PCNA | 3.083306878 | 0.391 | 0.057 | 8.45E-83 | 13_Proliferating_TCells |
| CHAF1A | 3.56538429 | 0.232 | 0.02 | 1.16E-80 | 13_Proliferating_TCells |
| EZH2 | 2.227796812 | 0.498 | 0.09 | 1.03E-77 | 13_Proliferating_TCells |
| CENPP | 2.929664372 | 0.357 | 0.05 | 6.94E-77 | 13_Proliferating_TCells |
| BARD1 | 2.718151903 | 0.348 | 0.05 | 1.08E-71 | 13_Proliferating_TCells |
| TUBB | 3.191240161 | 0.763 | 0.305 | 8.50E-69 | 13_Proliferating_TCells |
| NCAPD2 | 2.546402886 | 0.29 | 0.036 | 1.83E-67 | 13_Proliferating_TCells |
| LIG1 | 2.687284457 | 0.295 | 0.039 | 4.82E-66 | 13_Proliferating_TCells |
| MCM3 | 2.349124561 | 0.353 | 0.054 | 8.24E-66 | 13_Proliferating_TCells |
| MCM5 | 2.56859471 | 0.367 | 0.061 | 2.71E-63 | 13_Proliferating_TCells |
| TMPO | 2.215784 | 0.594 | 0.156 | 7.23E-63 | 13_Proliferating_TCells |
| SMC4 | 2.168423813 | 0.599 | 0.16 | 1.08E-61 | 13_Proliferating_TCells |
| HELLS | 2.593834146 | 0.275 | 0.037 | 2.07E-58 | 13_Proliferating_TCells |
| CENPU | 2.743853762 | 0.227 | 0.026 | 5.33E-58 | 13_Proliferating_TCells |
| TMEM106C | 2.357479256 | 0.329 | 0.054 | 2.35E-56 | 13_Proliferating_TCells |
| LMNB1 | 2.023295098 | 0.527 | 0.13 | 5.58E-54 | 13_Proliferating_TCells |
| TUBA1B | 2.674518988 | 0.85 | 0.499 | 7.73E-54 | 13_Proliferating_TCells |
| HMGN2 | 2.68006566 | 0.778 | 0.373 | 5.74E-53 | 13_Proliferating_TCells |
| PTTG1 | 2.262811383 | 0.493 | 0.124 | 5.13E-52 | 13_Proliferating_TCells |
| HMGB2 | 2.275687192 | 0.763 | 0.337 | 7.39E-51 | 13_Proliferating_TCells |
| HIST1H4C | 3.002114754 | 0.594 | 0.198 | 1.14E-50 | 13_Proliferating_TCells |

| | | | | | |
|---|---|---|---|---|---|
| NCAPD3 | 2.165928059 | 0.295 | 0.05 | 1.57E-47 | 13_Proliferating_TCells |
| SKA2 | 2.195291754 | 0.333 | 0.063 | 3.28E-47 | 13_Proliferating_TCells |
| GMNN | 2.27411016 | 0.242 | 0.035 | 8.76E-46 | 13_Proliferating_TCells |
| DUT | 2.310597383 | 0.565 | 0.187 | 2.60E-44 | 13_Proliferating_TCells |
| DNAJC9 | 2.017825351 | 0.391 | 0.089 | 1.57E-43 | 13_Proliferating_TCells |
| RBL1 | 2.182793791 | 0.314 | 0.06 | 4.26E-43 | 13_Proliferating_TCells |
| SAE1 | 2.015867593 | 0.42 | 0.102 | 7.06E-43 | 13_Proliferating_TCells |
| KIF20B | 2.02309125 | 0.367 | 0.079 | 3.26E-42 | 13_Proliferating_TCells |
| NUDT1 | 2.169787849 | 0.386 | 0.096 | 5.44E-39 | 13_Proliferating_TCells |
| CARHSP1 | 1.768625304 | 0.469 | 0.13 | 5.61E-39 | 13_Proliferating_TCells |
| RRM1 | 1.975743734 | 0.353 | 0.081 | 1.16E-38 | 13_Proliferating_TCells |
| PHF19 | 1.884465955 | 0.29 | 0.057 | 6.61E-38 | 13_Proliferating_TCells |
| CKAP5 | 2.125387613 | 0.324 | 0.071 | 2.87E-36 | 13_Proliferating_TCells |
| HIST1H1E | 1.921604631 | 0.454 | 0.132 | 1.06E-35 | 13_Proliferating_TCells |
| DNMT1 | 1.79551253 | 0.512 | 0.166 | 2.30E-35 | 13_Proliferating_TCells |
| DDX39A | 1.522858444 | 0.57 | 0.194 | 5.52E-35 | 13_Proliferating_TCells |
| CBX5 | 1.618067076 | 0.357 | 0.085 | 7.92E-35 | 13_Proliferating_TCells |
| SMC3 | 1.79760294 | 0.483 | 0.149 | 1.07E-34 | 13_Proliferating_TCells |
| DHFR | 2.047764865 | 0.285 | 0.06 | 4.98E-34 | 13_Proliferating_TCells |
| H2AFV | 1.653719726 | 0.7 | 0.339 | 1.15E-33 | 13_Proliferating_TCells |
| DEK | 1.472462613 | 0.681 | 0.307 | 3.87E-32 | 13_Proliferating_TCells |
| CKS2 | 1.587206296 | 0.493 | 0.16 | 4.54E-32 | 13_Proliferating_TCells |
| CEP192 | 1.786496493 | 0.329 | 0.078 | 9.97E-32 | 13_Proliferating_TCells |
| MCM6 | 1.800974396 | 0.338 | 0.084 | 1.12E-31 | 13_Proliferating_TCells |
| H2AFX | 1.66733704 | 0.483 | 0.157 | 2.34E-31 | 13_Proliferating_TCells |
| SMC1A | 1.715516875 | 0.435 | 0.131 | 2.96E-30 | 13_Proliferating_TCells |
| ANP32B | 1.419786127 | 0.71 | 0.338 | 6.06E-30 | 13_Proliferating_TCells |
| TFDP1 | 1.774679081 | 0.343 | 0.091 | 3.07E-29 | 13_Proliferating_TCells |
| H2AFZ | 1.634667106 | 0.778 | 0.475 | 4.39E-29 | 13_Proliferating_TCells |
| TACC3 | 1.602804188 | 0.367 | 0.103 | 8.54E-29 | 13_Proliferating_TCells |
| IDH2 | 1.434428549 | 0.575 | 0.228 | 3.44E-28 | 13_Proliferating_TCells |
| NUCKS1 | 1.402902934 | 0.667 | 0.316 | 4.47E-28 | 13_Proliferating_TCells |
| CENPX | 1.654333086 | 0.367 | 0.105 | 1.83E-27 | 13_Proliferating_TCells |
| H2AFY | 1.496990323 | 0.551 | 0.22 | 1.96E-26 | 13_Proliferating_TCells |
| USP1 | 1.706746146 | 0.348 | 0.099 | 2.45E-26 | 13_Proliferating_TCells |
| ANP32E | 1.442913908 | 0.473 | 0.165 | 8.61E-26 | 13_Proliferating_TCells |
| RANBP1 | 1.514206991 | 0.473 | 0.174 | 3.70E-24 | 13_Proliferating_TCells |
| FAM111A | 1.770342952 | 0.324 | 0.093 | 7.22E-24 | 13_Proliferating_TCells |
| BANF1 | 1.371070883 | 0.493 | 0.185 | 8.77E-24 | 13_Proliferating_TCells |
| HIST1H1D | 1.608562918 | 0.411 | 0.135 | 3.41E-23 | 13_Proliferating_TCells |
| TOPBP1 | 1.679197216 | 0.28 | 0.073 | 4.38E-23 | 13_Proliferating_TCells |
| SSRP1 | 1.520959142 | 0.329 | 0.095 | 1.16E-22 | 13_Proliferating_TCells |
| NUP210 | 1.250090074 | 0.459 | 0.163 | 1.27E-22 | 13_Proliferating_TCells |
| SNRPD1 | 1.219816525 | 0.444 | 0.157 | 7.07E-22 | 13_Proliferating_TCells |
| COX8A | 1.187591391 | 0.739 | 0.435 | 8.54E-22 | 13_Proliferating_TCells |
| FABP5 | 1.028705349 | 0.444 | 0.16 | 1.52E-20 | 13_Proliferating_TCells |

| | | | | | |
|---|---|---|---|---|---|
| PRIM2 | 1.309594151 | 0.3 | 0.087 | 3.41E-20 | 13_Proliferating_TCells |
| PPIH | 1.292204257 | 0.353 | 0.112 | 5.46E-20 | 13_Proliferating_TCells |
| HIST1H1C | 1.935519747 | 0.329 | 0.109 | 1.15E-19 | 13_Proliferating_TCells |
| SUGP2 | 1.351231841 | 0.314 | 0.095 | 1.56E-19 | 13_Proliferating_TCells |
| BAZ1B | 1.261163796 | 0.411 | 0.147 | 1.92E-19 | 13_Proliferating_TCells |
| MIS18BP1 | 1.000177042 | 0.459 | 0.17 | 2.17E-19 | 13_Proliferating_TCells |
| NUCB2 | 1.517067715 | 0.386 | 0.14 | 4.98E-19 | 13_Proliferating_TCells |
| HNRNPR | 1.124639977 | 0.56 | 0.244 | 5.27E-19 | 13_Proliferating_TCells |
| BLOC1S1 | 1.028086058 | 0.575 | 0.253 | 9.73E-19 | 13_Proliferating_TCells |
| NASP | 1.16067408 | 0.56 | 0.249 | 1.25E-18 | 13_Proliferating_TCells |
| HNRNPUL2 | 1.109820703 | 0.391 | 0.136 | 1.82E-18 | 13_Proliferating_TCells |
| SF3A2 | 1.004738012 | 0.507 | 0.208 | 3.79E-18 | 13_Proliferating_TCells |
| CALM3 | 1.322410594 | 0.604 | 0.309 | 7.44E-18 | 13_Proliferating_TCells |
| CKAP2 | 1.22215307 | 0.304 | 0.094 | 8.29E-18 | 13_Proliferating_TCells |
| DDB2 | 1.377841176 | 0.295 | 0.091 | 1.35E-17 | 13_Proliferating_TCells |
| MAD2L2 | 1.567947553 | 0.3 | 0.096 | 4.62E-17 | 13_Proliferating_TCells |
| AP2S1 | 1.008873229 | 0.473 | 0.193 | 6.52E-17 | 13_Proliferating_TCells |
| LSM5 | 1.155352888 | 0.357 | 0.124 | 8.95E-17 | 13_Proliferating_TCells |
| PA2G4 | 1.027784311 | 0.56 | 0.259 | 2.15E-16 | 13_Proliferating_TCells |
| COPS3 | 1.167739213 | 0.324 | 0.108 | 2.55E-16 | 13_Proliferating_TCells |
| LSM3 | 1.173273533 | 0.348 | 0.123 | 4.60E-16 | 13_Proliferating_TCells |
| RANGAP1 | 1.111774111 | 0.314 | 0.103 | 4.60E-16 | 13_Proliferating_TCells |
| KPNA2 | 1.274055085 | 0.386 | 0.146 | 5.33E-16 | 13_Proliferating_TCells |
| HSPB11 | 1.230576795 | 0.362 | 0.136 | 9.82E-16 | 13_Proliferating_TCells |
| PSIP1 | 1.038366588 | 0.464 | 0.195 | 1.14E-15 | 13_Proliferating_TCells |
| ARL6IP6 | 1.375781005 | 0.314 | 0.11 | 7.30E-15 | 13_Proliferating_TCells |
| TAF15 | 1.000142792 | 0.628 | 0.316 | 1.13E-14 | 13_Proliferating_TCells |
| SIVA1 | 1.121740107 | 0.498 | 0.231 | 2.14E-14 | 13_Proliferating_TCells |
| CNTRL | 1.110569806 | 0.411 | 0.17 | 3.66E-14 | 13_Proliferating_TCells |
| DAZAP1 | 1.02371937 | 0.401 | 0.157 | 4.07E-14 | 13_Proliferating_TCells |
| RPA3 | 1.301972547 | 0.333 | 0.125 | 4.36E-14 | 13_Proliferating_TCells |
| PPP6R1 | 1.037247011 | 0.391 | 0.157 | 1.18E-13 | 13_Proliferating_TCells |
| CTCF | 1.236408664 | 0.319 | 0.116 | 1.62E-13 | 13_Proliferating_TCells |
| SLF1 | 1.129593062 | 0.324 | 0.117 | 1.70E-13 | 13_Proliferating_TCells |
| DLEU2 | 1.028251425 | 0.522 | 0.25 | 3.61E-13 | 13_Proliferating_TCells |
| PRDX3 | 1.079820439 | 0.343 | 0.133 | 3.07E-12 | 13_Proliferating_TCells |
| LSM4 | 1.171052034 | 0.362 | 0.149 | 4.53E-12 | 13_Proliferating_TCells |
| XPO1 | 1.259579741 | 0.338 | 0.134 | 5.66E-12 | 13_Proliferating_TCells |
| PSMC3 | 1.139570239 | 0.329 | 0.126 | 5.75E-12 | 13_Proliferating_TCells |
| CCT5 | 1.029723591 | 0.377 | 0.157 | 1.48E-11 | 13_Proliferating_TCells |
| FAF1 | 1.140544136 | 0.478 | 0.24 | 1.15E-10 | 13_Proliferating_TCells |
| TUBA1C | 1.138801964 | 0.391 | 0.188 | 2.08E-09 | 13_Proliferating_TCells |
| ARL6IP1 | 1.207520929 | 0.638 | 0.401 | 5.50E-08 | 13_Proliferating_TCells |
