## Supplementary Table 7 for "MAIT cells have a negative impact on glioblastoma"

| Supplementary Table 7: Signature genes | |
|---|---|
| **NDsig genes** | **MDSCsig genes** |
| *A1BG* | *ADIPOR1* |
| *ABCA13* | *ALOX5AP* |
| *ACAA1* | *ARG2* |
| *ACLY* | *ASPRV1* |
| *ACP3* | *ATG3* |
| *ACTR10* | *ATP11B* |
| *ACTR1B* | *ATP6V1G1* |
| *ACTR2* | *BTG1* |
| *ADA2* | *C5AR1* |
| *ADAM10* | *CCR1* |
| *ADAM8* | *CD14* |
| *ADGRE3* | *CD33* |
| *ADGRE5* | *CD84* |
| *ADGRG3* | *CDK2AP2* |
| *AGA* | *CDKN2D* |
| *AGL* | *CLEC4D* |
| *AGPAT2* | *CLEC4E* |
| *AHSG* | *CSF3R* |
| *ALAD* | *CTSD* |
| *ALDH3B1* | *CXCL2* |
| *ALDOA* | *CXCR2* |
| *ALDOC* | *DUSP1* |
| *ALOX5* | *EIF4EBP1* |
| *AMPD3* | *FABP5* |
| *ANO6* | *FBXL5* |
| *ANPEP* | *FGL2* |
| *ANXA2* | *GCNT2* |
| *AOC1* | *GDA* |
| *AP1M1* | *GLIPR2* |
| *AP2A2* | *GPCPD1* |
| *APAF1* | *GRINA* |
| *APEH* | *GSR* |
| *APRT* | *HDC* |
| *ARG1* | *HP* |
| *ARHGAP45* | *IER2* |
| *ARHGAP9* | *IER3* |
| *ARL8A* | *IFITM1* |
| *ARMC8* | *IFITM2* |
| *ARPC5* | *IGFBP6* |
| *ARSA* | *IL1B* |
| *ARSB* | *JUNB* |
| *ASAH1* | *LITAF* |
| *ATAD3B* | *LMNB1* |
| *ATG7* | *LRG1* |

| | |
|---|---|
| *ATP11A* | *MAP1LC3B* |
| *ATP11B* | *MSRB1* |
| *ATP6AP2* | *MTUS1* |
| *ATP6V0A1* | *MXD1* |
| *ATP6V0C* | *MYD88* |
| *ATP6V1D* | *NPL* |
| *ATP8A1* | *OSM* |
| *ATP8B4* | *PICALM* |
| *AZU1* | *PLA2G7* |
| *B2M* | *PPT1* |
| *B4GALT1* | *PROK2* |
| *BIN2* | *RGS3* |
| *BPI* | *RND1* |
| *BRI3* | *RNF149* |
| *BST1* | *S100A11* |
| *BST2* | *S100A6* |
| *C1orf35* | *SELL* |
| *C3* | *SELPLG* |
| *C3AR1* | *SEPHS2* |
| *C5AR1* | *SFXN5* |
| *C6orf120* | *SKAP2* |
| *CAB39* | *SLC40A1* |
| *CALML5* | *SNAP23* |
| *CAMP* | *SOCS3* |
| *CAND1* | *SRGN* |
| *CANT1* | *STEAP4* |
| *CAP1* | *STK17B* |
| *CAPN1* | *TACSTD2* |
| *CAT* | *TALDO1* |
| *CCT2* | *TARM1* |
| *CCT8* | *TPD52* |
| *CD14* | *TSPO* |
| *CD177* | *UBB* |
| *CD300A* | *UPP1* |
| *CD33* | *YPEL3* |
| *CD36* | *ZYX* |
| *CD44* | |
| *CD47* | |
| *CD53* | |
| *CD55* | |
| *CD58* | |
| *CD59* | |
| *CD63* | |
| *CD68* | |
| *CD93* | |
| *CDA* | |

| |
|---|
| *CDK13* |
| *CEACAM1* |
| *CEACAM3* |
| *CEACAM6* |
| *CEACAM8* |
| *CEP290* |
| *CFD* |
| *CFP* |
| *CHI3L1* |
| *CHIT1* |
| *CHRNB4* |
| *CKAP4* |
| *CLEC12A* |
| *CLEC4C* |
| *CLEC4D* |
| *CLEC5A* |
| *CMTM6* |
| *CNN2* |
| *COMMD3* |
| *COMMD9* |
| *COPB1* |
| *COTL1* |
| *CPNE1* |
| *CPNE3* |
| *CPPED1* |
| *CR1* |
| *CRACR2A* |
| *CREG1* |
| *CRISP3* |
| *CRISPLD2* |
| *CSNK2B* |
| *CST3* |
| *CSTB* |
| *CTSA* |
| *CTSB* |
| *CTSC* |
| *CTSD* |
| *CTSG* |
| *CTSH* |
| *CTSS* |
| *CTSZ* |
| *CXCL1* |
| *CXCR1* |
| *CXCR2* |
| *CYB5R3* |
| *CYBA* |

| |
|---|
| *CYBB* |
| *CYFIP1* |
| *CYSTM1* |
| *DBNL* |
| *DDOST* |
| *DDX3X* |
| *DEFA1* |
| *DEFA1B* |
| *DEFA4* |
| *DEGS1* |
| *DERA* |
| *DGAT1* |
| *DIAPH1* |
| *DNAJC13* |
| *DNAJC3* |
| *DNAJC5* |
| *DNASE1L1* |
| *DOCK2* |
| *DOK3* |
| *DPP7* |
| *DSC1* |
| *DSG1* |
| *DSN1* |
| *DSP* |
| *DYNC1H1* |
| *DYNC1LI1* |
| *DYNLL1* |
| *DYNLT1* |
| *EEF1A1* |
| *EEF2* |
| *ELANE* |
| *ENPP4* |
| *EPX* |
| *ERP44* |
| *FABP5* |
| *FAF2* |
| *FCAR* |
| *FCER1G* |
| *FCGR2A* |
| *FCGR3B* |
| *FCN1* |
| *FGL2* |
| *FGR* |
| *FLG2* |
| *FOLR3* |
| *FPR1* |

| |
|---|
| *FPR2* |
| *FRK* |
| *FRMPD3* |
| *FTH1* |
| *FTL* |
| *FUCA1* |
| *FUCA2* |
| *GAA* |
| *GALNS* |
| *GCA* |
| *GDI2* |
| *GGH* |
| *GHDC* |
| *GLA* |
| *GLB1* |
| *GLIPR1* |
| *GM2A* |
| *GMFG* |
| *GNS* |
| *GOLGA7* |
| *GPI* |
| *GPR84* |
| *GRN* |
| *GSDMD* |
| *GSN* |
| *GSTP1* |
| *GUSB* |
| *GYG1* |
| *HBB* |
| *HEBP2* |
| *HEXB* |
| *HGSNAT* |
| *HK3* |
| *HLA-A* |
| *HLA-B* |
| *HLA-C* |
| *HMGB1* |
| *HMOX2* |
| *HP* |
| *HPSE* |
| *HRNR* |
| *HSP90AA1* |
| *HSP90AB1* |
| *HSPA1A* |
| *HSPA1B* |
| *HSPA6* |

| |
|---|
| *HSPA8* |
| *HUWE1* |
| *HVCN1* |
| *IDH1* |
| *IGF2R* |
| *ILF2* |
| *IMPDH1* |
| *IMPDH2* |
| *IQGAP1* |
| *IQGAP2* |
| *IRAG2* |
| *IST1* |
| *ITGAL* |
| *ITGAM* |
| *ITGAV* |
| *ITGAX* |
| *ITGB2* |
| *JUP* |
| *KCMF1* |
| *KCNAB2* |
| *KPNB1* |
| *KRT1* |
| *LAIR1* |
| *LAMP1* |
| *LAMP2* |
| *LAMTOR1* |
| *LAMTOR2* |
| *LAMTOR3* |
| *LCN2* |
| *LGALS3* |
| *LILRA3* |
| *LILRB2* |
| *LILRB3* |
| *LPCAT1* |
| *LRG1* |
| *LRRC7* |
| *LTA4H* |
| *LTF* |
| *LYZ* |
| *MAGT1* |
| *MAN2B1* |
| *MANBA* |
| *MAPK1* |
| *MAPK14* |
| *MCEMP1* |
| *METTL7A* |

| |
|---|
| *MGAM* |
| *MGST1* |
| *MIF* |
| *MLEC* |
| *MME* |
| *MMP25* |
| *MMP8* |
| *MMP9* |
| *MNDA* |
| *MOSPD2* |
| *MPO* |
| *MS4A3* |
| *MVP* |
| *NAPRT* |
| *NBEAL2* |
| *NCKAP1L* |
| *NCSTN* |
| *NDUFC2* |
| *NEU1* |
| *NFAM1* |
| *NFASC* |
| *NFKB1* |
| *NHLRC3* |
| *NIT2* |
| *NME2* |
| *NPC2* |
| *NRAS* |
| *OLFM4* |
| *OLR1* |
| *ORM1* |
| *ORM2* |
| *ORMDL3* |
| *OSCAR* |
| *OSTF1* |
| *P2RX1* |
| *PA2G4* |
| *PADI2* |
| *PAFAH1B2* |
| *PDAP1* |
| *PDXK* |
| *PECAM1* |
| *PFKL* |
| *PGAM1* |
| *PGLYRP1* |
| *PGM1* |
| *PGM2* |

| |
|---|
| *PGRMC1* |
| *PIGR* |
| *PKM* |
| *PKP1* |
| *PLAC8* |
| *PLAU* |
| *PLAUR* |
| *PLD1* |
| *PLEKHO2* |
| *PNP* |
| *PPBP* |
| *PPIA* |
| *PPIE* |
| *PRCP* |
| *PRDX4* |
| *PRDX6* |
| *PRG2* |
| *PRG3* |
| *PRKCD* |
| *PRSS2* |
| *PRSS3* |
| *PRTN3* |
| *PSAP* |
| *PSEN1* |
| *PSMA2* |
| *PSMA5* |
| *PSMB1* |
| *PSMB7* |
| *PSMC2* |
| *PSMC3* |
| *PSMD1* |
| *PSMD11* |
| *PSMD12* |
| *PSMD13* |
| *PSMD14* |
| *PSMD2* |
| *PSMD3* |
| *PSMD6* |
| *PSMD7* |
| *PTAFR* |
| *PTGES2* |
| *PTPN6* |
| *PTPRB* |
| *PTPRC* |
| *PTPRJ* |
| *PTPRN2* |

| |
|---|
| *PTX3* |
| *PYCARD* |
| *PYGB* |
| *PYGL* |
| *QPCT* |
| *QSOX1* |
| *RAB10* |
| *RAB14* |
| *RAB18* |
| *RAB24* |
| *RAB27A* |
| *RAB31* |
| *RAB37* |
| *RAB3A* |
| *RAB3D* |
| *RAB44* |
| *RAB4B* |
| *RAB5B* |
| *RAB5C* |
| *RAB6A* |
| *RAB7A* |
| *RAB9B* |
| *RAC1* |
| *RAP1A* |
| *RAP1B* |
| *RAP2B* |
| *RAP2C* |
| *RETN* |
| *RHOA* |
| *RHOF* |
| *RHOG* |
| *RNASE2* |
| *RNASE3* |
| *RNASET2* |
| *ROCK1* |
| *S100A11* |
| *S100A12* |
| *S100A7* |
| *S100A8* |
| *S100A9* |
| *S100P* |
| *SCAMP1* |
| *SDCBP* |
| *SELL* |
| *SERPINA1* |
| *SERPINA3* |

| |
|---|
| *SERPINB1* |
| *SERPINB10* |
| *SERPINB12* |
| *SERPINB3* |
| *SERPINB6* |
| *SIGLEC14* |
| *SIGLEC5* |
| *SIGLEC9* |
| *SIRPA* |
| *SIRPB1* |
| *SLC11A1* |
| *SLC15A4* |
| *SLC27A2* |
| *SLC2A3* |
| *SLC2A5* |
| *SLC44A2* |
| *SLCO4C1* |
| *SLPI* |
| *SNAP23* |
| *SNAP25* |
| *SNAP29* |
| *SPTAN1* |
| *SRP14* |
| *STBD1* |
| *STING1* |
| *STK10* |
| *STK11IP* |
| *STOM* |
| *SURF4* |
| *SVIP* |
| *SYNGR1* |
| *TARM1* |
| *TBC1D10C* |
| *TCIRG1* |
| *TCN1* |
| *TICAM2* |
| *TIMP2* |
| *TLR2* |
| *TMBIM1* |
| *TMC6* |
| *TMEM179B* |
| *TMEM30A* |
| *TMEM63A* |
| *TNFAIP6* |
| *TNFRSF1B* |
| *TOLLIP* |

| |
|---|
| *TOM1* |
| *TRAPPC1* |
| *TRPM2* |
| *TSPAN14* |
| *TTR* |
| *TUBB* |
| *TUBB4B* |
| *TXNDC5* |
| *TYROBP* |
| *UBR4* |
| *UNC13D* |
| *VAMP8* |
| *VAPA* |
| *VAT1* |
| *VCL* |
| *VCP* |
| *VNN1* |
| *VPS35L* |
| *XRCC5* |
| *XRCC6* |
| *YPEL5* |
