## Supplementary Table 8 for "MAIT cells have a negative impact on glioblastoma"

| Supplementary Table 8. Seurat based clustering of the Myeloid population in the scRNA-seq analysis | | | | | |
|---|---|---|---|---|---|
| **Genes** | **avg_log2FC** | **pct.1** | **pct.2** | **p_val_adj** | **Clusters** |
| AIF1 | -1.05987108 | 0.362 | 0.67 | 0 | 02_Inflam-TAM |
| SH3BGRL3 | -1.089550498 | 0.397 | 0.696 | 0 | 02_Inflam-TAM |
| UBB | -1.588373229 | 0.28 | 0.579 | 0 | 02_Inflam-TAM |
| NPC2 | -1.068596313 | 0.505 | 0.8 | 0 | 02_Inflam-TAM |
| COX6A1 | -1.183971782 | 0.28 | 0.566 | 0 | 02_Inflam-TAM |
| PFN1 | -1.049591336 | 0.497 | 0.776 | 0 | 02_Inflam-TAM |
| CST3 | -1.102690893 | 0.587 | 0.863 | 0 | 02_Inflam-TAM |
| SERF2 | -1.018941415 | 0.525 | 0.797 | 0 | 02_Inflam-TAM |
| DNAJB1 | -2.602537074 | 0.279 | 0.551 | 0 | 02_Inflam-TAM |
| MIF | -1.345074482 | 0.321 | 0.593 | 0 | 02_Inflam-TAM |
| ATP6V0C | -1.045817063 | 0.415 | 0.686 | 0 | 02_Inflam-TAM |
| COX6B1 | -1.1902319 | 0.265 | 0.536 | 0 | 02_Inflam-TAM |
| CFL1 | -1.023243882 | 0.51 | 0.774 | 0 | 02_Inflam-TAM |
| PPIB | -1.198074791 | 0.337 | 0.597 | 0 | 02_Inflam-TAM |
| SRP14 | -1.051960255 | 0.294 | 0.552 | 0 | 02_Inflam-TAM |
| IFITM3 | -1.280703235 | 0.283 | 0.538 | 0 | 02_Inflam-TAM |
| S100A11 | -1.308748417 | 0.569 | 0.824 | 0 | 02_Inflam-TAM |
| HSPA1A | -2.692059368 | 0.4 | 0.653 | 0 | 02_Inflam-TAM |
| MB21D2 | 1.623904982 | 0.529 | 0.281 | 0 | 02_Inflam-TAM |
| HSPA1B | -2.50555479 | 0.319 | 0.561 | 0 | 02_Inflam-TAM |
| TREM2 | -1.149954122 | 0.391 | 0.633 | 0 | 02_Inflam-TAM |
| LGALS1 | -1.410987131 | 0.308 | 0.549 | 0 | 02_Inflam-TAM |
| MARCKS | -1.296061708 | 0.405 | 0.646 | 0 | 02_Inflam-TAM |
| HSPB1 | -2.560766783 | 0.283 | 0.522 | 0 | 02_Inflam-TAM |
| HSPA8 | -1.34473637 | 0.398 | 0.636 | 0 | 02_Inflam-TAM |
| TUBA1B | -1.016882964 | 0.434 | 0.67 | 0 | 02_Inflam-TAM |
| CD63 | -1.093177287 | 0.479 | 0.712 | 0 | 02_Inflam-TAM |
| TMSB10 | -1.148554992 | 0.701 | 0.921 | 0 | 02_Inflam-TAM |
| UBASH3B | 1.480622934 | 0.618 | 0.399 | 0 | 02_Inflam-TAM |
| HSP90AB1 | -1.252861743 | 0.58 | 0.795 | 0 | 02_Inflam-TAM |
| C1QB | -1.224975771 | 0.602 | 0.813 | 0 | 02_Inflam-TAM |
| C1QA | -1.140862723 | 0.598 | 0.807 | 0 | 02_Inflam-TAM |
| HLA-DRB5 | -1.207331753 | 0.563 | 0.771 | 0 | 02_Inflam-TAM |
| TYROBP | -1.072980184 | 0.671 | 0.879 | 0 | 02_Inflam-TAM |
| HSP90AA1 | -1.880249457 | 0.637 | 0.843 | 0 | 02_Inflam-TAM |
| SIK3 | 1.247726836 | 0.794 | 0.591 | 0 | 02_Inflam-TAM |
| MALT1 | 1.244845014 | 0.616 | 0.414 | 0 | 02_Inflam-TAM |
| COX7C | -1.006838143 | 0.275 | 0.543 | 4.48E-303 | 02_Inflam-TAM |
| HSPE1 | -1.817962763 | 0.231 | 0.454 | 5.64E-282 | 02_Inflam-TAM |
| ATP6V1F | -1.000014079 | 0.326 | 0.565 | 1.91E-281 | 02_Inflam-TAM |
| PRDX1 | -1.085207599 | 0.265 | 0.503 | 5.10E-271 | 02_Inflam-TAM |
| TMA7 | -1.059755267 | 0.268 | 0.51 | 8.51E-271 | 02_Inflam-TAM |
| KRTCAP2 | -1.218779531 | 0.203 | 0.435 | 2.23E-270 | 02_Inflam-TAM |

| | | | | | |
|---|---|---|---|---|---|
| HCST | -1.197565369 | 0.231 | 0.466 | 3.93E-269 | 02_Inflam-TAM |
| CSTB | -1.543741699 | 0.294 | 0.523 | 9.47E-269 | 02_Inflam-TAM |
| EGR1 | -1.517348207 | 0.303 | 0.514 | 7.94E-266 | 02_Inflam-TAM |
| COX6C | -1.20327252 | 0.229 | 0.461 | 2.18E-265 | 02_Inflam-TAM |
| SSR4 | -1.18897938 | 0.236 | 0.468 | 5.04E-263 | 02_Inflam-TAM |
| COX5B | -1.1022252 | 0.248 | 0.481 | 2.45E-262 | 02_Inflam-TAM |
| CD14 | -1.085611493 | 0.433 | 0.638 | 4.61E-257 | 02_Inflam-TAM |
| MYL12A | -1.120908371 | 0.242 | 0.471 | 5.16E-256 | 02_Inflam-TAM |
| TXNIP | -1.423636834 | 0.237 | 0.456 | 1.76E-255 | 02_Inflam-TAM |
| HMOX1 | -1.545218316 | 0.383 | 0.584 | 2.74E-247 | 02_Inflam-TAM |
| VAMP8 | -1.002172612 | 0.254 | 0.486 | 2.19E-246 | 02_Inflam-TAM |
| UBL5 | -1.156911108 | 0.228 | 0.449 | 4.56E-246 | 02_Inflam-TAM |
| FCGR3A | -1.062655085 | 0.346 | 0.549 | 2.26E-226 | 02_Inflam-TAM |
| OST4 | -1.07399137 | 0.221 | 0.435 | 1.25E-221 | 02_Inflam-TAM |
| S100A4 | -1.307900107 | 0.203 | 0.408 | 1.03E-218 | 02_Inflam-TAM |
| UQCR10 | -1.036477752 | 0.211 | 0.418 | 1.73E-212 | 02_Inflam-TAM |
| UQCR11 | -1.01741629 | 0.235 | 0.44 | 2.59E-204 | 02_Inflam-TAM |
| CCDC200 | 3.394188407 | 0.674 | 0.211 | 0 | 03_Classical-TIM |
| CROCC | 3.209572457 | 0.595 | 0.163 | 0 | 03_Classical-TIM |
| WDR74 | 2.753958307 | 0.619 | 0.19 | 0 | 03_Classical-TIM |
| TEX14 | 2.876707445 | 0.895 | 0.472 | 0 | 03_Classical-TIM |
| LINC00910 | 2.758837842 | 0.651 | 0.241 | 0 | 03_Classical-TIM |
| AC253572.2 | 2.811348773 | 0.561 | 0.169 | 0 | 03_Classical-TIM |
| AC245014.3 | 2.654042881 | 0.564 | 0.183 | 0 | 03_Classical-TIM |
| AC012447.1 | 3.223557789 | 0.451 | 0.096 | 0 | 03_Classical-TIM |
| AC084871.1 | 2.823804825 | 0.52 | 0.177 | 0 | 03_Classical-TIM |
| AL021155.5 | 3.444922437 | 0.392 | 0.064 | 0 | 03_Classical-TIM |
| AC007952.4 | 2.629175474 | 0.466 | 0.152 | 0 | 03_Classical-TIM |
| TMEM107 | 2.620106662 | 0.45 | 0.138 | 0 | 03_Classical-TIM |
| EGR1 | 1.622858943 | 0.746 | 0.444 | 0 | 03_Classical-TIM |
| C12orf57 | 2.392477835 | 0.507 | 0.209 | 0 | 03_Classical-TIM |
| Z93241.1 | 2.932041799 | 0.345 | 0.078 | 0 | 03_Classical-TIM |
| AL691403.1 | 1.732959622 | 0.464 | 0.208 | 0 | 03_Classical-TIM |
| EIF4A3 | 1.940474661 | 0.614 | 0.359 | 0 | 03_Classical-TIM |
| AC018754.1 | 2.801538903 | 0.307 | 0.068 | 0 | 03_Classical-TIM |
| AC022217.3 | 1.205084632 | 0.512 | 0.277 | 0 | 03_Classical-TIM |
| AC020916.1 | 1.109425048 | 0.811 | 0.578 | 0 | 03_Classical-TIM |
| AL390957.1 | 1.442965391 | 0.461 | 0.228 | 0 | 03_Classical-TIM |
| AC103591.3 | 2.72768588 | 0.306 | 0.076 | 0 | 03_Classical-TIM |
| AL136987.1 | 1.636851503 | 0.416 | 0.194 | 0 | 03_Classical-TIM |
| C9orf72 | 1.277975615 | 0.742 | 0.52 | 0 | 03_Classical-TIM |
| ATF3 | 1.2071084 | 0.815 | 0.599 | 0 | 03_Classical-TIM |
| ATP2B1-AS1 | 1.433477946 | 0.467 | 0.252 | 0 | 03_Classical-TIM |
| MYLIP | 1.384567248 | 0.463 | 0.25 | 1.27E-299 | 03_Classical-TIM |
| KDM6B | -1.728859282 | 0.252 | 0.637 | 0 | 04_HSP-Macrophage |
| TNFAIP3 | -1.426762329 | 0.354 | 0.711 | 0 | 04_HSP-Macrophage |

| | | | | | |
|---|---|---|---|---|---|
| PLAUR | -1.807459469 | 0.42 | 0.764 | 0 | 04_HSP-Macrophage |
| NR4A2 | -1.011633707 | 0.459 | 0.786 | 0 | 04_HSP-Macrophage |
| NLRP3 | -1.647887741 | 0.304 | 0.613 | 0 | 04_HSP-Macrophage |
| GRASP | -1.308963699 | 0.32 | 0.628 | 0 | 04_HSP-Macrophage |
| GPR183 | -1.439825401 | 0.265 | 0.571 | 0 | 04_HSP-Macrophage |
| IL1B | -2.17133847 | 0.276 | 0.581 | 0 | 04_HSP-Macrophage |
| HSPA1B | 1.596285899 | 0.793 | 0.49 | 0 | 04_HSP-Macrophage |
| LMNA | -2.070054244 | 0.137 | 0.44 | 0 | 04_HSP-Macrophage |
| NFKB1 | -1.386781096 | 0.407 | 0.709 | 0 | 04_HSP-Macrophage |
| AC020916.1 | -1.059618232 | 0.343 | 0.643 | 0 | 04_HSP-Macrophage |
| SYTL3 | -1.285042425 | 0.331 | 0.629 | 0 | 04_HSP-Macrophage |
| NFKBID | -1.168519533 | 0.278 | 0.573 | 0 | 04_HSP-Macrophage |
| KLF4 | -1.254962663 | 0.254 | 0.546 | 0 | 04_HSP-Macrophage |
| C5AR1 | -1.303960425 | 0.377 | 0.669 | 0 | 04_HSP-Macrophage |
| CDKN1A | -1.153118912 | 0.313 | 0.598 | 0 | 04_HSP-Macrophage |
| VEGFA | -1.924328513 | 0.136 | 0.421 | 0 | 04_HSP-Macrophage |
| RGCC | -1.476960462 | 0.217 | 0.5 | 0 | 04_HSP-Macrophage |
| CXCR4 | -1.4357392 | 0.401 | 0.669 | 0 | 04_HSP-Macrophage |
| NFKBIA | -1.562954312 | 0.59 | 0.855 | 0 | 04_HSP-Macrophage |
| OLR1 | -1.257616965 | 0.578 | 0.833 | 0 | 04_HSP-Macrophage |
| ATP1B3 | -1.234244269 | 0.544 | 0.796 | 0 | 04_HSP-Macrophage |
| HSPA1A | 1.86358805 | 0.835 | 0.586 | 0 | 04_HSP-Macrophage |
| CEBPB | -1.268981076 | 0.427 | 0.67 | 0 | 04_HSP-Macrophage |
| HSPH1 | 2.310002527 | 0.684 | 0.443 | 0 | 04_HSP-Macrophage |
| CCL3 | -1.565536965 | 0.517 | 0.758 | 0 | 04_HSP-Macrophage |
| MAP3K8 | -1.096267698 | 0.534 | 0.774 | 0 | 04_HSP-Macrophage |
| BTG1 | -1.195022645 | 0.532 | 0.762 | 0 | 04_HSP-Macrophage |
| 1-Mar | 1.513102622 | 0.5 | 0.273 | 0 | 04_HSP-Macrophage |
| BAG3 | 2.164566658 | 0.462 | 0.238 | 0 | 04_HSP-Macrophage |
| DNAJB1 | 1.432841472 | 0.699 | 0.486 | 0 | 04_HSP-Macrophage |
| HSPB1 | 1.647014792 | 0.67 | 0.463 | 0 | 04_HSP-Macrophage |
| ZFAND2A | 1.959825164 | 0.375 | 0.175 | 0 | 04_HSP-Macrophage |
| TFRC | -1.418830659 | 0.239 | 0.516 | 2.55E-302 | 04_HSP-Macrophage |
| SERPINB9 | -1.026993446 | 0.498 | 0.724 | 7.92E-295 | 04_HSP-Macrophage |
| GK | -1.106450955 | 0.35 | 0.621 | 1.65E-291 | 04_HSP-Macrophage |
| HBEGF | -1.724983532 | 0.156 | 0.418 | 2.24E-288 | 04_HSP-Macrophage |
| RCSD1 | 1.287602375 | 0.565 | 0.362 | 6.40E-287 | 04_HSP-Macrophage |
| CTNNB1 | -1.034108373 | 0.429 | 0.674 | 3.92E-271 | 04_HSP-Macrophage |
| ID2 | -1.01909779 | 0.44 | 0.67 | 5.36E-269 | 04_HSP-Macrophage |
| BCL2A1 | -1.541853972 | 0.214 | 0.474 | 8.58E-268 | 04_HSP-Macrophage |
| ATM | 1.257516101 | 0.52 | 0.32 | 1.88E-265 | 04_HSP-Macrophage |
| IRAK2 | -1.305781758 | 0.22 | 0.482 | 1.27E-259 | 04_HSP-Macrophage |
| KLF2 | -1.551402127 | 0.37 | 0.592 | 3.72E-258 | 04_HSP-Macrophage |
| ICAM1 | -1.610514976 | 0.19 | 0.439 | 2.75E-257 | 04_HSP-Macrophage |
| IER3 | -1.132940237 | 0.447 | 0.669 | 2.57E-249 | 04_HSP-Macrophage |
| CD55 | -1.305663277 | 0.238 | 0.49 | 1.00E-248 | 04_HSP-Macrophage |

| | | | | | |
|---|---|---|---|---|---|
| MMP19 | -2.236481669 | 0.082 | 0.301 | 8.45E-239 | 04_HSP-Macrophage |
| RABGEF1 | -1.030313789 | 0.354 | 0.601 | 7.57E-238 | 04_HSP-Macrophage |
| ITGAX | -1.228346622 | 0.367 | 0.591 | 1.21E-223 | 04_HSP-Macrophage |
| DUSP2 | -1.798871509 | 0.12 | 0.339 | 1.43E-219 | 04_HSP-Macrophage |
| CSRNP1 | -1.259255495 | 0.173 | 0.408 | 6.70E-219 | 04_HSP-Macrophage |
| CCL3L1 | -1.809717708 | 0.239 | 0.46 | 8.24E-218 | 04_HSP-Macrophage |
| CD69 | -1.249739382 | 0.216 | 0.448 | 8.01E-214 | 04_HSP-Macrophage |
| MYADM | -1.501772512 | 0.154 | 0.374 | 7.09E-211 | 04_HSP-Macrophage |
| SLC31A2 | -1.091849549 | 0.267 | 0.494 | 1.34E-208 | 04_HSP-Macrophage |
| STX11 | -1.195701604 | 0.191 | 0.42 | 1.30E-207 | 04_HSP-Macrophage |
| XBP1 | -1.127719295 | 0.229 | 0.457 | 2.50E-207 | 04_HSP-Macrophage |
| SOCS3 | -1.502290806 | 0.125 | 0.339 | 6.78E-205 | 04_HSP-Macrophage |
| RFX2 | -1.199498304 | 0.195 | 0.421 | 2.51E-204 | 04_HSP-Macrophage |
| METRNL | -1.317289808 | 0.198 | 0.416 | 1.66E-203 | 04_HSP-Macrophage |
| PLK3 | -1.038577149 | 0.259 | 0.491 | 3.32E-203 | 04_HSP-Macrophage |
| TUBB4B | -1.356298526 | 0.167 | 0.381 | 1.44E-197 | 04_HSP-Macrophage |
| CHMP1B | -1.403753171 | 0.186 | 0.398 | 5.92E-194 | 04_HSP-Macrophage |
| MALT1 | -1.170076915 | 0.253 | 0.469 | 3.36E-189 | 04_HSP-Macrophage |
| TGIF1 | -1.357193582 | 0.145 | 0.351 | 4.47E-189 | 04_HSP-Macrophage |
| PLEKHG2 | -1.117446452 | 0.17 | 0.39 | 6.08E-189 | 04_HSP-Macrophage |
| FOSL2 | -1.000339954 | 0.203 | 0.423 | 1.44E-185 | 04_HSP-Macrophage |
| BHLHE40 | -1.104426887 | 0.183 | 0.386 | 1.20E-168 | 04_HSP-Macrophage |
| TMIGD3 | 1.405062332 | 0.689 | 0.341 | 0 | 05_Microgia |
| P2RY12 | 1.58503124 | 0.574 | 0.235 | 0 | 05_Microgia |
| SLC2A5 | 1.026214581 | 0.71 | 0.39 | 0 | 05_Microgia |
| APOC2 | 1.424241834 | 0.708 | 0.408 | 0 | 05_Microgia |
| SYNDIG1 | 1.059238253 | 0.498 | 0.198 | 0 | 05_Microgia |
| ADGRG1 | 1.347065125 | 0.5 | 0.209 | 0 | 05_Microgia |
| CH25H | 1.298953251 | 0.588 | 0.299 | 0 | 05_Microgia |
| TMEM119 | 1.495066499 | 0.427 | 0.161 | 0 | 05_Microgia |
| FCGR1A | 1.015632724 | 0.682 | 0.434 | 0 | 05_Microgia |
| KLF2 | 1.035754443 | 0.78 | 0.538 | 0 | 05_Microgia |
| CX3CR1 | 1.346392834 | 0.433 | 0.201 | 0 | 05_Microgia |
| MCF2L | 1.123565862 | 0.408 | 0.177 | 0 | 05_Microgia |
| NAV3 | 1.357994988 | 0.322 | 0.096 | 0 | 05_Microgia |
| LINC01736 | 1.456801661 | 0.352 | 0.14 | 0 | 05_Microgia |
| SLC16A10 | -2.406745565 | 0.162 | 0.425 | 3.09E-301 | 05_Microgia |
| AREG | -3.421195897 | 0.094 | 0.335 | 1.38E-289 | 05_Microgia |
| SELPLG | 1.17719344 | 0.495 | 0.265 | 1.50E-285 | 05_Microgia |
| OLFML3 | 1.045566762 | 0.496 | 0.271 | 3.82E-261 | 05_Microgia |
| RGS16 | 1.197662295 | 0.385 | 0.183 | 4.77E-247 | 05_Microgia |
| CD44 | -1.791503536 | 0.261 | 0.48 | 1.08E-223 | 05_Microgia |
| TGFBI | -2.112851373 | 0.138 | 0.352 | 2.92E-220 | 05_Microgia |
| IQGAP2 | -1.72341998 | 0.171 | 0.378 | 2.59E-197 | 05_Microgia |
| MCTP1 | -1.693854627 | 0.18 | 0.388 | 4.14E-192 | 05_Microgia |
| S100A10 | 2.158284725 | 0.746 | 0.359 | 0 | 06_VCAN-TIM |

| | | | | | |
|---|---|---|---|---|---|
| GPNMB | 2.000790659 | 0.657 | 0.276 | 0 | 06_VCAN-TIM |
| VCAN | 2.816734619 | 0.464 | 0.097 | 0 | 06_VCAN-TIM |
| CSGALNACT1 | -2.149883666 | 0.21 | 0.567 | 0 | 06_VCAN-TIM |
| LGALS3 | 2.137083933 | 0.621 | 0.266 | 0 | 06_VCAN-TIM |
| S100A6 | 1.819550741 | 0.676 | 0.33 | 0 | 06_VCAN-TIM |
| LGALS1 | 1.996673381 | 0.816 | 0.479 | 0 | 06_VCAN-TIM |
| ANXA2 | 1.485529295 | 0.668 | 0.351 | 0 | 06_VCAN-TIM |
| CSTB | 2.386080235 | 0.773 | 0.458 | 0 | 06_VCAN-TIM |
| CD109 | 1.821566584 | 0.439 | 0.139 | 0 | 06_VCAN-TIM |
| ADM | 2.257646349 | 0.432 | 0.133 | 0 | 06_VCAN-TIM |
| PALD1 | -1.680318093 | 0.248 | 0.544 | 0 | 06_VCAN-TIM |
| PLIN2 | 1.54359486 | 0.788 | 0.502 | 0 | 06_VCAN-TIM |
| ENO1 | 1.61722469 | 0.72 | 0.439 | 0 | 06_VCAN-TIM |
| LSP1 | 1.361319113 | 0.544 | 0.269 | 0 | 06_VCAN-TIM |
| MIF | 1.973710899 | 0.8 | 0.525 | 0 | 06_VCAN-TIM |
| PKM | 1.331043508 | 0.719 | 0.448 | 0 | 06_VCAN-TIM |
| CD44 | 1.065408467 | 0.697 | 0.428 | 0 | 06_VCAN-TIM |
| FABP5 | 1.946241746 | 0.497 | 0.229 | 0 | 06_VCAN-TIM |
| S100A9 | 2.000115609 | 0.435 | 0.168 | 0 | 06_VCAN-TIM |
| S100A4 | 1.282847317 | 0.617 | 0.351 | 0 | 06_VCAN-TIM |
| GPI | 1.742690438 | 0.504 | 0.238 | 0 | 06_VCAN-TIM |
| VIM | 1.633623838 | 0.908 | 0.646 | 0 | 06_VCAN-TIM |
| TXN | 1.405124516 | 0.527 | 0.267 | 0 | 06_VCAN-TIM |
| TGFBI | 1.47745682 | 0.56 | 0.301 | 0 | 06_VCAN-TIM |
| LUCAT1 | 1.496458072 | 0.468 | 0.209 | 0 | 06_VCAN-TIM |
| EMP3 | 1.207001314 | 0.608 | 0.35 | 0 | 06_VCAN-TIM |
| BNIP3L | 1.36552633 | 0.604 | 0.347 | 0 | 06_VCAN-TIM |
| PPDPF | 1.356737799 | 0.602 | 0.346 | 0 | 06_VCAN-TIM |
| FLNA | 1.560469851 | 0.409 | 0.153 | 0 | 06_VCAN-TIM |
| ANGPTL4 | 3.11616109 | 0.301 | 0.046 | 0 | 06_VCAN-TIM |
| TIMP1 | 1.43098386 | 0.561 | 0.308 | 0 | 06_VCAN-TIM |
| SRGAP2 | -1.419774279 | 0.481 | 0.733 | 0 | 06_VCAN-TIM |
| TNS1 | 2.35568585 | 0.324 | 0.072 | 0 | 06_VCAN-TIM |
| UPP1 | 1.490120346 | 0.47 | 0.219 | 0 | 06_VCAN-TIM |
| ADAM8 | 2.574124368 | 0.325 | 0.075 | 0 | 06_VCAN-TIM |
| ELMO1 | -1.446904457 | 0.598 | 0.845 | 0 | 06_VCAN-TIM |
| RNASE1 | 1.610735882 | 0.568 | 0.321 | 0 | 06_VCAN-TIM |
| GSTO1 | 1.300587988 | 0.529 | 0.29 | 0 | 06_VCAN-TIM |
| BNIP3 | 3.139379671 | 0.283 | 0.044 | 0 | 06_VCAN-TIM |
| NUPR1 | 2.269199332 | 0.428 | 0.19 | 0 | 06_VCAN-TIM |
| MEF2C | -1.294716578 | 0.472 | 0.708 | 0 | 06_VCAN-TIM |
| SNTB1 | 1.973706553 | 0.34 | 0.105 | 0 | 06_VCAN-TIM |
| CTSL | 1.478416468 | 0.676 | 0.445 | 0 | 06_VCAN-TIM |
| PGK1 | 1.285555394 | 0.685 | 0.455 | 0 | 06_VCAN-TIM |
| ERO1A | 2.01583551 | 0.359 | 0.132 | 0 | 06_VCAN-TIM |
| ALDOA | 1.421372846 | 0.633 | 0.406 | 0 | 06_VCAN-TIM |

| | | | | | |
|---|---|---|---|---|---|
| LDHA | 1.288897643 | 0.684 | 0.459 | 0 | 06_VCAN-TIM |
| CXCL3 | 2.528138935 | 0.299 | 0.077 | 0 | 06_VCAN-TIM |
| TPI1 | 1.163343031 | 0.704 | 0.482 | 0 | 06_VCAN-TIM |
| FRMD4A | -1.335204245 | 0.59 | 0.81 | 0 | 06_VCAN-TIM |
| C15orf48 | 1.318975687 | 0.338 | 0.12 | 0 | 06_VCAN-TIM |
| C3 | -1.116986411 | 0.553 | 0.768 | 0 | 06_VCAN-TIM |
| CXCL2 | 1.901536813 | 0.344 | 0.137 | 1.32E-303 | 06_VCAN-TIM |
| ANKRD28 | 1.541792 | 0.356 | 0.143 | 1.65E-298 | 06_VCAN-TIM |
| HK2 | 1.865731259 | 0.442 | 0.218 | 6.16E-296 | 06_VCAN-TIM |
| ZBTB16 | -1.297336759 | 0.366 | 0.63 | 1.02E-279 | 06_VCAN-TIM |
| ANXA1 | 1.146207232 | 0.618 | 0.364 | 1.47E-274 | 06_VCAN-TIM |
| SPTLC2 | -1.446896027 | 0.406 | 0.637 | 2.78E-270 | 06_VCAN-TIM |
| NHSL1 | -1.600303616 | 0.328 | 0.576 | 1.65E-260 | 06_VCAN-TIM |
| GBE1 | 1.504214752 | 0.493 | 0.268 | 4.24E-254 | 06_VCAN-TIM |
| SRGAP2B | -1.462561608 | 0.346 | 0.583 | 7.28E-249 | 06_VCAN-TIM |
| PLD4 | -1.80503986 | 0.146 | 0.392 | 6.25E-236 | 06_VCAN-TIM |
| DLEU1 | -1.270641514 | 0.387 | 0.617 | 2.99E-235 | 06_VCAN-TIM |
| RALA | 1.214310988 | 0.423 | 0.216 | 4.22E-232 | 06_VCAN-TIM |
| PLA2G4A | -1.882443609 | 0.168 | 0.412 | 3.59E-229 | 06_VCAN-TIM |
| ST6GAL1 | -1.476708746 | 0.316 | 0.552 | 1.90E-228 | 06_VCAN-TIM |
| SEC61G | 1.120321954 | 0.461 | 0.261 | 4.63E-209 | 06_VCAN-TIM |
| PADI2 | -1.378572709 | 0.325 | 0.552 | 2.38E-208 | 06_VCAN-TIM |
| EPB41L2 | -1.037008851 | 0.465 | 0.669 | 2.57E-208 | 06_VCAN-TIM |
| LYZ | 1.071797607 | 0.504 | 0.293 | 2.77E-203 | 06_VCAN-TIM |
| AHNAK | 1.034882644 | 0.426 | 0.223 | 7.02E-201 | 06_VCAN-TIM |
| MMP19 | 1.042854438 | 0.465 | 0.254 | 2.52E-200 | 06_VCAN-TIM |
| PDE3B | -1.377325178 | 0.342 | 0.562 | 1.22E-199 | 06_VCAN-TIM |
| SRGAP1 | -1.137078949 | 0.433 | 0.633 | 2.61E-194 | 06_VCAN-TIM |
| AOAH | -1.365703171 | 0.263 | 0.481 | 5.93E-187 | 06_VCAN-TIM |
| VASH1 | -1.23941473 | 0.279 | 0.498 | 9.95E-186 | 06_VCAN-TIM |
| SRGAP2C | -1.45955067 | 0.257 | 0.469 | 1.30E-185 | 06_VCAN-TIM |
| FAM149A | -2.142945367 | 0.096 | 0.298 | 3.26E-182 | 06_VCAN-TIM |
| CTTNBP2 | -1.912736136 | 0.115 | 0.318 | 5.00E-173 | 06_VCAN-TIM |
| ABCC4 | -1.768735931 | 0.177 | 0.378 | 4.00E-170 | 06_VCAN-TIM |
| CD69 | -1.388996768 | 0.239 | 0.441 | 7.12E-166 | 06_VCAN-TIM |
| BAG3 | 2.565826453 | 0.78 | 0.241 | 0 | 07_HSP-Macrophage2 |
| HSPH1 | 2.071692783 | 0.945 | 0.45 | 0 | 07_HSP-Macrophage2 |
| HSPA6 | 3.152145662 | 0.672 | 0.203 | 0 | 07_HSP-Macrophage2 |
| DNAJB1 | 2.563025341 | 0.956 | 0.491 | 0 | 07_HSP-Macrophage2 |
| HSPA1B | 2.403684387 | 0.968 | 0.506 | 0 | 07_HSP-Macrophage2 |
| HSPD1 | 2.546597543 | 0.899 | 0.449 | 0 | 07_HSP-Macrophage2 |
| HSPE1 | 2.392410785 | 0.835 | 0.403 | 0 | 07_HSP-Macrophage2 |
| ZFAND2A | 2.446815023 | 0.602 | 0.181 | 0 | 07_HSP-Macrophage2 |
| HSPB1 | 2.62468462 | 0.871 | 0.47 | 0 | 07_HSP-Macrophage2 |
| DNAJA4 | 2.266699497 | 0.556 | 0.159 | 0 | 07_HSP-Macrophage2 |
| HSPA1A | 2.257314568 | 0.983 | 0.599 | 0 | 07_HSP-Macrophage2 |

| | | | | | |
|---|---|---|---|---|---|
| IER5 | 2.142982734 | 0.685 | 0.341 | 0 | 07_HSP-Macrophage2 |
| FKBP4 | 2.515371299 | 0.356 | 0.09 | 0 | 07_HSP-Macrophage2 |
| CACYBP | 2.110490857 | 0.482 | 0.173 | 5.32E-295 | 07_HSP-Macrophage2 |
| CHORDC1 | 1.69973209 | 0.528 | 0.216 | 1.70E-257 | 07_HSP-Macrophage2 |
| SERPINH1 | 2.482414007 | 0.261 | 0.061 | 3.59E-250 | 07_HSP-Macrophage2 |
| DNAJA1 | 1.379908759 | 0.779 | 0.526 | 1.52E-237 | 07_HSP-Macrophage2 |
| P4HA1 | 1.401265339 | 0.698 | 0.411 | 9.71E-204 | 07_HSP-Macrophage2 |
| STIP1 | 1.586343292 | 0.485 | 0.21 | 5.88E-202 | 07_HSP-Macrophage2 |
| HSPA8 | 1.194738446 | 0.821 | 0.591 | 3.22E-194 | 07_HSP-Macrophage2 |
| DNAJB4 | 1.868622505 | 0.348 | 0.119 | 5.04E-194 | 07_HSP-Macrophage2 |
| MRPL18 | 1.782809519 | 0.383 | 0.156 | 2.16E-170 | 07_HSP-Macrophage2 |
| SNAP23 | 1.541863099 | 0.506 | 0.261 | 7.43E-153 | 07_HSP-Macrophage2 |
| TCP1 | 1.441885566 | 0.454 | 0.228 | 6.43E-134 | 07_HSP-Macrophage2 |
| ATP2C1 | 1.102588132 | 0.586 | 0.382 | 2.01E-97 | 07_HSP-Macrophage2 |
| AREG | 2.257239803 | 0.809 | 0.289 | 0 | 08_DC |
| CRYBG1 | 2.713454743 | 0.562 | 0.128 | 0 | 08_DC |
| PPA1 | 2.435695419 | 0.589 | 0.165 | 0 | 08_DC |
| LSP1 | 1.800160361 | 0.7 | 0.282 | 0 | 08_DC |
| FCER1A | 4.323700851 | 0.448 | 0.032 | 0 | 08_DC |
| CST7 | 3.726786983 | 0.466 | 0.071 | 0 | 08_DC |
| IL1R2 | 3.629650864 | 0.457 | 0.072 | 0 | 08_DC |
| CCSER1 | 3.244482376 | 0.449 | 0.067 | 0 | 08_DC |
| AFF3 | 2.810680796 | 0.427 | 0.054 | 0 | 08_DC |
| DUSP4 | 2.134994386 | 0.532 | 0.163 | 0 | 08_DC |
| GPAT3 | 2.657128402 | 0.444 | 0.076 | 0 | 08_DC |
| CNN2 | 2.739196901 | 0.436 | 0.073 | 0 | 08_DC |
| C15orf48 | 3.12947551 | 0.485 | 0.129 | 0 | 08_DC |
| DUSP5 | 2.886011831 | 0.421 | 0.093 | 0 | 08_DC |
| CRIP1 | 2.501430789 | 0.421 | 0.1 | 0 | 08_DC |
| CLEC10A | 3.553867365 | 0.348 | 0.033 | 0 | 08_DC |
| MCOLN2 | 3.080948874 | 0.354 | 0.047 | 0 | 08_DC |
| CD1C | 4.710686005 | 0.319 | 0.015 | 0 | 08_DC |
| PKIB | 2.325840359 | 0.398 | 0.098 | 0 | 08_DC |
| JAML | 2.783649396 | 0.351 | 0.051 | 0 | 08_DC |
| IL18R1 | 3.030503166 | 0.331 | 0.036 | 0 | 08_DC |
| HLA-DQA1 | 2.113880183 | 0.946 | 0.665 | 0 | 08_DC |
| IRF4 | 2.628443481 | 0.291 | 0.033 | 0 | 08_DC |
| FLT3 | 3.832866991 | 0.277 | 0.021 | 0 | 08_DC |
| SPIB | 2.447936712 | 0.311 | 0.058 | 0 | 08_DC |
| ADAM19 | 3.812763803 | 0.274 | 0.022 | 0 | 08_DC |
| TRAF1 | 2.292567268 | 0.317 | 0.065 | 0 | 08_DC |
| ICAM3 | 2.983387753 | 0.281 | 0.033 | 0 | 08_DC |
| CYP2S1 | 2.846408954 | 0.294 | 0.047 | 0 | 08_DC |
| CCR7 | 2.923835499 | 0.294 | 0.053 | 0 | 08_DC |
| IL1R1 | 2.581909762 | 0.28 | 0.047 | 0 | 08_DC |
| CCL22 | 5.002323967 | 0.25 | 0.018 | 0 | 08_DC |

| | | | | | |
|---|---|---|---|---|---|
| RHOF | 2.745184413 | 0.274 | 0.046 | 0 | 08_DC |
| GPR157 | 3.937746909 | 0.249 | 0.022 | 0 | 08_DC |
| SLC38A1 | 2.837388962 | 0.242 | 0.026 | 0 | 08_DC |
| FRY | 2.901741206 | 0.247 | 0.041 | 0 | 08_DC |
| ISG20 | 1.712573822 | 0.508 | 0.154 | 4.27E-298 | 08_DC |
| IL2RG | 2.157835674 | 0.342 | 0.077 | 2.57E-297 | 08_DC |
| FYN | 1.825821219 | 0.349 | 0.079 | 1.34E-290 | 08_DC |
| SH3BP4 | 2.033530605 | 0.333 | 0.074 | 2.19E-288 | 08_DC |
| SPP1 | -1.965753639 | 0.632 | 0.887 | 5.14E-286 | 08_DC |
| SLC1A3 | -1.522493789 | 0.544 | 0.891 | 4.29E-285 | 08_DC |
| SULF2 | 2.024942393 | 0.318 | 0.07 | 1.59E-274 | 08_DC |
| RFTN1 | 1.795172081 | 0.514 | 0.166 | 1.59E-273 | 08_DC |
| DAPP1 | 2.820875267 | 0.28 | 0.06 | 9.71E-262 | 08_DC |
| CD48 | 1.93563582 | 0.332 | 0.08 | 1.62E-257 | 08_DC |
| LYZ | 1.350434829 | 0.684 | 0.301 | 2.28E-256 | 08_DC |
| VDR | 2.246676851 | 0.287 | 0.062 | 4.14E-254 | 08_DC |
| SLC11A1 | -1.46908861 | 0.473 | 0.837 | 1.44E-249 | 08_DC |
| CYTIP | 1.560932092 | 0.585 | 0.226 | 6.29E-248 | 08_DC |
| VIM | 1.184912669 | 0.907 | 0.663 | 7.85E-245 | 08_DC |
| RPS5 | 1.391916016 | 0.818 | 0.557 | 6.69E-243 | 08_DC |
| RAB11FIP1 | 2.043396733 | 0.37 | 0.102 | 1.13E-242 | 08_DC |
| TIMP1 | 2.273354939 | 0.662 | 0.321 | 1.00E-239 | 08_DC |
| SATB1 | 1.941570103 | 0.373 | 0.105 | 1.13E-237 | 08_DC |
| FRMD4A | -1.792922334 | 0.463 | 0.8 | 2.04E-237 | 08_DC |
| RALA | 1.666307673 | 0.567 | 0.224 | 3.82E-236 | 08_DC |
| IFITM1 | 1.599034732 | 0.389 | 0.112 | 5.06E-236 | 08_DC |
| SLC41A2 | 2.074125333 | 0.269 | 0.058 | 6.67E-235 | 08_DC |
| APOE | -1.26195939 | 0.602 | 0.893 | 4.34E-233 | 08_DC |
| ARF6 | 1.755415075 | 0.582 | 0.251 | 8.69E-219 | 08_DC |
| AHNAK | 1.363742336 | 0.585 | 0.231 | 4.54E-218 | 08_DC |
| IFITM2 | 1.753244991 | 0.734 | 0.43 | 1.49E-213 | 08_DC |
| BCL3 | 1.983385085 | 0.459 | 0.168 | 2.52E-213 | 08_DC |
| KCNMA1 | -2.582009452 | 0.179 | 0.58 | 9.92E-210 | 08_DC |
| CDK2AP1 | 1.891578411 | 0.522 | 0.217 | 2.20E-208 | 08_DC |
| EEF1G | 1.224017009 | 0.808 | 0.516 | 2.66E-207 | 08_DC |
| AL133415.1 | 1.921654896 | 0.368 | 0.114 | 7.46E-206 | 08_DC |
| NRARP | 2.106536225 | 0.318 | 0.088 | 6.62E-205 | 08_DC |
| MERTK | -2.195607641 | 0.206 | 0.604 | 7.76E-205 | 08_DC |
| RPL10A | 1.187037873 | 0.818 | 0.571 | 8.17E-204 | 08_DC |
| SPINT2 | 1.521878405 | 0.501 | 0.196 | 3.01E-198 | 08_DC |
| EEF1B2 | 1.271867843 | 0.782 | 0.517 | 1.66E-197 | 08_DC |
| S100A10 | 1.264351361 | 0.719 | 0.386 | 3.36E-194 | 08_DC |
| BIRC3 | 2.070722571 | 0.372 | 0.12 | 6.04E-193 | 08_DC |
| RPSA | 1.105890597 | 0.821 | 0.569 | 4.17E-190 | 08_DC |
| C1QC | -1.200481792 | 0.559 | 0.84 | 1.99E-187 | 08_DC |
| S100A6 | 1.01791665 | 0.699 | 0.352 | 3.76E-186 | 08_DC |

| | | | | | |
|---|---|---|---|---|---|
| HLA-DQB2 | 2.054722862 | 0.327 | 0.098 | 1.98E-185 | 08_DC |
| INSIG1 | 1.626403992 | 0.566 | 0.257 | 2.36E-185 | 08_DC |
| RPL35A | 1.054846402 | 0.856 | 0.652 | 1.04E-184 | 08_DC |
| C1orf162 | 1.112325988 | 0.78 | 0.476 | 2.68E-178 | 08_DC |
| RPL23A | 1.071031015 | 0.824 | 0.602 | 4.17E-177 | 08_DC |
| ARHGAP24 | -1.573156355 | 0.388 | 0.718 | 6.29E-177 | 08_DC |
| ABCA1 | -1.876908888 | 0.353 | 0.667 | 9.39E-176 | 08_DC |
| SEL1L3 | 1.384080833 | 0.372 | 0.122 | 5.22E-175 | 08_DC |
| S100A4 | 1.185992268 | 0.689 | 0.367 | 1.16E-174 | 08_DC |
| RPL21 | 1.101080085 | 0.827 | 0.626 | 2.40E-173 | 08_DC |
| EMP3 | 1.403643997 | 0.674 | 0.364 | 1.67E-171 | 08_DC |
| RPL5 | 1.055544085 | 0.856 | 0.647 | 6.37E-171 | 08_DC |
| MTSS1 | -2.066341705 | 0.222 | 0.574 | 2.51E-170 | 08_DC |
| RPL4 | 1.029209671 | 0.812 | 0.572 | 1.36E-167 | 08_DC |
| GSTP1 | 1.184142555 | 0.727 | 0.422 | 1.61E-166 | 08_DC |
| TES | 1.320195247 | 0.427 | 0.158 | 2.53E-166 | 08_DC |
| CKLF | 1.39157031 | 0.613 | 0.309 | 5.51E-166 | 08_DC |
| LHFPL2 | -1.695743278 | 0.276 | 0.617 | 1.36E-163 | 08_DC |
| SERPINB1 | 1.53538444 | 0.698 | 0.413 | 2.31E-162 | 08_DC |
| STAB1 | -2.365947625 | 0.171 | 0.515 | 1.46E-160 | 08_DC |
| EIF3L | 1.299337514 | 0.547 | 0.255 | 6.54E-160 | 08_DC |
| FCGR2B | 1.264539639 | 0.546 | 0.248 | 4.09E-159 | 08_DC |
| BTF3 | 1.029323543 | 0.783 | 0.529 | 4.16E-159 | 08_DC |
| CYTOR | 1.562370915 | 0.349 | 0.119 | 2.25E-158 | 08_DC |
| ETV3 | 1.362802303 | 0.537 | 0.241 | 5.33E-158 | 08_DC |
| PLTP | -2.711197144 | 0.172 | 0.504 | 2.45E-157 | 08_DC |
| CXCL16 | 1.062636323 | 0.838 | 0.614 | 1.76E-153 | 08_DC |
| MYL12A | 1.054989444 | 0.715 | 0.427 | 9.63E-150 | 08_DC |
| TAGLN2 | 1.254522439 | 0.666 | 0.379 | 6.49E-149 | 08_DC |
| SLCO2B1 | -1.801019022 | 0.25 | 0.577 | 1.53E-148 | 08_DC |
| GPR183 | 1.030886003 | 0.804 | 0.525 | 1.63E-147 | 08_DC |
| HINT1 | 1.112191159 | 0.659 | 0.363 | 6.59E-146 | 08_DC |
| BNC2 | -2.252243799 | 0.192 | 0.519 | 1.22E-145 | 08_DC |
| MAML2 | -1.646349984 | 0.399 | 0.674 | 4.17E-143 | 08_DC |
| CMTM6 | 1.068936913 | 0.764 | 0.482 | 2.62E-142 | 08_DC |
| APOC1 | -1.231051613 | 0.513 | 0.758 | 3.56E-142 | 08_DC |
| FMNL2 | -1.406006845 | 0.569 | 0.785 | 4.68E-142 | 08_DC |
| EIF3K | 1.03306619 | 0.688 | 0.401 | 5.66E-142 | 08_DC |
| COMMD6 | 1.035423693 | 0.625 | 0.329 | 3.90E-137 | 08_DC |
| SLC25A5 | 1.039227435 | 0.666 | 0.375 | 7.57E-137 | 08_DC |
| EIF3F | 1.087661175 | 0.604 | 0.317 | 6.58E-136 | 08_DC |
| RARA | 1.362638804 | 0.358 | 0.134 | 8.14E-132 | 08_DC |
| DUSP2 | 1.051748733 | 0.589 | 0.303 | 3.64E-127 | 08_DC |
| C1QB | -1.070011769 | 0.555 | 0.79 | 1.05E-125 | 08_DC |
| FOXN2 | 1.49892008 | 0.409 | 0.174 | 3.15E-125 | 08_DC |
| NPL | -2.250573795 | 0.125 | 0.431 | 8.27E-125 | 08_DC |

| | | | | | |
|---|---|---|---|---|---|
| SDK1 | -2.309646638 | 0.152 | 0.454 | 2.57E-124 | 08_DC |
| PLP2 | 1.149911358 | 0.358 | 0.137 | 9.85E-124 | 08_DC |
| SIPA1L3 | 1.590193387 | 0.391 | 0.162 | 7.18E-123 | 08_DC |
| TUBA1A | 1.290059668 | 0.525 | 0.266 | 1.46E-120 | 08_DC |
| CIITA | 1.064391348 | 0.609 | 0.322 | 7.12E-120 | 08_DC |
| SNHG15 | 1.287702708 | 0.467 | 0.22 | 3.06E-119 | 08_DC |
| LIMD2 | 1.075156114 | 0.462 | 0.208 | 1.62E-117 | 08_DC |
| STAT4 | 1.078661744 | 0.325 | 0.121 | 4.07E-116 | 08_DC |
| HMGA1 | 1.28121556 | 0.383 | 0.16 | 1.73E-115 | 08_DC |
| RNASE6 | 1.271340896 | 0.559 | 0.302 | 1.39E-114 | 08_DC |
| CDK14 | 1.037538818 | 0.389 | 0.159 | 1.61E-114 | 08_DC |
| NDUFV2 | 1.151936536 | 0.559 | 0.297 | 3.30E-114 | 08_DC |
| LINC01010 | 1.347719131 | 0.381 | 0.161 | 1.07E-111 | 08_DC |
| PDK4 | -1.762187272 | 0.258 | 0.525 | 8.68E-110 | 08_DC |
| SERPINE1 | -2.340318831 | 0.218 | 0.477 | 8.71E-110 | 08_DC |
| ZBTB46 | 1.344939209 | 0.348 | 0.138 | 1.00E-109 | 08_DC |
| DMXL2 | -2.87826196 | 0.072 | 0.342 | 9.38E-109 | 08_DC |
| XYLT1 | 1.029778343 | 0.464 | 0.217 | 2.32E-108 | 08_DC |
| SFMBT2 | -1.248191389 | 0.462 | 0.69 | 6.01E-108 | 08_DC |
| KCNQ3 | -1.429442159 | 0.236 | 0.53 | 6.16E-108 | 08_DC |
| SH3RF3 | -2.294913066 | 0.124 | 0.401 | 8.94E-108 | 08_DC |
| HTRA1 | -1.829589144 | 0.187 | 0.464 | 1.51E-107 | 08_DC |
| TNFAIP8 | 1.182123533 | 0.422 | 0.194 | 6.47E-105 | 08_DC |
| BID | 1.060908495 | 0.527 | 0.276 | 3.60E-104 | 08_DC |
| SLC2A5 | -1.882338266 | 0.162 | 0.437 | 5.27E-104 | 08_DC |
| TNS3 | -1.724532268 | 0.258 | 0.508 | 1.95E-102 | 08_DC |
| GABARAPL2 | 1.00056905 | 0.539 | 0.287 | 9.15E-101 | 08_DC |
| LNCAROD | -2.16465163 | 0.198 | 0.456 | 1.95E-97 | 08_DC |
| G0S2 | 1.829994944 | 0.398 | 0.195 | 2.06E-97 | 08_DC |
| MAFB | -1.477619855 | 0.366 | 0.601 | 1.66E-96 | 08_DC |
| KLF2 | -1.326554226 | 0.332 | 0.575 | 3.57E-96 | 08_DC |
| FCGBP | -1.870859136 | 0.168 | 0.432 | 1.09E-95 | 08_DC |
| SUPT4H1 | 1.124353315 | 0.382 | 0.173 | 3.20E-94 | 08_DC |
| ARHGAP6 | -2.089457562 | 0.135 | 0.387 | 3.15E-93 | 08_DC |
| ATG7 | -1.554278195 | 0.316 | 0.539 | 1.55E-92 | 08_DC |
| FRMD4B | -1.536734108 | 0.301 | 0.528 | 2.12E-90 | 08_DC |
| SLC9A9 | -1.762346056 | 0.222 | 0.462 | 1.16E-89 | 08_DC |
| CTSL | -1.956879874 | 0.249 | 0.476 | 1.48E-89 | 08_DC |
| DLEU1 | -1.353947538 | 0.387 | 0.602 | 2.24E-88 | 08_DC |
| DENND1B | 1.139784346 | 0.381 | 0.175 | 2.23E-86 | 08_DC |
| FHIT | -1.371626753 | 0.331 | 0.551 | 1.88E-85 | 08_DC |
| ST6GAL1 | -1.520783351 | 0.316 | 0.536 | 2.85E-85 | 08_DC |
| VMO1 | 1.366042508 | 0.425 | 0.218 | 7.50E-85 | 08_DC |
| RNASE1 | -2.902603455 | 0.132 | 0.354 | 6.97E-84 | 08_DC |
| APOC2 | -1.723301638 | 0.221 | 0.45 | 4.79E-81 | 08_DC |
| MGAT5 | -1.855132259 | 0.258 | 0.47 | 6.16E-81 | 08_DC |

| | | | | | |
|---|---|---|---|---|---|
| GPNMB | -2.321785905 | 0.096 | 0.322 | 7.59E-81 | 08_DC |
| TMIGD3 | -1.768359847 | 0.157 | 0.389 | 9.65E-81 | 08_DC |
| FOLR2 | -2.36551538 | 0.099 | 0.322 | 5.45E-80 | 08_DC |
| TFRC | -1.745631909 | 0.279 | 0.491 | 5.12E-79 | 08_DC |
| TBC1D16 | -1.886954975 | 0.159 | 0.386 | 1.85E-78 | 08_DC |
| DIAPH2 | -1.535011801 | 0.266 | 0.478 | 2.45E-77 | 08_DC |
| MITF | -1.81913504 | 0.197 | 0.414 | 8.47E-74 | 08_DC |
| STARD13 | -2.097619958 | 0.122 | 0.337 | 1.67E-72 | 08_DC |
| LRP1 | -1.611330778 | 0.202 | 0.402 | 1.80E-66 | 08_DC |
| HLA-DRA | -1.414193298 | 0.593 | 0.932 | 2.74E-208 | 09_lowreads |
| HLA-DRB1 | -1.353621374 | 0.55 | 0.917 | 4.18E-194 | 09_lowreads |
| CST3 | -1.316080234 | 0.419 | 0.835 | 5.91E-181 | 09_lowreads |
| B2M | -1.086803374 | 0.704 | 0.961 | 4.91E-179 | 09_lowreads |
| HLA-DPA1 | -1.39573052 | 0.447 | 0.852 | 7.70E-178 | 09_lowreads |
| HLA-A | -1.103638151 | 0.39 | 0.852 | 1.10E-177 | 09_lowreads |
| TYROBP | -1.148503145 | 0.464 | 0.86 | 6.99E-177 | 09_lowreads |
| C1QC | -1.243189264 | 0.472 | 0.841 | 2.17E-165 | 09_lowreads |
| NPC2 | -1.119003919 | 0.309 | 0.771 | 3.65E-164 | 09_lowreads |
| C1QA | -1.2455532 | 0.384 | 0.789 | 4.05E-163 | 09_lowreads |
| TMSB4X | -1.079471508 | 0.712 | 0.945 | 4.99E-162 | 09_lowreads |
| HLA-DRB5 | -1.51968046 | 0.339 | 0.753 | 9.68E-162 | 09_lowreads |
| CYBA | -1.064306502 | 0.364 | 0.813 | 1.76E-161 | 09_lowreads |
| RPS23 | -1.19323661 | 0.379 | 0.804 | 4.26E-161 | 09_lowreads |
| RPLP1 | -1.059552309 | 0.582 | 0.899 | 4.54E-160 | 09_lowreads |
| ITM2B | -1.124226031 | 0.425 | 0.845 | 1.53E-158 | 09_lowreads |
| RPS19 | -1.068045113 | 0.443 | 0.847 | 3.18E-156 | 09_lowreads |
| FCER1G | -1.075468991 | 0.398 | 0.827 | 1.15E-153 | 09_lowreads |
| C1QB | -1.23957701 | 0.403 | 0.794 | 1.27E-151 | 09_lowreads |
| CD81 | -1.270580936 | 0.241 | 0.671 | 5.67E-150 | 09_lowreads |
| RPL32 | -1.10035803 | 0.384 | 0.802 | 2.55E-149 | 09_lowreads |
| RPS18 | -1.06806665 | 0.342 | 0.785 | 2.71E-149 | 09_lowreads |
| CFL1 | -1.021500888 | 0.292 | 0.749 | 4.60E-149 | 09_lowreads |
| RPL28 | -1.02178736 | 0.461 | 0.849 | 2.60E-148 | 09_lowreads |
| RPL34 | -1.115962223 | 0.333 | 0.768 | 2.56E-145 | 09_lowreads |
| FAU | -1.091567206 | 0.331 | 0.765 | 3.34E-143 | 09_lowreads |
| ITGB2 | -1.171762156 | 0.209 | 0.65 | 7.61E-141 | 09_lowreads |
| RPL37 | -1.071355808 | 0.335 | 0.768 | 2.13E-140 | 09_lowreads |
| HLA-C | -1.007181472 | 0.349 | 0.791 | 2.02E-138 | 09_lowreads |
| GRN | -1.24754395 | 0.195 | 0.616 | 1.96E-137 | 09_lowreads |
| RPS16 | -1.061723539 | 0.328 | 0.761 | 1.99E-137 | 09_lowreads |
| RPS14 | -1.036744089 | 0.331 | 0.758 | 2.15E-137 | 09_lowreads |
| RPLP2 | -1.044270342 | 0.339 | 0.766 | 3.00E-137 | 09_lowreads |
| RPS13 | -1.050379782 | 0.332 | 0.764 | 1.39E-136 | 09_lowreads |
| RPS12 | -1.056259966 | 0.423 | 0.815 | 8.21E-135 | 09_lowreads |
| RPS15A | -1.032317471 | 0.362 | 0.78 | 1.86E-134 | 09_lowreads |
| ATP6V0C | -1.12452002 | 0.231 | 0.66 | 2.13E-133 | 09_lowreads |

| | | | | | |
|---|---|---|---|---|---|
| CD14 | -1.354943348 | 0.215 | 0.62 | 3.06E-133 | 09_lowreads |
| CD63 | -1.026521767 | 0.255 | 0.691 | 2.24E-132 | 09_lowreads |
| RPL18A | -1.01446335 | 0.306 | 0.736 | 3.40E-131 | 09_lowreads |
| RPL36 | -1.010930682 | 0.292 | 0.725 | 5.37E-131 | 09_lowreads |
| S100A11 | -1.025764926 | 0.393 | 0.799 | 2.46E-130 | 09_lowreads |
| AIF1 | -1.081275379 | 0.216 | 0.638 | 2.09E-129 | 09_lowreads |
| TREM2 | -1.241348028 | 0.203 | 0.61 | 3.65E-128 | 09_lowreads |
| RPL39 | -1.062042211 | 0.311 | 0.734 | 8.33E-128 | 09_lowreads |
| RPL35A | -1.113571248 | 0.248 | 0.672 | 2.62E-127 | 09_lowreads |
| RPS28 | -1.038314247 | 0.336 | 0.755 | 8.28E-126 | 09_lowreads |
| RPS21 | -1.059278741 | 0.25 | 0.669 | 9.92E-126 | 09_lowreads |
| RPL36A | -1.083843015 | 0.226 | 0.646 | 1.18E-125 | 09_lowreads |
| RPL30 | -1.023529224 | 0.342 | 0.755 | 2.65E-125 | 09_lowreads |
| FCGR3A | -1.392800033 | 0.152 | 0.531 | 4.36E-125 | 09_lowreads |
| ATP5F1E | -1.076254854 | 0.22 | 0.638 | 2.02E-124 | 09_lowreads |
| RPL37A | -1.074652059 | 0.234 | 0.654 | 8.46E-124 | 09_lowreads |
| RPS27 | -1.042314727 | 0.31 | 0.728 | 3.71E-122 | 09_lowreads |
| CD68 | -1.035342138 | 0.218 | 0.627 | 2.87E-119 | 09_lowreads |
| HLA-DPB1 | -1.028290737 | 0.433 | 0.821 | 1.88E-117 | 09_lowreads |
| RPS26 | -1.032123549 | 0.27 | 0.674 | 1.96E-113 | 09_lowreads |
| DBI | -1.162744173 | 0.222 | 0.604 | 4.61E-113 | 09_lowreads |
| RPL21 | -1.08326774 | 0.255 | 0.645 | 9.82E-112 | 09_lowreads |
| RHOB | -1.122992052 | 0.412 | 0.769 | 1.84E-109 | 09_lowreads |
| RPL36AL | -1.056564622 | 0.164 | 0.532 | 8.02E-109 | 09_lowreads |
| RPL38 | -1.040591183 | 0.193 | 0.56 | 8.77E-106 | 09_lowreads |
| HCST | -1.478161157 | 0.114 | 0.442 | 4.64E-105 | 09_lowreads |
| ATP5MG | -1.024210732 | 0.156 | 0.519 | 2.79E-104 | 09_lowreads |
| HLA-DQA1 | -1.100323827 | 0.32 | 0.686 | 2.15E-103 | 09_lowreads |
| RPS29 | -1.073660485 | 0.158 | 0.501 | 1.86E-98 | 09_lowreads |
| FCGR1A | -1.113272726 | 0.141 | 0.472 | 3.61E-98 | 09_lowreads |
| CXCR4 | -1.06190173 | 0.295 | 0.648 | 9.04E-97 | 09_lowreads |
| RPS20 | -1.005575213 | 0.144 | 0.473 | 1.28E-95 | 09_lowreads |
| RPL23 | -1.073754222 | 0.15 | 0.476 | 2.63E-94 | 09_lowreads |
| SYNGR2 | -1.035151669 | 0.159 | 0.489 | 7.02E-92 | 09_lowreads |
| IFITM3 | -1.05631592 | 0.184 | 0.51 | 4.80E-91 | 09_lowreads |
| CHCHD10 | -1.122807722 | 0.125 | 0.428 | 5.96E-86 | 09_lowreads |
| KRTCAP2 | -1.124510743 | 0.118 | 0.41 | 9.55E-86 | 09_lowreads |
| IFITM2 | -1.165147461 | 0.145 | 0.45 | 1.06E-85 | 09_lowreads |
| COX6C | -1.086011109 | 0.135 | 0.436 | 2.35E-85 | 09_lowreads |
| CCL3 | -1.322111058 | 0.443 | 0.739 | 3.36E-83 | 09_lowreads |
| GPR34 | -1.092875371 | 0.15 | 0.445 | 3.90E-80 | 09_lowreads |
| ATP5MC3 | -1.202453471 | 0.112 | 0.391 | 4.27E-78 | 09_lowreads |
| BCAP31 | -1.184995581 | 0.117 | 0.393 | 1.02E-76 | 09_lowreads |
| SELENOK | -1.09502379 | 0.145 | 0.431 | 4.17E-76 | 09_lowreads |
| TMEM176B | -1.106888356 | 0.101 | 0.372 | 7.52E-76 | 09_lowreads |
| FOLR2 | -1.593758488 | 0.07 | 0.322 | 2.63E-75 | 09_lowreads |

| | | | | | |
|---|---|---|---|---|---|
| TOMM7 | -1.012457198 | 0.134 | 0.412 | 3.64E-75 | 09_lowreads |
| BCL2A1 | -1.20712148 | 0.166 | 0.452 | 4.64E-74 | 09_lowreads |
| HIGD2A | -1.011245742 | 0.112 | 0.386 | 1.46E-73 | 09_lowreads |
| IGSF6 | -1.280280456 | 0.093 | 0.347 | 4.78E-73 | 09_lowreads |
| PLD4 | -1.041596443 | 0.112 | 0.375 | 4.46E-69 | 09_lowreads |
| ATP5MF | -1.078441291 | 0.112 | 0.362 | 3.41E-67 | 09_lowreads |
| ARL6IP5 | -1.02863863 | 0.116 | 0.357 | 2.88E-64 | 09_lowreads |
| LIPA | -1.195636716 | 0.104 | 0.34 | 2.82E-63 | 09_lowreads |
| CH25H | -1.507292364 | 0.107 | 0.34 | 5.87E-60 | 09_lowreads |
| NOP10 | -1.023599366 | 0.097 | 0.324 | 1.04E-58 | 09_lowreads |
| RNASE6 | -1.102326483 | 0.099 | 0.318 | 1.27E-58 | 09_lowreads |
| OTUD1 | -1.055796826 | 0.104 | 0.339 | 7.22E-58 | 09_lowreads |
| CFD | -1.305469578 | 0.077 | 0.282 | 7.17E-54 | 09_lowreads |
| OLFML3 | -1.118641077 | 0.093 | 0.303 | 2.18E-52 | 09_lowreads |
| ANAPC11 | -1.102408248 | 0.089 | 0.294 | 2.62E-52 | 09_lowreads |
| CCL4 | -1.262869979 | 0.464 | 0.694 | 6.55E-49 | 09_lowreads |
| CCL3L1 | -1.266878023 | 0.209 | 0.442 | 1.61E-48 | 09_lowreads |
| CXCL8 | -1.089998851 | 0.184 | 0.393 | 1.70E-40 | 09_lowreads |
| FYN | 3.314016932 | 0.54 | 0.075 | 0 | 10_CD3_positive |
| IL32 | 4.849618838 | 0.481 | 0.03 | 0 | 10_CD3_positive |
| IFITM1 | 2.50211674 | 0.504 | 0.111 | 0 | 10_CD3_positive |
| CD3E | 5.388105304 | 0.399 | 0.013 | 0 | 10_CD3_positive |
| CCL5 | 3.277764557 | 0.456 | 0.093 | 0 | 10_CD3_positive |
| IL7R | 3.457983046 | 0.404 | 0.058 | 0 | 10_CD3_positive |
| CRIP1 | 2.364932364 | 0.438 | 0.102 | 0 | 10_CD3_positive |
| ETS1 | 3.387710277 | 0.373 | 0.047 | 0 | 10_CD3_positive |
| CD3D | 5.356856559 | 0.316 | 0.009 | 0 | 10_CD3_positive |
| CD52 | 3.105007911 | 0.359 | 0.061 | 0 | 10_CD3_positive |
| GZMK | 5.43420455 | 0.307 | 0.01 | 0 | 10_CD3_positive |
| CST7 | 2.529922108 | 0.371 | 0.077 | 0 | 10_CD3_positive |
| SYNE2 | 3.604388198 | 0.299 | 0.022 | 0 | 10_CD3_positive |
| CD2 | 5.183342259 | 0.285 | 0.009 | 0 | 10_CD3_positive |
| GZMA | 5.231139641 | 0.285 | 0.009 | 0 | 10_CD3_positive |
| CD96 | 4.643127669 | 0.282 | 0.011 | 0 | 10_CD3_positive |
| SPOCK2 | 4.194855727 | 0.283 | 0.012 | 0 | 10_CD3_positive |
| BCL11B | 5.018659859 | 0.277 | 0.009 | 0 | 10_CD3_positive |
| SKAP1 | 4.868785166 | 0.276 | 0.009 | 0 | 10_CD3_positive |
| TUBA4A | 3.855251524 | 0.295 | 0.028 | 0 | 10_CD3_positive |
| SLC38A1 | 3.555984298 | 0.289 | 0.027 | 0 | 10_CD3_positive |
| TC2N | 4.940343715 | 0.252 | 0.008 | 0 | 10_CD3_positive |
| PPP1R16B | 3.157842089 | 0.266 | 0.026 | 0 | 10_CD3_positive |
| CD247 | 3.86980955 | 0.257 | 0.017 | 0 | 10_CD3_positive |
| NKG7 | 3.804530281 | 0.272 | 0.033 | 0 | 10_CD3_positive |
| ITK | 4.697715725 | 0.246 | 0.009 | 0 | 10_CD3_positive |
| ITM2A | 3.394152742 | 0.229 | 0.013 | 0 | 10_CD3_positive |
| RORA | 3.345687157 | 0.237 | 0.023 | 0 | 10_CD3_positive |

| | | | | | |
|---|---|---|---|---|---|
| CAMK4 | 4.370966469 | 0.217 | 0.011 | 0 | 10_CD3_positive |
| CD7 | 3.046291579 | 0.272 | 0.058 | 5.23E-215 | 10_CD3_positive |
| IL2RG | 2.398001806 | 0.315 | 0.079 | 3.94E-197 | 10_CD3_positive |
| ISG20 | 1.855018378 | 0.435 | 0.159 | 1.22E-158 | 10_CD3_positive |
| RPS26 | 1.419617985 | 0.862 | 0.655 | 6.56E-153 | 10_CD3_positive |
| STAT4 | 1.600620947 | 0.373 | 0.121 | 1.06E-150 | 10_CD3_positive |
| CRYBG1 | 1.708356612 | 0.377 | 0.137 | 1.89E-133 | 10_CD3_positive |
| CNOT6L | 1.841934797 | 0.408 | 0.163 | 2.55E-128 | 10_CD3_positive |
| RUNX3 | 1.845954932 | 0.326 | 0.116 | 1.26E-117 | 10_CD3_positive |
| S100A4 | 1.22736412 | 0.651 | 0.37 | 4.09E-117 | 10_CD3_positive |
| RPS5 | 1.12435449 | 0.783 | 0.56 | 6.07E-116 | 10_CD3_positive |
| LEPROTL1 | 1.946794557 | 0.41 | 0.18 | 5.36E-114 | 10_CD3_positive |
| NIBAN1 | 1.60555055 | 0.415 | 0.185 | 4.19E-104 | 10_CD3_positive |
| S100A6 | 1.002638604 | 0.632 | 0.357 | 4.79E-99 | 10_CD3_positive |
| PARP8 | 1.566800115 | 0.517 | 0.281 | 3.51E-98 | 10_CD3_positive |
| PITPNC1 | 1.421784144 | 0.398 | 0.172 | 7.96E-98 | 10_CD3_positive |
| CBLB | 1.139032136 | 0.474 | 0.234 | 2.36E-88 | 10_CD3_positive |
| CEMIP2 | 1.301413239 | 0.43 | 0.211 | 7.81E-81 | 10_CD3_positive |
| DUSP2 | 1.008563431 | 0.547 | 0.307 | 4.26E-78 | 10_CD3_positive |
| TXNIP | 1.195491739 | 0.621 | 0.418 | 2.96E-74 | 10_CD3_positive |
| SRGAP2 | -1.005265238 | 0.51 | 0.714 | 2.63E-62 | 10_CD3_positive |
| PPDPF | 1.048446508 | 0.567 | 0.366 | 1.98E-61 | 10_CD3_positive |
| SLCO2B1 | -1.150818544 | 0.339 | 0.572 | 4.53E-59 | 10_CD3_positive |
| MERTK | -1.008777208 | 0.382 | 0.596 | 1.99E-50 | 10_CD3_positive |
| CSF3R | -1.028121839 | 0.329 | 0.535 | 5.04E-49 | 10_CD3_positive |
| ISG15 | 3.836679824 | 0.76 | 0.177 | 0 | 11_IFN-TAM |
| MX1 | 2.769230694 | 0.773 | 0.227 | 0 | 11_IFN-TAM |
| IFI44L | 2.050451359 | 0.778 | 0.272 | 0 | 11_IFN-TAM |
| IFIT3 | 3.911615872 | 0.563 | 0.066 | 0 | 11_IFN-TAM |
| IFI6 | 2.700279279 | 0.758 | 0.268 | 0 | 11_IFN-TAM |
| LY6E | 2.61331845 | 0.695 | 0.216 | 0 | 11_IFN-TAM |
| IFITM1 | 3.059456437 | 0.573 | 0.11 | 0 | 11_IFN-TAM |
| IFIT1 | 4.786916641 | 0.479 | 0.04 | 0 | 11_IFN-TAM |
| IFITM3 | 2.168401972 | 0.904 | 0.49 | 0 | 11_IFN-TAM |
| IFIT2 | 3.078434404 | 0.505 | 0.097 | 0 | 11_IFN-TAM |
| OAS1 | 2.637277276 | 0.495 | 0.104 | 0 | 11_IFN-TAM |
| OAS3 | 3.217307788 | 0.429 | 0.056 | 0 | 11_IFN-TAM |
| RSAD2 | 3.859267839 | 0.384 | 0.04 | 0 | 11_IFN-TAM |
| GBP1 | 3.041613375 | 0.421 | 0.078 | 0 | 11_IFN-TAM |
| HERC5 | 2.814151364 | 0.389 | 0.067 | 0 | 11_IFN-TAM |
| TNFSF10 | 3.167296201 | 0.377 | 0.071 | 0 | 11_IFN-TAM |
| CXCL10 | 5.340576242 | 0.29 | 0.025 | 0 | 11_IFN-TAM |
| MX2 | 2.187205859 | 0.647 | 0.205 | 8.55E-293 | 11_IFN-TAM |
| OAS2 | 2.46139065 | 0.368 | 0.069 | 2.28E-291 | 11_IFN-TAM |
| STAT1 | 1.92825056 | 0.647 | 0.207 | 3.89E-279 | 11_IFN-TAM |
| SIGLEC1 | 2.171968075 | 0.399 | 0.089 | 8.36E-249 | 11_IFN-TAM |

| | | | | | |
|---|---|---|---|---|---|
| OASL | 2.436443376 | 0.316 | 0.059 | 5.57E-243 | 11_IFN-TAM |
| IRF7 | 1.836048285 | 0.67 | 0.248 | 1.89E-241 | 11_IFN-TAM |
| IFI35 | 2.204838331 | 0.41 | 0.104 | 1.91E-223 | 11_IFN-TAM |
| GBP4 | 2.549432411 | 0.302 | 0.06 | 4.11E-219 | 11_IFN-TAM |
| CMPK2 | 2.545920821 | 0.253 | 0.043 | 4.60E-217 | 11_IFN-TAM |
| IFIH1 | 2.350127975 | 0.34 | 0.076 | 7.62E-211 | 11_IFN-TAM |
| XAF1 | 1.844219093 | 0.554 | 0.188 | 6.85E-203 | 11_IFN-TAM |
| SAMD9L | 2.037384631 | 0.422 | 0.119 | 1.50E-192 | 11_IFN-TAM |
| EPSTI1 | 1.696126981 | 0.679 | 0.291 | 1.82E-190 | 11_IFN-TAM |
| SP110 | 1.631019957 | 0.642 | 0.267 | 2.86E-183 | 11_IFN-TAM |
| PARP14 | 1.713770415 | 0.743 | 0.368 | 7.23E-176 | 11_IFN-TAM |
| DDX58 | 2.51267019 | 0.307 | 0.073 | 1.04E-173 | 11_IFN-TAM |
| UBE2L6 | 1.900553507 | 0.491 | 0.175 | 1.40E-170 | 11_IFN-TAM |
| TNFSF13B | 1.686405879 | 0.597 | 0.258 | 2.52E-159 | 11_IFN-TAM |
| IFI44 | 1.619881974 | 0.512 | 0.186 | 2.42E-155 | 11_IFN-TAM |
| LAP3 | 1.40437613 | 0.689 | 0.352 | 4.37E-145 | 11_IFN-TAM |
| SERPING1 | 1.781098686 | 0.475 | 0.176 | 4.12E-144 | 11_IFN-TAM |
| BST2 | 1.481042418 | 0.699 | 0.385 | 4.85E-144 | 11_IFN-TAM |
| EIF2AK2 | 1.632655983 | 0.489 | 0.189 | 1.76E-136 | 11_IFN-TAM |
| PSMB9 | 1.673380899 | 0.48 | 0.188 | 3.30E-135 | 11_IFN-TAM |
| TYMP | 1.07854841 | 0.873 | 0.609 | 5.76E-135 | 11_IFN-TAM |
| IFITM2 | 1.369324447 | 0.759 | 0.433 | 1.57E-134 | 11_IFN-TAM |
| LGALS3BP | 1.686524901 | 0.49 | 0.195 | 9.53E-133 | 11_IFN-TAM |
| PSME2 | 1.618254132 | 0.631 | 0.322 | 1.44E-132 | 11_IFN-TAM |
| ISG20 | 1.899326338 | 0.432 | 0.16 | 2.63E-127 | 11_IFN-TAM |
| TAP1 | 1.874373947 | 0.339 | 0.106 | 3.05E-127 | 11_IFN-TAM |
| PARP9 | 1.567341072 | 0.398 | 0.137 | 2.01E-123 | 11_IFN-TAM |
| VAMP5 | 1.89036814 | 0.373 | 0.131 | 2.80E-119 | 11_IFN-TAM |
| TRIM22 | 1.275521845 | 0.628 | 0.31 | 2.05E-117 | 11_IFN-TAM |
| RNF213 | 1.32891894 | 0.797 | 0.51 | 2.85E-115 | 11_IFN-TAM |
| WARS | 1.92152849 | 0.424 | 0.169 | 5.38E-108 | 11_IFN-TAM |
| APOL6 | 1.576696694 | 0.323 | 0.107 | 4.54E-105 | 11_IFN-TAM |
| CALHM6 | 1.748449583 | 0.483 | 0.219 | 7.40E-102 | 11_IFN-TAM |
| STAT2 | 1.423959872 | 0.398 | 0.154 | 8.86E-98 | 11_IFN-TAM |
| PML | 1.381068338 | 0.39 | 0.152 | 3.77E-94 | 11_IFN-TAM |
| SHFL | 1.494085788 | 0.342 | 0.134 | 3.39E-82 | 11_IFN-TAM |
| PSMB8 | 1.194589031 | 0.407 | 0.187 | 3.32E-72 | 11_IFN-TAM |
| SHISA5 | 1.233898055 | 0.344 | 0.142 | 5.50E-71 | 11_IFN-TAM |
| DDX60L | 1.228056365 | 0.455 | 0.22 | 5.26E-68 | 11_IFN-TAM |
| ADAR | 1.003740667 | 0.535 | 0.287 | 6.44E-68 | 11_IFN-TAM |
| GIMAP4 | 1.095742089 | 0.392 | 0.184 | 3.27E-63 | 11_IFN-TAM |
| SSB | 1.116502938 | 0.439 | 0.227 | 2.06E-59 | 11_IFN-TAM |
| PSMB10 | 1.026037459 | 0.485 | 0.264 | 2.12E-58 | 11_IFN-TAM |
| STMN1 | 3.882930261 | 0.727 | 0.138 | 0 | 12_Proliferating |
| MKI67 | 6.471957059 | 0.531 | 0.01 | 0 | 12_Proliferating |
| PCLAF | 5.985451556 | 0.501 | 0.012 | 0 | 12_Proliferating |

| | | | | | |
|---|---|---|---|---|---|
| DIAPH3 | 5.953105273 | 0.465 | 0.01 | 0 | 12_Proliferating |
| RRM2 | 6.753110499 | 0.457 | 0.007 | 0 | 12_Proliferating |
| NUSAP1 | 4.809990741 | 0.448 | 0.022 | 0 | 12_Proliferating |
| TYMS | 5.701595538 | 0.421 | 0.012 | 0 | 12_Proliferating |
| CENPF | 5.297865951 | 0.418 | 0.018 | 0 | 12_Proliferating |
| TK1 | 5.132488324 | 0.379 | 0.015 | 0 | 12_Proliferating |
| TOP2A | 5.951482406 | 0.367 | 0.008 | 0 | 12_Proliferating |
| CIT | 4.961902049 | 0.36 | 0.013 | 0 | 12_Proliferating |
| CLSPN | 5.042518766 | 0.338 | 0.011 | 0 | 12_Proliferating |
| ASPM | 6.661198237 | 0.33 | 0.004 | 0 | 12_Proliferating |
| UBE2C | 6.650354179 | 0.324 | 0.005 | 0 | 12_Proliferating |
| HELLS | 3.817531231 | 0.342 | 0.028 | 0 | 12_Proliferating |
| NCAPG2 | 3.780332296 | 0.335 | 0.025 | 0 | 12_Proliferating |
| TPX2 | 5.441565835 | 0.303 | 0.007 | 0 | 12_Proliferating |
| KNL1 | 5.439505719 | 0.299 | 0.008 | 0 | 12_Proliferating |
| GTSE1 | 5.86305394 | 0.297 | 0.006 | 0 | 12_Proliferating |
| PRC1 | 4.127025081 | 0.309 | 0.02 | 0 | 12_Proliferating |
| CDK1 | 3.407220757 | 0.318 | 0.03 | 0 | 12_Proliferating |
| LINC01572 | 3.30168362 | 0.321 | 0.035 | 0 | 12_Proliferating |
| BIRC5 | 5.534335097 | 0.291 | 0.006 | 0 | 12_Proliferating |
| UHRF1 | 4.929754633 | 0.294 | 0.011 | 0 | 12_Proliferating |
| PKMYT1 | 6.542188755 | 0.279 | 0.003 | 0 | 12_Proliferating |
| POLQ | 5.88100531 | 0.273 | 0.004 | 0 | 12_Proliferating |
| ZWINT | 5.041232639 | 0.27 | 0.009 | 0 | 12_Proliferating |
| BRIP1 | 4.774951525 | 0.271 | 0.011 | 0 | 12_Proliferating |
| HIST1H1B | 4.288644021 | 0.279 | 0.022 | 0 | 12_Proliferating |
| CENPM | 4.29968392 | 0.27 | 0.013 | 0 | 12_Proliferating |
| ANLN | 5.329514153 | 0.261 | 0.006 | 0 | 12_Proliferating |
| TROAP | 7.144041063 | 0.256 | 0.002 | 0 | 12_Proliferating |
| CENPK | 4.099083389 | 0.267 | 0.014 | 0 | 12_Proliferating |
| FANCI | 3.610089213 | 0.276 | 0.023 | 0 | 12_Proliferating |
| RAD51AP1 | 4.261275774 | 0.267 | 0.015 | 0 | 12_Proliferating |
| KIFC1 | 4.850326903 | 0.252 | 0.009 | 0 | 12_Proliferating |
| MELK | 5.899290695 | 0.237 | 0.004 | 0 | 12_Proliferating |
| CENPE | 4.034094485 | 0.247 | 0.019 | 0 | 12_Proliferating |
| AURKB | 6.618130592 | 0.226 | 0.003 | 0 | 12_Proliferating |
| NCAPH | 4.939999124 | 0.225 | 0.007 | 0 | 12_Proliferating |
| MYBL2 | 4.772404289 | 0.214 | 0.004 | 0 | 12_Proliferating |
| KIF11 | 5.079143822 | 0.21 | 0.006 | 0 | 12_Proliferating |
| Z94721.1 | 3.370048486 | 0.252 | 0.022 | 3.08E-295 | 12_Proliferating |
| FANCA | 3.642873901 | 0.241 | 0.021 | 9.02E-290 | 12_Proliferating |
| PTTG1 | 3.67174497 | 0.368 | 0.05 | 6.41E-288 | 12_Proliferating |
| CKS1B | 3.433737132 | 0.293 | 0.034 | 2.54E-267 | 12_Proliferating |
| TACC3 | 2.68235836 | 0.377 | 0.06 | 1.10E-237 | 12_Proliferating |
| ATAD2 | 2.924748571 | 0.33 | 0.047 | 7.44E-235 | 12_Proliferating |
| ATAD5 | 3.165721146 | 0.238 | 0.025 | 6.81E-230 | 12_Proliferating |

| | | | | | |
|---|---|---|---|---|---|
| SMC2 | 2.904996838 | 0.314 | 0.044 | 1.32E-228 | 12_Proliferating |
| LIG1 | 2.96295336 | 0.299 | 0.04 | 3.09E-224 | 12_Proliferating |
| CENPP | 2.433262846 | 0.519 | 0.123 | 8.96E-202 | 12_Proliferating |
| LMNB1 | 2.980176709 | 0.294 | 0.044 | 1.51E-195 | 12_Proliferating |
| KNTC1 | 2.934879173 | 0.24 | 0.03 | 1.05E-190 | 12_Proliferating |
| MCM7 | 2.946711169 | 0.261 | 0.036 | 9.39E-190 | 12_Proliferating |
| NCAPD2 | 3.055254735 | 0.243 | 0.031 | 1.92E-188 | 12_Proliferating |
| PCNA | 2.979227624 | 0.284 | 0.045 | 1.94E-177 | 12_Proliferating |
| SMC4 | 2.35781135 | 0.431 | 0.097 | 2.05E-174 | 12_Proliferating |
| DTYMK | 2.983346743 | 0.252 | 0.037 | 2.62E-171 | 12_Proliferating |
| BARD1 | 2.621552575 | 0.294 | 0.048 | 8.51E-171 | 12_Proliferating |
| KIF22 | 2.625490609 | 0.3 | 0.052 | 4.40E-164 | 12_Proliferating |
| H2AFZ | 2.467005062 | 0.75 | 0.365 | 1.41E-163 | 12_Proliferating |
| TUBB | 2.424015563 | 0.76 | 0.377 | 9.01E-158 | 12_Proliferating |
| NSD2 | 2.250397556 | 0.4 | 0.096 | 1.96E-144 | 12_Proliferating |
| KIF20B | 2.710843668 | 0.255 | 0.044 | 1.66E-138 | 12_Proliferating |
| BRCA2 | 2.224541197 | 0.297 | 0.059 | 1.79E-134 | 12_Proliferating |
| HMGN2 | 2.337167674 | 0.748 | 0.399 | 7.35E-134 | 12_Proliferating |
| DNMT1 | 1.993566782 | 0.501 | 0.163 | 1.10E-126 | 12_Proliferating |
| HMGB2 | 2.1100687 | 0.685 | 0.327 | 7.95E-125 | 12_Proliferating |
| HMGB1 | 1.489528832 | 0.867 | 0.645 | 2.52E-111 | 12_Proliferating |
| HIST1H4C | 2.909370766 | 0.452 | 0.153 | 7.13E-111 | 12_Proliferating |
| CEP128 | 1.896850896 | 0.371 | 0.101 | 1.41E-106 | 12_Proliferating |
| TUBA1B | 1.785707002 | 0.848 | 0.632 | 3.31E-104 | 12_Proliferating |
| H2AFV | 1.767708044 | 0.606 | 0.282 | 5.15E-93 | 12_Proliferating |
| EZH2 | 1.849008532 | 0.297 | 0.077 | 5.98E-90 | 12_Proliferating |
| H2AFX | 2.294185467 | 0.329 | 0.096 | 5.62E-89 | 12_Proliferating |
| CENPX | 2.146246396 | 0.299 | 0.085 | 6.11E-82 | 12_Proliferating |
| DUT | 1.847822217 | 0.471 | 0.192 | 4.29E-79 | 12_Proliferating |
| DEK | 1.455645272 | 0.624 | 0.33 | 1.43E-72 | 12_Proliferating |
| HIST1H1D | 2.151606455 | 0.288 | 0.085 | 8.27E-71 | 12_Proliferating |
| NUCKS1 | 1.365718435 | 0.664 | 0.368 | 5.76E-69 | 12_Proliferating |
| SMC1A | 1.708648628 | 0.409 | 0.156 | 1.31E-68 | 12_Proliferating |
| RANBP1 | 1.722701883 | 0.403 | 0.155 | 6.97E-68 | 12_Proliferating |
| PARP1 | 1.468208129 | 0.465 | 0.193 | 6.89E-67 | 12_Proliferating |
| CKAP5 | 1.783120528 | 0.33 | 0.111 | 4.00E-66 | 12_Proliferating |
| CCDC18 | 1.558772267 | 0.35 | 0.126 | 3.14E-61 | 12_Proliferating |
| NASP | 1.274669347 | 0.541 | 0.263 | 1.40E-60 | 12_Proliferating |
| TMPO | 1.329616395 | 0.452 | 0.195 | 3.23E-58 | 12_Proliferating |
| MCM5 | 1.354948518 | 0.368 | 0.142 | 6.61E-58 | 12_Proliferating |
| KPNA2 | 1.697211158 | 0.341 | 0.128 | 7.06E-57 | 12_Proliferating |
| PRIM2 | 1.438784933 | 0.357 | 0.136 | 9.46E-55 | 12_Proliferating |
| AC068587.4 | 1.579829717 | 0.421 | 0.185 | 4.82E-53 | 12_Proliferating |
| HIST1H1E | 2.037472868 | 0.383 | 0.168 | 6.53E-51 | 12_Proliferating |
| IDH2 | 1.374619038 | 0.41 | 0.181 | 8.87E-51 | 12_Proliferating |
| CKS2 | 1.172560759 | 0.48 | 0.228 | 2.27E-49 | 12_Proliferating |

| | | | | | |
|---|---|---|---|---|---|
| MZT2B | 1.338611616 | 0.526 | 0.282 | 3.42E-49 | 12_Proliferating |
| UQCC2 | 1.433496929 | 0.351 | 0.143 | 1.47E-48 | 12_Proliferating |
| SPATA5 | 1.165917403 | 0.367 | 0.147 | 2.34E-48 | 12_Proliferating |
| PRKDC | 1.235676092 | 0.466 | 0.229 | 1.07E-44 | 12_Proliferating |
| RAD21 | 1.204794142 | 0.436 | 0.207 | 1.29E-44 | 12_Proliferating |
| SIVA1 | 1.370247494 | 0.443 | 0.229 | 2.73E-43 | 12_Proliferating |
| HNRNPAB | 1.09725289 | 0.475 | 0.25 | 2.19E-38 | 12_Proliferating |
| UQCRQ | 1.232655065 | 0.522 | 0.3 | 2.41E-38 | 12_Proliferating |
| ANAPC11 | 1.250481702 | 0.489 | 0.285 | 2.10E-35 | 12_Proliferating |
| ANP32B | 1.198265806 | 0.496 | 0.286 | 3.79E-35 | 12_Proliferating |
| PAXX | 1.070316149 | 0.466 | 0.249 | 5.14E-35 | 12_Proliferating |
| RAN | 1.079607232 | 0.561 | 0.344 | 5.16E-35 | 12_Proliferating |
| IRF4 | 6.433386953 | 0.705 | 0.039 | 0 | 13_Angio-TAM |
| GZMB | 9.304744414 | 0.665 | 0.008 | 0 | 13_Angio-TAM |
| CXCR3 | 7.354679382 | 0.606 | 0.01 | 0 | 13_Angio-TAM |
| CLIC3 | 8.067924783 | 0.57 | 0.008 | 0 | 13_Angio-TAM |
| PTPRS | 7.47554396 | 0.546 | 0.006 | 0 | 13_Angio-TAM |
| RHEX | 6.912878379 | 0.53 | 0.009 | 0 | 13_Angio-TAM |
| JCHAIN | 6.418296851 | 0.522 | 0.003 | 0 | 13_Angio-TAM |
| PPP1R16B | 5.028427986 | 0.534 | 0.03 | 0 | 13_Angio-TAM |
| SEMA7A | 5.054066592 | 0.534 | 0.031 | 0 | 13_Angio-TAM |
| TSPAN13 | 6.526849615 | 0.506 | 0.014 | 0 | 13_Angio-TAM |
| IGKC | 4.758022282 | 0.47 | 0.006 | 0 | 13_Angio-TAM |
| DERL3 | 6.197221894 | 0.474 | 0.016 | 0 | 13_Angio-TAM |
| FAM160A1 | 7.169297547 | 0.45 | 0.008 | 0 | 13_Angio-TAM |
| PLXNA4 | 6.973993839 | 0.422 | 0.006 | 0 | 13_Angio-TAM |
| LILRA4 | 5.556917023 | 0.438 | 0.024 | 0 | 13_Angio-TAM |
| MAP1A | 6.430368415 | 0.406 | 0.005 | 0 | 13_Angio-TAM |
| COBLL1 | 5.390002047 | 0.402 | 0.009 | 0 | 13_Angio-TAM |
| PHEX | 5.557827619 | 0.39 | 0.017 | 0 | 13_Angio-TAM |
| CYFIP2 | 4.843359622 | 0.386 | 0.017 | 0 | 13_Angio-TAM |
| MZB1 | 7.11668908 | 0.359 | 0.001 | 0 | 13_Angio-TAM |
| RASD1 | 5.815389856 | 0.375 | 0.017 | 0 | 13_Angio-TAM |
| BCL11A | 5.243914864 | 0.371 | 0.015 | 0 | 13_Angio-TAM |
| SMPD3 | 8.198406168 | 0.355 | 0.002 | 0 | 13_Angio-TAM |
| P2RY14 | 5.963280252 | 0.355 | 0.009 | 0 | 13_Angio-TAM |
| NIBAN3 | 7.88706592 | 0.343 | 0.002 | 0 | 13_Angio-TAM |
| PPP1R14A | 4.814635099 | 0.355 | 0.014 | 0 | 13_Angio-TAM |
| LIME1 | 4.916093608 | 0.351 | 0.015 | 0 | 13_Angio-TAM |
| AC023590.1 | 6.917541834 | 0.339 | 0.005 | 0 | 13_Angio-TAM |
| MYBL2 | 6.781021047 | 0.335 | 0.006 | 0 | 13_Angio-TAM |
| LAMP5 | 7.983507418 | 0.303 | 0.002 | 0 | 13_Angio-TAM |
| CLEC4C | 9.974608444 | 0.287 | 0 | 0 | 13_Angio-TAM |
| LRRC26 | 10.56601279 | 0.259 | 0 | 0 | 13_Angio-TAM |
| PACSIN1 | 9.042557317 | 0.259 | 0.001 | 0 | 13_Angio-TAM |
| VASH2 | 10.31011753 | 0.251 | 0 | 0 | 13_Angio-TAM |

| | | | | | |
|---|---|---|---|---|---|
| LINC02812 | 5.955449324 | 0.255 | 0.005 | 0 | 13_Angio-TAM |
| LINC00996 | 5.9468387 | 0.247 | 0.006 | 0 | 13_Angio-TAM |
| SCT | 9.384110881 | 0.231 | 0 | 0 | 13_Angio-TAM |
| LINC01226 | 6.230385396 | 0.227 | 0.005 | 0 | 13_Angio-TAM |
| ANKRD53 | 7.741943963 | 0.223 | 0.002 | 0 | 13_Angio-TAM |
| CUX2 | 6.812799414 | 0.223 | 0.004 | 0 | 13_Angio-TAM |
| SCAMP5 | 6.620074065 | 0.219 | 0.003 | 0 | 13_Angio-TAM |
| INSYN2A | 6.629230334 | 0.207 | 0.003 | 0 | 13_Angio-TAM |
| TRAF4 | 4.519340624 | 0.442 | 0.033 | 5.44E-270 | 13_Angio-TAM |
| AFF3 | 4.313010062 | 0.602 | 0.064 | 1.61E-263 | 13_Angio-TAM |
| P2RY6 | 4.925112364 | 0.518 | 0.049 | 2.54E-260 | 13_Angio-TAM |
| PTGDS | 7.620517139 | 0.422 | 0.032 | 1.85E-258 | 13_Angio-TAM |
| CBFA2T3 | 4.696518713 | 0.223 | 0.01 | 5.84E-226 | 13_Angio-TAM |
| LTB | 4.679710124 | 0.335 | 0.023 | 7.81E-223 | 13_Angio-TAM |
| TPM2 | 4.209523728 | 0.386 | 0.031 | 1.09E-216 | 13_Angio-TAM |
| ADAM19 | 3.677834472 | 0.359 | 0.029 | 1.89E-195 | 13_Angio-TAM |
| COL24A1 | 4.884948569 | 0.247 | 0.014 | 1.94E-192 | 13_Angio-TAM |
| PLAC8 | 4.078313398 | 0.478 | 0.054 | 2.92E-192 | 13_Angio-TAM |
| SIDT1 | 4.053768453 | 0.47 | 0.054 | 2.02E-184 | 13_Angio-TAM |
| SELL | 3.633007849 | 0.462 | 0.052 | 2.89E-183 | 13_Angio-TAM |
| SEL1L3 | 4.287469944 | 0.689 | 0.128 | 4.86E-183 | 13_Angio-TAM |
| SULF2 | 3.772607375 | 0.55 | 0.076 | 7.27E-179 | 13_Angio-TAM |
| ATP2A3 | 4.315073279 | 0.291 | 0.021 | 8.66E-177 | 13_Angio-TAM |
| IL3RA | 4.081009481 | 0.669 | 0.124 | 2.03E-175 | 13_Angio-TAM |
| GLT1D1 | 3.99875909 | 0.386 | 0.038 | 4.05E-174 | 13_Angio-TAM |
| SPIB | 3.62545599 | 0.494 | 0.065 | 3.48E-168 | 13_Angio-TAM |
| ITM2C | 4.177695893 | 0.59 | 0.102 | 6.82E-161 | 13_Angio-TAM |
| LPIN1 | 4.04895793 | 0.526 | 0.079 | 2.82E-157 | 13_Angio-TAM |
| SLC38A1 | 3.494203877 | 0.339 | 0.032 | 1.77E-154 | 13_Angio-TAM |
| SOX4 | 4.523980253 | 0.426 | 0.053 | 3.75E-153 | 13_Angio-TAM |
| AC021594.2 | 4.206371247 | 0.279 | 0.023 | 3.30E-152 | 13_Angio-TAM |
| C12orf75 | 3.763879929 | 0.633 | 0.123 | 9.68E-152 | 13_Angio-TAM |
| EGLN3 | 3.210229807 | 0.386 | 0.046 | 6.12E-139 | 13_Angio-TAM |
| CCDC69 | 3.520051104 | 0.442 | 0.062 | 3.07E-136 | 13_Angio-TAM |
| POLB | 4.029365674 | 0.554 | 0.105 | 8.37E-134 | 13_Angio-TAM |
| LILRA5 | 3.602972849 | 0.267 | 0.024 | 6.59E-132 | 13_Angio-TAM |
| PLP2 | 3.294776879 | 0.633 | 0.142 | 5.84E-128 | 13_Angio-TAM |
| ENTPD7 | 4.132947178 | 0.291 | 0.029 | 2.69E-127 | 13_Angio-TAM |
| RAB11FIP1 | 3.344733153 | 0.562 | 0.109 | 6.98E-125 | 13_Angio-TAM |
| DUSP5 | 3.359687943 | 0.546 | 0.103 | 6.51E-123 | 13_Angio-TAM |
| PPP1R14B | 4.420178915 | 0.598 | 0.138 | 1.37E-121 | 13_Angio-TAM |
| PIM2 | 3.69180091 | 0.335 | 0.041 | 2.77E-115 | 13_Angio-TAM |
| CHAF1A | 3.679915071 | 0.315 | 0.037 | 1.89E-114 | 13_Angio-TAM |
| RUBCN | 3.222125641 | 0.633 | 0.157 | 2.23E-113 | 13_Angio-TAM |
| ZFAT | 4.006698549 | 0.367 | 0.05 | 7.02E-113 | 13_Angio-TAM |
| SPON2 | 3.904085736 | 0.275 | 0.029 | 1.36E-111 | 13_Angio-TAM |

| | | | | | |
|---|---|---|---|---|---|
| FLT3 | 3.477120699 | 0.271 | 0.028 | 1.97E-109 | 13_Angio-TAM |
| SLC41A2 | 3.21506088 | 0.41 | 0.064 | 1.26E-108 | 13_Angio-TAM |
| SEMA3C | 2.999358782 | 0.275 | 0.029 | 9.32E-108 | 13_Angio-TAM |
| TSEN54 | 3.12823929 | 0.327 | 0.042 | 4.39E-105 | 13_Angio-TAM |
| C12orf45 | 3.794727435 | 0.378 | 0.059 | 5.10E-103 | 13_Angio-TAM |
| BCL2L11 | 3.032101197 | 0.506 | 0.101 | 2.21E-102 | 13_Angio-TAM |
| CALCRL | 3.326586084 | 0.247 | 0.025 | 4.42E-101 | 13_Angio-TAM |
| P2RX1 | 3.897905606 | 0.235 | 0.024 | 3.73E-99 | 13_Angio-TAM |
| OPN3 | 3.096979265 | 0.514 | 0.107 | 4.52E-99 | 13_Angio-TAM |
| DSTN | 3.475164346 | 0.586 | 0.153 | 5.59E-99 | 13_Angio-TAM |
| RFTN1 | 2.720320508 | 0.645 | 0.176 | 4.82E-96 | 13_Angio-TAM |
| AGPAT5 | 3.310303328 | 0.398 | 0.068 | 1.66E-94 | 13_Angio-TAM |
| IRF7 | 3.368637456 | 0.717 | 0.256 | 8.04E-92 | 13_Angio-TAM |
| SRD5A1 | 3.539931466 | 0.283 | 0.038 | 7.50E-86 | 13_Angio-TAM |
| DOCK4 | -2.083892363 | 0.351 | 0.948 | 2.16E-84 | 13_Angio-TAM |
| CRIP1 | 2.36032566 | 0.498 | 0.109 | 6.04E-83 | 13_Angio-TAM |
| JAML | 2.790328966 | 0.355 | 0.06 | 1.55E-81 | 13_Angio-TAM |
| CYB561A3 | 3.139607468 | 0.355 | 0.062 | 1.33E-80 | 13_Angio-TAM |
| RBM38 | 3.112202621 | 0.378 | 0.07 | 1.89E-80 | 13_Angio-TAM |
| PLXDC2 | -2.098904478 | 0.402 | 0.949 | 4.41E-80 | 13_Angio-TAM |
| CSF2RB | 3.42141239 | 0.295 | 0.045 | 1.28E-77 | 13_Angio-TAM |
| EZR | 2.401400864 | 0.884 | 0.517 | 2.20E-75 | 13_Angio-TAM |
| PFKFB2 | 2.984049757 | 0.323 | 0.054 | 3.87E-75 | 13_Angio-TAM |
| TSPYL2 | 3.130738355 | 0.554 | 0.162 | 5.06E-75 | 13_Angio-TAM |
| SLC15A4 | 2.611251583 | 0.661 | 0.232 | 8.09E-75 | 13_Angio-TAM |
| SLC11A1 | -2.443329078 | 0.199 | 0.828 | 1.43E-72 | 13_Angio-TAM |
| SEC61B | 2.86218917 | 0.801 | 0.46 | 2.47E-71 | 13_Angio-TAM |
| SETBP1 | 3.550734761 | 0.239 | 0.033 | 1.43E-69 | 13_Angio-TAM |
| MCOLN2 | 2.693234721 | 0.323 | 0.057 | 4.42E-69 | 13_Angio-TAM |
| IL2RG | 2.450502808 | 0.398 | 0.084 | 8.29E-69 | 13_Angio-TAM |
| APOE | -2.50709235 | 0.478 | 0.885 | 2.00E-67 | 13_Angio-TAM |
| N4BP2L1 | 2.645504302 | 0.522 | 0.151 | 2.84E-66 | 13_Angio-TAM |
| AREG | 2.06504153 | 0.765 | 0.305 | 1.31E-65 | 13_Angio-TAM |
| SLC1A3 | -1.924044747 | 0.271 | 0.882 | 1.01E-64 | 13_Angio-TAM |
| GNA15 | 2.428509256 | 0.653 | 0.248 | 6.37E-64 | 13_Angio-TAM |
| C1QC | -2.42861514 | 0.327 | 0.833 | 1.16E-62 | 13_Angio-TAM |
| SEPHS1 | 3.279358154 | 0.259 | 0.042 | 1.02E-61 | 13_Angio-TAM |
| MSR1 | -2.83454955 | 0.135 | 0.732 | 1.02E-59 | 13_Angio-TAM |
| FRMD4A | -2.301047327 | 0.215 | 0.791 | 1.64E-57 | 13_Angio-TAM |
| LRMDA | -1.929718095 | 0.315 | 0.844 | 1.21E-56 | 13_Angio-TAM |
| FGD4 | -2.266893214 | 0.243 | 0.791 | 1.58E-56 | 13_Angio-TAM |
| PPM1K | 2.546153825 | 0.39 | 0.096 | 3.61E-56 | 13_Angio-TAM |
| FMNL2 | -2.603921615 | 0.267 | 0.78 | 5.93E-55 | 13_Angio-TAM |
| C1QA | -2.465502604 | 0.283 | 0.779 | 1.16E-54 | 13_Angio-TAM |
| FYTTD1 | 2.598849171 | 0.506 | 0.167 | 4.77E-54 | 13_Angio-TAM |
| ANTXR2 | 2.154172563 | 0.347 | 0.076 | 8.22E-54 | 13_Angio-TAM |

| | | | | | |
|---|---|---|---|---|---|
| HINT1 | 2.191787936 | 0.705 | 0.372 | 2.99E-53 | 13_Angio-TAM |
| C1QB | -2.40435298 | 0.335 | 0.784 | 6.67E-53 | 13_Angio-TAM |
| OLR1 | -2.000965636 | 0.279 | 0.807 | 8.31E-53 | 13_Angio-TAM |
| SPP1 | -2.431712035 | 0.566 | 0.879 | 1.63E-52 | 13_Angio-TAM |
| NR3C1 | 2.689612888 | 0.713 | 0.383 | 3.15E-52 | 13_Angio-TAM |
| APP | 2.189802928 | 0.649 | 0.294 | 8.55E-52 | 13_Angio-TAM |
| RAB31 | -1.847471476 | 0.375 | 0.861 | 2.33E-51 | 13_Angio-TAM |
| HYOU1 | 2.517235746 | 0.343 | 0.08 | 8.58E-51 | 13_Angio-TAM |
| DPYD | -2.689010109 | 0.207 | 0.713 | 3.49E-50 | 13_Angio-TAM |
| C3 | -2.310756069 | 0.243 | 0.749 | 3.56E-50 | 13_Angio-TAM |
| CD2AP | 2.214919333 | 0.606 | 0.249 | 1.06E-49 | 13_Angio-TAM |
| PTCRA | 2.743064997 | 0.315 | 0.07 | 1.44E-49 | 13_Angio-TAM |
| CXXC5 | 2.809518651 | 0.291 | 0.061 | 2.17E-49 | 13_Angio-TAM |
| SAMSN1 | -2.164380819 | 0.303 | 0.79 | 2.31E-49 | 13_Angio-TAM |
| SERPINF1 | 2.368243625 | 0.598 | 0.247 | 5.44E-49 | 13_Angio-TAM |
| RPS5 | 1.731096581 | 0.785 | 0.565 | 7.70E-49 | 13_Angio-TAM |
| MYL12A | 1.968710503 | 0.737 | 0.436 | 6.73E-48 | 13_Angio-TAM |
| PAXX | 2.519068616 | 0.594 | 0.251 | 7.93E-48 | 13_Angio-TAM |
| NAMPT | -2.36727783 | 0.398 | 0.795 | 8.20E-48 | 13_Angio-TAM |
| ATG101 | 2.541837318 | 0.382 | 0.104 | 1.97E-47 | 13_Angio-TAM |
| APOC1 | -2.535976633 | 0.307 | 0.752 | 6.30E-47 | 13_Angio-TAM |
| TAGLN2 | 1.885605592 | 0.717 | 0.387 | 2.15E-46 | 13_Angio-TAM |
| RPL10A | 1.629035252 | 0.805 | 0.578 | 3.74E-46 | 13_Angio-TAM |
| SPCS1 | 2.178362863 | 0.637 | 0.304 | 1.97E-45 | 13_Angio-TAM |
| MS4A7 | -1.861459695 | 0.299 | 0.768 | 2.51E-44 | 13_Angio-TAM |
| CD163 | -2.685488262 | 0.124 | 0.629 | 3.87E-44 | 13_Angio-TAM |
| SPINT2 | 2.073355022 | 0.534 | 0.205 | 4.54E-44 | 13_Angio-TAM |
| PBX3 | 2.420927945 | 0.538 | 0.213 | 5.95E-44 | 13_Angio-TAM |
| YPEL5 | 1.685852814 | 0.713 | 0.36 | 6.51E-44 | 13_Angio-TAM |
| VEGFB | 2.096892593 | 0.566 | 0.232 | 1.12E-43 | 13_Angio-TAM |
| EEF1B2 | 1.589693938 | 0.785 | 0.525 | 4.05E-43 | 13_Angio-TAM |
| PHB | 2.501099081 | 0.418 | 0.135 | 5.58E-43 | 13_Angio-TAM |
| ERN1 | 2.182893092 | 0.53 | 0.194 | 5.81E-43 | 13_Angio-TAM |
| RNF126 | 2.466762942 | 0.347 | 0.092 | 6.65E-43 | 13_Angio-TAM |
| HSP90B1 | 1.855025639 | 0.785 | 0.524 | 8.20E-43 | 13_Angio-TAM |
| ELL2 | -1.605310253 | 0.661 | 0.909 | 2.51E-42 | 13_Angio-TAM |
| PLD4 | 1.907947839 | 0.677 | 0.365 | 4.14E-42 | 13_Angio-TAM |
| IRF8 | 1.927885909 | 0.709 | 0.384 | 5.70E-42 | 13_Angio-TAM |
| SGK1 | -1.508278096 | 0.398 | 0.828 | 6.73E-42 | 13_Angio-TAM |
| ERP29 | 1.982646752 | 0.582 | 0.26 | 7.17E-42 | 13_Angio-TAM |
| PMEPA1 | 2.07969476 | 0.59 | 0.252 | 7.88E-42 | 13_Angio-TAM |
| ARHGAP26 | -1.848509595 | 0.402 | 0.807 | 1.48E-41 | 13_Angio-TAM |
| LIMS1 | -1.78536922 | 0.442 | 0.835 | 2.42E-41 | 13_Angio-TAM |
| ITGAE | 2.450097299 | 0.291 | 0.069 | 3.10E-41 | 13_Angio-TAM |
| CDYL | 2.360705016 | 0.494 | 0.18 | 3.89E-41 | 13_Angio-TAM |
| TLR2 | -1.993798202 | 0.211 | 0.704 | 4.40E-41 | 13_Angio-TAM |

| | | | | | |
|---|---|---|---|---|---|
| RHOB | -2.177343253 | 0.343 | 0.76 | 6.05E-41 | 13_Angio-TAM |
| RPSA | 1.401094422 | 0.809 | 0.576 | 6.11E-41 | 13_Angio-TAM |
| NUCB2 | 2.455883513 | 0.287 | 0.068 | 6.11E-41 | 13_Angio-TAM |
| PARK7 | 1.921718068 | 0.618 | 0.302 | 8.26E-41 | 13_Angio-TAM |
| UGCG | 1.909221416 | 0.614 | 0.281 | 1.06E-40 | 13_Angio-TAM |
| SLC8A1 | -1.896920701 | 0.299 | 0.736 | 1.54E-40 | 13_Angio-TAM |
| RRBP1 | 2.116791719 | 0.685 | 0.381 | 1.66E-40 | 13_Angio-TAM |
| SPN | 2.488643778 | 0.319 | 0.083 | 7.97E-40 | 13_Angio-TAM |
| PRKCB | 1.92582293 | 0.578 | 0.236 | 8.76E-40 | 13_Angio-TAM |
| TREM2 | -2.393847766 | 0.116 | 0.6 | 1.21E-39 | 13_Angio-TAM |
| INPP4A | 2.22523693 | 0.618 | 0.303 | 2.66E-39 | 13_Angio-TAM |
| GPR183 | 1.994504956 | 0.797 | 0.534 | 3.85E-39 | 13_Angio-TAM |
| FMN1 | -2.228833011 | 0.275 | 0.727 | 1.24E-38 | 13_Angio-TAM |
| HIF1A | -1.554962116 | 0.482 | 0.819 | 1.50E-38 | 13_Angio-TAM |
| CERS6 | 2.08829543 | 0.55 | 0.24 | 2.18E-38 | 13_Angio-TAM |
| C5AR1 | -2.312111349 | 0.183 | 0.637 | 2.34E-38 | 13_Angio-TAM |
| ZNF10 | 2.065969146 | 0.398 | 0.124 | 2.43E-38 | 13_Angio-TAM |
| ABCA1 | -2.554804723 | 0.211 | 0.658 | 3.57E-38 | 13_Angio-TAM |
| HIVEP1 | 1.86108227 | 0.598 | 0.261 | 4.06E-38 | 13_Angio-TAM |
| OSTC | 1.891760636 | 0.502 | 0.193 | 4.44E-38 | 13_Angio-TAM |
| ANXA11 | 1.747623049 | 0.614 | 0.288 | 4.88E-38 | 13_Angio-TAM |
| TNFRSF21 | 1.835226314 | 0.478 | 0.173 | 1.00E-37 | 13_Angio-TAM |
| SSR4 | 1.589939522 | 0.733 | 0.432 | 2.60E-37 | 13_Angio-TAM |
| TLE5 | 1.883024187 | 0.462 | 0.167 | 2.88E-37 | 13_Angio-TAM |
| FCHSD2 | 2.168496787 | 0.769 | 0.538 | 2.08E-35 | 13_Angio-TAM |
| CXCR4 | 1.192188686 | 0.896 | 0.636 | 2.87E-35 | 13_Angio-TAM |
| NUTM2B-AS1 | 2.064050125 | 0.526 | 0.226 | 5.97E-35 | 13_Angio-TAM |
| NSMCE3 | 2.411070615 | 0.275 | 0.07 | 6.75E-35 | 13_Angio-TAM |
| GLUL | -1.786407088 | 0.446 | 0.776 | 8.24E-35 | 13_Angio-TAM |
| ACSL1 | -1.460517144 | 0.641 | 0.882 | 8.85E-35 | 13_Angio-TAM |
| STAT4 | 1.824193218 | 0.398 | 0.127 | 9.42E-35 | 13_Angio-TAM |
| IL16 | 2.227982372 | 0.371 | 0.118 | 9.88E-35 | 13_Angio-TAM |
| KCNMA1 | -2.495787905 | 0.12 | 0.568 | 5.78E-34 | 13_Angio-TAM |
| MAML2 | -2.079718436 | 0.247 | 0.666 | 8.54E-34 | 13_Angio-TAM |
| IRAK3 | -1.695912248 | 0.235 | 0.683 | 1.11E-33 | 13_Angio-TAM |
| CD14 | -2.504135246 | 0.191 | 0.61 | 1.13E-33 | 13_Angio-TAM |
| SMC6 | 2.17474063 | 0.291 | 0.077 | 1.30E-33 | 13_Angio-TAM |
| SLC7A5 | 1.603384795 | 0.785 | 0.533 | 2.44E-33 | 13_Angio-TAM |
| ZFHX3 | -2.475254339 | 0.147 | 0.572 | 2.81E-33 | 13_Angio-TAM |
| HIGD1A | 2.686964337 | 0.299 | 0.085 | 2.84E-33 | 13_Angio-TAM |
| SNHG7 | 2.238546131 | 0.414 | 0.152 | 2.93E-33 | 13_Angio-TAM |
| COMMD6 | 1.604600158 | 0.641 | 0.338 | 7.20E-33 | 13_Angio-TAM |
| ALG2 | 2.093903282 | 0.355 | 0.112 | 7.23E-33 | 13_Angio-TAM |
| SLCO2B1 | -2.158782086 | 0.131 | 0.568 | 1.68E-32 | 13_Angio-TAM |
| EPB41L2 | -2.145800594 | 0.239 | 0.65 | 1.77E-32 | 13_Angio-TAM |
| MERTK | -1.987486923 | 0.151 | 0.592 | 2.76E-32 | 13_Angio-TAM |

| | | | | | |
|---|---|---|---|---|---|
| SMIM3 | 1.774517633 | 0.534 | 0.239 | 3.82E-32 | 13_Angio-TAM |
| SMIM14 | 2.119871241 | 0.355 | 0.114 | 5.54E-32 | 13_Angio-TAM |
| MDFIC | 2.253556967 | 0.271 | 0.07 | 5.99E-32 | 13_Angio-TAM |
| MARCKS | -2.179526508 | 0.203 | 0.613 | 7.17E-32 | 13_Angio-TAM |
| FCGR3A | -2.935959973 | 0.116 | 0.521 | 1.30E-31 | 13_Angio-TAM |
| CHCHD2 | 1.494278924 | 0.761 | 0.498 | 1.95E-31 | 13_Angio-TAM |
| ACTN4 | 1.760737897 | 0.494 | 0.203 | 2.47E-31 | 13_Angio-TAM |
| CSF1R | -1.806454495 | 0.191 | 0.626 | 2.53E-31 | 13_Angio-TAM |
| ISG20 | 1.603279829 | 0.45 | 0.165 | 3.97E-31 | 13_Angio-TAM |
| PTRHD1 | 2.125159167 | 0.347 | 0.113 | 4.19E-31 | 13_Angio-TAM |
| LSP1 | 1.406939228 | 0.622 | 0.295 | 4.69E-31 | 13_Angio-TAM |
| FCGR2A | -1.869412896 | 0.231 | 0.646 | 1.22E-30 | 13_Angio-TAM |
| KAT2B | 1.935053227 | 0.382 | 0.129 | 1.83E-30 | 13_Angio-TAM |
| VSIG4 | -2.886013177 | 0.096 | 0.506 | 2.03E-30 | 13_Angio-TAM |
| RELL1 | 1.786657386 | 0.478 | 0.189 | 2.33E-30 | 13_Angio-TAM |
| GAB1 | 2.17276017 | 0.446 | 0.177 | 2.67E-30 | 13_Angio-TAM |
| DNAJB9 | 2.163090882 | 0.319 | 0.096 | 3.03E-30 | 13_Angio-TAM |
| SRGAP1 | -2.174547771 | 0.207 | 0.616 | 3.44E-30 | 13_Angio-TAM |
| ALOX5AP | 1.333560235 | 0.777 | 0.549 | 4.92E-30 | 13_Angio-TAM |
| TANC2 | -2.533678662 | 0.135 | 0.547 | 9.96E-30 | 13_Angio-TAM |
| DDX24 | 1.447577855 | 0.721 | 0.459 | 1.18E-29 | 13_Angio-TAM |
| LCP2 | -1.759038023 | 0.295 | 0.688 | 1.29E-29 | 13_Angio-TAM |
| CAT | 1.791753875 | 0.394 | 0.142 | 1.70E-29 | 13_Angio-TAM |
| MIR4435-2HG | 1.913114294 | 0.347 | 0.112 | 3.62E-29 | 13_Angio-TAM |
| MEF2A | -1.317345401 | 0.49 | 0.806 | 4.26E-29 | 13_Angio-TAM |
| EIF2AK4 | 1.59770087 | 0.534 | 0.248 | 5.38E-29 | 13_Angio-TAM |
| CORO1C | 1.656371889 | 0.633 | 0.332 | 6.31E-29 | 13_Angio-TAM |
| CEBPD | -1.436187687 | 0.438 | 0.808 | 6.62E-29 | 13_Angio-TAM |
| TENT4A | 2.034171063 | 0.386 | 0.138 | 7.21E-29 | 13_Angio-TAM |
| SRP14 | 1.450681676 | 0.717 | 0.513 | 7.78E-29 | 13_Angio-TAM |
| JUN | -1.835765695 | 0.49 | 0.792 | 9.28E-29 | 13_Angio-TAM |
| TBXAS1 | -1.552477001 | 0.311 | 0.703 | 1.15E-28 | 13_Angio-TAM |
| ARL4C | 1.628842498 | 0.586 | 0.289 | 1.28E-28 | 13_Angio-TAM |
| EXT1 | 1.934966684 | 0.335 | 0.107 | 1.39E-28 | 13_Angio-TAM |
| A2M | -1.74348971 | 0.215 | 0.624 | 1.85E-28 | 13_Angio-TAM |
| CORO7 | 1.921037119 | 0.586 | 0.311 | 1.94E-28 | 13_Angio-TAM |
| CPM | -2.734195931 | 0.127 | 0.517 | 2.95E-28 | 13_Angio-TAM |
| OGT | 1.391132797 | 0.558 | 0.258 | 6.69E-28 | 13_Angio-TAM |
| FCGR1A | -3.183107055 | 0.076 | 0.464 | 7.53E-28 | 13_Angio-TAM |
| TCF4 | 1.668235019 | 0.693 | 0.457 | 9.03E-28 | 13_Angio-TAM |
| ASAH1 | -1.549731234 | 0.283 | 0.676 | 1.08E-27 | 13_Angio-TAM |
| ZBTB16 | -1.629861038 | 0.195 | 0.606 | 1.28E-27 | 13_Angio-TAM |
| CCL3 | -2.208905032 | 0.486 | 0.731 | 1.43E-27 | 13_Angio-TAM |
| HMOX1 | -2.769086338 | 0.171 | 0.556 | 1.43E-27 | 13_Angio-TAM |
| PDE8A | -2.395640734 | 0.175 | 0.562 | 1.51E-27 | 13_Angio-TAM |
| MAFB | -1.8871495 | 0.199 | 0.595 | 1.72E-27 | 13_Angio-TAM |

| | | | | | |
|---|---|---|---|---|---|
| JDP2 | -2.290432272 | 0.159 | 0.55 | 1.82E-27 | 13_Angio-TAM |
| PDE7A | 1.968385804 | 0.418 | 0.165 | 2.17E-27 | 13_Angio-TAM |
| DLEU1 | -1.977471778 | 0.227 | 0.596 | 2.76E-27 | 13_Angio-TAM |
| LPCAT2 | -2.184150914 | 0.139 | 0.533 | 3.28E-27 | 13_Angio-TAM |
| S100A6 | 1.047828782 | 0.693 | 0.363 | 3.66E-27 | 13_Angio-TAM |
| SUB1 | 1.403196132 | 0.669 | 0.41 | 4.20E-27 | 13_Angio-TAM |
| CYFIP1 | -1.828638739 | 0.167 | 0.574 | 6.35E-27 | 13_Angio-TAM |
| SLA | -1.955763089 | 0.183 | 0.572 | 7.51E-27 | 13_Angio-TAM |
| PLAUR | -1.817216311 | 0.446 | 0.725 | 7.53E-27 | 13_Angio-TAM |
| CCDC50 | 1.903242276 | 0.462 | 0.206 | 8.69E-27 | 13_Angio-TAM |
| RPL7 | 1.180872358 | 0.753 | 0.497 | 8.90E-27 | 13_Angio-TAM |
| IER3 | -1.953188504 | 0.291 | 0.645 | 1.38E-26 | 13_Angio-TAM |
| MTSS1 | -1.987456093 | 0.171 | 0.563 | 1.67E-26 | 13_Angio-TAM |
| SDCBP | -1.427343224 | 0.39 | 0.707 | 3.93E-26 | 13_Angio-TAM |
| STAB1 | -2.12611086 | 0.108 | 0.505 | 5.63E-26 | 13_Angio-TAM |
| CLEC7A | -1.847303273 | 0.171 | 0.557 | 6.05E-26 | 13_Angio-TAM |
| BNC2 | -2.540502796 | 0.131 | 0.509 | 9.99E-26 | 13_Angio-TAM |
| ZFP36L1 | -1.761752538 | 0.426 | 0.713 | 1.73E-25 | 13_Angio-TAM |
| PRKAG2 | -2.026451187 | 0.199 | 0.572 | 1.90E-25 | 13_Angio-TAM |
| USP9Y | 1.722373179 | 0.43 | 0.174 | 2.07E-25 | 13_Angio-TAM |
| CLINT1 | 1.561977017 | 0.498 | 0.23 | 2.31E-25 | 13_Angio-TAM |
| RASGEF1B | -1.436856678 | 0.498 | 0.763 | 2.52E-25 | 13_Angio-TAM |
| ARID5B | -2.213677607 | 0.195 | 0.56 | 5.25E-25 | 13_Angio-TAM |
| EEF1G | 1.154706666 | 0.741 | 0.526 | 1.06E-24 | 13_Angio-TAM |
| FAM177A1 | 1.923186649 | 0.49 | 0.234 | 1.26E-24 | 13_Angio-TAM |
| KCNQ3 | -1.901267352 | 0.139 | 0.522 | 1.39E-24 | 13_Angio-TAM |
| GORASP2 | 1.827023111 | 0.319 | 0.106 | 1.53E-24 | 13_Angio-TAM |
| RBM47 | -1.529981784 | 0.454 | 0.742 | 2.86E-24 | 13_Angio-TAM |
| EGR1 | -3.296596509 | 0.139 | 0.485 | 3.43E-24 | 13_Angio-TAM |
| ZC3HAV1 | 1.476635508 | 0.669 | 0.397 | 3.85E-24 | 13_Angio-TAM |
| CEBPB | -1.428411946 | 0.263 | 0.644 | 4.39E-24 | 13_Angio-TAM |
| TRAM1 | 1.644033399 | 0.522 | 0.255 | 4.42E-24 | 13_Angio-TAM |
| BAG1 | 1.686199342 | 0.41 | 0.169 | 4.73E-24 | 13_Angio-TAM |
| RNF11 | 1.748169146 | 0.371 | 0.142 | 5.45E-24 | 13_Angio-TAM |
| CTSL | -2.562347996 | 0.108 | 0.47 | 5.67E-24 | 13_Angio-TAM |
| NOP58 | 1.624615998 | 0.466 | 0.208 | 8.48E-24 | 13_Angio-TAM |
| GK | -1.845785216 | 0.231 | 0.592 | 1.65E-23 | 13_Angio-TAM |
| RABGAP1L | 1.843520214 | 0.657 | 0.424 | 1.86E-23 | 13_Angio-TAM |
| CNOT6L | 1.622879442 | 0.414 | 0.168 | 2.25E-23 | 13_Angio-TAM |
| N4BP2 | 1.848496355 | 0.311 | 0.105 | 2.67E-23 | 13_Angio-TAM |
| IL1RAP | -2.466506708 | 0.108 | 0.467 | 2.81E-23 | 13_Angio-TAM |
| CREB3L2 | 1.562733415 | 0.438 | 0.187 | 3.07E-23 | 13_Angio-TAM |
| SPTLC2 | -1.945862719 | 0.291 | 0.615 | 3.30E-23 | 13_Angio-TAM |
| TXN | 1.509811715 | 0.562 | 0.292 | 3.82E-23 | 13_Angio-TAM |
| GUK1 | 1.192726229 | 0.693 | 0.426 | 6.06E-23 | 13_Angio-TAM |
| ARHGAP24 | -1.435868307 | 0.398 | 0.708 | 6.53E-23 | 13_Angio-TAM |

| | | | | | |
|---|---|---|---|---|---|
| EPB41L3 | -2.112378839 | 0.171 | 0.531 | 7.20E-23 | 13_Angio-TAM |
| SNHG29 | 1.225316308 | 0.701 | 0.429 | 9.76E-23 | 13_Angio-TAM |
| SOD2 | -1.82141101 | 0.382 | 0.677 | 1.03E-22 | 13_Angio-TAM |
| SEC11C | 1.960909949 | 0.307 | 0.105 | 1.46E-22 | 13_Angio-TAM |
| SF3B5 | 1.4237446 | 0.482 | 0.226 | 1.74E-22 | 13_Angio-TAM |
| ADAM28 | -1.577361478 | 0.183 | 0.561 | 2.02E-22 | 13_Angio-TAM |
| PPARD | -1.967622171 | 0.151 | 0.506 | 2.12E-22 | 13_Angio-TAM |
| GRASP | 1.133256123 | 0.813 | 0.591 | 2.39E-22 | 13_Angio-TAM |
| SELENOS | 1.46435298 | 0.502 | 0.246 | 3.80E-22 | 13_Angio-TAM |
| SLC25A37 | -2.043233348 | 0.147 | 0.511 | 3.90E-22 | 13_Angio-TAM |
| KLF13 | 1.720370885 | 0.375 | 0.149 | 4.04E-22 | 13_Angio-TAM |
| MDN1 | 1.983805752 | 0.319 | 0.112 | 4.58E-22 | 13_Angio-TAM |
| ARID3A | 1.849654819 | 0.363 | 0.143 | 4.67E-22 | 13_Angio-TAM |
| ABR | -1.767998132 | 0.295 | 0.623 | 7.74E-22 | 13_Angio-TAM |
| ITGAX | -1.769172899 | 0.203 | 0.567 | 8.61E-22 | 13_Angio-TAM |
| FBRSL1 | 1.738516665 | 0.359 | 0.138 | 9.99E-22 | 13_Angio-TAM |
| NHSL1 | -1.754375378 | 0.183 | 0.553 | 1.06E-21 | 13_Angio-TAM |
| LDHB | 1.353165566 | 0.514 | 0.256 | 1.38E-21 | 13_Angio-TAM |
| EIF3F | 1.191831739 | 0.602 | 0.326 | 1.42E-21 | 13_Angio-TAM |
| LRP1 | -3.208818797 | 0.064 | 0.397 | 1.43E-21 | 13_Angio-TAM |
| IGFLR1 | 1.660000984 | 0.438 | 0.202 | 1.80E-21 | 13_Angio-TAM |
| PLTP | -2.558126484 | 0.159 | 0.494 | 2.46E-21 | 13_Angio-TAM |
| HTRA1 | -2.458691748 | 0.12 | 0.456 | 2.47E-21 | 13_Angio-TAM |
| CD47 | 1.695906752 | 0.41 | 0.18 | 2.59E-21 | 13_Angio-TAM |
| IGF2R | 1.708039936 | 0.522 | 0.267 | 2.77E-21 | 13_Angio-TAM |
| RNF138 | 1.636135634 | 0.343 | 0.128 | 3.11E-21 | 13_Angio-TAM |
| RIN3 | -1.578694279 | 0.227 | 0.58 | 3.71E-21 | 13_Angio-TAM |
| KPNA2 | 1.800349908 | 0.339 | 0.13 | 4.70E-21 | 13_Angio-TAM |
| PDE3B | -2.120916673 | 0.195 | 0.542 | 9.10E-21 | 13_Angio-TAM |
| DSE | -1.796314838 | 0.271 | 0.59 | 9.28E-21 | 13_Angio-TAM |
| PRKCH | -1.695064397 | 0.259 | 0.594 | 9.83E-21 | 13_Angio-TAM |
| CDK2AP2 | 1.484806541 | 0.367 | 0.147 | 1.20E-20 | 13_Angio-TAM |
| SPI1 | -1.205588339 | 0.307 | 0.647 | 2.04E-20 | 13_Angio-TAM |
| PADI2 | -1.970110992 | 0.203 | 0.531 | 2.28E-20 | 13_Angio-TAM |
| SLC9A9 | -2.235422099 | 0.127 | 0.455 | 3.30E-20 | 13_Angio-TAM |
| RALA | 1.380466149 | 0.49 | 0.235 | 3.32E-20 | 13_Angio-TAM |
| SSR2 | 1.155841417 | 0.57 | 0.309 | 3.34E-20 | 13_Angio-TAM |
| SORL1 | -1.765649543 | 0.223 | 0.562 | 3.85E-20 | 13_Angio-TAM |
| SLC20A1 | 1.416481831 | 0.454 | 0.21 | 3.96E-20 | 13_Angio-TAM |
| STMN1 | 1.539500302 | 0.363 | 0.146 | 4.32E-20 | 13_Angio-TAM |
| RASAL2 | -2.604625038 | 0.076 | 0.409 | 4.58E-20 | 13_Angio-TAM |
| FYB1 | -1.500026984 | 0.295 | 0.605 | 4.86E-20 | 13_Angio-TAM |
| COX7A2L | 1.610045492 | 0.414 | 0.192 | 5.29E-20 | 13_Angio-TAM |
| TCF12 | -1.460206844 | 0.398 | 0.693 | 6.39E-20 | 13_Angio-TAM |
| AC138123.1 | 1.486051446 | 0.382 | 0.164 | 1.26E-19 | 13_Angio-TAM |
| BANP | 1.598477128 | 0.49 | 0.243 | 1.49E-19 | 13_Angio-TAM |

| | | | | | |
|---|---|---|---|---|---|
| DOCK8 | -1.14332512 | 0.394 | 0.719 | 1.51E-19 | 13_Angio-TAM |
| SEC61G | 1.266825742 | 0.53 | 0.28 | 1.56E-19 | 13_Angio-TAM |
| CSGALNACT1 | -1.993751759 | 0.219 | 0.532 | 1.59E-19 | 13_Angio-TAM |
| ITGB2 | -1.30127176 | 0.343 | 0.639 | 1.65E-19 | 13_Angio-TAM |
| ATG7 | -1.911821672 | 0.207 | 0.532 | 2.14E-19 | 13_Angio-TAM |
| TGFB1 | 1.057563722 | 0.765 | 0.558 | 2.48E-19 | 13_Angio-TAM |
| AP3S1 | 1.873642497 | 0.351 | 0.144 | 2.52E-19 | 13_Angio-TAM |
| ANXA5 | -1.218215323 | 0.311 | 0.641 | 3.06E-19 | 13_Angio-TAM |
| ABL2 | -2.139129645 | 0.195 | 0.511 | 3.63E-19 | 13_Angio-TAM |
| RBM3 | 1.168659419 | 0.649 | 0.411 | 4.07E-19 | 13_Angio-TAM |
| RABGEF1 | -1.894778882 | 0.287 | 0.574 | 4.54E-19 | 13_Angio-TAM |
| ITGAV | -1.703229729 | 0.223 | 0.546 | 4.66E-19 | 13_Angio-TAM |
| CPVL | -1.78152965 | 0.183 | 0.514 | 5.35E-19 | 13_Angio-TAM |
| PLIN2 | -1.63758246 | 0.219 | 0.533 | 5.70E-19 | 13_Angio-TAM |
| IL18 | -1.619500584 | 0.183 | 0.515 | 5.82E-19 | 13_Angio-TAM |
| MANBA | -1.377246848 | 0.394 | 0.682 | 7.07E-19 | 13_Angio-TAM |
| PELI1 | -1.612858256 | 0.355 | 0.653 | 8.26E-19 | 13_Angio-TAM |
| BMP2K | -2.046793396 | 0.219 | 0.526 | 8.52E-19 | 13_Angio-TAM |
| MFSD1 | -1.170556907 | 0.287 | 0.611 | 9.06E-19 | 13_Angio-TAM |
| SERPINA1 | -2.608858075 | 0.12 | 0.425 | 1.20E-18 | 13_Angio-TAM |
| CARD11 | 1.522272082 | 0.482 | 0.241 | 1.25E-18 | 13_Angio-TAM |
| CSF3R | -1.423664222 | 0.187 | 0.532 | 1.46E-18 | 13_Angio-TAM |
| CD86 | -1.159828228 | 0.267 | 0.626 | 1.50E-18 | 13_Angio-TAM |
| CD81 | -1.442655809 | 0.386 | 0.659 | 1.57E-18 | 13_Angio-TAM |
| SBF2 | -1.829633245 | 0.207 | 0.52 | 1.70E-18 | 13_Angio-TAM |
| PDE4DIP | -1.71351432 | 0.155 | 0.491 | 2.49E-18 | 13_Angio-TAM |
| MED26 | 1.876071422 | 0.351 | 0.145 | 2.61E-18 | 13_Angio-TAM |
| DENND3 | -1.709933479 | 0.155 | 0.486 | 2.78E-18 | 13_Angio-TAM |
| FRMD4B | -1.606920996 | 0.191 | 0.522 | 3.33E-18 | 13_Angio-TAM |
| APLP2 | -1.690655906 | 0.227 | 0.544 | 3.81E-18 | 13_Angio-TAM |
| RHBDF2 | -1.173726869 | 0.359 | 0.662 | 4.47E-18 | 13_Angio-TAM |
| RAB20 | -1.198642451 | 0.263 | 0.599 | 4.64E-18 | 13_Angio-TAM |
| ODC1 | 1.571864983 | 0.371 | 0.159 | 5.37E-18 | 13_Angio-TAM |
| EIF3L | 1.180652885 | 0.506 | 0.265 | 7.38E-18 | 13_Angio-TAM |
| FPR1 | -1.552743354 | 0.195 | 0.534 | 7.45E-18 | 13_Angio-TAM |
| MITF | -2.400001739 | 0.104 | 0.408 | 7.46E-18 | 13_Angio-TAM |
| SLC12A7 | 1.818731143 | 0.347 | 0.143 | 1.25E-17 | 13_Angio-TAM |
| IRF2BP2 | 1.43870058 | 0.53 | 0.303 | 1.33E-17 | 13_Angio-TAM |
| XYLT1 | 1.697525535 | 0.454 | 0.225 | 1.94E-17 | 13_Angio-TAM |
| RPS17 | 1.121434086 | 0.598 | 0.359 | 2.12E-17 | 13_Angio-TAM |
| CD63 | -1.188919393 | 0.438 | 0.679 | 2.16E-17 | 13_Angio-TAM |
| BHLHE40 | 1.218268637 | 0.606 | 0.361 | 2.65E-17 | 13_Angio-TAM |
| GPR34 | -2.014812876 | 0.124 | 0.438 | 2.97E-17 | 13_Angio-TAM |
| G3BP2 | 1.582751628 | 0.414 | 0.202 | 3.32E-17 | 13_Angio-TAM |
| CDK17 | 1.612162647 | 0.41 | 0.19 | 3.66E-17 | 13_Angio-TAM |
| NPL | -1.833902042 | 0.108 | 0.421 | 3.83E-17 | 13_Angio-TAM |

| | | | | | |
|---|---|---|---|---|---|
| USP53 | -1.76691602 | 0.215 | 0.509 | 4.23E-17 | 13_Angio-TAM |
| ZNF706 | 1.340895644 | 0.534 | 0.305 | 4.79E-17 | 13_Angio-TAM |
| SEPTIN9 | 1.391409477 | 0.402 | 0.186 | 5.10E-17 | 13_Angio-TAM |
| PI4KA | 1.643211824 | 0.47 | 0.246 | 6.55E-17 | 13_Angio-TAM |
| SLC2A5 | -1.924383741 | 0.116 | 0.429 | 6.72E-17 | 13_Angio-TAM |
| TMEM176B | -2.730020491 | 0.072 | 0.365 | 7.34E-17 | 13_Angio-TAM |
| SLC16A10 | -3.852420412 | 0.12 | 0.397 | 8.86E-17 | 13_Angio-TAM |
| RPL23 | 1.053702754 | 0.669 | 0.464 | 1.23E-16 | 13_Angio-TAM |
| CAMK2D | -2.441448178 | 0.056 | 0.349 | 1.33E-16 | 13_Angio-TAM |
| BCL2 | -1.727594019 | 0.179 | 0.486 | 1.52E-16 | 13_Angio-TAM |
| SOAT1 | -2.185076477 | 0.12 | 0.418 | 1.68E-16 | 13_Angio-TAM |
| ATP8B4 | -1.625493634 | 0.171 | 0.473 | 1.77E-16 | 13_Angio-TAM |
| ST6GALNAC3 | -2.504367624 | 0.088 | 0.385 | 2.21E-16 | 13_Angio-TAM |
| ATF6 | -2.05766951 | 0.303 | 0.577 | 2.26E-16 | 13_Angio-TAM |
| MALT1 | 1.123452913 | 0.685 | 0.443 | 2.46E-16 | 13_Angio-TAM |
| FNDC3B | -1.842743734 | 0.291 | 0.567 | 4.17E-16 | 13_Angio-TAM |
| FNIP2 | -1.807705491 | 0.251 | 0.545 | 4.30E-16 | 13_Angio-TAM |
| TNS3 | -1.611263814 | 0.199 | 0.501 | 5.17E-16 | 13_Angio-TAM |
| HIF1A-AS3 | -2.915068163 | 0.084 | 0.368 | 6.07E-16 | 13_Angio-TAM |
| ETS2 | -1.130398386 | 0.359 | 0.648 | 6.18E-16 | 13_Angio-TAM |
| CAMK1D | -1.533543712 | 0.247 | 0.539 | 6.25E-16 | 13_Angio-TAM |
| HAVCR2 | -1.399811321 | 0.243 | 0.545 | 7.18E-16 | 13_Angio-TAM |
| UXT | 1.381091185 | 0.454 | 0.241 | 7.28E-16 | 13_Angio-TAM |
| PIK3R1 | -1.471747719 | 0.283 | 0.575 | 8.55E-16 | 13_Angio-TAM |
| PDK4 | -1.667236596 | 0.223 | 0.517 | 9.51E-16 | 13_Angio-TAM |
| ADAP2 | -1.551752625 | 0.151 | 0.462 | 1.12E-15 | 13_Angio-TAM |
| RASSF4 | -1.97810963 | 0.163 | 0.445 | 1.20E-15 | 13_Angio-TAM |
| AOAH | -1.61296438 | 0.155 | 0.461 | 1.42E-15 | 13_Angio-TAM |
| FAM110B | -4.385092439 | 0.016 | 0.279 | 1.50E-15 | 13_Angio-TAM |
| LNCAROD | -2.347447358 | 0.159 | 0.449 | 1.51E-15 | 13_Angio-TAM |
| SCIN | -2.921776155 | 0.052 | 0.328 | 2.36E-15 | 13_Angio-TAM |
| PTPRJ | -1.493599116 | 0.299 | 0.569 | 3.16E-15 | 13_Angio-TAM |
| ABCC4 | -2.124692057 | 0.076 | 0.359 | 3.35E-15 | 13_Angio-TAM |
| RHOQ | -1.634522853 | 0.187 | 0.478 | 3.82E-15 | 13_Angio-TAM |
| VASH1 | -1.239156314 | 0.171 | 0.477 | 3.97E-15 | 13_Angio-TAM |
| ARHGAP22 | -2.078284797 | 0.084 | 0.374 | 4.00E-15 | 13_Angio-TAM |
| IRAK2 | -2.058170587 | 0.171 | 0.452 | 4.21E-15 | 13_Angio-TAM |
| KDM5A | 1.189533811 | 0.482 | 0.252 | 4.37E-15 | 13_Angio-TAM |
| UBE2J1 | 1.294799674 | 0.442 | 0.226 | 5.40E-15 | 13_Angio-TAM |
| B3GNT5 | -1.221274303 | 0.239 | 0.541 | 5.48E-15 | 13_Angio-TAM |
| AIF1 | -1.22732724 | 0.371 | 0.626 | 7.68E-15 | 13_Angio-TAM |
| APOC2 | -2.28827891 | 0.167 | 0.444 | 9.56E-15 | 13_Angio-TAM |
| ZHX2 | 1.220700269 | 0.49 | 0.26 | 1.24E-14 | 13_Angio-TAM |
| SDK1 | -1.379694964 | 0.151 | 0.445 | 1.27E-14 | 13_Angio-TAM |
| VSIR | -1.491888806 | 0.263 | 0.531 | 1.70E-14 | 13_Angio-TAM |
| TPRG1 | -3.447505908 | 0.048 | 0.308 | 1.70E-14 | 13_Angio-TAM |

| | | | | | |
|---|---|---|---|---|---|
| RIN2 | -2.111267685 | 0.092 | 0.369 | 1.73E-14 | 13_Angio-TAM |
| WDFY3 | -2.325827609 | 0.076 | 0.353 | 1.78E-14 | 13_Angio-TAM |
| ARRB2 | -1.308544926 | 0.271 | 0.555 | 1.79E-14 | 13_Angio-TAM |
| SRSF3 | 1.006694209 | 0.669 | 0.443 | 1.88E-14 | 13_Angio-TAM |
| ST6GAL1 | -1.376616088 | 0.243 | 0.53 | 2.22E-14 | 13_Angio-TAM |
| PPARG | -2.875104955 | 0.052 | 0.317 | 2.46E-14 | 13_Angio-TAM |
| IRS2 | -1.022194598 | 0.175 | 0.468 | 2.51E-14 | 13_Angio-TAM |
| VMP1 | -1.243389146 | 0.323 | 0.61 | 2.89E-14 | 13_Angio-TAM |
| AP2A2 | -2.246656054 | 0.096 | 0.372 | 3.35E-14 | 13_Angio-TAM |
| NRP2 | -2.830457318 | 0.084 | 0.348 | 3.56E-14 | 13_Angio-TAM |
| ADGRE5 | 1.014884354 | 0.542 | 0.302 | 3.57E-14 | 13_Angio-TAM |
| DOCK10 | -1.371288775 | 0.247 | 0.543 | 4.11E-14 | 13_Angio-TAM |
| BCL3 | 1.33283334 | 0.378 | 0.177 | 4.30E-14 | 13_Angio-TAM |
| HBEGF | -2.343072557 | 0.12 | 0.388 | 4.38E-14 | 13_Angio-TAM |
| CREG1 | -1.449171758 | 0.151 | 0.442 | 5.94E-14 | 13_Angio-TAM |
| CTNNB1 | -1.103456285 | 0.402 | 0.646 | 6.07E-14 | 13_Angio-TAM |
| SIVA1 | 1.3405358 | 0.434 | 0.231 | 6.84E-14 | 13_Angio-TAM |
| NFIC | -1.734890126 | 0.147 | 0.429 | 7.59E-14 | 13_Angio-TAM |
| NLRP3 | -1.285767933 | 0.303 | 0.578 | 7.67E-14 | 13_Angio-TAM |
| GPNMB | -2.740418963 | 0.056 | 0.316 | 8.18E-14 | 13_Angio-TAM |
| KLF2 | -1.448959953 | 0.303 | 0.568 | 1.25E-13 | 13_Angio-TAM |
| CEP170 | -1.362317471 | 0.295 | 0.556 | 1.74E-13 | 13_Angio-TAM |
| LHFPL2 | -1.202360856 | 0.347 | 0.606 | 1.86E-13 | 13_Angio-TAM |
| TUT7 | -1.444429118 | 0.171 | 0.454 | 2.32E-13 | 13_Angio-TAM |
| RAPH1 | -2.018844859 | 0.127 | 0.396 | 2.49E-13 | 13_Angio-TAM |
| DENND1A | -1.692234857 | 0.112 | 0.389 | 2.57E-13 | 13_Angio-TAM |
| MYO1F | -1.733331832 | 0.147 | 0.416 | 3.15E-13 | 13_Angio-TAM |
| MAP2K3 | 1.191560377 | 0.506 | 0.296 | 3.26E-13 | 13_Angio-TAM |
| PHC2 | -1.816561894 | 0.139 | 0.412 | 3.41E-13 | 13_Angio-TAM |
| IFNGR2 | -1.092909845 | 0.351 | 0.635 | 3.77E-13 | 13_Angio-TAM |
| ARHGAP6 | -1.474957435 | 0.1 | 0.379 | 4.61E-13 | 13_Angio-TAM |
| LTC4S | -2.077096973 | 0.116 | 0.381 | 4.80E-13 | 13_Angio-TAM |
| FCGBP | -1.837189502 | 0.155 | 0.424 | 5.43E-13 | 13_Angio-TAM |
| PLD3 | -1.882882703 | 0.112 | 0.378 | 5.52E-13 | 13_Angio-TAM |
| BASP1 | -1.527392661 | 0.311 | 0.574 | 5.67E-13 | 13_Angio-TAM |
| AHI1 | 1.287174739 | 0.414 | 0.21 | 5.98E-13 | 13_Angio-TAM |
| SPRED1 | -2.138756347 | 0.068 | 0.328 | 6.96E-13 | 13_Angio-TAM |
| C3AR1 | -1.570966696 | 0.143 | 0.419 | 8.36E-13 | 13_Angio-TAM |
| CPEB4 | -1.838129101 | 0.163 | 0.432 | 8.67E-13 | 13_Angio-TAM |
| TSC22D2 | -1.610606283 | 0.303 | 0.561 | 9.23E-13 | 13_Angio-TAM |
| CCL3L1 | -2.346942299 | 0.183 | 0.436 | 1.04E-12 | 13_Angio-TAM |
| MPP1 | -1.786715977 | 0.116 | 0.382 | 1.24E-12 | 13_Angio-TAM |
| SOCS6 | -2.166544218 | 0.112 | 0.371 | 1.30E-12 | 13_Angio-TAM |
| TREM1 | -2.934856872 | 0.06 | 0.301 | 1.34E-12 | 13_Angio-TAM |
| GBP2 | -2.472600577 | 0.084 | 0.336 | 1.42E-12 | 13_Angio-TAM |
| GPCPD1 | -1.692669941 | 0.239 | 0.495 | 1.46E-12 | 13_Angio-TAM |

| | | | | | |
|---|---|---|---|---|---|
| MS4A4A | -1.48606143 | 0.259 | 0.509 | 1.50E-12 | 13_Angio-TAM |
| MGAT1 | -1.218739567 | 0.347 | 0.596 | 1.81E-12 | 13_Angio-TAM |
| ENG | -2.806039307 | 0.072 | 0.312 | 2.12E-12 | 13_Angio-TAM |
| MAT2A | -1.047338387 | 0.363 | 0.613 | 2.64E-12 | 13_Angio-TAM |
| ATAD2B | 1.263583915 | 0.43 | 0.222 | 2.84E-12 | 13_Angio-TAM |
| PLA2G4A | -2.008114279 | 0.131 | 0.389 | 2.89E-12 | 13_Angio-TAM |
| ITSN1 | -1.753090316 | 0.143 | 0.405 | 2.98E-12 | 13_Angio-TAM |
| CSNK1G2 | 1.235350608 | 0.41 | 0.207 | 3.08E-12 | 13_Angio-TAM |
| DMXL2 | -2.648857709 | 0.092 | 0.334 | 3.19E-12 | 13_Angio-TAM |
| PARP14 | -1.967341301 | 0.12 | 0.378 | 3.40E-12 | 13_Angio-TAM |
| ADAM17 | -1.416333041 | 0.263 | 0.528 | 3.68E-12 | 13_Angio-TAM |
| CADM1 | -2.742434452 | 0.064 | 0.305 | 4.18E-12 | 13_Angio-TAM |
| TMIGD3 | -1.559059562 | 0.116 | 0.382 | 4.44E-12 | 13_Angio-TAM |
| CD84 | -1.512239631 | 0.131 | 0.397 | 4.88E-12 | 13_Angio-TAM |
| LAT2 | -2.280742911 | 0.08 | 0.325 | 5.74E-12 | 13_Angio-TAM |
| GLDN | -2.57201711 | 0.068 | 0.313 | 5.80E-12 | 13_Angio-TAM |
| RGS10 | -1.191714493 | 0.339 | 0.576 | 6.39E-12 | 13_Angio-TAM |
| NEDD9 | -1.301845344 | 0.263 | 0.516 | 7.06E-12 | 13_Angio-TAM |
| L3MBTL4 | -2.24383165 | 0.052 | 0.296 | 7.53E-12 | 13_Angio-TAM |
| COX7A2 | 1.052303295 | 0.526 | 0.321 | 7.70E-12 | 13_Angio-TAM |
| KLF7 | -1.763179347 | 0.131 | 0.384 | 8.45E-12 | 13_Angio-TAM |
| CSTB | -1.531656199 | 0.227 | 0.491 | 9.09E-12 | 13_Angio-TAM |
| LINC01619 | -2.155404215 | 0.1 | 0.35 | 9.60E-12 | 13_Angio-TAM |
| MIR181A1HG | -1.428447993 | 0.211 | 0.479 | 1.00E-11 | 13_Angio-TAM |
| SIGLEC10 | -1.733690886 | 0.1 | 0.352 | 1.04E-11 | 13_Angio-TAM |
| OGFRL1 | -1.128243463 | 0.287 | 0.543 | 1.07E-11 | 13_Angio-TAM |
| ATP13A3 | -1.561033055 | 0.247 | 0.495 | 1.50E-11 | 13_Angio-TAM |
| SELENOF | 1.0402684 | 0.458 | 0.258 | 1.50E-11 | 13_Angio-TAM |
| SERPINE1 | -1.900681659 | 0.243 | 0.469 | 1.68E-11 | 13_Angio-TAM |
| SRGAP2C | -1.270866169 | 0.179 | 0.449 | 1.75E-11 | 13_Angio-TAM |
| KCTD12 | -1.563809196 | 0.175 | 0.428 | 1.89E-11 | 13_Angio-TAM |
| SDCCAG8 | -1.283143285 | 0.247 | 0.513 | 1.91E-11 | 13_Angio-TAM |
| MAML3 | -1.044066953 | 0.363 | 0.611 | 2.33E-11 | 13_Angio-TAM |
| SLC43A2 | -1.439049679 | 0.155 | 0.411 | 2.43E-11 | 13_Angio-TAM |
| SLC31A2 | -1.131538117 | 0.203 | 0.469 | 2.65E-11 | 13_Angio-TAM |
| FAM102B | -1.986213843 | 0.112 | 0.356 | 3.03E-11 | 13_Angio-TAM |
| MACF1 | -1.183779657 | 0.299 | 0.548 | 3.26E-11 | 13_Angio-TAM |
| ATF3 | -1.192389454 | 0.414 | 0.629 | 3.43E-11 | 13_Angio-TAM |
| WDFY4 | 1.163090428 | 0.466 | 0.266 | 3.62E-11 | 13_Angio-TAM |
| ACER3 | -1.679675087 | 0.147 | 0.399 | 3.66E-11 | 13_Angio-TAM |
| PLK3 | -1.327756734 | 0.207 | 0.466 | 3.81E-11 | 13_Angio-TAM |
| RAPGEF2 | 1.198000716 | 0.578 | 0.371 | 3.85E-11 | 13_Angio-TAM |
| EPS8 | -1.654749465 | 0.131 | 0.385 | 5.34E-11 | 13_Angio-TAM |
| ARL15 | -2.005792943 | 0.124 | 0.364 | 6.16E-11 | 13_Angio-TAM |
| OLFML3 | -2.339031951 | 0.064 | 0.298 | 6.39E-11 | 13_Angio-TAM |
| GPRIN3 | -1.748379397 | 0.116 | 0.363 | 7.14E-11 | 13_Angio-TAM |

| | | | | | |
|---|---|---|---|---|---|
| LRCH1 | -1.968391436 | 0.135 | 0.377 | 8.54E-11 | 13_Angio-TAM |
| FCGR1B | -4.099410614 | 0.012 | 0.219 | 8.64E-11 | 13_Angio-TAM |
| GAA | -1.39933494 | 0.139 | 0.393 | 9.77E-11 | 13_Angio-TAM |
| DTNA | -2.749814462 | 0.064 | 0.29 | 1.05E-10 | 13_Angio-TAM |
| AKR1B1 | -1.643577756 | 0.139 | 0.381 | 1.19E-10 | 13_Angio-TAM |
| OTUD1 | -2.192129054 | 0.1 | 0.333 | 1.56E-10 | 13_Angio-TAM |
| DISC1 | -1.446384348 | 0.187 | 0.437 | 1.61E-10 | 13_Angio-TAM |
| AL163541.1 | -2.358793335 | 0.08 | 0.308 | 1.80E-10 | 13_Angio-TAM |
| NUMB | -1.321199884 | 0.251 | 0.502 | 2.01E-10 | 13_Angio-TAM |
| CFD | -3.040411679 | 0.06 | 0.277 | 2.03E-10 | 13_Angio-TAM |
| KIF1B | -1.688535567 | 0.195 | 0.43 | 2.11E-10 | 13_Angio-TAM |
| TRIO | -1.315515183 | 0.291 | 0.53 | 2.45E-10 | 13_Angio-TAM |
| CH25H | -3.015231938 | 0.116 | 0.334 | 2.84E-10 | 13_Angio-TAM |
| PAPOLG | -2.065844929 | 0.12 | 0.349 | 3.45E-10 | 13_Angio-TAM |
| IGSF21 | -2.128164444 | 0.064 | 0.287 | 4.94E-10 | 13_Angio-TAM |
| LRRK2 | -3.049889728 | 0.044 | 0.257 | 4.95E-10 | 13_Angio-TAM |
| SLC4A7 | -1.57326042 | 0.231 | 0.464 | 6.01E-10 | 13_Angio-TAM |
| DST | -1.491896018 | 0.175 | 0.416 | 6.01E-10 | 13_Angio-TAM |
| CTTNBP2 | -1.855843652 | 0.072 | 0.299 | 6.55E-10 | 13_Angio-TAM |
| IL13RA1 | -1.194692106 | 0.239 | 0.486 | 6.69E-10 | 13_Angio-TAM |
| SAP30 | -1.903539465 | 0.12 | 0.35 | 7.51E-10 | 13_Angio-TAM |
| SUSD6 | -1.261453571 | 0.343 | 0.559 | 8.01E-10 | 13_Angio-TAM |
| MKNK1 | -1.853211801 | 0.112 | 0.337 | 8.10E-10 | 13_Angio-TAM |
| LY86 | -1.52663953 | 0.159 | 0.395 | 8.26E-10 | 13_Angio-TAM |
| BCAT1 | -2.110337752 | 0.088 | 0.31 | 1.12E-09 | 13_Angio-TAM |
| C9orf72 | -1.08068929 | 0.319 | 0.55 | 1.17E-09 | 13_Angio-TAM |
| RASA3 | -1.579937897 | 0.195 | 0.426 | 1.54E-09 | 13_Angio-TAM |
| DAGLB | -1.39168057 | 0.167 | 0.409 | 1.60E-09 | 13_Angio-TAM |
| IPCEF1 | -2.405230557 | 0.096 | 0.31 | 1.81E-09 | 13_Angio-TAM |
| RFX2 | -1.54176771 | 0.167 | 0.396 | 1.91E-09 | 13_Angio-TAM |
| VPS37B | -1.386274649 | 0.131 | 0.376 | 1.97E-09 | 13_Angio-TAM |
| P4HA1 | -1.685597797 | 0.195 | 0.424 | 2.10E-09 | 13_Angio-TAM |
| PRKN | -2.135890602 | 0.084 | 0.297 | 2.43E-09 | 13_Angio-TAM |
| 3-Mar | -1.575836392 | 0.179 | 0.408 | 2.89E-09 | 13_Angio-TAM |
| FCHO2 | -2.221266254 | 0.072 | 0.283 | 3.37E-09 | 13_Angio-TAM |
| MB21D2 | -1.644214821 | 0.096 | 0.319 | 3.55E-09 | 13_Angio-TAM |
| MSN | -1.162471401 | 0.307 | 0.54 | 3.81E-09 | 13_Angio-TAM |
| TTC7B | -2.571470219 | 0.036 | 0.237 | 3.86E-09 | 13_Angio-TAM |
| SWAP70 | -1.009288706 | 0.203 | 0.451 | 4.05E-09 | 13_Angio-TAM |
| PDXK | -1.600961102 | 0.171 | 0.395 | 4.47E-09 | 13_Angio-TAM |
| ALOX5 | -1.034772557 | 0.219 | 0.471 | 4.49E-09 | 13_Angio-TAM |
| TSPO | -1.060477467 | 0.247 | 0.474 | 5.05E-09 | 13_Angio-TAM |
| MMP19 | -2.667054037 | 0.072 | 0.276 | 6.88E-09 | 13_Angio-TAM |
| TBC1D12 | -1.882693288 | 0.092 | 0.306 | 7.07E-09 | 13_Angio-TAM |
| CD58 | -1.208170285 | 0.191 | 0.433 | 7.08E-09 | 13_Angio-TAM |
| STX4 | -2.129582217 | 0.116 | 0.332 | 7.44E-09 | 13_Angio-TAM |

| | | | | | |
|---|---|---|---|---|---|
| FAM135A | -2.080201129 | 0.076 | 0.287 | 8.04E-09 | 13_Angio-TAM |
| FHIT | -1.036506382 | 0.327 | 0.545 | 8.87E-09 | 13_Angio-TAM |
| VEGFA | -1.764451282 | 0.171 | 0.389 | 9.91E-09 | 13_Angio-TAM |
| FGL2 | -1.481508159 | 0.131 | 0.361 | 1.09E-08 | 13_Angio-TAM |
| SIRPA | -1.46657667 | 0.135 | 0.364 | 1.31E-08 | 13_Angio-TAM |
| LINC02798 | -1.729379801 | 0.068 | 0.278 | 1.40E-08 | 13_Angio-TAM |
| LINC00963 | -2.024974763 | 0.052 | 0.257 | 1.63E-08 | 13_Angio-TAM |
| BHLHE41 | -1.812380831 | 0.108 | 0.322 | 1.69E-08 | 13_Angio-TAM |
| UBXN2B | -2.186897705 | 0.068 | 0.272 | 1.96E-08 | 13_Angio-TAM |
| MTHFD1L | -1.203692407 | 0.175 | 0.41 | 2.17E-08 | 13_Angio-TAM |
| DLEU2 | -1.07767103 | 0.339 | 0.554 | 2.81E-08 | 13_Angio-TAM |
| LPIN2 | -1.67571076 | 0.131 | 0.344 | 3.04E-08 | 13_Angio-TAM |
| AL078590.2 | -1.671260784 | 0.151 | 0.361 | 3.62E-08 | 13_Angio-TAM |
| CTTNBP2NL | -1.551939095 | 0.151 | 0.37 | 3.63E-08 | 13_Angio-TAM |
| MNDA | -2.15360532 | 0.08 | 0.28 | 4.30E-08 | 13_Angio-TAM |
| GNB4 | -1.280598184 | 0.251 | 0.466 | 4.42E-08 | 13_Angio-TAM |
| IER2 | -1.120231496 | 0.442 | 0.645 | 4.91E-08 | 13_Angio-TAM |
| FOLR2 | -1.769148873 | 0.108 | 0.315 | 5.02E-08 | 13_Angio-TAM |
| PTK2B | -1.110581094 | 0.267 | 0.486 | 5.16E-08 | 13_Angio-TAM |
| TNFAIP2 | -2.037369327 | 0.1 | 0.306 | 5.17E-08 | 13_Angio-TAM |
| IVNS1ABP | -1.305551317 | 0.183 | 0.403 | 5.28E-08 | 13_Angio-TAM |
| ARHGAP21 | -1.482295532 | 0.147 | 0.359 | 6.37E-08 | 13_Angio-TAM |
| DENND4C | -1.257486776 | 0.187 | 0.404 | 7.84E-08 | 13_Angio-TAM |
| YBX3 | -1.093793104 | 0.171 | 0.396 | 9.66E-08 | 13_Angio-TAM |
| EVL | -1.06000189 | 0.147 | 0.369 | 1.27E-07 | 13_Angio-TAM |
| ERGIC1 | -1.217543899 | 0.203 | 0.429 | 1.83E-07 | 13_Angio-TAM |
| ABHD12 | -1.184361536 | 0.207 | 0.427 | 1.95E-07 | 13_Angio-TAM |
| ATP6V0A1 | -1.244625801 | 0.112 | 0.318 | 1.96E-07 | 13_Angio-TAM |
| ARAP1 | -1.57986811 | 0.12 | 0.327 | 2.96E-07 | 13_Angio-TAM |
| LIMK2 | -1.802284085 | 0.104 | 0.304 | 2.97E-07 | 13_Angio-TAM |
| SERPINB6 | -1.320851359 | 0.139 | 0.35 | 3.02E-07 | 13_Angio-TAM |
| BCL2A1 | -1.094350751 | 0.227 | 0.444 | 3.23E-07 | 13_Angio-TAM |
| ANKS1A | -1.220915995 | 0.219 | 0.429 | 4.37E-07 | 13_Angio-TAM |
| NAIP | -1.4491025 | 0.124 | 0.326 | 5.52E-07 | 13_Angio-TAM |
| TBC1D16 | -1.254210212 | 0.171 | 0.379 | 5.52E-07 | 13_Angio-TAM |
| SLC2A3 | -1.008164091 | 0.331 | 0.545 | 5.67E-07 | 13_Angio-TAM |
| CHSY1 | -1.157748498 | 0.191 | 0.405 | 6.79E-07 | 13_Angio-TAM |
| WDR91 | -1.281244097 | 0.104 | 0.305 | 8.18E-07 | 13_Angio-TAM |
| ASAP1 | -1.269907267 | 0.255 | 0.456 | 1.07E-06 | 13_Angio-TAM |
| ATP6V1B2 | -1.109229809 | 0.267 | 0.47 | 2.60E-06 | 13_Angio-TAM |
| STX11 | -1.035467091 | 0.183 | 0.395 | 3.35E-06 | 13_Angio-TAM |
| TM6SF1 | -1.023223559 | 0.131 | 0.333 | 7.00E-06 | 13_Angio-TAM |
| RGCC | -1.071976762 | 0.267 | 0.468 | 7.69E-06 | 13_Angio-TAM |
| IER5L | 1.788235422 | 0.727 | 0.193 | 1.45E-06 | 14_Contaminant |
| CDC14B | 1.618136454 | 0.682 | 0.182 | 1.84E-05 | 14_Contaminant |
| PAPOLG | 1.834294219 | 0.864 | 0.348 | 2.24E-05 | 14_Contaminant |

| | | | | | |
|---|---|---|---|---|---|
| CPA6 | 2.471175418 | 0.545 | 0.134 | 2.65E-05 | 14_Contaminant |
| AC046195.1 | 2.210408628 | 0.5 | 0.106 | 2.75E-05 | 14_Contaminant |
| PLEKHG5 | 3.281301789 | 0.318 | 0.049 | 2.77E-05 | 14_Contaminant |
| TNFAIP8L3 | 1.986952278 | 0.636 | 0.176 | 4.42E-05 | 14_Contaminant |
| JDP2 | 1.707853959 | 1 | 0.548 | 5.57E-05 | 14_Contaminant |
| HABP4 | 3.04777961 | 0.273 | 0.039 | 0.00024011 | 14_Contaminant |
| FOXP2 | 2.25392166 | 0.409 | 0.082 | 0.00027498 | 14_Contaminant |
| LIMK2 | 1.937527363 | 0.773 | 0.303 | 0.00037678 | 14_Contaminant |
| CCDC26 | 1.964055233 | 0.545 | 0.139 | 0.00038802 | 14_Contaminant |
| OXR1 | 1.875996988 | 0.773 | 0.326 | 0.00080066 | 14_Contaminant |
| CLASP1 | 1.881052715 | 0.727 | 0.27 | 0.00090172 | 14_Contaminant |
| PDE8A | 1.397045343 | 0.955 | 0.559 | 0.00153123 | 14_Contaminant |
| PLVAP | 2.11211804 | 0.455 | 0.109 | 0.00183646 | 14_Contaminant |
| USP15 | 1.224902952 | 1 | 0.609 | 0.00662941 | 14_Contaminant |
| RBM47 | 1.041271615 | 1 | 0.741 | 0.00877592 | 14_Contaminant |
| IPCEF1 | 1.675541239 | 0.727 | 0.309 | 0.00962045 | 14_Contaminant |
| PMEPA1 | 1.549809779 | 0.682 | 0.253 | 0.0108065 | 14_Contaminant |
| INPP5D | 1.165062237 | 0.909 | 0.506 | 0.01223369 | 14_Contaminant |
| MRC2 | 1.613436433 | 0.591 | 0.192 | 0.01285559 | 14_Contaminant |
| PLA2G4A | 1.541166545 | 0.818 | 0.387 | 0.01570988 | 14_Contaminant |
| IL1RAP | 1.420839128 | 0.909 | 0.465 | 0.01699973 | 14_Contaminant |
| VIM | -5.05254369 | 0.182 | 0.672 | 0.01707823 | 14_Contaminant |
| TTC7A | 1.231633197 | 0.864 | 0.395 | 0.0172844 | 14_Contaminant |
| SPRY1 | 1.820815336 | 0.545 | 0.169 | 0.01777366 | 14_Contaminant |
| UBN1 | 1.911473481 | 0.5 | 0.149 | 0.01892704 | 14_Contaminant |
| RASSF4 | 1.599288107 | 0.818 | 0.443 | 0.01941311 | 14_Contaminant |
| CSGALNACT1 | 1.11845935 | 1 | 0.53 | 0.02214541 | 14_Contaminant |
| MED26 | 1.799075824 | 0.5 | 0.146 | 0.02316645 | 14_Contaminant |
| KIFC3 | 1.862955739 | 0.545 | 0.186 | 0.03358243 | 14_Contaminant |
| FGD4 | 1.427081788 | 1 | 0.788 | 0.04000649 | 14_Contaminant |
| COX19 | 1.836039495 | 0.455 | 0.129 | 0.04196503 | 14_Contaminant |
| RANBP9 | 1.553606559 | 0.773 | 0.393 | 0.04388297 | 14_Contaminant |
| SPATS2L | 1.44251245 | 0.727 | 0.319 | 0.04773431 | 14_Contaminant |
| KIF1B | 1.237273811 | 0.864 | 0.429 | 0.04902727 | 14_Contaminant |
| MAP4K3 | 1.307256777 | 0.909 | 0.541 | 0.05029094 | 14_Contaminant |
| VPS37B | 1.343120535 | 0.773 | 0.375 | 0.05365756 | 14_Contaminant |
| DOCK4-AS1 | 2.198330874 | 0.273 | 0.052 | 0.06665097 | 14_Contaminant |
| GPATCH11 | 2.060514784 | 0.364 | 0.089 | 0.07272751 | 14_Contaminant |
| NFKBID | 1.478602273 | 0.864 | 0.539 | 0.07414636 | 14_Contaminant |
| MALT1 | 1.247548783 | 0.864 | 0.444 | 0.08585091 | 14_Contaminant |
| HEATR5B | 1.37508436 | 0.591 | 0.208 | 0.09385317 | 14_Contaminant |
| WDR91 | 1.534849678 | 0.682 | 0.303 | 0.09700054 | 14_Contaminant |
| RPL9 | -2.985702178 | 0.227 | 0.674 | 0.12696571 | 14_Contaminant |
| LGMN | 1.081889979 | 0.909 | 0.557 | 0.13630572 | 14_Contaminant |
| RPS3A | -3.001934565 | 0.364 | 0.724 | 0.13790398 | 14_Contaminant |
| MB21D2 | 1.337808052 | 0.727 | 0.317 | 0.19841628 | 14_Contaminant |

| | | | | | |
|---|---|---|---|---|---|
| RPS27 | -2.860283219 | 0.364 | 0.715 | 0.20250625 | 14_Contaminant |
| VCL | 2.11569129 | 0.409 | 0.119 | 0.20869249 | 14_Contaminant |
| STX17-AS1 | 1.366329444 | 0.727 | 0.346 | 0.22069837 | 14_Contaminant |
| KLF2 | 1.485232679 | 0.909 | 0.566 | 0.24908181 | 14_Contaminant |
| AKAP8L | 1.966329097 | 0.455 | 0.153 | 0.26108487 | 14_Contaminant |
| AC079015.1 | 1.787476035 | 0.455 | 0.143 | 0.26268558 | 14_Contaminant |
| PCNX2 | 1.228514464 | 0.636 | 0.25 | 0.26736338 | 14_Contaminant |
| RPL26 | -2.469885595 | 0.409 | 0.751 | 0.27096124 | 14_Contaminant |
| SRGAP3 | 1.596029317 | 0.5 | 0.17 | 0.2848543 | 14_Contaminant |
| MICU3 | 2.17586199 | 0.409 | 0.122 | 0.28671068 | 14_Contaminant |
| SOCS6 | 1.228750355 | 0.773 | 0.369 | 0.29507528 | 14_Contaminant |
| EPB41L2 | 1.027649907 | 0.955 | 0.648 | 0.29688607 | 14_Contaminant |
| RPL18A | -2.676301188 | 0.409 | 0.723 | 0.36589326 | 14_Contaminant |
| HBEGF | 1.062917658 | 0.818 | 0.387 | 0.38880057 | 14_Contaminant |
| RPLP0 | -2.600957913 | 0.364 | 0.724 | 0.39301819 | 14_Contaminant |
| DPYD-AS1 | 2.110703032 | 0.364 | 0.101 | 0.39326431 | 14_Contaminant |
| RPL39 | -2.520673286 | 0.364 | 0.721 | 0.40509052 | 14_Contaminant |
| LINC01374 | 1.376117942 | 0.682 | 0.296 | 0.45414645 | 14_Contaminant |
| ABCC4 | 1.286226987 | 0.727 | 0.357 | 0.4846456 | 14_Contaminant |
| PLCL1 | 1.12607211 | 0.545 | 0.192 | 0.50216741 | 14_Contaminant |
| SPTLC2 | 1.057591653 | 0.955 | 0.613 | 0.59872627 | 14_Contaminant |
| CCDC40 | 1.869384458 | 0.318 | 0.08 | 0.60594634 | 14_Contaminant |
| NRROS | 1.951794468 | 0.455 | 0.158 | 0.61318277 | 14_Contaminant |
| METTL22 | 1.991228213 | 0.364 | 0.106 | 0.6905645 | 14_Contaminant |
| CD86 | 1.149582908 | 0.909 | 0.623 | 0.6966238 | 14_Contaminant |
| BIRC6 | 1.132424469 | 0.864 | 0.473 | 0.69811165 | 14_Contaminant |
| MAP3K8 | 1.084347367 | 1 | 0.746 | 0.7890315 | 14_Contaminant |
| GALNT10 | 1.450992659 | 0.591 | 0.239 | 0.86512004 | 14_Contaminant |
| SH3RF3 | 1.010068123 | 0.773 | 0.391 | 0.89052571 | 14_Contaminant |
| TLE4 | 1.520856418 | 0.5 | 0.181 | 0.9050229 | 14_Contaminant |
| DLEU1 | 1.010263323 | 0.909 | 0.594 | 0.94650635 | 14_Contaminant |
| EIF3B | 1.650645099 | 0.455 | 0.159 | 0.95751207 | 14_Contaminant |
| AKT3 | 1.169753876 | 0.636 | 0.267 | 0.97470591 | 14_Contaminant |
| FAM118A | 2.160226085 | 0.409 | 0.138 | 0.97810174 | 14_Contaminant |
| FAM20A | 1.74548944 | 0.591 | 0.235 | 1 | 14_Contaminant |
| BTF3 | -3.717989221 | 0.091 | 0.539 | 1 | 14_Contaminant |
| STK32B | 1.905705147 | 0.273 | 0.063 | 1 | 14_Contaminant |
| RGS16 | 1.259152596 | 0.545 | 0.206 | 1 | 14_Contaminant |
| USP4 | 1.24193819 | 0.773 | 0.418 | 1 | 14_Contaminant |
| LYN | 1.123380648 | 0.909 | 0.688 | 1 | 14_Contaminant |
| VASH1 | 1.104550886 | 0.818 | 0.475 | 1 | 14_Contaminant |
| TOR1B | 2.255963112 | 0.318 | 0.087 | 1 | 14_Contaminant |
| RALGAPA2 | 1.504885018 | 0.591 | 0.259 | 1 | 14_Contaminant |
| ADAM17 | 1.327199471 | 0.818 | 0.526 | 1 | 14_Contaminant |
| MANBA | 1.232697884 | 0.909 | 0.681 | 1 | 14_Contaminant |
| NAA10 | 1.843705454 | 0.455 | 0.172 | 1 | 14_Contaminant |

| | | | | | |
|---|---|---|---|---|---|
| ARHGAP6 | 1.389504685 | 0.727 | 0.378 | 1 | 14_Contaminant |
| SUPT5H | 1.514539413 | 0.545 | 0.23 | 1 | 14_Contaminant |
| RPL41 | -2.90205505 | 0.273 | 0.675 | 1 | 14_Contaminant |
| GRASP | 1.049728208 | 0.955 | 0.592 | 1 | 14_Contaminant |
| DTNB | 1.27720822 | 0.591 | 0.248 | 1 | 14_Contaminant |
| SH2B3 | 1.147337798 | 0.818 | 0.488 | 1 | 14_Contaminant |
| TLN2 | 1.299819592 | 0.545 | 0.214 | 1 | 14_Contaminant |
| BCAS3 | 1.082118912 | 0.682 | 0.319 | 1 | 14_Contaminant |
| HLCS | 1.976438915 | 0.318 | 0.087 | 1 | 14_Contaminant |
| FANCL | 2.120865231 | 0.273 | 0.068 | 1 | 14_Contaminant |
| RPS4X | -2.445631276 | 0.455 | 0.71 | 1 | 14_Contaminant |
| XPC | 1.698406519 | 0.409 | 0.14 | 1 | 14_Contaminant |
| RPS6 | -1.991748713 | 0.545 | 0.752 | 1 | 14_Contaminant |
| CDC42SE1 | 1.634343126 | 0.636 | 0.347 | 1 | 14_Contaminant |
| VPS37A | 1.912033798 | 0.409 | 0.146 | 1 | 14_Contaminant |
| ANK2 | 1.661119837 | 0.318 | 0.089 | 1 | 14_Contaminant |
| NELFCD | 2.091169626 | 0.273 | 0.072 | 1 | 14_Contaminant |
| CENPU | 1.655337494 | 0.409 | 0.143 | 1 | 14_Contaminant |
| CYTH3 | 1.537644881 | 0.545 | 0.25 | 1 | 14_Contaminant |
| GSTM2 | 1.722099468 | 0.5 | 0.214 | 1 | 14_Contaminant |
| ADGRE5 | 1.562062214 | 0.591 | 0.303 | 1 | 14_Contaminant |
| RPL21 | -2.609385705 | 0.318 | 0.634 | 1 | 14_Contaminant |
| RPL10A | -2.496763677 | 0.182 | 0.58 | 1 | 14_Contaminant |
| EEF1B2 | -3.241111428 | 0.136 | 0.527 | 1 | 14_Contaminant |
| ATP2B4 | 1.799389994 | 0.455 | 0.176 | 1 | 14_Contaminant |
| STX4 | 1.631586887 | 0.636 | 0.331 | 1 | 14_Contaminant |
| ARMH1 | 1.780370512 | 0.318 | 0.095 | 1 | 14_Contaminant |
| RPL29 | -1.806037002 | 0.455 | 0.745 | 1 | 14_Contaminant |
| AKAP17A | 1.754420675 | 0.409 | 0.148 | 1 | 14_Contaminant |
| AMPD2 | 1.654579562 | 0.5 | 0.222 | 1 | 14_Contaminant |
| HAVCR2 | 1.097328412 | 0.818 | 0.543 | 1 | 14_Contaminant |
| NPL | 1.17299541 | 0.773 | 0.419 | 1 | 14_Contaminant |
| SH3TC1 | 1.11174472 | 0.818 | 0.522 | 1 | 14_Contaminant |
| CHKB | 1.761131212 | 0.455 | 0.188 | 1 | 14_Contaminant |
| DNM2 | 1.251112135 | 0.773 | 0.497 | 1 | 14_Contaminant |
| SLC31A2 | 1.014095423 | 0.773 | 0.467 | 1 | 14_Contaminant |
| NACA | -1.810332153 | 0.364 | 0.687 | 1 | 14_Contaminant |
| RPL4 | -2.458986446 | 0.227 | 0.581 | 1 | 14_Contaminant |
| RMDN1 | 2.263766531 | 0.318 | 0.103 | 1 | 14_Contaminant |
| TUBB | -11.67412724 | 0 | 0.383 | 1 | 14_Contaminant |
| BMPR1A | 1.274987154 | 0.318 | 0.097 | 1 | 14_Contaminant |
| RCC2 | 1.522161 | 0.591 | 0.312 | 1 | 14_Contaminant |
| BCO2 | 1.387681882 | 0.364 | 0.121 | 1 | 14_Contaminant |
| RPL8 | -2.037021977 | 0.5 | 0.716 | 1 | 14_Contaminant |
| USP9Y | 1.20962796 | 0.455 | 0.176 | 1 | 14_Contaminant |
| PLEKHG2 | 1.124497826 | 0.682 | 0.364 | 1 | 14_Contaminant |

| | | | | | |
|---|---|---|---|---|---|
| RPL5 | -2.148542748 | 0.364 | 0.655 | 1 | 14_Contaminant |
| SCAMP4 | 2.038685334 | 0.364 | 0.134 | 1 | 14_Contaminant |
| EMP3 | -12.10095659 | 0 | 0.376 | 1 | 14_Contaminant |
| CD46 | 1.133523989 | 0.545 | 0.245 | 1 | 14_Contaminant |
| SAFB2 | 1.419687021 | 0.5 | 0.227 | 1 | 14_Contaminant |
| SNX8 | 1.098270754 | 0.636 | 0.312 | 1 | 14_Contaminant |
| RPS7 | -1.854396203 | 0.455 | 0.716 | 1 | 14_Contaminant |
| FUT8 | 1.476115233 | 0.318 | 0.101 | 1 | 14_Contaminant |
| PIK3R1 | 1.263861672 | 0.773 | 0.573 | 1 | 14_Contaminant |
| RPSA | -2.149792349 | 0.182 | 0.578 | 1 | 14_Contaminant |
| PPIL4 | 1.593745748 | 0.5 | 0.236 | 1 | 14_Contaminant |
| STAT2 | 1.363292542 | 0.409 | 0.16 | 1 | 14_Contaminant |
| PDPN | 1.469594849 | 0.5 | 0.231 | 1 | 14_Contaminant |
| MLXIPL | 1.663943209 | 0.409 | 0.164 | 1 | 14_Contaminant |
| POM121 | 1.366734228 | 0.455 | 0.192 | 1 | 14_Contaminant |
| FAM20C | 1.306075153 | 0.5 | 0.223 | 1 | 14_Contaminant |
| CACNA1A | 1.35005142 | 0.364 | 0.125 | 1 | 14_Contaminant |
| SLC4A7 | 1.082737552 | 0.727 | 0.462 | 1 | 14_Contaminant |
| RALGAPA1 | 1.00915249 | 0.636 | 0.338 | 1 | 14_Contaminant |
| DCAF6 | 1.06781047 | 0.591 | 0.304 | 1 | 14_Contaminant |
| SIK2 | 1.1468047 | 0.636 | 0.362 | 1 | 14_Contaminant |
| GOLPH3 | 1.216936402 | 0.5 | 0.234 | 1 | 14_Contaminant |
| CALR | -2.918707734 | 0.136 | 0.49 | 1 | 14_Contaminant |
| MS4A4E | 1.486681925 | 0.5 | 0.235 | 1 | 14_Contaminant |
| INPP4A | 1.549509537 | 0.591 | 0.305 | 1 | 14_Contaminant |
| AL139807.1 | 1.343092038 | 0.364 | 0.13 | 1 | 14_Contaminant |
| RPL35A | -1.865356112 | 0.364 | 0.659 | 1 | 14_Contaminant |
| TCF7L2 | 1.257878274 | 0.455 | 0.195 | 1 | 14_Contaminant |
| SPRY2 | 1.448126617 | 0.364 | 0.133 | 1 | 14_Contaminant |
| RPL15 | -1.741332101 | 0.5 | 0.722 | 1 | 14_Contaminant |
| PLEKHM2 | 1.063452016 | 0.591 | 0.303 | 1 | 14_Contaminant |
| RPL14 | -1.580261625 | 0.5 | 0.734 | 1 | 14_Contaminant |
| TACC1 | 1.078703346 | 0.773 | 0.47 | 1 | 14_Contaminant |
| THRAP3 | 1.120464263 | 0.591 | 0.325 | 1 | 14_Contaminant |
| TRPC4AP | 1.370798848 | 0.545 | 0.283 | 1 | 14_Contaminant |
| AC087286.2 | 1.291210531 | 0.455 | 0.198 | 1 | 14_Contaminant |
| RPL3 | -1.564923181 | 0.5 | 0.736 | 1 | 14_Contaminant |
| BTBD1 | 1.489512998 | 0.5 | 0.244 | 1 | 14_Contaminant |
| AP2A2 | 1.146504439 | 0.636 | 0.371 | 1 | 14_Contaminant |
| EEF1D | -2.017046867 | 0.273 | 0.589 | 1 | 14_Contaminant |
| AP1B1 | 1.057266482 | 0.773 | 0.528 | 1 | 14_Contaminant |
| SECISBP2 | 1.355182211 | 0.5 | 0.238 | 1 | 14_Contaminant |
| SH3BGRL3 | -1.865092634 | 0.318 | 0.652 | 1 | 14_Contaminant |
| CD44 | -3.220323572 | 0.091 | 0.455 | 1 | 14_Contaminant |
| TBC1D8 | 1.023891557 | 0.636 | 0.352 | 1 | 14_Contaminant |
| S100A10 | -3.847969798 | 0.045 | 0.398 | 1 | 14_Contaminant |

| | | | | | |
|---|---|---|---|---|---|
| TTTY14 | 1.994304934 | 0.409 | 0.166 | 1 | 14_Contaminant |
| FCHSD2 | 1.193767788 | 0.818 | 0.539 | 1 | 14_Contaminant |
| CLEC2D | 1.41078175 | 0.318 | 0.109 | 1 | 14_Contaminant |
| NFATC1 | 1.182575086 | 0.545 | 0.278 | 1 | 14_Contaminant |
| SRRM1 | 1.146371087 | 0.682 | 0.425 | 1 | 14_Contaminant |
| IFI44 | 1.094825184 | 0.455 | 0.194 | 1 | 14_Contaminant |
| CCDC57 | 1.51699659 | 0.318 | 0.114 | 1 | 14_Contaminant |
| GADD45G | 1.324410913 | 0.364 | 0.141 | 1 | 14_Contaminant |
| SIGLEC8 | 1.442650581 | 0.409 | 0.18 | 1 | 14_Contaminant |
| PARVB | 1.098452232 | 0.636 | 0.427 | 1 | 14_Contaminant |
| KLHL5 | 1.226581713 | 0.636 | 0.374 | 1 | 14_Contaminant |
| SH3BP2 | 1.393249475 | 0.455 | 0.214 | 1 | 14_Contaminant |
| S100A4 | -4.190230335 | 0.045 | 0.378 | 1 | 14_Contaminant |
| CTDSP1 | 1.244085887 | 0.591 | 0.336 | 1 | 14_Contaminant |
| HOOK3 | 1.189073597 | 0.636 | 0.394 | 1 | 14_Contaminant |
| STRADA | 1.47326754 | 0.318 | 0.115 | 1 | 14_Contaminant |
| TIMP2 | 1.036229211 | 0.682 | 0.473 | 1 | 14_Contaminant |
| CPEB3 | 1.012572214 | 0.591 | 0.297 | 1 | 14_Contaminant |
| TRPM2 | 1.296431176 | 0.591 | 0.34 | 1 | 14_Contaminant |
| S100A6 | -4.914747932 | 0.045 | 0.365 | 1 | 14_Contaminant |
| FAM110B | 1.270819454 | 0.545 | 0.278 | 1 | 14_Contaminant |
| BNIP3L | -3.918931719 | 0.045 | 0.373 | 1 | 14_Contaminant |
| GPNMB | -12.01181831 | 0 | 0.314 | 1 | 14_Contaminant |
| EPC1 | 1.154587094 | 0.636 | 0.428 | 1 | 14_Contaminant |
| P2RY8 | 1.948153262 | 0.409 | 0.181 | 1 | 14_Contaminant |
| RPS21 | -1.712971444 | 0.455 | 0.657 | 1 | 14_Contaminant |
| NDUFS5 | -3.588789942 | 0.045 | 0.37 | 1 | 14_Contaminant |
| RPL23 | -2.418757153 | 0.136 | 0.466 | 1 | 14_Contaminant |
| MTRNR2L12 | -15.88302348 | 0 | 0.308 | 1 | 14_Contaminant |
| PABPN1 | 1.464589911 | 0.545 | 0.344 | 1 | 14_Contaminant |
| HGSNAT | 1.236136179 | 0.591 | 0.304 | 1 | 14_Contaminant |
| BICD1 | 1.680765512 | 0.364 | 0.155 | 1 | 14_Contaminant |
| SYNJ1 | 1.345185392 | 0.409 | 0.178 | 1 | 14_Contaminant |
| ARID4B | 1.075840791 | 0.682 | 0.439 | 1 | 14_Contaminant |
| MARK3 | 1.039649498 | 0.591 | 0.317 | 1 | 14_Contaminant |
| VPS26A | 1.425904155 | 0.5 | 0.263 | 1 | 14_Contaminant |
| UBE2F | 1.335778285 | 0.545 | 0.326 | 1 | 14_Contaminant |
| RPL31 | -1.916315494 | 0.273 | 0.556 | 1 | 14_Contaminant |
| FAM102B | 1.318740145 | 0.591 | 0.355 | 1 | 14_Contaminant |
| AC012150.1 | 1.115538313 | 0.364 | 0.146 | 1 | 14_Contaminant |
| RPL36 | -1.350148262 | 0.5 | 0.712 | 1 | 14_Contaminant |
| TAL1 | 1.160553813 | 0.409 | 0.171 | 1 | 14_Contaminant |
| ABI3 | 1.461794048 | 0.5 | 0.27 | 1 | 14_Contaminant |
| PCNX4 | 1.078445088 | 0.591 | 0.324 | 1 | 14_Contaminant |
| ZNF410 | 1.540021213 | 0.364 | 0.157 | 1 | 14_Contaminant |
| RYR1 | 1.000877241 | 0.455 | 0.213 | 1 | 14_Contaminant |

| | | | | | |
|---|---|---|---|---|---|
| DDIT3 | 1.167238157 | 0.455 | 0.225 | 1 | 14_Contaminant |
| UBA52 | -1.405278679 | 0.455 | 0.664 | 1 | 14_Contaminant |
| ARHGEF40 | 1.144751707 | 0.455 | 0.225 | 1 | 14_Contaminant |
| NFX1 | 1.095722759 | 0.455 | 0.219 | 1 | 14_Contaminant |
| TENT4B | 1.327175329 | 0.409 | 0.192 | 1 | 14_Contaminant |
| SLC25A5 | -2.880237705 | 0.091 | 0.386 | 1 | 14_Contaminant |
| RPL22 | -1.634438418 | 0.409 | 0.621 | 1 | 14_Contaminant |
| PLCB2 | 1.396636215 | 0.455 | 0.239 | 1 | 14_Contaminant |
| TIMP1 | -4.623271151 | 0.045 | 0.333 | 1 | 14_Contaminant |
| MAGT1 | 1.056332581 | 0.591 | 0.356 | 1 | 14_Contaminant |
| ZXDC | 1.324673656 | 0.409 | 0.191 | 1 | 14_Contaminant |
| TRIM33 | 1.082408066 | 0.409 | 0.191 | 1 | 14_Contaminant |
| USP22 | 1.00969476 | 0.5 | 0.259 | 1 | 14_Contaminant |
| SOD2 | -1.906853505 | 0.409 | 0.675 | 1 | 14_Contaminant |
| CIB1 | -3.446072422 | 0.045 | 0.331 | 1 | 14_Contaminant |
| LAMTOR4 | -2.659330549 | 0.091 | 0.382 | 1 | 14_Contaminant |
| NUP98 | 1.119290061 | 0.636 | 0.405 | 1 | 14_Contaminant |
| ZFAS1 | -1.367261511 | 0.182 | 0.518 | 1 | 14_Contaminant |
| AP2S1 | -3.50270981 | 0.045 | 0.326 | 1 | 14_Contaminant |
| COMMD6 | -2.766988706 | 0.045 | 0.339 | 1 | 14_Contaminant |
| VPS36 | 1.019383083 | 0.409 | 0.191 | 1 | 14_Contaminant |
| SBNO2 | 1.159284772 | 0.364 | 0.161 | 1 | 14_Contaminant |
| SERP1 | -1.498238863 | 0.227 | 0.542 | 1 | 14_Contaminant |
| PCMTD1 | 1.158294839 | 0.455 | 0.236 | 1 | 14_Contaminant |
| OST4 | -1.99256219 | 0.091 | 0.403 | 1 | 14_Contaminant |
| MYL12A | -2.062511669 | 0.136 | 0.438 | 1 | 14_Contaminant |
| ANXA2 | -2.586418646 | 0.091 | 0.383 | 1 | 14_Contaminant |
| TOMM7 | -1.670830039 | 0.091 | 0.404 | 1 | 14_Contaminant |
| FCGR2B | -11.11156983 | 0 | 0.259 | 1 | 14_Contaminant |
| SIPA1L2 | 1.235859617 | 0.5 | 0.297 | 1 | 14_Contaminant |
| NSA2 | -10.57413928 | 0 | 0.257 | 1 | 14_Contaminant |
| CDV3 | 1.168550813 | 0.591 | 0.345 | 1 | 14_Contaminant |
| STK4 | 1.111547298 | 0.727 | 0.526 | 1 | 14_Contaminant |
| IFITM10 | 1.42117071 | 0.455 | 0.249 | 1 | 14_Contaminant |
| ATP5PO | -2.846487767 | 0.045 | 0.327 | 1 | 14_Contaminant |
| AC012368.1 | 1.068239034 | 0.455 | 0.231 | 1 | 14_Contaminant |
| LYZ | -4.236264169 | 0.045 | 0.315 | 1 | 14_Contaminant |
| EIF3F | -2.622967696 | 0.045 | 0.328 | 1 | 14_Contaminant |
| ZDHHC14 | 1.034438934 | 0.455 | 0.238 | 1 | 14_Contaminant |
| LTC4S | 1.190506149 | 0.591 | 0.379 | 1 | 14_Contaminant |
| CXorf21 | 1.01065445 | 0.364 | 0.162 | 1 | 14_Contaminant |
| ARPC1B | -1.37507316 | 0.227 | 0.545 | 1 | 14_Contaminant |
| FXYD5 | -1.840568153 | 0.273 | 0.518 | 1 | 14_Contaminant |
| HNRNPL | 1.033810795 | 0.545 | 0.328 | 1 | 14_Contaminant |
| PPP4R3B | 1.170916059 | 0.409 | 0.208 | 1 | 14_Contaminant |
| MSRA | 1.151608038 | 0.455 | 0.243 | 1 | 14_Contaminant |

| | | | | | |
|---|---|---|---|---|---|
| ELOB | -2.222425124 | 0.091 | 0.373 | 1 | 14_Contaminant |
| PEA15 | -10.55352811 | 0 | 0.243 | 1 | 14_Contaminant |
| RAB5C | -2.287004121 | 0.091 | 0.37 | 1 | 14_Contaminant |
| DUSP2 | -2.882216479 | 0.045 | 0.314 | 1 | 14_Contaminant |
| HCST | -1.674486606 | 0.136 | 0.432 | 1 | 14_Contaminant |
| CRTAP | -10.49932708 | 0 | 0.24 | 1 | 14_Contaminant |
| RPS5 | -1.55212599 | 0.318 | 0.567 | 1 | 14_Contaminant |
| CLIC1 | -1.499833051 | 0.318 | 0.567 | 1 | 14_Contaminant |
