## Supplementary Table 9 for "MAIT cells have a negative impact on glioblastoma"

| Supplementary Table 9: Seurat based clustering of the DC subset of the myeloid compartment in the scRNA-seq analysis | | | | | |
|---|---|---|---|---|---|
| **Genes** | **avg_log2FC** | **pct.1** | **pct.2** | **p_val_adj** | **Cluster** |
| CCL22 | 4.00301177 | 0.784 | 0.081 | 3.23E-166 | 01_mDC1 |
| C15orf48 | 3.250695794 | 0.931 | 0.354 | 4.59E-134 | 01_mDC1 |
| CST7 | 2.594259224 | 0.934 | 0.331 | 5.30E-134 | 01_mDC1 |
| CCR7 | 2.787022446 | 0.789 | 0.14 | 1.93E-131 | 01_mDC1 |
| G0S2 | 3.295281427 | 0.863 | 0.25 | 7.06E-125 | 01_mDC1 |
| EMP3 | 2.063889166 | 0.975 | 0.585 | 2.80E-121 | 01_mDC1 |
| BCL3 | 2.444787524 | 0.893 | 0.329 | 2.18E-114 | 01_mDC1 |
| TIMP1 | 2.593561581 | 0.952 | 0.582 | 2.58E-111 | 01_mDC1 |
| TRAF1 | 2.541235917 | 0.769 | 0.176 | 6.06E-107 | 01_mDC1 |
| IL1R2 | 2.424445811 | 0.891 | 0.329 | 8.76E-107 | 01_mDC1 |
| CD44 | 2.10348052 | 0.977 | 0.59 | 1.14E-104 | 01_mDC1 |
| TNFAIP8 | 2.657796669 | 0.817 | 0.296 | 1.44E-100 | 01_mDC1 |
| THBS1 | 2.69123264 | 0.764 | 0.223 | 2.17E-95 | 01_mDC1 |
| ADAM19 | 2.866742208 | 0.68 | 0.15 | 1.30E-93 | 01_mDC1 |
| KYNU | 2.163095716 | 0.934 | 0.508 | 2.87E-93 | 01_mDC1 |
| TNFRSF4 | 3.876432797 | 0.508 | 0.054 | 1.13E-92 | 01_mDC1 |
| MT2A | 2.351489163 | 0.947 | 0.562 | 6.51E-92 | 01_mDC1 |
| EHD1 | 3.754523626 | 0.51 | 0.056 | 7.51E-91 | 01_mDC1 |
| ZNF331 | -3.48356306 | 0.317 | 0.817 | 8.82E-91 | 01_mDC1 |
| SERPINB1 | 1.829728965 | 0.954 | 0.628 | 4.09E-90 | 01_mDC1 |
| IL1R1 | 2.499748089 | 0.68 | 0.159 | 1.21E-87 | 01_mDC1 |
| CD1C | 1.897588408 | 0.754 | 0.191 | 2.01E-87 | 01_mDC1 |
| ARF6 | 1.75423943 | 0.909 | 0.488 | 1.88E-84 | 01_mDC1 |
| DUSP5 | 2.123978613 | 0.815 | 0.304 | 9.48E-83 | 01_mDC1 |
| NINJ1 | 1.851269446 | 0.893 | 0.435 | 1.20E-82 | 01_mDC1 |
| IL7R | 2.683326279 | 0.591 | 0.112 | 8.62E-82 | 01_mDC1 |
| CDK2AP1 | 1.839660492 | 0.868 | 0.423 | 1.02E-75 | 01_mDC1 |
| NFKB1 | 1.112635246 | 0.99 | 0.772 | 2.49E-74 | 01_mDC1 |
| MRC1 | 1.747099064 | 0.657 | 0.151 | 5.54E-73 | 01_mDC1 |
| ITGA5 | 2.112158085 | 0.728 | 0.27 | 1.98E-69 | 01_mDC1 |
| CRIP1 | 1.995569668 | 0.794 | 0.312 | 2.98E-69 | 01_mDC1 |
| BIRC3 | 1.186015823 | 0.756 | 0.255 | 1.93E-67 | 01_mDC1 |
| NR4A2 | -2.08313537 | 0.558 | 0.866 | 2.46E-67 | 01_mDC1 |
| FCER1A | 1.3792124 | 0.863 | 0.334 | 4.32E-67 | 01_mDC1 |
| BCL2A1 | 1.884299829 | 0.909 | 0.57 | 1.00E-66 | 01_mDC1 |
| RHOF | 2.639849348 | 0.604 | 0.174 | 1.71E-66 | 01_mDC1 |
| JARID2 | 1.429010075 | 0.959 | 0.688 | 4.29E-65 | 01_mDC1 |
| RALA | 1.649064447 | 0.873 | 0.483 | 6.15E-65 | 01_mDC1 |
| PIM3 | 1.795511134 | 0.782 | 0.343 | 7.52E-65 | 01_mDC1 |
| C3 | -4.17030105 | 0.162 | 0.642 | 3.81E-64 | 01_mDC1 |
| SEMA6B | 2.862586611 | 0.5 | 0.104 | 6.59E-64 | 01_mDC1 |
| TNFRSF18 | 4.447832316 | 0.322 | 0.021 | 7.29E-63 | 01_mDC1 |
| CFLAR | 1.247306714 | 0.962 | 0.673 | 7.98E-63 | 01_mDC1 |
| IFITM2 | 1.284533817 | 0.959 | 0.669 | 2.14E-62 | 01_mDC1 |

| | | | | | |
|---|---|---|---|---|---|
| ZFP36L2 | -2.2205595 | 0.487 | 0.838 | 2.65E-61 | 01_mDC1 |
| AKAP12 | 4.507209971 | 0.332 | 0.027 | 3.42E-61 | 01_mDC1 |
| MCOLN2 | 1.779448893 | 0.708 | 0.245 | 1.11E-60 | 01_mDC1 |
| S100A10 | 1.195050013 | 0.957 | 0.669 | 1.13E-59 | 01_mDC1 |
| SNHG15 | 1.740268458 | 0.787 | 0.364 | 1.32E-59 | 01_mDC1 |
| CMTM6 | 1.210057094 | 0.944 | 0.723 | 1.79E-59 | 01_mDC1 |
| MAP4K4 | 1.942324139 | 0.744 | 0.319 | 1.94E-59 | 01_mDC1 |
| B4GALT5 | 1.996181677 | 0.558 | 0.143 | 1.63E-58 | 01_mDC1 |
| RAP1B | 1.255639131 | 0.916 | 0.629 | 3.13E-57 | 01_mDC1 |
| SLC1A3 | -3.64575158 | 0.178 | 0.648 | 9.59E-57 | 01_mDC1 |
| BID | 1.49495275 | 0.815 | 0.441 | 1.61E-55 | 01_mDC1 |
| GPR137B | 1.556875641 | 0.784 | 0.366 | 2.01E-55 | 01_mDC1 |
| CD80 | 2.677344934 | 0.487 | 0.116 | 3.66E-55 | 01_mDC1 |
| LILRB2 | 2.535594872 | 0.541 | 0.162 | 4.80E-54 | 01_mDC1 |
| CRLF2 | 3.816505448 | 0.305 | 0.025 | 7.41E-54 | 01_mDC1 |
| CXCL8 | 1.966533393 | 0.749 | 0.337 | 1.64E-53 | 01_mDC1 |
| VSIR | -2.81188893 | 0.17 | 0.632 | 2.91E-53 | 01_mDC1 |
| SDC2 | 2.038798923 | 0.525 | 0.138 | 4.11E-53 | 01_mDC1 |
| VEGFA | 1.379142771 | 0.85 | 0.445 | 1.33E-52 | 01_mDC1 |
| DAPP1 | 2.000804229 | 0.581 | 0.188 | 1.42E-52 | 01_mDC1 |
| CD58 | 1.36229933 | 0.881 | 0.538 | 2.41E-52 | 01_mDC1 |
| IL1A | 3.122845601 | 0.447 | 0.096 | 4.69E-52 | 01_mDC1 |
| ETV3 | 1.387279506 | 0.838 | 0.45 | 1.89E-51 | 01_mDC1 |
| FLNA | 1.439966896 | 0.718 | 0.299 | 5.98E-51 | 01_mDC1 |
| FYB1 | -2.78451385 | 0.17 | 0.617 | 6.34E-51 | 01_mDC1 |
| LAMP3 | 1.833813146 | 0.444 | 0.091 | 2.91E-50 | 01_mDC1 |
| VMO1 | 1.82882088 | 0.716 | 0.325 | 3.21E-50 | 01_mDC1 |
| RELB | 1.592322067 | 0.711 | 0.324 | 5.16E-50 | 01_mDC1 |
| CYTOR | 1.823830689 | 0.66 | 0.26 | 6.90E-50 | 01_mDC1 |
| INSIG1 | 1.093401089 | 0.893 | 0.473 | 7.76E-50 | 01_mDC1 |
| METRNL | 1.203755762 | 0.898 | 0.538 | 1.41E-49 | 01_mDC1 |
| AL137857.1 | 1.933452102 | 0.546 | 0.154 | 1.64E-49 | 01_mDC1 |
| AL133415.1 | 1.896144326 | 0.668 | 0.278 | 1.88E-49 | 01_mDC1 |
| MAP3K14 | 2.25853169 | 0.48 | 0.121 | 3.57E-49 | 01_mDC1 |
| LMNA | 1.7941543 | 0.744 | 0.363 | 3.74E-49 | 01_mDC1 |
| APBB1IP | -2.61460789 | 0.175 | 0.614 | 4.85E-49 | 01_mDC1 |
| HOMER2 | 2.75150086 | 0.368 | 0.06 | 1.10E-48 | 01_mDC1 |
| FAM107B | 1.67102403 | 0.655 | 0.259 | 1.88E-48 | 01_mDC1 |
| MAMLD1 | 2.518008403 | 0.421 | 0.087 | 2.34E-48 | 01_mDC1 |
| USP12 | 1.79385963 | 0.609 | 0.212 | 3.92E-48 | 01_mDC1 |
| GPAT3 | 1.411930458 | 0.756 | 0.354 | 1.00E-47 | 01_mDC1 |
| IFNGR2 | 1.141099414 | 0.916 | 0.698 | 3.24E-47 | 01_mDC1 |
| NFKB2 | 1.71628033 | 0.635 | 0.25 | 4.97E-47 | 01_mDC1 |
| GNG4 | 3.597163719 | 0.249 | 0.015 | 6.82E-47 | 01_mDC1 |
| SLC11A1 | -3.44085769 | 0.147 | 0.562 | 1.76E-46 | 01_mDC1 |
| CRYBG1 | 1.023832194 | 0.873 | 0.481 | 7.62E-46 | 01_mDC1 |

| | | | | | |
|---|---|---|---|---|---|
| PDE4A | 1.241033436 | 0.802 | 0.426 | 8.08E-46 | 01_mDC1 |
| MEF2C | -2.08067832 | 0.373 | 0.753 | 1.11E-45 | 01_mDC1 |
| STK4 | 1.019252727 | 0.937 | 0.722 | 1.15E-45 | 01_mDC1 |
| TAGLN2 | 1.063658153 | 0.906 | 0.603 | 3.01E-45 | 01_mDC1 |
| MAP2K1 | 1.027905742 | 0.901 | 0.605 | 3.75E-45 | 01_mDC1 |
| PPIF | 1.58963316 | 0.723 | 0.336 | 4.87E-45 | 01_mDC1 |
| ICAM1 | 1.475170106 | 0.746 | 0.369 | 5.11E-45 | 01_mDC1 |
| LDHA | 1.072598288 | 0.904 | 0.66 | 1.16E-44 | 01_mDC1 |
| APOE | -4.2868754 | 0.348 | 0.674 | 2.54E-44 | 01_mDC1 |
| NECTIN2 | 1.575804504 | 0.713 | 0.356 | 1.57E-43 | 01_mDC1 |
| RHOB | -2.94892326 | 0.378 | 0.688 | 1.74E-43 | 01_mDC1 |
| LAMB3 | 3.942732637 | 0.251 | 0.021 | 1.85E-43 | 01_mDC1 |
| DUSP4 | 1.073560942 | 0.855 | 0.432 | 2.89E-43 | 01_mDC1 |
| MIR155HG | 1.657675113 | 0.546 | 0.174 | 2.90E-43 | 01_mDC1 |
| DPYD | -2.26531371 | 0.33 | 0.704 | 1.19E-42 | 01_mDC1 |
| CSGALNACT1 | -4.19823558 | 0.048 | 0.454 | 1.36E-42 | 01_mDC1 |
| C1QA | -3.25142545 | 0.223 | 0.605 | 1.86E-42 | 01_mDC1 |
| TREM2 | -4.24493508 | 0.094 | 0.487 | 2.22E-42 | 01_mDC1 |
| PTGIR | 4.422426877 | 0.221 | 0.012 | 2.50E-42 | 01_mDC1 |
| IRF4 | 1.288600597 | 0.594 | 0.203 | 3.35E-42 | 01_mDC1 |
| ALCAM | 1.575643427 | 0.868 | 0.582 | 4.28E-42 | 01_mDC1 |
| GPR157 | 1.457432445 | 0.543 | 0.164 | 6.05E-42 | 01_mDC1 |
| FOXN2 | 1.470526995 | 0.701 | 0.329 | 7.25E-42 | 01_mDC1 |
| GPR35 | 2.52246737 | 0.383 | 0.082 | 8.76E-42 | 01_mDC1 |
| HIVEP1 | 1.491466455 | 0.607 | 0.235 | 1.23E-41 | 01_mDC1 |
| PMAIP1 | 1.796956425 | 0.614 | 0.254 | 1.97E-41 | 01_mDC1 |
| RNF145 | 1.518061892 | 0.675 | 0.315 | 2.54E-41 | 01_mDC1 |
| CD163 | -2.45573994 | 0.188 | 0.603 | 3.21E-41 | 01_mDC1 |
| RFTN1 | 1.314011996 | 0.802 | 0.429 | 3.41E-41 | 01_mDC1 |
| C1QB | -3.01603214 | 0.274 | 0.639 | 5.37E-41 | 01_mDC1 |
| RUNX3 | 1.852829358 | 0.576 | 0.221 | 1.64E-40 | 01_mDC1 |
| SPP1 | -3.7134794 | 0.406 | 0.698 | 1.46E-39 | 01_mDC1 |
| CLEC10A | 1.181416115 | 0.673 | 0.26 | 3.35E-39 | 01_mDC1 |
| ANXA2 | 1.096613362 | 0.865 | 0.554 | 7.38E-39 | 01_mDC1 |
| MARCKSL1 | 1.418125217 | 0.487 | 0.15 | 7.60E-39 | 01_mDC1 |
| RNF149 | -1.42993993 | 0.642 | 0.846 | 1.17E-38 | 01_mDC1 |
| TRAF5 | 2.304904511 | 0.416 | 0.107 | 1.21E-38 | 01_mDC1 |
| C1QC | -3.29751429 | 0.297 | 0.634 | 1.42E-38 | 01_mDC1 |
| LPXN | 1.595144607 | 0.675 | 0.344 | 2.20E-38 | 01_mDC1 |
| MAP2K3 | 1.509003656 | 0.647 | 0.306 | 5.31E-38 | 01_mDC1 |
| CTSZ | 1.108160505 | 0.878 | 0.621 | 8.90E-38 | 01_mDC1 |
| TNIP2 | 2.059567108 | 0.459 | 0.145 | 1.12E-37 | 01_mDC1 |
| NDUFV2 | 1.2453566 | 0.789 | 0.496 | 1.28E-37 | 01_mDC1 |
| FURIN | 2.329647606 | 0.404 | 0.109 | 3.63E-37 | 01_mDC1 |
| HIP1 | 1.858655821 | 0.523 | 0.189 | 9.29E-37 | 01_mDC1 |
| IL10RA | 1.2845424 | 0.746 | 0.417 | 1.64E-36 | 01_mDC1 |

| | | | | | |
|---|---|---|---|---|---|
| ZC3H12A | 1.361251201 | 0.591 | 0.232 | 2.22E-36 | 01_mDC1 |
| ARHGAP31 | 1.496514328 | 0.678 | 0.339 | 4.45E-36 | 01_mDC1 |
| DENND5A | 1.515062828 | 0.665 | 0.329 | 6.04E-36 | 01_mDC1 |
| RASAL1 | 3.562250597 | 0.228 | 0.022 | 6.05E-36 | 01_mDC1 |
| TRABD2A | 2.552311208 | 0.299 | 0.054 | 8.27E-36 | 01_mDC1 |
| AC025580.2 | 2.60277327 | 0.365 | 0.089 | 2.34E-35 | 01_mDC1 |
| IER3 | 1.178531982 | 0.873 | 0.635 | 2.34E-35 | 01_mDC1 |
| PTP4A2 | 1.291221637 | 0.721 | 0.398 | 2.51E-35 | 01_mDC1 |
| FNBP1 | 1.100765992 | 0.812 | 0.522 | 3.51E-35 | 01_mDC1 |
| MTRNR2L12 | -6.46366161 | 0.013 | 0.356 | 5.51E-35 | 01_mDC1 |
| RNF19B | 1.799389085 | 0.475 | 0.158 | 7.58E-35 | 01_mDC1 |
| ARHGAP24 | -3.45487326 | 0.104 | 0.466 | 8.53E-35 | 01_mDC1 |
| EREG | 1.575983632 | 0.548 | 0.211 | 9.81E-35 | 01_mDC1 |
| GNA15 | 1.185763043 | 0.723 | 0.381 | 1.10E-34 | 01_mDC1 |
| NQO2 | 1.940402158 | 0.49 | 0.181 | 1.20E-34 | 01_mDC1 |
| ZEB2 | 1.078580885 | 0.886 | 0.661 | 1.41E-34 | 01_mDC1 |
| HDAC9 | -2.81624269 | 0.221 | 0.575 | 2.56E-34 | 01_mDC1 |
| CALCRL | 1.646828618 | 0.416 | 0.119 | 3.06E-34 | 01_mDC1 |
| ZBTB16 | -2.27152344 | 0.213 | 0.586 | 6.01E-34 | 01_mDC1 |
| LAPTM4A | 1.021983523 | 0.827 | 0.567 | 4.03E-33 | 01_mDC1 |
| IFITM1 | 1.084189382 | 0.685 | 0.308 | 5.91E-33 | 01_mDC1 |
| TNIP1 | 2.14710136 | 0.5 | 0.214 | 8.62E-33 | 01_mDC1 |
| OPN3 | 1.552870524 | 0.533 | 0.216 | 8.76E-33 | 01_mDC1 |
| RNF213-AS1 | 2.579390512 | 0.272 | 0.045 | 9.18E-33 | 01_mDC1 |
| STX11 | 1.003361067 | 0.825 | 0.536 | 1.65E-32 | 01_mDC1 |
| SATB1 | 1.400726361 | 0.629 | 0.3 | 1.69E-32 | 01_mDC1 |
| FCGR3A | -3.4465087 | 0.086 | 0.431 | 2.23E-32 | 01_mDC1 |
| BZW1 | 1.158694366 | 0.731 | 0.414 | 2.30E-32 | 01_mDC1 |
| LFNG | 2.189414912 | 0.335 | 0.079 | 3.05E-32 | 01_mDC1 |
| ANKRD44 | -1.92463422 | 0.294 | 0.644 | 3.50E-32 | 01_mDC1 |
| USP53 | -2.99397194 | 0.145 | 0.496 | 5.86E-32 | 01_mDC1 |
| SERPINB9P1 | 2.002673355 | 0.373 | 0.099 | 5.86E-32 | 01_mDC1 |
| ZMIZ1 | 1.19282237 | 0.761 | 0.471 | 7.58E-32 | 01_mDC1 |
| ENTPD1 | 1.240064966 | 0.759 | 0.488 | 8.16E-32 | 01_mDC1 |
| LILRA5 | 2.733767181 | 0.299 | 0.062 | 8.91E-32 | 01_mDC1 |
| SPHK1 | 2.35857506 | 0.386 | 0.117 | 9.89E-32 | 01_mDC1 |
| PDGFB | 1.288101721 | 0.635 | 0.288 | 1.00E-31 | 01_mDC1 |
| SYNJ2 | 2.773760685 | 0.294 | 0.06 | 1.00E-31 | 01_mDC1 |
| UPP1 | 1.25245853 | 0.668 | 0.352 | 1.82E-31 | 01_mDC1 |
| MYO1G | 1.740537817 | 0.523 | 0.221 | 2.16E-31 | 01_mDC1 |
| CFP | 1.759732501 | 0.409 | 0.119 | 2.24E-31 | 01_mDC1 |
| FRMD4A | -2.81621897 | 0.188 | 0.531 | 2.59E-31 | 01_mDC1 |
| SOD2 | 1.16082782 | 0.863 | 0.622 | 4.51E-31 | 01_mDC1 |
| TPRA1 | 2.288064547 | 0.401 | 0.13 | 5.08E-31 | 01_mDC1 |
| TXNRD1 | 1.365114722 | 0.635 | 0.323 | 5.41E-31 | 01_mDC1 |
| ZHX2 | 1.369531034 | 0.536 | 0.214 | 9.93E-31 | 01_mDC1 |

| | | | | | |
|---|---|---|---|---|---|
| H2AFZ | 1.17828984 | 0.817 | 0.568 | 1.03E-30 | 01_mDC1 |
| IL4I1 | 1.643137752 | 0.475 | 0.175 | 1.34E-30 | 01_mDC1 |
| INPP5D | -2.37770973 | 0.19 | 0.538 | 2.12E-30 | 01_mDC1 |
| IL1B | 1.389836199 | 0.904 | 0.64 | 2.22E-30 | 01_mDC1 |
| ALG2 | 1.83321236 | 0.449 | 0.161 | 2.27E-30 | 01_mDC1 |
| SLC41A2 | 1.258924087 | 0.51 | 0.199 | 2.79E-30 | 01_mDC1 |
| FGD4 | -1.79903757 | 0.495 | 0.765 | 2.95E-30 | 01_mDC1 |
| SMAD3 | 1.544830071 | 0.497 | 0.196 | 7.07E-30 | 01_mDC1 |
| LGALS3 | 1.052751102 | 0.703 | 0.385 | 7.91E-30 | 01_mDC1 |
| TNFAIP2 | 2.098315347 | 0.492 | 0.209 | 5.46E-29 | 01_mDC1 |
| SFMBT2 | -2.54343171 | 0.206 | 0.537 | 6.74E-29 | 01_mDC1 |
| APOC1 | -3.73022281 | 0.292 | 0.573 | 7.14E-29 | 01_mDC1 |
| PYCARD | -1.87011847 | 0.208 | 0.546 | 1.79E-28 | 01_mDC1 |
| PID1 | 2.205740922 | 0.411 | 0.14 | 1.99E-28 | 01_mDC1 |
| NCS1 | 2.751347761 | 0.256 | 0.047 | 2.44E-28 | 01_mDC1 |
| SRC | 2.148378642 | 0.365 | 0.109 | 2.66E-28 | 01_mDC1 |
| SSBP3 | 1.909817612 | 0.289 | 0.064 | 2.94E-28 | 01_mDC1 |
| LRRFIP2 | 1.53832399 | 0.528 | 0.236 | 3.10E-28 | 01_mDC1 |
| ST6GAL1 | -3.65680655 | 0.066 | 0.38 | 5.20E-28 | 01_mDC1 |
| MEF2A | -1.55833618 | 0.579 | 0.804 | 5.95E-28 | 01_mDC1 |
| TRIP10 | 2.062403426 | 0.358 | 0.107 | 9.06E-28 | 01_mDC1 |
| GPR34 | -4.26171279 | 0.036 | 0.338 | 1.16E-27 | 01_mDC1 |
| KCTD12 | -2.2521483 | 0.147 | 0.476 | 1.20E-27 | 01_mDC1 |
| KCNQ3 | -6.7676694 | 0.013 | 0.302 | 1.36E-27 | 01_mDC1 |
| UGCG | 1.416775049 | 0.584 | 0.286 | 1.53E-27 | 01_mDC1 |
| ZBTB7A | 1.505813492 | 0.477 | 0.193 | 2.55E-27 | 01_mDC1 |
| TBC1D8 | 1.128437179 | 0.711 | 0.423 | 9.96E-27 | 01_mDC1 |
| PNP | 1.696194985 | 0.47 | 0.192 | 1.15E-26 | 01_mDC1 |
| IL18 | -1.6650013 | 0.299 | 0.599 | 1.24E-26 | 01_mDC1 |
| CSF3R | -1.69975903 | 0.305 | 0.614 | 1.48E-26 | 01_mDC1 |
| CTSC | -2.12147838 | 0.198 | 0.51 | 1.80E-26 | 01_mDC1 |
| STAT4 | 1.460862159 | 0.556 | 0.256 | 2.93E-26 | 01_mDC1 |
| ARHGEF2 | 1.542217676 | 0.525 | 0.241 | 4.65E-26 | 01_mDC1 |
| CSF1R | -1.74850432 | 0.393 | 0.643 | 1.05E-25 | 01_mDC1 |
| HMOX1 | -1.61798448 | 0.185 | 0.518 | 1.19E-25 | 01_mDC1 |
| CYP2S1 | 1.257486991 | 0.513 | 0.233 | 1.28E-25 | 01_mDC1 |
| IL18R1 | 1.339191182 | 0.563 | 0.266 | 1.56E-25 | 01_mDC1 |
| SRGAP2 | -1.83660151 | 0.391 | 0.666 | 2.07E-25 | 01_mDC1 |
| HIPK2 | 1.142210962 | 0.65 | 0.355 | 2.43E-25 | 01_mDC1 |
| CTSD | -2.28747889 | 0.216 | 0.529 | 2.53E-25 | 01_mDC1 |
| ANXA11 | 1.140705885 | 0.695 | 0.437 | 2.65E-25 | 01_mDC1 |
| SPTLC2 | -2.16641253 | 0.287 | 0.585 | 2.74E-25 | 01_mDC1 |
| CXCL2 | 1.808418853 | 0.391 | 0.133 | 8.27E-25 | 01_mDC1 |
| FABP5 | 2.13764412 | 0.51 | 0.264 | 5.34E-24 | 01_mDC1 |
| LHFPL2 | -3.707253 | 0.053 | 0.334 | 7.11E-24 | 01_mDC1 |
| DLEU1 | -2.72297607 | 0.152 | 0.452 | 8.14E-24 | 01_mDC1 |

| | | | | | |
|---|---|---|---|---|---|
| DISC1 | -2.94695395 | 0.089 | 0.379 | 1.39E-23 | 01_mDC1 |
| SLCO2B1 | -4.32034723 | 0.033 | 0.301 | 1.75E-23 | 01_mDC1 |
| MFSD12 | 1.683638184 | 0.497 | 0.234 | 2.54E-23 | 01_mDC1 |
| SIGLEC10 | -2.07287792 | 0.15 | 0.459 | 4.30E-23 | 01_mDC1 |
| ALYREF | 1.51650755 | 0.437 | 0.175 | 5.33E-23 | 01_mDC1 |
| SLCO3A1 | 1.697797741 | 0.345 | 0.11 | 6.86E-23 | 01_mDC1 |
| LRMDA | -1.6277916 | 0.528 | 0.76 | 6.87E-23 | 01_mDC1 |
| EGR1 | -3.67850912 | 0.089 | 0.369 | 1.04E-22 | 01_mDC1 |
| HSPA1A | -3.26950906 | 0.398 | 0.629 | 1.07E-22 | 01_mDC1 |
| GALNT2 | -2.83055158 | 0.094 | 0.381 | 1.15E-22 | 01_mDC1 |
| FPR3 | 1.146918143 | 0.556 | 0.279 | 2.09E-22 | 01_mDC1 |
| TTYH2 | 1.527537735 | 0.431 | 0.175 | 2.26E-22 | 01_mDC1 |
| GRAMD2B | 1.854289221 | 0.363 | 0.126 | 2.39E-22 | 01_mDC1 |
| PSMB9 | -2.48157524 | 0.122 | 0.412 | 2.96E-22 | 01_mDC1 |
| TRAF3 | 1.358792253 | 0.396 | 0.146 | 8.62E-22 | 01_mDC1 |
| DAGLB | -2.11842183 | 0.168 | 0.467 | 9.98E-22 | 01_mDC1 |
| CORO1A | -1.40022025 | 0.388 | 0.629 | 1.45E-21 | 01_mDC1 |
| GNAQ | -1.32933426 | 0.464 | 0.726 | 2.25E-21 | 01_mDC1 |
| SP110 | -2.02506322 | 0.129 | 0.426 | 2.87E-21 | 01_mDC1 |
| RAB21 | 1.247866993 | 0.525 | 0.261 | 3.09E-21 | 01_mDC1 |
| DDAH2 | 1.07420893 | 0.65 | 0.387 | 4.75E-21 | 01_mDC1 |
| TBC1D5 | -1.83503611 | 0.193 | 0.495 | 5.19E-21 | 01_mDC1 |
| ZFY | 1.60873851 | 0.327 | 0.106 | 6.91E-21 | 01_mDC1 |
| CNPY3 | -1.45219529 | 0.297 | 0.581 | 1.22E-20 | 01_mDC1 |
| PTTG1 | 2.0916877 | 0.294 | 0.092 | 1.70E-20 | 01_mDC1 |
| HBEGF | -2.00595559 | 0.249 | 0.5 | 1.92E-20 | 01_mDC1 |
| MIR4435-2HG | 1.310562689 | 0.48 | 0.231 | 4.35E-20 | 01_mDC1 |
| IFNGR1 | -1.29504219 | 0.477 | 0.704 | 4.67E-20 | 01_mDC1 |
| MYCBP2 | -1.7142267 | 0.239 | 0.529 | 5.38E-20 | 01_mDC1 |
| SNN | 2.035352401 | 0.299 | 0.096 | 6.16E-20 | 01_mDC1 |
| FCGR1A | -2.69394324 | 0.119 | 0.379 | 1.19E-19 | 01_mDC1 |
| ATF3 | -1.67867969 | 0.416 | 0.64 | 1.26E-19 | 01_mDC1 |
| OLFML3 | -4.26326445 | 0.023 | 0.256 | 1.86E-19 | 01_mDC1 |
| PALD1 | -1.68889067 | 0.292 | 0.574 | 1.93E-19 | 01_mDC1 |
| TBXAS1 | -1.28653279 | 0.505 | 0.72 | 2.38E-19 | 01_mDC1 |
| SSH1 | 1.364049602 | 0.482 | 0.233 | 2.94E-19 | 01_mDC1 |
| KMO | 1.068383218 | 0.348 | 0.122 | 3.45E-19 | 01_mDC1 |
| ZBTB20 | -2.11796059 | 0.18 | 0.455 | 3.76E-19 | 01_mDC1 |
| RGS19 | -2.03215952 | 0.107 | 0.378 | 4.57E-19 | 01_mDC1 |
| ANKLE2 | 1.439646033 | 0.437 | 0.195 | 5.44E-19 | 01_mDC1 |
| FBRSL1 | 1.137304982 | 0.431 | 0.18 | 7.34E-19 | 01_mDC1 |
| MIR22HG | 1.519450581 | 0.411 | 0.18 | 8.30E-19 | 01_mDC1 |
| MAFB | -2.15141441 | 0.155 | 0.423 | 1.00E-18 | 01_mDC1 |
| N4BP1 | 1.365613177 | 0.416 | 0.176 | 1.31E-18 | 01_mDC1 |
| WTAP | 1.061677672 | 0.683 | 0.473 | 1.61E-18 | 01_mDC1 |
| MAN2A1 | -2.06747954 | 0.272 | 0.526 | 2.17E-18 | 01_mDC1 |

| | | | | | |
|---|---|---|---|---|---|
| DNAJB1 | -2.69483439 | 0.279 | 0.531 | 3.48E-18 | 01_mDC1 |
| GNG7 | -2.07639426 | 0.165 | 0.429 | 3.49E-18 | 01_mDC1 |
| NDUFA6 | 1.039857125 | 0.612 | 0.374 | 4.51E-18 | 01_mDC1 |
| MIR3945HG | 1.861090383 | 0.365 | 0.15 | 4.92E-18 | 01_mDC1 |
| ENTPD1-AS1 | 1.090980889 | 0.523 | 0.272 | 5.35E-18 | 01_mDC1 |
| SHTN1 | -2.03454329 | 0.147 | 0.421 | 5.59E-18 | 01_mDC1 |
| STK24 | 1.051202097 | 0.558 | 0.309 | 6.67E-18 | 01_mDC1 |
| HSPA1B | -3.12602833 | 0.272 | 0.5 | 7.82E-18 | 01_mDC1 |
| GLRX | -1.85080692 | 0.112 | 0.375 | 8.68E-18 | 01_mDC1 |
| TNFRSF1A | 1.099510014 | 0.584 | 0.369 | 1.30E-17 | 01_mDC1 |
| FAM53C | -2.34598488 | 0.109 | 0.362 | 1.32E-17 | 01_mDC1 |
| NAIP | -2.4504621 | 0.081 | 0.33 | 1.70E-17 | 01_mDC1 |
| 1-Mar | -1.94223553 | 0.15 | 0.417 | 1.71E-17 | 01_mDC1 |
| PRMT9 | -2.35257178 | 0.165 | 0.42 | 1.88E-17 | 01_mDC1 |
| CADM1 | -3.55666805 | 0.043 | 0.272 | 2.74E-17 | 01_mDC1 |
| NFKBIE | 1.297765456 | 0.424 | 0.193 | 2.75E-17 | 01_mDC1 |
| SLC25A37 | -1.72959101 | 0.175 | 0.442 | 3.29E-17 | 01_mDC1 |
| CD84 | -2.15449495 | 0.094 | 0.348 | 4.30E-17 | 01_mDC1 |
| ABCC4 | -2.90219183 | 0.071 | 0.308 | 4.67E-17 | 01_mDC1 |
| ZC3HAV1 | -2.10029673 | 0.274 | 0.517 | 4.68E-17 | 01_mDC1 |
| CD300A | -1.89136501 | 0.147 | 0.406 | 5.34E-17 | 01_mDC1 |
| IRS2 | -1.81455324 | 0.173 | 0.439 | 6.35E-17 | 01_mDC1 |
| SLC6A6 | 1.407924631 | 0.434 | 0.212 | 1.02E-16 | 01_mDC1 |
| DOK2 | 1.612542407 | 0.345 | 0.139 | 1.24E-16 | 01_mDC1 |
| CCSER2 | 1.332600804 | 0.429 | 0.2 | 1.35E-16 | 01_mDC1 |
| FGD2 | -2.70014773 | 0.074 | 0.312 | 1.44E-16 | 01_mDC1 |
| ATF5 | 1.265090565 | 0.464 | 0.241 | 1.48E-16 | 01_mDC1 |
| PXDC1 | 1.55716665 | 0.358 | 0.149 | 1.53E-16 | 01_mDC1 |
| SNX10 | -1.96554085 | 0.119 | 0.37 | 1.98E-16 | 01_mDC1 |
| ANTXR2 | 1.373923541 | 0.378 | 0.162 | 2.05E-16 | 01_mDC1 |
| LACTB | 1.121185462 | 0.492 | 0.261 | 3.11E-16 | 01_mDC1 |
| MGAT4A | -1.91231486 | 0.145 | 0.409 | 3.23E-16 | 01_mDC1 |
| FHIT | -2.32757966 | 0.145 | 0.389 | 5.26E-16 | 01_mDC1 |
| CHD9 | -1.91074241 | 0.168 | 0.419 | 5.86E-16 | 01_mDC1 |
| ADAM8 | 1.108020169 | 0.472 | 0.235 | 6.58E-16 | 01_mDC1 |
| FCHSD2 | -1.54199066 | 0.317 | 0.571 | 6.99E-16 | 01_mDC1 |
| RARA | 1.001241095 | 0.536 | 0.302 | 1.03E-15 | 01_mDC1 |
| CUL1 | 1.291592689 | 0.523 | 0.299 | 1.09E-15 | 01_mDC1 |
| ARHGAP15 | -1.13877974 | 0.487 | 0.704 | 1.23E-15 | 01_mDC1 |
| MSR1 | -1.72315288 | 0.386 | 0.588 | 1.29E-15 | 01_mDC1 |
| DNMBP | 1.046672408 | 0.439 | 0.208 | 1.31E-15 | 01_mDC1 |
| SBNO2 | 1.481572961 | 0.383 | 0.171 | 1.32E-15 | 01_mDC1 |
| IGSF6 | -1.61977146 | 0.221 | 0.471 | 2.08E-15 | 01_mDC1 |
| FNDC3A | -1.97315707 | 0.165 | 0.418 | 2.12E-15 | 01_mDC1 |
| BIRC2 | 1.065662571 | 0.5 | 0.266 | 2.88E-15 | 01_mDC1 |
| LINC02256 | -1.75975349 | 0.145 | 0.396 | 2.88E-15 | 01_mDC1 |

| | | | | | |
|---|---|---|---|---|---|
| TRPM2 | -2.68346378 | 0.051 | 0.268 | 4.46E-15 | 01_mDC1 |
| ANKRD28 | 1.08000222 | 0.401 | 0.177 | 4.66E-15 | 01_mDC1 |
| CAMK2D | -2.96678754 | 0.063 | 0.283 | 4.94E-15 | 01_mDC1 |
| SUMF1 | -2.16557691 | 0.119 | 0.359 | 5.12E-15 | 01_mDC1 |
| AC060765.2 | 1.078062655 | 0.348 | 0.137 | 7.28E-15 | 01_mDC1 |
| GPAT4 | 1.088565228 | 0.475 | 0.258 | 1.20E-14 | 01_mDC1 |
| SORL1 | -1.56474239 | 0.294 | 0.522 | 1.30E-14 | 01_mDC1 |
| PSMB8 | -1.73716842 | 0.132 | 0.375 | 1.34E-14 | 01_mDC1 |
| LUCAT1 | 1.188451832 | 0.355 | 0.149 | 1.37E-14 | 01_mDC1 |
| PRAM1 | -2.76790499 | 0.046 | 0.255 | 1.59E-14 | 01_mDC1 |
| ARHGAP22 | -2.8613091 | 0.061 | 0.275 | 1.71E-14 | 01_mDC1 |
| VASH1 | -1.42944957 | 0.388 | 0.606 | 2.73E-14 | 01_mDC1 |
| AL163541.1 | -3.72811867 | 0.048 | 0.25 | 2.82E-14 | 01_mDC1 |
| SMIM3 | 1.177095628 | 0.454 | 0.237 | 2.99E-14 | 01_mDC1 |
| TANC2 | -1.81242866 | 0.208 | 0.449 | 3.38E-14 | 01_mDC1 |
| DOCK10 | -1.44138355 | 0.307 | 0.552 | 3.71E-14 | 01_mDC1 |
| PLEKHA5 | 1.298159122 | 0.426 | 0.215 | 4.37E-14 | 01_mDC1 |
| SP100 | -1.17095616 | 0.348 | 0.588 | 4.59E-14 | 01_mDC1 |
| C9orf72 | -1.20509143 | 0.404 | 0.641 | 4.73E-14 | 01_mDC1 |
| SCPEP1 | -1.97945866 | 0.122 | 0.352 | 6.36E-14 | 01_mDC1 |
| CREB5 | 1.05421819 | 0.5 | 0.278 | 8.59E-14 | 01_mDC1 |
| SLC9A9 | -2.97480793 | 0.058 | 0.263 | 1.14E-13 | 01_mDC1 |
| HSPH1 | -2.34765427 | 0.249 | 0.476 | 1.27E-13 | 01_mDC1 |
| PARVG | -1.203409 | 0.317 | 0.554 | 1.30E-13 | 01_mDC1 |
| SAP30 | -1.94676473 | 0.201 | 0.421 | 1.62E-13 | 01_mDC1 |
| MNDA | -1.62117046 | 0.206 | 0.439 | 1.68E-13 | 01_mDC1 |
| SMNDC1 | 1.274871784 | 0.391 | 0.189 | 2.11E-13 | 01_mDC1 |
| RERE | -1.78055833 | 0.221 | 0.453 | 2.41E-13 | 01_mDC1 |
| PDCL3 | 1.213929944 | 0.447 | 0.24 | 2.54E-13 | 01_mDC1 |
| SKAP2 | -1.61032737 | 0.213 | 0.447 | 2.69E-13 | 01_mDC1 |
| PDK4 | -3.49179493 | 0.094 | 0.299 | 3.21E-13 | 01_mDC1 |
| LPCAT2 | -1.54122404 | 0.327 | 0.551 | 3.68E-13 | 01_mDC1 |
| SYK | -1.26249818 | 0.284 | 0.521 | 4.32E-13 | 01_mDC1 |
| ADAM28 | -1.16781666 | 0.396 | 0.626 | 5.09E-13 | 01_mDC1 |
| SMYD3 | -2.89745002 | 0.069 | 0.27 | 5.29E-13 | 01_mDC1 |
| RASA1 | -1.8903895 | 0.208 | 0.433 | 6.56E-13 | 01_mDC1 |
| ATM | -2.13134017 | 0.089 | 0.302 | 7.19E-13 | 01_mDC1 |
| PCNX2 | -2.33237048 | 0.091 | 0.302 | 8.62E-13 | 01_mDC1 |
| AOAH | -1.31143066 | 0.35 | 0.558 | 8.97E-13 | 01_mDC1 |
| CD69 | -1.55030982 | 0.274 | 0.495 | 1.06E-12 | 01_mDC1 |
| FLI1 | -2.20671931 | 0.091 | 0.304 | 1.36E-12 | 01_mDC1 |
| IFI44L | -2.43429127 | 0.069 | 0.271 | 1.38E-12 | 01_mDC1 |
| PCNX4 | -2.35600369 | 0.084 | 0.289 | 1.53E-12 | 01_mDC1 |
| IFRD1 | -1.54938949 | 0.332 | 0.537 | 2.50E-12 | 01_mDC1 |
| CALCOCO2 | -1.86729335 | 0.089 | 0.303 | 2.82E-12 | 01_mDC1 |
| CORO1B | -1.7015968 | 0.129 | 0.346 | 3.01E-12 | 01_mDC1 |

| | | | | | |
|---|---|---|---|---|---|
| SRGAP2B | -1.50565268 | 0.269 | 0.498 | 3.08E-12 | 01_mDC1 |
| EVL | -1.57234179 | 0.16 | 0.386 | 4.59E-12 | 01_mDC1 |
| ADAMTSL4-AS | -1.6908307 | 0.277 | 0.49 | 5.00E-12 | 01_mDC1 |
| EPB41L2 | -1.75928259 | 0.317 | 0.523 | 5.72E-12 | 01_mDC1 |
| CYBC1 | -2.01279697 | 0.071 | 0.273 | 6.10E-12 | 01_mDC1 |
| RABGAP1L | -1.96123941 | 0.206 | 0.422 | 6.54E-12 | 01_mDC1 |
| WWOX | -2.33793417 | 0.109 | 0.313 | 9.64E-12 | 01_mDC1 |
| NAF1 | -2.29589548 | 0.119 | 0.327 | 1.10E-11 | 01_mDC1 |
| EXOC4 | -1.70498631 | 0.188 | 0.407 | 1.16E-11 | 01_mDC1 |
| SLA | -1.48888146 | 0.254 | 0.473 | 1.26E-11 | 01_mDC1 |
| CLNS1A | -1.75536581 | 0.124 | 0.342 | 1.72E-11 | 01_mDC1 |
| DHRS7 | -1.45212205 | 0.261 | 0.47 | 1.73E-11 | 01_mDC1 |
| ARID5B | -1.3463363 | 0.299 | 0.524 | 1.97E-11 | 01_mDC1 |
| SLC4A7 | -2.0026034 | 0.175 | 0.388 | 2.11E-11 | 01_mDC1 |
| PLA2G4A | -1.78535165 | 0.18 | 0.394 | 2.17E-11 | 01_mDC1 |
| KLHL6 | -1.4412727 | 0.305 | 0.528 | 2.30E-11 | 01_mDC1 |
| BST2 | -1.24868076 | 0.345 | 0.545 | 2.48E-11 | 01_mDC1 |
| JAK2 | -1.80621957 | 0.16 | 0.371 | 4.92E-11 | 01_mDC1 |
| UBE2L6 | -1.82753371 | 0.122 | 0.327 | 4.95E-11 | 01_mDC1 |
| LAMTOR1 | -1.25811544 | 0.274 | 0.486 | 4.99E-11 | 01_mDC1 |
| AKR1B1 | -1.28589943 | 0.251 | 0.458 | 1.79E-10 | 01_mDC1 |
| WDFY4 | -2.29551913 | 0.157 | 0.364 | 1.90E-10 | 01_mDC1 |
| CLEC2B | -1.28861686 | 0.193 | 0.411 | 5.57E-10 | 01_mDC1 |
| PRKCH | -1.2091133 | 0.325 | 0.547 | 5.62E-10 | 01_mDC1 |
| CKS2 | -1.79371673 | 0.137 | 0.343 | 5.76E-10 | 01_mDC1 |
| CIAO2A | -1.3315286 | 0.173 | 0.386 | 6.56E-10 | 01_mDC1 |
| PAK1 | -1.08166835 | 0.32 | 0.537 | 1.08E-09 | 01_mDC1 |
| CHKA | -1.36719546 | 0.272 | 0.483 | 1.11E-09 | 01_mDC1 |
| ADAP2 | -1.37481472 | 0.203 | 0.406 | 2.81E-09 | 01_mDC1 |
| HEXA | -1.26080938 | 0.236 | 0.439 | 7.43E-09 | 01_mDC1 |
| SCAMP2 | -1.21741203 | 0.236 | 0.438 | 1.12E-08 | 01_mDC1 |
| TBC1D22A | -1.00664619 | 0.383 | 0.585 | 8.44E-08 | 01_mDC1 |
| SERPINA1 | 1.930865506 | 0.913 | 0.544 | 2.35E-66 | 02_HSP-DC |
| MS4A6A | 1.442730362 | 0.939 | 0.682 | 2.15E-57 | 02_HSP-DC |
| EEF1G | 1.02434477 | 0.994 | 0.77 | 8.72E-52 | 02_HSP-DC |
| VAMP8 | 1.253407167 | 0.907 | 0.523 | 3.32E-50 | 02_HSP-DC |
| HSPA8 | 1.005270959 | 0.952 | 0.672 | 2.71E-45 | 02_HSP-DC |
| GAPT | 2.104114999 | 0.502 | 0.121 | 1.41E-44 | 02_HSP-DC |
| UBE2L6 | 1.884717754 | 0.592 | 0.191 | 1.49E-42 | 02_HSP-DC |
| TKT | 1.409511902 | 0.749 | 0.328 | 1.38E-41 | 02_HSP-DC |
| BST2 | 1.236271015 | 0.839 | 0.403 | 2.07E-41 | 02_HSP-DC |
| PLAC8 | 2.147316887 | 0.527 | 0.152 | 3.34E-41 | 02_HSP-DC |
| LAMTOR1 | 1.282245303 | 0.797 | 0.336 | 3.66E-41 | 02_HSP-DC |
| PSMB9 | 1.469162557 | 0.675 | 0.249 | 1.30E-40 | 02_HSP-DC |
| CLNS1A | 1.470446954 | 0.617 | 0.2 | 4.89E-40 | 02_HSP-DC |
| SNX10 | 1.543155372 | 0.63 | 0.22 | 2.08E-39 | 02_HSP-DC |

| | | | | | |
|---|---|---|---|---|---|
| CALHM6 | 1.637011687 | 0.743 | 0.337 | 7.38E-37 | 02_HSP-DC |
| VSIR | 1.181777308 | 0.836 | 0.428 | 8.74E-35 | 02_HSP-DC |
| POLD4 | 1.095485408 | 0.804 | 0.388 | 1.17E-33 | 02_HSP-DC |
| VAMP5 | 1.981869913 | 0.514 | 0.17 | 1.37E-33 | 02_HSP-DC |
| CASP4 | 1.64720618 | 0.524 | 0.175 | 5.10E-32 | 02_HSP-DC |
| PSME1 | 1.097546865 | 0.859 | 0.5 | 8.74E-32 | 02_HSP-DC |
| RGS19 | 1.291775622 | 0.614 | 0.228 | 1.17E-30 | 02_HSP-DC |
| CASP1 | 1.17841048 | 0.556 | 0.191 | 3.24E-30 | 02_HSP-DC |
| AKR1C3 | 2.02560162 | 0.386 | 0.1 | 1.55E-29 | 02_HSP-DC |
| EIF3G | 1.01394021 | 0.801 | 0.426 | 5.01E-29 | 02_HSP-DC |
| NCOA4 | 1.187978915 | 0.73 | 0.333 | 8.45E-29 | 02_HSP-DC |
| ERP29 | 1.104418798 | 0.749 | 0.357 | 1.43E-28 | 02_HSP-DC |
| CEACAM4 | 1.888569405 | 0.344 | 0.079 | 4.16E-28 | 02_HSP-DC |
| SSR3 | 1.117387216 | 0.759 | 0.389 | 7.17E-28 | 02_HSP-DC |
| CD300A | 1.163111632 | 0.637 | 0.262 | 2.10E-27 | 02_HSP-DC |
| TMEM109 | 1.387765442 | 0.579 | 0.23 | 5.28E-27 | 02_HSP-DC |
| MNDA | 1.225578691 | 0.682 | 0.3 | 1.19E-26 | 02_HSP-DC |
| PRDX3 | 1.33005719 | 0.537 | 0.199 | 1.63E-26 | 02_HSP-DC |
| NDUFB5 | 1.270879582 | 0.514 | 0.18 | 2.89E-26 | 02_HSP-DC |
| HSPA1A | 1.743297797 | 0.801 | 0.509 | 3.83E-26 | 02_HSP-DC |
| DCK | 1.481098686 | 0.421 | 0.125 | 4.15E-26 | 02_HSP-DC |
| PSMB10 | 1.249194704 | 0.688 | 0.323 | 1.37E-25 | 02_HSP-DC |
| ETHE1 | 1.17305196 | 0.579 | 0.226 | 1.45E-25 | 02_HSP-DC |
| SERPING1 | 1.935106584 | 0.309 | 0.072 | 3.39E-25 | 02_HSP-DC |
| RTRAF | 1.1118951 | 0.666 | 0.309 | 4.14E-25 | 02_HSP-DC |
| HEBP1 | 1.697017146 | 0.341 | 0.087 | 4.67E-25 | 02_HSP-DC |
| PYCARD | 1.060048643 | 0.752 | 0.382 | 5.65E-25 | 02_HSP-DC |
| LDHB | 1.036087186 | 0.714 | 0.34 | 7.28E-25 | 02_HSP-DC |
| CLEC12A | 1.264595681 | 0.379 | 0.104 | 8.16E-25 | 02_HSP-DC |
| HLA-DQA2 | 1.247206445 | 0.412 | 0.132 | 8.36E-25 | 02_HSP-DC |
| ANKRD22 | 1.317590904 | 0.331 | 0.082 | 1.52E-24 | 02_HSP-DC |
| CAT | 1.047407478 | 0.611 | 0.247 | 1.63E-24 | 02_HSP-DC |
| LILRB3 | 1.259613353 | 0.441 | 0.141 | 4.59E-24 | 02_HSP-DC |
| DNAJB1 | 1.626945205 | 0.736 | 0.396 | 7.19E-24 | 02_HSP-DC |
| PCBP2 | 1.027559455 | 0.846 | 0.552 | 7.90E-24 | 02_HSP-DC |
| RAC2 | 1.242139433 | 0.54 | 0.207 | 9.91E-24 | 02_HSP-DC |
| OXA1L | 1.160848679 | 0.64 | 0.288 | 1.44E-23 | 02_HSP-DC |
| CSTA | 1.297248555 | 0.479 | 0.175 | 1.81E-23 | 02_HSP-DC |
| CRTAP | 1.096594393 | 0.698 | 0.346 | 2.00E-23 | 02_HSP-DC |
| CD1D | 1.984932139 | 0.302 | 0.073 | 2.44E-23 | 02_HSP-DC |
| HSD17B11 | 1.119503115 | 0.572 | 0.228 | 6.51E-23 | 02_HSP-DC |
| ATP5F1A | 1.12306766 | 0.624 | 0.288 | 9.21E-23 | 02_HSP-DC |
| APEX1 | 1.09775575 | 0.595 | 0.252 | 9.24E-23 | 02_HSP-DC |
| GSDMD | 1.315888899 | 0.405 | 0.126 | 1.28E-22 | 02_HSP-DC |
| HNRNPA1 | 1.018820416 | 0.839 | 0.613 | 1.71E-22 | 02_HSP-DC |
| PSMB8 | 1.181487298 | 0.572 | 0.244 | 2.78E-22 | 02_HSP-DC |

| | | | | | |
|---|---|---|---|---|---|
| IFITM3 | 1.043720884 | 0.91 | 0.682 | 3.10E-22 | 02_HSP-DC |
| LY6E | 1.456129381 | 0.633 | 0.319 | 4.09E-22 | 02_HSP-DC |
| CIAO2A | 1.026088125 | 0.601 | 0.26 | 4.20E-22 | 02_HSP-DC |
| CORO1B | 1.204110874 | 0.537 | 0.226 | 5.74E-22 | 02_HSP-DC |
| DENND6B | 1.706769159 | 0.325 | 0.087 | 6.05E-22 | 02_HSP-DC |
| CCT8 | 1.124330529 | 0.492 | 0.19 | 4.25E-21 | 02_HSP-DC |
| PLAAT4 | 1.670935782 | 0.399 | 0.135 | 6.50E-21 | 02_HSP-DC |
| LILRA2 | 1.228154871 | 0.389 | 0.123 | 1.03E-20 | 02_HSP-DC |
| HAUS4 | 1.365194862 | 0.395 | 0.131 | 1.13E-20 | 02_HSP-DC |
| SNX17 | 1.179266378 | 0.518 | 0.205 | 2.59E-20 | 02_HSP-DC |
| MIF4GD | 1.265505002 | 0.412 | 0.14 | 3.10E-20 | 02_HSP-DC |
| HSPA1B | 1.579287363 | 0.669 | 0.382 | 7.03E-20 | 02_HSP-DC |
| ECHS1 | 1.371337736 | 0.418 | 0.15 | 1.03E-19 | 02_HSP-DC |
| NUDT16 | 1.073095846 | 0.55 | 0.251 | 1.14E-19 | 02_HSP-DC |
| C1QBP | 1.025572083 | 0.55 | 0.234 | 1.51E-19 | 02_HSP-DC |
| FMNL2 | -2.11607493 | 0.383 | 0.613 | 1.59E-19 | 02_HSP-DC |
| CCDC115 | 1.513961004 | 0.296 | 0.077 | 1.61E-19 | 02_HSP-DC |
| PSME2 | 1.056399791 | 0.781 | 0.459 | 1.84E-19 | 02_HSP-DC |
| COMMD9 | 1.276712211 | 0.357 | 0.111 | 2.03E-19 | 02_HSP-DC |
| LAP3 | 1.001935558 | 0.752 | 0.445 | 2.46E-19 | 02_HSP-DC |
| PLBD1 | 1.189614846 | 0.457 | 0.171 | 2.77E-19 | 02_HSP-DC |
| CLEC4E | 1.415607312 | 0.363 | 0.12 | 4.08E-19 | 02_HSP-DC |
| ZFAND1 | 1.91662464 | 0.277 | 0.072 | 4.35E-19 | 02_HSP-DC |
| MAP4K4 | -2.72809358 | 0.228 | 0.481 | 6.18E-19 | 02_HSP-DC |
| TNFSF13 | 1.009432073 | 0.543 | 0.233 | 7.10E-19 | 02_HSP-DC |
| IMPDH2 | 1.16559816 | 0.466 | 0.175 | 7.72E-19 | 02_HSP-DC |
| MRPL3 | 1.166238787 | 0.437 | 0.162 | 7.86E-19 | 02_HSP-DC |
| IFI6 | 1.392508524 | 0.547 | 0.259 | 9.77E-19 | 02_HSP-DC |
| LEPROTL1 | 1.098132371 | 0.505 | 0.214 | 1.38E-18 | 02_HSP-DC |
| FCGR3A | 1.253612243 | 0.579 | 0.28 | 1.42E-18 | 02_HSP-DC |
| PDGFB | -2.60833344 | 0.164 | 0.433 | 2.09E-18 | 02_HSP-DC |
| PHB | 1.151951491 | 0.434 | 0.162 | 2.99E-18 | 02_HSP-DC |
| LYZ | 1.042407337 | 0.913 | 0.656 | 3.26E-18 | 02_HSP-DC |
| GIMAP4 | 1.448175635 | 0.35 | 0.111 | 3.73E-18 | 02_HSP-DC |
| TAP1 | 1.092279655 | 0.466 | 0.185 | 8.62E-18 | 02_HSP-DC |
| SCIMP | 1.173139275 | 0.363 | 0.118 | 1.49E-17 | 02_HSP-DC |
| PSMF1 | 1.156408982 | 0.502 | 0.209 | 2.08E-17 | 02_HSP-DC |
| CLEC4A | 1.167473397 | 0.437 | 0.166 | 2.58E-17 | 02_HSP-DC |
| SLC66A3 | 1.370976122 | 0.379 | 0.135 | 5.23E-17 | 02_HSP-DC |
| GNG10 | 1.062699381 | 0.543 | 0.247 | 5.49E-17 | 02_HSP-DC |
| APOO | 1.523053214 | 0.531 | 0.28 | 6.76E-17 | 02_HSP-DC |
| GLO1 | 1.311710877 | 0.415 | 0.165 | 2.31E-16 | 02_HSP-DC |
| BCKDHA | 1.451714519 | 0.305 | 0.097 | 3.87E-16 | 02_HSP-DC |
| SLC7A7 | 1.374966021 | 0.35 | 0.121 | 5.14E-16 | 02_HSP-DC |
| B3GAT3 | 1.193237261 | 0.322 | 0.101 | 6.98E-16 | 02_HSP-DC |
| POLR2G | 1.169055687 | 0.441 | 0.177 | 7.21E-16 | 02_HSP-DC |

| | | | | | |
|---|---|---|---|---|---|
| CDKN1B | 1.148067575 | 0.431 | 0.167 | 9.29E-16 | 02_HSP-DC |
| LSM6 | 1.232456948 | 0.334 | 0.112 | 1.30E-15 | 02_HSP-DC |
| HAX1 | 1.118453521 | 0.392 | 0.15 | 3.90E-15 | 02_HSP-DC |
| COA4 | 1.205651248 | 0.35 | 0.124 | 5.24E-15 | 02_HSP-DC |
| POLE4 | 1.314697224 | 0.344 | 0.121 | 6.83E-15 | 02_HSP-DC |
| S100A9 | 1.112349053 | 0.299 | 0.097 | 9.54E-15 | 02_HSP-DC |
| TAX1BP3 | 1.008792041 | 0.428 | 0.169 | 9.55E-15 | 02_HSP-DC |
| NDRG2 | 1.132366717 | 0.453 | 0.194 | 1.04E-14 | 02_HSP-DC |
| DENND5A | -2.10232293 | 0.232 | 0.464 | 1.12E-14 | 02_HSP-DC |
| CCL4 | -1.21074021 | 0.46 | 0.676 | 1.25E-14 | 02_HSP-DC |
| SELENOK | 1.120512808 | 0.826 | 0.546 | 1.41E-14 | 02_HSP-DC |
| TMEM70 | 1.396050403 | 0.405 | 0.172 | 1.92E-14 | 02_HSP-DC |
| SUMF2 | 1.089305745 | 0.354 | 0.126 | 2.00E-14 | 02_HSP-DC |
| LMNA | -1.9858808 | 0.277 | 0.51 | 2.97E-14 | 02_HSP-DC |
| HBEGF | 1.053620886 | 0.621 | 0.387 | 6.07E-14 | 02_HSP-DC |
| IMPDH1 | 1.127974591 | 0.331 | 0.115 | 6.31E-14 | 02_HSP-DC |
| TMEM176B | 1.385660461 | 0.399 | 0.17 | 7.66E-14 | 02_HSP-DC |
| IMP4 | 1.278168647 | 0.322 | 0.112 | 1.06E-13 | 02_HSP-DC |
| SVBP | 1.090335588 | 0.431 | 0.184 | 1.71E-13 | 02_HSP-DC |
| RNF167 | 1.12253437 | 0.347 | 0.127 | 1.96E-13 | 02_HSP-DC |
| CD33 | 1.041648568 | 0.363 | 0.14 | 2.66E-13 | 02_HSP-DC |
| SLC12A9 | 1.034606987 | 0.357 | 0.132 | 3.38E-13 | 02_HSP-DC |
| COMMD7 | 1.042462983 | 0.344 | 0.129 | 4.85E-13 | 02_HSP-DC |
| MPG | 1.268175807 | 0.35 | 0.135 | 5.07E-13 | 02_HSP-DC |
| DDRGK1 | 1.107475974 | 0.367 | 0.142 | 5.92E-13 | 02_HSP-DC |
| ECH1 | 1.133593576 | 0.428 | 0.186 | 8.24E-13 | 02_HSP-DC |
| CACYBP | 1.20108975 | 0.373 | 0.15 | 1.04E-12 | 02_HSP-DC |
| AKR7A2 | 1.354828904 | 0.334 | 0.127 | 1.94E-12 | 02_HSP-DC |
| PSMG2 | 1.020074702 | 0.45 | 0.208 | 2.03E-12 | 02_HSP-DC |
| CCT7 | 1.044661202 | 0.412 | 0.172 | 2.22E-12 | 02_HSP-DC |
| GRHPR | 1.020516636 | 0.373 | 0.146 | 2.36E-12 | 02_HSP-DC |
| CCL22 | -3.19959612 | 0.09 | 0.309 | 4.26E-12 | 02_HSP-DC |
| TSEN34 | 1.019751009 | 0.376 | 0.154 | 7.52E-12 | 02_HSP-DC |
| HIVEP1 | -2.32941385 | 0.158 | 0.377 | 1.33E-11 | 02_HSP-DC |
| LSM2 | 1.090385771 | 0.373 | 0.159 | 2.87E-11 | 02_HSP-DC |
| SHKBP1 | 1.009254993 | 0.399 | 0.17 | 4.12E-11 | 02_HSP-DC |
| CCR7 | -2.06622509 | 0.138 | 0.353 | 7.10E-11 | 02_HSP-DC |
| CNPY2 | 1.014624583 | 0.354 | 0.141 | 8.15E-11 | 02_HSP-DC |
| COPS3 | 1.013035741 | 0.363 | 0.151 | 1.27E-10 | 02_HSP-DC |
| G0S2 | -2.23026313 | 0.244 | 0.452 | 1.56E-10 | 02_HSP-DC |
| RASAL2 | -2.43431307 | 0.148 | 0.352 | 2.08E-10 | 02_HSP-DC |
| USP12 | -2.14591985 | 0.151 | 0.358 | 2.22E-10 | 02_HSP-DC |
| PEAK1 | -2.12677371 | 0.19 | 0.393 | 5.35E-10 | 02_HSP-DC |
| NDUFB3 | 1.010794499 | 0.363 | 0.16 | 1.86E-09 | 02_HSP-DC |
| MTRNR2L8 | 8.329858963 | 0.746 | 0.038 | 4.05E-176 | 03_Mito-DC |
| MTRNR2L12 | 2.586719363 | 0.89 | 0.167 | 1.29E-106 | 03_Mito-DC |

| | | | | | |
|---|---|---|---|---|---|
| XIST | 3.200976836 | 0.565 | 0.085 | 8.18E-72 | 03_Mito-DC |
| DOCK4 | 1.97385904 | 0.99 | 0.748 | 3.78E-62 | 03_Mito-DC |
| ACKR3 | 3.321287454 | 0.498 | 0.085 | 1.00E-56 | 03_Mito-DC |
| RPS5 | -1.99947377 | 0.55 | 0.878 | 1.43E-48 | 03_Mito-DC |
| RPL8 | -1.60234634 | 0.703 | 0.922 | 7.54E-45 | 03_Mito-DC |
| RPS27A | -1.50851249 | 0.718 | 0.933 | 1.01E-43 | 03_Mito-DC |
| ELL2 | 1.71087341 | 0.952 | 0.708 | 1.24E-43 | 03_Mito-DC |
| RPL11 | -1.48591305 | 0.699 | 0.94 | 1.32E-42 | 03_Mito-DC |
| AC007952.4 | 2.946439936 | 0.407 | 0.07 | 2.27E-42 | 03_Mito-DC |
| TANC2 | 2.318486733 | 0.732 | 0.331 | 2.73E-41 | 03_Mito-DC |
| RPL21 | -1.77306393 | 0.603 | 0.876 | 3.46E-41 | 03_Mito-DC |
| IFITM2 | -2.80477805 | 0.407 | 0.798 | 1.57E-40 | 03_Mito-DC |
| SFMBT2 | 1.806088794 | 0.823 | 0.392 | 1.64E-40 | 03_Mito-DC |
| RPL15 | -1.48567712 | 0.675 | 0.916 | 3.53E-40 | 03_Mito-DC |
| RPS13 | -1.44274202 | 0.675 | 0.93 | 1.08E-38 | 03_Mito-DC |
| EEF1B2 | -1.81006966 | 0.44 | 0.851 | 1.77E-38 | 03_Mito-DC |
| LRMDA | 1.636316234 | 0.947 | 0.66 | 2.34E-37 | 03_Mito-DC |
| SLC1A3 | 1.473981324 | 0.914 | 0.464 | 2.91E-37 | 03_Mito-DC |
| RPL10A | -1.57100038 | 0.55 | 0.87 | 5.09E-37 | 03_Mito-DC |
| MT2A | -3.61819209 | 0.258 | 0.726 | 5.38E-37 | 03_Mito-DC |
| ARHGAP24 | 1.67288676 | 0.742 | 0.313 | 3.00E-36 | 03_Mito-DC |
| KCNQ3 | 2.040332057 | 0.579 | 0.17 | 3.19E-36 | 03_Mito-DC |
| RPS7 | -1.34501336 | 0.703 | 0.92 | 6.08E-36 | 03_Mito-DC |
| RPL35A | -1.4205985 | 0.646 | 0.9 | 2.45E-35 | 03_Mito-DC |
| CCNH | 1.450543031 | 0.962 | 0.761 | 3.09E-35 | 03_Mito-DC |
| RPS3 | -1.32682683 | 0.67 | 0.934 | 4.52E-35 | 03_Mito-DC |
| MAN2A1 | 1.836465813 | 0.78 | 0.408 | 6.94E-35 | 03_Mito-DC |
| RPS16 | -1.35821483 | 0.727 | 0.928 | 7.27E-35 | 03_Mito-DC |
| RPL17 | -1.41978885 | 0.651 | 0.899 | 1.77E-34 | 03_Mito-DC |
| SLC8A1 | 1.49377399 | 0.938 | 0.711 | 1.92E-34 | 03_Mito-DC |
| RPS6 | -1.28053573 | 0.722 | 0.923 | 4.03E-34 | 03_Mito-DC |
| OLR1 | 1.384661776 | 0.89 | 0.628 | 7.03E-33 | 03_Mito-DC |
| S100A10 | -2.29019521 | 0.435 | 0.793 | 9.44E-33 | 03_Mito-DC |
| DLEU1 | 1.555192775 | 0.722 | 0.318 | 2.06E-32 | 03_Mito-DC |
| RPSA | -1.40765622 | 0.522 | 0.879 | 2.92E-32 | 03_Mito-DC |
| RASGEF1B | 1.392182109 | 0.923 | 0.651 | 6.85E-32 | 03_Mito-DC |
| CST7 | -4.44363117 | 0.077 | 0.554 | 2.48E-31 | 03_Mito-DC |
| ZNF331 | 1.528465732 | 0.933 | 0.648 | 3.05E-31 | 03_Mito-DC |
| ARPC1B | -1.61645766 | 0.536 | 0.829 | 4.18E-31 | 03_Mito-DC |
| SRGAP1 | 1.52023626 | 0.852 | 0.523 | 8.14E-31 | 03_Mito-DC |
| RPL7 | -1.62552908 | 0.416 | 0.807 | 2.08E-30 | 03_Mito-DC |
| RPLP0 | -1.25157603 | 0.713 | 0.913 | 2.12E-30 | 03_Mito-DC |
| RPS4Y1 | -2.93368768 | 0.067 | 0.567 | 3.93E-30 | 03_Mito-DC |
| RPL23A | -1.34393085 | 0.565 | 0.874 | 7.48E-30 | 03_Mito-DC |
| FRMD4A | 1.410353859 | 0.804 | 0.384 | 1.16E-29 | 03_Mito-DC |
| SLC11A1 | 1.412785502 | 0.794 | 0.4 | 2.14E-29 | 03_Mito-DC |

| | | | | | |
|---|---|---|---|---|---|
| MACC1 | 2.38240515 | 0.512 | 0.167 | 2.54E-29 | 03_Mito-DC |
| RACK1 | -1.26590919 | 0.608 | 0.896 | 2.99E-29 | 03_Mito-DC |
| RPS3A | -1.26117355 | 0.713 | 0.915 | 3.52E-29 | 03_Mito-DC |
| RPS14 | -1.16014078 | 0.718 | 0.926 | 7.62E-29 | 03_Mito-DC |
| FGD4 | 1.455835491 | 0.909 | 0.661 | 1.23E-28 | 03_Mito-DC |
| TAGLN2 | -2.1758632 | 0.359 | 0.733 | 1.55E-28 | 03_Mito-DC |
| RPL13A | -1.1732263 | 0.651 | 0.89 | 2.21E-28 | 03_Mito-DC |
| CLIC1 | -1.39826242 | 0.493 | 0.835 | 4.10E-28 | 03_Mito-DC |
| CLDN1 | 3.68679258 | 0.268 | 0.042 | 5.80E-28 | 03_Mito-DC |
| AL136987.1 | 2.199462712 | 0.541 | 0.201 | 1.14E-27 | 03_Mito-DC |
| PFDN5 | -1.22888178 | 0.603 | 0.871 | 4.95E-27 | 03_Mito-DC |
| RPL6 | -1.18486546 | 0.684 | 0.898 | 6.91E-27 | 03_Mito-DC |
| RPL5 | -1.26299785 | 0.66 | 0.897 | 7.26E-27 | 03_Mito-DC |
| UBA52 | -1.14350993 | 0.612 | 0.886 | 3.75E-26 | 03_Mito-DC |
| ACTG1 | -1.44138775 | 0.679 | 0.903 | 1.44E-25 | 03_Mito-DC |
| SLC25A6 | -1.51039307 | 0.44 | 0.807 | 2.38E-25 | 03_Mito-DC |
| MSR1 | 1.429135047 | 0.818 | 0.49 | 3.16E-25 | 03_Mito-DC |
| EEF1D | -1.27482175 | 0.569 | 0.843 | 5.60E-25 | 03_Mito-DC |
| C3 | 1.111065291 | 0.856 | 0.463 | 6.42E-25 | 03_Mito-DC |
| HINT1 | -1.69891554 | 0.34 | 0.73 | 7.16E-25 | 03_Mito-DC |
| PFN1 | -1.03757893 | 0.722 | 0.933 | 7.31E-25 | 03_Mito-DC |
| QKI | 1.265618229 | 0.904 | 0.676 | 8.02E-25 | 03_Mito-DC |
| FP671120.4 | -1.75927173 | 0.225 | 0.642 | 9.13E-25 | 03_Mito-DC |
| CRIP1 | -4.09105738 | 0.072 | 0.496 | 1.37E-24 | 03_Mito-DC |
| ALDOA | -1.98259823 | 0.249 | 0.67 | 1.41E-24 | 03_Mito-DC |
| EEF1G | -1.27777363 | 0.589 | 0.852 | 5.07E-24 | 03_Mito-DC |
| GSTP1 | -1.45051858 | 0.435 | 0.794 | 6.00E-24 | 03_Mito-DC |
| FAM110B | 2.343937616 | 0.416 | 0.126 | 1.00E-23 | 03_Mito-DC |
| AL163541.1 | 1.990194843 | 0.464 | 0.155 | 2.97E-23 | 03_Mito-DC |
| BTF3 | -1.22460762 | 0.526 | 0.838 | 3.31E-23 | 03_Mito-DC |
| MEIKIN | 2.435980843 | 0.464 | 0.166 | 4.95E-23 | 03_Mito-DC |
| DPYD | 1.303896802 | 0.861 | 0.566 | 5.94E-23 | 03_Mito-DC |
| DHX34 | 2.174013536 | 0.431 | 0.136 | 7.44E-23 | 03_Mito-DC |
| MS4A14 | 2.689379453 | 0.282 | 0.057 | 1.04E-22 | 03_Mito-DC |
| TUBA1B | -1.3197895 | 0.603 | 0.858 | 1.19E-22 | 03_Mito-DC |
| TAOK3 | 1.314551257 | 0.861 | 0.643 | 1.73E-22 | 03_Mito-DC |
| MYL6 | -1.0305108 | 0.656 | 0.905 | 2.62E-22 | 03_Mito-DC |
| SYNGR2 | -1.47401552 | 0.44 | 0.785 | 3.30E-22 | 03_Mito-DC |
| PADI2 | 1.564683191 | 0.775 | 0.519 | 3.41E-22 | 03_Mito-DC |
| PPA1 | -2.00963751 | 0.292 | 0.661 | 3.60E-22 | 03_Mito-DC |
| LDHA | -1.63871788 | 0.44 | 0.769 | 3.82E-22 | 03_Mito-DC |
| ARHGAP15 | 1.1528918 | 0.852 | 0.615 | 4.53E-22 | 03_Mito-DC |
| SERPINB1 | -1.84384935 | 0.426 | 0.759 | 5.53E-22 | 03_Mito-DC |
| KCNE1 | 2.487304334 | 0.301 | 0.069 | 2.43E-21 | 03_Mito-DC |
| USP53 | 1.29538022 | 0.689 | 0.359 | 3.94E-21 | 03_Mito-DC |
| CLEC10A | -4.08377142 | 0.038 | 0.42 | 7.16E-21 | 03_Mito-DC |

| | | | | | |
|---|---|---|---|---|---|
| ATF3 | 1.198617782 | 0.813 | 0.545 | 8.04E-21 | 03_Mito-DC |
| INSIG1 | -1.90952632 | 0.268 | 0.633 | 1.08E-20 | 03_Mito-DC |
| APOC1 | 1.227249598 | 0.785 | 0.454 | 1.16E-20 | 03_Mito-DC |
| ARHGAP26 | 1.327559715 | 0.842 | 0.612 | 1.62E-20 | 03_Mito-DC |
| ARF6 | -2.10020474 | 0.292 | 0.646 | 1.77E-20 | 03_Mito-DC |
| CRYBG1 | -1.83059502 | 0.258 | 0.635 | 3.70E-20 | 03_Mito-DC |
| DAGLB | 1.502907083 | 0.656 | 0.346 | 5.00E-20 | 03_Mito-DC |
| RPS11 | -1.10801076 | 0.517 | 0.841 | 5.00E-20 | 03_Mito-DC |
| RPL4 | -1.10570682 | 0.589 | 0.857 | 5.19E-20 | 03_Mito-DC |
| ARPC3 | -1.03194313 | 0.593 | 0.867 | 5.52E-20 | 03_Mito-DC |
| EIF3K | -1.35648838 | 0.392 | 0.747 | 1.37E-19 | 03_Mito-DC |
| FNDC3A | 1.855834087 | 0.612 | 0.31 | 1.91E-19 | 03_Mito-DC |
| BNC2 | 2.02660085 | 0.421 | 0.146 | 2.45E-19 | 03_Mito-DC |
| PDE8A | 1.557502591 | 0.694 | 0.394 | 2.73E-19 | 03_Mito-DC |
| CD1C | -4.57407296 | 0.029 | 0.387 | 3.46E-19 | 03_Mito-DC |
| PRDM1 | 1.741516834 | 0.55 | 0.246 | 5.19E-19 | 03_Mito-DC |
| IL1R2 | -2.63070252 | 0.163 | 0.526 | 5.47E-19 | 03_Mito-DC |
| COTL1 | -1.11152523 | 0.608 | 0.864 | 8.39E-19 | 03_Mito-DC |
| RNASEK | -1.0569531 | 0.526 | 0.834 | 8.93E-19 | 03_Mito-DC |
| FCER1A | -2.75047648 | 0.172 | 0.52 | 1.12E-18 | 03_Mito-DC |
| CD9 | 1.560925296 | 0.641 | 0.346 | 1.34E-18 | 03_Mito-DC |
| ADAM17 | 1.453129529 | 0.689 | 0.395 | 1.59E-18 | 03_Mito-DC |
| ZNF618 | 1.98239672 | 0.368 | 0.113 | 1.68E-18 | 03_Mito-DC |
| CSMD1 | 1.719826895 | 0.306 | 0.08 | 1.94E-18 | 03_Mito-DC |
| AC138123.1 | -3.01687389 | 0.043 | 0.405 | 3.47E-18 | 03_Mito-DC |
| TEX14 | 1.568621324 | 0.675 | 0.405 | 4.22E-18 | 03_Mito-DC |
| ATP8B4 | 1.443673535 | 0.708 | 0.421 | 4.36E-18 | 03_Mito-DC |
| TIMP1 | -2.51244074 | 0.435 | 0.717 | 4.93E-18 | 03_Mito-DC |
| SLC25A5 | -1.33414667 | 0.397 | 0.726 | 6.98E-18 | 03_Mito-DC |
| UTRN | 1.273129698 | 0.737 | 0.483 | 9.00E-18 | 03_Mito-DC |
| AL390957.1 | 1.736592014 | 0.579 | 0.284 | 1.02E-17 | 03_Mito-DC |
| MYL12A | -1.26835068 | 0.478 | 0.767 | 1.16E-17 | 03_Mito-DC |
| PRMT9 | 1.768114631 | 0.598 | 0.314 | 1.69E-17 | 03_Mito-DC |
| CSF3R | 1.250373626 | 0.746 | 0.5 | 1.69E-17 | 03_Mito-DC |
| TSC22D2 | 1.50731382 | 0.727 | 0.474 | 1.90E-17 | 03_Mito-DC |
| IFITM1 | -2.60986654 | 0.1 | 0.455 | 2.24E-17 | 03_Mito-DC |
| LSP1 | -1.30805687 | 0.493 | 0.758 | 7.57E-17 | 03_Mito-DC |
| TPST1 | 2.3971134 | 0.292 | 0.078 | 8.40E-17 | 03_Mito-DC |
| EIF3L | -1.71084427 | 0.273 | 0.601 | 1.05E-16 | 03_Mito-DC |
| MIF | -1.18966796 | 0.445 | 0.777 | 2.48E-16 | 03_Mito-DC |
| ADAM19 | -6.40712386 | 0.014 | 0.332 | 2.64E-16 | 03_Mito-DC |
| ARF5 | -1.64835549 | 0.249 | 0.598 | 2.78E-16 | 03_Mito-DC |
| CLEC5A | 1.481783429 | 0.584 | 0.306 | 3.98E-16 | 03_Mito-DC |
| KLHL6 | 1.293028247 | 0.703 | 0.432 | 4.44E-16 | 03_Mito-DC |
| PGLS | -1.96240194 | 0.187 | 0.523 | 7.59E-16 | 03_Mito-DC |
| MAP4K3 | 1.421986096 | 0.689 | 0.432 | 1.08E-15 | 03_Mito-DC |

| | | | | | |
|---|---|---|---|---|---|
| PAG1 | 1.552276102 | 0.617 | 0.364 | 1.22E-15 | 03_Mito-DC |
| SLC25A3 | -1.03802881 | 0.522 | 0.796 | 1.87E-15 | 03_Mito-DC |
| PRKAG2 | 1.369807393 | 0.708 | 0.471 | 1.94E-15 | 03_Mito-DC |
| SAP30 | 1.51023345 | 0.584 | 0.329 | 2.64E-15 | 03_Mito-DC |
| ARID5B | 1.434290321 | 0.684 | 0.431 | 3.91E-15 | 03_Mito-DC |
| CHCHD2 | -1.02716216 | 0.435 | 0.78 | 5.09E-15 | 03_Mito-DC |
| IFITM10 | 1.974961363 | 0.292 | 0.085 | 1.11E-14 | 03_Mito-DC |
| GAREM1 | -1.97432943 | 0.115 | 0.461 | 1.59E-14 | 03_Mito-DC |
| SMYD3 | 1.996667671 | 0.426 | 0.184 | 1.65E-14 | 03_Mito-DC |
| EIF3F | -1.33885072 | 0.335 | 0.659 | 1.69E-14 | 03_Mito-DC |
| RASAL2 | 1.401280672 | 0.555 | 0.271 | 2.03E-14 | 03_Mito-DC |
| CCDC138 | 1.629574029 | 0.397 | 0.152 | 2.36E-14 | 03_Mito-DC |
| IFITM3 | -1.5145405 | 0.531 | 0.761 | 2.74E-14 | 03_Mito-DC |
| SPINT2 | -1.66611762 | 0.244 | 0.561 | 2.94E-14 | 03_Mito-DC |
| AC020916.1 | 1.392730237 | 0.722 | 0.498 | 3.13E-14 | 03_Mito-DC |
| SKP1 | -1.2038579 | 0.349 | 0.687 | 3.28E-14 | 03_Mito-DC |
| CSGALNACT1 | 1.242700853 | 0.593 | 0.309 | 3.41E-14 | 03_Mito-DC |
| DBI | -1.15261397 | 0.498 | 0.785 | 4.22E-14 | 03_Mito-DC |
| BCL3 | -2.20705983 | 0.215 | 0.517 | 8.13E-14 | 03_Mito-DC |
| NDUFV2 | -1.5096254 | 0.297 | 0.617 | 9.67E-14 | 03_Mito-DC |
| NHSL1 | 1.568450097 | 0.665 | 0.412 | 1.39E-13 | 03_Mito-DC |
| ARIH1 | 1.180468988 | 0.737 | 0.529 | 1.58E-13 | 03_Mito-DC |
| MCTP1 | 1.462876954 | 0.641 | 0.397 | 1.86E-13 | 03_Mito-DC |
| FHIT | 1.528041902 | 0.545 | 0.29 | 2.32E-13 | 03_Mito-DC |
| USP36 | 1.772668424 | 0.617 | 0.377 | 2.59E-13 | 03_Mito-DC |
| PRKCH | 1.388040791 | 0.675 | 0.46 | 2.74E-13 | 03_Mito-DC |
| WWOX | 1.750959996 | 0.474 | 0.226 | 3.30E-13 | 03_Mito-DC |
| CYTH3 | 1.821841562 | 0.426 | 0.187 | 8.27E-13 | 03_Mito-DC |
| AC245014.3 | 1.850929751 | 0.344 | 0.126 | 1.27E-12 | 03_Mito-DC |
| ZBTB16 | 1.021105482 | 0.732 | 0.45 | 1.43E-12 | 03_Mito-DC |
| FOXO3 | 1.338922798 | 0.641 | 0.392 | 1.63E-12 | 03_Mito-DC |
| PSME1 | -1.51070821 | 0.325 | 0.613 | 2.25E-12 | 03_Mito-DC |
| CKB | 1.953792372 | 0.383 | 0.164 | 3.40E-12 | 03_Mito-DC |
| STX17-AS1 | 1.199025203 | 0.699 | 0.488 | 3.54E-12 | 03_Mito-DC |
| TXNRD1 | -2.10497406 | 0.144 | 0.446 | 4.01E-12 | 03_Mito-DC |
| DLEU2 | 1.26695186 | 0.665 | 0.459 | 4.26E-12 | 03_Mito-DC |
| EMP3 | -1.43936672 | 0.474 | 0.72 | 4.33E-12 | 03_Mito-DC |
| ANXA11 | -1.36423352 | 0.234 | 0.547 | 4.50E-12 | 03_Mito-DC |
| RERE | 1.288722456 | 0.612 | 0.357 | 5.90E-12 | 03_Mito-DC |
| TUBA1A | -1.76425805 | 0.282 | 0.577 | 8.11E-12 | 03_Mito-DC |
| RHBDF2 | 1.004133589 | 0.732 | 0.508 | 9.05E-12 | 03_Mito-DC |
| RPL23 | -1.02203178 | 0.44 | 0.743 | 1.00E-11 | 03_Mito-DC |
| DTNA | 1.702403348 | 0.44 | 0.21 | 1.13E-11 | 03_Mito-DC |
| SULF2 | -2.1930167 | 0.081 | 0.37 | 1.39E-11 | 03_Mito-DC |
| RPS20 | -1.04518407 | 0.407 | 0.697 | 1.50E-11 | 03_Mito-DC |
| RALGAPA1 | 1.148662997 | 0.565 | 0.326 | 1.56E-11 | 03_Mito-DC |

| | | | | | |
|---|---|---|---|---|---|
| DAPP1 | -3.60341149 | 0.057 | 0.328 | 1.85E-11 | 03_Mito-DC |
| RGS13 | 2.004333736 | 0.368 | 0.158 | 2.03E-11 | 03_Mito-DC |
| TOMM7 | -1.03929923 | 0.335 | 0.679 | 2.04E-11 | 03_Mito-DC |
| PLA2G4A | 1.296256521 | 0.545 | 0.305 | 2.33E-11 | 03_Mito-DC |
| PSME4 | 1.440112886 | 0.536 | 0.299 | 2.96E-11 | 03_Mito-DC |
| PIM3 | -1.86559814 | 0.206 | 0.497 | 3.81E-11 | 03_Mito-DC |
| AOAH | 1.134073812 | 0.684 | 0.475 | 3.85E-11 | 03_Mito-DC |
| JDP2 | 1.096599289 | 0.646 | 0.423 | 4.34E-11 | 03_Mito-DC |
| INPP5D | 1.100260012 | 0.641 | 0.417 | 4.72E-11 | 03_Mito-DC |
| APOO | -2.24272285 | 0.086 | 0.371 | 4.73E-11 | 03_Mito-DC |
| IL2RG | -2.40858877 | 0.115 | 0.391 | 4.95E-11 | 03_Mito-DC |
| LNCAROD | 1.335204288 | 0.378 | 0.157 | 6.96E-11 | 03_Mito-DC |
| LINC00910 | 1.768138607 | 0.383 | 0.166 | 8.80E-11 | 03_Mito-DC |
| FMNL2 | 1.081018141 | 0.742 | 0.537 | 9.39E-11 | 03_Mito-DC |
| SNHG29 | -1.09027179 | 0.435 | 0.7 | 1.07E-10 | 03_Mito-DC |
| DUSP5 | -2.16044932 | 0.196 | 0.475 | 1.16E-10 | 03_Mito-DC |
| COMMD6 | -1.04997457 | 0.354 | 0.675 | 1.32E-10 | 03_Mito-DC |
| ADAMTSL4-AS | 1.35717537 | 0.617 | 0.405 | 1.48E-10 | 03_Mito-DC |
| RASA1 | 1.098054766 | 0.569 | 0.343 | 1.55E-10 | 03_Mito-DC |
| WBP1L | 1.516215085 | 0.464 | 0.246 | 1.80E-10 | 03_Mito-DC |
| CCL22 | -4.43195742 | 0.048 | 0.299 | 2.07E-10 | 03_Mito-DC |
| CD99 | -1.32803511 | 0.268 | 0.562 | 2.15E-10 | 03_Mito-DC |
| AL137857.1 | -3.37228451 | 0.038 | 0.29 | 2.61E-10 | 03_Mito-DC |
| TNFSF13B | -1.47194946 | 0.206 | 0.503 | 3.25E-10 | 03_Mito-DC |
| H2AFZ | -1.26295386 | 0.383 | 0.673 | 4.32E-10 | 03_Mito-DC |
| SORL1 | 1.030476379 | 0.651 | 0.433 | 4.36E-10 | 03_Mito-DC |
| STK38L | 1.209721614 | 0.641 | 0.431 | 5.14E-10 | 03_Mito-DC |
| CNN2 | -1.50837069 | 0.211 | 0.492 | 5.85E-10 | 03_Mito-DC |
| RAB5C | -1.24370484 | 0.311 | 0.583 | 6.28E-10 | 03_Mito-DC |
| DISC1 | 1.362435776 | 0.493 | 0.273 | 8.15E-10 | 03_Mito-DC |
| FKBP1A | -1.28860022 | 0.244 | 0.535 | 1.17E-09 | 03_Mito-DC |
| KCNMA1 | 1.563689351 | 0.344 | 0.141 | 1.33E-09 | 03_Mito-DC |
| NDUFA11 | -1.13020608 | 0.306 | 0.601 | 1.55E-09 | 03_Mito-DC |
| ITPR1 | 1.334205388 | 0.517 | 0.297 | 1.64E-09 | 03_Mito-DC |
| RHOF | -2.94591536 | 0.072 | 0.32 | 1.64E-09 | 03_Mito-DC |
| GALNT2 | 1.308723854 | 0.502 | 0.275 | 1.81E-09 | 03_Mito-DC |
| TCF12 | 1.137540023 | 0.718 | 0.51 | 1.87E-09 | 03_Mito-DC |
| WDR83OS | -1.13889279 | 0.311 | 0.575 | 1.92E-09 | 03_Mito-DC |
| VDAC2 | -1.26916506 | 0.244 | 0.536 | 1.96E-09 | 03_Mito-DC |
| DDAH2 | -1.51626957 | 0.215 | 0.493 | 2.64E-09 | 03_Mito-DC |
| HIF1A-AS3 | 1.189057105 | 0.507 | 0.283 | 3.05E-09 | 03_Mito-DC |
| BRK1 | -1.15209027 | 0.349 | 0.617 | 3.22E-09 | 03_Mito-DC |
| AC004448.2 | -1.80774345 | 0.024 | 0.263 | 3.72E-09 | 03_Mito-DC |
| MRC1 | -1.3655603 | 0.057 | 0.319 | 3.82E-09 | 03_Mito-DC |
| HIGD2A | -1.0699955 | 0.388 | 0.648 | 3.83E-09 | 03_Mito-DC |
| PALD1 | 1.048924998 | 0.675 | 0.473 | 3.96E-09 | 03_Mito-DC |

| | | | | | |
|---|---|---|---|---|---|
| HMGA1 | -1.78285481 | 0.172 | 0.432 | 4.89E-09 | 03_Mito-DC |
| THBS1 | -1.87024806 | 0.139 | 0.4 | 5.39E-09 | 03_Mito-DC |
| RCOR1 | 1.191323736 | 0.569 | 0.358 | 6.11E-09 | 03_Mito-DC |
| ABCC4 | 1.331558094 | 0.435 | 0.216 | 7.49E-09 | 03_Mito-DC |
| ANXA2 | -1.10170729 | 0.421 | 0.669 | 1.09E-08 | 03_Mito-DC |
| RAN | -1.02352276 | 0.311 | 0.611 | 1.34E-08 | 03_Mito-DC |
| SUPT4H1 | -1.5869337 | 0.167 | 0.425 | 1.36E-08 | 03_Mito-DC |
| FAM53C | 1.216057439 | 0.478 | 0.267 | 1.43E-08 | 03_Mito-DC |
| FNIP1 | 1.142361107 | 0.565 | 0.361 | 1.48E-08 | 03_Mito-DC |
| ATP5PO | -1.04219938 | 0.301 | 0.583 | 1.48E-08 | 03_Mito-DC |
| AP001636.3 | 1.465559229 | 0.383 | 0.18 | 1.59E-08 | 03_Mito-DC |
| AP1S2 | -1.35769754 | 0.234 | 0.503 | 2.18E-08 | 03_Mito-DC |
| UBAP1 | 1.277800219 | 0.55 | 0.349 | 3.96E-08 | 03_Mito-DC |
| SLC4A7 | 1.327205844 | 0.517 | 0.303 | 4.10E-08 | 03_Mito-DC |
| PSMB1 | -1.05637053 | 0.297 | 0.575 | 4.45E-08 | 03_Mito-DC |
| IL7R | -4.25131142 | 0.048 | 0.267 | 4.82E-08 | 03_Mito-DC |
| CFP | -4.07082943 | 0.014 | 0.223 | 5.84E-08 | 03_Mito-DC |
| SNRPD3 | -2.51983649 | 0.048 | 0.27 | 6.25E-08 | 03_Mito-DC |
| PSMA2 | -1.54775834 | 0.177 | 0.425 | 7.81E-08 | 03_Mito-DC |
| GRINA | -1.0006843 | 0.455 | 0.679 | 9.95E-08 | 03_Mito-DC |
| RNASE6 | -1.23641087 | 0.354 | 0.601 | 1.01E-07 | 03_Mito-DC |
| CD69 | 1.023439836 | 0.632 | 0.406 | 1.20E-07 | 03_Mito-DC |
| UBE2F | -1.25924431 | 0.249 | 0.515 | 1.28E-07 | 03_Mito-DC |
| UQCRFS1 | -1.13489435 | 0.211 | 0.465 | 1.36E-07 | 03_Mito-DC |
| NINJ1 | -1.40812953 | 0.373 | 0.583 | 1.36E-07 | 03_Mito-DC |
| UXT | -1.13523939 | 0.234 | 0.493 | 1.38E-07 | 03_Mito-DC |
| ATP2C1 | 1.272713987 | 0.498 | 0.287 | 1.51E-07 | 03_Mito-DC |
| GPR137B | -1.17460705 | 0.249 | 0.511 | 1.52E-07 | 03_Mito-DC |
| TALDO1 | -1.20332415 | 0.211 | 0.473 | 1.56E-07 | 03_Mito-DC |
| SBF2 | 1.143554803 | 0.541 | 0.34 | 1.77E-07 | 03_Mito-DC |
| MAF1 | -1.85985396 | 0.096 | 0.333 | 1.90E-07 | 03_Mito-DC |
| ERP29 | -1.40969469 | 0.225 | 0.471 | 2.38E-07 | 03_Mito-DC |
| G0S2 | -2.51156388 | 0.22 | 0.44 | 2.88E-07 | 03_Mito-DC |
| LRRK2 | 1.387622343 | 0.522 | 0.322 | 3.04E-07 | 03_Mito-DC |
| CD1E | -3.72796206 | 0.038 | 0.241 | 3.60E-07 | 03_Mito-DC |
| NDUFA6 | -1.24477327 | 0.22 | 0.47 | 4.50E-07 | 03_Mito-DC |
| UCP2 | -1.53388093 | 0.239 | 0.477 | 4.94E-07 | 03_Mito-DC |
| TXN | -2.75177236 | 0.254 | 0.486 | 5.08E-07 | 03_Mito-DC |
| EIF3E | -1.05552642 | 0.34 | 0.58 | 5.33E-07 | 03_Mito-DC |
| EIF3G | -1.04162604 | 0.282 | 0.538 | 5.79E-07 | 03_Mito-DC |
| COX7A2L | -1.31472378 | 0.153 | 0.394 | 6.32E-07 | 03_Mito-DC |
| PRELID1 | -1.17188595 | 0.268 | 0.51 | 6.34E-07 | 03_Mito-DC |
| SLC41A2 | -2.29701962 | 0.086 | 0.311 | 6.95E-07 | 03_Mito-DC |
| GYPC | -1.35236288 | 0.115 | 0.355 | 8.20E-07 | 03_Mito-DC |
| HNRNPA0 | -1.03117899 | 0.368 | 0.608 | 1.07E-06 | 03_Mito-DC |
| UBE2I | -1.29465324 | 0.196 | 0.435 | 1.12E-06 | 03_Mito-DC |

| | | | | | |
|---|---|---|---|---|---|
| DUSP4 | -1.16601128 | 0.34 | 0.575 | 1.36E-06 | 03_Mito-DC |
| TOMM22 | -1.90138305 | 0.086 | 0.307 | 1.69E-06 | 03_Mito-DC |
| AL133415.1 | -1.94289584 | 0.196 | 0.408 | 1.72E-06 | 03_Mito-DC |
| PARK7 | -1.03027596 | 0.301 | 0.553 | 1.88E-06 | 03_Mito-DC |
| SNHG15 | -1.2208836 | 0.273 | 0.507 | 2.88E-06 | 03_Mito-DC |
| ERGIC3 | -1.31483531 | 0.153 | 0.391 | 3.29E-06 | 03_Mito-DC |
| ALKBH7 | -1.41517492 | 0.196 | 0.426 | 3.96E-06 | 03_Mito-DC |
| SNX3 | -1.28538289 | 0.397 | 0.628 | 3.99E-06 | 03_Mito-DC |
| BIRC3 | -2.17031092 | 0.201 | 0.415 | 4.07E-06 | 03_Mito-DC |
| APRT | -1.03181865 | 0.335 | 0.567 | 5.67E-06 | 03_Mito-DC |
| OSTF1 | -1.0934862 | 0.215 | 0.45 | 6.19E-06 | 03_Mito-DC |
| AC009093.2 | -1.59478025 | 0.129 | 0.349 | 6.41E-06 | 03_Mito-DC |
| ATP5PB | -1.11053293 | 0.258 | 0.482 | 7.60E-06 | 03_Mito-DC |
| TMEM219 | -1.17023649 | 0.278 | 0.501 | 7.78E-06 | 03_Mito-DC |
| APH1A | -1.37832662 | 0.215 | 0.433 | 8.39E-06 | 03_Mito-DC |
| CAT | -1.57414801 | 0.129 | 0.352 | 9.08E-06 | 03_Mito-DC |
| TUBA1C | -1.22457499 | 0.206 | 0.44 | 9.51E-06 | 03_Mito-DC |
| RTRAF | -1.26600083 | 0.187 | 0.414 | 1.01E-05 | 03_Mito-DC |
| AURKAIP1 | -1.33329455 | 0.177 | 0.401 | 1.23E-05 | 03_Mito-DC |
| RCSD1 | -1.12467767 | 0.182 | 0.418 | 1.58E-05 | 03_Mito-DC |
| GABARAPL2 | -1.05958883 | 0.349 | 0.576 | 1.78E-05 | 03_Mito-DC |
| FRY | -2.43783697 | 0.086 | 0.287 | 1.82E-05 | 03_Mito-DC |
| FKBP2 | -1.75562633 | 0.091 | 0.292 | 2.41E-05 | 03_Mito-DC |
| NDUFB9 | -1.15257507 | 0.172 | 0.399 | 2.43E-05 | 03_Mito-DC |
| AUTS2 | -1.14869075 | 0.172 | 0.407 | 2.64E-05 | 03_Mito-DC |
| MDH2 | -1.26095806 | 0.187 | 0.404 | 3.14E-05 | 03_Mito-DC |
| CRTAP | -1.11192116 | 0.23 | 0.448 | 3.55E-05 | 03_Mito-DC |
| SF3B5 | -1.27326658 | 0.196 | 0.398 | 3.83E-05 | 03_Mito-DC |
| EIF3M | -1.00821382 | 0.23 | 0.458 | 5.25E-05 | 03_Mito-DC |
| PGAM1 | -1.17040871 | 0.239 | 0.457 | 5.45E-05 | 03_Mito-DC |
| IRF4 | -1.68180051 | 0.124 | 0.333 | 6.48E-05 | 03_Mito-DC |
| OPN3 | -1.55195195 | 0.124 | 0.326 | 6.98E-05 | 03_Mito-DC |
| ATP5F1A | -1.26972938 | 0.182 | 0.385 | 8.96E-05 | 03_Mito-DC |
| DBNL | -1.15221976 | 0.211 | 0.418 | 0.0001199 | 03_Mito-DC |
| PSMA6 | -1.03101451 | 0.301 | 0.516 | 0.0001221 | 03_Mito-DC |
| FOXN2 | -1.02434692 | 0.244 | 0.454 | 0.0001759 | 03_Mito-DC |
| ADGRE5 | -1.17541153 | 0.258 | 0.467 | 0.0002706 | 03_Mito-DC |
| FLNA | -1.0317715 | 0.22 | 0.438 | 0.0002905 | 03_Mito-DC |
| C4orf3 | -1.14559788 | 0.239 | 0.442 | 0.0003385 | 03_Mito-DC |
| EIF3I | -1.00285943 | 0.201 | 0.415 | 0.0003628 | 03_Mito-DC |
| ARL4C | -1.14125553 | 0.297 | 0.507 | 0.0004732 | 03_Mito-DC |
| SSR3 | -1.01344088 | 0.287 | 0.493 | 0.0005145 | 03_Mito-DC |
| LAMTOR1 | -1.1640912 | 0.249 | 0.46 | 0.0005249 | 03_Mito-DC |
| JTB | -1.06328893 | 0.23 | 0.436 | 0.0005607 | 03_Mito-DC |
| OSTC | -1.09978627 | 0.182 | 0.384 | 0.0009867 | 03_Mito-DC |
| ARL4C | 1.812192168 | 0.852 | 0.444 | 1.06E-25 | 04-cDC2 |

| | | | | | |
|---|---|---|---|---|---|
| CD1C | 1.855094581 | 0.734 | 0.301 | 2.73E-22 | 04-cDC2 |
| FCER1A | 1.42311707 | 0.82 | 0.439 | 4.81E-17 | 04-cDC2 |
| METRNL | 1.193912332 | 0.875 | 0.61 | 1.02E-16 | 04-cDC2 |
| C3 | -3.71032282 | 0.141 | 0.552 | 2.81E-15 | 04-cDC2 |
| CD1E | 1.877237351 | 0.5 | 0.187 | 3.04E-15 | 04-cDC2 |
| IFITM2 | 1.018819443 | 0.961 | 0.724 | 8.67E-14 | 04-cDC2 |
| CAVIN2 | 2.559275751 | 0.297 | 0.076 | 4.99E-13 | 04-cDC2 |
| INSIG1 | 1.233033697 | 0.836 | 0.559 | 8.33E-13 | 04-cDC2 |
| CLEC10A | 1.294812519 | 0.688 | 0.338 | 3.53E-12 | 04-cDC2 |
| ZFP36L1 | -1.64628817 | 0.461 | 0.763 | 4.64E-12 | 04-cDC2 |
| C1QA | -3.57501179 | 0.188 | 0.535 | 2.22E-11 | 04-cDC2 |
| SLC1A3 | -3.32749311 | 0.203 | 0.556 | 2.61E-11 | 04-cDC2 |
| C1QC | -3.99466679 | 0.266 | 0.572 | 1.09E-10 | 04-cDC2 |
| SLC11A1 | -2.97427733 | 0.133 | 0.484 | 1.64E-10 | 04-cDC2 |
| TREM2 | -3.95441103 | 0.07 | 0.413 | 3.97E-10 | 04-cDC2 |
| C1QB | -3.19961321 | 0.266 | 0.57 | 5.65E-10 | 04-cDC2 |
| APOE | -4.48598258 | 0.328 | 0.613 | 1.51E-09 | 04-cDC2 |
| RNASE6 | 1.008602919 | 0.773 | 0.548 | 3.76E-09 | 04-cDC2 |
| SPP1 | -3.49258535 | 0.359 | 0.646 | 9.66E-09 | 04-cDC2 |
| DLEU1 | -3.44232577 | 0.078 | 0.401 | 1.52E-08 | 04-cDC2 |
| CKLF | 1.005863996 | 0.812 | 0.605 | 1.74E-08 | 04-cDC2 |
| APOC1 | -3.58656414 | 0.234 | 0.525 | 8.30E-08 | 04-cDC2 |
| AKR1C3 | 1.662669094 | 0.359 | 0.14 | 1.97E-07 | 04-cDC2 |
| FRMD4A | -2.97475933 | 0.164 | 0.468 | 2.01E-07 | 04-cDC2 |
| FMNL2 | -1.98777693 | 0.273 | 0.592 | 2.54E-07 | 04-cDC2 |
| RPS4Y1 | 1.002937414 | 0.719 | 0.478 | 3.04E-07 | 04-cDC2 |
| NRARP | 1.286670874 | 0.562 | 0.304 | 5.42E-07 | 04-cDC2 |
| SORL1 | -1.95713981 | 0.172 | 0.49 | 5.43E-07 | 04-cDC2 |
| USP53 | -2.29743991 | 0.125 | 0.43 | 1.61E-06 | 04-cDC2 |
| TCF12 | -1.85662223 | 0.258 | 0.565 | 3.04E-06 | 04-cDC2 |
| LHFPL2 | -5.04590456 | 0.023 | 0.283 | 3.73E-06 | 04-cDC2 |
| GAREM1 | 1.074290052 | 0.648 | 0.392 | 5.15E-06 | 04-cDC2 |
| IFITM1 | 1.090725823 | 0.641 | 0.385 | 5.20E-06 | 04-cDC2 |
| SOD2 | -1.28405962 | 0.469 | 0.705 | 7.17E-06 | 04-cDC2 |
| OLR1 | -1.47876599 | 0.477 | 0.681 | 8.83E-06 | 04-cDC2 |
| SATB1 | 1.047888769 | 0.617 | 0.364 | 1.63E-05 | 04-cDC2 |
| SPIB | 1.328132516 | 0.516 | 0.309 | 3.35E-05 | 04-cDC2 |
| JAML | 1.11495139 | 0.555 | 0.349 | 3.81E-05 | 04-cDC2 |
| TXNIP | -1.34312042 | 0.305 | 0.574 | 4.54E-05 | 04-cDC2 |
| FCGR3A | -2.80571616 | 0.109 | 0.363 | 5.11E-05 | 04-cDC2 |
| MSR1 | -1.45602392 | 0.273 | 0.56 | 5.80E-05 | 04-cDC2 |
| KCNQ3 | -6.06021397 | 0.016 | 0.246 | 6.05E-05 | 04-cDC2 |
| GPR34 | -3.21909927 | 0.047 | 0.278 | 0.0002579 | 04-cDC2 |
| MAFB | -2.18395395 | 0.117 | 0.375 | 0.0002785 | 04-cDC2 |
| APBB1IP | -1.66565149 | 0.281 | 0.52 | 0.0003299 | 04-cDC2 |
| MAML2 | -2.16162507 | 0.156 | 0.413 | 0.0004031 | 04-cDC2 |

| | | | | | |
|---|---|---|---|---|---|
| EBI3 | -2.22090508 | 0.094 | 0.34 | 0.0007319 | 04-cDC2 |
| CAMK2D | -3.8986475 | 0.031 | 0.244 | 0.0009592 | 04-cDC2 |
| ARID5B | -1.55328457 | 0.227 | 0.488 | 0.0011778 | 04-cDC2 |
| TNFAIP2 | -2.61571121 | 0.078 | 0.302 | 0.0016144 | 04-cDC2 |
| ADAM17 | -1.49971088 | 0.203 | 0.457 | 0.002684 | 04-cDC2 |
| CSGALNACT1 | -2.35522789 | 0.141 | 0.367 | 0.002863 | 04-cDC2 |
| A2M | -1.62170602 | 0.219 | 0.468 | 0.0035838 | 04-cDC2 |
| C5AR1 | -2.20433592 | 0.188 | 0.395 | 0.0052003 | 04-cDC2 |
| PLA2G4A | -2.26972968 | 0.133 | 0.357 | 0.0054249 | 04-cDC2 |
| MARCKS | -1.43751907 | 0.344 | 0.556 | 0.0054557 | 04-cDC2 |
| PTPRJ | -1.87272602 | 0.18 | 0.413 | 0.0063145 | 04-cDC2 |
| RASAL2 | -1.4148425 | 0.102 | 0.33 | 0.0075023 | 04-cDC2 |
| HMOX1 | -1.81013586 | 0.234 | 0.449 | 0.0084628 | 04-cDC2 |
| DLEU2 | -1.63402769 | 0.281 | 0.506 | 0.0088086 | 04-cDC2 |
| LRCH1 | -2.23832799 | 0.062 | 0.268 | 0.0097175 | 04-cDC2 |
| GALNT2 | -1.91173318 | 0.109 | 0.325 | 0.0100009 | 04-cDC2 |
| LRRK1 | -2.53748725 | 0.07 | 0.276 | 0.0102169 | 04-cDC2 |
| CD163 | -1.42152743 | 0.312 | 0.512 | 0.011905 | 04-cDC2 |
| ZBTB16 | -1.27361696 | 0.273 | 0.509 | 0.0125293 | 04-cDC2 |
| SFMBT2 | -1.98535008 | 0.258 | 0.469 | 0.0143636 | 04-cDC2 |
| ADAP2 | -1.46784006 | 0.148 | 0.372 | 0.016489 | 04-cDC2 |
| SDCCAG8 | -1.48917793 | 0.211 | 0.444 | 0.0167257 | 04-cDC2 |
| ALCAM | -1.13452575 | 0.453 | 0.675 | 0.018777 | 04-cDC2 |
| FNDC3A | -1.82081977 | 0.156 | 0.37 | 0.0229948 | 04-cDC2 |
| CYFIP1 | -1.78515698 | 0.172 | 0.385 | 0.0278356 | 04-cDC2 |
| CD69 | -1.39313102 | 0.242 | 0.455 | 0.0549196 | 04-cDC2 |
| TOM1 | -1.07741144 | 0.227 | 0.47 | 0.0570629 | 04-cDC2 |
| PDE8A | -1.58900828 | 0.242 | 0.453 | 0.0747191 | 04-cDC2 |
| EPB41L2 | -1.58330728 | 0.281 | 0.487 | 0.1106693 | 04-cDC2 |
| ZFHX3 | -1.29320511 | 0.203 | 0.42 | 0.1449303 | 04-cDC2 |
| NUMB | -1.24277295 | 0.227 | 0.449 | 0.2136572 | 04-cDC2 |
| VASH1 | -1.04271636 | 0.359 | 0.567 | 0.3491927 | 04-cDC2 |
| SRGAP1 | -1.06233334 | 0.383 | 0.585 | 0.3778808 | 04-cDC2 |
| IRAK2 | -1.08905269 | 0.242 | 0.451 | 0.4759071 | 04-cDC2 |
| FCGBP | 3.489328444 | 0.702 | 0.117 | 4.08E-57 | 05_mo-DC1 |
| HLA-DRB5 | -3.98090794 | 0.567 | 0.957 | 1.52E-47 | 05_mo-DC1 |
| RGS16 | 3.978341757 | 0.558 | 0.081 | 3.16E-47 | 05_mo-DC1 |
| SPP1 | 3.112769919 | 0.99 | 0.595 | 1.75E-46 | 05_mo-DC1 |
| CTSD | 2.764585868 | 0.933 | 0.411 | 7.35E-44 | 05_mo-DC1 |
| HLA-DQA1 | -3.09603342 | 0.712 | 0.972 | 1.08E-43 | 05_mo-DC1 |
| TREM2 | 2.497357594 | 0.913 | 0.345 | 2.97E-42 | 05_mo-DC1 |
| ADGRG1 | 2.87548983 | 0.567 | 0.094 | 3.19E-42 | 05_mo-DC1 |
| SLC2A5 | 2.504369488 | 0.625 | 0.111 | 2.25E-41 | 05_mo-DC1 |
| CYTL1 | 3.929115635 | 0.327 | 0.026 | 1.14E-40 | 05_mo-DC1 |
| RAB42 | 4.668293199 | 0.269 | 0.016 | 2.05E-40 | 05_mo-DC1 |
| APOE | 2.315093812 | 0.981 | 0.56 | 3.62E-35 | 05_mo-DC1 |

| | | | | | |
|---|---|---|---|---|---|
| BNIP3 | 4.048301922 | 0.269 | 0.02 | 1.91E-34 | 05_mo-DC1 |
| TMIGD3 | 3.153018102 | 0.558 | 0.118 | 3.39E-34 | 05_mo-DC1 |
| C1QC | 2.009958307 | 0.962 | 0.516 | 9.27E-34 | 05_mo-DC1 |
| HTRA1 | 2.38644496 | 0.625 | 0.146 | 6.90E-33 | 05_mo-DC1 |
| GPNMB | 3.579119673 | 0.433 | 0.072 | 2.70E-31 | 05_mo-DC1 |
| LINC01736 | 2.986754173 | 0.452 | 0.075 | 4.31E-31 | 05_mo-DC1 |
| TMEM119 | 3.456712375 | 0.308 | 0.033 | 1.88E-30 | 05_mo-DC1 |
| APOC1 | 1.795332918 | 0.933 | 0.468 | 8.50E-30 | 05_mo-DC1 |
| C1QB | 1.653501634 | 0.962 | 0.513 | 2.98E-29 | 05_mo-DC1 |
| NPL | 2.294411787 | 0.49 | 0.092 | 8.25E-29 | 05_mo-DC1 |
| APOO | 1.955839676 | 0.788 | 0.298 | 8.83E-29 | 05_mo-DC1 |
| GYPC | 2.249917891 | 0.76 | 0.289 | 1.33E-28 | 05_mo-DC1 |
| TAL1 | 2.960565513 | 0.298 | 0.032 | 2.57E-28 | 05_mo-DC1 |
| C5AR1 | 1.734256152 | 0.856 | 0.343 | 1.71E-27 | 05_mo-DC1 |
| C1QA | 1.577749421 | 0.942 | 0.474 | 4.09E-27 | 05_mo-DC1 |
| FSCN1 | 2.330324262 | 0.587 | 0.159 | 1.02E-26 | 05_mo-DC1 |
| C12orf75 | 2.378289255 | 0.49 | 0.106 | 8.92E-26 | 05_mo-DC1 |
| AC105402.3 | 2.301848936 | 0.481 | 0.103 | 2.17E-25 | 05_mo-DC1 |
| SLC29A1 | 2.66017942 | 0.481 | 0.111 | 5.09E-24 | 05_mo-DC1 |
| A2M | 1.810203538 | 0.827 | 0.418 | 1.49E-23 | 05_mo-DC1 |
| RHOB | 1.711635025 | 0.933 | 0.584 | 3.35E-23 | 05_mo-DC1 |
| PLA2G7 | 3.046481426 | 0.221 | 0.021 | 8.60E-23 | 05_mo-DC1 |
| CD276 | 2.910040289 | 0.317 | 0.048 | 1.66E-22 | 05_mo-DC1 |
| SLC11A1 | 1.327197731 | 0.923 | 0.42 | 2.55E-22 | 05_mo-DC1 |
| GAL3ST4 | 2.919893016 | 0.26 | 0.033 | 5.20E-21 | 05_mo-DC1 |
| MCRIP2 | 2.232466246 | 0.356 | 0.066 | 5.83E-20 | 05_mo-DC1 |
| PCED1B-AS1 | 1.7239587 | 0.644 | 0.216 | 1.05E-19 | 05_mo-DC1 |
| CD68 | 1.542390708 | 0.875 | 0.544 | 1.08E-19 | 05_mo-DC1 |
| CD14 | 1.924556265 | 0.798 | 0.414 | 1.51E-19 | 05_mo-DC1 |
| SLCO2B1 | 1.527373829 | 0.625 | 0.202 | 1.08E-18 | 05_mo-DC1 |
| XIST | 1.630079872 | 0.49 | 0.126 | 3.02E-18 | 05_mo-DC1 |
| MERTK | 1.372184941 | 0.567 | 0.166 | 5.14E-18 | 05_mo-DC1 |
| LINC01094 | 2.581993833 | 0.346 | 0.072 | 2.46E-17 | 05_mo-DC1 |
| CAPG | 1.395062641 | 0.885 | 0.646 | 2.58E-17 | 05_mo-DC1 |
| APOC2 | 2.459629288 | 0.558 | 0.19 | 5.34E-17 | 05_mo-DC1 |
| SNHG12 | 2.85538399 | 0.558 | 0.205 | 1.56E-16 | 05_mo-DC1 |
| SLC35F1 | 2.673579791 | 0.26 | 0.041 | 1.89E-16 | 05_mo-DC1 |
| STAB1 | 1.637756441 | 0.5 | 0.141 | 2.73E-16 | 05_mo-DC1 |
| OLFML3 | 1.688960581 | 0.548 | 0.17 | 3.71E-16 | 05_mo-DC1 |
| FCER1A | -7.27479667 | 0.01 | 0.506 | 7.53E-16 | 05_mo-DC1 |
| LILRB4 | 1.350273114 | 0.788 | 0.41 | 1.21E-15 | 05_mo-DC1 |
| LIPA | 2.038619871 | 0.577 | 0.217 | 3.19E-15 | 05_mo-DC1 |
| TSPO | 1.518350643 | 0.837 | 0.587 | 8.39E-15 | 05_mo-DC1 |
| GLRX | 1.538406142 | 0.663 | 0.28 | 1.87E-14 | 05_mo-DC1 |
| HMOX1 | 1.968008794 | 0.75 | 0.408 | 2.32E-14 | 05_mo-DC1 |
| AREG | -1.85793678 | 0.558 | 0.852 | 2.87E-14 | 05_mo-DC1 |

| | | | | | |
|---|---|---|---|---|---|
| TPI1 | 1.517141658 | 0.894 | 0.67 | 3.86E-14 | 05_mo-DC1 |
| SORT1 | 1.230123221 | 0.577 | 0.194 | 8.00E-14 | 05_mo-DC1 |
| CTSL | 1.665288512 | 0.587 | 0.217 | 1.75E-13 | 05_mo-DC1 |
| EIF4EBP1 | 1.971974172 | 0.519 | 0.188 | 2.07E-13 | 05_mo-DC1 |
| P4HB | 1.708001279 | 0.74 | 0.384 | 5.12E-13 | 05_mo-DC1 |
| GATM | 2.296460707 | 0.365 | 0.097 | 5.16E-13 | 05_mo-DC1 |
| SLC7A7 | 1.659633989 | 0.462 | 0.147 | 8.23E-13 | 05_mo-DC1 |
| CRYBG1 | -2.96449352 | 0.212 | 0.611 | 1.13E-12 | 05_mo-DC1 |
| RPS4Y1 | -2.8940896 | 0.067 | 0.53 | 2.51E-12 | 05_mo-DC1 |
| IL1R2 | -4.66446149 | 0.087 | 0.504 | 3.57E-12 | 05_mo-DC1 |
| ISCU | 1.747604872 | 0.577 | 0.268 | 6.46E-12 | 05_mo-DC1 |
| GLUL | 1.060300957 | 0.923 | 0.714 | 1.08E-11 | 05_mo-DC1 |
| RABAC1 | 1.419104546 | 0.692 | 0.354 | 1.43E-11 | 05_mo-DC1 |
| VAT1 | 2.295471079 | 0.25 | 0.05 | 1.99E-11 | 05_mo-DC1 |
| CRIP1 | -5.09874355 | 0.058 | 0.465 | 2.50E-11 | 05_mo-DC1 |
| SUSD3 | 1.757615107 | 0.404 | 0.122 | 2.66E-11 | 05_mo-DC1 |
| FCGR3A | 1.120707455 | 0.692 | 0.316 | 4.20E-11 | 05_mo-DC1 |
| PYCARD | 1.2361655 | 0.76 | 0.436 | 5.93E-11 | 05_mo-DC1 |
| CYFIP1 | 1.159578647 | 0.712 | 0.341 | 7.31E-11 | 05_mo-DC1 |
| SCIN | 1.602959432 | 0.394 | 0.12 | 7.68E-11 | 05_mo-DC1 |
| SLC25A37 | 1.489887644 | 0.702 | 0.348 | 7.80E-11 | 05_mo-DC1 |
| AFF3 | -3.87496538 | 0.058 | 0.47 | 1.11E-10 | 05_mo-DC1 |
| PDPN | 1.974563762 | 0.308 | 0.077 | 2.15E-10 | 05_mo-DC1 |
| STK4 | -1.53905687 | 0.538 | 0.796 | 2.68E-10 | 05_mo-DC1 |
| ASAH1 | 1.0384116 | 0.846 | 0.557 | 3.55E-10 | 05_mo-DC1 |
| EVL | 1.22374084 | 0.644 | 0.304 | 3.58E-10 | 05_mo-DC1 |
| IFITM1 | -4.79991916 | 0.048 | 0.433 | 5.50E-10 | 05_mo-DC1 |
| ALDOA | 1.559888228 | 0.817 | 0.597 | 7.23E-10 | 05_mo-DC1 |
| PLD3 | 1.727187424 | 0.529 | 0.225 | 7.48E-10 | 05_mo-DC1 |
| CCSER1 | -3.76143907 | 0.096 | 0.489 | 7.67E-10 | 05_mo-DC1 |
| MPP1 | 1.295074777 | 0.548 | 0.228 | 8.73E-10 | 05_mo-DC1 |
| SCARB2 | 1.406555525 | 0.442 | 0.153 | 9.24E-10 | 05_mo-DC1 |
| BSG | 1.17920071 | 0.692 | 0.367 | 1.08E-09 | 05_mo-DC1 |
| BIN1 | 1.858183137 | 0.74 | 0.528 | 1.18E-09 | 05_mo-DC1 |
| GLDN | 1.93553443 | 0.385 | 0.13 | 1.95E-09 | 05_mo-DC1 |
| CD99 | 1.20450054 | 0.76 | 0.504 | 2.08E-09 | 05_mo-DC1 |
| CALM3 | 1.514320216 | 0.519 | 0.209 | 2.36E-09 | 05_mo-DC1 |
| ATP6V1F | 1.027727745 | 0.837 | 0.604 | 2.65E-09 | 05_mo-DC1 |
| NFIC | 1.23484857 | 0.577 | 0.258 | 2.88E-09 | 05_mo-DC1 |
| CHCHD10 | 1.22217135 | 0.75 | 0.465 | 3.22E-09 | 05_mo-DC1 |
| LGALS9 | 1.309625828 | 0.712 | 0.428 | 4.20E-09 | 05_mo-DC1 |
| NUDT5 | 1.657113081 | 0.385 | 0.122 | 6.94E-09 | 05_mo-DC1 |
| DNASE2 | 1.475322745 | 0.442 | 0.158 | 7.06E-09 | 05_mo-DC1 |
| TCEAL3 | 1.675397045 | 0.404 | 0.14 | 7.12E-09 | 05_mo-DC1 |
| MEA1 | 1.548323408 | 0.471 | 0.189 | 7.32E-09 | 05_mo-DC1 |
| IGSF21 | 1.331546643 | 0.327 | 0.09 | 7.58E-09 | 05_mo-DC1 |

| | | | | | |
|---|---|---|---|---|---|
| GPR34 | 1.128335461 | 0.567 | 0.236 | 8.63E-09 | 05_mo-DC1 |
| FAM110B | 1.332907616 | 0.433 | 0.146 | 9.73E-09 | 05_mo-DC1 |
| MMP14 | 1.737425583 | 0.346 | 0.104 | 1.04E-08 | 05_mo-DC1 |
| PYGL | 1.235642955 | 0.337 | 0.097 | 1.28E-08 | 05_mo-DC1 |
| CLEC10A | -4.23342638 | 0.029 | 0.392 | 1.54E-08 | 05_mo-DC1 |
| CCND1 | 1.459686072 | 0.394 | 0.135 | 1.66E-08 | 05_mo-DC1 |
| EEPD1 | 1.45389281 | 0.308 | 0.084 | 1.72E-08 | 05_mo-DC1 |
| LHFPL2 | 1.11360846 | 0.558 | 0.239 | 1.84E-08 | 05_mo-DC1 |
| S100A6 | -1.23142835 | 0.452 | 0.741 | 1.94E-08 | 05_mo-DC1 |
| RNASE1 | 2.87838389 | 0.356 | 0.12 | 2.03E-08 | 05_mo-DC1 |
| CNN2 | -2.52452427 | 0.106 | 0.479 | 2.36E-08 | 05_mo-DC1 |
| DUSP5 | -3.69519786 | 0.115 | 0.46 | 2.85E-08 | 05_mo-DC1 |
| TOLLIP | 1.496740889 | 0.365 | 0.116 | 2.99E-08 | 05_mo-DC1 |
| CCL4L2 | -2.79999538 | 0.135 | 0.513 | 3.88E-08 | 05_mo-DC1 |
| CREG1 | 1.207390884 | 0.625 | 0.315 | 4.16E-08 | 05_mo-DC1 |
| FCGR1A | 1.200160672 | 0.606 | 0.29 | 4.56E-08 | 05_mo-DC1 |
| ARL6IP4 | 1.104567043 | 0.76 | 0.51 | 4.90E-08 | 05_mo-DC1 |
| LPAR5 | 1.580435897 | 0.308 | 0.088 | 5.02E-08 | 05_mo-DC1 |
| ADAP2 | 1.139601537 | 0.663 | 0.33 | 5.41E-08 | 05_mo-DC1 |
| SH3BP1 | 1.487824954 | 0.442 | 0.174 | 6.90E-08 | 05_mo-DC1 |
| MYDGF | 1.2866211 | 0.644 | 0.354 | 7.06E-08 | 05_mo-DC1 |
| INSIG1 | -2.46177375 | 0.317 | 0.602 | 7.17E-08 | 05_mo-DC1 |
| RNASEH2C | 1.458570543 | 0.413 | 0.146 | 7.17E-08 | 05_mo-DC1 |
| PLIN2 | 1.684646904 | 0.683 | 0.409 | 9.50E-08 | 05_mo-DC1 |
| RAB13 | 1.446008819 | 0.365 | 0.122 | 1.27E-07 | 05_mo-DC1 |
| INO80D | -2.72923353 | 0.106 | 0.458 | 1.38E-07 | 05_mo-DC1 |
| STX10 | 1.436828488 | 0.385 | 0.134 | 1.39E-07 | 05_mo-DC1 |
| CREBL2 | 1.426719778 | 0.365 | 0.122 | 1.51E-07 | 05_mo-DC1 |
| ABCA1 | 1.518321446 | 0.663 | 0.327 | 1.58E-07 | 05_mo-DC1 |
| CDK2AP1 | -2.37710635 | 0.269 | 0.559 | 1.80E-07 | 05_mo-DC1 |
| CST7 | -2.97353941 | 0.202 | 0.509 | 2.52E-07 | 05_mo-DC1 |
| IL17RA | 1.237647866 | 0.615 | 0.316 | 2.88E-07 | 05_mo-DC1 |
| LYZ | -1.41717148 | 0.519 | 0.723 | 3.60E-07 | 05_mo-DC1 |
| CNPY3 | 1.024301826 | 0.74 | 0.49 | 4.03E-07 | 05_mo-DC1 |
| CD1C | -4.95907533 | 0.038 | 0.36 | 4.69E-07 | 05_mo-DC1 |
| DHRS7 | 1.095971145 | 0.673 | 0.396 | 5.03E-07 | 05_mo-DC1 |
| LRP1 | 1.322329648 | 0.452 | 0.183 | 5.28E-07 | 05_mo-DC1 |
| PRDX1 | 1.064717611 | 0.75 | 0.526 | 5.55E-07 | 05_mo-DC1 |
| MRC2 | 1.472643561 | 0.346 | 0.116 | 7.12E-07 | 05_mo-DC1 |
| IFITM10 | 1.463152811 | 0.317 | 0.099 | 7.91E-07 | 05_mo-DC1 |
| CIITA | -1.80345383 | 0.365 | 0.637 | 8.62E-07 | 05_mo-DC1 |
| FAM13A | 1.785414372 | 0.356 | 0.121 | 8.70E-07 | 05_mo-DC1 |
| AP1B1 | 1.051910057 | 0.692 | 0.386 | 8.93E-07 | 05_mo-DC1 |
| SELPLG | 1.417787345 | 0.462 | 0.198 | 1.19E-06 | 05_mo-DC1 |
| FKBP8 | 1.078808078 | 0.702 | 0.407 | 1.32E-06 | 05_mo-DC1 |
| OSM | 1.558206542 | 0.481 | 0.213 | 1.34E-06 | 05_mo-DC1 |

| | | | | | |
|---|---|---|---|---|---|
| ARF6 | -1.84248504 | 0.337 | 0.616 | 1.70E-06 | 05_mo-DC1 |
| SELENOW | 1.149785789 | 0.635 | 0.343 | 1.80E-06 | 05_mo-DC1 |
| AGO2 | -2.12078491 | 0.183 | 0.506 | 1.87E-06 | 05_mo-DC1 |
| HM13 | 1.295863601 | 0.654 | 0.394 | 1.88E-06 | 05_mo-DC1 |
| TNFAIP8 | -3.26314152 | 0.154 | 0.452 | 1.96E-06 | 05_mo-DC1 |
| DDX3Y | -2.82664762 | 0.048 | 0.364 | 2.02E-06 | 05_mo-DC1 |
| CCDC85B | 1.245712134 | 0.529 | 0.247 | 2.52E-06 | 05_mo-DC1 |
| FEZ2 | 1.145247011 | 0.471 | 0.205 | 2.65E-06 | 05_mo-DC1 |
| ADAM19 | -7.55009297 | 0.01 | 0.309 | 2.87E-06 | 05_mo-DC1 |
| MRPL40 | 1.40303146 | 0.327 | 0.109 | 3.21E-06 | 05_mo-DC1 |
| CDK14 | -2.36016549 | 0.096 | 0.418 | 3.43E-06 | 05_mo-DC1 |
| MRPL12 | 1.405177302 | 0.346 | 0.12 | 3.71E-06 | 05_mo-DC1 |
| PDE4DIP | 1.269232701 | 0.615 | 0.319 | 4.01E-06 | 05_mo-DC1 |
| ETV5 | 1.307124484 | 0.433 | 0.173 | 4.01E-06 | 05_mo-DC1 |
| CDCP1 | 1.273434893 | 0.356 | 0.126 | 4.88E-06 | 05_mo-DC1 |
| LINC01578 | -1.71204695 | 0.317 | 0.596 | 5.13E-06 | 05_mo-DC1 |
| FYN | -3.39814443 | 0.077 | 0.384 | 5.66E-06 | 05_mo-DC1 |
| TRAF1 | -4.01821836 | 0.048 | 0.351 | 5.94E-06 | 05_mo-DC1 |
| HEBP1 | 1.535432488 | 0.346 | 0.124 | 6.40E-06 | 05_mo-DC1 |
| RGS19 | 1.257882805 | 0.567 | 0.288 | 6.48E-06 | 05_mo-DC1 |
| VSIG4 | 1.392231216 | 0.615 | 0.374 | 7.50E-06 | 05_mo-DC1 |
| AHR | -1.91277449 | 0.173 | 0.5 | 9.22E-06 | 05_mo-DC1 |
| SLC25A39 | 1.305746177 | 0.519 | 0.245 | 1.02E-05 | 05_mo-DC1 |
| RFTN1 | -2.05062463 | 0.25 | 0.547 | 1.04E-05 | 05_mo-DC1 |
| IL1R1 | -5.51337663 | 0.029 | 0.314 | 1.15E-05 | 05_mo-DC1 |
| DAB2 | 1.394595038 | 0.356 | 0.126 | 1.28E-05 | 05_mo-DC1 |
| CBWD5 | 1.539661724 | 0.308 | 0.106 | 1.34E-05 | 05_mo-DC1 |
| CTSA | 1.312831013 | 0.5 | 0.232 | 1.37E-05 | 05_mo-DC1 |
| FLT3 | -4.40975257 | 0.019 | 0.309 | 1.77E-05 | 05_mo-DC1 |
| SPATA13 | 1.024922618 | 0.423 | 0.172 | 1.78E-05 | 05_mo-DC1 |
| ARHGAP35 | 1.388363281 | 0.298 | 0.097 | 2.34E-05 | 05_mo-DC1 |
| LGMN | 1.033465 | 0.615 | 0.35 | 2.55E-05 | 05_mo-DC1 |
| IFNLR1 | 1.125616554 | 0.481 | 0.217 | 2.63E-05 | 05_mo-DC1 |
| DAPP1 | -4.70228201 | 0.029 | 0.31 | 2.80E-05 | 05_mo-DC1 |
| CISD3 | 1.548320014 | 0.327 | 0.121 | 3.17E-05 | 05_mo-DC1 |
| THBS1 | -3.70761019 | 0.096 | 0.384 | 3.35E-05 | 05_mo-DC1 |
| MKNK1 | 1.027175289 | 0.433 | 0.184 | 3.37E-05 | 05_mo-DC1 |
| BIRC3 | -3.27678797 | 0.115 | 0.406 | 3.65E-05 | 05_mo-DC1 |
| NEK6 | 1.420145932 | 0.375 | 0.151 | 4.04E-05 | 05_mo-DC1 |
| EPN1 | 1.005408389 | 0.548 | 0.275 | 4.12E-05 | 05_mo-DC1 |
| LMAN1 | 1.416996273 | 0.394 | 0.166 | 4.65E-05 | 05_mo-DC1 |
| LPCAT1 | 1.373214412 | 0.308 | 0.106 | 5.12E-05 | 05_mo-DC1 |
| H1F0 | 1.193459159 | 0.337 | 0.124 | 5.14E-05 | 05_mo-DC1 |
| IER5 | -2.40011271 | 0.173 | 0.455 | 5.31E-05 | 05_mo-DC1 |
| PLTP | 1.095617255 | 0.385 | 0.157 | 6.16E-05 | 05_mo-DC1 |
| NAXE | 1.336810543 | 0.327 | 0.118 | 6.33E-05 | 05_mo-DC1 |

| | | | | | |
|---|---|---|---|---|---|
| TBC1D9B | 1.195377288 | 0.404 | 0.167 | 6.77E-05 | 05_mo-DC1 |
| VKORC1 | 1.326726595 | 0.51 | 0.26 | 7.77E-05 | 05_mo-DC1 |
| CD44 | -1.4914555 | 0.5 | 0.705 | 8.32E-05 | 05_mo-DC1 |
| MT2A | -2.5840086 | 0.462 | 0.677 | 8.46E-05 | 05_mo-DC1 |
| MCOLN2 | -2.70833852 | 0.106 | 0.384 | 9.02E-05 | 05_mo-DC1 |
| PSTPIP2 | -2.01815936 | 0.125 | 0.424 | 0.0001094 | 05_mo-DC1 |
| TGFBI | 1.544219806 | 0.596 | 0.359 | 0.0001142 | 05_mo-DC1 |
| PRAM1 | 1.009000809 | 0.433 | 0.184 | 0.0001183 | 05_mo-DC1 |
| BIN2 | 1.143080076 | 0.365 | 0.147 | 0.0001193 | 05_mo-DC1 |
| PTMS | 1.049436083 | 0.644 | 0.381 | 0.000125 | 05_mo-DC1 |
| JAML | -2.20580578 | 0.096 | 0.387 | 0.0001504 | 05_mo-DC1 |
| LILRB1 | 1.034415519 | 0.433 | 0.196 | 0.0001686 | 05_mo-DC1 |
| SCPEP1 | 1.091854409 | 0.529 | 0.274 | 0.0001765 | 05_mo-DC1 |
| DYNLRB1 | 1.131215027 | 0.481 | 0.225 | 0.00018 | 05_mo-DC1 |
| GPR157 | -4.5136258 | 0.019 | 0.281 | 0.0001888 | 05_mo-DC1 |
| GLS | -1.49044986 | 0.433 | 0.648 | 0.0002078 | 05_mo-DC1 |
| ISG20 | -1.69597921 | 0.279 | 0.54 | 0.0002109 | 05_mo-DC1 |
| TAF10 | -1.63867451 | 0.231 | 0.493 | 0.0002552 | 05_mo-DC1 |
| FOXN2 | -2.49485364 | 0.183 | 0.443 | 0.0002706 | 05_mo-DC1 |
| CCR7 | -4.09423096 | 0.067 | 0.327 | 0.0002759 | 05_mo-DC1 |
| NDUFS2 | 1.100117257 | 0.481 | 0.23 | 0.0003174 | 05_mo-DC1 |
| AC009093.2 | -3.2852503 | 0.077 | 0.337 | 0.0003628 | 05_mo-DC1 |
| IL18R1 | -2.42884598 | 0.096 | 0.362 | 0.0004135 | 05_mo-DC1 |
| SERPINE1 | 1.15816871 | 0.413 | 0.193 | 0.0004155 | 05_mo-DC1 |
| SDF2L1 | 1.695767275 | 0.375 | 0.165 | 0.0004186 | 05_mo-DC1 |
| NAA10 | 1.098176388 | 0.462 | 0.223 | 0.0004257 | 05_mo-DC1 |
| FAM107B | -2.53354489 | 0.115 | 0.38 | 0.0004867 | 05_mo-DC1 |
| PIH1D1 | 1.330958762 | 0.346 | 0.143 | 0.0005535 | 05_mo-DC1 |
| SLC1A5 | 1.072619296 | 0.442 | 0.201 | 0.0005753 | 05_mo-DC1 |
| FAM174C | 1.438486598 | 0.442 | 0.217 | 0.000579 | 05_mo-DC1 |
| LIMK2 | 1.101988229 | 0.404 | 0.179 | 0.0006427 | 05_mo-DC1 |
| IL2RG | -2.18471427 | 0.115 | 0.37 | 0.0007302 | 05_mo-DC1 |
| LAMP1 | 1.077140545 | 0.481 | 0.224 | 0.0007464 | 05_mo-DC1 |
| TMEM163 | -3.48965172 | 0.048 | 0.301 | 0.0007527 | 05_mo-DC1 |
| AL138963.4 | -4.08390143 | 0.029 | 0.28 | 0.0007601 | 05_mo-DC1 |
| GATD3B | 1.20949442 | 0.365 | 0.157 | 0.00089 | 05_mo-DC1 |
| SNCA | 1.03732462 | 0.413 | 0.189 | 0.0009438 | 05_mo-DC1 |
| EREG | -3.56433573 | 0.067 | 0.316 | 0.0009769 | 05_mo-DC1 |
| TMEM134 | 1.309929108 | 0.346 | 0.145 | 0.0009894 | 05_mo-DC1 |
| RAB11FIP1 | -2.27244397 | 0.135 | 0.393 | 0.0009987 | 05_mo-DC1 |
| SLC38A1 | -1.97902135 | 0.019 | 0.268 | 0.0010913 | 05_mo-DC1 |
| AL137857.1 | -3.70659773 | 0.029 | 0.272 | 0.0011416 | 05_mo-DC1 |
| UTY | -4.67792083 | 0.019 | 0.255 | 0.0011975 | 05_mo-DC1 |
| TRPS1 | -1.88057438 | 0.356 | 0.565 | 0.0012111 | 05_mo-DC1 |
| PMAIP1 | -2.53037996 | 0.106 | 0.366 | 0.0012272 | 05_mo-DC1 |
| PDE8A | 1.056374152 | 0.663 | 0.418 | 0.0012581 | 05_mo-DC1 |

| | | | | | |
|---|---|---|---|---|---|
| CYTOR | -2.17493697 | 0.115 | 0.382 | 0.0016172 | 05_mo-DC1 |
| DENND4A | -1.26687763 | 0.327 | 0.582 | 0.0026036 | 05_mo-DC1 |
| BAX | 1.002442573 | 0.615 | 0.386 | 0.0028035 | 05_mo-DC1 |
| C15orf48 | -2.44231311 | 0.308 | 0.519 | 0.002907 | 05_mo-DC1 |
| ZNF428 | 1.336422655 | 0.385 | 0.181 | 0.0032675 | 05_mo-DC1 |
| GPI | 1.585815375 | 0.538 | 0.322 | 0.0034169 | 05_mo-DC1 |
| CD1E | -6.03315023 | 0.01 | 0.228 | 0.0035824 | 05_mo-DC1 |
| HPCAL1 | 1.178524074 | 0.404 | 0.195 | 0.003829 | 05_mo-DC1 |
| BCL3 | -2.01926054 | 0.279 | 0.49 | 0.0039614 | 05_mo-DC1 |
| SIPA1L3 | -2.42871662 | 0.173 | 0.417 | 0.0042396 | 05_mo-DC1 |
| ODF3B | 1.067339164 | 0.606 | 0.387 | 0.0046598 | 05_mo-DC1 |
| AL133415.1 | -2.54170554 | 0.163 | 0.395 | 0.00505 | 05_mo-DC1 |
| CALCRL | -13.4811332 | 0 | 0.211 | 0.005146 | 05_mo-DC1 |
| IRF4 | -2.38811758 | 0.077 | 0.321 | 0.0052402 | 05_mo-DC1 |
| GPAT3 | -1.73220781 | 0.24 | 0.474 | 0.0055407 | 05_mo-DC1 |
| JAK1 | -1.14454256 | 0.442 | 0.645 | 0.0059566 | 05_mo-DC1 |
| MNDA | -2.01596833 | 0.144 | 0.396 | 0.0061886 | 05_mo-DC1 |
| AHNAK | -1.14726706 | 0.375 | 0.611 | 0.006295 | 05_mo-DC1 |
| PKIB | -1.69817198 | 0.183 | 0.426 | 0.0063967 | 05_mo-DC1 |
| MRPS34 | 1.059498299 | 0.385 | 0.183 | 0.0065734 | 05_mo-DC1 |
| NOSIP | 1.104741171 | 0.433 | 0.209 | 0.0067656 | 05_mo-DC1 |
| KYNU | -1.52906424 | 0.433 | 0.633 | 0.0071833 | 05_mo-DC1 |
| DAPK1 | -2.05847936 | 0.115 | 0.356 | 0.0075954 | 05_mo-DC1 |
| AC060765.2 | -12.468787 | 0 | 0.206 | 0.0076276 | 05_mo-DC1 |
| NUDT14 | 1.15022732 | 0.365 | 0.164 | 0.0076697 | 05_mo-DC1 |
| LSM2 | 1.012707447 | 0.404 | 0.188 | 0.0076842 | 05_mo-DC1 |
| TMEM131 | -1.8742611 | 0.154 | 0.403 | 0.0076891 | 05_mo-DC1 |
| TP53BP2 | -2.22631212 | 0.106 | 0.345 | 0.0105504 | 05_mo-DC1 |
| CERS6 | -1.80437217 | 0.144 | 0.38 | 0.0106781 | 05_mo-DC1 |
| FRY | -3.0629167 | 0.048 | 0.274 | 0.0120965 | 05_mo-DC1 |
| ARID1B | -1.41172557 | 0.25 | 0.495 | 0.0124157 | 05_mo-DC1 |
| SATB1 | -2.03499795 | 0.173 | 0.401 | 0.0140166 | 05_mo-DC1 |
| BHLHE40 | -1.12868009 | 0.337 | 0.577 | 0.0147779 | 05_mo-DC1 |
| SIGLEC9 | 1.020033865 | 0.375 | 0.167 | 0.0163886 | 05_mo-DC1 |
| USP9Y | -4.14616113 | 0.019 | 0.226 | 0.0170889 | 05_mo-DC1 |
| SMARCA2 | -1.36531975 | 0.317 | 0.54 | 0.0179269 | 05_mo-DC1 |
| NCF1 | 1.295364436 | 0.5 | 0.289 | 0.0183906 | 05_mo-DC1 |
| UQCRC1 | 1.032249892 | 0.548 | 0.323 | 0.0229944 | 05_mo-DC1 |
| RALA | -1.30043576 | 0.385 | 0.599 | 0.0254987 | 05_mo-DC1 |
| TES | -1.62819627 | 0.231 | 0.455 | 0.0255385 | 05_mo-DC1 |
| ANTXR2 | -4.09539098 | 0.029 | 0.232 | 0.0256139 | 05_mo-DC1 |
| H2AFJ | 1.079274989 | 0.462 | 0.253 | 0.0278546 | 05_mo-DC1 |
| GPR132 | -1.29203534 | 0.356 | 0.569 | 0.0291986 | 05_mo-DC1 |
| CLIC2 | -2.67880841 | 0.058 | 0.27 | 0.0302836 | 05_mo-DC1 |
| DUSP23 | 1.041007595 | 0.433 | 0.223 | 0.0326145 | 05_mo-DC1 |
| MS4A4E | -2.10606258 | 0.154 | 0.376 | 0.0354305 | 05_mo-DC1 |

| | | | | | |
|---|---|---|---|---|---|
| IDH3G | 1.039555163 | 0.433 | 0.226 | 0.0378853 | 05_mo-DC1 |
| CCL22 | -2.9361572 | 0.067 | 0.279 | 0.0405813 | 05_mo-DC1 |
| HLA-DQB2 | -2.53721631 | 0.144 | 0.345 | 0.0478514 | 05_mo-DC1 |
| PILRA | 1.105104969 | 0.452 | 0.229 | 0.0585323 | 05_mo-DC1 |
| VAV3 | -2.68191333 | 0.058 | 0.262 | 0.0750441 | 05_mo-DC1 |
| AUTS2 | -1.80418615 | 0.183 | 0.389 | 0.1012304 | 05_mo-DC1 |
| GAREM1 | -1.43963293 | 0.221 | 0.428 | 0.1085083 | 05_mo-DC1 |
| INSR | -1.79435547 | 0.173 | 0.379 | 0.1392775 | 05_mo-DC1 |
| CDC42SE2 | -1.18306746 | 0.25 | 0.473 | 0.1460108 | 05_mo-DC1 |
| AC004817.3 | 1.03600997 | 0.519 | 0.292 | 0.1639849 | 05_mo-DC1 |
| SLC41A2 | -1.93317135 | 0.087 | 0.294 | 0.190593 | 05_mo-DC1 |
| CD48 | -1.54068402 | 0.144 | 0.36 | 0.1924533 | 05_mo-DC1 |
| HIVEP1 | -2.06181783 | 0.144 | 0.345 | 0.1949528 | 05_mo-DC1 |
| HGSNAT | -1.00573097 | 0.298 | 0.503 | 0.4264832 | 05_mo-DC1 |
| PABPN1 | -1.07743429 | 0.298 | 0.499 | 0.5207405 | 05_mo-DC1 |
| ANKRD36B | 6.539934711 | 0.918 | 0.082 | 1.29E-135 | 06_mo-DC2 |
| AC092683.1 | 6.342397371 | 0.694 | 0.029 | 1.24E-134 | 06_mo-DC2 |
| LINGO1 | 4.409473498 | 0.765 | 0.093 | 4.76E-82 | 06_mo-DC2 |
| AC092683.2 | 6.670364833 | 0.337 | 0.006 | 5.18E-81 | 06_mo-DC2 |
| CACNA1A | 4.173959494 | 0.429 | 0.032 | 9.87E-56 | 06_mo-DC2 |
| APOC1 | 2.711084643 | 0.959 | 0.468 | 3.72E-38 | 06_mo-DC2 |
| LINC02642 | 3.768827146 | 0.367 | 0.037 | 7.56E-38 | 06_mo-DC2 |
| SLC1A3 | 2.009656409 | 0.99 | 0.494 | 1.85E-35 | 06_mo-DC2 |
| APOE | 2.355104128 | 0.98 | 0.562 | 8.82E-34 | 06_mo-DC2 |
| KCNQ3 | 2.201909373 | 0.735 | 0.191 | 3.14E-32 | 06_mo-DC2 |
| DLEU1 | 2.111809913 | 0.867 | 0.34 | 3.30E-32 | 06_mo-DC2 |
| MT3 | 3.253808909 | 0.316 | 0.031 | 6.72E-32 | 06_mo-DC2 |
| MT-ND6 | 1.718670899 | 1 | 0.78 | 1.31E-31 | 06_mo-DC2 |
| LHFPL2 | 1.742597477 | 0.776 | 0.225 | 2.72E-30 | 06_mo-DC2 |
| EPB41L2 | 2.01424431 | 0.908 | 0.439 | 2.14E-29 | 06_mo-DC2 |
| FP236383.3 | 1.700625884 | 0.99 | 0.746 | 3.35E-29 | 06_mo-DC2 |
| FRMD4A | 1.589647084 | 0.939 | 0.407 | 1.21E-28 | 06_mo-DC2 |
| CSGALNACT1 | 1.763968395 | 0.867 | 0.312 | 1.80E-28 | 06_mo-DC2 |
| C3 | 1.582172749 | 0.959 | 0.487 | 3.76E-28 | 06_mo-DC2 |
| MICU3 | 3.459380664 | 0.306 | 0.034 | 7.46E-28 | 06_mo-DC2 |
| SAP30 | 2.06640144 | 0.816 | 0.333 | 1.35E-27 | 06_mo-DC2 |
| PLA2G4A | 1.98370675 | 0.827 | 0.304 | 2.30E-27 | 06_mo-DC2 |
| ARHGAP22 | 2.409843003 | 0.653 | 0.189 | 7.32E-27 | 06_mo-DC2 |
| LRRK1 | 2.322109861 | 0.704 | 0.227 | 2.03E-26 | 06_mo-DC2 |
| SLC22A15 | 3.209085931 | 0.265 | 0.027 | 1.46E-25 | 06_mo-DC2 |
| USP53 | 2.113136615 | 0.837 | 0.374 | 1.03E-24 | 06_mo-DC2 |
| SFMBT2 | 1.670825197 | 0.898 | 0.42 | 3.64E-24 | 06_mo-DC2 |
| MEF2A | 1.501585764 | 0.98 | 0.73 | 7.88E-24 | 06_mo-DC2 |
| SYNDIG1 | 3.286741435 | 0.316 | 0.043 | 7.94E-24 | 06_mo-DC2 |
| ADGRB3 | 3.082668358 | 0.276 | 0.032 | 8.12E-24 | 06_mo-DC2 |
| SORL1 | 1.632317925 | 0.888 | 0.434 | 1.14E-23 | 06_mo-DC2 |

| | | | | | |
|---|---|---|---|---|---|
| ARHGAP24 | 1.771439717 | 0.827 | 0.34 | 6.77E-23 | 06_mo-DC2 |
| SLC11A1 | 1.544985745 | 0.908 | 0.423 | 8.68E-23 | 06_mo-DC2 |
| SCIN | 2.414874329 | 0.49 | 0.114 | 1.38E-22 | 06_mo-DC2 |
| AL163541.1 | 2.016652511 | 0.592 | 0.17 | 1.65E-22 | 06_mo-DC2 |
| PCNX2 | 1.831304807 | 0.673 | 0.218 | 4.83E-22 | 06_mo-DC2 |
| CD163 | 1.368011429 | 0.929 | 0.465 | 7.36E-22 | 06_mo-DC2 |
| ELL2 | 1.437892035 | 0.969 | 0.726 | 9.11E-22 | 06_mo-DC2 |
| EZR | -2.94221084 | 0.276 | 0.787 | 1.08E-21 | 06_mo-DC2 |
| FN1 | 2.539527305 | 0.531 | 0.146 | 1.29E-21 | 06_mo-DC2 |
| SERPINE1 | 3.493601188 | 0.582 | 0.182 | 1.75E-21 | 06_mo-DC2 |
| L3MBTL4 | 2.207011581 | 0.541 | 0.144 | 2.62E-21 | 06_mo-DC2 |
| ZBTB16 | 1.64074851 | 0.888 | 0.461 | 8.44E-21 | 06_mo-DC2 |
| SPRED1 | 1.720645644 | 0.531 | 0.137 | 1.27E-20 | 06_mo-DC2 |
| MEF2C | 1.436475693 | 0.98 | 0.631 | 1.46E-20 | 06_mo-DC2 |
| GFAP | 2.873909245 | 0.357 | 0.066 | 2.64E-20 | 06_mo-DC2 |
| JDP2 | 1.496670225 | 0.867 | 0.425 | 2.77E-20 | 06_mo-DC2 |
| LINC00937 | 2.529689368 | 0.367 | 0.069 | 4.18E-20 | 06_mo-DC2 |
| C1QC | 1.440995588 | 0.908 | 0.521 | 4.69E-20 | 06_mo-DC2 |
| MSR1 | 1.429887568 | 0.918 | 0.509 | 1.09E-19 | 06_mo-DC2 |
| SSH2 | 1.57364057 | 0.898 | 0.559 | 1.15E-19 | 06_mo-DC2 |
| MARCKS | 1.382858765 | 0.918 | 0.512 | 1.91E-19 | 06_mo-DC2 |
| LINC02432 | 3.670386111 | 0.235 | 0.028 | 3.29E-19 | 06_mo-DC2 |
| ACKR3 | 2.054548383 | 0.48 | 0.119 | 3.56E-19 | 06_mo-DC2 |
| CD69 | 1.451378714 | 0.847 | 0.409 | 1.82E-18 | 06_mo-DC2 |
| RPS6KA2 | 1.975993067 | 0.602 | 0.197 | 2.04E-18 | 06_mo-DC2 |
| GLDN | 2.13630415 | 0.48 | 0.125 | 2.49E-18 | 06_mo-DC2 |
| DPYD | 1.401131939 | 0.939 | 0.584 | 6.23E-18 | 06_mo-DC2 |
| SPATA13 | 2.495800099 | 0.531 | 0.165 | 6.56E-18 | 06_mo-DC2 |
| MLXIPL | 2.6983127 | 0.286 | 0.046 | 1.37E-17 | 06_mo-DC2 |
| C1QA | 1.433007042 | 0.878 | 0.48 | 1.42E-17 | 06_mo-DC2 |
| APBB1IP | 1.361087304 | 0.847 | 0.476 | 1.98E-17 | 06_mo-DC2 |
| KCNIP1 | 2.933208336 | 0.316 | 0.059 | 2.08E-17 | 06_mo-DC2 |
| LINC01684 | 2.686344234 | 0.286 | 0.047 | 2.17E-17 | 06_mo-DC2 |
| RPS5 | -1.77401167 | 0.612 | 0.848 | 2.93E-17 | 06_mo-DC2 |
| FCGBP | 1.541110945 | 0.5 | 0.133 | 3.67E-17 | 06_mo-DC2 |
| TREM2 | 1.409901892 | 0.786 | 0.357 | 4.65E-17 | 06_mo-DC2 |
| LPCAT2 | 1.795774401 | 0.837 | 0.469 | 8.42E-17 | 06_mo-DC2 |
| FAM177B | 3.113167461 | 0.265 | 0.042 | 1.82E-16 | 06_mo-DC2 |
| PPARG | 2.247917294 | 0.449 | 0.119 | 2.07E-16 | 06_mo-DC2 |
| AL078590.2 | 2.152985376 | 0.714 | 0.35 | 2.09E-16 | 06_mo-DC2 |
| OLR1 | 1.128034192 | 0.98 | 0.642 | 3.52E-16 | 06_mo-DC2 |
| FP671120.4 | 1.417194215 | 0.908 | 0.562 | 3.79E-16 | 06_mo-DC2 |
| CCDC26 | 1.999246252 | 0.449 | 0.12 | 4.18E-16 | 06_mo-DC2 |
| RPL8 | -1.40275199 | 0.704 | 0.905 | 5.86E-16 | 06_mo-DC2 |
| ZFP36L2 | 1.077301417 | 1 | 0.729 | 6.82E-16 | 06_mo-DC2 |
| LPAR1 | 2.401968955 | 0.327 | 0.065 | 7.25E-16 | 06_mo-DC2 |

| | | | | | |
|---|---|---|---|---|---|
| HSPA8 | -2.25463412 | 0.337 | 0.756 | 1.13E-15 | 06_mo-DC2 |
| FAM168A | 2.061375562 | 0.48 | 0.139 | 1.36E-15 | 06_mo-DC2 |
| SLCO2B1 | 1.528333191 | 0.602 | 0.206 | 2.48E-15 | 06_mo-DC2 |
| DUSP4 | -3.92488826 | 0.071 | 0.575 | 2.72E-15 | 06_mo-DC2 |
| C15orf48 | -6.66156524 | 0.041 | 0.537 | 2.93E-15 | 06_mo-DC2 |
| ZNF331 | 1.188854365 | 0.939 | 0.669 | 3.18E-15 | 06_mo-DC2 |
| LNCAROD | 1.575408785 | 0.531 | 0.164 | 3.91E-15 | 06_mo-DC2 |
| EGR3 | 2.138302763 | 0.418 | 0.108 | 4.68E-15 | 06_mo-DC2 |
| AC245297.3 | 2.348683097 | 0.449 | 0.126 | 5.92E-15 | 06_mo-DC2 |
| OSBPL1A | 2.22292339 | 0.357 | 0.081 | 6.70E-15 | 06_mo-DC2 |
| RPL21 | -1.57713824 | 0.602 | 0.855 | 6.83E-15 | 06_mo-DC2 |
| IL1R2 | -6.68642432 | 0.01 | 0.508 | 8.79E-15 | 06_mo-DC2 |
| FAM149A | 1.667197499 | 0.571 | 0.196 | 9.13E-15 | 06_mo-DC2 |
| LSP1 | -2.10099161 | 0.367 | 0.746 | 2.03E-14 | 06_mo-DC2 |
| MIR646HG | 1.828813869 | 0.459 | 0.135 | 5.77E-14 | 06_mo-DC2 |
| AC012368.1 | 2.157569663 | 0.398 | 0.107 | 1.26E-13 | 06_mo-DC2 |
| CST7 | -5.41457802 | 0.051 | 0.518 | 1.34E-13 | 06_mo-DC2 |
| DEPTOR | 1.900001827 | 0.459 | 0.139 | 1.85E-13 | 06_mo-DC2 |
| VASH1 | 1.273968322 | 0.867 | 0.528 | 1.99E-13 | 06_mo-DC2 |
| P2RY13 | 2.157808427 | 0.439 | 0.135 | 2.79E-13 | 06_mo-DC2 |
| CD44 | -2.67614308 | 0.306 | 0.718 | 3.21E-13 | 06_mo-DC2 |
| AC079015.1 | 2.286572714 | 0.265 | 0.049 | 4.44E-13 | 06_mo-DC2 |
| TIMP1 | -3.67625271 | 0.357 | 0.701 | 5.92E-13 | 06_mo-DC2 |
| LINC01374 | 1.78972546 | 0.418 | 0.121 | 8.43E-13 | 06_mo-DC2 |
| LYZ | -2.54849823 | 0.398 | 0.73 | 8.96E-13 | 06_mo-DC2 |
| ST6GAL1 | 1.460161065 | 0.633 | 0.275 | 1.02E-12 | 06_mo-DC2 |
| BNC2 | 1.653261058 | 0.49 | 0.163 | 1.18E-12 | 06_mo-DC2 |
| AC243960.1 | 2.519105916 | 0.276 | 0.056 | 1.48E-12 | 06_mo-DC2 |
| RALA | -2.7784455 | 0.163 | 0.614 | 1.79E-12 | 06_mo-DC2 |
| TAGLN2 | -2.37126501 | 0.367 | 0.703 | 1.94E-12 | 06_mo-DC2 |
| CRYBG1 | -2.99007896 | 0.184 | 0.611 | 2.21E-12 | 06_mo-DC2 |
| CCNI | -1.56987073 | 0.439 | 0.792 | 3.34E-12 | 06_mo-DC2 |
| SLC2A5 | 1.770908241 | 0.429 | 0.127 | 3.62E-12 | 06_mo-DC2 |
| TRIM56 | 2.486380925 | 0.316 | 0.076 | 4.90E-12 | 06_mo-DC2 |
| ADORA3 | 2.097360708 | 0.347 | 0.089 | 5.02E-12 | 06_mo-DC2 |
| AC005670.3 | 2.130434616 | 0.327 | 0.078 | 5.12E-12 | 06_mo-DC2 |
| EEF1B2 | -1.52997416 | 0.592 | 0.809 | 7.64E-12 | 06_mo-DC2 |
| JARID2 | -2.15609652 | 0.459 | 0.779 | 8.29E-12 | 06_mo-DC2 |
| EGR2 | 2.048468648 | 0.388 | 0.112 | 9.31E-12 | 06_mo-DC2 |
| ANKRD36C | 2.192350328 | 0.316 | 0.076 | 1.08E-11 | 06_mo-DC2 |
| CASS4 | 1.323681306 | 0.622 | 0.266 | 1.11E-11 | 06_mo-DC2 |
| FYB1 | 1.124846482 | 0.857 | 0.476 | 1.17E-11 | 06_mo-DC2 |
| MCF2L | 2.093784588 | 0.347 | 0.091 | 1.21E-11 | 06_mo-DC2 |
| ZNF804A | 2.034665745 | 0.378 | 0.108 | 2.06E-11 | 06_mo-DC2 |
| CTDSP1 | 1.76288946 | 0.571 | 0.238 | 2.07E-11 | 06_mo-DC2 |
| CRIP1 | -5.13339962 | 0.031 | 0.465 | 2.25E-11 | 06_mo-DC2 |

| | | | | | |
|---|---|---|---|---|---|
| AP001636.3 | 1.826837321 | 0.5 | 0.188 | 2.73E-11 | 06_mo-DC2 |
| EPC1 | 1.40386821 | 0.786 | 0.47 | 3.00E-11 | 06_mo-DC2 |
| OGFRL1 | 1.258402365 | 0.857 | 0.656 | 3.02E-11 | 06_mo-DC2 |
| DUSP6 | 1.891517818 | 0.378 | 0.11 | 3.04E-11 | 06_mo-DC2 |
| TPST1 | 1.899628519 | 0.347 | 0.091 | 3.30E-11 | 06_mo-DC2 |
| HTRA1 | 1.790652486 | 0.469 | 0.159 | 3.49E-11 | 06_mo-DC2 |
| G0S2 | -7.12980334 | 0.01 | 0.437 | 3.73E-11 | 06_mo-DC2 |
| GALNT2 | 1.479285891 | 0.643 | 0.283 | 3.90E-11 | 06_mo-DC2 |
| AC046195.1 | 2.336691734 | 0.265 | 0.057 | 5.53E-11 | 06_mo-DC2 |
| INSIG1 | -2.54013424 | 0.204 | 0.609 | 5.96E-11 | 06_mo-DC2 |
| LDHA | -1.84114917 | 0.408 | 0.745 | 7.76E-11 | 06_mo-DC2 |
| SUMF1 | 1.527325729 | 0.612 | 0.275 | 7.91E-11 | 06_mo-DC2 |
| RHBDF2 | 1.063416559 | 0.837 | 0.518 | 8.02E-11 | 06_mo-DC2 |
| AC012150.1 | 2.438272822 | 0.255 | 0.053 | 8.25E-11 | 06_mo-DC2 |
| INPP5D | 1.322363897 | 0.755 | 0.427 | 8.36E-11 | 06_mo-DC2 |
| HSPA1A | -4.85802058 | 0.204 | 0.594 | 8.86E-11 | 06_mo-DC2 |
| BCL3 | -3.83783308 | 0.092 | 0.502 | 1.04E-10 | 06_mo-DC2 |
| IFITM2 | -2.24135331 | 0.51 | 0.761 | 1.06E-10 | 06_mo-DC2 |
| YPEL3 | 1.524222056 | 0.551 | 0.224 | 1.79E-10 | 06_mo-DC2 |
| CD55 | -1.76334299 | 0.398 | 0.756 | 2.10E-10 | 06_mo-DC2 |
| FAM135A | 1.788825767 | 0.51 | 0.203 | 2.28E-10 | 06_mo-DC2 |
| NFIA | 1.914695395 | 0.316 | 0.081 | 2.46E-10 | 06_mo-DC2 |
| TGFBR1 | 1.856945797 | 0.633 | 0.321 | 2.48E-10 | 06_mo-DC2 |
| DLEU7 | 2.420719913 | 0.265 | 0.059 | 2.62E-10 | 06_mo-DC2 |
| CSF3R | 1.138501155 | 0.816 | 0.514 | 2.97E-10 | 06_mo-DC2 |
| PALD1 | 1.121377312 | 0.776 | 0.482 | 3.01E-10 | 06_mo-DC2 |
| EEF1G | -1.26169914 | 0.561 | 0.833 | 3.22E-10 | 06_mo-DC2 |
| QKI | 1.290547021 | 0.898 | 0.694 | 3.82E-10 | 06_mo-DC2 |
| ADAM17 | 1.223848264 | 0.745 | 0.414 | 5.98E-10 | 06_mo-DC2 |
| IPCEF1 | 2.16177627 | 0.439 | 0.161 | 6.25E-10 | 06_mo-DC2 |
| LITAF | -1.40675941 | 0.622 | 0.851 | 6.40E-10 | 06_mo-DC2 |
| SYNGR2 | -1.61180265 | 0.459 | 0.757 | 6.89E-10 | 06_mo-DC2 |
| RPL41 | -2.10108077 | 0.612 | 0.868 | 7.96E-10 | 06_mo-DC2 |
| FAR2 | 1.644424007 | 0.429 | 0.142 | 8.88E-10 | 06_mo-DC2 |
| CPEB3 | 1.798658801 | 0.429 | 0.155 | 1.28E-09 | 06_mo-DC2 |
| BAZ2B | 1.239162054 | 0.673 | 0.347 | 1.49E-09 | 06_mo-DC2 |
| LINC02256 | 1.463199364 | 0.643 | 0.309 | 2.12E-09 | 06_mo-DC2 |
| UNC5B | 2.063103725 | 0.286 | 0.071 | 2.52E-09 | 06_mo-DC2 |
| CLEC10A | -7.08489402 | 0.01 | 0.392 | 3.30E-09 | 06_mo-DC2 |
| LYPLAL1 | 1.966852942 | 0.347 | 0.102 | 3.47E-09 | 06_mo-DC2 |
| CLEC12A | 1.972109156 | 0.408 | 0.143 | 4.30E-09 | 06_mo-DC2 |
| CLIC1 | -1.2821736 | 0.561 | 0.804 | 4.30E-09 | 06_mo-DC2 |
| DOCK10 | 1.156780738 | 0.755 | 0.47 | 4.77E-09 | 06_mo-DC2 |
| SPATA6 | 1.906214262 | 0.327 | 0.094 | 5.52E-09 | 06_mo-DC2 |
| AL691403.1 | 1.47552492 | 0.469 | 0.173 | 5.54E-09 | 06_mo-DC2 |
| FRMD4B | 1.280548358 | 0.592 | 0.267 | 5.85E-09 | 06_mo-DC2 |

| | | | | | |
|---|---|---|---|---|---|
| TMOD1 | 2.040114825 | 0.316 | 0.087 | 6.13E-09 | 06_mo-DC2 |
| FCER1A | -4.09960272 | 0.153 | 0.494 | 6.44E-09 | 06_mo-DC2 |
| JAK2 | 1.607066478 | 0.602 | 0.297 | 6.71E-09 | 06_mo-DC2 |
| MAN2A1 | 1.058856263 | 0.765 | 0.439 | 6.78E-09 | 06_mo-DC2 |
| RPL7 | -1.38031009 | 0.551 | 0.767 | 6.95E-09 | 06_mo-DC2 |
| EMP3 | -2.12104898 | 0.429 | 0.704 | 7.55E-09 | 06_mo-DC2 |
| TRPM2 | 1.716951749 | 0.48 | 0.193 | 8.72E-09 | 06_mo-DC2 |
| MITF | 1.596760781 | 0.459 | 0.175 | 1.06E-08 | 06_mo-DC2 |
| CFLAR | -1.60874046 | 0.49 | 0.766 | 1.16E-08 | 06_mo-DC2 |
| FLNA | -3.72154292 | 0.051 | 0.433 | 1.36E-08 | 06_mo-DC2 |
| THADA | 1.511186762 | 0.531 | 0.23 | 1.48E-08 | 06_mo-DC2 |
| UBA7 | 1.932687768 | 0.367 | 0.121 | 1.71E-08 | 06_mo-DC2 |
| PIM3 | -3.10546991 | 0.112 | 0.481 | 2.25E-08 | 06_mo-DC2 |
| SLC4A7 | 1.297671257 | 0.622 | 0.313 | 2.53E-08 | 06_mo-DC2 |
| CD1C | -14.2424978 | 0 | 0.361 | 2.79E-08 | 06_mo-DC2 |
| BLNK | 2.01486745 | 0.286 | 0.077 | 2.96E-08 | 06_mo-DC2 |
| PSME4 | 1.171206604 | 0.633 | 0.311 | 3.09E-08 | 06_mo-DC2 |
| C20orf194 | 1.646622094 | 0.48 | 0.194 | 3.52E-08 | 06_mo-DC2 |
| ZNRF2 | 1.856040006 | 0.388 | 0.132 | 3.58E-08 | 06_mo-DC2 |
| ARHGAP25 | 1.803045604 | 0.388 | 0.134 | 3.86E-08 | 06_mo-DC2 |
| ANXA2 | -1.86604425 | 0.347 | 0.655 | 4.04E-08 | 06_mo-DC2 |
| HINT1 | -1.59743333 | 0.378 | 0.697 | 4.08E-08 | 06_mo-DC2 |
| AC084871.1 | 1.679858831 | 0.357 | 0.112 | 4.22E-08 | 06_mo-DC2 |
| CNN2 | -2.81528748 | 0.112 | 0.477 | 4.88E-08 | 06_mo-DC2 |
| AC020916.1 | 1.141487337 | 0.796 | 0.511 | 4.96E-08 | 06_mo-DC2 |
| CEBPA | 2.086871621 | 0.357 | 0.119 | 6.57E-08 | 06_mo-DC2 |
| TTC33 | 1.65972317 | 0.408 | 0.153 | 7.62E-08 | 06_mo-DC2 |
| ADAMTS10 | 2.245085221 | 0.306 | 0.088 | 8.07E-08 | 06_mo-DC2 |
| 3-Mar | 1.45555179 | 0.592 | 0.276 | 8.45E-08 | 06_mo-DC2 |
| SQSTM1 | -1.62004994 | 0.459 | 0.761 | 9.09E-08 | 06_mo-DC2 |
| PPAN | 1.908317579 | 0.306 | 0.09 | 1.30E-07 | 06_mo-DC2 |
| CHD9 | 1.23291374 | 0.633 | 0.334 | 1.52E-07 | 06_mo-DC2 |
| SERPINB1 | -1.93849115 | 0.5 | 0.727 | 1.73E-07 | 06_mo-DC2 |
| STX17-AS1 | 1.2036944 | 0.776 | 0.499 | 2.08E-07 | 06_mo-DC2 |
| IFITM1 | -3.77257941 | 0.082 | 0.429 | 2.16E-07 | 06_mo-DC2 |
| TRAF1 | -6.54061181 | 0.01 | 0.352 | 2.19E-07 | 06_mo-DC2 |
| CD14 | 1.040923314 | 0.724 | 0.421 | 2.27E-07 | 06_mo-DC2 |
| GCLC | 1.612160604 | 0.388 | 0.139 | 2.28E-07 | 06_mo-DC2 |
| NCF1 | 1.142646207 | 0.592 | 0.283 | 2.45E-07 | 06_mo-DC2 |
| TMEM117 | 1.419453276 | 0.296 | 0.085 | 3.13E-07 | 06_mo-DC2 |
| BIRC3 | -4.44757958 | 0.071 | 0.407 | 3.66E-07 | 06_mo-DC2 |
| PCED1B | 2.170047501 | 0.306 | 0.095 | 3.81E-07 | 06_mo-DC2 |
| VEGFA | -2.19352992 | 0.235 | 0.572 | 4.01E-07 | 06_mo-DC2 |
| AC068587.4 | 1.527791885 | 0.347 | 0.115 | 5.20E-07 | 06_mo-DC2 |
| B3GNT5 | 1.213511206 | 0.806 | 0.566 | 5.27E-07 | 06_mo-DC2 |
| SCPEP1 | 1.492909654 | 0.561 | 0.273 | 7.83E-07 | 06_mo-DC2 |

| | | | | | |
|---|---|---|---|---|---|
| S100A6 | -1.58901858 | 0.51 | 0.736 | 8.93E-07 | 06_mo-DC2 |
| MXD1 | -2.17152391 | 0.143 | 0.509 | 9.15E-07 | 06_mo-DC2 |
| LINC00623 | 1.60447004 | 0.408 | 0.16 | 9.32E-07 | 06_mo-DC2 |
| PMS1 | 1.855290743 | 0.378 | 0.144 | 1.01E-06 | 06_mo-DC2 |
| DNAJB1 | -3.69814386 | 0.143 | 0.488 | 1.04E-06 | 06_mo-DC2 |
| PWWP3A | 2.188702972 | 0.316 | 0.103 | 1.04E-06 | 06_mo-DC2 |
| TEX14 | 1.34630125 | 0.673 | 0.426 | 1.11E-06 | 06_mo-DC2 |
| ANKRD36 | 1.83090315 | 0.296 | 0.09 | 1.20E-06 | 06_mo-DC2 |
| CCSER1 | -2.95546534 | 0.143 | 0.484 | 1.28E-06 | 06_mo-DC2 |
| FYN | -3.94377778 | 0.051 | 0.384 | 1.34E-06 | 06_mo-DC2 |
| DUSP5 | -3.57025802 | 0.143 | 0.457 | 1.35E-06 | 06_mo-DC2 |
| GPAT3 | -1.84510084 | 0.143 | 0.48 | 1.50E-06 | 06_mo-DC2 |
| THBS1 | -5.02834728 | 0.061 | 0.385 | 1.53E-06 | 06_mo-DC2 |
| CYTIP | -1.63878044 | 0.296 | 0.612 | 1.56E-06 | 06_mo-DC2 |
| BHLHE41 | 1.352195111 | 0.51 | 0.239 | 2.00E-06 | 06_mo-DC2 |
| ABCA1 | 1.073900742 | 0.602 | 0.333 | 2.11E-06 | 06_mo-DC2 |
| MIS18BP1 | 1.419017325 | 0.673 | 0.41 | 2.21E-06 | 06_mo-DC2 |
| PTGER4 | 1.257311105 | 0.673 | 0.407 | 2.54E-06 | 06_mo-DC2 |
| HSPA1B | -3.86428226 | 0.122 | 0.463 | 2.76E-06 | 06_mo-DC2 |
| TMCC1 | 1.380690883 | 0.459 | 0.201 | 2.94E-06 | 06_mo-DC2 |
| GPR137B | -2.58297363 | 0.173 | 0.496 | 2.95E-06 | 06_mo-DC2 |
| TUT7 | 1.330831489 | 0.561 | 0.292 | 2.95E-06 | 06_mo-DC2 |
| COMMD6 | -1.48955697 | 0.337 | 0.651 | 3.27E-06 | 06_mo-DC2 |
| CDKN1A | -1.3669913 | 0.52 | 0.742 | 3.36E-06 | 06_mo-DC2 |
| RNASEK | -1.01944741 | 0.592 | 0.805 | 3.66E-06 | 06_mo-DC2 |
| GRINA | -1.58726794 | 0.418 | 0.664 | 4.84E-06 | 06_mo-DC2 |
| NINJ1 | -2.03905282 | 0.286 | 0.573 | 4.92E-06 | 06_mo-DC2 |
| SLC9A9 | 1.407427611 | 0.449 | 0.194 | 5.45E-06 | 06_mo-DC2 |
| LMNA | -2.51674202 | 0.163 | 0.483 | 5.52E-06 | 06_mo-DC2 |
| IL1R1 | -6.7655407 | 0.01 | 0.314 | 6.14E-06 | 06_mo-DC2 |
| LPAR6 | 1.246202071 | 0.469 | 0.198 | 6.98E-06 | 06_mo-DC2 |
| PRKN | 1.783432802 | 0.316 | 0.106 | 7.43E-06 | 06_mo-DC2 |
| GIMAP4 | 1.496862231 | 0.378 | 0.145 | 7.66E-06 | 06_mo-DC2 |
| CCR7 | -4.79327766 | 0.02 | 0.329 | 8.17E-06 | 06_mo-DC2 |
| TES | -1.88238148 | 0.133 | 0.46 | 8.60E-06 | 06_mo-DC2 |
| GARS-DT | 1.787685968 | 0.316 | 0.109 | 8.79E-06 | 06_mo-DC2 |
| AC004687.1 | 1.10056647 | 0.704 | 0.458 | 9.52E-06 | 06_mo-DC2 |
| TENM4 | 1.243016383 | 0.418 | 0.177 | 9.94E-06 | 06_mo-DC2 |
| TNS3 | 1.013114135 | 0.51 | 0.234 | 1.01E-05 | 06_mo-DC2 |
| ARAP1 | 1.24928511 | 0.541 | 0.273 | 1.01E-05 | 06_mo-DC2 |
| TNK2 | 1.748116003 | 0.347 | 0.126 | 1.12E-05 | 06_mo-DC2 |
| VDR | -4.56776702 | 0.01 | 0.314 | 1.19E-05 | 06_mo-DC2 |
| GBP2 | 1.433138062 | 0.571 | 0.309 | 1.24E-05 | 06_mo-DC2 |
| CLEC9A | 1.084272106 | 0.337 | 0.115 | 1.24E-05 | 06_mo-DC2 |
| ATP2B1 | -1.77961025 | 0.316 | 0.609 | 1.29E-05 | 06_mo-DC2 |
| ADAM19 | -6.54115873 | 0.01 | 0.307 | 1.36E-05 | 06_mo-DC2 |

| | | | | | |
|---|---|---|---|---|---|
| ATP13A3 | -1.83418112 | 0.265 | 0.585 | 1.42E-05 | 06_mo-DC2 |
| DUSP2 | -1.67129615 | 0.306 | 0.622 | 1.48E-05 | 06_mo-DC2 |
| LGALS3 | -2.47300145 | 0.173 | 0.488 | 1.49E-05 | 06_mo-DC2 |
| GNA12 | -2.54507654 | 0.163 | 0.47 | 1.49E-05 | 06_mo-DC2 |
| RASAL2 | 1.083005724 | 0.582 | 0.292 | 1.54E-05 | 06_mo-DC2 |
| IRF4 | -4.26003846 | 0.02 | 0.324 | 1.55E-05 | 06_mo-DC2 |
| ETV3 | -1.84916524 | 0.245 | 0.572 | 1.60E-05 | 06_mo-DC2 |
| PDCD4 | 1.262814995 | 0.551 | 0.286 | 1.76E-05 | 06_mo-DC2 |
| IL2RG | -3.50901708 | 0.071 | 0.372 | 1.83E-05 | 06_mo-DC2 |
| SPINT2 | -1.69944998 | 0.214 | 0.538 | 1.92E-05 | 06_mo-DC2 |
| SEC11A | -1.30227501 | 0.378 | 0.662 | 1.99E-05 | 06_mo-DC2 |
| LCP1 | -1.58416106 | 0.418 | 0.689 | 2.15E-05 | 06_mo-DC2 |
| MRC1 | -6.09708855 | 0.01 | 0.302 | 2.33E-05 | 06_mo-DC2 |
| IQGAP2 | -2.30015748 | 0.194 | 0.508 | 2.42E-05 | 06_mo-DC2 |
| SLC16A10 | -3.33914792 | 0.122 | 0.427 | 2.81E-05 | 06_mo-DC2 |
| MGAT4A | 1.081706721 | 0.582 | 0.323 | 2.91E-05 | 06_mo-DC2 |
| FAM110B | 1.120624019 | 0.388 | 0.15 | 3.16E-05 | 06_mo-DC2 |
| ZNF618 | 1.589226706 | 0.347 | 0.135 | 3.61E-05 | 06_mo-DC2 |
| TBC1D2B | 1.420737927 | 0.378 | 0.149 | 3.66E-05 | 06_mo-DC2 |
| ATF6B | 1.176495565 | 0.408 | 0.17 | 3.99E-05 | 06_mo-DC2 |
| GPR157 | -13.49305 | 0 | 0.281 | 4.05E-05 | 06_mo-DC2 |
| PTPRJ | 1.053526197 | 0.633 | 0.376 | 4.17E-05 | 06_mo-DC2 |
| ANXA11 | -1.78098227 | 0.224 | 0.523 | 4.51E-05 | 06_mo-DC2 |
| SH3BP5 | -2.16683115 | 0.173 | 0.479 | 4.70E-05 | 06_mo-DC2 |
| ORMDL1 | 1.270232018 | 0.602 | 0.345 | 4.82E-05 | 06_mo-DC2 |
| HIVEP2 | -2.06250827 | 0.245 | 0.545 | 4.84E-05 | 06_mo-DC2 |
| SLC26A3 | 1.594768688 | 0.418 | 0.178 | 5.24E-05 | 06_mo-DC2 |
| SOD2 | -1.59613059 | 0.429 | 0.703 | 5.46E-05 | 06_mo-DC2 |
| SNHG15 | -1.98523209 | 0.184 | 0.494 | 5.82E-05 | 06_mo-DC2 |
| SLC25A5 | -1.24921222 | 0.429 | 0.698 | 5.85E-05 | 06_mo-DC2 |
| PDK4 | 1.004729657 | 0.48 | 0.23 | 5.97E-05 | 06_mo-DC2 |
| MCOLN2 | -2.83327224 | 0.082 | 0.385 | 6.16E-05 | 06_mo-DC2 |
| FAM13A | 1.370346939 | 0.337 | 0.124 | 6.60E-05 | 06_mo-DC2 |
| RERE | 1.146623469 | 0.633 | 0.376 | 7.30E-05 | 06_mo-DC2 |
| CHD7 | 1.500705289 | 0.327 | 0.123 | 7.63E-05 | 06_mo-DC2 |
| H2AFZ | -1.24349817 | 0.378 | 0.65 | 7.99E-05 | 06_mo-DC2 |
| ADAM8 | -3.90427221 | 0.031 | 0.315 | 9.64E-05 | 06_mo-DC2 |
| CCL22 | -8.86020071 | 0.01 | 0.282 | 0.0001026 | 06_mo-DC2 |
| WDR70 | 1.241544941 | 0.418 | 0.185 | 0.0001139 | 06_mo-DC2 |
| SLC38A1 | -12.8868278 | 0 | 0.268 | 0.0001178 | 06_mo-DC2 |
| WDR1 | -1.74207767 | 0.214 | 0.508 | 0.000123 | 06_mo-DC2 |
| MYL12B | -1.06647465 | 0.52 | 0.736 | 0.0001319 | 06_mo-DC2 |
| WDFY3 | 1.347586884 | 0.408 | 0.18 | 0.0001493 | 06_mo-DC2 |
| HIVEP1 | -3.37971152 | 0.071 | 0.35 | 0.0001836 | 06_mo-DC2 |
| EPS8 | 1.340648342 | 0.582 | 0.33 | 0.0001847 | 06_mo-DC2 |
| SKI | 1.148907665 | 0.51 | 0.271 | 0.0002029 | 06_mo-DC2 |

| | | | | | |
|---|---|---|---|---|---|
| ITFG1 | 1.134712519 | 0.48 | 0.237 | 0.0002157 | 06_mo-DC2 |
| GSAP | 1.541889996 | 0.418 | 0.196 | 0.0002513 | 06_mo-DC2 |
| TUT4 | 1.36727198 | 0.48 | 0.239 | 0.0002526 | 06_mo-DC2 |
| SUCLG2 | 1.671042992 | 0.378 | 0.167 | 0.00026 | 06_mo-DC2 |
| PKN2 | 1.17617371 | 0.52 | 0.284 | 0.0002873 | 06_mo-DC2 |
| TMEM259 | 1.007520981 | 0.592 | 0.336 | 0.0003156 | 06_mo-DC2 |
| ZHX2 | -3.45070904 | 0.041 | 0.316 | 0.000337 | 06_mo-DC2 |
| APOC2 | 2.167980456 | 0.418 | 0.201 | 0.0003398 | 06_mo-DC2 |
| RAB8B | -1.51708359 | 0.204 | 0.499 | 0.0003559 | 06_mo-DC2 |
| PIK3IP1 | 1.577648376 | 0.357 | 0.148 | 0.0003577 | 06_mo-DC2 |
| LINC01480 | 2.205597074 | 0.337 | 0.133 | 0.0003588 | 06_mo-DC2 |
| RUNX3 | -3.52286311 | 0.061 | 0.331 | 0.0003613 | 06_mo-DC2 |
| HSPH1 | -3.13994654 | 0.153 | 0.435 | 0.0003694 | 06_mo-DC2 |
| IL7R | -14.1500263 | 0 | 0.253 | 0.0004213 | 06_mo-DC2 |
| EREG | -4.2489399 | 0.051 | 0.316 | 0.0004547 | 06_mo-DC2 |
| ZC3H12A | -2.86948226 | 0.071 | 0.343 | 0.0005003 | 06_mo-DC2 |
| UBASH3B | -2.09918616 | 0.173 | 0.462 | 0.0005106 | 06_mo-DC2 |
| CPVL | -1.6017579 | 0.459 | 0.662 | 0.0005287 | 06_mo-DC2 |
| CYP2S1 | -2.91601194 | 0.051 | 0.323 | 0.0005484 | 06_mo-DC2 |
| FHIT | 1.033928466 | 0.541 | 0.311 | 0.0005771 | 06_mo-DC2 |
| ARF6 | -1.51080702 | 0.357 | 0.614 | 0.00061 | 06_mo-DC2 |
| LAPTM4A | -1.26000782 | 0.398 | 0.651 | 0.0006313 | 06_mo-DC2 |
| LRCH1 | 1.101315527 | 0.469 | 0.235 | 0.0007177 | 06_mo-DC2 |
| ARHGEF6 | 1.330587401 | 0.408 | 0.189 | 0.0007205 | 06_mo-DC2 |
| MYLIP | 1.210505449 | 0.418 | 0.197 | 0.0007279 | 06_mo-DC2 |
| CLIC2 | -4.71180334 | 0.02 | 0.272 | 0.0007432 | 06_mo-DC2 |
| PPIF | -2.06221125 | 0.163 | 0.456 | 0.0007572 | 06_mo-DC2 |
| RASA3 | 1.022348801 | 0.51 | 0.259 | 0.0008075 | 06_mo-DC2 |
| SRGAP2C | 1.122960126 | 0.541 | 0.297 | 0.0008453 | 06_mo-DC2 |
| RPL23 | -1.09279393 | 0.5 | 0.715 | 0.0008708 | 06_mo-DC2 |
| RFTN1 | -1.82389092 | 0.286 | 0.543 | 0.000888 | 06_mo-DC2 |
| PDE4A | -1.37874374 | 0.255 | 0.542 | 0.0009529 | 06_mo-DC2 |
| OLFML3 | 1.392549834 | 0.398 | 0.181 | 0.0009542 | 06_mo-DC2 |
| BMP2K | 1.23035924 | 0.551 | 0.323 | 0.0010245 | 06_mo-DC2 |
| HMGN1 | -1.12480737 | 0.429 | 0.647 | 0.0011144 | 06_mo-DC2 |
| AC245014.3 | 1.382021987 | 0.347 | 0.143 | 0.0011696 | 06_mo-DC2 |
| SULF2 | -2.88854255 | 0.082 | 0.347 | 0.001285 | 06_mo-DC2 |
| MTRNR2L12 | -2.48575448 | 0.031 | 0.283 | 0.0013009 | 06_mo-DC2 |
| PKM | -1.14204424 | 0.449 | 0.692 | 0.0013678 | 06_mo-DC2 |
| EIF3L | -1.2951174 | 0.296 | 0.573 | 0.0014591 | 06_mo-DC2 |
| TGFBR2 | 1.23187879 | 0.48 | 0.249 | 0.0017841 | 06_mo-DC2 |
| RAB5C | -1.28318683 | 0.286 | 0.564 | 0.0018147 | 06_mo-DC2 |
| MAFF | -2.07167173 | 0.133 | 0.401 | 0.0018393 | 06_mo-DC2 |
| ABCC4 | 1.049686148 | 0.469 | 0.231 | 0.0019085 | 06_mo-DC2 |
| LIMK2 | 1.361069041 | 0.388 | 0.181 | 0.0022206 | 06_mo-DC2 |
| PPA1 | -1.43953416 | 0.388 | 0.625 | 0.0023924 | 06_mo-DC2 |

| | | | | | |
|---|---|---|---|---|---|
| BTBD9 | 1.242679044 | 0.561 | 0.359 | 0.0024495 | 06_mo-DC2 |
| MSN | -1.08542249 | 0.48 | 0.718 | 0.0024766 | 06_mo-DC2 |
| TNFAIP8 | -2.40752075 | 0.194 | 0.448 | 0.0024989 | 06_mo-DC2 |
| RHOF | -3.13967881 | 0.051 | 0.302 | 0.0025083 | 06_mo-DC2 |
| CNBP | -1.24136048 | 0.316 | 0.594 | 0.0025901 | 06_mo-DC2 |
| FGD2 | 1.125272232 | 0.449 | 0.236 | 0.0027181 | 06_mo-DC2 |
| PLCL2 | 1.059519248 | 0.357 | 0.156 | 0.0028323 | 06_mo-DC2 |
| RANBP9 | 1.192340913 | 0.52 | 0.306 | 0.0029146 | 06_mo-DC2 |
| CD1E | -12.9762 | 0 | 0.228 | 0.0029832 | 06_mo-DC2 |
| MOB1A | -1.67091155 | 0.235 | 0.497 | 0.0030182 | 06_mo-DC2 |
| IL1RAP | 1.079801436 | 0.51 | 0.269 | 0.0030681 | 06_mo-DC2 |
| DNAJA1 | -1.25064526 | 0.347 | 0.617 | 0.0033887 | 06_mo-DC2 |
| HK2 | 1.108722988 | 0.357 | 0.152 | 0.003653 | 06_mo-DC2 |
| ICAM1 | -1.89832782 | 0.224 | 0.484 | 0.0040239 | 06_mo-DC2 |
| MARCKSL1 | -4.88343817 | 0.02 | 0.253 | 0.0040514 | 06_mo-DC2 |
| GAA | 1.193175188 | 0.49 | 0.281 | 0.004322 | 06_mo-DC2 |
| B4GALT5 | -3.81073328 | 0.031 | 0.266 | 0.0043608 | 06_mo-DC2 |
| BID | -1.54424537 | 0.316 | 0.554 | 0.0048005 | 06_mo-DC2 |
| HSPA5 | -1.50879699 | 0.306 | 0.567 | 0.0051694 | 06_mo-DC2 |
| TFEC | 1.365846916 | 0.418 | 0.206 | 0.0054736 | 06_mo-DC2 |
| PCNX4 | 1.114975479 | 0.439 | 0.222 | 0.0058608 | 06_mo-DC2 |
| C3AR1 | 1.012367368 | 0.5 | 0.29 | 0.0067868 | 06_mo-DC2 |
| MYO1G | -2.72813813 | 0.071 | 0.316 | 0.0069584 | 06_mo-DC2 |
| TMEM59 | -1.65893057 | 0.194 | 0.441 | 0.0074326 | 06_mo-DC2 |
| EIF3F | -1.11332016 | 0.398 | 0.629 | 0.0075379 | 06_mo-DC2 |
| MGRN1 | 1.01490038 | 0.418 | 0.21 | 0.0081858 | 06_mo-DC2 |
| DIP2B | 1.071375174 | 0.439 | 0.221 | 0.0086229 | 06_mo-DC2 |
| CXCL2 | -13.8203909 | 0 | 0.214 | 0.0088031 | 06_mo-DC2 |
| EIF3E | -1.26559904 | 0.316 | 0.563 | 0.0092776 | 06_mo-DC2 |
| SMIM3 | -2.7526944 | 0.071 | 0.309 | 0.0095689 | 06_mo-DC2 |
| RARA | -1.91228921 | 0.122 | 0.379 | 0.0110858 | 06_mo-DC2 |
| PBX3 | 1.128830011 | 0.429 | 0.216 | 0.0115834 | 06_mo-DC2 |
| CALCRL | -13.3892772 | 0 | 0.21 | 0.0121187 | 06_mo-DC2 |
| CFP | -12.4071403 | 0 | 0.208 | 0.0142068 | 06_mo-DC2 |
| SKAP2 | 1.038162462 | 0.582 | 0.373 | 0.0152648 | 06_mo-DC2 |
| PID1 | -4.62533579 | 0.01 | 0.225 | 0.0155297 | 06_mo-DC2 |
| AC009093.2 | -2.47646875 | 0.092 | 0.335 | 0.018643 | 06_mo-DC2 |
| GDI2 | -1.29020148 | 0.286 | 0.539 | 0.0191 | 06_mo-DC2 |
| SDC2 | -3.2958664 | 0.031 | 0.253 | 0.0194336 | 06_mo-DC2 |
| HSPB1 | -2.81183685 | 0.194 | 0.43 | 0.0207119 | 06_mo-DC2 |
| MYH9 | -1.02750659 | 0.48 | 0.706 | 0.0216274 | 06_mo-DC2 |
| TUBA1A | -1.48806244 | 0.306 | 0.552 | 0.022195 | 06_mo-DC2 |
| ITSN1 | -2.42396526 | 0.071 | 0.305 | 0.0223543 | 06_mo-DC2 |
| PDIA6 | -2.13509075 | 0.122 | 0.35 | 0.0225755 | 06_mo-DC2 |
| SKP1 | -1.00224745 | 0.418 | 0.656 | 0.0225972 | 06_mo-DC2 |
| MBOAT7 | -2.80803572 | 0.051 | 0.277 | 0.0243072 | 06_mo-DC2 |

| | | | | | |
|---|---|---|---|---|---|
| LY6E | -1.98190016 | 0.153 | 0.4 | 0.0249431 | 06_mo-DC2 |
| SATB1 | -1.84877029 | 0.153 | 0.402 | 0.0254196 | 06_mo-DC2 |
| AL137857.1 | -3.17071067 | 0.051 | 0.27 | 0.0264347 | 06_mo-DC2 |
| HMGA1 | -1.56036358 | 0.163 | 0.412 | 0.0279804 | 06_mo-DC2 |
| SPIB | -2.05725068 | 0.102 | 0.342 | 0.0281704 | 06_mo-DC2 |
| RAB11FIP1 | -2.24998853 | 0.163 | 0.39 | 0.0288164 | 06_mo-DC2 |
| MOB3B | -2.25456123 | 0.051 | 0.285 | 0.0294782 | 06_mo-DC2 |
| FLT1 | -2.429267 | 0.112 | 0.347 | 0.0324879 | 06_mo-DC2 |
| PGK1 | -1.11689237 | 0.459 | 0.674 | 0.0332768 | 06_mo-DC2 |
| VDAC1 | -1.32857701 | 0.214 | 0.469 | 0.0372366 | 06_mo-DC2 |
| GYPC | -2.32056445 | 0.102 | 0.337 | 0.0379087 | 06_mo-DC2 |
| TNIP2 | -3.49702794 | 0.031 | 0.24 | 0.0385561 | 06_mo-DC2 |
| VDAC2 | -1.27893948 | 0.276 | 0.511 | 0.040012 | 06_mo-DC2 |
| PTPN2 | -1.20255617 | 0.255 | 0.512 | 0.0456027 | 06_mo-DC2 |
| ELOVL5 | -1.51609644 | 0.194 | 0.441 | 0.0463639 | 06_mo-DC2 |
| SUPT4H1 | -1.65784319 | 0.163 | 0.405 | 0.0468545 | 06_mo-DC2 |
| SLC16A3 | -1.05904378 | 0.357 | 0.59 | 0.0513619 | 06_mo-DC2 |
| RAN | -1.08138941 | 0.347 | 0.585 | 0.0538327 | 06_mo-DC2 |
| ZMIZ1 | -1.26178246 | 0.337 | 0.561 | 0.0539661 | 06_mo-DC2 |
| TBC1D8 | -1.44200522 | 0.276 | 0.513 | 0.0554664 | 06_mo-DC2 |
| HSPD1 | -1.71034486 | 0.316 | 0.547 | 0.0584965 | 06_mo-DC2 |
| GABARAPL2 | -1.12978802 | 0.327 | 0.56 | 0.068864 | 06_mo-DC2 |
| ACOT9 | -1.42614462 | 0.163 | 0.41 | 0.0693747 | 06_mo-DC2 |
| POLR2E | -1.49927683 | 0.214 | 0.442 | 0.0754291 | 06_mo-DC2 |
| ST13 | -1.39559357 | 0.224 | 0.464 | 0.0803263 | 06_mo-DC2 |
| CYTOR | -2.14781561 | 0.153 | 0.379 | 0.0838681 | 06_mo-DC2 |
| UPP1 | -1.28318166 | 0.214 | 0.449 | 0.0862802 | 06_mo-DC2 |
| COX7A2L | -1.85044923 | 0.163 | 0.374 | 0.0872695 | 06_mo-DC2 |
| PMAIP1 | -1.93859643 | 0.133 | 0.363 | 0.0924278 | 06_mo-DC2 |
| USP12 | -2.42433219 | 0.122 | 0.329 | 0.0953877 | 06_mo-DC2 |
| DDAH2 | -1.24802892 | 0.235 | 0.47 | 0.0961235 | 06_mo-DC2 |
| VMO1 | -2.11424339 | 0.235 | 0.44 | 0.1023084 | 06_mo-DC2 |
| EIF4B | -1.19160928 | 0.316 | 0.542 | 0.1029675 | 06_mo-DC2 |
| BANP | -1.399522 | 0.173 | 0.414 | 0.1199026 | 06_mo-DC2 |
| GPBP1 | -1.18213104 | 0.337 | 0.565 | 0.1209717 | 06_mo-DC2 |
| FRY | -2.65087035 | 0.061 | 0.273 | 0.1282038 | 06_mo-DC2 |
| NFKB2 | -1.49880596 | 0.133 | 0.365 | 0.1397324 | 06_mo-DC2 |
| H2AFV | -1.32630447 | 0.286 | 0.498 | 0.1472847 | 06_mo-DC2 |
| AC007384.1 | -1.45683853 | 0.163 | 0.407 | 0.1567569 | 06_mo-DC2 |
| ARHGAP31 | -1.62788136 | 0.235 | 0.441 | 0.1673986 | 06_mo-DC2 |
| UBE2I | -1.22871792 | 0.163 | 0.419 | 0.1699069 | 06_mo-DC2 |
| TPP1 | -1.20085838 | 0.255 | 0.488 | 0.1758923 | 06_mo-DC2 |
| ACTR3 | -1.03306863 | 0.357 | 0.593 | 0.1815129 | 06_mo-DC2 |
| BCOR | -2.29835575 | 0.092 | 0.302 | 0.1838128 | 06_mo-DC2 |
| RAPGEF6 | -1.27299234 | 0.276 | 0.501 | 0.2142214 | 06_mo-DC2 |
| MAP2K3 | -1.69053962 | 0.204 | 0.408 | 0.2236519 | 06_mo-DC2 |

| | | | | | |
|---|---|---|---|---|---|
| PPDPF | -1.03400378 | 0.398 | 0.599 | 0.2275891 | 06_mo-DC2 |
| PDCL3 | -2.09852065 | 0.102 | 0.307 | 0.2655785 | 06_mo-DC2 |
| ACTN4 | -1.59710848 | 0.153 | 0.369 | 0.2749562 | 06_mo-DC2 |
| GLIPR1 | -1.09318181 | 0.347 | 0.56 | 0.2806009 | 06_mo-DC2 |
| GNA15 | -1.24581937 | 0.276 | 0.484 | 0.3112235 | 06_mo-DC2 |
| AUTS2 | -1.86447483 | 0.173 | 0.388 | 0.3301539 | 06_mo-DC2 |
| RNASE6 | -1.20970486 | 0.378 | 0.58 | 0.3601466 | 06_mo-DC2 |
| RELB | -1.29106094 | 0.224 | 0.439 | 0.3677143 | 06_mo-DC2 |
| APOO | -2.14877828 | 0.143 | 0.345 | 0.3682214 | 06_mo-DC2 |
| SEC14L1 | -1.05636085 | 0.388 | 0.595 | 0.4115015 | 06_mo-DC2 |
| CSTB | -1.07929058 | 0.388 | 0.607 | 0.4163541 | 06_mo-DC2 |
| FKBP1A | -1.15204869 | 0.306 | 0.508 | 0.4166783 | 06_mo-DC2 |
| JAML | -1.48688962 | 0.173 | 0.38 | 0.42178 | 06_mo-DC2 |
| PPT1 | -1.07129673 | 0.327 | 0.561 | 0.4313146 | 06_mo-DC2 |
| KCTD20 | -1.15977436 | 0.133 | 0.36 | 0.5001291 | 06_mo-DC2 |
| SLC41A2 | -2.05152758 | 0.092 | 0.293 | 0.5726494 | 06_mo-DC2 |
| EIF4E | -1.49302513 | 0.245 | 0.456 | 0.6022372 | 06_mo-DC2 |
| TMEM123 | -1.18552421 | 0.245 | 0.456 | 0.667274 | 06_mo-DC2 |
| UGCG | -1.55671793 | 0.173 | 0.376 | 0.7364431 | 06_mo-DC2 |
| STK10 | -1.25075269 | 0.184 | 0.396 | 0.8962972 | 06_mo-DC2 |
| RBM17 | -1.21514498 | 0.163 | 0.383 | 1 | 06_mo-DC2 |
| NECTIN2 | -1.06030889 | 0.255 | 0.463 | 1 | 06_mo-DC2 |
| OXSR1 | -1.05870911 | 0.204 | 0.424 | 1 | 06_mo-DC2 |
| EMILIN2 | -1.14222792 | 0.214 | 0.419 | 1 | 06_mo-DC2 |
| PEA15 | -1.14731511 | 0.204 | 0.407 | 1 | 06_mo-DC2 |
| CLNK | 5.640041735 | 0.758 | 0.03 | 3.64E-145 | 07_cDC1 |
| DNASE1L3 | 3.31291578 | 0.549 | 0.019 | 1.67E-103 | 07_cDC1 |
| RAB7B | 4.338488178 | 0.802 | 0.072 | 2.19E-99 | 07_cDC1 |
| IDO1 | 4.131365115 | 0.769 | 0.065 | 1.56E-97 | 07_cDC1 |
| NCALD | 5.678359012 | 0.418 | 0.009 | 3.91E-93 | 07_cDC1 |
| TOX | 4.052856929 | 0.626 | 0.046 | 3.14E-83 | 07_cDC1 |
| DYSF | 5.79195583 | 0.374 | 0.008 | 4.56E-82 | 07_cDC1 |
| AC099560.1 | 6.588213908 | 0.396 | 0.013 | 7.77E-78 | 07_cDC1 |
| SLC24A4 | 5.679292521 | 0.407 | 0.015 | 5.07E-76 | 07_cDC1 |
| TACSTD2 | 5.485901046 | 0.462 | 0.023 | 4.52E-75 | 07_cDC1 |
| C1orf21 | 5.127373497 | 0.451 | 0.024 | 4.19E-70 | 07_cDC1 |
| XCR1 | 3.901412952 | 0.275 | 0.004 | 1.69E-67 | 07_cDC1 |
| CPNE3 | 3.769926324 | 0.912 | 0.219 | 2.16E-64 | 07_cDC1 |
| CLEC9A | 3.420210125 | 0.692 | 0.093 | 9.00E-63 | 07_cDC1 |
| WDFY4 | 3.845511304 | 0.945 | 0.27 | 2.04E-61 | 07_cDC1 |
| RUBCNL | 3.986292602 | 0.67 | 0.096 | 1.57E-57 | 07_cDC1 |
| VAC14 | 3.4527829 | 0.659 | 0.089 | 5.98E-57 | 07_cDC1 |
| NEGR1 | 4.443146554 | 0.549 | 0.058 | 1.31E-55 | 07_cDC1 |
| THRB | 4.358754508 | 0.593 | 0.07 | 5.53E-55 | 07_cDC1 |
| CADM1 | 3.085912803 | 0.835 | 0.173 | 1.06E-53 | 07_cDC1 |
| SLAMF7 | 3.732852946 | 0.505 | 0.051 | 1.24E-52 | 07_cDC1 |

| | | | | | |
|---|---|---|---|---|---|
| GNAO1 | 3.757010139 | 0.352 | 0.019 | 2.89E-52 | 07_cDC1 |
| LGALS2 | 3.474872438 | 0.571 | 0.072 | 3.61E-52 | 07_cDC1 |
| PPM1H | 4.557541924 | 0.374 | 0.024 | 1.72E-51 | 07_cDC1 |
| RHEX | 4.47625196 | 0.363 | 0.023 | 2.02E-49 | 07_cDC1 |
| TSPAN13 | 4.436806646 | 0.308 | 0.015 | 4.37E-47 | 07_cDC1 |
| C1orf54 | 3.003079728 | 0.802 | 0.23 | 3.09E-42 | 07_cDC1 |
| PLPP3 | 5.338622261 | 0.209 | 0.006 | 1.82E-41 | 07_cDC1 |
| BTLA | 5.036879339 | 0.253 | 0.013 | 9.63E-39 | 07_cDC1 |
| TYROBP | -3.3983979 | 0.473 | 0.933 | 1.31E-38 | 07_cDC1 |
| DPP4 | 3.480201572 | 0.209 | 0.007 | 1.46E-37 | 07_cDC1 |
| MIR924HG | 4.411409227 | 0.407 | 0.046 | 3.52E-37 | 07_cDC1 |
| CYYR1 | 3.900473921 | 0.286 | 0.019 | 7.87E-37 | 07_cDC1 |
| EGLN3 | 2.994596576 | 0.418 | 0.048 | 1.53E-36 | 07_cDC1 |
| HABP4 | 3.617251483 | 0.33 | 0.028 | 4.04E-36 | 07_cDC1 |
| PTK2 | 2.772380318 | 0.582 | 0.107 | 7.38E-35 | 07_cDC1 |
| FCER1G | -3.19320989 | 0.363 | 0.895 | 3.96E-33 | 07_cDC1 |
| NLRC5 | 3.40751255 | 0.495 | 0.086 | 5.04E-32 | 07_cDC1 |
| CPVL | 2.302897719 | 0.945 | 0.63 | 1.56E-31 | 07_cDC1 |
| TMEM14A | 3.726354225 | 0.341 | 0.037 | 4.12E-31 | 07_cDC1 |
| BCL2L11 | 2.540888396 | 0.747 | 0.218 | 3.34E-30 | 07_cDC1 |
| HLA-DOB | 2.672121188 | 0.505 | 0.094 | 9.05E-30 | 07_cDC1 |
| DBN1 | 3.375776854 | 0.253 | 0.02 | 8.70E-29 | 07_cDC1 |
| PNMA1 | 3.154034141 | 0.385 | 0.051 | 1.01E-28 | 07_cDC1 |
| CAMK2D | 2.767211502 | 0.692 | 0.196 | 3.50E-28 | 07_cDC1 |
| CCDC126 | 3.620064945 | 0.341 | 0.041 | 3.63E-28 | 07_cDC1 |
| ZEB2 | -5.4078942 | 0.066 | 0.761 | 7.66E-28 | 07_cDC1 |
| FCGR2A | -5.27068236 | 0.033 | 0.75 | 8.94E-28 | 07_cDC1 |
| ALOX5AP | -5.15848007 | 0.077 | 0.758 | 1.85E-27 | 07_cDC1 |
| BX664727.3 | 3.605513431 | 0.385 | 0.055 | 2.22E-27 | 07_cDC1 |
| ENOX1 | 2.760130734 | 0.242 | 0.019 | 9.78E-27 | 07_cDC1 |
| LDLRAD3 | 3.000372331 | 0.681 | 0.197 | 1.18E-26 | 07_cDC1 |
| ATP6V0A2 | 2.791893952 | 0.516 | 0.11 | 1.17E-25 | 07_cDC1 |
| FGD6 | 3.332000474 | 0.253 | 0.022 | 1.62E-25 | 07_cDC1 |
| CCSER1 | 3.072198885 | 0.901 | 0.434 | 2.41E-25 | 07_cDC1 |
| HSH2D | 2.870971519 | 0.407 | 0.067 | 3.29E-25 | 07_cDC1 |
| DOCK4 | -4.36247917 | 0.264 | 0.814 | 3.43E-25 | 07_cDC1 |
| AIM2 | 3.854492572 | 0.242 | 0.021 | 4.54E-25 | 07_cDC1 |
| IRAK3 | -4.55085699 | 0.055 | 0.739 | 4.74E-25 | 07_cDC1 |
| RNF130 | -3.87005553 | 0.077 | 0.75 | 1.09E-24 | 07_cDC1 |
| ZEB1 | 2.582334107 | 0.593 | 0.149 | 3.92E-24 | 07_cDC1 |
| RAB30 | 3.405967177 | 0.264 | 0.027 | 1.65E-23 | 07_cDC1 |
| CTSB | -2.54285704 | 0.429 | 0.859 | 2.71E-23 | 07_cDC1 |
| HDAC9 | 2.583601688 | 0.901 | 0.456 | 2.90E-23 | 07_cDC1 |
| NAAA | 2.772362518 | 0.626 | 0.211 | 4.89E-23 | 07_cDC1 |
| ECE1 | 3.121654096 | 0.33 | 0.047 | 5.28E-23 | 07_cDC1 |
| BATF3 | 2.594035032 | 0.396 | 0.067 | 8.11E-23 | 07_cDC1 |

| | | | | | |
|---|---|---|---|---|---|
| SAMSN1 | -2.74366214 | 0.253 | 0.833 | 1.23E-22 | 07_cDC1 |
| SNX3 | 2.708984495 | 0.824 | 0.582 | 2.41E-22 | 07_cDC1 |
| TLR2 | -3.98251556 | 0.099 | 0.72 | 3.63E-22 | 07_cDC1 |
| MPEG1 | 2.173554734 | 0.473 | 0.102 | 4.21E-22 | 07_cDC1 |
| PLXDC2 | -2.25027048 | 0.484 | 0.919 | 1.21E-21 | 07_cDC1 |
| KIF16B | 2.394086773 | 0.725 | 0.254 | 3.54E-21 | 07_cDC1 |
| FPR1 | -3.87661803 | 0.099 | 0.702 | 4.17E-21 | 07_cDC1 |
| LRRCC1 | 2.997645991 | 0.264 | 0.032 | 1.14E-20 | 07_cDC1 |
| ABL1 | 3.022728502 | 0.637 | 0.209 | 1.65E-20 | 07_cDC1 |
| DENND1B | 2.052582689 | 0.813 | 0.357 | 4.20E-20 | 07_cDC1 |
| UPF2 | 2.43096544 | 0.659 | 0.233 | 6.83E-20 | 07_cDC1 |
| MYO9A | 2.308331629 | 0.473 | 0.11 | 8.89E-20 | 07_cDC1 |
| KAT2B | 2.412931532 | 0.604 | 0.193 | 1.62E-19 | 07_cDC1 |
| ASAP1 | 1.836449457 | 0.89 | 0.516 | 1.66E-19 | 07_cDC1 |
| NET1 | 1.92409449 | 0.67 | 0.228 | 1.80E-19 | 07_cDC1 |
| FLNB | 2.965115535 | 0.308 | 0.047 | 2.62E-19 | 07_cDC1 |
| CSF1R | -5.54976544 | 0.011 | 0.614 | 3.62E-19 | 07_cDC1 |
| RAB11FIP1 | 2.318273452 | 0.769 | 0.35 | 5.13E-19 | 07_cDC1 |
| ALOX5 | -4.17663374 | 0.077 | 0.652 | 5.65E-19 | 07_cDC1 |
| FCGR2B | -8.22623086 | 0.011 | 0.593 | 1.41E-18 | 07_cDC1 |
| ACTN1 | 2.433638124 | 0.571 | 0.181 | 1.55E-18 | 07_cDC1 |
| TAP1 | 2.002446876 | 0.626 | 0.219 | 1.98E-18 | 07_cDC1 |
| GLUL | -2.6222561 | 0.275 | 0.758 | 2.01E-18 | 07_cDC1 |
| ZNF250 | 2.508638116 | 0.374 | 0.074 | 2.68E-18 | 07_cDC1 |
| ITM2B | -1.65040163 | 0.714 | 0.918 | 2.98E-18 | 07_cDC1 |
| NUBPL | 2.501941514 | 0.396 | 0.082 | 4.70E-18 | 07_cDC1 |
| OLR1 | -3.71918429 | 0.176 | 0.695 | 4.84E-18 | 07_cDC1 |
| GPR157 | 2.260853517 | 0.659 | 0.238 | 1.07E-17 | 07_cDC1 |
| TXN | 2.713582221 | 0.802 | 0.431 | 1.69E-17 | 07_cDC1 |
| DMD | 3.012732241 | 0.275 | 0.04 | 3.16E-17 | 07_cDC1 |
| CCDC6 | 2.116353202 | 0.681 | 0.269 | 9.37E-17 | 07_cDC1 |
| CAPG | -2.96951729 | 0.209 | 0.691 | 1.19E-16 | 07_cDC1 |
| AC007991.3 | 2.98760347 | 0.253 | 0.035 | 1.21E-16 | 07_cDC1 |
| FLT3 | 1.954792959 | 0.681 | 0.264 | 1.37E-16 | 07_cDC1 |
| REEP3 | 1.767070547 | 0.67 | 0.254 | 1.90E-16 | 07_cDC1 |
| SEMA7A | 2.23069916 | 0.473 | 0.125 | 2.20E-16 | 07_cDC1 |
| SAMD12 | 3.233162412 | 0.231 | 0.03 | 5.08E-16 | 07_cDC1 |
| FRY | 3.057725589 | 0.626 | 0.235 | 6.56E-16 | 07_cDC1 |
| PLD4 | -5.46497849 | 0.022 | 0.559 | 9.77E-16 | 07_cDC1 |
| KYNU | -3.73238013 | 0.143 | 0.649 | 1.05E-15 | 07_cDC1 |
| S100A4 | -2.19822615 | 0.286 | 0.736 | 1.24E-15 | 07_cDC1 |
| SERPINF2 | 2.684209314 | 0.385 | 0.09 | 1.39E-15 | 07_cDC1 |
| CLIC2 | 1.815011581 | 0.637 | 0.231 | 1.69E-15 | 07_cDC1 |
| HIF1A | -2.12919507 | 0.374 | 0.806 | 2.51E-15 | 07_cDC1 |
| SHTN1 | 2.045436776 | 0.714 | 0.326 | 3.31E-15 | 07_cDC1 |
| BTG1 | -1.57212251 | 0.637 | 0.902 | 3.76E-15 | 07_cDC1 |

| | | | | | |
|---|---|---|---|---|---|
| DEF8 | 2.302656863 | 0.352 | 0.073 | 4.42E-15 | 07_cDC1 |
| BCL2 | -3.02068359 | 0.187 | 0.683 | 5.02E-15 | 07_cDC1 |
| CCDC26 | 2.571138589 | 0.451 | 0.122 | 1.01E-14 | 07_cDC1 |
| IL13RA1 | -3.13935105 | 0.154 | 0.639 | 1.04E-14 | 07_cDC1 |
| LYRM4 | 2.437559343 | 0.407 | 0.103 | 1.10E-14 | 07_cDC1 |
| ATR | 2.151375897 | 0.516 | 0.16 | 1.11E-14 | 07_cDC1 |
| IER3 | -2.79615124 | 0.286 | 0.723 | 1.14E-14 | 07_cDC1 |
| ZNF516 | 2.868545387 | 0.516 | 0.173 | 1.26E-14 | 07_cDC1 |
| CEBPB | -2.86621966 | 0.231 | 0.681 | 1.77E-14 | 07_cDC1 |
| MS4A7 | -2.18026915 | 0.33 | 0.761 | 2.30E-14 | 07_cDC1 |
| BCL2L14 | 2.976547778 | 0.242 | 0.037 | 2.45E-14 | 07_cDC1 |
| IFITM3 | -2.24062447 | 0.308 | 0.756 | 2.80E-14 | 07_cDC1 |
| FMN1 | -3.39096504 | 0.11 | 0.618 | 4.56E-14 | 07_cDC1 |
| FCER1A | -15.0105732 | 0 | 0.502 | 7.45E-14 | 07_cDC1 |
| NCKAP5 | 2.472687692 | 0.429 | 0.115 | 9.27E-14 | 07_cDC1 |
| DAPP1 | 1.457320644 | 0.703 | 0.264 | 1.02E-13 | 07_cDC1 |
| SULF2 | 1.46939979 | 0.714 | 0.306 | 1.02E-13 | 07_cDC1 |
| S100B | 2.933282933 | 0.418 | 0.119 | 1.36E-13 | 07_cDC1 |
| KIAA1958 | 2.399122925 | 0.352 | 0.077 | 1.50E-13 | 07_cDC1 |
| MSR1 | -4.05715491 | 0.077 | 0.565 | 1.97E-13 | 07_cDC1 |
| RHBDF2 | -3.87917691 | 0.088 | 0.568 | 2.03E-13 | 07_cDC1 |
| CCND1 | 2.25664352 | 0.451 | 0.134 | 2.10E-13 | 07_cDC1 |
| IRF8 | 1.957101232 | 0.736 | 0.382 | 2.73E-13 | 07_cDC1 |
| OXSR1 | 1.983323303 | 0.769 | 0.387 | 4.46E-13 | 07_cDC1 |
| TAP2 | 1.797812884 | 0.593 | 0.228 | 5.38E-13 | 07_cDC1 |
| PTPN22 | 2.359923894 | 0.505 | 0.163 | 7.46E-13 | 07_cDC1 |
| SIPA1L3 | 2.257910495 | 0.725 | 0.379 | 7.67E-13 | 07_cDC1 |
| APBA1 | 2.476927871 | 0.275 | 0.051 | 8.32E-13 | 07_cDC1 |
| PLPP1 | 2.532748511 | 0.352 | 0.084 | 1.80E-12 | 07_cDC1 |
| P2RY14 | 2.302589157 | 0.341 | 0.079 | 2.19E-12 | 07_cDC1 |
| ARHGAP26 | -2.22542768 | 0.198 | 0.673 | 2.82E-12 | 07_cDC1 |
| TMEM50B | 2.117096377 | 0.538 | 0.194 | 2.84E-12 | 07_cDC1 |
| NBEAL2 | 2.673686378 | 0.385 | 0.103 | 3.07E-12 | 07_cDC1 |
| RABGAP1L | 1.775173895 | 0.725 | 0.343 | 3.22E-12 | 07_cDC1 |
| FNBP1 | 1.503715623 | 0.868 | 0.58 | 7.81E-12 | 07_cDC1 |
| VEGFA | -2.90696662 | 0.11 | 0.578 | 1.50E-11 | 07_cDC1 |
| NCOA7 | 1.899200784 | 0.44 | 0.136 | 1.84E-11 | 07_cDC1 |
| FARS2 | 2.180781974 | 0.429 | 0.134 | 1.88E-11 | 07_cDC1 |
| ANPEP | 1.855288253 | 0.385 | 0.104 | 3.14E-11 | 07_cDC1 |
| SORL1 | -4.68895642 | 0.044 | 0.49 | 3.18E-11 | 07_cDC1 |
| CYFIP2 | 2.380244187 | 0.275 | 0.056 | 3.24E-11 | 07_cDC1 |
| PLEKHM3 | 2.562708363 | 0.286 | 0.061 | 3.36E-11 | 07_cDC1 |
| MZT2A | 1.674034721 | 0.615 | 0.271 | 3.54E-11 | 07_cDC1 |
| SLC11A1 | -5.66569821 | 0.044 | 0.481 | 4.77E-11 | 07_cDC1 |
| DST | 1.659156776 | 0.78 | 0.42 | 5.36E-11 | 07_cDC1 |
| ITGB7 | 2.34877891 | 0.275 | 0.058 | 5.79E-11 | 07_cDC1 |

| | | | | | |
|---|---|---|---|---|---|
| TRERF1 | 2.694313614 | 0.363 | 0.098 | 7.45E-11 | 07_cDC1 |
| BCR | 2.368180068 | 0.363 | 0.095 | 7.68E-11 | 07_cDC1 |
| ERN1 | 2.413602393 | 0.516 | 0.193 | 8.04E-11 | 07_cDC1 |
| SRGAP1 | -2.60138647 | 0.143 | 0.595 | 1.03E-10 | 07_cDC1 |
| OGFRL1 | -2.01280265 | 0.286 | 0.694 | 1.89E-10 | 07_cDC1 |
| MYO1D | 2.110599431 | 0.429 | 0.136 | 2.01E-10 | 07_cDC1 |
| CD163 | -3.71798018 | 0.088 | 0.521 | 2.16E-10 | 07_cDC1 |
| RHOB | -3.37678958 | 0.264 | 0.63 | 2.37E-10 | 07_cDC1 |
| LILRB4 | -4.48859252 | 0.033 | 0.462 | 3.18E-10 | 07_cDC1 |
| SLAMF8 | 1.691005154 | 0.505 | 0.186 | 3.95E-10 | 07_cDC1 |
| FOXO3 | -5.45110719 | 0.033 | 0.451 | 4.47E-10 | 07_cDC1 |
| VDR | 2.420470671 | 0.615 | 0.274 | 5.74E-10 | 07_cDC1 |
| C3 | -4.19137058 | 0.154 | 0.54 | 5.95E-10 | 07_cDC1 |
| CD226 | 2.14566577 | 0.527 | 0.198 | 6.00E-10 | 07_cDC1 |
| GPR183 | -1.61863123 | 0.571 | 0.824 | 6.49E-10 | 07_cDC1 |
| CYP2S1 | 1.699225993 | 0.637 | 0.285 | 7.76E-10 | 07_cDC1 |
| PFKFB3 | 1.359936877 | 0.901 | 0.692 | 8.21E-10 | 07_cDC1 |
| ATG3 | 1.549293419 | 0.648 | 0.344 | 1.06E-09 | 07_cDC1 |
| ATP6V1B2 | -3.83576836 | 0.066 | 0.482 | 1.13E-09 | 07_cDC1 |
| PSTPIP2 | -5.56793481 | 0.011 | 0.429 | 1.27E-09 | 07_cDC1 |
| VAV3 | 1.892841387 | 0.56 | 0.228 | 1.28E-09 | 07_cDC1 |
| PTK2B | -2.95500023 | 0.11 | 0.539 | 1.33E-09 | 07_cDC1 |
| IL1B | -2.72527061 | 0.396 | 0.729 | 1.35E-09 | 07_cDC1 |
| C12orf45 | 2.082246649 | 0.495 | 0.187 | 1.52E-09 | 07_cDC1 |
| DPYD | -2.00882815 | 0.22 | 0.632 | 1.59E-09 | 07_cDC1 |
| SLC1A3 | -3.58499563 | 0.143 | 0.55 | 2.52E-09 | 07_cDC1 |
| AUTS2 | 2.425394592 | 0.681 | 0.355 | 2.55E-09 | 07_cDC1 |
| CTSH | -1.82876485 | 0.319 | 0.688 | 3.10E-09 | 07_cDC1 |
| C1QC | -4.53714015 | 0.209 | 0.568 | 3.34E-09 | 07_cDC1 |
| LST1 | -1.83662516 | 0.209 | 0.597 | 3.38E-09 | 07_cDC1 |
| B3GNT5 | -2.23153832 | 0.187 | 0.607 | 3.46E-09 | 07_cDC1 |
| ACSL1 | -1.71859931 | 0.571 | 0.83 | 3.58E-09 | 07_cDC1 |
| NME4 | 1.61677998 | 0.505 | 0.194 | 3.84E-09 | 07_cDC1 |
| GSN | -1.24548441 | 0.538 | 0.805 | 3.86E-09 | 07_cDC1 |
| ELOVL5 | 1.597867056 | 0.758 | 0.403 | 4.21E-09 | 07_cDC1 |
| MS4A4A | -5.41770628 | 0.022 | 0.424 | 4.24E-09 | 07_cDC1 |
| CYB5R3 | 1.53271708 | 0.637 | 0.337 | 5.01E-09 | 07_cDC1 |
| HCST | -2.18209375 | 0.253 | 0.621 | 5.05E-09 | 07_cDC1 |
| FARP2 | 1.975347397 | 0.505 | 0.197 | 5.43E-09 | 07_cDC1 |
| LRMDA | -1.96018194 | 0.363 | 0.721 | 7.71E-09 | 07_cDC1 |
| ZBTB46 | 1.894741893 | 0.659 | 0.335 | 8.88E-09 | 07_cDC1 |
| SFT2D2 | 1.807149017 | 0.516 | 0.207 | 1.02E-08 | 07_cDC1 |
| CLEC10A | -13.8633046 | 0 | 0.391 | 1.02E-08 | 07_cDC1 |
| LPXN | -4.1744687 | 0.066 | 0.453 | 1.03E-08 | 07_cDC1 |
| UCP2 | 1.794935972 | 0.681 | 0.429 | 1.38E-08 | 07_cDC1 |
| MAP3K1 | 2.362989957 | 0.462 | 0.174 | 1.48E-08 | 07_cDC1 |

| | | | | | |
|---|---|---|---|---|---|
| MS4A6A | -1.43569568 | 0.407 | 0.755 | 1.48E-08 | 07_cDC1 |
| MIR4435-2HG | 1.386800741 | 0.626 | 0.275 | 1.75E-08 | 07_cDC1 |
| RAB31 | -1.39785232 | 0.67 | 0.885 | 1.90E-08 | 07_cDC1 |
| FILIP1L | -1.6015966 | 0.242 | 0.647 | 2.53E-08 | 07_cDC1 |
| SMCHD1 | 1.448943419 | 0.857 | 0.55 | 2.90E-08 | 07_cDC1 |
| IGSF6 | -4.08803846 | 0.044 | 0.429 | 2.95E-08 | 07_cDC1 |
| IDO2 | 2.481667513 | 0.286 | 0.074 | 3.19E-08 | 07_cDC1 |
| SLC25A29 | 2.052846758 | 0.319 | 0.089 | 3.38E-08 | 07_cDC1 |
| CD14 | -3.93025242 | 0.088 | 0.463 | 4.02E-08 | 07_cDC1 |
| LAIR1 | -3.61346766 | 0.055 | 0.443 | 4.16E-08 | 07_cDC1 |
| GAS7 | 2.555040543 | 0.473 | 0.191 | 4.42E-08 | 07_cDC1 |
| RNF213 | -3.62477237 | 0.077 | 0.461 | 4.63E-08 | 07_cDC1 |
| DUSP4 | 1.148592849 | 0.846 | 0.523 | 4.65E-08 | 07_cDC1 |
| SMCO4 | 1.405907504 | 0.505 | 0.197 | 4.82E-08 | 07_cDC1 |
| ARHGAP18 | -3.33662666 | 0.088 | 0.47 | 4.91E-08 | 07_cDC1 |
| PACS1 | 1.722631219 | 0.659 | 0.333 | 4.93E-08 | 07_cDC1 |
| PLEKHA5 | 1.98550965 | 0.571 | 0.251 | 5.07E-08 | 07_cDC1 |
| NHSL1 | -3.52300233 | 0.077 | 0.471 | 5.33E-08 | 07_cDC1 |
| HBEGF | -3.66701564 | 0.077 | 0.457 | 5.91E-08 | 07_cDC1 |
| MCOLN2 | 1.245876299 | 0.681 | 0.345 | 6.27E-08 | 07_cDC1 |
| C1QB | -3.93032647 | 0.242 | 0.564 | 6.61E-08 | 07_cDC1 |
| MT2A | -2.88550184 | 0.396 | 0.679 | 7.20E-08 | 07_cDC1 |
| TRAF4 | 1.821352284 | 0.319 | 0.091 | 1.04E-07 | 07_cDC1 |
| ETS2 | -1.60299977 | 0.396 | 0.725 | 1.05E-07 | 07_cDC1 |
| MARCKS | -2.53458793 | 0.198 | 0.56 | 1.08E-07 | 07_cDC1 |
| SLC7A5 | -2.03955183 | 0.297 | 0.632 | 1.18E-07 | 07_cDC1 |
| FUT8 | 1.869440413 | 0.319 | 0.092 | 1.28E-07 | 07_cDC1 |
| IL18 | -2.49868796 | 0.187 | 0.543 | 1.42E-07 | 07_cDC1 |
| LSM6 | 2.034415816 | 0.396 | 0.143 | 1.60E-07 | 07_cDC1 |
| TRPS1 | 1.996150543 | 0.791 | 0.535 | 1.66E-07 | 07_cDC1 |
| CD1C | -14.1284682 | 0 | 0.359 | 1.97E-07 | 07_cDC1 |
| CYBB | -2.80297126 | 0.11 | 0.483 | 2.12E-07 | 07_cDC1 |
| SDCBP | -1.53405452 | 0.429 | 0.733 | 2.13E-07 | 07_cDC1 |
| RUNX1 | 1.749558194 | 0.846 | 0.576 | 2.35E-07 | 07_cDC1 |
| CALCRL | 2.560602442 | 0.462 | 0.179 | 2.75E-07 | 07_cDC1 |
| MVB12B | 1.777764052 | 0.407 | 0.147 | 2.93E-07 | 07_cDC1 |
| CLEC5A | -6.41069867 | 0.011 | 0.365 | 3.22E-07 | 07_cDC1 |
| CD44 | -1.90973086 | 0.418 | 0.708 | 3.66E-07 | 07_cDC1 |
| FYB1 | -2.43108506 | 0.154 | 0.523 | 3.77E-07 | 07_cDC1 |
| TPMT | 1.669799558 | 0.33 | 0.1 | 3.87E-07 | 07_cDC1 |
| PSMB9 | 1.396560988 | 0.604 | 0.319 | 4.76E-07 | 07_cDC1 |
| VSIG4 | -3.72319342 | 0.055 | 0.413 | 5.48E-07 | 07_cDC1 |
| LINC02245 | 1.604101772 | 0.495 | 0.204 | 5.61E-07 | 07_cDC1 |
| TRIM22 | 1.259988095 | 0.681 | 0.387 | 5.99E-07 | 07_cDC1 |
| ATP1A1 | 1.214860402 | 0.791 | 0.557 | 6.56E-07 | 07_cDC1 |
| IGF2R | 1.344586445 | 0.396 | 0.133 | 6.63E-07 | 07_cDC1 |

| | | | | | |
|---|---|---|---|---|---|
| TNFRSF1B | -1.41063517 | 0.407 | 0.71 | 9.02E-07 | 07_cDC1 |
| APAF1 | 1.625220973 | 0.571 | 0.279 | 9.32E-07 | 07_cDC1 |
| PIM2 | 1.41578008 | 0.374 | 0.126 | 9.44E-07 | 07_cDC1 |
| DENND3 | -3.19530881 | 0.077 | 0.437 | 9.58E-07 | 07_cDC1 |
| TREM2 | -4.02556706 | 0.055 | 0.405 | 9.75E-07 | 07_cDC1 |
| CNN2 | 1.119338892 | 0.725 | 0.436 | 1.13E-06 | 07_cDC1 |
| SIRPA | -3.60915299 | 0.022 | 0.377 | 1.13E-06 | 07_cDC1 |
| SYTL3 | -1.22057142 | 0.44 | 0.747 | 1.19E-06 | 07_cDC1 |
| RTN1 | -5.21591444 | 0.022 | 0.363 | 1.49E-06 | 07_cDC1 |
| ICAM3 | 1.407884212 | 0.549 | 0.278 | 2.00E-06 | 07_cDC1 |
| RAB8B | 1.239172426 | 0.78 | 0.46 | 2.38E-06 | 07_cDC1 |
| SERPINA1 | -1.83949731 | 0.319 | 0.639 | 2.56E-06 | 07_cDC1 |
| OAZ2 | 1.861981838 | 0.352 | 0.123 | 3.03E-06 | 07_cDC1 |
| ELL2 | -1.58220188 | 0.484 | 0.758 | 3.13E-06 | 07_cDC1 |
| SFMBT2 | -2.95211413 | 0.121 | 0.472 | 3.60E-06 | 07_cDC1 |
| BACH1 | 1.458826817 | 0.747 | 0.422 | 3.65E-06 | 07_cDC1 |
| ACER3 | 1.346569838 | 0.714 | 0.395 | 3.86E-06 | 07_cDC1 |
| STIM2 | 1.763771105 | 0.484 | 0.207 | 4.16E-06 | 07_cDC1 |
| PEA15 | -2.83395457 | 0.077 | 0.415 | 4.52E-06 | 07_cDC1 |
| AKAP6 | 2.770467656 | 0.308 | 0.098 | 5.12E-06 | 07_cDC1 |
| ATP8B4 | -2.52589432 | 0.132 | 0.481 | 5.19E-06 | 07_cDC1 |
| FCGR3A | -4.71105267 | 0.033 | 0.361 | 5.64E-06 | 07_cDC1 |
| PSTPIP1 | 1.987521351 | 0.33 | 0.112 | 5.78E-06 | 07_cDC1 |
| FPR3 | -3.53000728 | 0.033 | 0.371 | 6.06E-06 | 07_cDC1 |
| C5AR1 | -2.94862234 | 0.055 | 0.398 | 7.23E-06 | 07_cDC1 |
| SH3BP5 | -2.57820519 | 0.154 | 0.479 | 7.98E-06 | 07_cDC1 |
| C1QA | -3.9093335 | 0.253 | 0.522 | 8.42E-06 | 07_cDC1 |
| FMNL3 | -4.35636797 | 0.033 | 0.356 | 1.03E-05 | 07_cDC1 |
| IRS2 | -2.94426147 | 0.055 | 0.39 | 1.15E-05 | 07_cDC1 |
| TSPO | -1.67348773 | 0.341 | 0.621 | 1.50E-05 | 07_cDC1 |
| CD300A | -4.12282105 | 0.044 | 0.358 | 1.57E-05 | 07_cDC1 |
| CSF3R | -1.92041685 | 0.231 | 0.553 | 1.59E-05 | 07_cDC1 |
| SLC41A2 | 1.407873726 | 0.549 | 0.263 | 1.76E-05 | 07_cDC1 |
| NFATC2 | 1.539599409 | 0.495 | 0.231 | 1.78E-05 | 07_cDC1 |
| NINJ1 | -1.61625361 | 0.231 | 0.575 | 1.79E-05 | 07_cDC1 |
| PIK3CB | 1.545126995 | 0.429 | 0.173 | 2.76E-05 | 07_cDC1 |
| SPPL3 | 1.73896602 | 0.473 | 0.221 | 2.79E-05 | 07_cDC1 |
| MRC1 | -13.7250095 | 0 | 0.301 | 3.05E-05 | 07_cDC1 |
| UTRN | -1.77668927 | 0.187 | 0.539 | 3.08E-05 | 07_cDC1 |
| PKP2 | 1.884862074 | 0.418 | 0.171 | 3.18E-05 | 07_cDC1 |
| ARID1B | 1.429030683 | 0.725 | 0.462 | 3.43E-05 | 07_cDC1 |
| LINC00963 | -4.23323127 | 0.044 | 0.349 | 3.56E-05 | 07_cDC1 |
| SAP30 | -3.83238008 | 0.077 | 0.382 | 3.59E-05 | 07_cDC1 |
| FOXN2 | 1.259998002 | 0.692 | 0.408 | 4.09E-05 | 07_cDC1 |
| ANXA6 | 1.318301155 | 0.341 | 0.122 | 4.23E-05 | 07_cDC1 |
| PPP1R14B | 1.46828261 | 0.505 | 0.242 | 4.67E-05 | 07_cDC1 |

| | | | | | |
|---|---|---|---|---|---|
| IRF4 | -5.01068745 | 0.022 | 0.323 | 5.50E-05 | 07_cDC1 |
| LPCAT2 | -2.30391388 | 0.209 | 0.511 | 5.88E-05 | 07_cDC1 |
| SMIM3 | -4.38131586 | 0.011 | 0.311 | 6.62E-05 | 07_cDC1 |
| PDE3B | -2.44861696 | 0.11 | 0.433 | 6.76E-05 | 07_cDC1 |
| NLRP3 | -1.07264306 | 0.407 | 0.743 | 7.48E-05 | 07_cDC1 |
| DAGLB | -2.65091286 | 0.088 | 0.408 | 8.24E-05 | 07_cDC1 |
| APOC1 | -3.67185333 | 0.231 | 0.517 | 8.32E-05 | 07_cDC1 |
| N4BP2 | 1.985133036 | 0.374 | 0.157 | 8.33E-05 | 07_cDC1 |
| ITGAV | -2.46776156 | 0.154 | 0.462 | 9.87E-05 | 07_cDC1 |
| HSD17B12 | 1.437631318 | 0.44 | 0.198 | 0.0001071 | 07_cDC1 |
| GBP2 | -3.89134281 | 0.044 | 0.344 | 0.0001113 | 07_cDC1 |
| NRARP | 1.5910944 | 0.549 | 0.311 | 0.0001195 | 07_cDC1 |
| EREG | -4.28348221 | 0.022 | 0.316 | 0.000122 | 07_cDC1 |
| CASP1 | -12.5352243 | 0 | 0.283 | 0.000126 | 07_cDC1 |
| PPM1M | 1.72794436 | 0.363 | 0.142 | 0.0001273 | 07_cDC1 |
| LPP | 1.544652527 | 0.802 | 0.566 | 0.0001284 | 07_cDC1 |
| CTSD | -2.59846728 | 0.165 | 0.465 | 0.0001316 | 07_cDC1 |
| MS4A4E | -3.03220793 | 0.077 | 0.379 | 0.0001358 | 07_cDC1 |
| DIPK2A | 1.544199081 | 0.319 | 0.111 | 0.0001392 | 07_cDC1 |
| SWAP70 | -1.95787642 | 0.187 | 0.496 | 0.0001612 | 07_cDC1 |
| A2M | -2.68941529 | 0.176 | 0.464 | 0.0001664 | 07_cDC1 |
| MAFB | -3.50636347 | 0.077 | 0.371 | 0.0001742 | 07_cDC1 |
| IRAK2 | 1.092327749 | 0.714 | 0.415 | 0.0001747 | 07_cDC1 |
| HMGB2 | -2.84045363 | 0.11 | 0.406 | 0.0001754 | 07_cDC1 |
| NDEL1 | 1.412852697 | 0.736 | 0.463 | 0.000177 | 07_cDC1 |
| CARD16 | -3.24542896 | 0.044 | 0.342 | 0.0001824 | 07_cDC1 |
| ARHGAP5 | 2.024271688 | 0.495 | 0.256 | 0.000188 | 07_cDC1 |
| ZC3HAV1 | 1.295241565 | 0.681 | 0.439 | 0.0001977 | 07_cDC1 |
| TNFSF13 | -4.83674364 | 0.033 | 0.313 | 0.0002047 | 07_cDC1 |
| FCGR1A | -4.29357205 | 0.044 | 0.329 | 0.0002175 | 07_cDC1 |
| PLIN2 | -2.91214478 | 0.165 | 0.445 | 0.0002217 | 07_cDC1 |
| GPX4 | -1.02266126 | 0.527 | 0.765 | 0.0002393 | 07_cDC1 |
| FLOT2 | 1.043377655 | 0.484 | 0.226 | 0.0002496 | 07_cDC1 |
| ERCC1 | -2.30154662 | 0.176 | 0.456 | 0.000267 | 07_cDC1 |
| KLHL2 | 1.963111853 | 0.363 | 0.151 | 0.0002802 | 07_cDC1 |
| TREM1 | -3.06595544 | 0.044 | 0.344 | 0.0002892 | 07_cDC1 |
| ARHGAP24 | -3.42273046 | 0.099 | 0.389 | 0.0003064 | 07_cDC1 |
| SPTLC2 | -2.16429149 | 0.231 | 0.525 | 0.0003747 | 07_cDC1 |
| PRKCH | -2.02774833 | 0.209 | 0.507 | 0.0003874 | 07_cDC1 |
| RASGEF1B | -1.35755244 | 0.473 | 0.703 | 0.0003991 | 07_cDC1 |
| APOE | -4.19197933 | 0.396 | 0.602 | 0.000406 | 07_cDC1 |
| FRMD4B | -3.77210776 | 0.022 | 0.305 | 0.0004413 | 07_cDC1 |
| TAGAP | -2.09513958 | 0.165 | 0.456 | 0.0004426 | 07_cDC1 |
| G3BP2 | 1.212752097 | 0.582 | 0.309 | 0.0004748 | 07_cDC1 |
| PTPN2 | 1.438396107 | 0.703 | 0.482 | 0.0004983 | 07_cDC1 |
| MGAT1 | -1.12966052 | 0.516 | 0.746 | 0.0005076 | 07_cDC1 |

| | | | | | |
|---|---|---|---|---|---|
| TOM1 | -2.01724059 | 0.176 | 0.467 | 0.0005326 | 07_cDC1 |
| RAB32 | 1.45983976 | 0.495 | 0.263 | 0.0005364 | 07_cDC1 |
| LTC4S | -2.46284534 | 0.11 | 0.402 | 0.0005582 | 07_cDC1 |
| AP001011.1 | 1.381266452 | 0.319 | 0.116 | 0.0005615 | 07_cDC1 |
| MAPK8 | 1.186449659 | 0.516 | 0.252 | 0.0005726 | 07_cDC1 |
| SLC25A33 | 1.751775384 | 0.451 | 0.215 | 0.0005834 | 07_cDC1 |
| EPS8 | -2.76074037 | 0.077 | 0.363 | 0.0006573 | 07_cDC1 |
| PIK3R1 | -2.21628199 | 0.176 | 0.469 | 0.0006801 | 07_cDC1 |
| SCPEP1 | -3.61872131 | 0.033 | 0.309 | 0.0007254 | 07_cDC1 |
| C3AR1 | -3.52138364 | 0.044 | 0.32 | 0.000763 | 07_cDC1 |
| PTAFR | -5.75066337 | 0.011 | 0.273 | 0.0008121 | 07_cDC1 |
| CD9 | -2.14031021 | 0.11 | 0.404 | 0.0008303 | 07_cDC1 |
| DAPK1 | 1.160817606 | 0.593 | 0.323 | 0.000848 | 07_cDC1 |
| PAPSS2 | 1.987685209 | 0.429 | 0.199 | 0.0008656 | 07_cDC1 |
| SLCO3A1 | 1.885038514 | 0.374 | 0.158 | 0.0008792 | 07_cDC1 |
| RCSD1 | -2.29682474 | 0.11 | 0.403 | 0.0009067 | 07_cDC1 |
| LAMP1 | -12.4252179 | 0 | 0.257 | 0.0009515 | 07_cDC1 |
| TXNIP | -1.42836719 | 0.275 | 0.569 | 0.000959 | 07_cDC1 |
| PRKAG2 | -1.66588139 | 0.242 | 0.521 | 0.0010334 | 07_cDC1 |
| JDP2 | -1.93350329 | 0.176 | 0.472 | 0.001065 | 07_cDC1 |
| P2RY6 | 1.355321783 | 0.407 | 0.171 | 0.0011105 | 07_cDC1 |
| FRMD4A | -2.99786916 | 0.198 | 0.457 | 0.0011181 | 07_cDC1 |
| LBR | 1.371480131 | 0.341 | 0.139 | 0.0012128 | 07_cDC1 |
| SRI | 1.093595148 | 0.549 | 0.292 | 0.0013133 | 07_cDC1 |
| KIF2A | 1.511039428 | 0.385 | 0.169 | 0.0013737 | 07_cDC1 |
| ALCAM | -1.53965165 | 0.396 | 0.673 | 0.0013944 | 07_cDC1 |
| CD99 | -1.6815427 | 0.264 | 0.538 | 0.0015217 | 07_cDC1 |
| GRINA | -1.15337631 | 0.451 | 0.661 | 0.0017339 | 07_cDC1 |
| CTSC | -2.22548929 | 0.176 | 0.445 | 0.0017897 | 07_cDC1 |
| PAG1 | -1.43322729 | 0.121 | 0.417 | 0.001812 | 07_cDC1 |
| IFI6 | -3.18570265 | 0.066 | 0.335 | 0.0018198 | 07_cDC1 |
| CD68 | -1.71611251 | 0.374 | 0.579 | 0.0018494 | 07_cDC1 |
| CBL | 1.341258384 | 0.44 | 0.219 | 0.0020858 | 07_cDC1 |
| CARD11 | 1.198708934 | 0.418 | 0.19 | 0.0021984 | 07_cDC1 |
| SLC15A3 | -4.4730716 | 0.022 | 0.275 | 0.0022672 | 07_cDC1 |
| WBP1L | -3.76295617 | 0.033 | 0.292 | 0.0022767 | 07_cDC1 |
| GPR34 | -4.94372808 | 0.022 | 0.274 | 0.002389 | 07_cDC1 |
| TMEM51 | -12.2278867 | 0 | 0.245 | 0.0024632 | 07_cDC1 |
| MFSD14B | 1.208324944 | 0.396 | 0.17 | 0.0025278 | 07_cDC1 |
| PRKAG2-AS1 | -4.23014693 | 0.022 | 0.275 | 0.0026193 | 07_cDC1 |
| LRBA | 1.384123448 | 0.429 | 0.208 | 0.0026601 | 07_cDC1 |
| KCNK6 | 1.682771035 | 0.352 | 0.146 | 0.002688 | 07_cDC1 |
| TNFSF12 | -12.051941 | 0 | 0.242 | 0.0030344 | 07_cDC1 |
| UBASH3B | 1.472385328 | 0.703 | 0.427 | 0.0031522 | 07_cDC1 |
| GPSM3 | -1.43471777 | 0.264 | 0.548 | 0.0031759 | 07_cDC1 |
| EPB41L2 | -2.01782938 | 0.231 | 0.485 | 0.0033777 | 07_cDC1 |

| | | | | | |
|---|---|---|---|---|---|
| NEK7 | 1.471356135 | 0.473 | 0.249 | 0.0033909 | 07_cDC1 |
| PRDM1 | -3.48328326 | 0.044 | 0.304 | 0.0034749 | 07_cDC1 |
| OPN3 | -3.38519894 | 0.055 | 0.314 | 0.003589 | 07_cDC1 |
| F13A1 | -12.5849389 | 0 | 0.239 | 0.0037344 | 07_cDC1 |
| JPT1 | 1.099084419 | 0.626 | 0.411 | 0.003762 | 07_cDC1 |
| LILRB2 | -3.1853025 | 0.022 | 0.276 | 0.0042029 | 07_cDC1 |
| DTNA | -3.71765052 | 0.011 | 0.257 | 0.004861 | 07_cDC1 |
| SLC3A2 | -2.28399366 | 0.165 | 0.417 | 0.0049761 | 07_cDC1 |
| TMCC3 | -3.2752548 | 0.033 | 0.287 | 0.0051175 | 07_cDC1 |
| BIRC3 | 1.066342771 | 0.626 | 0.37 | 0.0051308 | 07_cDC1 |
| CHD1 | -1.48200321 | 0.385 | 0.621 | 0.0057082 | 07_cDC1 |
| NIBAN1 | 1.302874453 | 0.593 | 0.363 | 0.0058351 | 07_cDC1 |
| ATP6AP1 | -1.49177922 | 0.341 | 0.557 | 0.0058705 | 07_cDC1 |
| PDGFB | -2.35595783 | 0.132 | 0.394 | 0.0063594 | 07_cDC1 |
| PPARD | -1.17979072 | 0.275 | 0.551 | 0.0064415 | 07_cDC1 |
| SCFD1 | 1.325992121 | 0.407 | 0.187 | 0.0067088 | 07_cDC1 |
| TLE3 | 1.313896533 | 0.374 | 0.162 | 0.0067989 | 07_cDC1 |
| AP3D1 | 1.273607247 | 0.418 | 0.196 | 0.0069351 | 07_cDC1 |
| CD302 | -1.32462136 | 0.297 | 0.559 | 0.0069445 | 07_cDC1 |
| TGFB1 | -1.14156542 | 0.462 | 0.682 | 0.0079008 | 07_cDC1 |
| MACC1 | -13.1135291 | 0 | 0.228 | 0.0080615 | 07_cDC1 |
| YBX3 | 1.148674468 | 0.648 | 0.422 | 0.0085663 | 07_cDC1 |
| TTYH3 | -3.09634847 | 0.044 | 0.291 | 0.0088921 | 07_cDC1 |
| VPS37B | -2.82629914 | 0.077 | 0.33 | 0.0091659 | 07_cDC1 |
| VMO1 | 1.091307881 | 0.659 | 0.412 | 0.0094355 | 07_cDC1 |
| SLC16A3 | -1.22180437 | 0.319 | 0.591 | 0.0094855 | 07_cDC1 |
| SLC9A7 | 1.075518215 | 0.429 | 0.2 | 0.009948 | 07_cDC1 |
| KCNQ3 | -6.5897259 | 0.011 | 0.24 | 0.0102138 | 07_cDC1 |
| ENG | -2.37629644 | 0.066 | 0.321 | 0.0102141 | 07_cDC1 |
| TRPM2 | -12.2885015 | 0 | 0.225 | 0.0103871 | 07_cDC1 |
| ABL2 | -2.03773307 | 0.143 | 0.405 | 0.0106215 | 07_cDC1 |
| NDUFS4 | 1.206632217 | 0.418 | 0.207 | 0.0107105 | 07_cDC1 |
| ABRACL | 1.392832767 | 0.418 | 0.217 | 0.0109212 | 07_cDC1 |
| PID1 | -13.1002793 | 0 | 0.224 | 0.0109252 | 07_cDC1 |
| NCOA4 | -1.92716636 | 0.176 | 0.43 | 0.0117801 | 07_cDC1 |
| RIPK2 | -1.48315631 | 0.286 | 0.538 | 0.0121898 | 07_cDC1 |
| RAB5A | 1.131005904 | 0.56 | 0.355 | 0.0128386 | 07_cDC1 |
| CTNND1 | -1.8158738 | 0.143 | 0.415 | 0.0129259 | 07_cDC1 |
| GABARAPL1 | -1.64948531 | 0.242 | 0.491 | 0.0148544 | 07_cDC1 |
| IGFBP7 | 1.376389097 | 0.396 | 0.182 | 0.0153248 | 07_cDC1 |
| CCR7 | -1.67203601 | 0.066 | 0.325 | 0.0154286 | 07_cDC1 |
| SLA | -1.9474829 | 0.176 | 0.431 | 0.0155151 | 07_cDC1 |
| DSTN | 1.221567206 | 0.407 | 0.2 | 0.0155302 | 07_cDC1 |
| ITGAM | -4.66784234 | 0.011 | 0.237 | 0.0161418 | 07_cDC1 |
| SNCA | -12.2807471 | 0 | 0.218 | 0.0171668 | 07_cDC1 |
| FGD4 | -1.3068706 | 0.462 | 0.71 | 0.0172793 | 07_cDC1 |

| | | | | | |
|---|---|---|---|---|---|
| IFI44L | -6.23244913 | 0.011 | 0.231 | 0.0181904 | 07_cDC1 |
| HTT | 1.300728116 | 0.385 | 0.183 | 0.0186559 | 07_cDC1 |
| ATP6V1H | 1.818501455 | 0.418 | 0.204 | 0.0194526 | 07_cDC1 |
| TRAK1 | 1.830089628 | 0.505 | 0.283 | 0.0194819 | 07_cDC1 |
| RB1 | -1.48822582 | 0.275 | 0.531 | 0.0200398 | 07_cDC1 |
| LPAR6 | -5.84155219 | 0.011 | 0.229 | 0.021626 | 07_cDC1 |
| CSTA | -3.28492938 | 0.022 | 0.251 | 0.022487 | 07_cDC1 |
| PRKAR1A | 1.032491818 | 0.604 | 0.384 | 0.0239205 | 07_cDC1 |
| DLGAP4 | 1.421886821 | 0.429 | 0.221 | 0.0241015 | 07_cDC1 |
| IL1RAP | -2.15984044 | 0.055 | 0.299 | 0.0246688 | 07_cDC1 |
| AKR1B1 | -1.95664324 | 0.187 | 0.418 | 0.0250179 | 07_cDC1 |
| 6-Mar | 1.086535463 | 0.462 | 0.238 | 0.0251186 | 07_cDC1 |
| XYLT1 | -1.19859562 | 0.22 | 0.486 | 0.0252328 | 07_cDC1 |
| GAPT | -12.0428446 | 0 | 0.212 | 0.0255503 | 07_cDC1 |
| ST6GALNAC3 | -2.27728082 | 0.055 | 0.297 | 0.0265881 | 07_cDC1 |
| SLC37A2 | -12.2264396 | 0 | 0.212 | 0.0268454 | 07_cDC1 |
| CDC42SE1 | -1.34008757 | 0.242 | 0.495 | 0.0268524 | 07_cDC1 |
| SRP19 | 1.144978504 | 0.462 | 0.246 | 0.0275518 | 07_cDC1 |
| MYD88 | 1.119513718 | 0.462 | 0.257 | 0.0304862 | 07_cDC1 |
| IDH2 | 1.078233463 | 0.451 | 0.247 | 0.0307939 | 07_cDC1 |
| FNDC3B | -1.53106584 | 0.198 | 0.468 | 0.0309675 | 07_cDC1 |
| DLEU1 | -2.87754979 | 0.165 | 0.387 | 0.0331736 | 07_cDC1 |
| CREB5 | -1.49063993 | 0.099 | 0.351 | 0.0340596 | 07_cDC1 |
| LCP2 | -1.39536839 | 0.352 | 0.595 | 0.035235 | 07_cDC1 |
| TTYH2 | 1.143301923 | 0.451 | 0.228 | 0.0352708 | 07_cDC1 |
| CD1E | -4.70908814 | 0.011 | 0.226 | 0.0354972 | 07_cDC1 |
| AL078590.2 | -2.32230306 | 0.154 | 0.387 | 0.0377775 | 07_cDC1 |
| RASSF4 | -1.88753092 | 0.22 | 0.447 | 0.0390328 | 07_cDC1 |
| ZFHX3 | 1.36355709 | 0.593 | 0.389 | 0.0391502 | 07_cDC1 |
| ST8SIA4 | 1.19164698 | 0.626 | 0.416 | 0.0403539 | 07_cDC1 |
| AC004817.3 | -2.2314231 | 0.077 | 0.323 | 0.0411309 | 07_cDC1 |
| SSBP2 | -2.47975099 | 0.099 | 0.34 | 0.0411463 | 07_cDC1 |
| DOK2 | -12.0353815 | 0 | 0.205 | 0.0417837 | 07_cDC1 |
| SLC25A37 | -2.13657918 | 0.154 | 0.387 | 0.04398 | 07_cDC1 |
| AP001636.3 | -5.18161728 | 0.011 | 0.221 | 0.0458358 | 07_cDC1 |
| MPP1 | -3.52905769 | 0.044 | 0.264 | 0.0467419 | 07_cDC1 |
| IFITM1 | -1.8640278 | 0.187 | 0.42 | 0.0549308 | 07_cDC1 |
| TFEC | -4.36871852 | 0.022 | 0.232 | 0.0603602 | 07_cDC1 |
| SORT1 | -4.07456142 | 0.022 | 0.233 | 0.0653151 | 07_cDC1 |
| PRAM1 | -5.90631025 | 0.011 | 0.213 | 0.0672484 | 07_cDC1 |
| MEIKIN | -3.60982241 | 0.011 | 0.219 | 0.0679222 | 07_cDC1 |
| BHLHE41 | -2.35731991 | 0.044 | 0.27 | 0.0703314 | 07_cDC1 |
| N4BP2L2 | -1.26591163 | 0.22 | 0.501 | 0.0710009 | 07_cDC1 |
| ITGA5 | -2.17160838 | 0.187 | 0.402 | 0.0794447 | 07_cDC1 |
| LRP1 | -4.79083573 | 0.011 | 0.214 | 0.0805174 | 07_cDC1 |
| CKS2 | 1.444215065 | 0.484 | 0.277 | 0.081578 | 07_cDC1 |

| | | | | | |
|---|---|---|---|---|---|
| XBP1 | -1.36087718 | 0.286 | 0.517 | 0.083315 | 07_cDC1 |
| PMEPA1 | -3.18583797 | 0.044 | 0.257 | 0.0879004 | 07_cDC1 |
| TMEM176B | -3.90882781 | 0.022 | 0.23 | 0.088777 | 07_cDC1 |
| FBXO34 | 1.446971994 | 0.549 | 0.323 | 0.0911182 | 07_cDC1 |
| AZI2 | -3.7568436 | 0.033 | 0.244 | 0.1014942 | 07_cDC1 |
| PTDSS1 | 1.182494526 | 0.473 | 0.271 | 0.108938 | 07_cDC1 |
| OSM | -3.1831085 | 0.033 | 0.244 | 0.1137066 | 07_cDC1 |
| EBI3 | -1.59453044 | 0.099 | 0.333 | 0.1334738 | 07_cDC1 |
| PLA2G4A | -2.48302402 | 0.132 | 0.351 | 0.138532 | 07_cDC1 |
| CD69 | -1.1297632 | 0.209 | 0.452 | 0.1401247 | 07_cDC1 |
| LINC02256 | -2.13953929 | 0.121 | 0.344 | 0.1412029 | 07_cDC1 |
| MED13 | 1.118543122 | 0.527 | 0.313 | 0.1496052 | 07_cDC1 |
| PPP3R1 | 1.193569891 | 0.527 | 0.318 | 0.1680578 | 07_cDC1 |
| LHFPL2 | -2.97382636 | 0.066 | 0.273 | 0.1775114 | 07_cDC1 |
| CLEC4A | -3.1041622 | 0.033 | 0.234 | 0.1819774 | 07_cDC1 |
| AL136987.1 | -3.01362008 | 0.055 | 0.26 | 0.193341 | 07_cDC1 |
| NECTIN2 | -1.16706454 | 0.231 | 0.463 | 0.2513438 | 07_cDC1 |
| RUNX2 | -1.61905166 | 0.143 | 0.377 | 0.2522482 | 07_cDC1 |
| GSTO1 | -1.56578617 | 0.198 | 0.417 | 0.2747345 | 07_cDC1 |
| RHOQ | -1.23682697 | 0.363 | 0.583 | 0.282105 | 07_cDC1 |
| PCED1B-AS1 | -2.62598658 | 0.055 | 0.258 | 0.2878905 | 07_cDC1 |
| TMEM131 | 1.13789325 | 0.593 | 0.372 | 0.3049315 | 07_cDC1 |
| CCDC50 | -1.92565359 | 0.055 | 0.264 | 0.3124846 | 07_cDC1 |
| ARID5A | -1.1436467 | 0.242 | 0.484 | 0.3392029 | 07_cDC1 |
| TRIB1 | -1.463278 | 0.286 | 0.491 | 0.3641514 | 07_cDC1 |
| HIF1A-AS3 | -2.24303346 | 0.121 | 0.326 | 0.3802562 | 07_cDC1 |
| EHBP1L1 | 1.097263141 | 0.44 | 0.233 | 0.3984071 | 07_cDC1 |
| FYN | 1.098110316 | 0.571 | 0.349 | 0.4212118 | 07_cDC1 |
| AC025164.1 | -2.29469094 | 0.055 | 0.257 | 0.4777824 | 07_cDC1 |
| ABCC4 | -1.974696 | 0.055 | 0.259 | 0.5257994 | 07_cDC1 |
| AL390957.1 | -1.53554187 | 0.121 | 0.338 | 0.5823898 | 07_cDC1 |
| SSH1 | -2.16516902 | 0.11 | 0.31 | 0.6063116 | 07_cDC1 |
| PFDN1 | -1.94934384 | 0.088 | 0.289 | 0.6573223 | 07_cDC1 |
| BCAT1 | -1.30084293 | 0.121 | 0.331 | 0.7077676 | 07_cDC1 |
| DNM2 | -1.58776619 | 0.231 | 0.432 | 0.7203498 | 07_cDC1 |
| MAGT1 | -1.27220966 | 0.154 | 0.384 | 0.7687038 | 07_cDC1 |
| MAN2A1 | -1.32981279 | 0.264 | 0.472 | 0.857214 | 07_cDC1 |
| LY6E | -1.39180853 | 0.187 | 0.396 | 0.9565331 | 07_cDC1 |
| INSR | -1.19536904 | 0.165 | 0.378 | 1 | 07_cDC1 |
| ITSN1 | -1.82134284 | 0.099 | 0.302 | 1 | 07_cDC1 |
| BEST1 | -1.51884001 | 0.154 | 0.363 | 1 | 07_cDC1 |
| TSPAN14 | -1.22616683 | 0.11 | 0.319 | 1 | 07_cDC1 |
| UBAC2 | -1.27907439 | 0.253 | 0.469 | 1 | 07_cDC1 |
| PRKCE | -1.14408921 | 0.264 | 0.467 | 1 | 07_cDC1 |
| HOOK2 | -1.22634319 | 0.198 | 0.398 | 1 | 07_cDC1 |
| RNF13 | -1.05721716 | 0.22 | 0.421 | 1 | 07_cDC1 |

| | | | | | |
|---|---|---|---|---|---|
| SPRY1 | 4.573023521 | 0.457 | 0.027 | 3.26E-63 | 08_mo-DC3 |
| CTTNBP2 | 4.592877771 | 0.642 | 0.072 | 3.74E-61 | 08_mo-DC3 |
| MERTK | 2.991325426 | 0.864 | 0.156 | 8.66E-58 | 08_mo-DC3 |
| SDK1 | 3.749935495 | 0.728 | 0.111 | 1.07E-55 | 08_mo-DC3 |
| IGSF21 | 3.761154639 | 0.605 | 0.078 | 3.92E-50 | 08_mo-DC3 |
| CDC14B | 5.755452764 | 0.309 | 0.015 | 1.98E-47 | 08_mo-DC3 |
| FRMD4A | 2.55649956 | 1 | 0.41 | 3.67E-39 | 08_mo-DC3 |
| KCNMA1 | 3.247789947 | 0.691 | 0.14 | 1.49E-38 | 08_mo-DC3 |
| SLC2A5 | 2.694229782 | 0.654 | 0.118 | 1.58E-38 | 08_mo-DC3 |
| TAL1 | 3.921699627 | 0.37 | 0.032 | 2.60E-38 | 08_mo-DC3 |
| KLF2 | 3.736596705 | 0.877 | 0.285 | 5.59E-38 | 08_mo-DC3 |
| LINC02798 | 3.663908389 | 0.519 | 0.078 | 4.69E-36 | 08_mo-DC3 |
| LHFPL2 | 2.210289624 | 0.84 | 0.228 | 1.33E-35 | 08_mo-DC3 |
| SLC1A3 | 2.104301563 | 1 | 0.499 | 1.52E-34 | 08_mo-DC3 |
| ARHGAP24 | 2.443370347 | 0.938 | 0.34 | 1.83E-34 | 08_mo-DC3 |
| CSGALNACT1 | 2.242066934 | 0.938 | 0.315 | 6.71E-34 | 08_mo-DC3 |
| TMEM119 | 3.295112874 | 0.346 | 0.035 | 1.20E-30 | 08_mo-DC3 |
| SIGLEC8 | 3.081717823 | 0.444 | 0.064 | 4.66E-30 | 08_mo-DC3 |
| PLTP | 3.093300869 | 0.63 | 0.147 | 5.65E-30 | 08_mo-DC3 |
| ARHGAP6 | 3.193645697 | 0.531 | 0.096 | 5.92E-30 | 08_mo-DC3 |
| PDPN | 2.87417721 | 0.469 | 0.072 | 6.21E-30 | 08_mo-DC3 |
| SYNDIG1 | 3.26762283 | 0.37 | 0.043 | 2.84E-29 | 08_mo-DC3 |
| VIM | -2.71105914 | 0.457 | 0.955 | 3.26E-29 | 08_mo-DC3 |
| SLCO2B1 | 2.182693342 | 0.741 | 0.202 | 3.99E-29 | 08_mo-DC3 |
| CPA6 | 3.835537465 | 0.333 | 0.036 | 1.63E-28 | 08_mo-DC3 |
| GPR34 | 2.367449814 | 0.765 | 0.23 | 2.73E-28 | 08_mo-DC3 |
| ITGA9 | 2.693572954 | 0.444 | 0.066 | 6.90E-28 | 08_mo-DC3 |
| C1QC | 2.105172569 | 0.963 | 0.523 | 9.70E-28 | 08_mo-DC3 |
| NPL | 2.892988114 | 0.519 | 0.097 | 1.44E-27 | 08_mo-DC3 |
| C1QB | 2.061529024 | 0.975 | 0.52 | 1.63E-27 | 08_mo-DC3 |
| ST6GAL1 | 2.447676145 | 0.802 | 0.27 | 6.81E-27 | 08_mo-DC3 |
| C3 | 2.042828136 | 0.963 | 0.492 | 1.57E-26 | 08_mo-DC3 |
| C5AR1 | 2.03679008 | 0.901 | 0.348 | 3.25E-26 | 08_mo-DC3 |
| PLVAP | 4.241408418 | 0.284 | 0.028 | 5.40E-26 | 08_mo-DC3 |
| SPTLC2 | 2.569388971 | 0.889 | 0.486 | 1.46E-25 | 08_mo-DC3 |
| IL1RAP | 2.432977082 | 0.778 | 0.257 | 2.14E-25 | 08_mo-DC3 |
| RHOB | 2.32076678 | 0.975 | 0.587 | 2.17E-25 | 08_mo-DC3 |
| PDK4 | 2.798850994 | 0.716 | 0.219 | 1.93E-24 | 08_mo-DC3 |
| EPB41L2 | 1.986426322 | 0.914 | 0.445 | 2.51E-24 | 08_mo-DC3 |
| AREG | -2.82668392 | 0.259 | 0.865 | 3.82E-24 | 08_mo-DC3 |
| IPCEF1 | 2.880747041 | 0.605 | 0.155 | 4.19E-24 | 08_mo-DC3 |
| APOE | 1.733746112 | 0.975 | 0.567 | 4.41E-24 | 08_mo-DC3 |
| MLXIPL | 3.247387898 | 0.346 | 0.045 | 4.41E-24 | 08_mo-DC3 |
| LGMN | 2.365129606 | 0.827 | 0.343 | 6.77E-24 | 08_mo-DC3 |
| SCIN | 2.389875637 | 0.543 | 0.116 | 7.53E-24 | 08_mo-DC3 |
| STAB1 | 2.13367113 | 0.593 | 0.142 | 1.95E-23 | 08_mo-DC3 |

| | | | | | |
|---|---|---|---|---|---|
| SPRED1 | 2.518639223 | 0.58 | 0.139 | 2.87E-23 | 08_mo-DC3 |
| ZDHHC14 | 2.519444368 | 0.481 | 0.094 | 2.89E-23 | 08_mo-DC3 |
| DOCK4 | 1.61792048 | 1 | 0.769 | 1.53E-22 | 08_mo-DC3 |
| S100A10 | -4.02788356 | 0.148 | 0.777 | 2.45E-22 | 08_mo-DC3 |
| CX3CR1 | 2.288503097 | 0.556 | 0.13 | 2.87E-22 | 08_mo-DC3 |
| FOLR2 | 3.101401873 | 0.42 | 0.075 | 2.89E-22 | 08_mo-DC3 |
| EGR3 | 2.539526638 | 0.494 | 0.107 | 3.59E-22 | 08_mo-DC3 |
| ABCC4 | 2.352338707 | 0.704 | 0.221 | 6.95E-22 | 08_mo-DC3 |
| S100A6 | -4.21790999 | 0.148 | 0.754 | 1.23E-21 | 08_mo-DC3 |
| MCF2L | 2.536451284 | 0.457 | 0.088 | 1.33E-21 | 08_mo-DC3 |
| MTSS1 | 2.430908878 | 0.642 | 0.189 | 1.54E-21 | 08_mo-DC3 |
| KCNQ3 | 1.960837586 | 0.679 | 0.201 | 4.71E-21 | 08_mo-DC3 |
| LSP1 | -3.80514402 | 0.198 | 0.751 | 9.86E-21 | 08_mo-DC3 |
| S100A4 | -4.28712955 | 0.16 | 0.74 | 1.45E-20 | 08_mo-DC3 |
| LPCAT2 | 1.56303846 | 0.926 | 0.468 | 1.71E-20 | 08_mo-DC3 |
| SPP1 | 1.917555941 | 0.938 | 0.604 | 2.27E-20 | 08_mo-DC3 |
| CHD7 | 2.241879445 | 0.506 | 0.115 | 2.28E-20 | 08_mo-DC3 |
| APBB1IP | 1.747491455 | 0.914 | 0.477 | 2.31E-20 | 08_mo-DC3 |
| MSR1 | 1.649191704 | 0.951 | 0.512 | 2.98E-20 | 08_mo-DC3 |
| CH25H | 2.826353364 | 0.543 | 0.142 | 3.25E-20 | 08_mo-DC3 |
| FCAR | 3.105735945 | 0.296 | 0.039 | 3.58E-20 | 08_mo-DC3 |
| RYR1 | 2.685513183 | 0.432 | 0.089 | 9.25E-20 | 08_mo-DC3 |
| CCDC26 | 2.649073388 | 0.506 | 0.121 | 1.09E-19 | 08_mo-DC3 |
| C1QA | 1.666355345 | 0.926 | 0.482 | 1.41E-19 | 08_mo-DC3 |
| ELL2 | 1.592566452 | 0.988 | 0.728 | 1.80E-19 | 08_mo-DC3 |
| C3AR1 | 2.053109597 | 0.728 | 0.28 | 2.87E-19 | 08_mo-DC3 |
| SLC9A9 | 2.026288268 | 0.63 | 0.186 | 4.95E-19 | 08_mo-DC3 |
| FGD4 | 1.652799341 | 0.975 | 0.679 | 7.07E-19 | 08_mo-DC3 |
| NBL1 | 3.882829799 | 0.222 | 0.022 | 9.84E-19 | 08_mo-DC3 |
| LYZ | -3.80899248 | 0.185 | 0.738 | 1.38E-18 | 08_mo-DC3 |
| APPL2 | 2.32498825 | 0.457 | 0.1 | 1.78E-18 | 08_mo-DC3 |
| GNB4 | 1.826081307 | 0.79 | 0.336 | 1.89E-18 | 08_mo-DC3 |
| GALNT2 | 1.648225105 | 0.741 | 0.282 | 1.90E-18 | 08_mo-DC3 |
| SLC11A1 | 1.629359573 | 0.889 | 0.43 | 3.09E-18 | 08_mo-DC3 |
| TCF12 | 1.756391417 | 0.901 | 0.518 | 3.73E-18 | 08_mo-DC3 |
| A2M | 2.01413322 | 0.852 | 0.424 | 4.29E-18 | 08_mo-DC3 |
| OXR1 | 3.309801937 | 0.519 | 0.149 | 8.15E-18 | 08_mo-DC3 |
| SH3RF3 | 2.337572746 | 0.444 | 0.098 | 8.30E-18 | 08_mo-DC3 |
| PIK3IP1 | 2.099043367 | 0.531 | 0.141 | 1.06E-17 | 08_mo-DC3 |
| PDE3B | 2.242957042 | 0.815 | 0.391 | 1.40E-17 | 08_mo-DC3 |
| DLEU1 | 1.65771933 | 0.827 | 0.348 | 1.47E-17 | 08_mo-DC3 |
| SOCS6 | 2.481191859 | 0.531 | 0.15 | 2.33E-17 | 08_mo-DC3 |
| CYFIP1 | 1.695396212 | 0.79 | 0.343 | 2.61E-17 | 08_mo-DC3 |
| USP53 | 1.770794823 | 0.827 | 0.38 | 2.83E-17 | 08_mo-DC3 |
| JDP2 | 1.917241446 | 0.84 | 0.432 | 3.76E-17 | 08_mo-DC3 |
| TREM2 | 1.548982142 | 0.827 | 0.359 | 8.91E-17 | 08_mo-DC3 |

| | | | | | |
|---|---|---|---|---|---|
| PLA2G4A | 2.126879684 | 0.753 | 0.315 | 1.26E-16 | 08_mo-DC3 |
| PRKN | 2.085091548 | 0.444 | 0.101 | 1.42E-16 | 08_mo-DC3 |
| MAP4K3 | 1.675115284 | 0.852 | 0.446 | 1.98E-16 | 08_mo-DC3 |
| PAG1 | 1.7152834 | 0.815 | 0.375 | 1.99E-16 | 08_mo-DC3 |
| ARHGAP12 | 2.620697406 | 0.309 | 0.051 | 2.20E-16 | 08_mo-DC3 |
| ARHGAP26 | 1.746683103 | 0.938 | 0.627 | 5.06E-16 | 08_mo-DC3 |
| CENPU | 2.445474714 | 0.333 | 0.059 | 8.25E-16 | 08_mo-DC3 |
| BMP2K | 1.675596617 | 0.741 | 0.315 | 9.41E-16 | 08_mo-DC3 |
| SLC4A7 | 2.10986577 | 0.716 | 0.311 | 9.89E-16 | 08_mo-DC3 |
| GSTM2 | 2.532989389 | 0.481 | 0.137 | 1.18E-15 | 08_mo-DC3 |
| APOC1 | 1.067587856 | 0.914 | 0.477 | 1.26E-15 | 08_mo-DC3 |
| TBC1D16 | 2.390377868 | 0.481 | 0.131 | 1.28E-15 | 08_mo-DC3 |
| ST6GALNAC3 | 1.976602237 | 0.667 | 0.26 | 2.21E-15 | 08_mo-DC3 |
| ANXA2 | -3.45214478 | 0.111 | 0.664 | 3.40E-15 | 08_mo-DC3 |
| PIK3R1 | 1.741218585 | 0.802 | 0.432 | 3.55E-15 | 08_mo-DC3 |
| IFITM10 | 2.193516359 | 0.42 | 0.096 | 5.31E-15 | 08_mo-DC3 |
| MS4A7 | 1.255947864 | 0.926 | 0.724 | 7.34E-15 | 08_mo-DC3 |
| LINC01374 | 2.151465832 | 0.469 | 0.122 | 8.82E-15 | 08_mo-DC3 |
| MAFB | 1.782522934 | 0.753 | 0.331 | 9.85E-15 | 08_mo-DC3 |
| ATP6V0A1 | 1.739051867 | 0.58 | 0.182 | 1.04E-14 | 08_mo-DC3 |
| ADRB2 | 3.025645104 | 0.284 | 0.047 | 1.58E-14 | 08_mo-DC3 |
| RNASE1 | 2.711152191 | 0.444 | 0.119 | 1.63E-14 | 08_mo-DC3 |
| BTG2 | 1.546194539 | 0.914 | 0.616 | 1.81E-14 | 08_mo-DC3 |
| LILRB4 | 1.47673973 | 0.852 | 0.412 | 2.06E-14 | 08_mo-DC3 |
| CTSD | 1.618920039 | 0.827 | 0.426 | 2.09E-14 | 08_mo-DC3 |
| DAB2 | 2.287957016 | 0.457 | 0.124 | 2.27E-14 | 08_mo-DC3 |
| TMIGD3 | 2.045871591 | 0.481 | 0.13 | 2.65E-14 | 08_mo-DC3 |
| AC008957.2 | 2.390072646 | 0.284 | 0.046 | 3.17E-14 | 08_mo-DC3 |
| ARG2 | 3.007585834 | 0.272 | 0.044 | 5.01E-14 | 08_mo-DC3 |
| CSF1R | 1.406646489 | 0.914 | 0.559 | 5.10E-14 | 08_mo-DC3 |
| ZFP36L2 | 1.482978926 | 0.963 | 0.734 | 8.45E-14 | 08_mo-DC3 |
| TIMP1 | -4.31512975 | 0.259 | 0.702 | 9.68E-14 | 08_mo-DC3 |
| SFMBT2 | 1.760899475 | 0.815 | 0.431 | 1.19E-13 | 08_mo-DC3 |
| TTC7B | 2.597800357 | 0.37 | 0.084 | 1.25E-13 | 08_mo-DC3 |
| CCL3 | 2.314362035 | 0.877 | 0.576 | 1.42E-13 | 08_mo-DC3 |
| AC046195.1 | 2.05376775 | 0.309 | 0.057 | 1.58E-13 | 08_mo-DC3 |
| HEATR5B | 2.279155109 | 0.519 | 0.162 | 1.95E-13 | 08_mo-DC3 |
| TMEM117 | 2.284197196 | 0.37 | 0.083 | 4.77E-13 | 08_mo-DC3 |
| SLA | 1.475998366 | 0.79 | 0.395 | 5.16E-13 | 08_mo-DC3 |
| OLR1 | 1.367720434 | 0.914 | 0.65 | 5.43E-13 | 08_mo-DC3 |
| BNC2 | 2.011508597 | 0.531 | 0.165 | 5.46E-13 | 08_mo-DC3 |
| FCGBP | 1.765511662 | 0.481 | 0.139 | 1.23E-12 | 08_mo-DC3 |
| P2RY12 | 2.31773266 | 0.481 | 0.15 | 1.55E-12 | 08_mo-DC3 |
| DUSP6 | 2.704531447 | 0.42 | 0.111 | 1.87E-12 | 08_mo-DC3 |
| DIAPH2 | 1.721613728 | 0.63 | 0.239 | 2.13E-12 | 08_mo-DC3 |
| CCL4 | 2.227202527 | 0.926 | 0.615 | 2.15E-12 | 08_mo-DC3 |

| | | | | | |
|---|---|---|---|---|---|
| AP2A2 | 1.837691191 | 0.494 | 0.158 | 2.62E-12 | 08_mo-DC3 |
| ADORA3 | 2.658457557 | 0.37 | 0.091 | 2.72E-12 | 08_mo-DC3 |
| SUSD6 | 1.424210297 | 0.815 | 0.464 | 2.76E-12 | 08_mo-DC3 |
| DAGLB | 1.568595456 | 0.753 | 0.368 | 2.84E-12 | 08_mo-DC3 |
| LRMDA | 1.27627236 | 0.926 | 0.687 | 3.04E-12 | 08_mo-DC3 |
| ANKRD22 | 1.841586311 | 0.432 | 0.117 | 3.18E-12 | 08_mo-DC3 |
| KIF21B | 2.286397124 | 0.383 | 0.094 | 3.59E-12 | 08_mo-DC3 |
| MICU3 | 2.601078602 | 0.247 | 0.04 | 4.08E-12 | 08_mo-DC3 |
| GSDME | 2.916071503 | 0.272 | 0.051 | 4.10E-12 | 08_mo-DC3 |
| GSTM3 | 2.556953846 | 0.383 | 0.099 | 5.04E-12 | 08_mo-DC3 |
| CD276 | 2.592987455 | 0.284 | 0.054 | 5.37E-12 | 08_mo-DC3 |
| ARHGAP22 | 2.162912849 | 0.556 | 0.2 | 7.29E-12 | 08_mo-DC3 |
| SRGAP2 | 1.659076514 | 0.84 | 0.581 | 7.38E-12 | 08_mo-DC3 |
| VSIR | 1.539619003 | 0.84 | 0.493 | 7.69E-12 | 08_mo-DC3 |
| SAMD4A | 3.196853867 | 0.272 | 0.052 | 8.44E-12 | 08_mo-DC3 |
| RIN2 | 1.81550127 | 0.568 | 0.208 | 8.71E-12 | 08_mo-DC3 |
| MAN1A1 | 1.480239735 | 0.827 | 0.503 | 9.10E-12 | 08_mo-DC3 |
| ZBTB16 | 1.283703072 | 0.827 | 0.47 | 9.52E-12 | 08_mo-DC3 |
| CRYBG1 | -3.71790814 | 0.148 | 0.608 | 9.82E-12 | 08_mo-DC3 |
| IL1R2 | -8.82065385 | 0.012 | 0.502 | 1.11E-11 | 08_mo-DC3 |
| PEAK1 | 1.767065198 | 0.704 | 0.331 | 1.12E-11 | 08_mo-DC3 |
| SH3BGRL3 | -1.71853351 | 0.617 | 0.84 | 1.18E-11 | 08_mo-DC3 |
| QKI | 1.48317925 | 0.914 | 0.696 | 1.34E-11 | 08_mo-DC3 |
| MAF | 2.224215717 | 0.284 | 0.056 | 2.45E-11 | 08_mo-DC3 |
| ADAM17 | 1.419042704 | 0.802 | 0.415 | 3.13E-11 | 08_mo-DC3 |
| IRS2 | 1.409309527 | 0.741 | 0.349 | 3.26E-11 | 08_mo-DC3 |
| MEF2A | 1.030700268 | 0.963 | 0.733 | 3.71E-11 | 08_mo-DC3 |
| PPA1 | -2.96478187 | 0.21 | 0.632 | 6.16E-11 | 08_mo-DC3 |
| CHSY1 | 1.940287915 | 0.667 | 0.314 | 7.21E-11 | 08_mo-DC3 |
| TFRC | 1.669122497 | 0.605 | 0.249 | 9.52E-11 | 08_mo-DC3 |
| FOSB | 1.321956641 | 0.938 | 0.734 | 1.03E-10 | 08_mo-DC3 |
| SRGAP2C | 1.753833187 | 0.642 | 0.294 | 1.33E-10 | 08_mo-DC3 |
| CD81 | 1.354844147 | 0.901 | 0.664 | 1.39E-10 | 08_mo-DC3 |
| C15orf48 | -5.50031312 | 0.074 | 0.529 | 1.49E-10 | 08_mo-DC3 |
| FCER1A | -4.99969545 | 0.025 | 0.497 | 1.50E-10 | 08_mo-DC3 |
| ADGRG1 | 2.282345799 | 0.395 | 0.112 | 1.90E-10 | 08_mo-DC3 |
| SRGAP2B | 1.359000529 | 0.79 | 0.419 | 2.20E-10 | 08_mo-DC3 |
| STARD13 | 2.133096312 | 0.383 | 0.103 | 2.31E-10 | 08_mo-DC3 |
| CD69 | 1.583447935 | 0.765 | 0.419 | 2.37E-10 | 08_mo-DC3 |
| EGR2 | 2.656731039 | 0.395 | 0.115 | 2.50E-10 | 08_mo-DC3 |
| MGAT5 | 1.910965841 | 0.593 | 0.232 | 3.14E-10 | 08_mo-DC3 |
| ITSN1 | 1.754132956 | 0.63 | 0.271 | 3.47E-10 | 08_mo-DC3 |
| LPAR5 | 2.390247185 | 0.346 | 0.089 | 3.86E-10 | 08_mo-DC3 |
| GNAQ | 1.050087002 | 0.938 | 0.642 | 4.00E-10 | 08_mo-DC3 |
| FMN1 | 1.447347783 | 0.889 | 0.571 | 6.14E-10 | 08_mo-DC3 |
| OTUD1 | 1.779811176 | 0.568 | 0.231 | 7.61E-10 | 08_mo-DC3 |

| | | | | | |
|---|---|---|---|---|---|
| RBMS3 | 1.885140876 | 0.333 | 0.083 | 9.81E-10 | 08_mo-DC3 |
| CD84 | 1.411849754 | 0.63 | 0.262 | 1.08E-09 | 08_mo-DC3 |
| DOP1B | 1.77518531 | 0.432 | 0.13 | 1.09E-09 | 08_mo-DC3 |
| CTDSP1 | 1.74208412 | 0.58 | 0.241 | 1.11E-09 | 08_mo-DC3 |
| SPRY2 | 2.188493752 | 0.346 | 0.09 | 1.25E-09 | 08_mo-DC3 |
| SPATS2L | 1.824909676 | 0.469 | 0.165 | 1.38E-09 | 08_mo-DC3 |
| VSIG4 | 1.475899124 | 0.728 | 0.372 | 1.75E-09 | 08_mo-DC3 |
| RALA | -2.69715161 | 0.173 | 0.608 | 1.85E-09 | 08_mo-DC3 |
| MFSD1 | 1.145582254 | 0.84 | 0.573 | 2.08E-09 | 08_mo-DC3 |
| MARCKS | 1.018193246 | 0.877 | 0.519 | 2.30E-09 | 08_mo-DC3 |
| SLC19A2 | 2.132332512 | 0.481 | 0.181 | 2.33E-09 | 08_mo-DC3 |
| RANBP9 | 1.877285357 | 0.642 | 0.302 | 2.73E-09 | 08_mo-DC3 |
| CRIP1 | -6.43842811 | 0.025 | 0.461 | 2.81E-09 | 08_mo-DC3 |
| EMP3 | -2.19672206 | 0.321 | 0.707 | 3.03E-09 | 08_mo-DC3 |
| LIMK2 | 1.689367288 | 0.494 | 0.177 | 3.12E-09 | 08_mo-DC3 |
| SPATA6 | 2.628461938 | 0.346 | 0.096 | 4.70E-09 | 08_mo-DC3 |
| RERE | 1.582267773 | 0.716 | 0.374 | 5.17E-09 | 08_mo-DC3 |
| MS4A4A | 1.24019167 | 0.704 | 0.382 | 5.24E-09 | 08_mo-DC3 |
| PLCL1 | 2.689526595 | 0.272 | 0.061 | 6.00E-09 | 08_mo-DC3 |
| TNFAIP8L3 | 2.655733281 | 0.358 | 0.106 | 6.01E-09 | 08_mo-DC3 |
| GPATCH11 | 2.346664097 | 0.284 | 0.066 | 6.98E-09 | 08_mo-DC3 |
| DIP2A | 1.808240975 | 0.407 | 0.126 | 7.06E-09 | 08_mo-DC3 |
| TMEM176B | 1.461022689 | 0.531 | 0.2 | 7.81E-09 | 08_mo-DC3 |
| ANKS1A | 1.664088582 | 0.679 | 0.349 | 8.33E-09 | 08_mo-DC3 |
| GSTP1 | -1.70560781 | 0.469 | 0.76 | 9.73E-09 | 08_mo-DC3 |
| CST7 | -3.87313964 | 0.086 | 0.511 | 1.14E-08 | 08_mo-DC3 |
| RHBDF2 | 1.184661426 | 0.802 | 0.524 | 1.14E-08 | 08_mo-DC3 |
| DSCAM | 2.597316521 | 0.296 | 0.073 | 1.28E-08 | 08_mo-DC3 |
| FHIT | 1.569979689 | 0.654 | 0.307 | 1.29E-08 | 08_mo-DC3 |
| SLC29A1 | 1.791689617 | 0.395 | 0.121 | 1.42E-08 | 08_mo-DC3 |
| FCGR3A | 1.572780623 | 0.667 | 0.323 | 1.55E-08 | 08_mo-DC3 |
| TIMP2 | 1.382155216 | 0.654 | 0.328 | 1.83E-08 | 08_mo-DC3 |
| GPAT3 | -3.49988444 | 0.049 | 0.482 | 2.10E-08 | 08_mo-DC3 |
| ITGB5 | 2.322074264 | 0.259 | 0.057 | 2.12E-08 | 08_mo-DC3 |
| LGALS1 | -2.25368706 | 0.321 | 0.685 | 3.01E-08 | 08_mo-DC3 |
| NCK2 | 1.558645372 | 0.494 | 0.184 | 3.15E-08 | 08_mo-DC3 |
| MAML2 | 1.433439287 | 0.704 | 0.374 | 3.18E-08 | 08_mo-DC3 |
| USP4 | 1.464573789 | 0.679 | 0.368 | 3.29E-08 | 08_mo-DC3 |
| FCGR1A | 1.456861777 | 0.642 | 0.293 | 4.00E-08 | 08_mo-DC3 |
| SPATA13 | 1.419349977 | 0.481 | 0.172 | 4.33E-08 | 08_mo-DC3 |
| NHSL1 | 1.474918491 | 0.753 | 0.43 | 5.75E-08 | 08_mo-DC3 |
| CYTIP | -2.39512166 | 0.222 | 0.613 | 5.84E-08 | 08_mo-DC3 |
| OTULINL | 1.563366007 | 0.58 | 0.267 | 6.24E-08 | 08_mo-DC3 |
| TRPM2 | 1.32754582 | 0.519 | 0.194 | 6.59E-08 | 08_mo-DC3 |
| FMNL2 | 1.261834307 | 0.84 | 0.55 | 6.90E-08 | 08_mo-DC3 |
| WASF2 | 1.246989664 | 0.802 | 0.518 | 7.06E-08 | 08_mo-DC3 |

| | | | | | |
|---|---|---|---|---|---|
| MEF2C | 1.089640757 | 0.926 | 0.639 | 7.60E-08 | 08_mo-DC3 |
| AP1B1 | 1.194561128 | 0.716 | 0.389 | 8.04E-08 | 08_mo-DC3 |
| OSBPL1A | 1.83753615 | 0.321 | 0.087 | 8.49E-08 | 08_mo-DC3 |
| ADAMTSL4-AS | 1.185081319 | 0.741 | 0.417 | 8.54E-08 | 08_mo-DC3 |
| LNCAROD | 1.786483558 | 0.469 | 0.172 | 8.80E-08 | 08_mo-DC3 |
| WWOX | 1.621641922 | 0.556 | 0.244 | 1.05E-07 | 08_mo-DC3 |
| HTRA1 | 1.627352631 | 0.457 | 0.163 | 1.09E-07 | 08_mo-DC3 |
| MYL12A | -1.5744784 | 0.469 | 0.742 | 1.16E-07 | 08_mo-DC3 |
| RASA3 | 1.61025339 | 0.568 | 0.259 | 1.21E-07 | 08_mo-DC3 |
| ITPR2 | 1.545005068 | 0.704 | 0.38 | 1.67E-07 | 08_mo-DC3 |
| BCL3 | -3.5482951 | 0.111 | 0.496 | 2.02E-07 | 08_mo-DC3 |
| PADI2 | 1.458275397 | 0.79 | 0.541 | 2.22E-07 | 08_mo-DC3 |
| CLEC10A | -13.685257 | 0 | 0.388 | 2.27E-07 | 08_mo-DC3 |
| SIPA1L2 | 1.798165844 | 0.432 | 0.152 | 2.35E-07 | 08_mo-DC3 |
| FLNA | -4.44820816 | 0.037 | 0.429 | 2.53E-07 | 08_mo-DC3 |
| DEPTOR | 1.6391903 | 0.42 | 0.145 | 2.68E-07 | 08_mo-DC3 |
| SNX5 | 1.344147573 | 0.593 | 0.286 | 3.12E-07 | 08_mo-DC3 |
| CHKA | 1.456167207 | 0.704 | 0.412 | 3.33E-07 | 08_mo-DC3 |
| PDE8A | 1.118874744 | 0.741 | 0.418 | 3.37E-07 | 08_mo-DC3 |
| CNN2 | -3.26525473 | 0.086 | 0.474 | 3.45E-07 | 08_mo-DC3 |
| DOCK11 | 1.622766793 | 0.519 | 0.216 | 4.03E-07 | 08_mo-DC3 |
| AFF4 | 1.081577789 | 0.852 | 0.625 | 4.20E-07 | 08_mo-DC3 |
| RPL7 | -1.41550027 | 0.556 | 0.764 | 4.67E-07 | 08_mo-DC3 |
| EGR1 | 1.713862927 | 0.593 | 0.279 | 4.85E-07 | 08_mo-DC3 |
| EVI5 | 1.358460972 | 0.543 | 0.235 | 5.03E-07 | 08_mo-DC3 |
| CCL3L1 | 1.870204617 | 0.605 | 0.301 | 5.46E-07 | 08_mo-DC3 |
| CTSC | 1.230039082 | 0.716 | 0.412 | 5.72E-07 | 08_mo-DC3 |
| MMP2 | 2.006292407 | 0.321 | 0.093 | 5.88E-07 | 08_mo-DC3 |
| SKAP2 | 1.190491642 | 0.679 | 0.37 | 6.28E-07 | 08_mo-DC3 |
| NKTR | 1.242588645 | 0.58 | 0.24 | 6.58E-07 | 08_mo-DC3 |
| GALNT10 | 1.547680541 | 0.395 | 0.132 | 8.73E-07 | 08_mo-DC3 |
| FMNL3 | 1.075920318 | 0.667 | 0.318 | 9.83E-07 | 08_mo-DC3 |
| DUSP5 | -4.13424109 | 0.086 | 0.456 | 1.04E-06 | 08_mo-DC3 |
| AHNAK | -1.74795048 | 0.222 | 0.615 | 1.14E-06 | 08_mo-DC3 |
| MACF1 | 1.049120307 | 0.753 | 0.445 | 1.21E-06 | 08_mo-DC3 |
| KLF7 | 1.632530584 | 0.494 | 0.208 | 1.36E-06 | 08_mo-DC3 |
| HINT1 | -1.6843377 | 0.42 | 0.691 | 1.37E-06 | 08_mo-DC3 |
| PAPOLG | 1.717187673 | 0.568 | 0.288 | 1.45E-06 | 08_mo-DC3 |
| ARHGAP21 | 1.273268329 | 0.543 | 0.233 | 1.56E-06 | 08_mo-DC3 |
| CCSER1 | -3.06845738 | 0.099 | 0.482 | 1.57E-06 | 08_mo-DC3 |
| EXOC4 | 1.480854342 | 0.642 | 0.334 | 1.60E-06 | 08_mo-DC3 |
| LPAR1 | 1.997132932 | 0.272 | 0.071 | 1.66E-06 | 08_mo-DC3 |
| DISP1 | 2.218474215 | 0.272 | 0.072 | 1.85E-06 | 08_mo-DC3 |
| LINC02256 | 1.175985176 | 0.605 | 0.315 | 2.58E-06 | 08_mo-DC3 |
| USP6NL | 2.136872101 | 0.346 | 0.115 | 2.79E-06 | 08_mo-DC3 |
| CD1C | -13.9504206 | 0 | 0.357 | 3.13E-06 | 08_mo-DC3 |

| | | | | | |
|---|---|---|---|---|---|
| CAMK2D | 1.167613593 | 0.506 | 0.21 | 3.71E-06 | 08_mo-DC3 |
| ATF3 | 1.093717333 | 0.84 | 0.567 | 4.02E-06 | 08_mo-DC3 |
| JAML | -4.30004271 | 0.025 | 0.386 | 4.05E-06 | 08_mo-DC3 |
| AL133415.1 | -4.19712326 | 0.037 | 0.398 | 4.08E-06 | 08_mo-DC3 |
| PARP8 | 1.572558312 | 0.494 | 0.205 | 4.19E-06 | 08_mo-DC3 |
| DHRS7 | 1.277840423 | 0.679 | 0.401 | 4.47E-06 | 08_mo-DC3 |
| SLC25A37 | 1.121260882 | 0.667 | 0.356 | 4.61E-06 | 08_mo-DC3 |
| LGALS3 | -2.90683974 | 0.123 | 0.487 | 5.21E-06 | 08_mo-DC3 |
| LCP2 | 1.18856812 | 0.84 | 0.565 | 5.36E-06 | 08_mo-DC3 |
| SSH2 | 1.000694457 | 0.815 | 0.568 | 5.48E-06 | 08_mo-DC3 |
| ZFHX3 | 1.3708629 | 0.667 | 0.387 | 5.89E-06 | 08_mo-DC3 |
| SLC44A2 | 2.227446673 | 0.333 | 0.112 | 6.06E-06 | 08_mo-DC3 |
| SERPINB1 | -1.92623217 | 0.457 | 0.727 | 6.41E-06 | 08_mo-DC3 |
| RBM47 | 1.153860696 | 0.84 | 0.629 | 6.50E-06 | 08_mo-DC3 |
| CASS4 | 1.508296006 | 0.556 | 0.274 | 6.81E-06 | 08_mo-DC3 |
| NEDD9 | 1.397560234 | 0.765 | 0.507 | 7.77E-06 | 08_mo-DC3 |
| JUN | 1.281870462 | 0.901 | 0.682 | 8.08E-06 | 08_mo-DC3 |
| CYTH3 | 1.598713919 | 0.481 | 0.205 | 8.65E-06 | 08_mo-DC3 |
| ZNF330 | 1.77071047 | 0.333 | 0.11 | 8.79E-06 | 08_mo-DC3 |
| LINC01480 | 1.678818415 | 0.37 | 0.134 | 9.65E-06 | 08_mo-DC3 |
| TAGLN2 | -1.75689756 | 0.432 | 0.696 | 1.00E-05 | 08_mo-DC3 |
| KYNU | -2.13114607 | 0.321 | 0.636 | 1.03E-05 | 08_mo-DC3 |
| SRGAP1 | 1.018837979 | 0.84 | 0.553 | 1.03E-05 | 08_mo-DC3 |
| SCPEP1 | 1.211424483 | 0.568 | 0.276 | 1.14E-05 | 08_mo-DC3 |
| PPARG | 1.770221372 | 0.37 | 0.128 | 1.16E-05 | 08_mo-DC3 |
| TM6SF1 | 1.362753364 | 0.617 | 0.356 | 1.28E-05 | 08_mo-DC3 |
| THBS1 | -4.51108199 | 0.037 | 0.382 | 1.57E-05 | 08_mo-DC3 |
| LDHA | -1.60118511 | 0.506 | 0.736 | 1.60E-05 | 08_mo-DC3 |
| CITED2 | 2.098228375 | 0.457 | 0.198 | 1.60E-05 | 08_mo-DC3 |
| RPS29 | -1.21787072 | 0.506 | 0.752 | 1.69E-05 | 08_mo-DC3 |
| SLC15A4 | 1.483696828 | 0.506 | 0.223 | 1.96E-05 | 08_mo-DC3 |
| RPL31 | -1.16771612 | 0.568 | 0.783 | 2.00E-05 | 08_mo-DC3 |
| TNS3 | 1.361956213 | 0.519 | 0.237 | 2.02E-05 | 08_mo-DC3 |
| CTR9 | 1.641383119 | 0.358 | 0.126 | 2.38E-05 | 08_mo-DC3 |
| PTPRJ | 1.123430274 | 0.691 | 0.376 | 2.42E-05 | 08_mo-DC3 |
| PDGFB | 1.103832661 | 0.691 | 0.36 | 2.56E-05 | 08_mo-DC3 |
| RNF122 | 2.004718929 | 0.309 | 0.1 | 3.45E-05 | 08_mo-DC3 |
| ULK3 | 1.911082972 | 0.296 | 0.093 | 3.48E-05 | 08_mo-DC3 |
| EIF3K | -1.38522993 | 0.481 | 0.71 | 3.60E-05 | 08_mo-DC3 |
| ISG20 | -1.99615843 | 0.185 | 0.541 | 3.77E-05 | 08_mo-DC3 |
| G0S2 | -4.76354448 | 0.099 | 0.427 | 3.86E-05 | 08_mo-DC3 |
| LAIR1 | 1.214151385 | 0.679 | 0.405 | 3.99E-05 | 08_mo-DC3 |
| CYBB | 1.336930598 | 0.679 | 0.449 | 4.34E-05 | 08_mo-DC3 |
| CD44 | -1.72111098 | 0.42 | 0.706 | 4.36E-05 | 08_mo-DC3 |
| AFF3 | -3.0534827 | 0.123 | 0.459 | 4.38E-05 | 08_mo-DC3 |
| IRF4 | -13.4188977 | 0 | 0.322 | 4.88E-05 | 08_mo-DC3 |

| | | | | | |
|---|---|---|---|---|---|
| AAK1 | 1.051498813 | 0.519 | 0.239 | 5.25E-05 | 08_mo-DC3 |
| OLFML3 | 1.375973899 | 0.444 | 0.181 | 5.92E-05 | 08_mo-DC3 |
| SORT1 | 1.687862555 | 0.457 | 0.207 | 6.04E-05 | 08_mo-DC3 |
| INSIG1 | -1.99744916 | 0.284 | 0.599 | 6.15E-05 | 08_mo-DC3 |
| TRAF1 | -5.4880369 | 0.025 | 0.348 | 6.16E-05 | 08_mo-DC3 |
| 3-Mar | 1.316951274 | 0.556 | 0.282 | 6.19E-05 | 08_mo-DC3 |
| NFIC | 1.442524518 | 0.531 | 0.266 | 7.29E-05 | 08_mo-DC3 |
| NUFIP2 | 1.006517984 | 0.704 | 0.416 | 7.56E-05 | 08_mo-DC3 |
| MAP3K5 | 1.151266312 | 0.457 | 0.191 | 7.58E-05 | 08_mo-DC3 |
| ATM | 1.208563151 | 0.506 | 0.232 | 8.03E-05 | 08_mo-DC3 |
| SLC2A13 | 1.400434242 | 0.37 | 0.136 | 8.21E-05 | 08_mo-DC3 |
| RAB3GAP2 | 1.59817308 | 0.346 | 0.121 | 8.35E-05 | 08_mo-DC3 |
| MIR29B2CHG | 1.299537723 | 0.395 | 0.149 | 8.77E-05 | 08_mo-DC3 |
| FRMD4B | 1.413340584 | 0.556 | 0.273 | 9.19E-05 | 08_mo-DC3 |
| ADAM8 | -12.6283143 | 0 | 0.313 | 9.20E-05 | 08_mo-DC3 |
| PLK3 | 1.165068502 | 0.716 | 0.458 | 9.36E-05 | 08_mo-DC3 |
| SATB1 | -3.15108076 | 0.074 | 0.403 | 9.74E-05 | 08_mo-DC3 |
| DENND3 | 1.122367036 | 0.654 | 0.402 | 0.0001062 | 08_mo-DC3 |
| RAB1A | 1.030784607 | 0.815 | 0.569 | 0.0001076 | 08_mo-DC3 |
| IL1R1 | -13.2609327 | 0 | 0.311 | 0.0001077 | 08_mo-DC3 |
| SH3TC1 | 1.025582234 | 0.741 | 0.53 | 0.0001134 | 08_mo-DC3 |
| BHLHE41 | 1.523241643 | 0.506 | 0.242 | 0.0001219 | 08_mo-DC3 |
| PUM1 | 1.083355354 | 0.617 | 0.339 | 0.0001426 | 08_mo-DC3 |
| HIVEP3 | 1.637356402 | 0.346 | 0.124 | 0.0001464 | 08_mo-DC3 |
| FAM110B | 1.361958789 | 0.395 | 0.153 | 0.0001538 | 08_mo-DC3 |
| IL18R1 | -4.08176537 | 0.049 | 0.36 | 0.0001735 | 08_mo-DC3 |
| PSME4 | 1.201432817 | 0.605 | 0.316 | 0.0001879 | 08_mo-DC3 |
| CD14 | 1.369151593 | 0.679 | 0.427 | 0.0002009 | 08_mo-DC3 |
| FAM149A | 1.233102509 | 0.469 | 0.206 | 0.0002295 | 08_mo-DC3 |
| CKLF | -1.53892262 | 0.321 | 0.639 | 0.0002586 | 08_mo-DC3 |
| CCNI | -1.07386182 | 0.556 | 0.781 | 0.0002652 | 08_mo-DC3 |
| SORL1 | 1.018371379 | 0.728 | 0.448 | 0.0002727 | 08_mo-DC3 |
| MPZL1 | 1.793852667 | 0.333 | 0.126 | 0.0002842 | 08_mo-DC3 |
| FCGR2B | -1.83964012 | 0.272 | 0.574 | 0.0002942 | 08_mo-DC3 |
| CCR7 | -5.83538741 | 0.025 | 0.325 | 0.0003187 | 08_mo-DC3 |
| RASSF4 | 1.434004294 | 0.654 | 0.421 | 0.0003373 | 08_mo-DC3 |
| MCOLN2 | -3.07973871 | 0.062 | 0.382 | 0.0003426 | 08_mo-DC3 |
| EREG | -5.12669019 | 0.012 | 0.315 | 0.0003564 | 08_mo-DC3 |
| KIF1B | 1.609546546 | 0.519 | 0.272 | 0.0003596 | 08_mo-DC3 |
| TNRC18 | 1.098425127 | 0.519 | 0.239 | 0.0003628 | 08_mo-DC3 |
| LRP1 | 1.370334396 | 0.432 | 0.188 | 0.0003942 | 08_mo-DC3 |
| CHN2 | 1.767887754 | 0.321 | 0.114 | 0.0003977 | 08_mo-DC3 |
| CLASP1 | 1.303617246 | 0.407 | 0.169 | 0.0004287 | 08_mo-DC3 |
| SELPLG | 1.737533402 | 0.444 | 0.203 | 0.0004433 | 08_mo-DC3 |
| ICAM3 | -5.03511718 | 0.012 | 0.311 | 0.0004477 | 08_mo-DC3 |
| TBC1D12 | 1.101885047 | 0.506 | 0.236 | 0.0004702 | 08_mo-DC3 |

| | | | | | |
|---|---|---|---|---|---|
| PRKCA | 1.452604101 | 0.358 | 0.138 | 0.0004727 | 08_mo-DC3 |
| DUSP4 | -1.81115221 | 0.247 | 0.559 | 0.0005151 | 08_mo-DC3 |
| MAGT1 | 1.049790225 | 0.617 | 0.356 | 0.0005539 | 08_mo-DC3 |
| IFITM1 | -3.20176392 | 0.123 | 0.422 | 0.0005853 | 08_mo-DC3 |
| CIB1 | -1.39178218 | 0.259 | 0.579 | 0.0005996 | 08_mo-DC3 |
| PKIB | -2.85900017 | 0.123 | 0.425 | 0.0007003 | 08_mo-DC3 |
| UPP1 | -2.38375863 | 0.136 | 0.451 | 0.0007556 | 08_mo-DC3 |
| SNRPD2 | -1.6405977 | 0.235 | 0.55 | 0.0008211 | 08_mo-DC3 |
| FYN | -3.23351392 | 0.074 | 0.379 | 0.000832 | 08_mo-DC3 |
| ARHGEF7 | 1.270769969 | 0.506 | 0.258 | 0.0008793 | 08_mo-DC3 |
| ATG7 | 1.469049805 | 0.556 | 0.297 | 0.0009053 | 08_mo-DC3 |
| MYO1E | 1.775459457 | 0.506 | 0.275 | 0.0009456 | 08_mo-DC3 |
| TPRG1 | 1.72960114 | 0.321 | 0.119 | 0.0009468 | 08_mo-DC3 |
| RLF | 1.668780539 | 0.556 | 0.31 | 0.0009999 | 08_mo-DC3 |
| FER | 1.626887849 | 0.37 | 0.15 | 0.0010117 | 08_mo-DC3 |
| ARHGEF40 | 1.520582354 | 0.432 | 0.204 | 0.0010761 | 08_mo-DC3 |
| CYTOR | -2.12302069 | 0.074 | 0.38 | 0.0011822 | 08_mo-DC3 |
| BIRC3 | -2.13445451 | 0.099 | 0.402 | 0.0011862 | 08_mo-DC3 |
| TXN | -3.37881587 | 0.16 | 0.47 | 0.001188 | 08_mo-DC3 |
| SULF2 | -3.29311988 | 0.049 | 0.346 | 0.001196 | 08_mo-DC3 |
| ACIN1 | 1.230628959 | 0.469 | 0.225 | 0.0012002 | 08_mo-DC3 |
| WDR70 | 1.645730203 | 0.42 | 0.188 | 0.0012847 | 08_mo-DC3 |
| PCCA | 1.320961923 | 0.321 | 0.115 | 0.0012925 | 08_mo-DC3 |
| ZBTB20 | 1.013759802 | 0.642 | 0.369 | 0.0015068 | 08_mo-DC3 |
| ADAM19 | -5.74844511 | 0.025 | 0.303 | 0.0016098 | 08_mo-DC3 |
| CPEB4 | 1.527136197 | 0.469 | 0.241 | 0.0016549 | 08_mo-DC3 |
| TMEM156 | 1.728167523 | 0.407 | 0.186 | 0.0020251 | 08_mo-DC3 |
| MKLN1 | 1.847802837 | 0.63 | 0.415 | 0.0020387 | 08_mo-DC3 |
| DENND4C | 1.447079758 | 0.506 | 0.262 | 0.0020518 | 08_mo-DC3 |
| TMEM176A | 1.154488626 | 0.321 | 0.117 | 0.0020755 | 08_mo-DC3 |
| GPCPD1 | 1.077487007 | 0.667 | 0.414 | 0.0022432 | 08_mo-DC3 |
| LPIN2 | 1.548241372 | 0.494 | 0.259 | 0.002271 | 08_mo-DC3 |
| SPIB | -3.15248389 | 0.049 | 0.342 | 0.0022718 | 08_mo-DC3 |
| PCNX4 | 1.375878874 | 0.469 | 0.223 | 0.002328 | 08_mo-DC3 |
| CTTNBP2NL | 1.180334257 | 0.63 | 0.393 | 0.0024962 | 08_mo-DC3 |
| MB21D2 | 1.346111669 | 0.481 | 0.244 | 0.0025389 | 08_mo-DC3 |
| ZFAS1 | -1.15896291 | 0.407 | 0.691 | 0.0025674 | 08_mo-DC3 |
| KDM3B | 1.62214472 | 0.407 | 0.195 | 0.0026416 | 08_mo-DC3 |
| IL2RG | -2.67215325 | 0.074 | 0.368 | 0.0027287 | 08_mo-DC3 |
| AC138123.1 | -2.43012059 | 0.074 | 0.371 | 0.0030058 | 08_mo-DC3 |
| ZNF644 | 1.088933989 | 0.58 | 0.331 | 0.0030555 | 08_mo-DC3 |
| LAMP1 | 1.326409234 | 0.469 | 0.229 | 0.0031293 | 08_mo-DC3 |
| MIF | -1.27963427 | 0.531 | 0.742 | 0.0031648 | 08_mo-DC3 |
| CCL22 | -7.4930358 | 0.012 | 0.278 | 0.0033814 | 08_mo-DC3 |
| IGF2R | 1.00256888 | 0.346 | 0.137 | 0.0036033 | 08_mo-DC3 |
| LRCH1 | 1.034643627 | 0.481 | 0.237 | 0.0037421 | 08_mo-DC3 |

| | | | | | |
|---|---|---|---|---|---|
| OSBPL3 | 1.058445162 | 0.494 | 0.255 | 0.004041 | 08_mo-DC3 |
| PMEPA1 | 1.312816915 | 0.457 | 0.232 | 0.0041622 | 08_mo-DC3 |
| AC022217.3 | 1.175772437 | 0.494 | 0.26 | 0.0042124 | 08_mo-DC3 |
| FAM135A | 1.033633553 | 0.457 | 0.21 | 0.0042823 | 08_mo-DC3 |
| PDE4DIP | 1.035678156 | 0.58 | 0.326 | 0.0046637 | 08_mo-DC3 |
| GOLGB1 | 1.181000619 | 0.605 | 0.374 | 0.0055929 | 08_mo-DC3 |
| EML4 | 1.176269058 | 0.494 | 0.258 | 0.0067866 | 08_mo-DC3 |
| SPECC1 | 1.050296375 | 0.444 | 0.214 | 0.0068878 | 08_mo-DC3 |
| ZNF609 | 1.274602897 | 0.494 | 0.253 | 0.0071133 | 08_mo-DC3 |
| WDFY3 | 1.262729942 | 0.407 | 0.183 | 0.0071623 | 08_mo-DC3 |
| MYO1G | -2.9827893 | 0.037 | 0.315 | 0.0071841 | 08_mo-DC3 |
| CPEB3 | 1.123205715 | 0.383 | 0.161 | 0.0074767 | 08_mo-DC3 |
| AP2S1 | -1.54980112 | 0.259 | 0.525 | 0.0075036 | 08_mo-DC3 |
| DIP2B | 1.308916008 | 0.444 | 0.223 | 0.0075218 | 08_mo-DC3 |
| SPINT2 | -1.59334213 | 0.259 | 0.532 | 0.0078369 | 08_mo-DC3 |
| FAM53B | 1.165301701 | 0.494 | 0.267 | 0.0083447 | 08_mo-DC3 |
| SMAD7 | 1.36164533 | 0.42 | 0.205 | 0.0085271 | 08_mo-DC3 |
| SLC16A10 | -2.53987103 | 0.136 | 0.423 | 0.0085276 | 08_mo-DC3 |
| MKNK1 | 1.347821734 | 0.407 | 0.19 | 0.0099184 | 08_mo-DC3 |
| TGFBR2 | 1.14197887 | 0.481 | 0.252 | 0.0103505 | 08_mo-DC3 |
| PKM | -1.23218947 | 0.469 | 0.688 | 0.0107959 | 08_mo-DC3 |
| CAPG | -1.17294085 | 0.444 | 0.675 | 0.0110286 | 08_mo-DC3 |
| TNFAIP8 | -2.32760945 | 0.173 | 0.447 | 0.0115659 | 08_mo-DC3 |
| SBF2 | 1.051861714 | 0.605 | 0.355 | 0.0126009 | 08_mo-DC3 |
| RHOF | -3.21134334 | 0.037 | 0.3 | 0.0134793 | 08_mo-DC3 |
| MIS18BP1 | 1.10183089 | 0.617 | 0.416 | 0.0134968 | 08_mo-DC3 |
| ETV3 | -1.56751891 | 0.284 | 0.566 | 0.0136833 | 08_mo-DC3 |
| LIPA | 1.298794337 | 0.444 | 0.23 | 0.0138975 | 08_mo-DC3 |
| PPIF | -2.42693589 | 0.198 | 0.45 | 0.0140641 | 08_mo-DC3 |
| TPI1 | -1.23745614 | 0.481 | 0.697 | 0.0141021 | 08_mo-DC3 |
| RPS6KA2 | 1.05513511 | 0.444 | 0.211 | 0.0147396 | 08_mo-DC3 |
| SNHG29 | -1.21144944 | 0.457 | 0.675 | 0.0159505 | 08_mo-DC3 |
| TBRG1 | 1.137787689 | 0.531 | 0.308 | 0.0160073 | 08_mo-DC3 |
| FEZ2 | 1.171039701 | 0.432 | 0.211 | 0.0163914 | 08_mo-DC3 |
| SETX | 1.020353431 | 0.494 | 0.257 | 0.0166104 | 08_mo-DC3 |
| RELCH | 1.411671462 | 0.346 | 0.145 | 0.0166594 | 08_mo-DC3 |
| EIF3E | -1.43225995 | 0.321 | 0.56 | 0.0167939 | 08_mo-DC3 |
| COX5A | -1.3882618 | 0.321 | 0.585 | 0.017656 | 08_mo-DC3 |
| GAA | 1.163715663 | 0.494 | 0.283 | 0.0209903 | 08_mo-DC3 |
| RFX2 | 1.130177706 | 0.494 | 0.26 | 0.0214743 | 08_mo-DC3 |
| SCAMP2 | 1.00073889 | 0.593 | 0.373 | 0.0215407 | 08_mo-DC3 |
| VDR | -3.35848729 | 0.049 | 0.308 | 0.0216418 | 08_mo-DC3 |
| PSMA6 | -1.54498237 | 0.222 | 0.501 | 0.0216662 | 08_mo-DC3 |
| FOXN2 | -2.05978833 | 0.16 | 0.44 | 0.022535 | 08_mo-DC3 |
| COMMD6 | -1.21302872 | 0.432 | 0.642 | 0.0255872 | 08_mo-DC3 |
| MIR4435-2HG | -3.40107104 | 0.062 | 0.309 | 0.032706 | 08_mo-DC3 |

| | | | | | |
|---|---|---|---|---|---|
| PIK3AP1 | 1.011533708 | 0.432 | 0.214 | 0.0330989 | 08_mo-DC3 |
| DAPP1 | -3.81630743 | 0.062 | 0.304 | 0.0361353 | 08_mo-DC3 |
| ANXA1 | -1.14331358 | 0.333 | 0.602 | 0.0416696 | 08_mo-DC3 |
| CD1E | -12.6841228 | 0 | 0.225 | 0.0421204 | 08_mo-DC3 |
| PSME1 | -1.37417606 | 0.346 | 0.586 | 0.0488072 | 08_mo-DC3 |
| TXNRD1 | -2.07300713 | 0.16 | 0.418 | 0.0493479 | 08_mo-DC3 |
| XYLT1 | -1.95055045 | 0.222 | 0.484 | 0.0520958 | 08_mo-DC3 |
| HIPK3 | 1.259724005 | 0.568 | 0.362 | 0.0531048 | 08_mo-DC3 |
| STOM | 1.190364345 | 0.407 | 0.205 | 0.0531469 | 08_mo-DC3 |
| GPR157 | -3.89148783 | 0.037 | 0.276 | 0.0538252 | 08_mo-DC3 |
| SEC63 | 1.027557824 | 0.407 | 0.197 | 0.0555576 | 08_mo-DC3 |
| UXT | -1.16839433 | 0.198 | 0.472 | 0.0566663 | 08_mo-DC3 |
| SKI | 1.234574604 | 0.481 | 0.276 | 0.0629585 | 08_mo-DC3 |
| NT5C2 | 1.075047985 | 0.506 | 0.285 | 0.0629699 | 08_mo-DC3 |
| PRKCD | 1.059298881 | 0.457 | 0.24 | 0.0638755 | 08_mo-DC3 |
| PLP2 | -2.11057621 | 0.123 | 0.382 | 0.0643916 | 08_mo-DC3 |
| C20orf194 | 1.179253414 | 0.407 | 0.201 | 0.0677797 | 08_mo-DC3 |
| HLA-DQB2 | -2.68766556 | 0.099 | 0.344 | 0.0687166 | 08_mo-DC3 |
| GAK | 1.085646813 | 0.457 | 0.249 | 0.0722605 | 08_mo-DC3 |
| PIM3 | -1.82403286 | 0.222 | 0.47 | 0.0746569 | 08_mo-DC3 |
| NCOA2 | 1.106365764 | 0.432 | 0.213 | 0.0810141 | 08_mo-DC3 |
| GNA15 | -1.533887 | 0.222 | 0.484 | 0.0822797 | 08_mo-DC3 |
| CTSA | 1.044257498 | 0.444 | 0.239 | 0.0957318 | 08_mo-DC3 |
| THBD | -3.35623985 | 0.074 | 0.306 | 0.0966785 | 08_mo-DC3 |
| CD48 | -1.93820664 | 0.111 | 0.359 | 0.1008001 | 08_mo-DC3 |
| ARHGEF6 | 1.031049188 | 0.395 | 0.192 | 0.1140797 | 08_mo-DC3 |
| PPDPF | -1.23224161 | 0.383 | 0.597 | 0.1198205 | 08_mo-DC3 |
| ARF6 | -1.41639547 | 0.395 | 0.609 | 0.1243323 | 08_mo-DC3 |
| CALCRL | -13.0972001 | 0 | 0.207 | 0.1334234 | 08_mo-DC3 |
| CDK14 | -1.81963281 | 0.148 | 0.41 | 0.1334865 | 08_mo-DC3 |
| FABP5 | -3.36735847 | 0.111 | 0.341 | 0.1371913 | 08_mo-DC3 |
| SLC38A1 | -3.03329549 | 0.037 | 0.263 | 0.1496834 | 08_mo-DC3 |
| CFP | -12.1150631 | 0 | 0.205 | 0.1520691 | 08_mo-DC3 |
| ETHE1 | -2.30871949 | 0.074 | 0.311 | 0.1522595 | 08_mo-DC3 |
| CSTA | -2.92523441 | 0.025 | 0.249 | 0.1638037 | 08_mo-DC3 |
| DOK2 | -11.8573339 | 0 | 0.204 | 0.165884 | 08_mo-DC3 |
| MXD1 | -1.7684221 | 0.272 | 0.498 | 0.1704494 | 08_mo-DC3 |
| PLEKHG2 | 1.028074198 | 0.568 | 0.356 | 0.1826537 | 08_mo-DC3 |
| ARL4C | -1.82175065 | 0.259 | 0.491 | 0.1827733 | 08_mo-DC3 |
| TES | -1.77784989 | 0.222 | 0.452 | 0.1829153 | 08_mo-DC3 |
| RAC2 | -2.81929893 | 0.062 | 0.288 | 0.1851322 | 08_mo-DC3 |
| FRY | -3.26648279 | 0.049 | 0.271 | 0.2021495 | 08_mo-DC3 |
| VPS37B | 1.084205353 | 0.506 | 0.304 | 0.2103036 | 08_mo-DC3 |
| IL7R | -4.79826632 | 0.037 | 0.248 | 0.2314736 | 08_mo-DC3 |
| SOAT1 | 1.062025333 | 0.556 | 0.342 | 0.236399 | 08_mo-DC3 |
| TOMM7 | -1.06275299 | 0.42 | 0.643 | 0.260365 | 08_mo-DC3 |

| | | | | | |
|---|---|---|---|---|---|
| FKBP1A | -1.32735651 | 0.247 | 0.509 | 0.2626633 | 08_mo-DC3 |
| PLEKHA5 | -3.03615981 | 0.062 | 0.282 | 0.2778777 | 08_mo-DC3 |
| MARK3 | 1.026740391 | 0.481 | 0.28 | 0.2780451 | 08_mo-DC3 |
| DUSP2 | -1.37898425 | 0.407 | 0.613 | 0.3105296 | 08_mo-DC3 |
| RUNX3 | -2.41956723 | 0.099 | 0.326 | 0.3426177 | 08_mo-DC3 |
| CYP2S1 | -2.278546 | 0.086 | 0.318 | 0.3481008 | 08_mo-DC3 |
| CD109 | -2.19973798 | 0.062 | 0.292 | 0.3573309 | 08_mo-DC3 |
| BANP | -1.99841967 | 0.185 | 0.41 | 0.3906213 | 08_mo-DC3 |
| FBL | -1.66438929 | 0.086 | 0.317 | 0.4100829 | 08_mo-DC3 |
| RARA | -1.7135611 | 0.136 | 0.375 | 0.4622905 | 08_mo-DC3 |
| OPN3 | -2.36888857 | 0.086 | 0.311 | 0.4719592 | 08_mo-DC3 |
| HIVEP2 | -1.50210037 | 0.321 | 0.537 | 0.5388819 | 08_mo-DC3 |
| GDI2 | -1.15508443 | 0.296 | 0.535 | 0.5521389 | 08_mo-DC3 |
| SNX29 | 1.053272532 | 0.494 | 0.294 | 0.5680633 | 08_mo-DC3 |
| ZMYM2 | 1.060121373 | 0.617 | 0.41 | 0.5967596 | 08_mo-DC3 |
| GNA12 | -1.26967669 | 0.222 | 0.463 | 0.6755505 | 08_mo-DC3 |
| SATB1-AS1 | -3.13736249 | 0.037 | 0.239 | 0.6877103 | 08_mo-DC3 |
| CAPN2 | -2.93020549 | 0.025 | 0.225 | 0.6939121 | 08_mo-DC3 |
| SNRPB | -1.06855901 | 0.259 | 0.514 | 0.8061684 | 08_mo-DC3 |
| APRT | -1.0634522 | 0.309 | 0.548 | 0.8133731 | 08_mo-DC3 |
| APOO | -2.35940013 | 0.136 | 0.343 | 0.8695362 | 08_mo-DC3 |
| SNHG15 | -1.4351613 | 0.259 | 0.486 | 0.9055296 | 08_mo-DC3 |
| RHOC | -1.47395141 | 0.198 | 0.437 | 0.9386071 | 08_mo-DC3 |
| PDCL3 | -2.43416852 | 0.099 | 0.305 | 0.9596726 | 08_mo-DC3 |
| LPXN | -1.66697842 | 0.222 | 0.442 | 1 | 08_mo-DC3 |
| FLT3 | -2.40442295 | 0.086 | 0.301 | 1 | 08_mo-DC3 |
| CDK2AP1 | -1.20519896 | 0.333 | 0.551 | 1 | 08_mo-DC3 |
| PARK7 | -1.13320756 | 0.296 | 0.531 | 1 | 08_mo-DC3 |
| ST18 | -2.10951391 | 0.062 | 0.27 | 1 | 08_mo-DC3 |
| ACTN4 | -1.67257198 | 0.136 | 0.368 | 1 | 08_mo-DC3 |
| CARD16 | -1.91174927 | 0.123 | 0.336 | 1 | 08_mo-DC3 |
| STAT4 | -1.3966219 | 0.123 | 0.346 | 1 | 08_mo-DC3 |
| PTRHD1 | -1.79576902 | 0.099 | 0.311 | 1 | 08_mo-DC3 |
| HPS5 | -1.61915116 | 0.185 | 0.403 | 1 | 08_mo-DC3 |
| CLEC5A | -1.55457621 | 0.136 | 0.356 | 1 | 08_mo-DC3 |
| RAB11FIP1 | -1.7807445 | 0.173 | 0.387 | 1 | 08_mo-DC3 |
| RBM8A | -1.1269889 | 0.296 | 0.518 | 1 | 08_mo-DC3 |
| AP1S2 | -1.36450868 | 0.259 | 0.477 | 1 | 08_mo-DC3 |
| C17orf49 | -1.43164702 | 0.136 | 0.373 | 1 | 08_mo-DC3 |
| LAMTOR1 | -1.1868987 | 0.21 | 0.443 | 1 | 08_mo-DC3 |
| RFTN1 | -1.30518693 | 0.309 | 0.539 | 1 | 08_mo-DC3 |
| PMAIP1 | -1.70665552 | 0.148 | 0.359 | 1 | 08_mo-DC3 |
| C19orf53 | -1.30445384 | 0.185 | 0.41 | 1 | 08_mo-DC3 |
| DAPK1 | -1.59073754 | 0.136 | 0.351 | 1 | 08_mo-DC3 |
| TMEM123 | -1.2970415 | 0.247 | 0.453 | 1 | 08_mo-DC3 |
| PEA15 | -1.42956687 | 0.198 | 0.405 | 1 | 08_mo-DC3 |

| | | | | | |
|---|---|---|---|---|---|
| SIPA1L3 | -1.60169723 | 0.198 | 0.412 | 1 | 08_mo-DC3 |
| UFC1 | -1.19573689 | 0.21 | 0.415 | 1 | 08_mo-DC3 |
| BORCS5 | -1.65757573 | 0.185 | 0.386 | 1 | 08_mo-DC3 |
| RSL24D1 | -1.3821457 | 0.148 | 0.353 | 1 | 08_mo-DC3 |
| TOMM6 | -1.07926854 | 0.272 | 0.485 | 1 | 08_mo-DC3 |
| ZNF385A | -1.20927777 | 0.148 | 0.353 | 1 | 08_mo-DC3 |
| IGFLR1 | -1.11959145 | 0.21 | 0.413 | 1 | 08_mo-DC3 |
| CEBPD | -3.03958219 | 0.192 | 0.91 | 2.27E-24 | 09_lowread |
| JUND | -2.79549251 | 0.151 | 0.829 | 3.80E-20 | 09_lowread |
| PNRC1 | -2.00091268 | 0.205 | 0.927 | 9.77E-19 | 09_lowread |
| DBI | -1.81130664 | 0.082 | 0.779 | 1.79E-16 | 09_lowread |
| MTRNR2L12 | 2.293425455 | 0.699 | 0.245 | 2.21E-16 | 09_lowread |
| RPS16 | -1.73446123 | 0.397 | 0.926 | 8.24E-16 | 09_lowread |
| GNG5 | -1.78423101 | 0.11 | 0.817 | 8.37E-16 | 09_lowread |
| RBM3 | -2.51313478 | 0.068 | 0.723 | 1.38E-15 | 09_lowread |
| BTG1 | -1.90623835 | 0.288 | 0.917 | 2.16E-15 | 09_lowread |
| RAC1 | -1.68623773 | 0.233 | 0.895 | 3.81E-15 | 09_lowread |
| CEBPB | -2.62866273 | 0.055 | 0.684 | 1.25E-14 | 09_lowread |
| GNAS | -1.67781003 | 0.192 | 0.862 | 1.35E-14 | 09_lowread |
| RPS2 | -1.70399018 | 0.507 | 0.913 | 1.56E-14 | 09_lowread |
| RPL12 | -1.5567485 | 0.466 | 0.921 | 1.68E-14 | 09_lowread |
| RPL13 | -1.39174531 | 0.726 | 0.951 | 3.59E-14 | 09_lowread |
| RPL36AL | -1.7371261 | 0.123 | 0.782 | 4.68E-14 | 09_lowread |
| DNASE1L3 | 4.689004493 | 0.26 | 0.04 | 6.89E-14 | 09_lowread |
| CXCL16 | -1.20685358 | 0.205 | 0.88 | 7.56E-14 | 09_lowread |
| NME2 | -1.67343358 | 0.164 | 0.823 | 8.54E-14 | 09_lowread |
| ALOX5AP | -2.35892053 | 0.137 | 0.746 | 1.47E-13 | 09_lowread |
| HLA-DMB | -1.4376082 | 0.096 | 0.731 | 1.61E-13 | 09_lowread |
| METRNL | -3.08406167 | 0.068 | 0.661 | 2.05E-13 | 09_lowread |
| RPL10A | -1.6317181 | 0.233 | 0.856 | 3.23E-13 | 09_lowread |
| ID2 | -1.66598609 | 0.192 | 0.829 | 4.28E-13 | 09_lowread |
| CYBA | -1.46534551 | 0.37 | 0.931 | 5.18E-13 | 09_lowread |
| RPS9 | -1.46316386 | 0.452 | 0.927 | 6.39E-13 | 09_lowread |
| ARF6 | -3.63082261 | 0.055 | 0.625 | 8.00E-13 | 09_lowread |
| SRSF5 | -1.4773706 | 0.123 | 0.752 | 1.10E-12 | 09_lowread |
| RPL14 | -1.42673453 | 0.397 | 0.915 | 1.61E-12 | 09_lowread |
| RPL27A | -1.55915761 | 0.123 | 0.772 | 1.90E-12 | 09_lowread |
| CDK2AP1 | -4.25712429 | 0.014 | 0.566 | 2.61E-12 | 09_lowread |
| TYROBP | -1.32531461 | 0.521 | 0.924 | 2.98E-12 | 09_lowread |
| COX6A1 | -1.58348505 | 0.137 | 0.743 | 4.01E-12 | 09_lowread |
| GNAI2 | -1.60366058 | 0.123 | 0.742 | 4.30E-12 | 09_lowread |
| CD81 | -2.24477609 | 0.123 | 0.704 | 8.05E-12 | 09_lowread |
| BTF3 | -1.20438758 | 0.178 | 0.826 | 1.21E-11 | 09_lowread |
| COX8A | -1.27149298 | 0.068 | 0.652 | 2.02E-11 | 09_lowread |
| S100A11 | -1.27618587 | 0.329 | 0.915 | 2.13E-11 | 09_lowread |
| HIGD2A | -2.01822997 | 0.068 | 0.64 | 3.79E-11 | 09_lowread |

| | | | | | |
|---|---|---|---|---|---|
| MZT2B | -13.4283123 | 0 | 0.513 | 4.12E-11 | 09_lowread |
| RPS19 | -1.15685959 | 0.699 | 0.94 | 4.54E-11 | 09_lowread |
| IL1B | -2.46213627 | 0.178 | 0.736 | 4.73E-11 | 09_lowread |
| GPX4 | -1.41761164 | 0.164 | 0.781 | 4.92E-11 | 09_lowread |
| RPL32 | -1.24104594 | 0.616 | 0.918 | 5.41E-11 | 09_lowread |
| RPS20 | -1.44656007 | 0.096 | 0.686 | 6.86E-11 | 09_lowread |
| H2AFZ | -1.2571581 | 0.082 | 0.661 | 7.74E-11 | 09_lowread |
| PCBP1 | -1.07273423 | 0.11 | 0.695 | 1.19E-10 | 09_lowread |
| RHOG | -1.22754631 | 0.096 | 0.663 | 1.55E-10 | 09_lowread |
| EDF1 | -1.29383788 | 0.082 | 0.656 | 1.97E-10 | 09_lowread |
| SLC25A5 | -1.07146354 | 0.11 | 0.71 | 2.14E-10 | 09_lowread |
| RPL27 | -1.21789636 | 0.205 | 0.838 | 2.67E-10 | 09_lowread |
| EIF5A | -3.57989722 | 0.027 | 0.541 | 2.84E-10 | 09_lowread |
| MGAT1 | -1.16652586 | 0.151 | 0.762 | 2.89E-10 | 09_lowread |
| ATP6V1G1 | -2.2891696 | 0.041 | 0.57 | 3.39E-10 | 09_lowread |
| PSMB1 | -2.0972096 | 0.041 | 0.561 | 3.95E-10 | 09_lowread |
| CHCHD2 | -1.24834355 | 0.164 | 0.761 | 4.01E-10 | 09_lowread |
| FAU | -1.17050049 | 0.479 | 0.91 | 4.71E-10 | 09_lowread |
| YBX1 | -1.1842593 | 0.205 | 0.838 | 5.70E-10 | 09_lowread |
| NINJ1 | -2.41388318 | 0.055 | 0.579 | 5.76E-10 | 09_lowread |
| CHCHD10 | -4.53262453 | 0.014 | 0.509 | 6.01E-10 | 09_lowread |
| SEC61B | -1.32602866 | 0.137 | 0.721 | 6.04E-10 | 09_lowread |
| CTSZ | -1.30857112 | 0.137 | 0.715 | 8.11E-10 | 09_lowread |
| RPS4X | -1.23617419 | 0.466 | 0.915 | 9.37E-10 | 09_lowread |
| RPL3 | -1.18633709 | 0.438 | 0.925 | 1.02E-09 | 09_lowread |
| RPS14 | -1.24711027 | 0.603 | 0.912 | 1.05E-09 | 09_lowread |
| EIF5 | -1.21243648 | 0.137 | 0.727 | 1.08E-09 | 09_lowread |
| PSMA7 | -1.7656334 | 0.096 | 0.643 | 1.11E-09 | 09_lowread |
| RPL31 | -1.09693632 | 0.205 | 0.8 | 1.46E-09 | 09_lowread |
| RPL26 | -1.24484642 | 0.589 | 0.922 | 1.54E-09 | 09_lowread |
| ALDOA | -1.86417227 | 0.096 | 0.638 | 1.59E-09 | 09_lowread |
| RPL37 | -1.13323307 | 0.397 | 0.913 | 1.63E-09 | 09_lowread |
| HCST | -1.11844679 | 0.082 | 0.625 | 2.23E-09 | 09_lowread |
| RPL29 | -1.12390624 | 0.479 | 0.916 | 2.97E-09 | 09_lowread |
| CSNK2B | -1.34713362 | 0.082 | 0.627 | 3.13E-09 | 09_lowread |
| RPS23 | -1.10747868 | 0.644 | 0.933 | 3.45E-09 | 09_lowread |
| RPS25 | -1.09069728 | 0.384 | 0.899 | 4.36E-09 | 09_lowread |
| EMD | -2.60331859 | 0.041 | 0.539 | 5.05E-09 | 09_lowread |
| RPL28 | -1.10204801 | 0.712 | 0.94 | 5.22E-09 | 09_lowread |
| VEGFA | -2.2320623 | 0.068 | 0.575 | 5.30E-09 | 09_lowread |
| TRIR | -1.19684835 | 0.096 | 0.63 | 5.63E-09 | 09_lowread |
| HINT1 | -1.17699162 | 0.137 | 0.704 | 6.76E-09 | 09_lowread |
| SPI1 | -1.25116957 | 0.219 | 0.801 | 7.30E-09 | 09_lowread |
| CCL3 | -2.09952017 | 0.096 | 0.617 | 8.41E-09 | 09_lowread |
| FCER1G | -1.07760957 | 0.384 | 0.887 | 9.55E-09 | 09_lowread |
| CXCL8 | -6.47106556 | 0.014 | 0.466 | 1.42E-08 | 09_lowread |

| | | | | | |
|---|---|---|---|---|---|
| RPL23A | -1.23664472 | 0.37 | 0.854 | 1.54E-08 | 09_lowread |
| RPL7A | -1.12171118 | 0.548 | 0.91 | 1.65E-08 | 09_lowread |
| KLF4 | -1.49556621 | 0.11 | 0.632 | 1.72E-08 | 09_lowread |
| TUBB4B | -2.72446143 | 0.041 | 0.517 | 1.86E-08 | 09_lowread |
| VDAC2 | -1.46908959 | 0.041 | 0.518 | 2.19E-08 | 09_lowread |
| UQCR10 | -1.60291475 | 0.041 | 0.523 | 2.41E-08 | 09_lowread |
| ELOB | -1.03814569 | 0.055 | 0.551 | 2.59E-08 | 09_lowread |
| RPS17 | -1.23155428 | 0.096 | 0.623 | 2.89E-08 | 09_lowread |
| NPC2 | -1.04246203 | 0.301 | 0.879 | 2.93E-08 | 09_lowread |
| UBXN1 | -1.84985832 | 0.068 | 0.554 | 3.32E-08 | 09_lowread |
| CIB1 | -1.72155821 | 0.082 | 0.586 | 3.35E-08 | 09_lowread |
| RPS18 | -1.0359025 | 0.548 | 0.93 | 3.45E-08 | 09_lowread |
| PABPC4 | -1.86818411 | 0.082 | 0.582 | 3.79E-08 | 09_lowread |
| ASAH1 | -1.69774713 | 0.096 | 0.602 | 4.42E-08 | 09_lowread |
| RPL21 | -1.06043165 | 0.301 | 0.865 | 4.49E-08 | 09_lowread |
| EEF1B2 | -1.08010468 | 0.288 | 0.82 | 4.65E-08 | 09_lowread |
| ZFAS1 | -1.02273328 | 0.151 | 0.702 | 6.17E-08 | 09_lowread |
| SLC31A2 | -1.01836382 | 0.055 | 0.539 | 6.39E-08 | 09_lowread |
| ARRB2 | -1.17109896 | 0.123 | 0.658 | 6.89E-08 | 09_lowread |
| NDUFA6 | -3.15093625 | 0.014 | 0.457 | 7.32E-08 | 09_lowread |
| TAF10 | -3.20260578 | 0.041 | 0.497 | 7.49E-08 | 09_lowread |
| UXT | -3.09161837 | 0.027 | 0.48 | 8.71E-08 | 09_lowread |
| SH3BGRL3 | -1.15556671 | 0.329 | 0.854 | 8.84E-08 | 09_lowread |
| STX11 | -1.07638679 | 0.123 | 0.636 | 9.82E-08 | 09_lowread |
| EMP3 | -1.22729084 | 0.178 | 0.712 | 1.19E-07 | 09_lowread |
| ANXA1 | -1.42557393 | 0.11 | 0.611 | 1.31E-07 | 09_lowread |
| PRELID1 | -2.38594257 | 0.041 | 0.498 | 1.45E-07 | 09_lowread |
| PPIA | -1.05248325 | 0.411 | 0.917 | 1.51E-07 | 09_lowread |
| ARF4 | -1.33820001 | 0.055 | 0.527 | 1.51E-07 | 09_lowread |
| LST1 | -1.49110695 | 0.096 | 0.598 | 1.62E-07 | 09_lowread |
| MARCKS | -1.8182793 | 0.082 | 0.561 | 1.64E-07 | 09_lowread |
| PRR13 | -1.3913472 | 0.082 | 0.567 | 1.93E-07 | 09_lowread |
| XBP1 | -1.3107764 | 0.055 | 0.525 | 1.96E-07 | 09_lowread |
| CSTB | -1.1308108 | 0.11 | 0.617 | 1.99E-07 | 09_lowread |
| CORO1A | -1.30170381 | 0.096 | 0.59 | 2.05E-07 | 09_lowread |
| ANXA2 | -1.0728889 | 0.137 | 0.66 | 2.08E-07 | 09_lowread |
| COX7A2 | -1.3660946 | 0.055 | 0.524 | 2.28E-07 | 09_lowread |
| BIN1 | -1.26739576 | 0.082 | 0.566 | 2.34E-07 | 09_lowread |
| RPL18 | -1.03530663 | 0.548 | 0.913 | 2.52E-07 | 09_lowread |
| RPS12 | -1.0577341 | 0.726 | 0.931 | 2.56E-07 | 09_lowread |
| PLK3 | -2.18337567 | 0.041 | 0.493 | 2.69E-07 | 09_lowread |
| RPS13 | -1.0333491 | 0.589 | 0.91 | 3.37E-07 | 09_lowread |
| GPR183 | -1.24708395 | 0.288 | 0.835 | 3.59E-07 | 09_lowread |
| RPS3A | -1.08231484 | 0.534 | 0.905 | 4.41E-07 | 09_lowread |
| GNB2 | -1.31169511 | 0.082 | 0.558 | 4.54E-07 | 09_lowread |
| GAREM1 | -3.43527214 | 0.014 | 0.434 | 5.03E-07 | 09_lowread |

| | | | | | |
|---|---|---|---|---|---|
| BCL3 | -2.38175156 | 0.055 | 0.497 | 5.40E-07 | 09_lowread |
| CCL4 | -2.29963147 | 0.178 | 0.654 | 5.43E-07 | 09_lowread |
| PIM3 | -2.43799742 | 0.041 | 0.478 | 5.51E-07 | 09_lowread |
| TALDO1 | -2.04743914 | 0.027 | 0.457 | 5.60E-07 | 09_lowread |
| DYNLT1 | -1.1398089 | 0.082 | 0.561 | 5.97E-07 | 09_lowread |
| GAS5 | -1.6710644 | 0.055 | 0.511 | 6.46E-07 | 09_lowread |
| CHMP4B | -2.39801825 | 0.027 | 0.452 | 6.52E-07 | 09_lowread |
| SEC61G | -3.69054647 | 0.014 | 0.43 | 6.54E-07 | 09_lowread |
| FP671120.4 | -2.08810565 | 0.151 | 0.607 | 7.08E-07 | 09_lowread |
| GPSM3 | -1.53591245 | 0.082 | 0.554 | 7.18E-07 | 09_lowread |
| WDR83OS | -1.13088874 | 0.082 | 0.561 | 7.58E-07 | 09_lowread |
| NECTIN2 | -3.02868351 | 0.041 | 0.47 | 8.82E-07 | 09_lowread |
| MTHFD2 | -1.26750724 | 0.041 | 0.48 | 9.11E-07 | 09_lowread |
| H2AFV | -1.19602855 | 0.055 | 0.506 | 1.03E-06 | 09_lowread |
| PRDX5 | -3.74887394 | 0.014 | 0.423 | 1.21E-06 | 09_lowread |
| TIMP1 | -2.13010645 | 0.233 | 0.701 | 1.72E-06 | 09_lowread |
| MYL12A | -1.0133197 | 0.219 | 0.753 | 1.87E-06 | 09_lowread |
| IFITM2 | -1.34220651 | 0.274 | 0.768 | 1.89E-06 | 09_lowread |
| SPCS1 | -1.22625623 | 0.041 | 0.471 | 1.94E-06 | 09_lowread |
| LINC01578 | -1.08569207 | 0.123 | 0.6 | 2.11E-06 | 09_lowread |
| PEA15 | -1.49139728 | 0.014 | 0.414 | 2.53E-06 | 09_lowread |
| NACA | -1.03178276 | 0.37 | 0.883 | 3.24E-06 | 09_lowread |
| PTP4A2 | -1.4232224 | 0.068 | 0.503 | 4.03E-06 | 09_lowread |
| FCER1A | -2.84079803 | 0.082 | 0.491 | 4.05E-06 | 09_lowread |
| SLC16A3 | -1.02220757 | 0.123 | 0.598 | 4.12E-06 | 09_lowread |
| MAN2B1 | -1.13684451 | 0.151 | 0.641 | 4.50E-06 | 09_lowread |
| AP2S1 | -1.07918247 | 0.082 | 0.532 | 5.62E-06 | 09_lowread |
| REX1BD | -1.55181055 | 0.027 | 0.43 | 5.66E-06 | 09_lowread |
| IER3 | -1.11935315 | 0.233 | 0.72 | 6.43E-06 | 09_lowread |
| APOC1 | -2.65815726 | 0.096 | 0.52 | 6.64E-06 | 09_lowread |
| DYNLL1 | -1.12430867 | 0.055 | 0.482 | 6.87E-06 | 09_lowread |
| C15orf48 | -2.65292928 | 0.11 | 0.525 | 6.88E-06 | 09_lowread |
| ARL4C | -2.13012158 | 0.082 | 0.498 | 7.12E-06 | 09_lowread |
| EIF1B | -3.0834728 | 0.041 | 0.441 | 7.56E-06 | 09_lowread |
| RHOB | -1.86431493 | 0.178 | 0.629 | 7.87E-06 | 09_lowread |
| ZNF706 | -1.7450693 | 0.068 | 0.492 | 8.07E-06 | 09_lowread |
| CXCR4 | -1.05432181 | 0.37 | 0.878 | 9.02E-06 | 09_lowread |
| FKBP8 | -1.67510355 | 0.041 | 0.447 | 1.06E-05 | 09_lowread |
| VIM | -1.05240444 | 0.603 | 0.944 | 1.36E-05 | 09_lowread |
| HNRNPL | -1.34121767 | 0.068 | 0.492 | 1.39E-05 | 09_lowread |
| SSR1 | -1.70849396 | 0.041 | 0.444 | 1.46E-05 | 09_lowread |
| TNFRSF1A | -1.34666004 | 0.041 | 0.444 | 1.50E-05 | 09_lowread |
| ARID5A | -1.09903021 | 0.068 | 0.49 | 1.61E-05 | 09_lowread |
| SNHG15 | -1.93311213 | 0.082 | 0.494 | 1.73E-05 | 09_lowread |
| CAPNS1 | -1.99097981 | 0.027 | 0.414 | 1.84E-05 | 09_lowread |
| CCL4L2 | -2.82466961 | 0.096 | 0.507 | 1.86E-05 | 09_lowread |

| | | | | | |
|---|---|---|---|---|---|
| IER5 | -2.15557743 | 0.055 | 0.455 | 1.93E-05 | 09_lowread |
| DRAP1 | -1.37730544 | 0.123 | 0.577 | 2.35E-05 | 09_lowread |
| CD164 | -1.19080454 | 0.068 | 0.487 | 2.42E-05 | 09_lowread |
| CRTAP | -1.4306649 | 0.041 | 0.437 | 2.51E-05 | 09_lowread |
| CD44 | -1.01057297 | 0.233 | 0.714 | 2.74E-05 | 09_lowread |
| LGALS3 | -1.05028323 | 0.068 | 0.488 | 2.80E-05 | 09_lowread |
| EFHD2 | -1.67938228 | 0.11 | 0.534 | 3.10E-05 | 09_lowread |
| OGDH | -1.9320383 | 0.041 | 0.43 | 3.48E-05 | 09_lowread |
| HMGN3 | -1.29383383 | 0.068 | 0.484 | 3.66E-05 | 09_lowread |
| AL133415.1 | -2.79127213 | 0.027 | 0.397 | 4.44E-05 | 09_lowread |
| INSIG1 | -1.0903355 | 0.151 | 0.604 | 4.64E-05 | 09_lowread |
| NBDY | -1.65724026 | 0.041 | 0.427 | 4.78E-05 | 09_lowread |
| ANXA7 | -2.35456377 | 0.014 | 0.374 | 4.94E-05 | 09_lowread |
| AC138123.1 | -4.10525146 | 0.014 | 0.372 | 4.95E-05 | 09_lowread |
| MKNK2 | -3.57209609 | 0.014 | 0.373 | 5.10E-05 | 09_lowread |
| OSTC | -1.95184116 | 0.014 | 0.373 | 5.20E-05 | 09_lowread |
| PDGFB | -2.96546767 | 0.027 | 0.396 | 5.39E-05 | 09_lowread |
| PTMS | -2.07187841 | 0.041 | 0.417 | 5.53E-05 | 09_lowread |
| SRSF3 | -1.08097658 | 0.151 | 0.608 | 5.64E-05 | 09_lowread |
| OLR1 | -1.19413877 | 0.247 | 0.685 | 5.91E-05 | 09_lowread |
| NAPA | -1.1745553 | 0.082 | 0.49 | 6.77E-05 | 09_lowread |
| VMO1 | -1.53998809 | 0.055 | 0.446 | 7.16E-05 | 09_lowread |
| G0S2 | -3.98562903 | 0.068 | 0.427 | 7.46E-05 | 09_lowread |
| POLR2E | -1.06950211 | 0.055 | 0.446 | 7.53E-05 | 09_lowread |
| NFKBIA | -1.09413832 | 0.466 | 0.917 | 7.55E-05 | 09_lowread |
| IL18 | -1.18688417 | 0.11 | 0.542 | 7.68E-05 | 09_lowread |
| TMED2 | -1.91894171 | 0.014 | 0.368 | 7.76E-05 | 09_lowread |
| PHB2 | -1.42101774 | 0.041 | 0.415 | 8.86E-05 | 09_lowread |
| PSMB10 | -2.05418581 | 0.041 | 0.416 | 9.26E-05 | 09_lowread |
| AHR | -1.51117426 | 0.096 | 0.497 | 9.31E-05 | 09_lowread |
| SF3B5 | -2.60215923 | 0.027 | 0.387 | 9.76E-05 | 09_lowread |
| NANS | -4.57467937 | 0.014 | 0.353 | 0.0001706 | 09_lowread |
| ABHD12 | -1.11048856 | 0.096 | 0.506 | 0.0001903 | 09_lowread |
| GLIPR2 | -1.54531992 | 0.068 | 0.453 | 0.0001932 | 09_lowread |
| SUPT4H1 | -1.23036921 | 0.041 | 0.407 | 0.0002147 | 09_lowread |
| NDUFA2 | -1.50235955 | 0.027 | 0.38 | 0.0002291 | 09_lowread |
| LY6E | -2.29122715 | 0.041 | 0.401 | 0.000235 | 09_lowread |
| CRIP1 | -2.20261633 | 0.082 | 0.455 | 0.000238 | 09_lowread |
| LPXN | -1.42174357 | 0.068 | 0.448 | 0.0002458 | 09_lowread |
| OXA1L | -1.63591696 | 0.027 | 0.378 | 0.0002561 | 09_lowread |
| UPP1 | -1.05184715 | 0.068 | 0.452 | 0.0002655 | 09_lowread |
| BNIP2 | -1.46910707 | 0.041 | 0.4 | 0.0003138 | 09_lowread |
| SF3B6 | -12.0605018 | 0 | 0.317 | 0.0003998 | 09_lowread |
| ANAPC11 | -1.26570271 | 0.068 | 0.443 | 0.0004367 | 09_lowread |
| MYDGF | -1.61311696 | 0.041 | 0.391 | 0.0004604 | 09_lowread |
| UBE2V1 | -1.23159951 | 0.068 | 0.441 | 0.0004637 | 09_lowread |

| | | | | | |
|---|---|---|---|---|---|
| NRARP | -3.04507975 | 0.014 | 0.341 | 0.0004778 | 09_lowread |
| AZIN1 | -2.04798241 | 0.055 | 0.413 | 0.0004973 | 09_lowread |
| PPP1R18 | -1.05021966 | 0.055 | 0.418 | 0.0005758 | 09_lowread |
| SAP30 | -2.89823729 | 0.041 | 0.38 | 0.0005941 | 09_lowread |
| CD9 | -1.51045356 | 0.055 | 0.403 | 0.0008222 | 09_lowread |
| CST7 | -1.53289103 | 0.123 | 0.507 | 0.0008667 | 09_lowread |
| TAGAP | -1.01634817 | 0.082 | 0.457 | 0.0009464 | 09_lowread |
| TNIP1 | -12.4690478 | 0 | 0.303 | 0.0010243 | 09_lowread |
| JDP2 | -1.21014659 | 0.096 | 0.472 | 0.0010733 | 09_lowread |
| UQCRC1 | -1.82209488 | 0.027 | 0.355 | 0.001248 | 09_lowread |
| SIVA1 | -2.36212163 | 0.027 | 0.353 | 0.0012608 | 09_lowread |
| TAF1D | -2.48161842 | 0.027 | 0.353 | 0.0013228 | 09_lowread |
| JOSD1 | -1.97791331 | 0.041 | 0.378 | 0.0013287 | 09_lowread |
| CHURC1 | -2.65550631 | 0.014 | 0.326 | 0.0013715 | 09_lowread |
| MAFF | -1.57904423 | 0.055 | 0.4 | 0.0014535 | 09_lowread |
| CCDC107 | -1.68996049 | 0.041 | 0.373 | 0.0017421 | 09_lowread |
| GLRX | -3.28833262 | 0.014 | 0.321 | 0.0018229 | 09_lowread |
| STX4 | -2.16410519 | 0.041 | 0.371 | 0.0019849 | 09_lowread |
| ERGIC3 | -1.15877898 | 0.041 | 0.374 | 0.0020409 | 09_lowread |
| DDOST | -1.66918232 | 0.055 | 0.393 | 0.0021658 | 09_lowread |
| UBE2A | -1.27862915 | 0.055 | 0.397 | 0.0022272 | 09_lowread |
| VKORC1 | -12.0259814 | 0 | 0.291 | 0.0022383 | 09_lowread |
| GSTO1 | -1.2548366 | 0.068 | 0.421 | 0.0024648 | 09_lowread |
| LAMTOR5 | -1.84870788 | 0.041 | 0.366 | 0.0024933 | 09_lowread |
| SMIM26 | -2.26696379 | 0.027 | 0.339 | 0.0026352 | 09_lowread |
| TREM1 | -1.46095965 | 0.027 | 0.341 | 0.0030544 | 09_lowread |
| PMAIP1 | -1.49479077 | 0.041 | 0.364 | 0.0032561 | 09_lowread |
| ABL2 | -1.46675123 | 0.068 | 0.406 | 0.0033349 | 09_lowread |
| LSM7 | -1.27277494 | 0.041 | 0.362 | 0.0033869 | 09_lowread |
| CTSD | -1.51055442 | 0.11 | 0.464 | 0.0035655 | 09_lowread |
| CLEC5A | -1.83992871 | 0.041 | 0.359 | 0.0037253 | 09_lowread |
| CCR7 | -4.12474926 | 0.027 | 0.323 | 0.0039465 | 09_lowread |
| NUDT3 | -1.43011602 | 0.055 | 0.384 | 0.0040738 | 09_lowread |
| CD1C | -2.64005252 | 0.041 | 0.353 | 0.0041314 | 09_lowread |
| PCGF5 | -1.29912796 | 0.055 | 0.382 | 0.0050387 | 09_lowread |
| CCL22 | -14.7599661 | 0 | 0.278 | 0.0055008 | 09_lowread |
| DNAJC7 | -2.56153232 | 0.027 | 0.325 | 0.006364 | 09_lowread |
| MAPKAPK2 | -2.46799763 | 0.027 | 0.325 | 0.00713 | 09_lowread |
| PRKAG2-AS1 | -12.1135593 | 0 | 0.273 | 0.0071737 | 09_lowread |
| SATB1 | -1.10934181 | 0.068 | 0.402 | 0.0072212 | 09_lowread |
| PPP1R15B | -2.55443731 | 0.014 | 0.3 | 0.0072715 | 09_lowread |
| CHMP4A | -11.828494 | 0 | 0.272 | 0.0078342 | 09_lowread |
| LTC4S | -1.16924544 | 0.068 | 0.4 | 0.0078654 | 09_lowread |
| PPP1R14B | -12.2234984 | 0 | 0.271 | 0.0081863 | 09_lowread |
| BHLHE41 | -12.4471636 | 0 | 0.269 | 0.0093374 | 09_lowread |
| AMPD2 | -1.24057845 | 0.055 | 0.375 | 0.0094317 | 09_lowread |

| | | | | | |
|---|---|---|---|---|---|
| AL137857.1 | -12.5258829 | 0 | 0.269 | 0.009755 | 09_lowread |
| DUSP5 | -1.37399779 | 0.11 | 0.453 | 0.0109314 | 09_lowread |
| SBDS | -3.80094 | 0.014 | 0.292 | 0.0110142 | 09_lowread |
| STUB1 | -11.8103156 | 0 | 0.266 | 0.0111196 | 09_lowread |
| RAB11B | -1.19825294 | 0.027 | 0.321 | 0.0112484 | 09_lowread |
| EIF1AX | -1.13607515 | 0.055 | 0.371 | 0.0113793 | 09_lowread |
| HADHB | -1.42176023 | 0.027 | 0.319 | 0.0119457 | 09_lowread |
| DCXR | -3.5395948 | 0.014 | 0.291 | 0.012278 | 09_lowread |
| FDX1 | -11.9868328 | 0 | 0.264 | 0.0126691 | 09_lowread |
| FBL | -2.12947059 | 0.027 | 0.319 | 0.0127007 | 09_lowread |
| PNPLA2 | -1.78537734 | 0.027 | 0.318 | 0.0132164 | 09_lowread |
| METTL26 | -1.58106577 | 0.014 | 0.291 | 0.0134871 | 09_lowread |
| POLR1D | -1.56323085 | 0.041 | 0.343 | 0.0138249 | 09_lowread |
| SNHG16 | -1.94627428 | 0.027 | 0.315 | 0.0147253 | 09_lowread |
| PTRHD1 | -1.64578927 | 0.027 | 0.314 | 0.0163032 | 09_lowread |
| GPS2 | -3.45068996 | 0.014 | 0.287 | 0.0166513 | 09_lowread |
| ATP5MC1 | -2.18773581 | 0.014 | 0.287 | 0.0167415 | 09_lowread |
| KLF2 | -2.67824172 | 0.041 | 0.331 | 0.0168593 | 09_lowread |
| CD151 | -11.7842321 | 0 | 0.26 | 0.0171456 | 09_lowread |
| MAF1 | -1.61102172 | 0.027 | 0.314 | 0.0173572 | 09_lowread |
| NUTF2 | -1.46973019 | 0.014 | 0.287 | 0.0174783 | 09_lowread |
| MLF2 | -1.03145326 | 0.068 | 0.388 | 0.0187623 | 09_lowread |
| NDUFAF3 | -1.38840027 | 0.027 | 0.312 | 0.0189773 | 09_lowread |
| ATP1B1 | -2.3098608 | 0.027 | 0.31 | 0.0191402 | 09_lowread |
| SCP2 | -1.08507224 | 0.041 | 0.338 | 0.0201939 | 09_lowread |
| MIR155HG | -1.78210327 | 0.014 | 0.284 | 0.0205691 | 09_lowread |
| SNRPF | -1.6239026 | 0.014 | 0.283 | 0.0216617 | 09_lowread |
| ILK | -1.73170895 | 0.014 | 0.282 | 0.0226081 | 09_lowread |
| TREM2 | -1.34781336 | 0.082 | 0.4 | 0.0239176 | 09_lowread |
| MGST2 | -2.11202634 | 0.014 | 0.281 | 0.0246228 | 09_lowread |
| KRAS | -2.93839467 | 0.027 | 0.303 | 0.0249274 | 09_lowread |
| COX16 | -11.6733706 | 0 | 0.253 | 0.0252055 | 09_lowread |
| CMPK1 | -4.08123057 | 0.014 | 0.278 | 0.0254346 | 09_lowread |
| TNFAIP2 | -3.78330092 | 0.027 | 0.296 | 0.0276281 | 09_lowread |
| NFIL3 | -1.32138602 | 0.082 | 0.402 | 0.0279226 | 09_lowread |
| LILRB2 | -4.64342005 | 0.014 | 0.273 | 0.0287332 | 09_lowread |
| RHOF | -3.10858477 | 0.027 | 0.299 | 0.0296067 | 09_lowread |
| FKBP2 | -3.44417136 | 0.014 | 0.277 | 0.0299603 | 09_lowread |
| CREB5 | -1.21475964 | 0.055 | 0.35 | 0.0305481 | 09_lowread |
| PNP | -1.92499527 | 0.014 | 0.277 | 0.0317139 | 09_lowread |
| FUNDC2 | -3.51629219 | 0.014 | 0.274 | 0.0351026 | 09_lowread |
| NENF | -1.73810443 | 0.027 | 0.302 | 0.035313 | 09_lowread |
| ARRDC1 | -11.6908935 | 0 | 0.246 | 0.0384885 | 09_lowread |
| ATP2B1-AS1 | -1.74036283 | 0.041 | 0.323 | 0.0404785 | 09_lowread |
| CTDSP1 | -2.90997351 | 0.014 | 0.272 | 0.0414087 | 09_lowread |
| MRPL14 | -11.7598467 | 0 | 0.245 | 0.0418636 | 09_lowread |

| | | | | | |
|---|---|---|---|---|---|
| BCL7B | -11.7600407 | 0 | 0.243 | 0.0474714 | 09_lowread |
| CD72 | -1.59678796 | 0.041 | 0.322 | 0.047714 | 09_lowread |
| TRAF1 | -1.13260909 | 0.055 | 0.344 | 0.048661 | 09_lowread |
| LINC01619 | -1.00558743 | 0.068 | 0.369 | 0.0491479 | 09_lowread |
| AC004448.2 | -12.2447565 | 0 | 0.241 | 0.0516086 | 09_lowread |
| SPG21 | -1.02453291 | 0.041 | 0.321 | 0.051658 | 09_lowread |
| MRPL57 | -1.08979498 | 0.014 | 0.268 | 0.0548403 | 09_lowread |
| DDT | -1.79368708 | 0.027 | 0.294 | 0.0560948 | 09_lowread |
| NHP2 | -1.82586632 | 0.027 | 0.293 | 0.0590071 | 09_lowread |
| JMJD6 | -2.36945422 | 0.027 | 0.289 | 0.0590637 | 09_lowread |
| TNIP2 | -11.9701691 | 0 | 0.238 | 0.0635435 | 09_lowread |
| DDIT3 | -11.9757349 | 0 | 0.238 | 0.0635435 | 09_lowread |
| MYO1G | -1.49763372 | 0.041 | 0.313 | 0.063997 | 09_lowread |
| LAMTOR2 | -1.17220645 | 0.041 | 0.316 | 0.0713083 | 09_lowread |
| CMTM3 | -1.76312214 | 0.027 | 0.289 | 0.0745623 | 09_lowread |
| TMEM70 | -11.8237861 | 0 | 0.231 | 0.0959795 | 09_lowread |
| CNIH1 | -1.53672903 | 0.014 | 0.258 | 0.0977921 | 09_lowread |
| ALYREF | -4.01051568 | 0.014 | 0.255 | 0.0996464 | 09_lowread |
| VEGFB | -2.11678478 | 0.027 | 0.283 | 0.1039513 | 09_lowread |
| TOMM5 | -11.6055259 | 0 | 0.229 | 0.1085179 | 09_lowread |
| OGFR | -1.48456695 | 0.027 | 0.282 | 0.1125205 | 09_lowread |
| NCF4 | -2.04513501 | 0.027 | 0.282 | 0.1144685 | 09_lowread |
| SNX17 | -1.1006386 | 0.027 | 0.282 | 0.1148524 | 09_lowread |
| TMED9 | -1.53037661 | 0.055 | 0.33 | 0.1176398 | 09_lowread |
| MRPL33 | -11.5035843 | 0 | 0.228 | 0.117746 | 09_lowread |
| PTGER4 | -1.02896352 | 0.123 | 0.44 | 0.1190705 | 09_lowread |
| DYNLRB1 | -3.28055498 | 0.014 | 0.254 | 0.1200474 | 09_lowread |
| LIPA | -2.64246676 | 0.014 | 0.253 | 0.1274401 | 09_lowread |
| COX14 | -1.17924263 | 0.014 | 0.253 | 0.1301177 | 09_lowread |
| UBE2M | -11.6300091 | 0 | 0.226 | 0.1330326 | 09_lowread |
| HNRNPH2 | -1.59423243 | 0.027 | 0.279 | 0.1336975 | 09_lowread |
| MIR22HG | -2.53872161 | 0.014 | 0.252 | 0.1401458 | 09_lowread |
| GINM1 | -11.5215342 | 0 | 0.224 | 0.1442768 | 09_lowread |
| SNRPD3 | -2.35500357 | 0.014 | 0.251 | 0.1464456 | 09_lowread |
| RAB5IF | -1.39954351 | 0.041 | 0.304 | 0.1484072 | 09_lowread |
| APOE | -2.51007593 | 0.301 | 0.604 | 0.1487327 | 09_lowread |
| NAA10 | -2.94919282 | 0.014 | 0.251 | 0.1506597 | 09_lowread |
| FABP5 | -1.66327469 | 0.068 | 0.341 | 0.15819 | 09_lowread |
| NME3 | -1.73552298 | 0.041 | 0.3 | 0.1586735 | 09_lowread |
| PRXL2C | -2.84380745 | 0.014 | 0.249 | 0.1630567 | 09_lowread |
| CSTA | -2.86953772 | 0.014 | 0.248 | 0.1691826 | 09_lowread |
| SLC25A39 | -1.0835439 | 0.027 | 0.276 | 0.170452 | 09_lowread |
| NQO2 | -2.3659426 | 0.027 | 0.273 | 0.1711334 | 09_lowread |
| ZC3H15 | -1.08354867 | 0.068 | 0.346 | 0.172622 | 09_lowread |
| ARL4A | -1.42923787 | 0.041 | 0.299 | 0.1801182 | 09_lowread |
| CCL3L1 | -2.48907722 | 0.068 | 0.33 | 0.180472 | 09_lowread |

| | | | | | |
|---|---|---|---|---|---|
| ECH1 | -2.89419079 | 0.014 | 0.247 | 0.1855922 | 09_lowread |
| AC004817.3 | -1.11091843 | 0.055 | 0.321 | 0.188544 | 09_lowread |
| CLEC10A | -1.06604562 | 0.096 | 0.381 | 0.20027 | 09_lowread |
| MRC1 | -1.05436299 | 0.041 | 0.295 | 0.2025211 | 09_lowread |
| FBP1 | -1.59049268 | 0.041 | 0.296 | 0.2113829 | 09_lowread |
| SRP19 | -1.92247943 | 0.027 | 0.271 | 0.2199272 | 09_lowread |
| FAM174C | -1.95701866 | 0.014 | 0.244 | 0.2283403 | 09_lowread |
| MRPS36 | -11.4629201 | 0 | 0.216 | 0.2338376 | 09_lowread |
| RAB34 | -11.3673149 | 0 | 0.215 | 0.2433658 | 09_lowread |
| OSM | -2.06001635 | 0.014 | 0.242 | 0.2452075 | 09_lowread |
| ITPRIP | -1.55924439 | 0.041 | 0.294 | 0.2486649 | 09_lowread |
| EREG | -1.73453722 | 0.055 | 0.311 | 0.2599299 | 09_lowread |
| PXDC1 | -11.816009 | 0 | 0.214 | 0.2635664 | 09_lowread |
| TSPAN33 | -2.71396903 | 0.027 | 0.264 | 0.273055 | 09_lowread |
| ABRACL | -2.21574353 | 0.014 | 0.24 | 0.2786786 | 09_lowread |
| FBXO7 | -1.01601927 | 0.014 | 0.24 | 0.2786786 | 09_lowread |
| CARD19 | -3.44739128 | 0.014 | 0.238 | 0.2921648 | 09_lowread |
| IL4I1 | -1.99186579 | 0.027 | 0.264 | 0.2950467 | 09_lowread |
| IFI6 | -1.66785028 | 0.068 | 0.331 | 0.3107669 | 09_lowread |
| MRPS24 | -3.15442349 | 0.014 | 0.237 | 0.3166863 | 09_lowread |
| RAB32 | -1.63868131 | 0.041 | 0.289 | 0.3183405 | 09_lowread |
| BUD31 | -3.1347983 | 0.014 | 0.237 | 0.329459 | 09_lowread |
| CCT2 | -1.38020679 | 0.041 | 0.289 | 0.3313953 | 09_lowread |
| DARS | -1.31021765 | 0.055 | 0.311 | 0.3472507 | 09_lowread |
| RTL8C | -11.4891188 | 0 | 0.209 | 0.3479135 | 09_lowread |
| PSMA3 | -1.13963662 | 0.041 | 0.285 | 0.3716583 | 09_lowread |
| SNHG5 | -1.66847523 | 0.055 | 0.308 | 0.3878346 | 09_lowread |
| AL136987.1 | -2.22031471 | 0.027 | 0.259 | 0.3959613 | 09_lowread |
| RBIS | -1.17780247 | 0.041 | 0.284 | 0.3981245 | 09_lowread |
| RFX2 | -1.11819932 | 0.041 | 0.284 | 0.4066939 | 09_lowread |
| MCRIP1 | -11.3078195 | 0 | 0.206 | 0.4073256 | 09_lowread |
| PLB1 | -11.4279689 | 0 | 0.205 | 0.4236532 | 09_lowread |
| CLEC4A | -2.25573948 | 0.014 | 0.232 | 0.4301992 | 09_lowread |
| GOLPH3 | -1.14107107 | 0.055 | 0.31 | 0.4491165 | 09_lowread |
| POLR2J | -3.74083209 | 0.014 | 0.229 | 0.4614795 | 09_lowread |
| CYTH3 | -3.0194974 | 0.014 | 0.23 | 0.4637654 | 09_lowread |
| EIF2AK3 | -1.16846734 | 0.068 | 0.329 | 0.4849514 | 09_lowread |
| GRHPR | -11.2777916 | 0 | 0.203 | 0.4955545 | 09_lowread |
| PSMF1 | -1.71126994 | 0.041 | 0.281 | 0.5005446 | 09_lowread |
| AC025164.1 | -1.15261197 | 0.027 | 0.256 | 0.5034988 | 09_lowread |
| PILRA | -1.963783 | 0.027 | 0.255 | 0.5058853 | 09_lowread |
| PUF60 | -1.07388822 | 0.027 | 0.256 | 0.5059526 | 09_lowread |
| VMA21 | -2.98239034 | 0.014 | 0.229 | 0.5134671 | 09_lowread |
| PYM1 | -11.3275402 | 0 | 0.202 | 0.5153038 | 09_lowread |
| PIGT | -11.3155602 | 0 | 0.202 | 0.5153038 | 09_lowread |
| LSM4 | -1.32876562 | 0.041 | 0.279 | 0.525047 | 09_lowread |

| | | | | | |
|---|---|---|---|---|---|
| SAFB2 | -1.59629898 | 0.041 | 0.28 | 0.5324885 | 09_lowread |
| SNHG32 | -3.58026939 | 0.014 | 0.227 | 0.5357446 | 09_lowread |
| EMC4 | -1.66328255 | 0.014 | 0.228 | 0.5448025 | 09_lowread |
| SHKBP1 | -2.66521385 | 0.014 | 0.228 | 0.5645181 | 09_lowread |
| RNF5 | -2.99225194 | 0.014 | 0.227 | 0.573207 | 09_lowread |
| MARCKSL1 | -1.40117917 | 0.027 | 0.248 | 0.5816014 | 09_lowread |
| UBALD2 | -2.7156154 | 0.027 | 0.249 | 0.5923966 | 09_lowread |
| ZBTB7A | -1.03102765 | 0.041 | 0.278 | 0.6057294 | 09_lowread |
| RIT1 | -1.84351092 | 0.014 | 0.226 | 0.6364062 | 09_lowread |
| GTF3C6 | -1.04685577 | 0.014 | 0.224 | 0.6876459 | 09_lowread |
| CHRNE | -1.94252532 | 0.027 | 0.25 | 0.6932622 | 09_lowread |
| DUSP23 | -2.36010358 | 0.027 | 0.248 | 0.6984499 | 09_lowread |
| TRIM8 | -1.55763228 | 0.027 | 0.25 | 0.715932 | 09_lowread |
| MSRA | -1.7764917 | 0.055 | 0.294 | 0.7366242 | 09_lowread |
| CNPPD1 | -1.21745078 | 0.041 | 0.276 | 0.742458 | 09_lowread |
| MAPKAP1 | -1.12784324 | 0.041 | 0.273 | 0.7616466 | 09_lowread |
| ARL8A | -2.31661425 | 0.014 | 0.222 | 0.7707917 | 09_lowread |
| TIMM17A | -1.1363383 | 0.027 | 0.247 | 0.8002881 | 09_lowread |
| BCL2L13 | -3.02286125 | 0.014 | 0.221 | 0.8077551 | 09_lowread |
| EIF4EBP1 | -2.58587573 | 0.014 | 0.221 | 0.8186575 | 09_lowread |
| MAP2K2 | -3.05332972 | 0.014 | 0.22 | 0.8450626 | 09_lowread |
| MRPL55 | -1.39840902 | 0.014 | 0.219 | 0.9356485 | 09_lowread |
| PLAC8 | -2.61382773 | 0.027 | 0.239 | 0.969819 | 09_lowread |
| VTI1B | -1.17735801 | 0.014 | 0.218 | 0.9721757 | 09_lowread |
| RANBP1 | -1.12736956 | 0.041 | 0.27 | 0.9966468 | 09_lowread |
| SLC36A4 | -1.34839983 | 0.027 | 0.244 | 1 | 09_lowread |
| CCDC200 | -2.14399586 | 0.027 | 0.241 | 1 | 09_lowread |
| MEIKIN | -2.37385578 | 0.014 | 0.217 | 1 | 09_lowread |
| RNPEP | -1.22953409 | 0.014 | 0.216 | 1 | 09_lowread |
| POLR2G | -1.43879077 | 0.027 | 0.241 | 1 | 09_lowread |
| C12orf57 | -1.29671052 | 0.041 | 0.264 | 1 | 09_lowread |
| DDX39A | -1.30368357 | 0.055 | 0.289 | 1 | 09_lowread |
| SNHG12 | -1.31466141 | 0.027 | 0.239 | 1 | 09_lowread |
| DGUOK | -1.03557347 | 0.055 | 0.289 | 1 | 09_lowread |
| PPIL4 | -1.3233325 | 0.041 | 0.263 | 1 | 09_lowread |
| ADAM8 | -1.39645893 | 0.068 | 0.308 | 1 | 09_lowread |
| AL118516.1 | -1.97097627 | 0.041 | 0.26 | 1 | 09_lowread |
| RAMP1 | -1.23764005 | 0.041 | 0.259 | 1 | 09_lowread |
| CCNT1 | -1.11056355 | 0.027 | 0.235 | 1 | 09_lowread |
| AMZ2 | -1.36528607 | 0.041 | 0.262 | 1 | 09_lowread |
| ITGAM | -1.32276548 | 0.027 | 0.233 | 1 | 09_lowread |
| SLC12A6 | -1.14353644 | 0.027 | 0.23 | 1 | 09_lowread |
| TBC1D7 | -1.68166836 | 0.027 | 0.231 | 1 | 09_lowread |
| CDKN1B | -1.21309556 | 0.027 | 0.231 | 1 | 09_lowread |
| ARRDC2 | -1.61677359 | 0.027 | 0.23 | 1 | 09_lowread |
| IL1RAP | -1.46443522 | 0.068 | 0.296 | 1 | 09_lowread |

| | | | | | |
|---|---|---|---|---|---|
| CCDC85B | -1.02532744 | 0.055 | 0.277 | 1 | 09_lowread |
| CD52 | -1.07779933 | 0.027 | 0.228 | 1 | 09_lowread |
| DHX15 | -1.0682417 | 0.027 | 0.227 | 1 | 09_lowread |
| CREB1 | -1.44270065 | 0.055 | 0.272 | 1 | 09_lowread |
| RNF19B | -1.22196787 | 0.041 | 0.251 | 1 | 09_lowread |
| SLC16A10 | -1.09952588 | 0.151 | 0.421 | 1 | 09_lowread |
| PPP4R3B | -1.93970672 | 0.041 | 0.245 | 1 | 09_lowread |
| TRIM28 | -1.14154406 | 0.041 | 0.247 | 1 | 09_lowread |
| ZNF655 | -1.20087198 | 0.041 | 0.248 | 1 | 09_lowread |
| NFKBIE | -1.1131373 | 0.055 | 0.263 | 1 | 09_lowread |
| ADGRE5 | 1.099881457 | 0.151 | 0.453 | 1 | 09_lowread |
| THBS1 | -1.50623522 | 0.151 | 0.375 | 1 | 09_lowread |
| TCIRG1 | 1.008386454 | 0.137 | 0.412 | 1 | 09_lowread |
| ADA2 | 1.002744627 | 0.137 | 0.393 | 1 | 09_lowread |
| STX17-AS1 | 1.05823928 | 0.219 | 0.532 | 1 | 09_lowread |
| RNPS1 | 1.188793568 | 0.123 | 0.34 | 1 | 09_lowread |
| GNAI3 | 1.004257266 | 0.151 | 0.385 | 1 | 09_lowread |
| OS9 | 1.052485654 | 0.178 | 0.433 | 1 | 09_lowread |
| IL2RG | 1.142639675 | 0.151 | 0.363 | 1 | 09_lowread |
| RNASE6 | 1.023566548 | 0.274 | 0.582 | 1 | 09_lowread |
| MT2A | -1.36233344 | 0.466 | 0.672 | 1 | 09_lowread |
| PTPRC | 1.06651557 | 0.616 | 0.835 | 1 | 09_lowread |
| PPP1R10 | 1.072801558 | 0.219 | 0.473 | 1 | 09_lowread |
| ENOX1 | 5.794944347 | 0.72 | 0.021 | 5.84E-82 | 10_mDC2 |
| ADAM12 | 6.316336643 | 0.44 | 0.009 | 1.02E-61 | 10_mDC2 |
| S1PR1 | 5.23953586 | 0.32 | 0.005 | 1.09E-51 | 10_mDC2 |
| SLC22A23 | 5.167755744 | 0.64 | 0.03 | 4.46E-50 | 10_mDC2 |
| LIMCH1 | 6.39335576 | 0.52 | 0.019 | 1.17E-49 | 10_mDC2 |
| CD200 | 6.704533267 | 0.28 | 0.003 | 1.26E-49 | 10_mDC2 |
| CCL19 | 8.515013992 | 0.24 | 0.003 | 3.00E-39 | 10_mDC2 |
| PLEKHG1 | 4.655979101 | 0.52 | 0.026 | 7.36E-37 | 10_mDC2 |
| CASC15 | 6.300074022 | 0.32 | 0.009 | 2.00E-35 | 10_mDC2 |
| ARNTL2 | 4.647512258 | 0.6 | 0.038 | 2.10E-35 | 10_mDC2 |
| SLCO5A1 | 5.454345679 | 0.52 | 0.031 | 1.19E-32 | 10_mDC2 |
| ACHE | 5.247474329 | 0.32 | 0.011 | 2.09E-30 | 10_mDC2 |
| LAMP3 | 4.159993045 | 0.96 | 0.17 | 2.42E-27 | 10_mDC2 |
| TBC1D4 | 4.077497814 | 0.72 | 0.079 | 8.31E-27 | 10_mDC2 |
| PDCD1LG2 | 6.001798083 | 0.32 | 0.013 | 9.17E-27 | 10_mDC2 |
| TSPAN13 | 4.276736537 | 0.44 | 0.026 | 1.27E-26 | 10_mDC2 |
| IL32 | 4.899718034 | 0.52 | 0.04 | 1.25E-25 | 10_mDC2 |
| IGKC | 5.74740801 | 0.24 | 0.007 | 2.53E-25 | 10_mDC2 |
| ANKRD33B | 4.580279855 | 0.64 | 0.066 | 1.25E-24 | 10_mDC2 |
| H2AFY2 | 4.977940808 | 0.48 | 0.035 | 1.77E-24 | 10_mDC2 |
| GCSAM | 4.943153214 | 0.28 | 0.011 | 2.41E-23 | 10_mDC2 |
| IDO1 | 3.208450005 | 0.76 | 0.097 | 1.60E-22 | 10_mDC2 |
| MGLL | 4.190343095 | 0.64 | 0.073 | 2.59E-22 | 10_mDC2 |

| | | | | | |
|---|---|---|---|---|---|
| GPC5 | 5.811796518 | 0.24 | 0.009 | 9.40E-21 | 10_mDC2 |
| ARAP2 | 3.857751426 | 0.72 | 0.101 | 1.40E-20 | 10_mDC2 |
| MBOAT2 | 3.598946981 | 0.44 | 0.04 | 1.40E-16 | 10_mDC2 |
| ARHGAP10 | 3.563775414 | 0.72 | 0.123 | 1.87E-16 | 10_mDC2 |
| UBD | 5.84924574 | 0.24 | 0.011 | 3.30E-16 | 10_mDC2 |
| BIRC3 | 3.984618578 | 1 | 0.375 | 7.71E-16 | 10_mDC2 |
| FSCN1 | 3.983803297 | 0.8 | 0.178 | 1.08E-14 | 10_mDC2 |
| CCR7 | 3.166625386 | 0.96 | 0.298 | 1.20E-14 | 10_mDC2 |
| PLCL1 | 4.100153203 | 0.52 | 0.064 | 1.81E-14 | 10_mDC2 |
| IL7R | 3.227966472 | 0.84 | 0.226 | 3.96E-13 | 10_mDC2 |
| PALM2-AKAP2 | 3.611996296 | 0.76 | 0.177 | 8.11E-13 | 10_mDC2 |
| BCL2L14 | 3.14560123 | 0.4 | 0.044 | 4.38E-12 | 10_mDC2 |
| KDM2B | 4.003396118 | 0.76 | 0.197 | 9.45E-12 | 10_mDC2 |
| GRSF1 | 3.40320509 | 0.8 | 0.215 | 1.03E-11 | 10_mDC2 |
| DAPP1 | 2.716139559 | 0.92 | 0.28 | 2.25E-11 | 10_mDC2 |
| C4orf46 | 3.688803473 | 0.24 | 0.016 | 2.37E-11 | 10_mDC2 |
| LY75 | 3.474454933 | 0.72 | 0.192 | 3.39E-10 | 10_mDC2 |
| ZEB1 | 3.306907306 | 0.72 | 0.167 | 7.11E-10 | 10_mDC2 |
| ANXA6 | 3.649611024 | 0.6 | 0.127 | 1.27E-09 | 10_mDC2 |
| NUB1 | 3.070566785 | 0.72 | 0.189 | 1.50E-09 | 10_mDC2 |
| DSG2 | 3.576600098 | 0.32 | 0.034 | 2.16E-09 | 10_mDC2 |
| MARCKSL1 | 3.729524736 | 0.8 | 0.228 | 2.45E-09 | 10_mDC2 |
| UVRAG | 3.147383343 | 0.92 | 0.4 | 3.19E-09 | 10_mDC2 |
| ZNF260 | 3.992971518 | 0.28 | 0.026 | 3.26E-09 | 10_mDC2 |
| TXN | 4.45192774 | 0.88 | 0.447 | 4.61E-09 | 10_mDC2 |
| CCNG2 | 3.8612398 | 0.48 | 0.079 | 4.73E-09 | 10_mDC2 |
| PLAUR | -7.66459188 | 0.04 | 0.856 | 6.63E-09 | 10_mDC2 |
| RAB9A | 2.693539546 | 0.68 | 0.17 | 1.07E-08 | 10_mDC2 |
| LAD1 | 3.433426816 | 0.24 | 0.02 | 1.52E-08 | 10_mDC2 |
| GPR157 | 2.140287247 | 0.84 | 0.253 | 1.74E-08 | 10_mDC2 |
| IL15 | 3.197225457 | 0.68 | 0.187 | 4.49E-08 | 10_mDC2 |
| AC010654.1 | 2.922715105 | 0.32 | 0.037 | 4.55E-08 | 10_mDC2 |
| RAB30 | 3.001578162 | 0.32 | 0.037 | 5.05E-08 | 10_mDC2 |
| CERS6 | 3.236830997 | 0.88 | 0.355 | 6.78E-08 | 10_mDC2 |
| VOPP1 | 2.42586162 | 0.96 | 0.5 | 7.35E-08 | 10_mDC2 |
| TUBB6 | 2.839915779 | 0.72 | 0.208 | 1.20E-07 | 10_mDC2 |
| C1orf162 | -13.6044857 | 0 | 0.797 | 1.38E-07 | 10_mDC2 |
| KIF2A | 3.269602798 | 0.64 | 0.174 | 1.63E-07 | 10_mDC2 |
| ST3GAL6 | 2.830625284 | 0.6 | 0.138 | 2.31E-07 | 10_mDC2 |
| POGLUT1 | 3.200078156 | 0.56 | 0.129 | 2.70E-07 | 10_mDC2 |
| MREG | 2.965065782 | 0.32 | 0.04 | 4.12E-07 | 10_mDC2 |
| CD274 | 3.615984561 | 0.28 | 0.031 | 5.63E-07 | 10_mDC2 |
| NUAK2 | 2.519521515 | 0.44 | 0.077 | 5.67E-07 | 10_mDC2 |
| CLIC2 | 2.707248382 | 0.72 | 0.248 | 7.28E-07 | 10_mDC2 |
| NRP2 | 2.753059498 | 0.64 | 0.15 | 7.63E-07 | 10_mDC2 |
| DUSP5 | 2.471179995 | 0.92 | 0.428 | 7.96E-07 | 10_mDC2 |

| | | | | | |
|---|---|---|---|---|---|
| FCER1G | -3.20195773 | 0.4 | 0.87 | 1.03E-06 | 10_mDC2 |
| REPIN1 | 3.109897787 | 0.44 | 0.081 | 1.04E-06 | 10_mDC2 |
| CSF2RB | 2.921537422 | 0.4 | 0.065 | 1.15E-06 | 10_mDC2 |
| AIF1 | -5.05792246 | 0.12 | 0.784 | 1.57E-06 | 10_mDC2 |
| TMEM176A | 3.126124112 | 0.52 | 0.122 | 2.44E-06 | 10_mDC2 |
| ADORA2A | 2.58756339 | 0.4 | 0.066 | 3.60E-06 | 10_mDC2 |
| PVR | 4.305417089 | 0.24 | 0.026 | 3.76E-06 | 10_mDC2 |
| LINC00299 | 3.582933487 | 0.24 | 0.026 | 4.23E-06 | 10_mDC2 |
| RPS27L | 2.610151143 | 0.8 | 0.378 | 5.82E-06 | 10_mDC2 |
| IDO2 | 2.600945726 | 0.44 | 0.081 | 6.63E-06 | 10_mDC2 |
| CRIP1 | 2.733792989 | 0.88 | 0.43 | 8.60E-06 | 10_mDC2 |
| CBLB | 2.207787812 | 0.72 | 0.24 | 9.23E-06 | 10_mDC2 |
| SH3PXD2A | 3.657959425 | 0.28 | 0.036 | 1.04E-05 | 10_mDC2 |
| NAV1 | 1.961381795 | 0.68 | 0.193 | 1.09E-05 | 10_mDC2 |
| LAPTM5 | -1.78444781 | 0.56 | 0.94 | 1.36E-05 | 10_mDC2 |
| MS4A6A | -4.95919226 | 0.08 | 0.745 | 1.96E-05 | 10_mDC2 |
| SPECC1 | 3.019128185 | 0.68 | 0.218 | 2.24E-05 | 10_mDC2 |
| ITGB2 | -3.95845292 | 0.16 | 0.765 | 2.52E-05 | 10_mDC2 |
| CYB5A | 2.842771127 | 0.4 | 0.075 | 2.74E-05 | 10_mDC2 |
| TMEM131L | 1.748571373 | 0.52 | 0.114 | 2.85E-05 | 10_mDC2 |
| ABTB2 | 2.423426082 | 0.36 | 0.062 | 3.35E-05 | 10_mDC2 |
| RNASET2 | -3.22624125 | 0.36 | 0.823 | 4.00E-05 | 10_mDC2 |
| SESN1 | 2.069058027 | 0.76 | 0.285 | 4.05E-05 | 10_mDC2 |
| ATP8A1 | 2.605603728 | 0.4 | 0.072 | 5.02E-05 | 10_mDC2 |
| ARHGDIB | -3.64857569 | 0.12 | 0.761 | 5.23E-05 | 10_mDC2 |
| CTSB | -3.02179114 | 0.4 | 0.841 | 5.86E-05 | 10_mDC2 |
| PLXDC2 | -2.8786357 | 0.4 | 0.901 | 5.87E-05 | 10_mDC2 |
| TCAF2 | 2.912278368 | 0.24 | 0.028 | 6.05E-05 | 10_mDC2 |
| HSD11B1-AS1 | 4.874055519 | 0.24 | 0.03 | 7.40E-05 | 10_mDC2 |
| NET1 | 1.55308447 | 0.76 | 0.246 | 8.57E-05 | 10_mDC2 |
| RASSF4 | 2.212847668 | 0.8 | 0.427 | 0.0001143 | 10_mDC2 |
| SIAH2 | 2.347786942 | 0.56 | 0.152 | 0.0001158 | 10_mDC2 |
| IL2RG | 2.018765492 | 0.8 | 0.345 | 0.0001175 | 10_mDC2 |
| FCGRT | -5.37330637 | 0.08 | 0.69 | 0.0001565 | 10_mDC2 |
| NLRP3 | -4.23578969 | 0.08 | 0.733 | 0.0001889 | 10_mDC2 |
| OLR1 | -5.97968579 | 0.04 | 0.674 | 0.0002885 | 10_mDC2 |
| DCAKD | 2.891892194 | 0.24 | 0.031 | 0.0002961 | 10_mDC2 |
| S100A4 | -4.53766425 | 0.16 | 0.718 | 0.0003058 | 10_mDC2 |
| P2RY10 | 2.172832582 | 0.36 | 0.064 | 0.0003061 | 10_mDC2 |
| MAP3K1 | 2.292161901 | 0.6 | 0.184 | 0.0003146 | 10_mDC2 |
| ZFAND5 | 1.675067716 | 0.96 | 0.752 | 0.0004313 | 10_mDC2 |
| NDE1 | 2.751712005 | 0.48 | 0.124 | 0.0004417 | 10_mDC2 |
| LGALS2 | 1.978757622 | 0.44 | 0.096 | 0.0004455 | 10_mDC2 |
| RDX | 2.091642999 | 0.72 | 0.259 | 0.0004697 | 10_mDC2 |
| ALOX5AP | -3.63913687 | 0.12 | 0.727 | 0.0005064 | 10_mDC2 |
| TSPAN33 | 2.242112096 | 0.72 | 0.244 | 0.0005131 | 10_mDC2 |

| | | | | | |
|---|---|---|---|---|---|
| AC008105.3 | 2.004155741 | 0.64 | 0.21 | 0.0006201 | 10_mDC2 |
| TBC1D8 | 2.223164693 | 0.88 | 0.492 | 0.0006217 | 10_mDC2 |
| CST3 | -1.12906327 | 0.68 | 0.943 | 0.0006783 | 10_mDC2 |
| MS4A7 | -3.33954757 | 0.24 | 0.743 | 0.0008022 | 10_mDC2 |
| TBXAS1 | -4.52037696 | 0.08 | 0.674 | 0.0008113 | 10_mDC2 |
| PSEN2 | 3.029469885 | 0.24 | 0.033 | 0.0008267 | 10_mDC2 |
| TRADD | 2.366482169 | 0.48 | 0.128 | 0.0008711 | 10_mDC2 |
| TNFRSF11A | 3.30184041 | 0.28 | 0.046 | 0.0010265 | 10_mDC2 |
| SLC2A3 | -2.25860042 | 0.08 | 0.731 | 0.001057 | 10_mDC2 |
| MMD | 2.061724857 | 0.52 | 0.137 | 0.001082 | 10_mDC2 |
| PTGES2 | 2.309391807 | 0.52 | 0.15 | 0.0010911 | 10_mDC2 |
| TFCP2 | 2.169706566 | 0.44 | 0.105 | 0.0012155 | 10_mDC2 |
| NECAP2 | 2.235001689 | 0.56 | 0.18 | 0.0014466 | 10_mDC2 |
| ST8SIA4 | 1.859008547 | 0.84 | 0.422 | 0.0017031 | 10_mDC2 |
| FAS | 2.517277013 | 0.32 | 0.058 | 0.001728 | 10_mDC2 |
| PSAP | -1.79247139 | 0.52 | 0.897 | 0.0018412 | 10_mDC2 |
| COMMD1 | 1.678252679 | 0.72 | 0.258 | 0.0018623 | 10_mDC2 |
| TMEM39A | 1.957989667 | 0.72 | 0.264 | 0.0019918 | 10_mDC2 |
| CEBPB | -4.48455972 | 0.12 | 0.663 | 0.0022034 | 10_mDC2 |
| H6PD | 2.442599012 | 0.44 | 0.111 | 0.0023299 | 10_mDC2 |
| ZFP36L2 | -3.08111625 | 0.2 | 0.756 | 0.0025719 | 10_mDC2 |
| TTN-AS1 | 2.279569763 | 0.4 | 0.087 | 0.0026662 | 10_mDC2 |
| ATP11C | 2.527673427 | 0.44 | 0.106 | 0.0027534 | 10_mDC2 |
| SLC8A1 | -2.94830223 | 0.2 | 0.752 | 0.003019 | 10_mDC2 |
| CD53 | -2.77422909 | 0.2 | 0.74 | 0.0030192 | 10_mDC2 |
| BET1 | 2.946644452 | 0.28 | 0.049 | 0.0034609 | 10_mDC2 |
| L3MBTL4 | 2.100935301 | 0.56 | 0.163 | 0.0038074 | 10_mDC2 |
| ZFAS1 | 1.788888659 | 0.92 | 0.672 | 0.0041391 | 10_mDC2 |
| TMTC2 | 3.535777933 | 0.32 | 0.063 | 0.0047087 | 10_mDC2 |
| CPVL | -3.9543072 | 0.08 | 0.658 | 0.0053062 | 10_mDC2 |
| SPPL2A | 1.82941352 | 0.72 | 0.296 | 0.0053303 | 10_mDC2 |
| CCNH | -2.21280218 | 0.28 | 0.797 | 0.005512 | 10_mDC2 |
| RNASE6 | -12.557654 | 0 | 0.576 | 0.0056043 | 10_mDC2 |
| PPM1K | 2.343700341 | 0.4 | 0.095 | 0.0058393 | 10_mDC2 |
| CREM | -2.09159035 | 0.32 | 0.818 | 0.0058473 | 10_mDC2 |
| CHD3 | 2.123017245 | 0.52 | 0.155 | 0.0064221 | 10_mDC2 |
| CFLAR | 1.421163042 | 1 | 0.744 | 0.0074975 | 10_mDC2 |
| FNBP1 | 2.048457094 | 0.92 | 0.592 | 0.0077133 | 10_mDC2 |
| FCGR2B | -12.5457923 | 0 | 0.567 | 0.0078944 | 10_mDC2 |
| C5orf56 | 2.155590099 | 0.32 | 0.063 | 0.0086161 | 10_mDC2 |
| GRK3 | 1.841422014 | 0.76 | 0.294 | 0.0088353 | 10_mDC2 |
| LIMS1 | -2.26219115 | 0.4 | 0.821 | 0.0089187 | 10_mDC2 |
| TUBA1C | 2.529118643 | 0.76 | 0.402 | 0.0089522 | 10_mDC2 |
| CSF1R | -5.78139655 | 0.04 | 0.587 | 0.0090468 | 10_mDC2 |
| BTN2A2 | 2.222221651 | 0.44 | 0.12 | 0.0109469 | 10_mDC2 |
| IRF1 | 2.363215054 | 0.72 | 0.304 | 0.0119334 | 10_mDC2 |

| | | | | | |
|---|---|---|---|---|---|
| CHST7 | 2.480876612 | 0.28 | 0.05 | 0.0120951 | 10_mDC2 |
| TLR2 | -2.96390081 | 0.16 | 0.691 | 0.0121299 | 10_mDC2 |
| RNF115 | 2.487974655 | 0.6 | 0.214 | 0.0135698 | 10_mDC2 |
| FCGR2A | -2.80581156 | 0.28 | 0.714 | 0.0156657 | 10_mDC2 |
| YWHAH | -3.26130362 | 0.2 | 0.677 | 0.018012 | 10_mDC2 |
| PEX14 | 2.659688456 | 0.36 | 0.083 | 0.0182191 | 10_mDC2 |
| NMRK1 | 2.882755183 | 0.28 | 0.052 | 0.0187633 | 10_mDC2 |
| HCLS1 | -1.74366579 | 0.48 | 0.843 | 0.0194056 | 10_mDC2 |
| LYST | 2.57807413 | 0.56 | 0.203 | 0.0210409 | 10_mDC2 |
| GBP1 | 1.553165957 | 0.4 | 0.097 | 0.0248604 | 10_mDC2 |
| PLD4 | -12.2641849 | 0 | 0.536 | 0.0260957 | 10_mDC2 |
| JAK1 | 2.007350768 | 0.88 | 0.627 | 0.0269539 | 10_mDC2 |
| FXYD5 | -2.15894594 | 0.4 | 0.76 | 0.0272151 | 10_mDC2 |
| SGK1 | -2.15944868 | 0.56 | 0.858 | 0.0285171 | 10_mDC2 |
| TRAF1 | 1.583226772 | 0.76 | 0.323 | 0.028873 | 10_mDC2 |
| HAVCR2 | -3.18165078 | 0.08 | 0.622 | 0.0292116 | 10_mDC2 |
| ISCU | 1.895536318 | 0.64 | 0.283 | 0.030493 | 10_mDC2 |
| FPR1 | -3.06549324 | 0.24 | 0.673 | 0.0307067 | 10_mDC2 |
| SINHCAF | 1.97197773 | 0.48 | 0.144 | 0.0309334 | 10_mDC2 |
| MAP3K14 | 1.528439052 | 0.6 | 0.208 | 0.0337271 | 10_mDC2 |
| GMFG | -11.8584879 | 0 | 0.529 | 0.0341956 | 10_mDC2 |
| RGCC | -3.61924639 | 0.2 | 0.662 | 0.0342677 | 10_mDC2 |
| ANTXR2 | 2.619291018 | 0.56 | 0.212 | 0.0380119 | 10_mDC2 |
| PTPRE | -3.15161895 | 0.08 | 0.632 | 0.0388369 | 10_mDC2 |
| C17orf49 | 2.144837548 | 0.68 | 0.355 | 0.0393351 | 10_mDC2 |
| SLC41A2 | 2.649581347 | 0.64 | 0.274 | 0.0413452 | 10_mDC2 |
| TXNL1 | 1.815613265 | 0.64 | 0.255 | 0.0437202 | 10_mDC2 |
| PLSCR1 | -1.94269422 | 0.4 | 0.797 | 0.0443807 | 10_mDC2 |
| NR6A1 | 1.949511525 | 0.36 | 0.085 | 0.0470024 | 10_mDC2 |
| BLVRA | 2.135924451 | 0.4 | 0.111 | 0.0558992 | 10_mDC2 |
| ARPC3 | -1.5990351 | 0.6 | 0.833 | 0.0574645 | 10_mDC2 |
| CKLF | -3.32277093 | 0.16 | 0.63 | 0.0587914 | 10_mDC2 |
| ZBTB10 | 2.340319871 | 0.4 | 0.111 | 0.0594215 | 10_mDC2 |
| PSME1 | 1.397723421 | 0.8 | 0.57 | 0.0607203 | 10_mDC2 |
| HLA-DMB | -2.3303531 | 0.32 | 0.707 | 0.0615881 | 10_mDC2 |
| METRNL | -3.33535631 | 0.2 | 0.639 | 0.0677155 | 10_mDC2 |
| FLT3 | 1.446230866 | 0.68 | 0.283 | 0.0705634 | 10_mDC2 |
| SLC20A1 | 2.297438023 | 0.6 | 0.214 | 0.071687 | 10_mDC2 |
| TBC1D9 | 2.053957787 | 0.64 | 0.27 | 0.0815976 | 10_mDC2 |
| AXL | -3.77167811 | 0.08 | 0.575 | 0.092343 | 10_mDC2 |
| PTPRC | -1.34715164 | 0.4 | 0.831 | 0.0984685 | 10_mDC2 |
| USP12 | 1.730137237 | 0.68 | 0.31 | 0.0995848 | 10_mDC2 |
| GPR137B | 1.651497504 | 0.84 | 0.469 | 0.1088755 | 10_mDC2 |
| VASH1 | -4.34683395 | 0.08 | 0.557 | 0.1089778 | 10_mDC2 |
| PSD3 | 2.258069206 | 0.48 | 0.161 | 0.1090597 | 10_mDC2 |
| HCST | -3.23196654 | 0.16 | 0.606 | 0.1101591 | 10_mDC2 |

| | | | | | |
|---|---|---|---|---|---|
| IRAK3 | -2.46392664 | 0.2 | 0.706 | 0.1125352 | 10_mDC2 |
| EIF4A1 | -1.28325859 | 0.64 | 0.876 | 0.1139347 | 10_mDC2 |
| MCM5 | 1.844898478 | 0.56 | 0.216 | 0.1224421 | 10_mDC2 |
| VAC14 | 2.258360473 | 0.4 | 0.119 | 0.1229285 | 10_mDC2 |
| LCP2 | -3.25678751 | 0.08 | 0.588 | 0.1279848 | 10_mDC2 |
| TMEM176B | 2.463198904 | 0.52 | 0.212 | 0.1289197 | 10_mDC2 |
| CORO1A | -3.5905864 | 0.12 | 0.574 | 0.1472542 | 10_mDC2 |
| MTSS1 | 2.209483131 | 0.56 | 0.208 | 0.1496094 | 10_mDC2 |
| ITGAX | -2.91571447 | 0.08 | 0.602 | 0.1535188 | 10_mDC2 |
| MBP | -3.0085308 | 0.16 | 0.615 | 0.1573745 | 10_mDC2 |
| SPTLC2 | -5.60047654 | 0.04 | 0.515 | 0.1580248 | 10_mDC2 |
| MYO1G | 1.406187655 | 0.68 | 0.293 | 0.1824166 | 10_mDC2 |
| FCER1A | -13.0812267 | 0 | 0.48 | 0.1885042 | 10_mDC2 |
| DOCK4 | -2.42829414 | 0.4 | 0.788 | 0.1961224 | 10_mDC2 |
| WHAMM | 1.799166907 | 0.52 | 0.189 | 0.2069914 | 10_mDC2 |
| PALD1 | -5.28083316 | 0.04 | 0.508 | 0.2139623 | 10_mDC2 |
| PLGRKT | 2.11186975 | 0.36 | 0.096 | 0.2148571 | 10_mDC2 |
| TRAFD1 | 2.455939103 | 0.28 | 0.062 | 0.2161137 | 10_mDC2 |
| PACS1 | 2.013220956 | 0.68 | 0.347 | 0.2195969 | 10_mDC2 |
| N4BP2L1 | 2.498795267 | 0.56 | 0.257 | 0.2221789 | 10_mDC2 |
| C3 | -6.18047119 | 0.08 | 0.525 | 0.2275412 | 10_mDC2 |
| GK | -2.36222169 | 0.36 | 0.73 | 0.2324041 | 10_mDC2 |
| RASSF2 | 1.476949789 | 0.8 | 0.403 | 0.2352068 | 10_mDC2 |
| RAB31 | -1.68690923 | 0.64 | 0.876 | 0.2673217 | 10_mDC2 |
| CYBB | -11.9418292 | 0 | 0.469 | 0.2687671 | 10_mDC2 |
| HIF1A | -1.96548816 | 0.4 | 0.786 | 0.2769677 | 10_mDC2 |
| SPG11 | 2.315656944 | 0.52 | 0.202 | 0.2808953 | 10_mDC2 |
| MOB1B | 2.505862128 | 0.64 | 0.26 | 0.2820098 | 10_mDC2 |
| SAMHD1 | -2.57247175 | 0.16 | 0.619 | 0.2824959 | 10_mDC2 |
| WSB2 | 2.61511694 | 0.32 | 0.083 | 0.2908491 | 10_mDC2 |
| CSF3R | -3.65943653 | 0.08 | 0.541 | 0.307736 | 10_mDC2 |
| ACSL1 | -2.14914233 | 0.56 | 0.819 | 0.3213066 | 10_mDC2 |
| TMEM131 | 1.4288739 | 0.72 | 0.38 | 0.3288723 | 10_mDC2 |
| LST1 | -3.04619727 | 0.16 | 0.58 | 0.3338766 | 10_mDC2 |
| TTYH2 | 1.335815709 | 0.6 | 0.236 | 0.3503581 | 10_mDC2 |
| VPS8 | 1.708854851 | 0.52 | 0.176 | 0.3599017 | 10_mDC2 |
| ST3GAL1 | 1.188397146 | 0.76 | 0.416 | 0.3977389 | 10_mDC2 |
| SLCO3A1 | 1.866254923 | 0.48 | 0.166 | 0.4372796 | 10_mDC2 |
| MEF2C | -2.41438656 | 0.2 | 0.662 | 0.4554108 | 10_mDC2 |
| RNF149 | -1.82907516 | 0.56 | 0.797 | 0.4820116 | 10_mDC2 |
| PIKFYVE | 1.741570898 | 0.56 | 0.233 | 0.4825531 | 10_mDC2 |
| TNFRSF1B | -2.00476246 | 0.28 | 0.699 | 0.5013189 | 10_mDC2 |
| AREG | -1.27136645 | 0.48 | 0.838 | 0.5180048 | 10_mDC2 |
| ARL17A | 2.273983672 | 0.28 | 0.066 | 0.5186658 | 10_mDC2 |
| ZNF331 | -2.2608203 | 0.24 | 0.694 | 0.5263662 | 10_mDC2 |
| PLK3 | -4.8836036 | 0.04 | 0.479 | 0.5782825 | 10_mDC2 |

| | | | | | |
|---|---|---|---|---|---|
| ITPKB | 2.028554873 | 0.36 | 0.102 | 0.5871203 | 10_mDC2 |
| WAS | -2.83499951 | 0.08 | 0.543 | 0.6060102 | 10_mDC2 |
| FMN1 | -2.79249981 | 0.12 | 0.596 | 0.6096997 | 10_mDC2 |
| TAP1 | 1.51161526 | 0.6 | 0.237 | 0.6234795 | 10_mDC2 |
| HBEGF | -12.482864 | 0 | 0.442 | 0.6337636 | 10_mDC2 |
| LRRFIP2 | 1.225635881 | 0.68 | 0.306 | 0.6428849 | 10_mDC2 |
| TRIP10 | 1.627560782 | 0.48 | 0.167 | 0.6912185 | 10_mDC2 |
| UTRN | -3.7068591 | 0.08 | 0.525 | 0.6991401 | 10_mDC2 |
| RAB20 | -3.45905426 | 0.08 | 0.521 | 0.7061276 | 10_mDC2 |
| DSC2 | 2.241862711 | 0.36 | 0.107 | 0.710179 | 10_mDC2 |
| LIMD2 | -4.70624651 | 0.04 | 0.472 | 0.7215377 | 10_mDC2 |
| MAT2A | -2.14730112 | 0.4 | 0.721 | 0.7471424 | 10_mDC2 |
| SLC7A5 | -2.6361028 | 0.2 | 0.619 | 0.7926862 | 10_mDC2 |
| HEATR5A | 2.100959269 | 0.4 | 0.124 | 0.8013663 | 10_mDC2 |
| SPPL3 | 1.594757566 | 0.56 | 0.231 | 0.8288366 | 10_mDC2 |
| SH3BP5 | -5.12412496 | 0.04 | 0.466 | 0.8333605 | 10_mDC2 |
| APBB1IP | -2.22809295 | 0.04 | 0.508 | 0.8529785 | 10_mDC2 |
| ST3GAL5 | 2.732748102 | 0.28 | 0.071 | 0.9173092 | 10_mDC2 |
| RELB | 1.463945005 | 0.76 | 0.419 | 0.9228142 | 10_mDC2 |
| SLC43A2 | -5.26236218 | 0.04 | 0.459 | 0.9826776 | 10_mDC2 |
| CD68 | -2.44479443 | 0.12 | 0.574 | 0.9910694 | 10_mDC2 |
| PRELID1 | -4.3235765 | 0.08 | 0.483 | 1 | 10_mDC2 |
| LAIR1 | -11.5840971 | 0 | 0.426 | 1 | 10_mDC2 |
| LDLRAD4 | -1.6091307 | 0.64 | 0.916 | 1 | 10_mDC2 |
| LPCAT2 | -3.92198553 | 0.08 | 0.5 | 1 | 10_mDC2 |
| RFTN1 | 1.695141313 | 0.84 | 0.521 | 1 | 10_mDC2 |
| HSPB1 | 2.62276123 | 0.68 | 0.41 | 1 | 10_mDC2 |
| JDP2 | -4.8549395 | 0.04 | 0.461 | 1 | 10_mDC2 |
| IL1B | -2.64841079 | 0.32 | 0.715 | 1 | 10_mDC2 |
| RHOQ | -2.49941005 | 0.16 | 0.576 | 1 | 10_mDC2 |
| PDE4B | -1.39652389 | 0.32 | 0.806 | 1 | 10_mDC2 |
| GPAT3 | -4.69668893 | 0.04 | 0.465 | 1 | 10_mDC2 |
| PHKB | 1.881906619 | 0.52 | 0.215 | 1 | 10_mDC2 |
| MBNL2 | 2.373127467 | 0.52 | 0.251 | 1 | 10_mDC2 |
| TBL1X | 1.963937328 | 0.44 | 0.163 | 1 | 10_mDC2 |
| ARHGAP15 | -2.12949 | 0.2 | 0.655 | 1 | 10_mDC2 |
| ATP13A3 | -2.79427379 | 0.16 | 0.572 | 1 | 10_mDC2 |
| VASP | -3.98208766 | 0.04 | 0.468 | 1 | 10_mDC2 |
| DCAF6 | 1.176532046 | 0.6 | 0.253 | 1 | 10_mDC2 |
| SYMPK | 1.613724665 | 0.48 | 0.174 | 1 | 10_mDC2 |
| CD163 | -3.49240753 | 0.08 | 0.502 | 1 | 10_mDC2 |
| MRTFA | 1.611536696 | 0.8 | 0.46 | 1 | 10_mDC2 |
| AKR1B1 | -11.3607252 | 0 | 0.411 | 1 | 10_mDC2 |
| SELPLG | 1.656286942 | 0.52 | 0.211 | 1 | 10_mDC2 |
| BMP2K | 1.82138764 | 0.64 | 0.332 | 1 | 10_mDC2 |
| CCL22 | 2.26064053 | 0.56 | 0.259 | 1 | 10_mDC2 |

| | | | | | |
|---|---|---|---|---|---|
| PSTPIP2 | -11.7900968 | 0 | 0.41 | 1 | 10_mDC2 |
| RASGEF1B | -2.24376289 | 0.32 | 0.695 | 1 | 10_mDC2 |
| PPP1R16B | 1.728206117 | 0.44 | 0.155 | 1 | 10_mDC2 |
| CST7 | 1.244414279 | 0.8 | 0.483 | 1 | 10_mDC2 |
| ASMTL | 1.814295118 | 0.44 | 0.156 | 1 | 10_mDC2 |
| MSR1 | -3.06284798 | 0.12 | 0.543 | 1 | 10_mDC2 |
| NEDD9 | -3.17580528 | 0.12 | 0.528 | 1 | 10_mDC2 |
| LRRK1 | 1.506503628 | 0.6 | 0.253 | 1 | 10_mDC2 |
| ARID1B | 1.775384909 | 0.8 | 0.473 | 1 | 10_mDC2 |
| CLEC7A | -1.99180375 | 0.24 | 0.648 | 1 | 10_mDC2 |
| RNF145 | 1.189946657 | 0.76 | 0.403 | 1 | 10_mDC2 |
| FOSB | -1.84716162 | 0.36 | 0.752 | 1 | 10_mDC2 |
| MVP | 1.598013255 | 0.56 | 0.259 | 1 | 10_mDC2 |
| SESN3 | 1.896637912 | 0.44 | 0.158 | 1 | 10_mDC2 |
| EPSTI1 | 1.831761394 | 0.64 | 0.287 | 1 | 10_mDC2 |
| CDC14A | 1.910682796 | 0.36 | 0.115 | 1 | 10_mDC2 |
| TRABD2A | 2.632480298 | 0.36 | 0.113 | 1 | 10_mDC2 |
| RAP2B | 1.732558986 | 0.56 | 0.242 | 1 | 10_mDC2 |
| PYCARD | -3.01149254 | 0.04 | 0.465 | 1 | 10_mDC2 |
| FLOT2 | 1.11780922 | 0.56 | 0.236 | 1 | 10_mDC2 |
| RNF130 | -1.63859509 | 0.32 | 0.716 | 1 | 10_mDC2 |
| HNRNPA3 | -1.67027069 | 0.28 | 0.683 | 1 | 10_mDC2 |
| SSR1 | -4.42718767 | 0.04 | 0.431 | 1 | 10_mDC2 |
| RCOR1 | -11.6778112 | 0 | 0.394 | 1 | 10_mDC2 |
| IL1R2 | -3.65283825 | 0.08 | 0.482 | 1 | 10_mDC2 |
| SATB1 | -11.5941491 | 0 | 0.392 | 1 | 10_mDC2 |
| GADD45A | 2.371716176 | 0.28 | 0.079 | 1 | 10_mDC2 |
| IL21R | 1.673428567 | 0.32 | 0.093 | 1 | 10_mDC2 |
| MFSD1 | -2.3841296 | 0.24 | 0.593 | 1 | 10_mDC2 |
| LTC4S | -11.3057772 | 0 | 0.391 | 1 | 10_mDC2 |
| CD40 | 2.747361062 | 0.4 | 0.149 | 1 | 10_mDC2 |
| ARPC1B | -1.37982436 | 0.52 | 0.793 | 1 | 10_mDC2 |
| TTF2 | 1.333234902 | 0.28 | 0.071 | 1 | 10_mDC2 |
| EBI3 | 2.391746955 | 0.56 | 0.315 | 1 | 10_mDC2 |
| DPYSL2 | 1.514183223 | 0.56 | 0.247 | 1 | 10_mDC2 |
| PARVG | -3.02006682 | 0.12 | 0.498 | 1 | 10_mDC2 |
| ARHGAP4 | -11.0841121 | 0 | 0.383 | 1 | 10_mDC2 |
| ATP8B4 | -3.72524397 | 0.08 | 0.467 | 1 | 10_mDC2 |
| FGL2 | -1.54057413 | 0.12 | 0.539 | 1 | 10_mDC2 |
| HLA-F | 1.221346417 | 0.64 | 0.334 | 1 | 10_mDC2 |
| SYAP1 | -1.79331126 | 0.36 | 0.69 | 1 | 10_mDC2 |
| OSTF1 | 1.343311242 | 0.68 | 0.413 | 1 | 10_mDC2 |
| ANXA1 | -1.68116603 | 0.2 | 0.594 | 1 | 10_mDC2 |
| ITGAV | -4.01063497 | 0.08 | 0.449 | 1 | 10_mDC2 |
| PIP4K2A | 1.874094719 | 0.6 | 0.312 | 1 | 10_mDC2 |
| TM6SF1 | -11.1991872 | 0 | 0.376 | 1 | 10_mDC2 |

| | | | | | |
|---|---|---|---|---|---|
| CD14 | -4.60625373 | 0.08 | 0.447 | 1 | 10_mDC2 |
| LINC01619 | 1.127281423 | 0.64 | 0.35 | 1 | 10_mDC2 |
| SOCS1 | 3.786601503 | 0.36 | 0.125 | 1 | 10_mDC2 |
| C1QC | -3.89555244 | 0.2 | 0.552 | 1 | 10_mDC2 |
| TAPBP | 1.15093338 | 0.76 | 0.48 | 1 | 10_mDC2 |
| SMIM14 | 1.760824351 | 0.48 | 0.21 | 1 | 10_mDC2 |
| AP1B1 | -4.50464855 | 0.04 | 0.413 | 1 | 10_mDC2 |
| PDXK | -11.1628715 | 0 | 0.373 | 1 | 10_mDC2 |
| PICALM | -1.09911726 | 0.36 | 0.808 | 1 | 10_mDC2 |
| RAP1B | 1.142865082 | 0.92 | 0.7 | 1 | 10_mDC2 |
| CRTAP | -3.78279013 | 0.04 | 0.424 | 1 | 10_mDC2 |
| BCL11A | 1.969969771 | 0.32 | 0.096 | 1 | 10_mDC2 |
| GABARAPL1 | -3.19223922 | 0.12 | 0.482 | 1 | 10_mDC2 |
| RUNX1 | -1.93441591 | 0.24 | 0.598 | 1 | 10_mDC2 |
| LINC-PINT | 2.282253036 | 0.56 | 0.253 | 1 | 10_mDC2 |
| TGFB1 | -1.66464208 | 0.28 | 0.676 | 1 | 10_mDC2 |
| MS4A4E | -11.6595235 | 0 | 0.367 | 1 | 10_mDC2 |
| TOM1 | -3.3837877 | 0.08 | 0.455 | 1 | 10_mDC2 |
| AMPD2 | -11.0741073 | 0 | 0.366 | 1 | 10_mDC2 |
| FAM102B | -11.5020654 | 0 | 0.365 | 1 | 10_mDC2 |
| LRCH3 | 2.026270645 | 0.44 | 0.169 | 1 | 10_mDC2 |
| SLC25A24 | 1.528354869 | 0.36 | 0.124 | 1 | 10_mDC2 |
| ANKRD10 | 1.080358506 | 0.68 | 0.355 | 1 | 10_mDC2 |
| MARCKS | 1.258335973 | 0.76 | 0.535 | 1 | 10_mDC2 |
| CLCN7 | 1.883662355 | 0.32 | 0.101 | 1 | 10_mDC2 |
| HMGB2 | -5.42507504 | 0.04 | 0.394 | 1 | 10_mDC2 |
| LPXN | -3.80532229 | 0.08 | 0.436 | 1 | 10_mDC2 |
| TBCA | 1.263578993 | 0.68 | 0.424 | 1 | 10_mDC2 |
| TNFAIP2 | 1.211197202 | 0.6 | 0.277 | 1 | 10_mDC2 |
| SERPINA1 | -2.23846321 | 0.32 | 0.625 | 1 | 10_mDC2 |
| NT5DC1 | 1.948914908 | 0.28 | 0.079 | 1 | 10_mDC2 |
| ELL2 | -1.77129323 | 0.36 | 0.748 | 1 | 10_mDC2 |
| GPBP1 | 1.288923075 | 0.8 | 0.546 | 1 | 10_mDC2 |
| GNA13 | -1.04878925 | 0.48 | 0.815 | 1 | 10_mDC2 |
| DMXL1 | 1.375160669 | 0.52 | 0.238 | 1 | 10_mDC2 |
| CHD1 | -2.02120895 | 0.24 | 0.613 | 1 | 10_mDC2 |
| CLN8 | -11.1521921 | 0 | 0.354 | 1 | 10_mDC2 |
| CD9 | -5.02172538 | 0.04 | 0.392 | 1 | 10_mDC2 |
| FAM168A | 1.788876423 | 0.4 | 0.157 | 1 | 10_mDC2 |
| SEC22C | 1.531002818 | 0.28 | 0.079 | 1 | 10_mDC2 |
| TNIP2 | 1.764174384 | 0.52 | 0.222 | 1 | 10_mDC2 |
| ADGRE2 | -2.2600602 | 0.12 | 0.521 | 1 | 10_mDC2 |
| MCTP1 | -3.75340654 | 0.08 | 0.437 | 1 | 10_mDC2 |
| UBTF | 1.437542752 | 0.28 | 0.079 | 1 | 10_mDC2 |
| CLEC5A | -11.4924645 | 0 | 0.35 | 1 | 10_mDC2 |
| UGCG | 1.25583622 | 0.68 | 0.358 | 1 | 10_mDC2 |

| | | | | | |
|---|---|---|---|---|---|
| EXOSC8 | 1.445962777 | 0.32 | 0.099 | 1 | 10_mDC2 |
| PPARD | -2.6085817 | 0.2 | 0.54 | 1 | 10_mDC2 |
| ARHGAP18 | -2.983662 | 0.08 | 0.453 | 1 | 10_mDC2 |
| LY86 | -1.45138122 | 0.12 | 0.502 | 1 | 10_mDC2 |
| SIGLEC10 | -4.5402933 | 0.04 | 0.384 | 1 | 10_mDC2 |
| DAGLB | -4.0933561 | 0.04 | 0.395 | 1 | 10_mDC2 |
| FOXO1 | 2.303346245 | 0.4 | 0.15 | 1 | 10_mDC2 |
| GRB2 | -1.35493346 | 0.4 | 0.772 | 1 | 10_mDC2 |
| PPP1R15B | 1.321810624 | 0.56 | 0.282 | 1 | 10_mDC2 |
| ERCC1 | -3.0941682 | 0.12 | 0.445 | 1 | 10_mDC2 |
| ENTPD4 | 1.551543388 | 0.36 | 0.128 | 1 | 10_mDC2 |
| COA1 | 1.452374259 | 0.44 | 0.182 | 1 | 10_mDC2 |
| RAB8B | 1.587259575 | 0.72 | 0.475 | 1 | 10_mDC2 |
| VSIG4 | -3.42500141 | 0.04 | 0.397 | 1 | 10_mDC2 |
| POU2F2 | -11.1258933 | 0 | 0.336 | 1 | 10_mDC2 |
| HEXB | -2.98054331 | 0.12 | 0.449 | 1 | 10_mDC2 |
| SKAP2 | -3.54835042 | 0.04 | 0.392 | 1 | 10_mDC2 |
| PDLIM5 | 1.323120533 | 0.56 | 0.265 | 1 | 10_mDC2 |
| BIN1 | -2.38125749 | 0.24 | 0.547 | 1 | 10_mDC2 |
| NAMPT | -1.37731672 | 0.6 | 0.863 | 1 | 10_mDC2 |
| DOCK10 | 1.52840521 | 0.76 | 0.484 | 1 | 10_mDC2 |
| PADI2 | -2.44202456 | 0.2 | 0.56 | 1 | 10_mDC2 |
| UBE2F | 1.002242503 | 0.76 | 0.473 | 1 | 10_mDC2 |
| PHF21A | 2.305316154 | 0.44 | 0.169 | 1 | 10_mDC2 |
| GAB1 | 1.850848516 | 0.4 | 0.158 | 1 | 10_mDC2 |
| PIK3R1 | -2.3102291 | 0.08 | 0.458 | 1 | 10_mDC2 |
| FOS | -1.24571319 | 0.64 | 0.904 | 1 | 10_mDC2 |
| EVI2B | -2.44993361 | 0.12 | 0.473 | 1 | 10_mDC2 |
| CLEC10A | -4.32329579 | 0.04 | 0.373 | 1 | 10_mDC2 |
| EZH2 | 2.378678905 | 0.32 | 0.109 | 1 | 10_mDC2 |
| INSIG1 | -2.23526135 | 0.28 | 0.588 | 1 | 10_mDC2 |
| ELMO1 | -1.2675708 | 0.56 | 0.827 | 1 | 10_mDC2 |
| BTG2 | -1.19676668 | 0.32 | 0.637 | 1 | 10_mDC2 |
| FBXW5 | 1.570702464 | 0.44 | 0.203 | 1 | 10_mDC2 |
| KCTD12 | -3.95981004 | 0.08 | 0.396 | 1 | 10_mDC2 |
| SKIL | -1.13410215 | 0.32 | 0.76 | 1 | 10_mDC2 |
| IFITM2 | -1.52424662 | 0.48 | 0.749 | 1 | 10_mDC2 |
| TSPO | -1.75334679 | 0.32 | 0.608 | 1 | 10_mDC2 |
| IGSF6 | -3.19589703 | 0.08 | 0.411 | 1 | 10_mDC2 |
| NIBAN1 | 1.233492877 | 0.64 | 0.372 | 1 | 10_mDC2 |
| PHF20 | -1.26669906 | 0.4 | 0.679 | 1 | 10_mDC2 |
| MYO1F | -3.40844748 | 0.04 | 0.38 | 1 | 10_mDC2 |
| PHACTR2 | 1.276296342 | 0.44 | 0.177 | 1 | 10_mDC2 |
| AHCYL2 | 1.546977748 | 0.32 | 0.107 | 1 | 10_mDC2 |
| PHACTR1 | -2.32016251 | 0.2 | 0.524 | 1 | 10_mDC2 |
| CBX5 | 1.611300668 | 0.32 | 0.105 | 1 | 10_mDC2 |

| | | | | | |
|---|---|---|---|---|---|
| SREBF2 | 1.437084441 | 0.32 | 0.108 | 1 | 10_mDC2 |
| ARL4C | -1.94035776 | 0.12 | 0.484 | 1 | 10_mDC2 |
| GTDC1 | 1.217871904 | 0.44 | 0.181 | 1 | 10_mDC2 |
| SMARCA4 | 1.102758555 | 0.4 | 0.154 | 1 | 10_mDC2 |
| PSME2 | 1.374461696 | 0.76 | 0.521 | 1 | 10_mDC2 |
| YEATS2 | 1.346087649 | 0.32 | 0.108 | 1 | 10_mDC2 |
| MAPK8 | 2.251763438 | 0.52 | 0.263 | 1 | 10_mDC2 |
| ITGA4 | 1.434871159 | 0.56 | 0.304 | 1 | 10_mDC2 |
| LYZ | -1.73525558 | 0.44 | 0.713 | 1 | 10_mDC2 |
| DHFR | 1.556143881 | 0.32 | 0.114 | 1 | 10_mDC2 |
| ENTPD1 | -1.45985174 | 0.2 | 0.564 | 1 | 10_mDC2 |
| ATP6AP2 | -2.36448045 | 0.08 | 0.422 | 1 | 10_mDC2 |
| FAM107B | 1.94511429 | 0.6 | 0.358 | 1 | 10_mDC2 |
| ATOX1 | 1.634533846 | 0.52 | 0.306 | 1 | 10_mDC2 |
| KDM5B | 2.404516388 | 0.36 | 0.133 | 1 | 10_mDC2 |
| ZNRD1 | 1.680978215 | 0.36 | 0.145 | 1 | 10_mDC2 |
| SLA | -2.18081834 | 0.08 | 0.422 | 1 | 10_mDC2 |
| IL18 | -2.0792909 | 0.24 | 0.526 | 1 | 10_mDC2 |
| CNN2 | 1.229257916 | 0.68 | 0.449 | 1 | 10_mDC2 |
| FGR | -1.69318686 | 0.2 | 0.555 | 1 | 10_mDC2 |
| TGFBI | -4.39232101 | 0.08 | 0.38 | 1 | 10_mDC2 |
| TAGAP | -2.8285557 | 0.12 | 0.444 | 1 | 10_mDC2 |
| C3AR1 | -11.1298864 | 0 | 0.309 | 1 | 10_mDC2 |
| TREM2 | -4.16787933 | 0.08 | 0.39 | 1 | 10_mDC2 |
| CD302 | -1.28215658 | 0.2 | 0.549 | 1 | 10_mDC2 |
| SPP1 | -1.71005636 | 0.36 | 0.627 | 1 | 10_mDC2 |
| ARL6IP5 | 1.224690312 | 0.72 | 0.495 | 1 | 10_mDC2 |
| PLIN2 | -3.30196038 | 0.12 | 0.433 | 1 | 10_mDC2 |
| INSR | -3.09175011 | 0.04 | 0.371 | 1 | 10_mDC2 |
| FYB1 | -2.21379983 | 0.2 | 0.506 | 1 | 10_mDC2 |
| NUMB | -1.96797828 | 0.08 | 0.436 | 1 | 10_mDC2 |
| EREG | -12.3361038 | 0 | 0.304 | 1 | 10_mDC2 |
| FOXN3 | -1.13937119 | 0.64 | 0.844 | 1 | 10_mDC2 |
| EIF4G3 | -2.09450816 | 0.08 | 0.427 | 1 | 10_mDC2 |
| ABI3 | -10.790196 | 0 | 0.302 | 1 | 10_mDC2 |
| TXNDC11 | 1.123932172 | 0.48 | 0.222 | 1 | 10_mDC2 |
| FAM118A | 1.219458123 | 0.44 | 0.187 | 1 | 10_mDC2 |
| RAP1A | -1.46368176 | 0.32 | 0.649 | 1 | 10_mDC2 |
| NETO2 | 1.822666257 | 0.36 | 0.143 | 1 | 10_mDC2 |
| CD37 | -1.20729412 | 0.48 | 0.729 | 1 | 10_mDC2 |
| ICAM3 | -10.7108262 | 0 | 0.3 | 1 | 10_mDC2 |
| C1QA | -3.22447352 | 0.24 | 0.51 | 1 | 10_mDC2 |
| THBD | -11.5899218 | 0 | 0.299 | 1 | 10_mDC2 |
| RUFY3 | 1.258015353 | 0.52 | 0.271 | 1 | 10_mDC2 |
| MAML3 | -1.34731081 | 0.2 | 0.604 | 1 | 10_mDC2 |
| ARAP1 | -10.7692001 | 0 | 0.295 | 1 | 10_mDC2 |

| | | | | | |
|---|---|---|---|---|---|
| DUSP2 | -1.32696419 | 0.32 | 0.606 | 1 | 10_mDC2 |
| RANBP2 | -1.96775448 | 0.48 | 0.706 | 1 | 10_mDC2 |
| CD44 | -1.41393713 | 0.4 | 0.696 | 1 | 10_mDC2 |
| ARF1 | -1.24563412 | 0.32 | 0.647 | 1 | 10_mDC2 |
| DDX21 | -1.98695822 | 0.2 | 0.514 | 1 | 10_mDC2 |
| HMGB1 | -1.06244333 | 0.56 | 0.772 | 1 | 10_mDC2 |
| AUTS2 | 1.299398395 | 0.64 | 0.37 | 1 | 10_mDC2 |
| PDE4A | -1.69617953 | 0.2 | 0.529 | 1 | 10_mDC2 |
| RILPL2 | -1.11370982 | 0.56 | 0.778 | 1 | 10_mDC2 |
| PRMT2 | 1.021699607 | 0.6 | 0.34 | 1 | 10_mDC2 |
| FBP1 | -10.6358081 | 0 | 0.289 | 1 | 10_mDC2 |
| LINC00963 | -3.9667204 | 0.04 | 0.335 | 1 | 10_mDC2 |
| TRAF5 | 1.047946341 | 0.44 | 0.183 | 1 | 10_mDC2 |
| BZW2 | 1.40181446 | 0.36 | 0.14 | 1 | 10_mDC2 |
| CD1C | -3.76393164 | 0.04 | 0.343 | 1 | 10_mDC2 |
| LILRB4 | -2.64085051 | 0.16 | 0.441 | 1 | 10_mDC2 |
| SLC11A1 | -2.66189281 | 0.16 | 0.459 | 1 | 10_mDC2 |
| DPP7 | -1.78928453 | 0.12 | 0.473 | 1 | 10_mDC2 |
| CASC4 | 1.394108714 | 0.4 | 0.179 | 1 | 10_mDC2 |
| TREM1 | -4.28381229 | 0.04 | 0.33 | 1 | 10_mDC2 |
| INPP5D | -2.55777328 | 0.16 | 0.453 | 1 | 10_mDC2 |
| SMC5 | 2.039573779 | 0.48 | 0.254 | 1 | 10_mDC2 |
| PPP3CA | 1.940872659 | 0.56 | 0.307 | 1 | 10_mDC2 |
| ITGA5 | -3.3695501 | 0.12 | 0.394 | 1 | 10_mDC2 |
| RASSF5 | -1.72354151 | 0.24 | 0.52 | 1 | 10_mDC2 |
| NBN | 1.066056806 | 0.4 | 0.159 | 1 | 10_mDC2 |
| MAP4 | 1.063262682 | 0.56 | 0.281 | 1 | 10_mDC2 |
| THBS1 | -4.7294884 | 0.08 | 0.369 | 1 | 10_mDC2 |
| CMTM3 | -10.5081888 | 0 | 0.281 | 1 | 10_mDC2 |
| RAC2 | -10.5489288 | 0 | 0.28 | 1 | 10_mDC2 |
| SLC16A3 | -1.61766484 | 0.32 | 0.579 | 1 | 10_mDC2 |
| MAP3K13 | 1.28069784 | 0.52 | 0.257 | 1 | 10_mDC2 |
| CLEC2B | -3.54791444 | 0.08 | 0.359 | 1 | 10_mDC2 |
| ATF4 | 1.23757324 | 0.64 | 0.392 | 1 | 10_mDC2 |
| ARHGAP26 | -1.33143318 | 0.36 | 0.649 | 1 | 10_mDC2 |
| SPOP | 1.303571914 | 0.44 | 0.204 | 1 | 10_mDC2 |
| IL16 | -10.5333754 | 0 | 0.274 | 1 | 10_mDC2 |
| MSN | -1.14101378 | 0.44 | 0.707 | 1 | 10_mDC2 |
| STX4 | -3.6349878 | 0.08 | 0.359 | 1 | 10_mDC2 |
| RPS6KA3 | -1.71720688 | 0.2 | 0.537 | 1 | 10_mDC2 |
| NDUFA9 | 1.379130361 | 0.32 | 0.119 | 1 | 10_mDC2 |
| PIK3C3 | 1.332133286 | 0.32 | 0.116 | 1 | 10_mDC2 |
| HERC4 | 1.213948858 | 0.4 | 0.185 | 1 | 10_mDC2 |
| KDM4C | 1.164179823 | 0.52 | 0.272 | 1 | 10_mDC2 |
| MCUB | -10.6293726 | 0 | 0.269 | 1 | 10_mDC2 |
| OSER1 | 1.413186824 | 0.44 | 0.214 | 1 | 10_mDC2 |

| | | | | | |
|---|---|---|---|---|---|
| MTHFD1L | -1.51664908 | 0.08 | 0.4 | 1 | 10_mDC2 |
| UBE2Z | 1.623764612 | 0.4 | 0.184 | 1 | 10_mDC2 |
| SMDT1 | -2.08661583 | 0.16 | 0.461 | 1 | 10_mDC2 |
| SELENOK | -1.42846025 | 0.4 | 0.607 | 1 | 10_mDC2 |
| SLC16A10 | -1.80109295 | 0.12 | 0.412 | 1 | 10_mDC2 |
| RARA | -2.83353021 | 0.08 | 0.367 | 1 | 10_mDC2 |
| ST14 | -10.5205698 | 0 | 0.266 | 1 | 10_mDC2 |
| LINC01128 | -10.5902338 | 0 | 0.265 | 1 | 10_mDC2 |
| SFMBT2 | -2.81050456 | 0.2 | 0.455 | 1 | 10_mDC2 |
| EIF4E | -2.28070013 | 0.16 | 0.447 | 1 | 10_mDC2 |
| GUSB | -3.79326141 | 0.08 | 0.342 | 1 | 10_mDC2 |
| MAFB | -3.4885576 | 0.08 | 0.358 | 1 | 10_mDC2 |
| SDCBP | -1.09287669 | 0.48 | 0.719 | 1 | 10_mDC2 |
| MAP4K3 | -1.87265351 | 0.16 | 0.473 | 1 | 10_mDC2 |
| TNFRSF21 | -10.7408464 | 0 | 0.263 | 1 | 10_mDC2 |
| GRAMD4 | -10.5900005 | 0 | 0.263 | 1 | 10_mDC2 |
| ZNF706 | -1.74868944 | 0.16 | 0.477 | 1 | 10_mDC2 |
| HSPA1B | 1.402828843 | 0.68 | 0.437 | 1 | 10_mDC2 |
| PTAFR | -10.6235622 | 0 | 0.262 | 1 | 10_mDC2 |
| A2M | -2.43781378 | 0.16 | 0.451 | 1 | 10_mDC2 |
| ANXA5 | -1.00874603 | 0.52 | 0.768 | 1 | 10_mDC2 |
| SAP30 | -2.95749994 | 0.08 | 0.369 | 1 | 10_mDC2 |
| HMGN4 | 1.140908952 | 0.4 | 0.176 | 1 | 10_mDC2 |
| FCGR1A | -3.73564728 | 0.04 | 0.316 | 1 | 10_mDC2 |
| AOAH | -1.79968394 | 0.2 | 0.509 | 1 | 10_mDC2 |
| SSNA1 | 1.521205055 | 0.44 | 0.223 | 1 | 10_mDC2 |
| USP24 | 1.738873721 | 0.4 | 0.191 | 1 | 10_mDC2 |
| FCGR3A | -3.70853804 | 0.08 | 0.346 | 1 | 10_mDC2 |
| B3GNT5 | -1.29599411 | 0.24 | 0.588 | 1 | 10_mDC2 |
| JAK2 | 1.66161808 | 0.56 | 0.312 | 1 | 10_mDC2 |
| CPM | -2.80263929 | 0.12 | 0.404 | 1 | 10_mDC2 |
| TRABD | -1.51534045 | 0.24 | 0.524 | 1 | 10_mDC2 |
| SLC25A37 | -2.36120406 | 0.08 | 0.377 | 1 | 10_mDC2 |
| PPP1R2 | 1.06299377 | 0.68 | 0.402 | 1 | 10_mDC2 |
| CTSS | -1.04140529 | 0.44 | 0.719 | 1 | 10_mDC2 |
| RIN3 | -1.23477211 | 0.44 | 0.674 | 1 | 10_mDC2 |
| SLC38A1 | 1.181388913 | 0.48 | 0.247 | 1 | 10_mDC2 |
| CD48 | -2.96024167 | 0.08 | 0.35 | 1 | 10_mDC2 |
| MGAT4A | -3.26318604 | 0.08 | 0.345 | 1 | 10_mDC2 |
| ADAM17 | -2.3151586 | 0.16 | 0.44 | 1 | 10_mDC2 |
| TCIRG1 | -1.71908111 | 0.12 | 0.403 | 1 | 10_mDC2 |
| ABCA1 | -3.3874401 | 0.08 | 0.355 | 1 | 10_mDC2 |
| FLT1 | -3.86375571 | 0.08 | 0.336 | 1 | 10_mDC2 |
| AL136987.1 | -10.8294655 | 0 | 0.252 | 1 | 10_mDC2 |
| IQSEC1 | -10.6859042 | 0 | 0.251 | 1 | 10_mDC2 |
| AL135905.2 | -10.4258798 | 0 | 0.251 | 1 | 10_mDC2 |

| | | | | | |
|---|---|---|---|---|---|
| NDRG2 | -10.3353328 | 0 | 0.251 | 1 | 10_mDC2 |
| CFP | 1.391117461 | 0.4 | 0.191 | 1 | 10_mDC2 |
| ENG | -3.23531675 | 0.04 | 0.31 | 1 | 10_mDC2 |
| ARHGAP24 | -2.04822096 | 0.08 | 0.377 | 1 | 10_mDC2 |
| MAP3K20 | 1.31565329 | 0.36 | 0.157 | 1 | 10_mDC2 |
| PCED1B-AS1 | -10.2589781 | 0 | 0.25 | 1 | 10_mDC2 |
| CD93 | -3.42263852 | 0.04 | 0.31 | 1 | 10_mDC2 |
| JAML | -2.16863667 | 0.08 | 0.371 | 1 | 10_mDC2 |
| RASAL2 | -2.84281959 | 0.04 | 0.315 | 1 | 10_mDC2 |
| RBM47 | -1.42771395 | 0.36 | 0.645 | 1 | 10_mDC2 |
| NDUFB10 | -2.46400634 | 0.16 | 0.396 | 1 | 10_mDC2 |
| KMT2A | 1.18178653 | 0.44 | 0.22 | 1 | 10_mDC2 |
| FAM135A | 1.717090558 | 0.44 | 0.22 | 1 | 10_mDC2 |
| DTNA | -11.0462321 | 0 | 0.246 | 1 | 10_mDC2 |
| EPS8 | -2.90012224 | 0.08 | 0.351 | 1 | 10_mDC2 |
| C7orf50 | -3.67549159 | 0.04 | 0.294 | 1 | 10_mDC2 |
| CLINT1 | 1.585594515 | 0.52 | 0.294 | 1 | 10_mDC2 |
| HSD17B11 | -3.29588668 | 0.04 | 0.303 | 1 | 10_mDC2 |
| HADHA | -1.59838299 | 0.16 | 0.443 | 1 | 10_mDC2 |
| MAP2K3 | -2.14381273 | 0.12 | 0.4 | 1 | 10_mDC2 |
| CD69 | -2.18575894 | 0.16 | 0.442 | 1 | 10_mDC2 |
| CHRNE | -10.4030731 | 0 | 0.243 | 1 | 10_mDC2 |
| FRMD4B | -4.26797344 | 0.04 | 0.292 | 1 | 10_mDC2 |
| CNPY3 | -1.49134616 | 0.24 | 0.512 | 1 | 10_mDC2 |
| MRC1 | -4.8370871 | 0.04 | 0.287 | 1 | 10_mDC2 |
| ZNF710 | -10.7782708 | 0 | 0.242 | 1 | 10_mDC2 |
| PHC2 | -2.03809788 | 0.08 | 0.371 | 1 | 10_mDC2 |
| PPT1 | -1.44790577 | 0.28 | 0.55 | 1 | 10_mDC2 |
| ABHD2 | -4.73595455 | 0.04 | 0.284 | 1 | 10_mDC2 |
| MS4A4A | -2.57577706 | 0.16 | 0.404 | 1 | 10_mDC2 |
| CD164 | -1.68473845 | 0.24 | 0.471 | 1 | 10_mDC2 |
| OTULINL | -3.85240657 | 0.04 | 0.288 | 1 | 10_mDC2 |
| RNF166 | -10.2525249 | 0 | 0.238 | 1 | 10_mDC2 |
| SUPT4H1 | -2.46436895 | 0.16 | 0.394 | 1 | 10_mDC2 |
| HACD4 | -1.83192115 | 0.04 | 0.313 | 1 | 10_mDC2 |
| CD109 | -4.29817447 | 0.04 | 0.284 | 1 | 10_mDC2 |
| DIAPH1 | -3.71717425 | 0.04 | 0.287 | 1 | 10_mDC2 |
| RGS19 | -1.91443232 | 0.04 | 0.312 | 1 | 10_mDC2 |
| MPRIP | -2.73101545 | 0.08 | 0.341 | 1 | 10_mDC2 |
| EPN1 | -3.07070528 | 0.04 | 0.298 | 1 | 10_mDC2 |
| NGLY1 | -4.14791992 | 0.04 | 0.285 | 1 | 10_mDC2 |
| FAM53B | 1.048107392 | 0.52 | 0.275 | 1 | 10_mDC2 |
| RASA1 | -2.43951039 | 0.12 | 0.379 | 1 | 10_mDC2 |
| GLIPR1 | -1.25056304 | 0.24 | 0.551 | 1 | 10_mDC2 |
| SORL1 | -1.73764883 | 0.2 | 0.467 | 1 | 10_mDC2 |
| KLHL24 | 1.028717945 | 0.52 | 0.277 | 1 | 10_mDC2 |

| | | | | | |
|---|---|---|---|---|---|
| CACYBP | 1.462452713 | 0.4 | 0.192 | 1 | 10_mDC2 |
| NANS | -2.39527932 | 0.08 | 0.34 | 1 | 10_mDC2 |
| TIMP1 | -1.4567193 | 0.4 | 0.683 | 1 | 10_mDC2 |
| BOD1L1 | -3.46409912 | 0.04 | 0.289 | 1 | 10_mDC2 |
| SATB1-AS1 | -10.6876741 | 0 | 0.232 | 1 | 10_mDC2 |
| KLF2 | -4.48091444 | 0.08 | 0.321 | 1 | 10_mDC2 |
| MTMR14 | -3.06506707 | 0.04 | 0.294 | 1 | 10_mDC2 |
| RIN2 | -10.9841696 | 0 | 0.231 | 1 | 10_mDC2 |
| KCNQ3 | -11.9746928 | 0 | 0.23 | 1 | 10_mDC2 |
| FPR3 | -2.02415203 | 0.08 | 0.355 | 1 | 10_mDC2 |
| CNOT4 | 1.126702385 | 0.44 | 0.234 | 1 | 10_mDC2 |
| LRRK2 | -3.05631168 | 0.12 | 0.353 | 1 | 10_mDC2 |
| ATP6AP1 | -1.35953226 | 0.32 | 0.548 | 1 | 10_mDC2 |
| WDFY2 | -10.5983622 | 0 | 0.228 | 1 | 10_mDC2 |
| CAPZA2 | -2.40956596 | 0.2 | 0.411 | 1 | 10_mDC2 |
| AFF3 | -1.6439218 | 0.16 | 0.446 | 1 | 10_mDC2 |
| SMG1 | 1.112566445 | 0.64 | 0.397 | 1 | 10_mDC2 |
| PITPNB | 1.088949373 | 0.48 | 0.254 | 1 | 10_mDC2 |
| ITGAM | -10.2348766 | 0 | 0.227 | 1 | 10_mDC2 |
| IL18R1 | -1.9140396 | 0.08 | 0.348 | 1 | 10_mDC2 |
| CSNK1G2 | -3.55237066 | 0.04 | 0.276 | 1 | 10_mDC2 |
| ARL5A | -2.05004102 | 0.16 | 0.409 | 1 | 10_mDC2 |
| MAN2A1 | -2.39589587 | 0.24 | 0.463 | 1 | 10_mDC2 |
| FAM149A | -10.7867678 | 0 | 0.224 | 1 | 10_mDC2 |
| CYTH3 | -10.6984881 | 0 | 0.224 | 1 | 10_mDC2 |
| SORT1 | -10.507815 | 0 | 0.224 | 1 | 10_mDC2 |
| TTC7A | -2.19253808 | 0.16 | 0.414 | 1 | 10_mDC2 |
| SRI | 1.0811655 | 0.52 | 0.304 | 1 | 10_mDC2 |
| TUT7 | -3.34777279 | 0.08 | 0.314 | 1 | 10_mDC2 |
| GTF3A | -2.08607832 | 0.16 | 0.399 | 1 | 10_mDC2 |
| CD99 | -1.33022015 | 0.24 | 0.527 | 1 | 10_mDC2 |
| SKI | -2.56130465 | 0.04 | 0.291 | 1 | 10_mDC2 |
| C20orf27 | -2.46885685 | 0.08 | 0.327 | 1 | 10_mDC2 |
| SOAT1 | -2.68698339 | 0.12 | 0.357 | 1 | 10_mDC2 |
| TOP1 | -1.10854782 | 0.44 | 0.678 | 1 | 10_mDC2 |
| KDM2A | 1.033800471 | 0.68 | 0.477 | 1 | 10_mDC2 |
| SMNDC1 | 1.195631387 | 0.44 | 0.238 | 1 | 10_mDC2 |
| SEPTIN9 | -2.91654479 | 0.04 | 0.289 | 1 | 10_mDC2 |
| MAPKAPK2 | -3.08581664 | 0.08 | 0.315 | 1 | 10_mDC2 |
| ARHGEF40 | -10.3382197 | 0 | 0.22 | 1 | 10_mDC2 |
| QSOX1 | -3.48388303 | 0.04 | 0.272 | 1 | 10_mDC2 |
| RABGEF1 | -1.61430471 | 0.32 | 0.545 | 1 | 10_mDC2 |
| KIF1B | -2.53117556 | 0.04 | 0.289 | 1 | 10_mDC2 |
| MIDN | -1.03585458 | 0.2 | 0.464 | 1 | 10_mDC2 |
| EVI2A | -3.61355809 | 0.04 | 0.269 | 1 | 10_mDC2 |
| HOMER1 | -11.1880937 | 0 | 0.218 | 1 | 10_mDC2 |

| | | | | | |
|---|---|---|---|---|---|
| MACC1 | -11.1841825 | 0 | 0.218 | 1 | 10_mDC2 |
| TEX14 | -2.00449103 | 0.2 | 0.446 | 1 | 10_mDC2 |
| RNF13 | -1.37641749 | 0.16 | 0.413 | 1 | 10_mDC2 |
| UQCRQ | -2.41647506 | 0.12 | 0.363 | 1 | 10_mDC2 |
| PLP2 | -2.11979788 | 0.12 | 0.373 | 1 | 10_mDC2 |
| REEP3 | 1.713970276 | 0.48 | 0.275 | 1 | 10_mDC2 |
| INPP4A | -3.99146632 | 0.04 | 0.269 | 1 | 10_mDC2 |
| EPB41L3 | -1.4866889 | 0.44 | 0.647 | 1 | 10_mDC2 |
| PTPN6 | -2.06019797 | 0.16 | 0.4 | 1 | 10_mDC2 |
| SQOR | -10.0862024 | 0 | 0.216 | 1 | 10_mDC2 |
| PIAS1 | 1.289156914 | 0.72 | 0.506 | 1 | 10_mDC2 |
| CITED2 | -10.8821624 | 0 | 0.216 | 1 | 10_mDC2 |
| TRPM2 | -10.359155 | 0 | 0.215 | 1 | 10_mDC2 |
| FBRSL1 | 1.186509957 | 0.48 | 0.242 | 1 | 10_mDC2 |
| MYOF | -10.3931176 | 0 | 0.214 | 1 | 10_mDC2 |
| PID1 | -11.1709328 | 0 | 0.214 | 1 | 10_mDC2 |
