## Supplementary Table 10 for "MAIT cells have a negative impact on glioblastoma"

**Supplementary Table 10.** Details of the antibodies used for CODEX experiment

| Antigen | Clone | Vendor | Catalogue number | RRID | Oligonucleotide tag# | Catalogue Number (Oligo) | Working Dilution | Exposure time (msec) |
|---|---|---|---|---|---|---|---|---|
| **CD45** | HI30 | Akoya Biosciences | 4150003 | AB_2895052 | 1 | 5450013 | 1:200 | 250 |
| **CD15** | HI98 | Biolegend | 301902 | AB_314194 | 28 | 5150005 | 1:250 | 120 |
| **CD66b** | G10F5 | Biolegend | 305102 | AB_314494 | 49 | 5150012 | 1:267 | 200 |
| **CD161** | HP-3G10 | Biolegend | 339902 | AB_1501090 | 29 | 5250005 | 1:125 | 300 |
| **CD3** | UCHT1 | Akoya Biosciences | 4350008 | AB_2895047 | 15 | 5350001 | 1:200 | 400 |
| **TCR Vα7.2** | 3C10 | Biolegend | 351702 | AB_10900258 | 33 | 5350006 | 1:125 | 400 |
