## Supplementary Table 11 for "MAIT cells have a negative impact on glioblastoma"

**Supplementary Table 11.** Specimens from Glioblastoma patient tumor used in CODEX study

| ID | Diagnosis | IDH status | Primary vs. Recurrent | Age at surgery | Sex | Anatomic location | Laterality | Methylation subtype |
|---|---|---|---|---|---|---|---|---|
| S1 | Glioblastoma, IDH-wildtype | Wildtype | Recurrent | 39 | M | Frontal lobe | Right | RTK2 subtype |
| S2 | Glioblastoma, IDH-wildtype | Wildtype | Recurrent | 78 | M | Temporal lobe | Right | RTK2 subtype |
| S3 | Glioblastoma, IDH-wildtype | Wildtype | Recurrent (of S4) | 55 | M | Frontal lobe | Right | RTK2 subtype |
| S4 | Glioblastoma, IDH-wildtype | Wildtype | Primary (of S3) | 50 | M | Frontal lobe | Right | No methylation (prior to assay) |
