## Supplementary Table 12 for "MAIT cells have a negative impact on glioblastoma"

**Supplementary Table 12.** Summary of number of cells identified in the CODEX data. Data were calculated from 4 GBM patient specimens.

| Analysis Region | Total Cells | MAIT | Neutrophil | CD45 | CD3 | Area | Sample ID | Type |
|---|---|---|---|---|---|---|---|---|
| Cellular Tumor 10 | 13910 | **2** | **5** | 766 | 47 | 3658686.75 | S4 | CT |
| Cellular Tumor 11 | 23534 | **2** | **0** | 892 | 41 | 5342696.50 | S4 | CT |
| Cellular Tumor 01 | 2820 | **0** | **12** | 261 | 31 | 893603.88 | S1 | CT |
| Cellular Tumor 02 | 2437 | **0** | **9** | 377 | 46 | 676032.94 | S1 | CT |
| Cellular Tumor 03 | 3907 | **0** | **35** | 331 | 96 | 976378.25 | S1 | CT |
| Cellular Tumor 04 | 4464 | **0** | **49** | 336 | 27 | 1246583.13 | S1 | CT |
| Cellular Tumor 05 | 1904 | **0** | **53** | 94 | 19 | 551108.56 | S1 | CT |
| Cellular Tumor 06 | 4484 | **0** | **37** | 322 | 31 | 1612965.88 | S1 | CT |
| Cellular Tumor 07 | 4351 | **1** | **266** | 505 | 42 | 1492950.88 | S1 | CT |
| Cellular Tumor 08 | 3367 | **3** | **287** | 469 | 80 | 958041.94 | S1 | CT |
| Cellular Tumor 09 | 14094 | **4** | **214** | 803 | 683 | 4527915.00 | S1 | CT |
| Cellular Tumor 12 | 3400 | **2** | **0** | 300 | 4 | 809932.88 | S3 | CT |
| Cellular Tumor 13 | 52504 | **2** | **8** | 3639 | 56 | 19172028.00 | S3 | CT |
| Cellular Tumor 14 | 5233 | **0** | **13** | 102 | 23 | 5130981.00 | S2 | CT |
| Cellular Tumor 15 | 1129 | **0** | **0** | 35 | 2 | 265003.69 | S2 | CT |
| Cellular Tumor 16 | 939 | **0** | **1** | 3 | 3 | 306470.91 | S2 | CT |
| Cellular Tumor 17 | 19424 | **1** | **11** | 380 | 150 | 5487941.50 | S2 | CT |
| Leading Edge 1 | 166 | **1** | **0** | 22 | 2 | 76452.45 | S3 | LE |
| Leading Edge 2 | 2901 | **0** | **55** | 269 | 66 | 2029063.75 | S3 | LE |
| Leading Edge 3 | 228 | **0** | **0** | 31 | 0 | 256650.02 | S3 | LE |
| Leading Edge 4 | 810 | **0** | **1** | 86 | 2 | 600840.94 | S3 | LE |
| Leading Edge 5 | 816 | **0** | **8** | 27 | 8 | 287649.75 | S3 | LE |
| Leading Edge 6 | 828 | **0** | **5** | 132 | 8 | 2018858.75 | S2 | LE |
| Leading Edge 7 | 6433 | **1** | **9** | 538 | 312 | 7309501.50 | S2 | LE |
| Leading Edge 8 | 2272 | **1** | **7** | 78 | 30 | 2063397.13 | S2 | LE |
| Leading Edge 9 | 415 | **0** | **1** | 7 | 1 | 1334600.88 | S2 | LE |
| Leading Edge 10 | 1216 | **0** | **150** | 206 | 9 | 1053511.38 | S2 | LE |
| Leading Edge 11 | 12191 | **0** | **88** | 978 | 137 | 14010879.00 | S2 | LE |
| Leading Edge 12 | 4535 | **0** | **41** | 176 | 53 | 7912806.00 | S2 | LE |
| Leading Edge 13 | 4498 | **0** | **8** | 392 | 156 | 2829959.00 | S2 | LE |
| Leading Edge 14 | 6906 | **0** | **14** | 1856 | 229 | 5362600.00 | S2 | LE |
| Leading Edge 15 | 2279 | **0** | **9** | 88 | 21 | 5236061.00 | S2 | LE |
| Leading Edge 16 | 4133 | **0** | **31** | 294 | 37 | 4632118.00 | S2 | LE |
| Leading Edge 17 | 1085 | **0** | **2** | 34 | 7 | 2397420.25 | S2 | LE |
| Leading Edge 18 | 3380 | **1** | **28** | 159 | 68 | 8399206.00 | S2 | LE |
| Necrosis | 63309 | **9** | **102** | 6279 | 1154 | 28592428.00 | S4 | NR |
